# Richness and composition of phyllosphere *Methylobacterium* communities cause variation in *Arabidopsis thaliana* growth

**DOI:** 10.1101/2024.08.30.610551

**Authors:** Jocelyn Lauzon, Jérémie Pelletier, Élanore Favron, Zihui Wang, Steven W. Kembel

**Author notes:** All data and scripts used for this article are available on a public GitHub repository: https://github.com/lauzonj/methylobacterium-arabidopsis-growth.

## Abstract

The phyllosphere – the aerial parts of plants – forms a vast microbial habitat that harbors diverse bacterial communities playing key roles in ecosystem function. The foliar surface is thus a promising study system to investigate biodiversity-ecosystem function relationships. Researchers have found a positive correlation between leaf bacterial diversity and ecosystem productivity, but the causality of this relationship has yet to be demonstrated. To understand how the diversity and composition of phyllosphere bacterial communities could cause variation in the growth of their host plants, we assembled synthetic communities composed of different diversity and compositions of *Methylobacterium* strains – a plant growth-promoting bacterial genus ubiquitous in the phyllosphere – that we inoculated on *Arabidopsis thaliana* grown in gnotobiotic conditions. We hypothesized that (1) increasing *Methylobacterium* diversity should cause an increase in host growth; (2) strains should differ in their impact on host growth; and (3) the relationship between bacterial diversity and plant productivity should be strain-dependent. Our results supported our three hypotheses but revealed unpredicted patterns in how *A. thaliana* leaf biomass varied according to inoculated *Methylobacterium* strain richness and identity. Increasing bacterial richness induced a higher host leaf biomass, but only after an initial reduction in biomass, suggesting competition alleviation by multispecies interactions. Two *Methylobacterium* strains showed beneficial effects on *A. thaliana* growth, and one strain was detrimental for the plant. Community composition shaped the relationship between diversity and productivity, highlighting the importance of community mutualistic and antagonistic interactions. Furthermore, niche complementarity was likely the main ecological mechanism driving the diversity-productivity relationship in our study system. By demonstrating the causal effects of *Methylobacterium* community diversity and composition on host plant growth, our experiment shed light on the importance of phyllosphere bacteria in terrestrial ecosystem functioning.

## INTRODUCTION

The role of biodiversity as one of the cornerstones of ecosystem dynamics and functioning is well known, whether through its effects on productivity, stability, or resilience (Tilman et al. 2014). A notable example of the link between biodiversity and ecosystem function is the positive effect of diversity on ecosystem productivity, as demonstrated by the well-studied positive relationship between plant community diversity and biomass production (Tilman et al. 2001). This biodiversity effect can be explained by two mechanisms, namely complementarity and selection, depending on whether the ecosystem response to diversity is dictated by the complementarity of species niches versus the selection of highly productive species (Loreau and Hector 2001). While the relationships between biodiversity and ecosystem function have been well studied within communities of organisms from the same trophic level (Van Der Plas 2019), including bacteria (Jiang 2007; Bell et al. 2005), we understand less about how the biodiversity of microorganisms can influence the functioning of their hosts.

Plant leaves harbor diverse microbial communities, most often dominated by bacteria (Vorholt 2012). These bacteria exert an influence on plant and terrestrial ecosystems. While bacteria are important plant pathogens, bacterial taxa can also promote host growth, assist in nutrient acquisition, stimulate host protection against pathogens and abiotic stresses, and play a critical role in carbon and nitrogen biogeochemical cycles, atmosphere purification, and climate regulation (Bashir et al. 2022). The high diversity and ecological importance of the plant microbiome thus make the phyllosphere a promising ecosystem to study ecological theory including biodiversity-ecosystem functioning associations (Meyer and Leveau 2012). A previous study (Laforest-Lapointe et al. 2017) showed that phyllosphere bacterial diversity was positively correlated with plant community productivity, acting as a key explanatory variable for models linking plant diversity with ecosystem productivity, and thus explaining a significant part of the plant biodiversity-productivity relationship. However, their study could not determine the causality of the relationship between microbial biodiversity and host plant productivity, since it was based on an experiment where plant diversity but not microbial diversity was manipulated. Flores-Núñez et al. (2023) modified the *Agave* leaf microbiome by inoculating bacteria and fungi consortia in a field experiment. They did not find a direct effect of microbial diversity on plant growth, but found that diverse communities mitigated the negative effect of deleterious microorganisms on plant growth, as well as an effect of diversity on leaf development pattern. The causal relation between phyllosphere bacterial diversity and plant productivity has not yet been clearly demonstrated.

The objectives of our study were to test for causal relationships between leaf bacterial diversity and host plant productivity; to reveal the role of bacterial taxa and their community interactions in driving this causation; and to shed light on the ecological mechanisms at play in the diversity-productivity relationship. To address these objectives, we investigated how the diversity and composition of phyllosphere bacterial communities influence the growth of *Arabidopsis thaliana* host plants by assembling and inoculating synthetic communities of phylogenetically diverse strains of *Methylobacterium* in a gnotobiotic experimental setup. Our choice to work with synthetic communities was motivated by the need to move beyond a correlational perspective to understand causal links between the phyllosphere microbiome and host and ecosystem function (Vorholt et al. 2017). *Methylobacterium* is a diverse and ubiquitous bacterial genus of plant leaves, known for its methylotrophy and plant growth-promoting characteristic (Abanda-Nkpwatt et al. 2006), and is often employed in synthetic communities experiments (Carlström et al. 2019; Innerebner et al. 2011).

We tested three hypotheses, using strain richness at the moment of inoculation as a diversity measure, and rosette leaf dry biomass at the end of the vegetative growth period as a measure of plant growth and productivity. Considering the previously observed positive correlation between leaf bacterial diversity and plant community productivity (Laforest-Lapointe et al. 2017), as well as the demonstrated positive effect of inoculating *Methylobacterium* strains on plant growth (Madhaiyan et al. 2004), our first hypothesis was that a greater *Methylobacterium* taxonomic diversity on plant leaves should be associated with increased host plant productivity. We predicted a positive logarithmic correlation between the number of strains contained in inoculated synthetic communities and *A. thaliana* leaf biomass, in line with the redundancy hypothesis (Naeem et al. 1995). Our second hypothesis was that *Methylobacterium* taxa would differentially influence plant growth. *Methylobacterium* strains exhibit genetic, phenotypic, and ecological differences – including dissimilarities in their interactions with other microorganisms (Carlström et al. 2019), host preferences (Knief et al. 2010; Leducq et al. 2022), substrate utilization, and metabolic pathways (Alessa et al. 2021). These differences could result in variation in the growth-promoting effect of different individual strains on the one hand, and on the other, in bacterial- and host-dependent mutualistic and antagonistic interactions driving ecosystem functions. We thus predicted that the presence of some specific *Methylobacterium* strains would lead to a higher-than-average leaf biomass of *A. thaliana*. Thirdly, considering the potential importance of microbial taxa and their community interactions in determining plant growth (Bell et al. 2005), our third hypothesis was that some *Methylobacterium* strains present in communities would influence the relation between bacterial diversity and plant growth. In other words, the strength of the effect of bacterial diversity on plant growth should be dependent on the strains composing the leaf community. We thus predicted that strains having a larger-than-average positive effect on *A. thaliana* leaf biomass (while testing our second hypothesis) would also, when present in communities, increase the positive effects of strain richness on leaf biomass. Finally, without having an *a priori* hypothesis, we tried to disentangle the relative importance of complementarity and selection in driving the biodiversity effect of *Methylobacterium* strains on *A. thaliana* vegetative growth.

## MATERIAL AND METHODS

### Arabidopsis thaliana growth

Seeds of *Arabidopsis thaliana* Col-0 were surface sterilized and stratified prior to seeding. For sterilization, seeds were soaked 2 min in 70% ethanol, rinsed with sterile water, then soaked 8 min in a 2.5% sodium hypochlorite solution containing 0.2% Triton, and finally rinsed 12 times with sterile water (protocol adapted from Innerebner et al. [2011] and Lindsey et al. [2017]). Stratification occurred at 4°C for 4 days. Plants were grown inside sterile Magenta boxes with vented lids (1 seed/box), on a sterile minimal salts medium (1 cm thick) supplemented with 1% sucrose, 0.7% plant agar, and 0.2µm-filtered Gamborg’s vitamins solution (pH = 5.6), at 20°C under a 16-hour photoperiod regime (PAR 400-700nm = 120 µmol photons m^-2^s^-1^).

### *Methylobacterium* strains and synthetic community design

Twelve *Methylobacterium* strains, representing diverse clades of the genus, were chosen to form the different synthetic communities (Appendix S1: Table S1). These strains had been previously isolated from tree leaves as part of a temperate phyllosphere study (Leducq et al. 2022). Our experimental design involved synthetic communities of six richness values (number of strains) for the diversity treatment. We assembled twelve monocultures (richness = 1), twelve different communities composed of a pair of strains (richness = 2), and six different communities for each of the other richness values (4, 6, 8, and 10) for a total of 48 distinct community compositions (Appendix S1: Table S2). All twelve strains were present the same number of times among the different communities of a given richness value.

### Bacterial cultures and community assembly

Strains were grown for three weeks at 22°C on an R2A Agar medium supplemented with succinate 3.5 mM as a carbon source. Just before community synthesis and inoculation, bacterial colonies were washed with 2 mL 10mM MgCl2 and gently detached from the medium’s surface with a sterile glass loop. For every strain, cells from petri dish were collected in a single sterile 50 mL tube and the suspension was homogenized by pipetting up and down. An extra 20 mL of 10 mM MgCl2 was added, and the suspension was gently vortexed. As a surrogate for bacterial cell concentration, the optical density at 600 nm (OD600) was measured in triplicate on a BioTek Eon Microplate Spectrophotometer plate reader (Richmond Scientific). All twelve liquid strains were then diluted to OD600 0.2 prior to building the communities. Strains were assembled in synthetic communities in equal relative abundance (1:1 ratio between every strain of a community). Those newly formed communities were finally diluted in 10 mM MgCl2 to an OD600 of 0.02 before inoculation (Innerebner et al. 2011).

### Bacterial inoculation on *A. thaliana*

Synthetic communities were inoculated on *A. thaliana* leaves after 14 days of plant growth post-stratification (Innerebner et al. 2011). At the moment of inoculation, the rosettes contained a mean of 5 true leaves longer than 1 mm and had reached 15-20% of their final size (growth stage #1.05; Boyes et al. 2001). We inoculated synthetic communities on plants by distributing a total of 40 µL of bacterial suspension over the leaves’ surface. Every plant was inoculated with the same quantity of bacterial cells, approximately 4 x 10^5^ cells. Each of the 48 unique community compositions was inoculated on three plants, and we dropped sterile 10mM MgCl2 on six control plants – for a total of 150 plant samples.

### Harvest and dry leaf biomass measurement

Plants were harvested after 38 days of growth, to ensure the rosettes had reached their final size. Plants were cut into three parts: roots, rosette (leaves) and stem. Roots and stems were discarded. We dried rosettes in a Heratherm IMH750-S incubator (Thermo Scientific) at 70°C for 48 hours. The dry biomass was weighted immediately after drying.

### Statistical analyses

All statistical analyses were performed using R (v4.2.2; R Core Team, 2021). Data was manipulated with the help of packages *dplyr* (v1.1.2; Wickham, François, et al. 2023) and *tidyr* (v1.3.0; Wickham, Vaughan, et al. 2023). Figures were made using the base R *graphics* package and the *ggplot2* package (v3.4.3; Wickham 2016). Model selection, as well as model exploration and averaging, were performed with the *MuMIn* package (v1.47.5; Bartoń 2023). Samples whose box contained fungal contaminants on the agar at the moment of harvesting (*n*=5) were discarded, as well as 2 mislabeled samples. We did not consider control plants (*n*=6) as samples having a “strain richness” of zero in our models, but we plotted their data in figures as a visual reference for the interpretation of the results. We used the remaining 137 samples for all statistical analyses. All models used dry leaf biomass as a continuous numerical response variable. To address our hypotheses, candidate models were evaluated based on the corrected Akaike Information Criterion (AICc) and assumptions of the selected models were visually assessed by plotting standardized residuals by fitted values (to test for the homoscedasticity of residuals) and by plotting sample quantiles by theoretical quantiles (Q-Q plot to test for the normality of the residuals’ distribution). All models’ assumptions were met (Appendix S1: Figure S1 to S3).

We quantified the relationship between leaf biomass and strain richness by constructing two linear models: a linear regression (additive biodiversity effect hypothesis) and a second-degree polynomial regression (redundant biodiversity effect). Both models were compared to a null model to assess their fit to the observed data. Then, to determine whether specific strains had a higher-than-average positive influence on leaf biomass, without consideration of diversity effects, we built a saturated linear model that included twelve binary variables coding the presence or absence of each strain in samples, without taking into account interactions between strains. We used the function *dredge* in *MuMIn* to generate every possible model based on the saturated model (Appendix S1: Table S3a). We reported coefficients and standard errors of included parameters in all models with a ΔAICc ≤ 2, to attest the effect of individual strains’ presence on leaf biomass and to reveal which strains had a higher-than-average impact. We selected the three strains having a higher-than-average effect on leaf biomass to test if these would interact with diversity to influence the polynomial regression fit linking strain richness and dry leaf biomass. We explored the full model space with the function *dredge*, starting from a saturated linear model that included strain richness (with second-degree polynomial terms), the presence or absence of each of the three strains, and all possible interactions between those four variables (Appendix S1: Table S3b). To overcome model uncertainty, we performed multi-model inference with the function *model.avg* in *MuMIn* to generate fully averaged parameter estimates based on all models with a ΔAICc ≤ 2. We used the function *emmip* in package *emmeans* (v1.8.2; Lenth 2022) to extract fitted values and 95% confidence intervals for each combination of the three strains’ presence or absence, over the entire range of richness values.

Finally, to analyze whether complementarity or selection was driving the leaf biomass response to strain richness, we built a conceptual framework based on three alternative models (Appendix S1: Figure S4). We first calculated expected biomass values assuming that, for every mixed bacterial community, all strains’ effects on biomass in mixtures would correspond to their mean effects in monoculture. We fitted a linear model (termed the “no diversity effect model”) to these expected values (Appendix S1: Figure S5a). Secondly, we calculated expected biomass values assuming that the effect of mixed communities on biomass would correspond to the effect of the occurring strain that has the highest mean effect in monoculture – a pure positive selection effect. The “positive selection model” fitted on those expected values is a second-degree polynomial regression (Appendix S1: Figure S5b). Similarly, we built a “negative selection model” based on the occurring strain having the lowest associated biomass in monoculture – a pure negative selection effect (Appendix S1: Figure S5c). Predictions with higher values than the positive selection model would imply positive complementarity, while predictions lower than the negative selection model would imply negative complementarity. By plotting our predictions based on observed data (Appendix S1: Figure S5d) with these alternative models, we qualitatively assessed how the relative contribution of complementarity and selection varied with increasing strain richness.

## RESULTS

### *Methylobacterium* strain richness causes variation in *A. thaliana* leaf biomass

The relation between strain richness and leaf biomass was best modeled by a second-degree polynomial function (AICc = 1055.4, weight = 0.965) (Appendix S1: Table S4). Strain richness explained close to 8% of the variation in leaf biomass (adj. R^2^ *=* 0.078). We observed an initial decrease in dry leaf biomass at low strain richness (from an average of 28.0 mg for monocultures to 23.4 mg at four strains), but a gradual increase at higher richness (from 23.4 mg at four strains to 36.2 mg at ten strains) (Figure 1).

**Figure 1.**
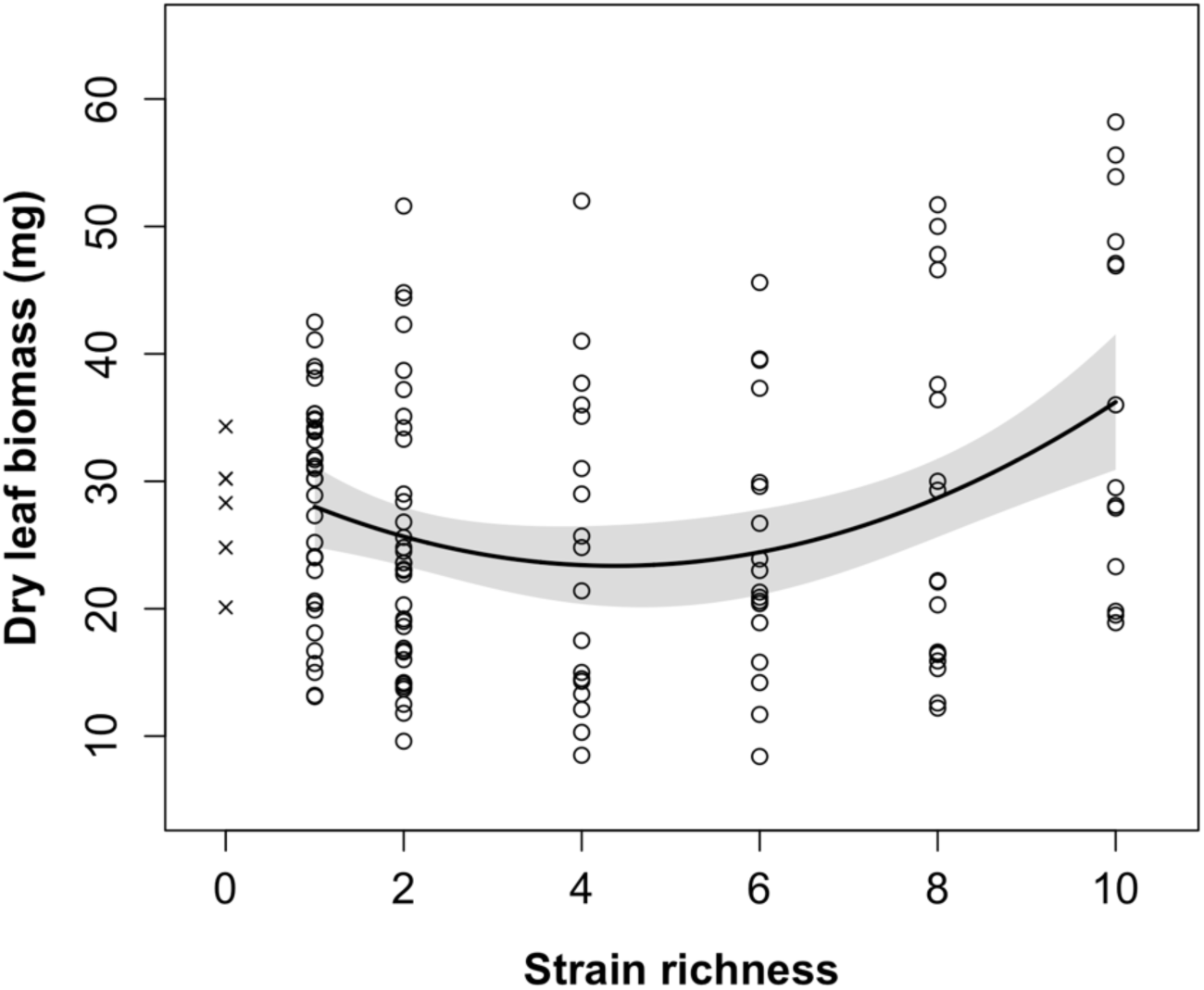
*A. thaliana* leaf biomass initially decreases with increasing taxonomic richness, but the relation becomes positive once *Methylobacterium* communities contain more than four strains. Solid line indicates fitted values of the models for the range of richness values used in the experiment. Shaded area indicates 95% confidence interval. *n*=137. “x” data points represent control plants; those samples were not used in the model but were included as a visual assessment of the effect of *Methylobacterium* inoculation on host growth.

### *Methylobacterium* strains influence *A. thaliana* leaf biomass

We identified three strains with a higher-than-average effect on *A. thaliana* leaf biomass: *Methylobacterium sp.* J-067, *Methylobacterium sp.* E-045, and *Methylobacterium sp.* J-092, henceforth referred to as “high-impact strains” (the other nine strains will be referred to as “low-impact strains”). All three high-impact strains were present in every model with a ΔAICc ≤ 2 (Table 1). The first model (ID 1) in Table 1 showed that, when the other two high-impact strains were absent (but notwithstanding the presence or absence of the low-impact strains), the presence of J-067 increased biomass by a mean of 5.8 mg (standard error: ± 2.0 mg); J-092 increased biomass by 8.1 (± 2.1) mg, while E-045 decreased biomass by 5.3 (± 2.1) mg. Based on the adjusted R² of the six best models, the prevalence of the three high-impact strains explained around 15.7% of the variation in leaf biomass. The results of all models explored are presented in Appendix S2: Table S1. Figure 2 illustrates that communities including J-067 and/or J-092 were often associated with an increased leaf biomass, while the ones containing E-045 tended to show relatively low leaf biomass. In monocultures, only the inoculation of strain J-092 caused an obvious increase in leaf biomass. Furthermore, the effect on plant growth of most monocultures and mixed communities was highly variable, including those containing high-impact strains.

**Table 1.** The presence of *Methylobacterium* sp. J-067 or J-092 in communities enhances *A. thaliana* growth, while E-045 has a detrimental effect. Parameters (intercept and strains) and their coefficients ± SE (mg) are shown for all models with a ΔAICc ≤ 2 (see Appendix S1: Table S3a for saturated model). Coefficients indicate, for a given strain in a given model, its positive or negative impact on host dry leaf biomass (mg) when the other strains included in the model are absent from communities, notwithstanding the presence or absent of strains not included in the model. Intercept indicates the mean biomass of samples when all strains included in the corresponding model are absent. *n*=137. ID: model number; SE: standard error; *df*: degrees of freedom; *LL*: log-likelihood; Δ: ΔAICc; wgt: model weight.

| ID | Intercept | E-046 | J-078 | J-088 | J-059 | J-043 | J-067 | J-048 | J-076 | E-045 | J-092 | E-005 | J-068 | df | LL | AICc | $\Delta$ | wgt | ajd. R<sup>2</sup> |
| --- | --- | --- | --- | --- | --- | --- | --- | --- | --- | --- | --- | --- | --- | --- | --- | --- | --- | --- | --- |
| 1 | 24.3<br>±1.3 |  |  |  |  |  | 5.8<br>±2.0 |  |  | -5.3<br>±2.1 | 8.1<br>±2.1 |  |  | 5 | -518.75 | 1048.0 | 0.00 | 0.035 | 0.153 |
| 2 | 24.8<br>±1.4 |  |  |  |  | -2.2<br>±2.0 | 6.0<br>±2.0 |  |  | -5.2<br>±2.1 | 8.7<br>±2.1 |  |  | 6 | -518.11 | 1048.9 | 0.92 | 0.022 | 0.161 |
| 3 | 24.5<br>±1.4 |  |  |  |  |  | 6.9<br>±2.3 | -2.4<br>±2.4 |  | -5.8<br>±2.2 | 9.0<br>±2.2 |  |  | 6 | -518.27 | 1049.2 | 1.22 | 0.019 | 0.159 |
| 4 | 24.7<br>±1.4 |  |  |  |  |  | 6.0<br>±2.0 |  | -1.5<br>±2.0 | -5.2<br>±2.1 | 8.1<br>±2.1 |  |  | 6 | -518.46 | 1049.6 | 1.61 | 0.016 | 0.156 |
| 5 | 24.5<br>±1.4 |  |  | -1.6<br>±2.2 |  |  | 6.0<br>±2.0 |  |  | -5.2<br>±2.1 | 8.7<br>±2.2 |  |  | 6 | -518.47 | 1049.6 | 1.62 | 0.015 | 0.156 |
| 6 | 24.0<br>±1.4 |  | 1.3<br>±2.0 |  |  |  | 5.6<br>±2.0 |  |  | -5.5<br>±2.2 | 7.9<br>±2.1 |  |  | 6 | -518.56 | 1049.8 | 1.80 | 0.014 | 0.155 |

**Figure 2.**
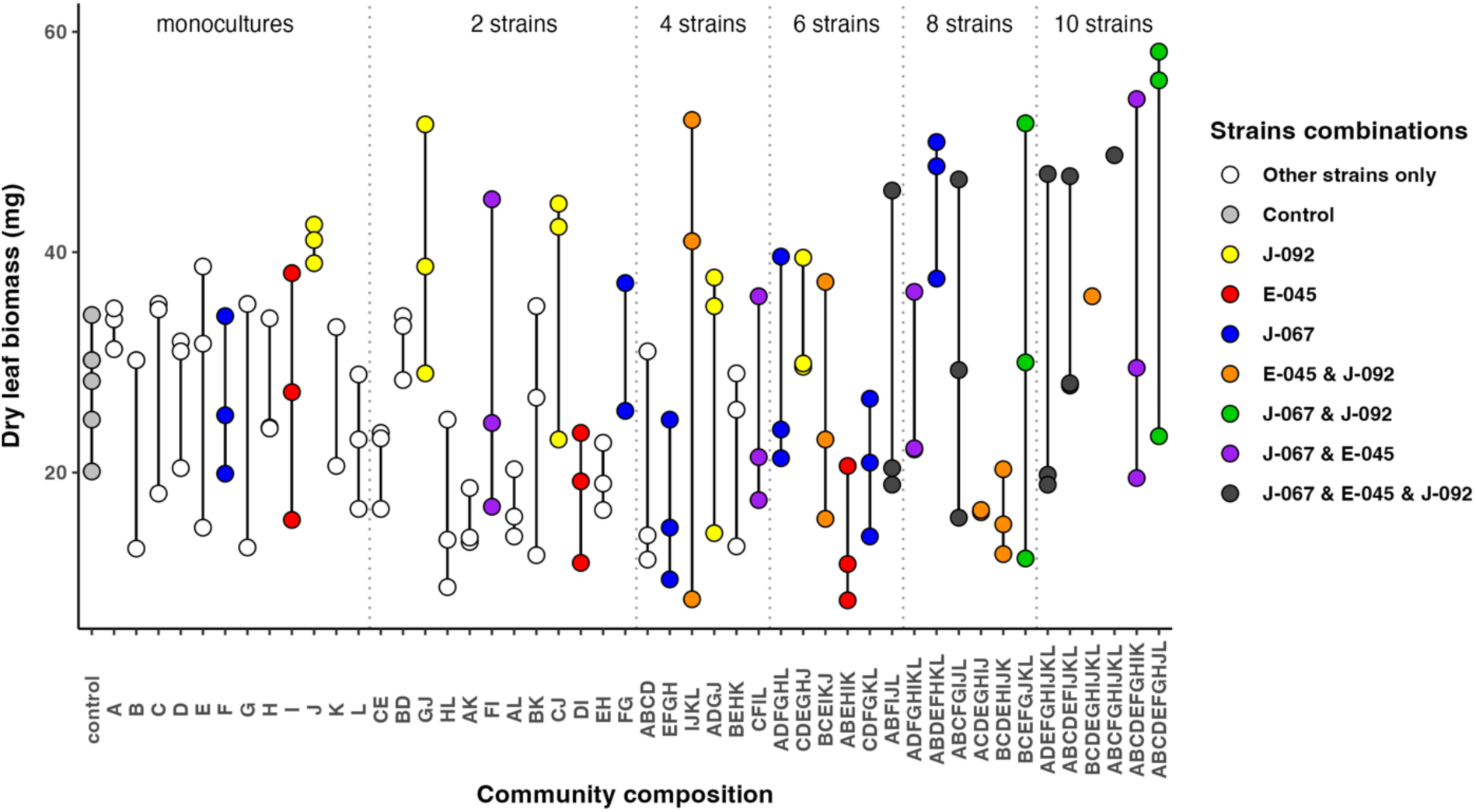
Dry leaf biomass of A. thaliana at the time of harvest in function of all the different compositions of synthetic communities. J-092 is the only strain in monoculture that cause an increase in leaf biomass. Most monocultures and mixed communities show high variability in their effect on plant growth. Dots show observed data (*n*=143) and are colored according to the different high-impact strains combinations (see legend on the right). Grey dotted lines separate community compositions according to strain richness. Letters on x axis refers to the strains composing the communities: A: E-046, B: J-078, C: J-088, D: J-059, E: J-043, F: J-067, G: J-048, H: J-076, I: E-045, J: J-092, K: E-005, L: J-068 (Appendix S1: Table S1).

### Interactions among strains and between strains and richness determined the biomass response to richness

To investigate the effect of specific strains on the relation between richness and leaf biomass, we explored all models that could be derived from a saturated model which included the three high-impact strains, the polynomial effect of richness and all possible interactions between those variables. The six best models (ΔAICc ≤ 2) all had a lower AICc (1032.4 to 1034.2) than our two previous hypotheses’ models and explained between 31.3% and 35.8% of leaf biomass variation based on their adjusted R^2^ (Appendix S1: Table S5a). The results of all models explored are presented in Appendix S3: Table S1.

Figure 3 shows predicted leaf biomass values in function of strain richness, under different composition scenarios, based on the averaged model (Appendix S1: Table S5b). The general response pattern of biomass with increasing richness (an initial decrease in biomass followed by an increase) was found throughout all high-impact strains combinations, as well as in communities inoculated with only low-impact strains. At high diversity values, communities containing J-067 (without E-045 nor J-092), and those containing J-067 and J-092 (without E-045), showed the steepest positive richness-biomass relationships. E-045 was neutral in monoculture but, when inoculated into mixed assemblages without J-067 nor J-092, provoked a steep decrease in the richness-biomass function. E-045 also diminished the strength of the relationship (the curve flattened) when communities harbored J-067 or J-092. The weakest relations between diversity and productivity occurred when only low-impact strains were inoculated, as well as when communities contained all three high-impact strains.

**Figure 3.**
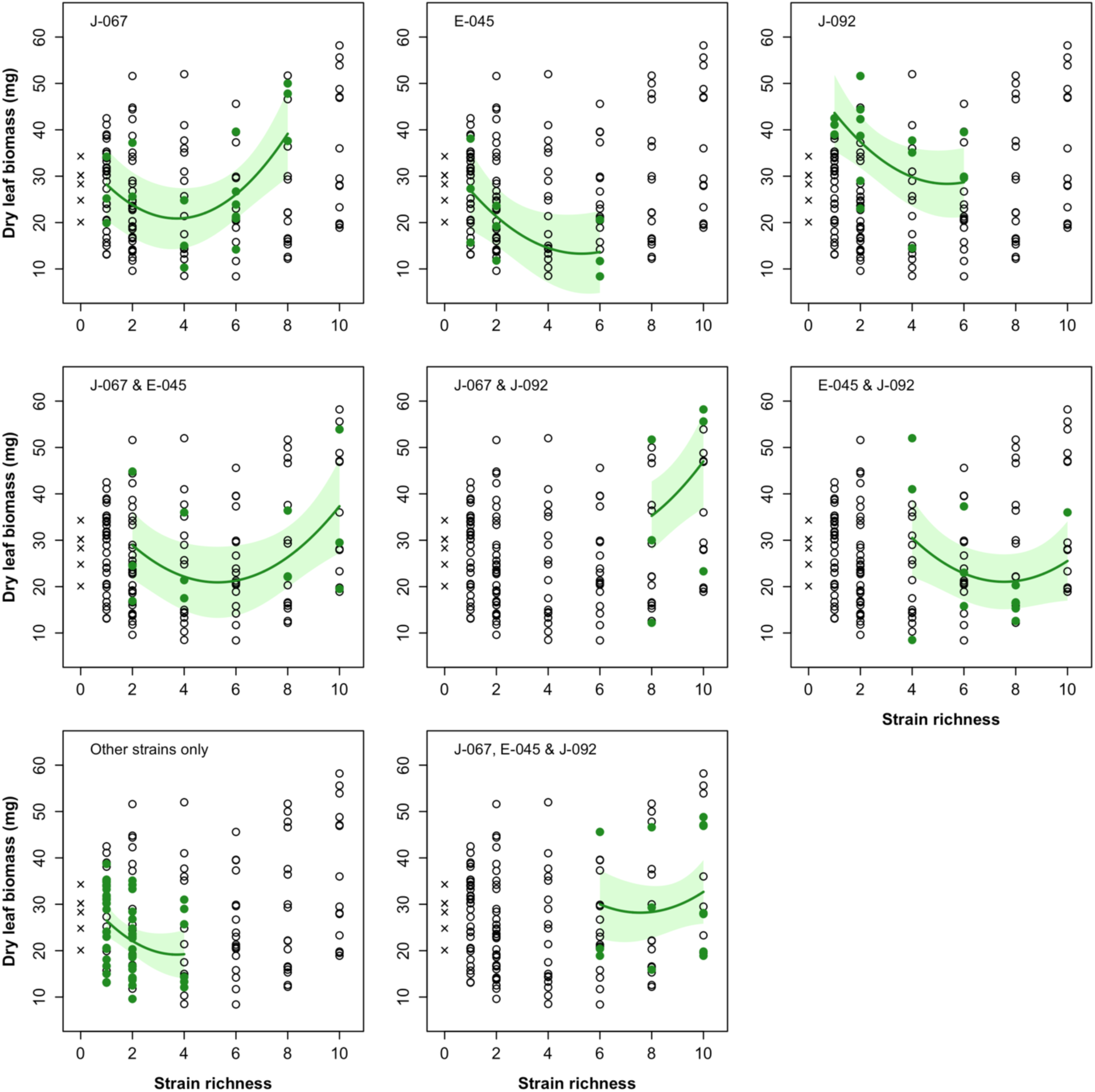
The effect of *Methylobacterium* taxonomic richness on *A. thaliana* leaf biomass is influenced by the presence (or absence) of strains J-067, E-045, and J-092. Model averaging was performed (Appendix S1: Table S5b) to reveal how richness and those three high-impact strains interact with each other to shape the observed diversity-productivity pattern. Eight panels illustrate the relation between strain richness and leaf biomass for samples (green points) containing a specific combination of strains (top left of each panels). Solid lines indicate fitted values of the averaged model based on fully averaged coefficients. Shaded areas indicate 95% confidence intervals. *n*=137. “x” data points represent control plants; those samples were not used in the model but were included in the figure as a visual assessment of the effect of *Methylobacterium* inoculation on host growth.

### The strain richness-leaf biomass relationship deviates from the alternative models

In communities composed of two to six *Methylobacterium* strains, our data-fitted model revealed that dry leaf biomass of *A. thaliana* was lower than expected if all strains had the same effect on biomass in mixed assemblages and in monocultures, given their relative abundance (Figure 4). At richnesses of two and four strains, the curve of the biomass response to richness was similar to the negative selection model predictions, but started to deviate from these predictions with a slope becoming positive beyond four strains. At a richness of eight strains, the observed biomass corresponded to the no diversity effect model, and when communities reached ten strains, most observed biomass values were higher than those expected by the no diversity effect model. The model we fitted on observed data did not match the positive selection model’s response curve at any richness values.

**Figure 4.**
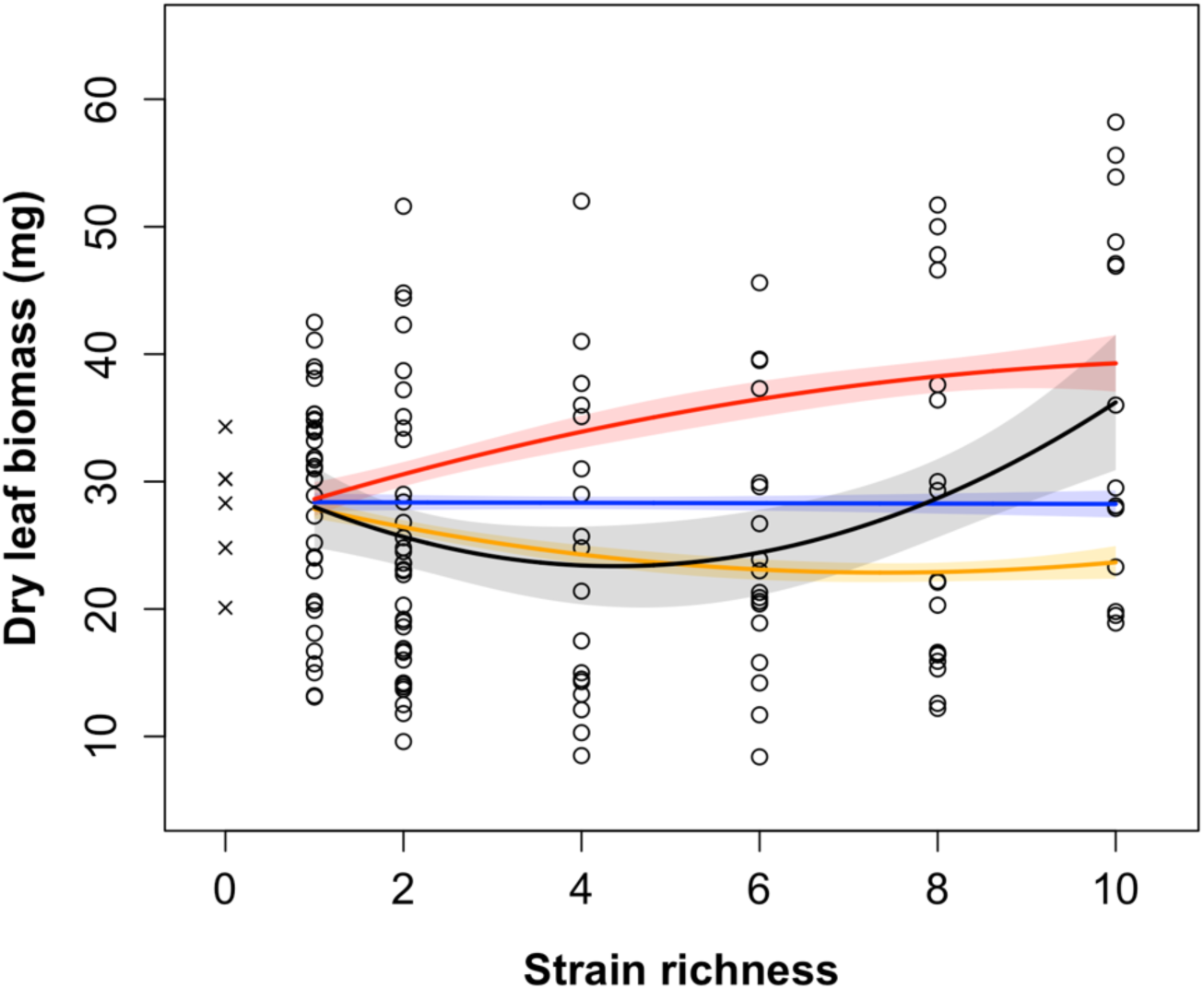
When two to six strains were inoculated, measured *A. thaliana* leaf biomass associated with mixed *Methylobacterium* communities was generally lower than expected if all strains were influencing plant growth in the same manner than in monocultures; however, ten-strains assemblages often had a higher than expected influence on plant growth. Except at a ten-strains richness, leaf biomass predicted by the data-fitted model was consistently lower than expected if mixed communities’ influence on plant growth was similar to the effect of the most growth-promoting strain in monoculture. Black lines represent predicted values of the regression model fitted on the observed data. Blue lines represent the no diversity effect model (Appendix S1: Figure S5a), red lines the positive selection model (Appendix S1: Figure S5b), and orange lines the negative selection model (Appendix S1: Figure S5c). Solid lines indicate fitted values of the different models for the range of richness values used in the experiment. Shaded areas indicate 95% confidence interval. *n*=137. “x” data points represent control plants; those samples were not used in the models but were included as a visual assessment of the effect of *Methylobacterium* inoculation on host growth.

## DISCUSSION

### Causality revealed: leaf bacterial diversity impacted individual plant productivity

We detected an effect of leaf bacterial diversity on plant growth in a fully controlled gnotobiotic experimental environment, indicating that foliar *Methylobacterium* taxonomic diversity causes variation in host plant productivity. However, we had predicted that functional redundancy among strains would result in a positive logarithmic relationship between inoculated strain richness and harvested leaf biomass, but the data portrayed an alternative pattern: increasing diversity caused an increase in plant growth, but only after an initial decrease at low strain richness compared with single strain effects. This pattern suggests an underlying ecological mechanism by which *Methylobacterium* diversity might have a small inherent negative effect on plant growth that can be overcome when diversity reaches a certain threshold. Direct or indirect inter-strain competition could explain this phenomenon. Schäfer et al. (2022) found that most bacterial pairwise interactions in the phyllosphere were negative, and Becker et al. (2012) demonstrated that increased strain richness in the rhizosphere could lead to a decrease in plant protection against pathogens. However, Grilli et al. (2017) demonstrated that multi-species interactions can stabilize competition dynamics. Likewise, by assembling synthetic bacterial communities in the zebrafish gut, Sundarraman et al. (2020) revealed that strong pairwise competition was reduced in communities containing four to five strains, resulting in higher-than-predicted cell abundance of every strain. In the light of those findings, we hypothesize that the observed initial negative effect of diversity on host biomass could be due to strong pairwise competition that ultimately reduced bacterial abundance and thus the potential for growth enhancement, for example through production of growth-producing chemical compounds such as phytohormones. Multi-species competition alleviation could then explain the diversity-driven increase in leaf biomass occurring when more than four strains were inoculated.

### *Methylobacterium* strains differed in their effect on plant growth and shaped bacterial diversity-host productivity relationship

*Methylobacterium* strains had varying influences on plant growth, validating our second hypothesis. We identified three strains exhibiting a high impact on *A. thaliana* vegetative growth. Two of these strains, *Methylobacterium* sp. J-067 and *Methylobacterium* sp. J-092, had a higher-than-average positive impact on leaf biomass, notwithstanding community diversity. However, we had not predicted to find a strain, *Methylobacterium* sp. E-045, with a global detrimental effect on its host, since *Methylobacterium* is generally considered a beneficial growth-promoting genus – although Flores-Núñez et al. (2023) identified a *Methylobacterium* strain reducing the growth of *Agave tequilana* in a field inoculation experiment. Furthermore, even if those three high-impact strains showed a general tendency to either be beneficial or detrimental to their host, a noticeable fraction of plant samples harboring these strains in monocultures or in mixed communities had a leaf biomass opposite to the expected effect of its symbionts. The fact that the effect of a distinct inoculated community composition on plant growth is highly variable hints at the importance of stochastic events in population and community dynamics. Indeed, Le Moigne et al. (2023) have shown that communities developing from an identical initial bacterial source can result in different compositions over time. This could in turn lead to a variation in community functions and impacts on the host plant.

The way bacterial diversity drove host productivity was shaped in part by which *Methylobacterium* taxa were present in the communities, thus supporting our third hypothesis. Interestingly, even in the presence of beneficial strains J-067 and J-092, the biodiversity-productivity function showed an initial negative relation, implying that pairwise microbial interactions can overcome and restrain beneficial bacteria-host symbioses. In addition, our prediction that high-impact strains would steepen the regression slope was validated. At high richness values, when the beneficial strains J-067 and J-092 were inoculated in mixed communities, the associated plant biomass increased faster with increasing diversity. At low richness values, the high-impact detrimental E-045 strain exacerbated the initial negative effect of diversity, in addition to neutralizing the effect of diversity on plant growth in rich assemblages. Furthermore, although the effects of J-067 and E-045 in monoculture were globally neutral, these strains behaved respectively as good and bad partners in mixed communities, corroborating a study by Garbeva et al. (2011) that showed that bacterial responses to competition are taxon-specific. This highlights the importance of considering how individual strains could interact with other bacteria in the phyllosphere before assessing their effects on the host. Furthermore, our finding that the biodiversity-ecosystem function relationship was dependent on composition in our study system was supported by a few studies showing the importance of composition on diversity-driven bacterial community functions (Bell et al. 2005; Jousset et al. 2014; Armitage 2016).

### The impact of *Methylobacterium* diversity on *A. thaliana* growth was plausibly driven by complementarity

Compared to most studies analyzing the effect of diversity of a group of organisms on the functions of these organisms, our experiment investigated the effect of symbiont diversity on the functioning of their host. Without being able to measure the growth of the inoculated bacterial strains, we could not apply the classical equation from Loreau & Hector (2001) to measure the relative contribution of complementarity and selection to the biodiversity effect. However, by analyzing how host growth responded to increasing bacterial diversity in a conceptual framework based on alternative models, we were able to qualitatively disentangle the mechanisms at play in different parts of the tested richness spectrum. The host growth response curve to diversity matched the negative selection model’s prediction curve for two and four-strain communities. This pattern could indicate negative species selection, although species selection alone could not explain the fact that the observed biomass-richness relationship clearly diverged from the negative selection model predictions when more than four strains were inoculated. If one or two particular host-detrimental strains were driving the biodiversity effect, it is difficult to conceive a diversity-dependent process that could relieve the bacterial assemblages of the dominance of these strains. On the other hand, complementarity, first through negative pairwise interactions then through positive multispecies competition alleviation, could explain the observed biomass response pattern to diversity at these relatively low richness values. In communities of six to ten strains, it was difficult to confidently separate complementarity versus selection effects, as the observed pattern did not fit any of the alternative selection models, nor did it go beyond their predictions to suggest complementarity. However, it is worth noting that the general trend in our data did not resemble at all the predictions of the positive selection model, ruling out the possibility of pure positive species selection driving the diversity-productivity relationship.

The detection of three high-impact strains might suggest a certain underlying effect of species selection since the presence of those specific strains was associated with higher or lower than average plant biomass across richness values. However, out of those three strains, only J-092 showed an effect in monoculture, suggesting that J-067 and E-045 effects on their host might be mostly due to how they interact with other partners (positive or negative complementarity) rather than due to their intrinsic growth-promoting or growth-inhibiting characteristics – for example J-067 might be a facilitator or community stabilizer while E-045 might be a strong competitor. Furthermore, the three strains associated with the lowest plant biomass in monocultures and thus driving the negative selection model’s predictions (J-068, J-048 and J-078) were not statistically associated with low biomass across the entire range of richness values. Finally, another observation supporting complementarity – and the importance of diversity *per se* – is the increase in productivity of plants when richness jumped from eight to ten strains, since all high-impact strains were already found interacting in communities at both of these diversity levels. Interestingly, only communities of ten strains showed a net positive biodiversity effect, and if the predicted biomass response to richness based on observed data was to be extrapolated over ten-strain richness, the values would be higher than expected by the positive selection model, indicating a positive complementarity effect. This highlighted that seemingly low-impact strains could actually be important actors driving diversity-productivity relationships. Overall, our results suggest that complementarity effect is most plausibly the key mechanism generally shaping the diversity-productivity relationship – a conclusion shared by Laforest-Lapointe et al. (2017) – although it is likely that both complementarity and selection act in richness- and taxa-dependent ways to drive the overall effects of *Methylobacterium* diversity on plant host productivity.

## Conclusion

Our study showed that increasing *Methylobacterium* diversity caused an increase in host vegetative growth, but only after reaching a certain richness threshold. The varying plant productivity response to bacterial biodiversity in our study highlights the fact that there is a need to better understand competition and coexistence dynamics in diverse microbial communities (Levine et al. 2017). Some *Methylobacterium* strains had beneficial or detrimental impacts on plant productivity, but their effect was highly variable in monocultures and in mixed assemblages. Strains also differed in their influence on the diversity-productivity relationship, their impact depending on associations with other strains. Finally, even if species selection likely contributes to explaining the net biodiversity effect we observed, our results suggest complementarity was the main ecological mechanism driving plant productivity in our experiment. What is especially interesting about our study is that even though some biodiversity-ecosystem functioning studies showed positive (Laforest-Lapointe et al. 2017; Bell et al. 2005), neutral (Jiang 2007) or negative (Becker et al. 2012) bacterial biodiversity effects, few, if any, have found that this relationship changes with varying values of microbial diversity. Our study emphasizes the importance of a better understanding of bacterial community dynamics in the phyllosphere and of how bacteria impact the productivity and function of plant hosts. This knowledge will be crucial to better understand plant-microbes symbioses in the context of global change (Perreault and Laforest-Lapointe 2022) – a topic of special interest for applied sciences like forestry, agriculture, and conservation.

## ACKNOWLEDGMENTS

We would like to thank François Ouellet for providing *A. thaliana* seeds and plant growth medium ingredients, as well as for his advice on growing *A. thaliana*; Geneviève Bourret for her help in the laboratory; Sylvain Dallaire for the maintenance of the growth chamber; Daniel Schönig for his help with the analyses; and Jean-Baptiste Leducq for his previous work on isolating and characterizing the *Methylobacterium* strains used in the experiment.

## AUTHOR CONTRIBUTIONS

Conceptualization and experimental design: J.L. and S.K.; laboratory experiment: J.L., E.F. and Z.W.; statistical analyses: J.L. and J.P.; manuscript: J.L., J.P. and S.K.; funding: S.K.

## SUPPORTING INFORMATION

Appendix S1: Tables S1-S5; Figures S1-S5

Appendix S2: Results for all models testing the effect of individual strains on leaf biomass

Appendix S3: Results for all models testing the effect of the three high-impact strains, strain richness, and all possible interactions

