## Appendix S1 for "Richness and composition of phyllosphere *Methylobacterium* communities cause variation in *Arabidopsis thaliana* growth"

**Table S1** – *Methylobacterium* strains used in experiment. ID: coding letters used in synthetic communities design and R script; forest: forest of isolation (MSH: Réserve Gault at Mont-Saint-Hilaire, QC, CAN; SBL: Station de Biologie des Laurentides, QC, CAN); host sp.: tree species from which the strain was isolated (ACSA: *Acer saccharum*; FAGR: *Fagus grandifolia*); sampling date: YY-MM-DD; T°C iso.: temperature at which the strain was isolated.

| ID | strain | clade[1] | forest | host sp. | sampling date | T°C iso. |
|---|---|---|---|---|---|---|
| A | E-046 | A | MSH | ACSA | 18-06-27 | 20 |
| B | J-078 | D | MSH | FAGR | 18-10-18 | 30 |
| C | J-088 | A | SBL | ACSA | 18-09-20 | 20 |
| D | J-059 | A | SBL | FAGR | 18-07-16 | 20 |
| E | J-043 | B | MSH | FAGR | 18-10-18 | 20 |
| F | J-067 | A | MSH | FAGR | 18-08-06 | 30 |
| G | J-048 | A | SBL | FAGR | 18-07-16 | 30 |
| H | J-076 | A | MSH | FAGR | 18-10-18 | 30 |
| I | E-045 | D | MSH | ACSA | 18-06-27 | 20 |
| J | J-092 | A | SBL | ACSA | 18-07-16 | 20 |
| K | E-005 | A | MSH | FAGR | 18-09-07 | 30 |
| L | J-068 | D | MSH | FAGR | 18-08-06 | 30 |

[1] based on Leducq et al., 2022

**Table S2** – Synthetic community compositions and experimental design. 48 distinct community compositions were constructed based on 12 strains. Each letter (A to L) represent a *Methylobacterium* strain (Appendix S1: Table S1).

| **Strain richness** | **1** | **2** | **4** | **6** | **8** | **10** |
|---|---|---|---|---|---|---|
| | A | CE | ABCD | ADFGHL | ADFGHIKL | ADEFGHIJKL |
| | B | BD | EFGH | CDEGHJ | ABDEFHKL | ABCDEFIJKL |
| | C | GJ | IJKL | BCEIKJ | ABCFGIJL | BCDEGHIJKL |
| | D | HL | ADGJ | ABEHIK | ACDEGHIJ | ABCFGHIJKL |
| | E | AK | BEHK | CDFGKL | BCDEHIJK | ABCDEFGHIK |
| **Community composition** | F | FI | CFIL | ABFIJL | BCEFGJKL | ABCDEFGHJL |
| | G | AL | | | | |
| | H | BK | | | | |
| | I | CJ | | | | |
| | J | DI | | | | |
| | K | EH | | | | |
| | L | FG | | | | |
| ***n*** | 12 | 12 | 6 | 6 | 6 | 6 |

**Table S3** – Saturated model equations used for model exploration in the context of our 2nd hypothesis (a) and 3rd hypothesis (b).

| **ID** | **Saturated global model** |
|---|---|
| a | dry leaf biomass ~ E-046 + J-078 + J-088 + J-059 + J-043 + J-067 + J-048 + J-076 + E-045 + J-092 + E-005 + J-068 |
| b | dry leaf biomass ~ strain richness + J-067 + E-045 + J-092 + strain richness:J-067 + strain richness:E-045 + strain richness:J-092 + J-067:E-045 + J-067:J-092 + E-045:J-092 + strain richness:J-067:E-045 + strain richness:J-067:J-092 + strain richness:E-045:J-092 + J-067:E-045:J-092 + strain richness:J-067:E-045:J-092. |

**Table S4** – The second-degree polynomial regression (model ID 1) best explains the relation between *Methylobacterium* strain diversity and *A. thaliana* vegetative growth, compared to a linear regression (model ID 2) and a null model. *n*=137. SE: standard error; *df*: degrees of freedom; *LL*: log-likelihood.

| **Model ID** | **Parameters** | **Coefficients ± SE (mg)** | ***df*** | ***LL*** | **AICc** | **ΔAICc** | **weight** | **adj.$R^2$** |
|---|---|---|---|---|---|---|---|---|
| 1 | - | - | 4 | -523.56 | 1055.4 | 0.00 | 0.965 | 0.0776 |
| | intercept | 31.17 ± 2.61 | | | | | | |
| | richness | -3.56 ± 1.38 | | | | | | |
| | richness^2 | 0.41 ± 0.13 | | | | | | |
| 2 | - | - | 3 | -528.31 | 1062.8 | 7.38 | 0.024 | 0.0187 |
| | intercept | 24.76 ± 1.65 | | | | | | |
| | richness | 0.60 ± 0.32 | | | | | | |
| null | - | - | 2 | -530.10 | 1064.3 | 8.88 | 0.011 | 0 |
| | intercept | 27.27 ± 0.99 | | | | | | |

**Table S5** – *Methylobacterium* sp. J-067, E-045, and J-092 interact among themselves, as well as with strain richness, to modulate *A. thaliana* growth response to bacterial diversity. (a) Models with a ΔAICc ≤ 2 (see Table S3b for saturated model). (b) Averaged model's full coefficients indicating the impact of diversity, strains, and their interactions on host biomass, to predict dry leaf biomass at different richness values and strain combinations. *n*=137. $D^2$ (in a): polynomial strain richness; $D^2$ and D (in b): respectively 2nd and 1st-degree terms in polynomial function of strain richness; "+" shows which parameters were selected in the different models; *df* : degrees of freedom; *LL* : log-likelihood; adj. SE: adjusted standard error.

a) Model exploration

| | **Parameters (strains)** | | | | | | | | | | | | | | | | | | | | |
|---|---|---|---|---|---|---|---|---|---|---|---|---|---|---|---|---|---|---|---|---|---|
| **ID** | **J-067** | **E-045** | **J-092** | **$D^2$** | **J-067:E-045** | **J-067:J-092** | **J-067:$D^2$** | **E-045:J-092** | **E-045:$D^2$** | **J-092:$D^2$** | **J-067:E-045:J-092** | **J-067:E-045:$D^2$** | **J-067:J-092:$D^2$** | **E-045:J-092:$D^2$** | **J-067:E-045:J-092:$D^2$** | ***df*** | ***LL*** | **AICc** | **ΔAICc** | **weight** | **ajd. $R^2$** |
| 1 | + | + | + | + | + | | | + | + | + | | | | | | 13 | -501.72 | 1032.4 | 0.00 | 0.110 | 0.339 |
| 2 | + | + | + | + | | | | | + | + | | | | | | 11 | -504.35 | 1032.8 | 0.41 | 0.089 | 0.313 |
| 3 | + | + | + | + | + | | | | + | + | | | | | | 12 | -503.27 | 1033.1 | 0.65 | 0.079 | 0.324 |
| 4 | + | + | + | + | + | | | + | + | + | | | | + | | 15 | -499.74 | 1033.5 | 1.05 | 0.065 | 0.358 |
| 5 | + | + | + | + | | | | + | + | + | | | | | | 12 | -503.69 | 1033.9 | 1.49 | 0.052 | 0.320 |
| 6 | + | + | + | + | + | + | | + | + | + | | | | | | 14 | -501.36 | 1034.2 | 1.76 | 0.045 | 0.343 |

b) Averaged model parameters and coefficients

| **Parameters** | **coefficients ± adj. SE (mg)** |
|---|---|
| Intercept | 32.92 ± 3.35 |
| J-067 | 1.73 ± 3.87 |
| E-045 | 1.10 ± 7.31 |
| J-092 | 18.55 ± 7.20 |
| D | -7.40 ± 2.26 |
| $D^2$ | 1.00 ± 0.28 |
| J-067:E-045 | 5.90 ± 5.86 |
| J-067:J-092 | -0.48 ± 2.33 |
| E-045:J-092 | 11.24 ± 19.35 |
| E-045:D | -0.47 ± 3.60 |
| E-045:$D^2$ | -0.26 ± 0.37 |
| J-092:D | -1.23 ± 3.60 |
| J-092:$D^2$ | -0.20 ± 0.36 |
| E-045:J-092:D | -2.26 ± 6.48 |
| E-045:J-092:$D^2$ | 0.20 ± 0.55 |

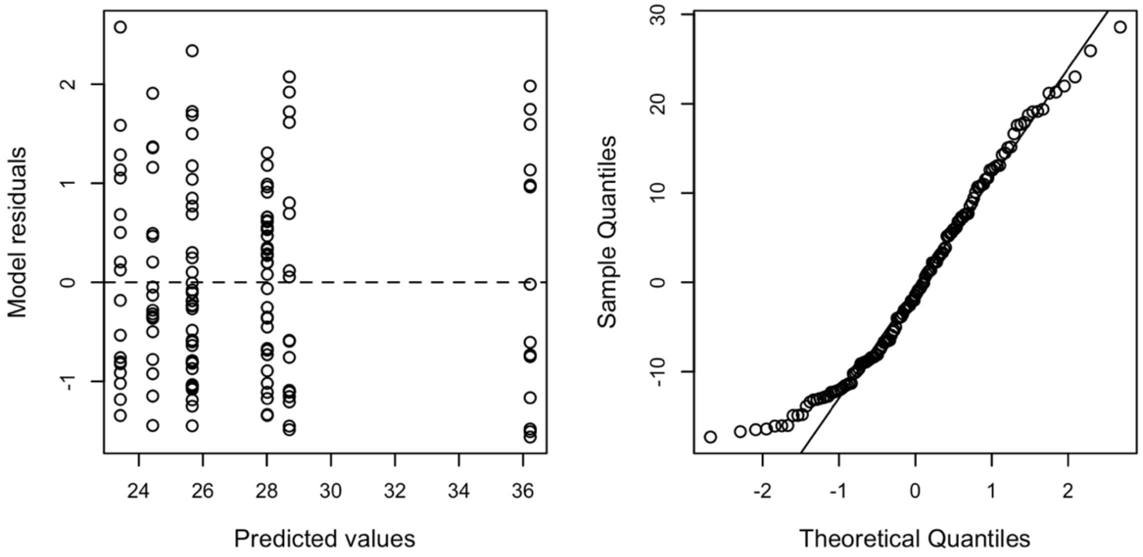


**Figure S1** – The assumptions of homoscedasticity and normality of residuals for the second-degree polynomial regression model are met (Model ID 1, Appendix S1: Table S4). The model can be used to draw inferences on the effect of *Methylobacterium* strain richness at the moment of inoculation on *A. thaliana* dry leaf biomass at the time of harvest. Left panel shows the homogeneity in the variance of scaled residuals across the response variable values. Right panel compares the model's residuals quantiles (*y* axis) to the quantiles of a normal distribution (*x* axis).

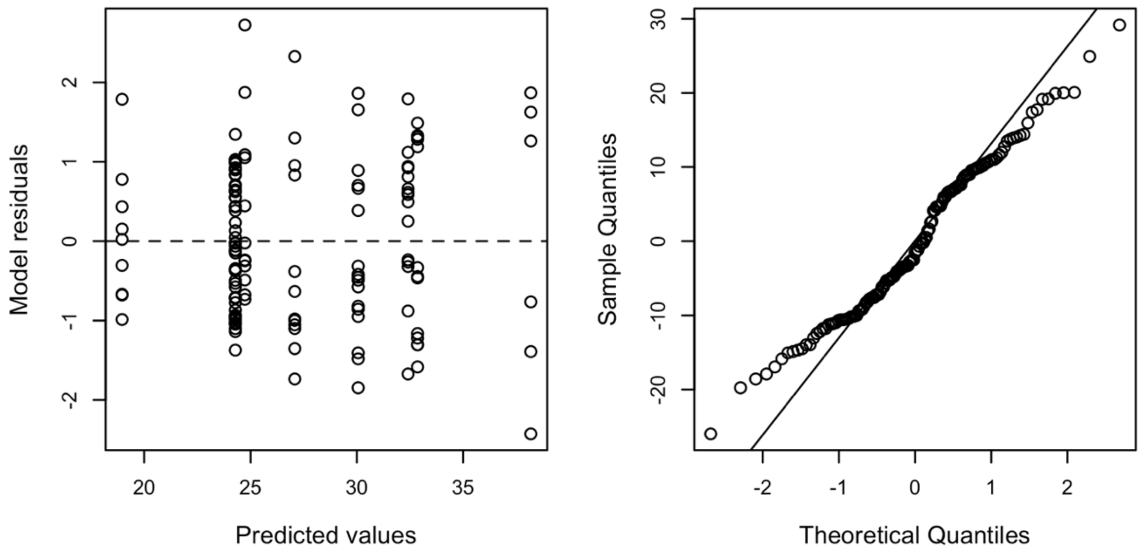


**Figure S2** – The assumptions of homoscedasticity and normality of residuals are met for the model ID 1 of Table 1 (leaf biomass ~ J-067 + E-045 + J-092) This model could be used to draw inferences on the effect of the included *Methylobacterium* strains on *A. thaliana* dry leaf biomass at the time of harvest. Left panel shows the homogeneity in the variance of scaled residuals across the response variable values. Right panel compares the model's residuals quantiles (*y* axis) to the quantiles of a normal distribution (*x* axis).

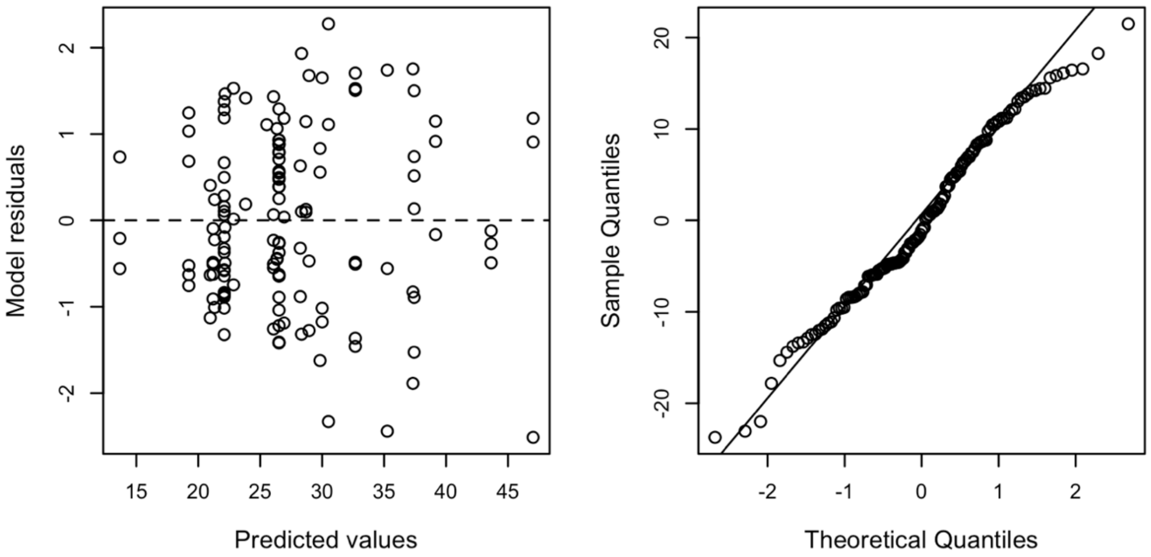


**Figure S3** – The assumptions of homoscedasticity and normality of residuals for the averaged model including polynomial strain richness, J-067 + E-045 + J-092, and some of their interactions (Appendix S1: Table S5b). This averaged model can be used to draw inferences on the effect of these three *Methylobacterium* strains and their interactions, on the effect of strain richness on *A. thaliana* dry leaf biomass at the time of harvest. Left panel shows the homogeneity in the variance of scaled residuals across the response variable values. Right panel compares the model's residuals quantiles (*y* axis) to the quantiles of a normal distribution (*x* axis).

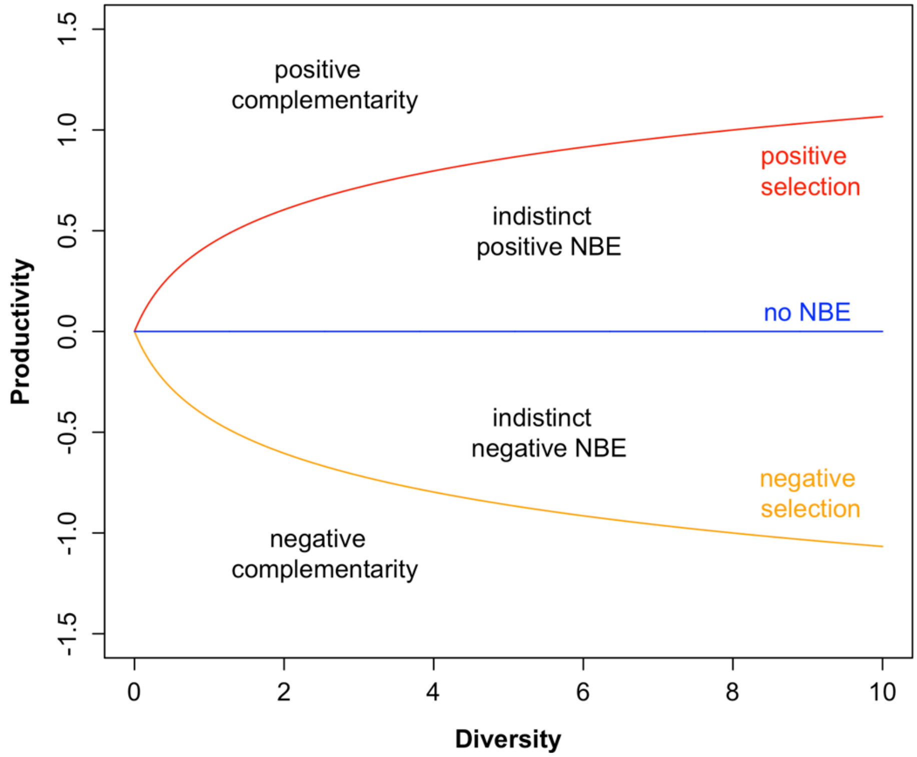


**Figure S4** – Conceptual framework to test whether complementarity or selection drives the diversity-productivity relationship. Predictions of a model built on observed data that would closely follow the positive selection (red) or the negative selection (orange) models predictions would be a strong argument in favor of positive or negative selection respectively. Measured biomass values higher than those predicted by the positive selection model would represent positive complementarity, while values under those predicted by the negative selection model would represent negative complementarity. Values between the mean and the positive selection model, and those between the mean and the negative selection model, would respectively represent indistinct positive or negative biodiversity effect. Values following the no diversity effect model (blue) would imply no net biodiversity effect (NBE).

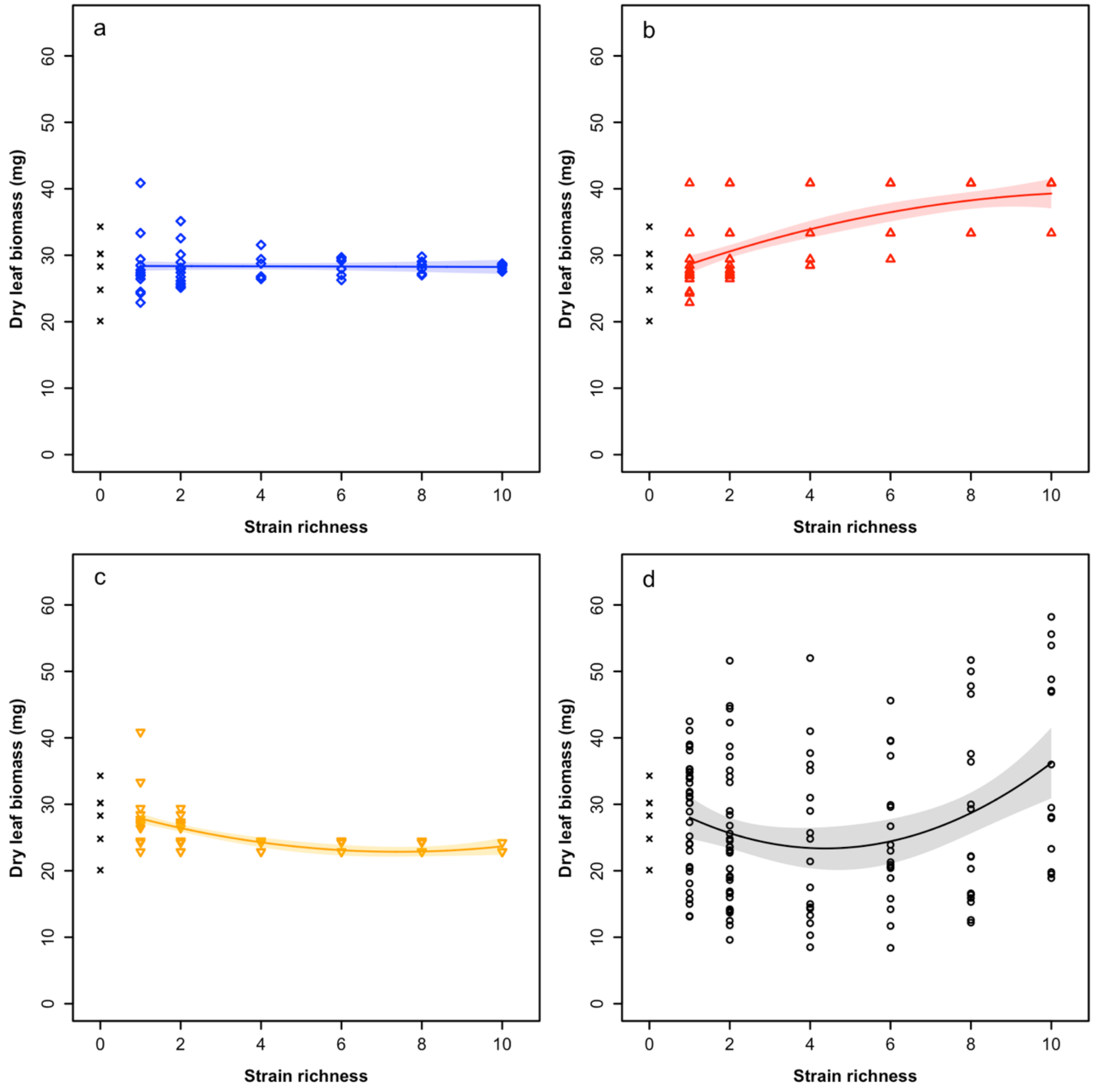


**Figure S5** – (a) Linear regression "no diversity effect model" ($y = -0.02x + 28.39$) representing the leaf biomass response to strain richness assuming that, for every mixed bacterial community composed of equally abundant inoculated strains, all strains' effect on biomass corresponds to their mean effect in monoculture. (b) Second-degree polynomial regression "positive selection model" ($y = -0.10x^2 + 2.24x + 26.47$) representing the biomass response to richness assuming that the effect of mixed communities on biomass corresponds to the effect of the occurring strain that has the highest mean effect in monoculture. (c) Second-degree polynomial regression "negative selection model" ($y = 0.12x^2 - 1.80x + 29.55$) representing the biomass response to richness assuming that the effect of mixed communities on biomass corresponds to the effect of the occurring strain that has the lowest mean effect in monoculture. (d) Second-degree polynomial regression model fitted on observed data ($y = 0.41x^2 - 3.56x + 31.17$). In all panels, solid lines indicate fitted values of the models for the range of richness values used in the experiment. Shaded areas indicate 95% confidence intervals. $n$=137. "x" data points represent control plants; those samples were not used in the models but were included as a visual assessment of the effect of *Methylobacterium* inoculation on host growth.
