## Appendix S2 for "Richness and composition of phyllosphere *Methylobacterium* communities cause variation in *Arabidopsis thaliana* growth"

**Table S1** – Parameters and statistical results for all models explored (*n* = 4096) in the context of our second hypothesis, based on the saturated model: dry leaf biomass ~ E-046 + J-078 + J-088 + J-059 + J-043 + J-067 + J-048 + J-076 + E-045 + J-092 + E-005 + J-068. Intercept indicates the mean biomass of samples when all strains included in the corresponding model are absent. ID: model number; +: parameters considered in the corresponding model; adj.R2: adjusted $R^2$; df: degrees of freedom; LL: log-likelihood.

(beginning on the next page)

models.H2

| ID | Intercept | E-046 | J-078 | J-088 | J-059 | J-043 | J-067 | J-048 | J-076 | E-045 | J-092 | E-005 | J-068 | adj.R2 | df | LL | AICc | ΔAICc | weight |
|---|---|---|---|---|---|---|---|---|---|---|---|---|---|---|---|---|---|---|---|
| 801 | 24,2851289472049 | NA | NA | NA | NA | NA | + | NA | NA | + | + | NA | NA | 0,152770250944867 | 5 | -518,751778824058 | 1047,96157291529 | 0 | 0.0347838502078085 |
| 817 | 24,7835695831331 | NA | NA | NA | NA | + | + | NA | NA | + | + | NA | NA | 0,160587766617823 | 6 | -518,117112947519 | 1048,88037974119 | 0,918806825901584 | 0.0219715798146781 |
| 865 | 24,5267289669618 | NA | NA | NA | NA | NA | + | + | NA | + | + | NA | NA | 0,158738207754769 | 6 | -518,267801491078 | 1049,18175682831 | 1,22018391301845 | 0.0188980973572024 |
| 929 | 24,6780337493185 | NA | NA | NA | NA | NA | + | NA | + | + | + | NA | NA | 0,156363663642873 | 6 | -518,460777285067 | 1049,56770841629 | 1,60613550099652 | 0.0155815177783647 |
| 805 | 24,5167653730408 | NA | NA | + | NA | NA | + | NA | NA | + | + | NA | NA | 0,156265294318751 | 6 | -518,468759903685 | 1049,58367365352 | 1,62210073823258 | 0.0154576315904029 |
| 803 | 24,0417650398524 | NA | + | NA | NA | NA | + | NA | NA | + | + | NA | NA | 0,15515748833508 | 6 | -518,558593603809 | 1049,76334105377 | 1,80176813848084 | 0.0141295611097913 |
| 2849 | 24,3479493491499 | NA | NA | NA | NA | NA | + | NA | NA | + | + | NA | + | 0,153331516414335 | 6 | -518,706407842647 | 1050,05896953145 | 2,09739661615754 | 0.012188036995195 |
| 819 | 24,510962772484 | NA | + | NA | NA | + | + | NA | NA | + | + | NA | NA | 0,166885777495989 | 7 | -517,601496508548 | 1050,07121007136 | 2,10963715606908 | 0.012113670721521 |
| 809 | 24,1899023315678 | NA | NA | NA | + | NA | + | NA | NA | + | + | NA | NA | 0,153061769276035 | 6 | -518,728217125893 | 1050,10258809794 | 2,14101518264943 | 0.0119251022749287 |
| 1825 | 24,3606196193321 | NA | NA | NA | NA | NA | + | NA | NA | + | + | + | NA | 0,15300802154772 | 6 | -518,73256184549 | 1050,11127753713 | 2,1497046218426 | 0.0118734034391806 |
| 802 | 24,3507012465142 | + | NA | NA | NA | NA | + | NA | NA | + | + | NA | NA | 0,152988597193207 | 6 | -518,734131953341 | 1050,11441775284 | 2,15284483754454 | 0.011854775542953 |
| 881 | 24,9680089756315 | NA | NA | NA | NA | + | + | + | NA | + | + | NA | NA | 0,165455838857388 | 7 | -517,718906228086 | 1050,30602951044 | 2,34445659514449 | 0.0107717278171317 |
| 2865 | 24,9353943001067 | NA | NA | NA | NA | + | + | NA | NA | + | + | NA | + | 0,162231034036783 | 7 | -517,982952863971 | 1050,83412278221 | 2,87254986691573 | 0.00827201615814629 |
| 821 | 24,8748377175022 | NA | NA | + | NA | + | + | NA | NA | + | + | NA | NA | 0,161930435038372 | 7 | -518,007514093501 | 1050,88324524127 | 2,92167232597558 | 0.00807132003155327 |
| 825 | 24,6271229859217 | NA | NA | NA | + | + | + | NA | NA | + | + | NA | NA | 0,161732997454348 | 7 | -518,023641458671 | 1050,91549997161 | 2,9539270563148 | 0.00794219492853549 |
| 869 | 24,7263368437626 | NA | NA | + | NA | NA | + | + | NA | + | + | NA | NA | 0,161636186671237 | 7 | -518,031547901549 | 1050,93131285736 | 2,96973994207178 | 0.0078796480056662 |
| 1841 | 24,7352289491656 | NA | NA | NA | NA | + | + | NA | NA | + | + | + | NA | 0,160817855430713 | 7 | -518,098343786766 | 1051,0649046278 | 3,10333171250568 | 0.0073705132803133 |
| 945 | 24,8374072438763 | NA | NA | NA | NA | + | + | NA | + | + | + | NA | NA | 0,160758675668157 | 7 | -518,103171781237 | 1051,07456061674 | 3,11298770144776 | 0.00733501424647221 |
| 818 | 24,8364193891437 | + | NA | NA | NA | + | + | NA | NA | + | + | NA | NA | 0,160739995857254 | 7 | -518,104695644102 | 1051,07760834247 | 3,11603542717694 | 0.00732384520285755 |
| 873 | 24,3125702511716 | NA | NA | NA | + | NA | + | + | NA | + | + | NA | NA | 0,160707125509554 | 7 | -518,107377061219 | 1051,0829711767 | 3,1213982614106 | 0.00730423322466742 |
| 807 | 24,2476945855311 | NA | + | + | NA | NA | + | NA | NA | + | + | NA | NA | 0,160386481611393 | 7 | -518,133528263265 | 1051,13527358079 | 3,17370066550279 | 0.00711569474453096 |
| 993 | 24,7339348451462 | NA | NA | NA | NA | NA | + | + | + | + | + | NA | NA | 0,159999528617728 | 7 | -518,165074240398 | 1051,19836553506 | 3,23679261976804 | 0.00689472684927768 |
| 933 | 24,9194820641939 | NA | NA | + | NA | NA | + | NA | + | + | + | NA | NA | 0,159970443753308 | 7 | -518,16744476957 | 1051,2031065934 | 3,24153367811323 | 0.00687840205498958 |
| 867 | 24,3375727965816 | NA | + | NA | NA | NA | + | + | NA | + | + | NA | NA | 0,159793575792109 | 7 | -518,181858427899 | 1051,23193391006 | 3,27036099477141 | 0.00677997020358092 |
| 2913 | 24,6213527950223 | NA | NA | NA | NA | NA | + | + | NA | + | + | NA | + | 0,159743239372756 | 7 | -518,185959984219 | 1051,2401370227 | 3,27856410741128 | 0.00675221872497437 |
| 931 | 24,4377000323524 | NA | + | NA | NA | NA | + | NA | + | + | + | NA | NA | 0,159308657201139 | 7 | -518,221360781047 | 1051,31093861636 | 3,34936570106697 | 0.00651736630930974 |
| 1889 | 24,6106157551883 | NA | NA | NA | NA | NA | + | + | NA | + | + | + | NA | 0,159024556920895 | 7 | -518,244493529932 | 1051,35720411413 | 3,39563119883655 | 0.00636833214199178 |
| 866 | 24,539849987258 | + | NA | NA | NA | NA | + | + | NA | + | + | NA | NA | 0,158748886010986 | 7 | -518,266932455473 | 1051,40208196521 | 3,44050904991946 | 0.00622702493125315 |
| 937 | 24,5367314054864 | NA | NA | NA | + | NA | + | NA | + | + | + | NA | NA | 0,157896240069917 | 7 | -518,336289221523 | 1051,54079549731 | 3,57922258201847 | 0.00580977540544683 |
| 813 | 24,3759510281905 | NA | NA | + | + | NA | + | NA | NA | + | + | NA | NA | 0,15707890445845 | 7 | -518,402707882587 | 1051,67363281944 | 3,71205990414637 | 0.00543643357207015 |
| 2853 | 24,5974478611863 | NA | NA | + | NA | NA | + | NA | NA | + | + | NA | + | 0,157030327856765 | 7 | -518,406653307517 | 1051,6815236693 | 3,71995075400741 | 0.00541502678871897 |
| 2977 | 24,732792756774 | NA | NA | NA | NA | NA | + | NA | + | + | + | NA | + | 0,156852088454736 | 7 | -518,42112808774 | 1051,71047322974 | 3,74890031445307 | 0.00533721001507099 |
| 806 | 24,6054110741201 | + | NA | + | NA | NA | + | NA | NA | + | + | NA | NA | 0,156619095252242 | 7 | -518,440044800091 | 1051,74830665445 | 3,78673373915444 | 0.005237196494364 |
| 1827 | 24,1609449878558 | NA | + | NA | NA | NA | + | NA | NA | + | + | + | NA | 0,156500888486429 | 7 | -518,449640007382 | 1051,76749706903 | 3,80592415373621 | 0.00518718482823481 |
| 1829 | 24,576810408944 | NA | NA | + | NA | NA | + | NA | NA | + | + | + | NA | 0,156427073635196 | 7 | -518,455631104592 | 1051,77947926345 | 3,81790634815661 | 0.00515620080650104 |
| 1953 | 24,6923070198444 | NA | NA | NA | NA | NA | + | NA | + | + | + | + | NA | 0,156378591019664 | 7 | -518,459565855039 | 1051,78734876434 | 3,82577584905175 | 0.00513595230559328 |
| 930 | 24,6852195102899 | + | NA | NA | NA | NA | + | NA | + | + | + | NA | NA | 0,156368913222645 | 7 | -518,460351258297 | 1051,78891957086 | 3,82734665556768 | 0.00513192009558243 |
| 804 | 24,1277914504488 | + | + | NA | NA | NA | + | NA | NA | + | + | NA | NA | 0,155724944671604 | 7 | -518,512592409857 | 1051,89340187398 | 3,93182895868745 | 0.00487070516928116 |
| 2851 | 24,1043114329489 | NA | + | NA | NA | NA | + | NA | NA | + | + | NA | + | 0,155677119536074 | 7 | -518,516470575931 | 1051,90115820613 | 3,93958529083488 | 0.00485185234655644 |
| 883 | 24,6973056085767 | NA | + | NA | NA | + | + | + | NA | + | + | NA | NA | 0,169459494281209 | 8 | -517,389664789623 | 1051,90432957925 | 3,94275666395561 | 0.00484416492602813 |
| 823 | 24,6092875854116 | NA | + | + | NA | + | + | NA | NA | + | + | NA | NA | 0,169401426355888 | 8 | -517,394451344672 | 1051,91390268934 | 3,95232977405271 | 0.0048210334681077 |
| 811 | 23,9780601153478 | NA | + | NA | + | NA | + | NA | NA | + | + | NA | NA | 0,155312155628793 | 7 | -518,54605846754 | 1051,96033398934 | 3,99876107405339 | 0.00471039923426431 |
| 889 | 24,7336984907855 | NA | NA | NA | + | + | + | + | NA | + | + | NA | NA | 0,168893629306031 | 8 | -517,436294949885 | 1051,99758989977 | 4,03601698447892 | 0.00462346633318844 |
| 2867 | 24,6688386667283 | NA | + | NA | NA | + | + | NA | NA | + | + | NA | + | 0,168795519831761 | 8 | -517,444376443113 | 1052,01375288623 | 4,0521799709345 | 0.004586252395957 |
| 545 | 23,469998397228 | NA | NA | NA | NA | NA | + | NA | NA | NA | + | NA | NA | 0,11299343679854 | 4 | -521,893011899873 | 1052,08905410278 | 4,1274811874855 | 0.00441678745120807 |
| 827 | 24,3746954316217 | NA | + | NA | + | + | + | NA | NA | + | + | NA | NA | 0,167811694296186 | 8 | -517,525363642286 | 1052,17572728457 | 4,21415436928078 | 0.00422946716725467 |
| 2929 | 25,1548568005133 | NA | NA | NA | NA | + | + | + | NA | + | + | NA | + | 0,167647142481252 | 8 | -517,538899985199 | 1052,2027999704 | 4,24122705510695 | 0.00417260139493865 |
| 820 | 24,6111764590408 | + | + | NA | NA | + | + | NA | NA | + | + | NA | NA | 0,167606353502275 | 8 | -517,542254949937 | 1052,20950989987 | 4,24793698458279 | 0.00415862592111291 |
| 2857 | 24,2509434661433 | NA | NA | NA | + | NA | + | NA | NA | + | + | NA | + | 0,153639834434281 | 7 | -518,681471564159 | 1052,23116018258 | 4,26958726729163 | 0.00411385099238311 |
| 810 | 24,2494076250295 | + | NA | NA | + | NA | + | NA | NA | + | + | NA | NA | 0,153528677412136 | 7 | -518,690462816436 | 1052,24914268714 | 4,2875697718448 | 0.00407702811026483 |
| 3873 | 24,4045074232384 | NA | NA | NA | NA | NA | + | NA | NA | + | + | + | + | 0,153483960108229 | 7 | -518,694079570043 | 1052,25637619435 | 4,29480327905912 | 0.004062309137634 |
| 2850 | 24,3904214451837 | + | NA | NA | NA | NA | + | NA | NA | + | + | NA | + | 0,153443347836374 | 7 | -518,697364141536 | 1052,26294533734 | 4,30137242204523 | 0.00404898808180075 |
| 1833 | 24,264868236332 | NA | NA | NA | + | NA | + | NA | NA | + | + | + | NA | 0,153366025963765 | 7 | -518,703617215465 | 1052,27545148519 | 4,31387856990273 | 0.00402374845485628 |
| 1826 | 24,4171262828717 | + | NA | NA | NA | NA | + | NA | NA | + | + | + | NA | 0,153197017435732 | 7 | -518,717283066268 | 1052,3027831868 | 4,34121027150832 | 0.00396913453161906 |
| 1843 | 24,5328639609467 | NA | + | NA | NA | + | + | NA | NA | + | + | + | NA | 0,16698170391344 | 8 | -517,59361295338 | 1052,31222590676 | 4,35065299146868 | 0.00395043898761105 |
| 947 | 24,5332777653151 | NA | + | NA | NA | + | + | NA | + | + | + | NA | NA | 0,166910434912873 | 8 | -517,599470165847 | 1052,32394033169 | 4,36236741640323 | 0.0039273680587454 |
| 885 | 25,0467720818186 | NA | NA | + | NA | + | + | + | NA | + | + | NA | NA | 0,166564618957182 | 8 | -517,627883794113 | 1052,38076758823 | 4,41919467293565 | 0.00381734772295379 |
| 1009 | 24,9249882783349 | NA | NA | NA | NA | + | + | + | + | + | + | NA | NA | 0,165616073458508 | 8 | -517,705759657194 | 1052,53651931439 | 4,57494639909669 | 0.00353134919108155 |
| 1905 | 24,9295533129808 | NA | NA | NA | NA | + | + | + | NA | + | + | + | NA | 0,165589279286219 | 8 | -517,707958181321 | 1052,54091636264 | 4,57934344735054 | 0.00352359396283631 |
| 882 | 24,9761124294338 | + | NA | NA | NA | + | + | + | NA | + | + | NA | NA | 0,165460076923606 | 8 | -517,718558545171 | 1052,56211709034 | 4,60054417505125 | 0.00348643985620326 |
| 561 | 24,0664336058463 | NA | NA | NA | NA | + | + | NA | NA | NA | + | NA | NA | 0,123241050996875 | 5 | -521,097415519013 | 1052,6528463052 | 4,69127338991052 | 0.00333181269793892 |
| 935 | 24,6674646720264 | NA | + | + | NA | NA | + | NA | + | + | + | NA | NA | 0,164891989741171 | 8 | -517,765147611531 | 1052,65529522306 | 4,6937223077723 | 0.00332773552679858 |
| 877 | 24,494811833413 | NA | NA | + | + | NA | + | + | NA | + | + | NA | NA | 0,164602374684813 | 8 | -517,788886873656 | 1052,70277374731 | 4,7412008320207 | 0.00324966784170378 |
| 871 | 24,4857752587194 | NA | + | + | NA | NA | + | + | NA | + | + | NA | NA | 0,163906028825531 | 8 | -517,845931531466 | 1052,81686306293 | 4,85529014764052 | 0.00306947989137418 |
| 2869 | 25,0309734679686 | NA | NA | + | NA | + | + | NA | NA | + | + | NA | + | 0,163627452544206 | 8 | -517,868739205352 | 1052,8624784107 | 4,90090549541333 | 0.0030002645162653 |
| 2873 | 24,7755461522552 | NA | NA | NA | + | + | + | NA | NA | + | + | NA | + | 0,163538147573994 | 8 | -517,87604919927 | 1052,87709839854 | 4,91552548324967 | 0.00297841256701322 |
| 829 | 24,7045401363036 | NA | NA | + | + | + | + | NA | NA | + | + | NA | NA | 0,163491689208427 | 8 | -517,879851706827 | 1052,88470341365 | 4,92313049836366 | 0.00296710863598124 |
| 997 | 24,9531915780154 | NA | NA | + | NA | NA | + | + | + | + | + | NA | NA | 0,163067258873868 | 8 | -517,914580558587 | 1052,95416111717 | 4,99258820188379 | 0.00286583312979671 |
| 1001 | 24,5424198625057 | NA | NA | NA | + | NA | + | + | + | + | + | NA | NA | 0,16301213994397 | 8 | -517,919089352716 | 1052,96317870543 | 5,00160579014187 | 0.00285294076455761 |
| 2917 | 24,8385963936012 | NA | NA | + | NA | NA | + | + | NA | + | + | NA | + | 0,162858609776678 | 8 | -517,931646738907 | 1052,98829347781 | 5,02672056252368 | 0.00281733928419204 |
| 3889 | 24,875874728525 | NA | NA | NA | NA | + | + | NA | NA | + | + | + | + | 0,162821164005518 | 8 | -517,934709116877 | 1052,99441823375 | 5,0328453184643 | 0.00280872472368579 |
| 941 | 24,7699374336727 | NA | NA | + | + | NA | + | NA | + | + | + | NA | NA | 0,16271106196551 | 8 | -517,943712653371 | 1053,01242530674 | 5,05085239145228 | 0.00278354977000589 |
| 949 | 24,9746068097189 | NA | NA | + | NA | + | + | NA | + | + | + | NA | NA | 0,162374741308653 | 8 | -517,971207772231 | 1053,06741554446 | 5,105842629172 | 0.00270805831724611 |
| 826 | 24,6998751875242 | + | NA | NA | + | + | + | NA | NA | + | + | NA | NA | 0,162353276694294 | 8 | -517,972962187628 | 1053,07092437526 | 5,10935145996518 | 0.00270331142326898 |

| | | | | | | | | | | | | | | | | | | |
|---|---|---|---|---|---|---|---|---|---|---|---|---|---|---|---|---|---|---|
| 953 | 24,6962396794505 | NA | NA | NA | + | + | + | NA | + | + | + | NA | NA | 0,162286438776152 | 8 | -517,978424913094 | 1053,08184982619 | 5,12027691089747 | 0.0026885842370251 |
| 2993 | 24,959732424419 | NA | NA | NA | NA | + | + | NA | + | + | + | NA | + | 0,162277366119429 | 8 | -517,979166396221 | 1053,08333279244 | 5,12175987715136 | 0.00268659143608342 |
| 2866 | 24,95201978382 | + | NA | NA | NA | + | + | NA | NA | + | + | NA | + | 0,162252039790314 | 8 | -517,98123620454 | 1053,08747240908 | 5,12589949378844 | 0.00268103645763398 |
| 822 | 24,9432673706063 | + | NA | + | NA | + | + | NA | NA | + | + | NA | NA | 0,162162603177996 | 8 | -517,98854496102 | 1053,10208992204 | 5,14051700674872 | 0.00266151284861186 |
| 1845 | 24,8305001877348 | NA | NA | + | NA | + | + | NA | NA | + | + | + | NA | 0,162109479875107 | 8 | -517,992885826131 | 1053,11077165226 | 5,14919873697204 | 0.00264998461968876 |
| 1831 | 24,3831559108926 | NA | + | + | NA | NA | + | NA | NA | + | + | + | NA | 0,1619872362272 | 8 | -518,002873678175 | 1053,13074735635 | 5,16917444105843 | 0.00262364870393825 |
| 1849 | 24,5879099183973 | NA | NA | NA | + | + | + | NA | NA | + | + | + | NA | 0,161911011820721 | 8 | -518,009100815576 | 1053,14320163115 | 5,18162871586128 | 0.00260736164646868 |
| 2921 | 24,4056494345903 | NA | NA | NA | + | NA | + | + | NA | + | + | NA | + | 0,161880792789298 | 8 | -518,011569396518 | 1053,14813879304 | 5,1865658777449 | 0.00260093310115659 |
| 1893 | 24,7952052493824 | NA | NA | + | NA | NA | + | + | NA | + | + | + | NA | 0,161843163549621 | 8 | -518,01464319014 | 1053,15428638028 | 5,19271346498999 | 0.00259295064407541 |
| 870 | 24,7589557298659 | + | NA | + | NA | NA | + | + | NA | + | + | NA | NA | 0,161693147592879 | 8 | -518,026896068765 | 1053,17879213753 | 5,21721922224015 | 0.00256137338577419 |
| 995 | 24,5421908577196 | NA | + | NA | NA | NA | + | + | + | + | + | NA | NA | 0,161556716512157 | 8 | -518,0380374705 | 1053,201074941 | 5,23950202570882 | 0.00253299447984501 |
| 808 | 24,3752029645034 | + | + | + | NA | NA | + | NA | NA | + | + | NA | NA | 0,161423849499873 | 8 | -518,048886077459 | 1053,22277215492 | 5,26119923962801 | 0.0025056635376799 |
| 875 | 24,1877193241279 | NA | + | NA | + | NA | + | + | NA | + | + | NA | NA | 0,161323582793904 | 8 | -518,05707172827 | 1053,23914345654 | 5,27757054124868 | 0.0024852367681711 |
| 1897 | 24,4119537835418 | NA | NA | NA | + | NA | + | + | NA | + | + | + | NA | 0,161220028124811 | 8 | -518,065524777665 | 1053,25604955533 | 5,29447664003987 | 0.00246431747946421 |
| 2855 | 24,3281406755882 | NA | + | + | NA | NA | + | NA | NA | + | + | NA | + | 0,161139900146968 | 8 | -518,072064817233 | 1053,26912963447 | 5,30755671917586 | 0.00244825333297533 |
| 815 | 24,130056269013 | NA | + | + | + | NA | + | NA | NA | + | + | NA | NA | 0,161010313920206 | 8 | -518,082640314089 | 1053,29028062818 | 5,32870771288754 | 0.00242249826391532 |
| 874 | 24,3593802097786 | + | NA | NA | + | NA | + | + | NA | + | + | NA | NA | 0,161010107132193 | 8 | -518,082657188698 | 1053,2903143774 | 5,32874146210543 | 0.00242245738554935 |
| 1842 | 24,7897263384696 | + | NA | NA | NA | + | + | NA | NA | + | + | + | NA | 0,160995395332074 | 8 | -518,083857711228 | 1053,29271542246 | 5,33114250716585 | 0.00241955091586977 |
| 1969 | 24,7892672891435 | NA | NA | NA | NA | + | + | NA | + | + | + | + | NA | 0,160992617562752 | 8 | -518,084084382328 | 1053,29316876466 | 5,33159584936561 | 0.00241900253575576 |
| 3041 | 24,8105906663146 | NA | NA | NA | NA | NA | + | + | + | + | + | NA | + | 0,160865199202871 | 8 | -518,09448114926 | 1053,31396229852 | 5,35238938323027 | 0.00239398301660614 |
| 946 | 24,8700629903367 | + | NA | NA | NA | + | + | NA | + | + | + | NA | NA | 0,160849964373729 | 8 | -518,095724137409 | 1053,31644827482 | 5,35487535952711 | 0.00239100917269824 |
| 1891 | 24,4314093074172 | NA | + | NA | NA | NA | + | + | NA | + | + | + | NA | 0,160807440409701 | 8 | -518,099193517534 | 1053,32338703507 | 5,36181411977782 | 0.00238272822617678 |
| 2915 | 24,4357389255339 | NA | + | NA | NA | NA | + | + | NA | + | + | NA | + | 0,160715050132202 | 8 | -518,106730615268 | 1053,33846123054 | 5,37688831524474 | 0.00236483687977434 |
| 2981 | 24,9912115061942 | NA | NA | + | NA | NA | + | NA | + | + | + | NA | + | 0,160651834744165 | 8 | -518,111887199772 | 1053,34877439954 | 5,3872014842525 | 0.00235267378552929 |
| 673 | 23,9901387367373 | NA | NA | NA | NA | NA | + | NA | + | NA | + | NA | NA | 0,118739022465902 | 5 | -521,448079124098 | 1053,35417351537 | 5,39260060008087 | 0.00234633117137107 |
| 939 | 24,32101496378 | NA | + | NA | + | NA | + | NA | + | + | + | NA | NA | 0,160558405748915 | 8 | -518,119507647803 | 1053,36401529561 | 5,40244238031596 | 0.00233481349534829 |
| 2017 | 24,7702016596899 | NA | NA | NA | NA | NA | + | + | + | + | + | + | NA | 0,160090722748565 | 8 | -518,1576410334 | 1053,4402820668 | 5,47870915150952 | 0.00224745536836974 |
| 549 | 23,7702623517341 | NA | NA | + | NA | NA | + | NA | NA | NA | + | NA | NA | 0,118166271396746 | 5 | -521,492562342078 | 1053,44313995133 | 5,48156703604127 | 0.00224424617782328 |
| 934 | 24,9407839578096 | + | NA | + | NA | NA | + | NA | + | + | + | NA | NA | 0,160009518097567 | 8 | -518,164260040148 | 1053,4535200803 | 5,49194716500574 | 0.00223262856959268 |
| 994 | 24,7253268284532 | + | NA | NA | NA | NA | + | + | + | + | + | NA | NA | 0,160007395737781 | 8 | -518,164433025527 | 1053,45386605105 | 5,49229313576348 | 0.00223224239089629 |
| 1955 | 24,4947399663145 | NA | + | NA | NA | NA | + | NA | + | + | + | + | NA | 0,159973766591718 | 8 | -518,167173949512 | 1053,45934789902 | 5,49777498373396 | 0.00222613236157438 |
| 1957 | 24,9225239794699 | NA | NA | + | NA | NA | + | NA | + | + | + | + | NA | 0,159971199920232 | 8 | -518,167383140074 | 1053,45976628015 | 5,49819336485712 | 0.00222566672440067 |
| 3937 | 24,6803056667745 | NA | NA | NA | NA | NA | + | + | NA | + | + | + | + | 0,15990743069491 | 8 | -518,172580296668 | 1053,47016059334 | 5,50858767804471 | 0.00221412959201875 |
| 868 | 24,3674332441509 | + | + | NA | NA | NA | + | + | NA | + | + | NA | NA | 0,159906422414071 | 8 | -518,172662467832 | 1053,47032493566 | 5,50875202037309 | 0.0022139476618873 |
| 2914 | 24,6100276867235 | + | NA | NA | NA | NA | + | + | NA | + | + | NA | + | 0,159753826573264 | 8 | -518,185097329046 | 1053,49519465809 | 5,53362174280119 | 0.00218658798938343 |
| 2979 | 24,4916169175534 | NA | + | NA | NA | NA | + | NA | + | + | + | NA | + | 0,159748630470533 | 8 | -518,185520713799 | 1053,4960414276 | 5,53446851230751 | 0.00218566241731765 |
| 1569 | 23,8747714545239 | NA | NA | NA | NA | NA | + | NA | NA | NA | + | + | NA | 0,117817367234638 | 5 | -521,519646140568 | 1053,49730754831 | 5,53573463302018 | 0.00218427919896546 |
| 932 | 24,4653644863643 | + | + | NA | NA | NA | + | NA | + | + | + | NA | NA | 0,159436652131075 | 8 | -518,210936293198 | 1053,5468725864 | 5,58529967110553 | 0.0021308125141589 |
| 1890 | 24,6181113268109 | + | NA | NA | NA | NA | + | + | NA | + | + | + | NA | 0,159028739234754 | 8 | -518,244153043389 | 1053,61330608678 | 5,65173317148674 | 0.00206119645754428 |
| 2985 | 24,5916874300696 | NA | NA | NA | + | NA | + | NA | + | + | + | NA | + | 0,158406040379638 | 8 | -518,294828984576 | 1053,71465796915 | 5,75308505386192 | 0.00195934586980862 |
| 938 | 24,5674100918155 | + | NA | NA | + | NA | + | NA | + | + | + | NA | NA | 0,158089279514792 | 8 | -518,320592965359 | 1053,76618593072 | 5,80461301542732 | 0.00190951006152415 |
| 891 | 24,5236295759427 | NA | + | NA | + | + | + | + | NA | + | + | NA | NA | 0,172004143152263 | 9 | -517,179579524727 | 1053,7764818841 | 5,81490896880882 | 0.00189970520747047 |
| 814 | 24,4738782278677 | + | NA | + | + | NA | + | NA | NA | + | + | NA | NA | 0,158020499377578 | 8 | -518,32618596877 | 1053,77737193754 | 5,81579902224871 | 0.00189885997598238 |
| 1961 | 24,5574286093552 | NA | NA | NA | + | NA | + | NA | + | + | + | + | NA | 0,157930957551633 | 8 | -518,333466568961 | 1053,79193313792 | 5,8303602226315 | 0.0018850853403314 |
| 2861 | 24,4547932177607 | NA | NA | + | + | NA | + | NA | NA | + | + | NA | + | 0,157898154927693 | 8 | -518,336133539926 | 1053,79726707985 | 5,83569416456135 | 0.00188006457056188 |
| 2931 | 24,8850274592948 | NA | + | NA | NA | + | + | + | NA | + | + | NA | + | 0,171754451845889 | 9 | -517,200222479324 | 1053,81776779329 | 5,85619487800227 | 0.00186089167046294 |
| 2937 | 24,9243309878054 | NA | NA | NA | + | + | + | + | NA | + | + | NA | + | 0,171614040318882 | 9 | -517,211828115993 | 1053,84097906663 | 5,87940615134085 | 0.00183941967693848 |
| 887 | 24,7709026163822 | NA | + | + | NA | + | + | + | NA | + | + | NA | NA | 0,171531385515239 | 9 | -517,218658983509 | 1053,85464080166 | 5,8930678863735 | 0.00182689766162273 |
| 2871 | 24,7738589450389 | NA | + | + | NA | + | + | NA | NA | + | + | NA | + | 0,171421282472783 | 9 | -517,227757206731 | 1053,87283724811 | 5,91126433281624 | 0.00181035152326221 |
| 1837 | 24,4427532547588 | NA | NA | + | + | NA | + | NA | NA | + | + | + | NA | 0,157329063400375 | 8 | -518,382386202389 | 1053,88977240478 | 5,92819948948681 | 0.00179508694812304 |
| 2854 | 24,6571123874425 | + | NA | + | NA | NA | + | NA | NA | + | + | NA | + | 0,157227022827739 | 8 | -518,390676203916 | 1053,90635240783 | 5,94477949254042 | 0.0017802671873748 |
| 3877 | 24,6366468348927 | NA | NA | + | NA | NA | + | NA | NA | + | + | + | + | 0,157111153244435 | 8 | -518,40008848854 | 1053,92517697708 | 5,96360406178928 | 0.00176358941698558 |
| 1828 | 24,246587406658 | + | + | NA | NA | NA | + | NA | NA | + | + | + | NA | 0,15706514977095 | 8 | -518,403825070809 | 1053,93265014162 | 5,97107722632836 | 0.00175701191637834 |
| 546 | 23,6913189172813 | + | NA | NA | NA | NA | + | NA | NA | NA | + | NA | NA | 0,114955524068769 | 5 | -521,741394421124 | 1053,94080410942 | 5,97923119413372 | 0.00174986318960251 |
| 2978 | 24,7278910214455 | + | NA | NA | NA | NA | + | NA | + | + | + | NA | + | 0,156855317744702 | 8 | -518,42086586506 | 1053,96673173012 | 6,00515881482966 | 0.00172732470306146 |
| 4001 | 24,736049762625 | NA | NA | NA | NA | NA | + | NA | + | + | + | + | + | 0,156853007554264 | 8 | -518,421053455727 | 1053,96710691145 | 6,00553399616297 | 0.00172700070345948 |
| 831 | 24,4489776010791 | NA | + | + | + | + | + | NA | NA | + | + | NA | NA | 0,170825834088296 | 9 | -517,276940389124 | 1053,97120361289 | 6,00963069760382 | 0.00172346682087544 |
| 3875 | 24,2005023994603 | NA | + | NA | NA | NA | + | NA | NA | + | + | + | + | 0,156803597790988 | 8 | -518,425065474026 | 1053,97513094805 | 6,01355803276124 | 0.00172008582561812 |
| 1830 | 24,6554420141798 | + | NA | + | NA | NA | + | NA | NA | + | + | + | NA | 0,156748854807494 | 8 | -518,429510269547 | 1053,98402053909 | 6,02244762380315 | 0.0017124573618846 |
| 1835 | 24,0854348271603 | NA | + | NA | + | NA | + | NA | NA | + | + | + | NA | 0,156741123017695 | 8 | -518,430138020357 | 1053,98527604071 | 6,02370312542416 | 0.00171138270273208 |
| 769 | 25,7709296077378 | NA | NA | NA | NA | NA | NA | NA | NA | + | + | NA | NA | 0,100513293995182 | 4 | -522,84961977992 | 1054,00226986287 | 6,04069694757868 | 0.00169690284025712 |
| 893 | 24,810332667007 | NA | NA | + | + | + | + | + | NA | + | + | NA | NA | 0,17062577371688 | 9 | -517,293457166419 | 1054,00423716748 | 6,04266425219384 | 0.00169523449853117 |
| 824 | 24,7385683989937 | + | + | + | NA | + | + | NA | NA | + | + | NA | NA | 0,170460589342221 | 9 | -517,307091616487 | 1054,03150606762 | 6,06993315232808 | 0.00167227776525922 |
| 1954 | 24,6989042982578 | + | NA | NA | NA | NA | + | NA | + | + | + | + | NA | 0,15638333587273 | 8 | -518,459180782394 | 1054,04336156479 | 6,08178864949832 | 0.00166239424555299 |
| 812 | 24,0392371648195 | + | + | NA | + | NA | + | NA | NA | + | + | NA | NA | 0,15617173833765 | 8 | -518,476351059073 | 1054,07770211815 | 6,11612920285506 | 0.00163409413207696 |
| 2852 | 24,1666474647121 | + | + | NA | NA | NA | + | NA | NA | + | + | NA | + | 0,156070151909274 | 8 | -518,484592854383 | 1054,09418570877 | 6,13261279347444 | 0.00162068161027608 |
| 609 | 23,53643918616 | NA | NA | NA | NA | NA | + | + | NA | NA | + | NA | NA | 0,113955998510839 | 5 | -521,818673257738 | 1054,09536178265 | 6,13378886736041 | 0.00161972886976718 |
| 2593 | 23,5453743316119 | NA | NA | NA | NA | NA | + | NA | NA | NA | + | NA | + | 0,113755045109812 | 5 | -521,834199553665 | 1054,12641437451 | 6,16484145921595 | 0.00159477470418989 |
| 2875 | 24,5279848639102 | NA | + | NA | + | + | + | NA | NA | + | + | NA | + | 0,169873921807831 | 9 | -517,355493699424 | 1054,12831023349 | 6,16673731820242 | 0.00159326368649499 |
| 2859 | 24,0387705386085 | NA | + | NA | + | NA | + | NA | NA | + | + | NA | + | 0,155844711307571 | 8 | -518,50287950514 | 1054,13075901028 | 6,16918609498839 | 0.00159131410669527 |
| 884 | 24,7485652361116 | + | + | NA | NA | + | + | + | NA | + | + | NA | NA | 0,169742590562193 | 9 | -517,366324293986 | 1054,14997142262 | 6,18839850732661 | 0.00157610080335983 |
| 547 | 23,3433710994083 | NA | + | NA | NA | NA | + | NA | NA | NA | + | NA | NA | 0,113531578301457 | 5 | -521,851461175025 | 1054,16093761723 | 6,19936470193443 | 0.00156748253827912 |
| 1011 | 24,6515994467522 | NA | + | NA | NA | + | + | + | + | + | + | NA | NA | 0,16963593245022 | 9 | -517,375118890401 | 1054,16756061545 | 6,20598770015681 | 0.00156230040629515 |
| 1847 | 24,6465857841256 | NA | + | + | NA | + | + | NA | NA | + | + | + | NA | 0,169631855921103 | 9 | -517,375455002098 | 1054,16823283884 | 6,20665992355157 | 0.00156177538709126 |
| 951 | 24,683429423895 | NA | + | + | NA | + | + | NA | + | + | + | NA | NA | 0,169624343483124 | 9 | -517,376074401739 | 1054,16947163812 | 6,20789872283262 | 0.00156080832350825 |
| 1907 | 24,7147100712092 | NA | + | NA | NA | + | + | + | NA | + | + | + | NA | 0,169526785106929 | 9 | -517,384117569954 | 1054,18555797455 | 6,22398505926299 | 0.00154830483086476 |

| | | | | | | | | | | | | | | | | | | |
|---|---|---|---|---|---|---|---|---|---|---|---|---|---|---|---|---|---|---|
| 828 | 24,4681354183328 | + | + | NA | + | + | + | NA | NA | + | + | NA | NA | 0,169330895354168 | 9 | -517,400264784717 | 1054,21785240408 | 6,25627948878969 | 0.00152350478492156 |
| 890 | 24,7930895560791 | + | NA | NA | + | + | + | + | NA | + | + | NA | NA | 0,169324139066421 | 9 | -517,400821638392 | 1054,21896611143 | 6,25739319613854 | 0.001522656651849 |
| 553 | 23,4524134666119 | NA | NA | NA | + | NA | + | NA | NA | NA | + | NA | NA | 0,113002469754912 | 5 | -521,892314659495 | 1054,24264458617 | 6,28107167087524 | 0.00150473585194492 |
| 2868 | 24,7275186913722 | + | + | NA | NA | + | + | NA | NA | + | + | NA | + | 0,169152764263642 | 9 | -517,414944847294 | 1054,24721252923 | 6,2856396139432 | 0.00150130299985344 |
| 1913 | 24,7118448447042 | NA | NA | NA | + | + | + | + | NA | + | + | + | NA | 0,16894534248448 | 9 | -517,432034837856 | 1054,28139251036 | 6,31981959506697 | 0.00147586374324566 |
| 1017 | 24,7296500229213 | NA | NA | NA | + | + | + | + | + | + | + | NA | NA | 0,168895901187068 | 9 | -517,436107798751 | 1054,28953843215 | 6,32796551685601 | 0.00146986483289042 |
| 2995 | 24,6604087676752 | NA | + | NA | NA | + | + | NA | + | + | + | NA | + | 0,168800047033771 | 9 | -517,444003548506 | 1054,30532993166 | 6,34375701636623 | 0.00145830484543697 |
| 2933 | 25,2369835641892 | NA | NA | + | NA | + | + | + | NA | + | + | NA | + | 0,168798944363693 | 9 | -517,444094372942 | 1054,30551158053 | 6,34393866523942 | 0.00145817240173565 |
| 3891 | 24,6698106999758 | NA | + | NA | NA | + | + | NA | NA | + | + | + | + | 0,16879593309399 | 9 | -517,444342403793 | 1054,30600764223 | 6,34443472694124 | 0.00145781077484324 |
| 563 | 23,847523739706 | NA | + | NA | NA | + | + | NA | NA | NA | + | NA | NA | 0,126507814602706 | 6 | -520,841838273843 | 1054,32983039384 | 6,36825747854937 | 0.00144054925118999 |
| 3057 | 25,0951004557155 | NA | NA | NA | NA | + | + | + | + | + | + | NA | + | 0,168156613985145 | 9 | -517,496981214715 | 1054,41128526408 | 6,44971234878471 | 0.00138305805610281 |
| 3953 | 25,0992535273693 | NA | NA | NA | NA | + | + | + | NA | + | + | + | + | 0,168120933654562 | 9 | -517,499917789661 | 1054,41715841397 | 6,45558549867724 | 0.00137900256001451 |
| 955 | 24,4226019766298 | NA | + | NA | + | + | + | NA | + | + | + | NA | NA | 0,168016294551362 | 9 | -517,508529107541 | 1054,43438104973 | 6,47280813443786 | 0.00136717851398933 |
| 2858 | 24,2911890815315 | + | NA | NA | + | NA | + | NA | NA | + | + | NA | + | 0,15394183837315 | 8 | -518,657037154812 | 1054,43907430962 | 6,47750139433265 | 0.00136397401330411 |
| 1851 | 24,3974313565252 | NA | + | NA | + | + | + | NA | NA | + | + | + | NA | 0,167935421322342 | 9 | -517,51518386148 | 1054,44769055761 | 6,48611764231487 | 0.00135811048360951 |
| 3881 | 24,3083683810599 | NA | NA | NA | + | NA | + | NA | NA | + | + | + | + | 0,153846395606753 | 8 | -518,664760140583 | 1054,45452028117 | 6,49294736587512 | 0.0013534806336738 |
| 1834 | 24,3197291126695 | + | NA | NA | + | NA | + | NA | NA | + | + | + | NA | 0,153808146727235 | 8 | -518,667854898201 | 1054,4607097964 | 6,49913688111178 | 0.00134929841398754 |
| 2817 | 25,165134921531 | NA | NA | NA | NA | NA | NA | NA | NA | + | + | NA | + | 0,111534079706204 | 5 | -522,005564397234 | 1054,46914406164 | 6,50757114635212 | 0.00134362022488498 |
| 2930 | 25,1297436108658 | + | NA | NA | NA | + | + | + | NA | + | + | NA | + | 0,167709876237661 | 9 | -517,533739703475 | 1054,48480224159 | 6,52322932630432 | 0.00133314197236323 |
| 1844 | 24,6322081829312 | + | + | NA | NA | + | + | NA | NA | + | + | + | NA | 0,167697559262891 | 9 | -517,534752889795 | 1054,48682861424 | 6,52525569894533 | 0.00133179193518862 |
| 3874 | 24,4427468298493 | + | NA | NA | NA | NA | + | NA | NA | + | + | + | + | 0,153584517300765 | 8 | -518,685946195549 | 1054,4968923911 | 6,53531947580836 | 0.00132510733893248 |
| 2609 | 24,2468082622659 | NA | NA | NA | NA | + | + | NA | NA | NA | + | NA | + | 0,125422197334175 | 6 | -520,926878039029 | 1054,49990992421 | 6,53833700892119 | 0.00132310956875923 |
| 948 | 24,6038965034295 | + | + | NA | NA | + | + | NA | + | + | + | NA | NA | 0,167610313444226 | 9 | -517,541929244999 | 1054,50118132464 | 6,53960840935315 | 0.00132226873500802 |
| 565 | 24,1894194127824 | NA | NA | + | NA | + | + | NA | NA | NA | + | NA | NA | 0,125363356746692 | 6 | -520,93148418988 | 1054,50912222591 | 6,54754931062371 | 0.00131702914092721 |
| 1005 | 24,7674067487548 | NA | NA | + | + | NA | + | + | + | + | + | NA | NA | 0,167495533482119 | 9 | -517,551369260488 | 1054,52006135562 | 6,55848844033062 | 0.00130984522897275 |
| 943 | 24,527725181372 | NA | + | + | + | NA | + | NA | + | + | + | NA | NA | 0,16744725479737 | 9 | -517,555339525982 | 1054,52800188661 | 6,56642897131815 | 0.0013046551055596 |
| 562 | 24,2604876668387 | + | NA | NA | NA | + | + | NA | NA | NA | + | NA | NA | 0,124887854248143 | 6 | -520,968696050907 | 1054,58354594797 | 6,62197303267749 | 0.0012689206893507 |
| 1971 | 24,5528560999649 | NA | + | NA | NA | + | + | NA | + | + | + | + | NA | 0,167002343105485 | 9 | -517,591916636644 | 1054,60115610793 | 6,63958319264361 | 0.0012577967864851 |
| 1585 | 24,2177808369358 | NA | NA | NA | NA | + | + | NA | NA | NA | + | + | NA | 0,124626911301875 | 6 | -520,989108329262 | 1054,62437050468 | 6,6627975893864 | 0.00124328169242582 |
| 1909 | 25,0123251994681 | NA | NA | + | NA | + | + | + | NA | + | + | + | NA | 0,166664683873277 | 9 | -517,619663270286 | 1054,65664937522 | 6,69507645992644 | 0.00122337688626123 |
| 886 | 25,0673093911384 | + | NA | + | NA | + | + | + | NA | + | + | NA | NA | 0,166589047221401 | 9 | -517,625877056606 | 1054,66907694786 | 6,70750403256648 | 0.00121579865284638 |
| 1013 | 25,0270661599683 | NA | NA | + | NA | + | + | + | + | + | + | NA | NA | 0,166587903404084 | 9 | -517,625971020431 | 1054,66926487551 | 6,70769196021774 | 0.00121568441712086 |
| 677 | 24,2987682065518 | NA | NA | + | NA | NA | + | NA | + | NA | + | NA | NA | 0,124024453981662 | 6 | -521,036212370677 | 1054,71857858751 | 6,7570056722177 | 0.00118607598695392 |
| 999 | 24,7313608884183 | NA | + | + | NA | NA | + | + | + | + | + | NA | NA | 0,166218425199043 | 9 | -517,656316663821 | 1054,72995616229 | 6,76838324699793 | 0.00117934780856873 |
| 879 | 24,3149047985469 | NA | + | + | + | NA | + | + | NA | + | + | NA | NA | 0,166215333552346 | 9 | -517,656570527393 | 1054,73046388943 | 6,7688909741405 | 0.00117904845312159 |
| 2925 | 24,6083073140918 | NA | NA | + | + | NA | + | + | NA | + | + | NA | + | 0,166103360754946 | 9 | -517,665764286188 | 1054,74885140702 | 6,78727849173015 | 0.00116825824333286 |
| 625 | 24,1077819907569 | NA | NA | NA | NA | + | + | + | NA | NA | + | NA | NA | 0,123813141482286 | 6 | -521,052726489578 | 1054,75160682531 | 6,79003391001925 | 0.00116664983148409 |
| 689 | 24,1670721837262 | NA | NA | NA | NA | + | + | NA | + | NA | + | NA | NA | 0,12378627230733 | 6 | -521,054826036315 | 1054,75580591878 | 6,79423300349276 | 0.00116420296519137 |
| 569 | 23,9585205715726 | NA | NA | NA | + | + | + | NA | NA | NA | + | NA | NA | 0,123739212839346 | 6 | -521,058503089542 | 1054,76316002524 | 6,80158710994624 | 0.00115992998971524 |
| 2033 | 24,8889209652247 | NA | NA | NA | NA | + | + | + | + | + | + | + | NA | 0,165740933935395 | 9 | -517,695513634134 | 1054,80835010291 | 6,8467771876235 | 0.00113401520189236 |
| 1959 | 24,7347854198937 | NA | + | + | NA | NA | + | NA | + | + | + | + | NA | 0,1657218960128 | 9 | -517,69707598087 | 1054,81147479639 | 6,84990188109441 | 0.00113224486024758 |
| 1010 | 24,9404005504576 | + | NA | NA | NA | + | + | + | + | + | + | NA | NA | 0,165637404343812 | 9 | -517,704009358016 | 1054,82534155068 | 6,86376863538635 | 0.00112442173130087 |
| 1906 | 24,9402877266161 | + | NA | NA | NA | + | + | + | NA | + | + | + | NA | 0,165597665697575 | 9 | -517,707270064359 | 1054,83186296336 | 6,87029004807232 | 0.00112076129327937 |
| 2983 | 24,7386240055434 | NA | + | + | NA | NA | + | NA | + | + | + | NA | + | 0,165552688531689 | 9 | -517,710960423822 | 1054,83924368229 | 6,87767076699993 | 0.00111663290354042 |
| 2877 | 24,8589859674091 | NA | NA | + | + | + | + | NA | NA | + | + | NA | + | 0,165392244510745 | 9 | -517,724123175769 | 1054,86556918618 | 6,90399627089209 | 0.00110203125139937 |
| 878 | 24,5772049927799 | + | NA | + | + | NA | + | + | NA | + | + | NA | NA | 0,165296936160016 | 9 | -517,73194103066 | 1054,88120489596 | 6,91963198067401 | 0.00109344932084428 |
| 936 | 24,721959519267 | + | + | + | NA | NA | + | NA | + | + | + | NA | NA | 0,165272406077369 | 9 | -517,733953014561 | 1054,88522886377 | 6,92365594847774 | 0.00109125153011442 |
| 1895 | 24,5950172154409 | NA | + | + | NA | NA | + | + | NA | + | + | + | NA | 0,16516372571231 | 9 | -517,742866384764 | 1054,90305560417 | 6,94148268888353 | 0.00108156802170105 |
| 1901 | 24,5853926767383 | NA | NA | + | + | NA | + | + | NA | + | + | + | NA | 0,165043536964444 | 9 | -517,752722259329 | 1054,9227673533 | 6,9611944380124 | 0.0010709605816113 |
| 2919 | 24,5985982004597 | NA | + | + | NA | NA | + | + | NA | + | + | NA | + | 0,165037134378915 | 9 | -517,753247252722 | 1054,92381734009 | 6,96224442479934 | 0.0010703984819435 |
| 957 | 24,8305958208044 | NA | NA | + | + | + | + | NA | + | + | + | NA | NA | 0,164760999623022 | 9 | -517,775885671616 | 1054,96909417788 | 7,00752126258612 | 0.00104643858311422 |
| 1573 | 24,1337393295698 | NA | NA | + | NA | NA | + | NA | NA | NA | + | + | NA | 0,122408629199281 | 6 | -521,162388381592 | 1054,97093060934 | 7,00935769404668 | 0.00104547816774816 |
| 513 | 24,9033333333334 | NA | NA | NA | NA | NA | NA | NA | NA | NA | + | NA | NA | 0,0796930196936916 | 3 | -524,41635579295 | 1055,01316271372 | 7,0515897984294 | 0.00102363324712738 |
| 830 | 24,8030456097494 | + | NA | + | + | + | + | NA | NA | + | + | NA | NA | 0,164440738394741 | 9 | -517,802132353995 | 1055,02158754264 | 7,06001462734503 | 0.00101933034879596 |
| 872 | 24,5490468519232 | + | + | + | NA | NA | + | + | NA | + | + | NA | NA | 0,16430392879049 | 9 | -517,813341380697 | 1055,04400559604 | 7,08243268074966 | 0.00100796844460262 |
| 3893 | 24,9729044433702 | NA | NA | + | NA | + | + | NA | NA | + | + | + | + | 0,164137218236263 | 9 | -517,826997760546 | 1055,07131835574 | 7,10974544044689 | 0.00099429680964337 |
| 3045 | 25,0456669421089 | NA | NA | + | NA | NA | + | + | + | + | + | NA | + | 0,1641314064542 | 9 | -517,827473793495 | 1055,07227042163 | 7,1106975063442 | 0.000993823604240855 |
| 1697 | 24,2370687889011 | NA | NA | NA | NA | NA | + | NA | + | NA | + | + | NA | 0,1217516399884 | 6 | -521,213624758764 | 1055,07340336368 | 7,11183044839208 | 0.000993260791389777 |
| 3897 | 24,7242131852347 | NA | NA | NA | + | + | + | NA | NA | + | + | + | + | 0,16405477160154 | 9 | -517,833750511441 | 1055,08482385753 | 7,12325094223684 | 0.00098760518982099 |
| 1003 | 24,4021587468139 | NA | + | NA | + | NA | + | + | + | + | + | NA | NA | 0,164027732405025 | 9 | -517,835964998525 | 1055,08925283169 | 7,1276799164043 | 0.000985420570682508 |
| 3049 | 24,6205517031586 | NA | NA | NA | + | NA | + | + | + | + | + | NA | + | 0,164018445444669 | 9 | -517,836725576123 | 1055,09077398689 | 7,12920107160176 | 0.000984671366821076 |
| 2997 | 25,0930955113137 | NA | NA | + | NA | + | + | NA | + | + | + | NA | + | 0,163847053138218 | 9 | -517,850760638855 | 1055,11884411236 | 7,15727119706571 | 0.000970947972042865 |
| 3001 | 24,8183396778216 | NA | NA | NA | + | + | + | NA | + | + | + | NA | + | 0,163843098606222 | 9 | -517,851084435699 | 1055,11949170604 | 7,15791879075368 | 0.000970633633047576 |
| 2874 | 24,8155869373688 | + | NA | NA | + | + | + | NA | NA | + | + | NA | + | 0,16383795380072 | 9 | -517,851505689776 | 1055,1203342142 | 7,15876129890717 | 0.000970224835782435 |
| 2870 | 25,059485390038 | + | NA | + | NA | + | + | NA | NA | + | + | NA | + | 0,163682797327643 | 9 | -517,864208606472 | 1055,14574004759 | 7,18416713229931 | 0.000957978099831831 |
| 1853 | 24,6711699552193 | NA | NA | + | + | + | + | + | + | + | + | + | NA | 0,163611337902509 | 9 | -517,870058315407 | 1055,15743946546 | 7,19586655016883 | 0.000952390565440221 |
| 2989 | 24,8434193474843 | NA | NA | + | + | NA | + | NA | + | + | + | NA | + | 0,163465983392673 | 9 | -517,881955576464 | 1055,18123398757 | 7,21966107228263 | 0.000941126862720496 |
| 1571 | 23,6853153696877 | NA | + | NA | NA | NA | + | NA | NA | NA | + | + | NA | 0,120865908676286 | 6 | -521,282639357402 | 1055,21143256096 | 7,24985964566758 | 0.000927023263504074 |
| 942 | 24,8317316365338 | + | NA | + | + | NA | + | NA | + | + | + | NA | NA | 0,163246887418634 | 9 | -517,899884669731 | 1055,21709217411 | 7,25551925881678 | 0.000924403675190419 |
| 2025 | 24,5911900270657 | NA | NA | NA | + | NA | + | + | + | + | + | + | NA | 0,163201682358547 | 9 | -517,903583312783 | 1055,22448946021 | 7,26291654492047 | 0.000920990951074736 |
| 1002 | 24,5700474682443 | + | NA | NA | + | NA | + | + | + | + | + | NA | NA | 0,163168851383145 | 9 | -517,906269392612 | 1055,22986161987 | 7,26828870457939 | 0.000918520415371414 |
| 771 | 25,3227390764367 | NA | + | NA | NA | NA | NA | NA | NA | + | + | NA | NA | 0,106545650407894 | 5 | -522,388905041666 | 1055,23582535051 | 7,27425243521679 | 0.000915785590666164 |
| 2021 | 24,9759152798648 | NA | NA | + | NA | NA | + | + | + | + | + | + | NA | 0,163107666384576 | 9 | -517,911274987532 | 1055,23987280971 | 7,27829989441852 | 0.000913934162285371 |
| 998 | 24,9573699549747 | + | NA | + | NA | + | + | + | + | + | + | NA | NA | 0,163068798997645 | 9 | -517,914454570365 | 1055,24623197538 | 7,28465906008546 | 0.000911032847841145 |
| 1832 | 24,5115706252342 | + | + | + | NA | NA | + | NA | NA | + | + | + | NA | 0,163033339434756 | 9 | -517,917355244236 | 1055,25203332312 | 7,29046040782737 | 0.000908394067634435 |
| 3941 | 24,8812173844362 | NA | NA | + | NA | NA | + | + | NA | + | + | + | + | 0,16295286606509 | 9 | -517,923937695467 | 1055,26519822558 | 7,30362531028936 | 0.000902434244627098 |

| | | | | | | | | | | | | | | | | | | |
|---|---|---|---|---|---|---|---|---|---|---|---|---|---|---|---|---|---|---|
| 550 | 24,0243804826378 | + | NA | + | NA | NA | + | NA | NA | NA | + | NA | NA | 0,120515653538753 | 6 | -521,30991143767 | 1055,26597672149 | 7,30440380620234 | 0.000902083042298338 |
| 816 | 24,2382069749267 | + | + | + | + | NA | + | NA | NA | + | + | NA | NA | 0,162902560224755 | 9 | -517,928052222894 | 1055,27342728043 | 7,31185436514261 | 0.000898728782517179 |
| 4017 | 24,8984812782627 | NA | NA | NA | NA | + | + | NA | + | + | + | + | + | 0,162859813082644 | 9 | -517,931548328257 | 1055,28041949116 | 7,31884657586806 | 0.000895592218075651 |
| 2918 | 24,8407434469728 | + | NA | + | NA | NA | + | + | NA | + | + | NA | + | 0,16285893652856 | 9 | -517,931620015988 | 1055,28056286662 | 7,3189899513302 | 0.000895528017402806 |
| 3890 | 24,8942697490276 | + | NA | NA | NA | + | + | NA | NA | + | + | + | + | 0,162847693854557 | 9 | -517,932539475465 | 1055,28240178557 | 7,32082887028423 | 0.000894704994106537 |
| 1839 | 24,2561076207022 | NA | + | + | + | NA | + | NA | NA | + | + | + | NA | 0,162813679807844 | 9 | -517,9353211707 | 1055,28796517605 | 7,32639226075503 | 0.000892219655816776 |
| 954 | 24,7499602029943 | + | NA | NA | + | + | + | NA | + | + | + | NA | NA | 0,162749596453161 | 9 | -517,940561650492 | 1055,29844613563 | 7,33687322033916 | 0.000887556226715865 |
| 1965 | 24,7804013760122 | NA | NA | + | + | NA | + | NA | + | + | + | + | NA | 0,162720646953893 | 9 | -517,942928893006 | 1055,30318062066 | 7,34160770536664 | 0.000885457650781738 |
| 1850 | 24,659092742873 | + | NA | NA | + | + | + | NA | NA | + | + | + | NA | 0,162565116126444 | 9 | -517,955645473987 | 1055,32861378262 | 7,36704086732811 | 0.000874268948656893 |
| 1973 | 24,9302728132396 | NA | NA | + | NA | + | + | NA | + | + | + | + | NA | 0,162553481596504 | 9 | -517,956596646723 | 1055,33051612809 | 7,36894321280147 | 0.000873437763231531 |
| 1899 | 24,2809389842956 | NA | + | NA | + | NA | + | + | NA | + | + | + | NA | 0,162498765214537 | 9 | -517,961069768581 | 1055,33946237181 | 7,37788945651732 | 0.000869539494895908 |
| 950 | 25,012761619111 | + | NA | + | NA | + | + | NA | + | + | + | NA | NA | 0,162491801620468 | 9 | -517,961639028779 | 1055,3406008922 | 7,37902797691231 | 0.000869044641534839 |
| 1977 | 24,6572058850616 | NA | NA | NA | + | + | + | NA | + | + | + | + | NA | 0,16246074271941 | 9 | -517,964177975599 | 1055,34567878584 | 7,38410587055387 | 0.000866840982075967 |
| 3879 | 24,4362592622628 | NA | + | + | NA | NA | + | NA | NA | + | + | + | + | 0,162457207620272 | 9 | -517,96446695052 | 1055,34625673568 | 7,384683820394 | 0.000866590522962404 |
| 2923 | 24,2863326222954 | NA | + | NA | + | NA | + | + | NA | + | + | NA | + | 0,162411088176997 | 9 | -517,96823684977 | 1055,35379653419 | 7,39222361889574 | 0.000863329714321027 |
| 1846 | 24,8995912317635 | + | NA | + | NA | + | + | NA | NA | + | + | + | NA | 0,162367541330817 | 9 | -517,971796269153 | 1055,36091537295 | 7,39934245766062 | 0.000860262224282805 |
| 551 | 23,5799303695423 | NA | + | + | NA | NA | + | NA | NA | NA | + | NA | NA | 0,119876101679459 | 6 | -521,359681167854 | 1055,36551618186 | 7,40394326657133 | 0.000858285547678816 |
| 3043 | 24,6207145264969 | NA | + | NA | NA | NA | + | + | + | + | + | NA | + | 0,162301493957055 | 9 | -517,977194477111 | 1055,37171178887 | 7,41013887357758 | 0.000855630861673289 |
| 2994 | 24,9692447998941 | + | NA | NA | NA | + | + | NA | + | + | + | NA | + | 0,162287802383213 | 9 | -517,978313468569 | 1055,37394977178 | 7,4123768564923 | 0.000854673953534288 |
| 3945 | 24,4807817544536 | NA | NA | NA | + | NA | + | + | NA | + | + | + | + | 0,162220643285173 | 9 | -517,983802014547 | 1055,38492686374 | 7,42335394844804 | 0.000849995885882482 |
| 2019 | 24,5980989069413 | NA | + | NA | NA | NA | + | + | + | + | + | + | NA | 0,162204730712414 | 9 | -517,985102397696 | 1055,38752763004 | 7,42595471474715 | 0.000848891283913839 |
| 675 | 23,8425013770604 | NA | + | NA | NA | NA | + | NA | + | NA | + | NA | NA | 0,11967647172518 | 6 | -521,375208907524 | 1055,3965716182 | 7,43499874591112 | 0.000845061250562392 |
| 681 | 23,8764919645957 | NA | NA | NA | + | NA | + | NA | + | NA | + | NA | NA | 0,119648073480271 | 6 | -521,3774175113 | 1055,40098886875 | 7,43941595346337 | 0.000843196904651897 |
| 674 | 24,0921591918543 | + | NA | NA | NA | NA | + | NA | + | NA | + | NA | NA | 0,119548327665148 | 6 | -521,38517443291 | 1055,41650271197 | 7,45492979668393 | 0.000836681594404637 |
| 2922 | 24,4296320671502 | + | NA | NA | + | NA | + | + | NA | + | + | NA | + | 0,161999689122153 | 9 | -518,001856287653 | 1055,42103540995 | 7,45946249466101 | 0.000834787529036705 |
| 785 | 26,1951385844158 | NA | NA | NA | NA | + | NA | NA | NA | + | + | NA | NA | 0,105246257228945 | 5 | -522,488406977864 | 1055,4348292229 | 7,47325630761338 | 0.000829049886257945 |
| 2856 | 24,4240123287229 | + | + | + | NA | NA | + | NA | NA | + | + | NA | + | 0,161901081707952 | 9 | -518,009912012445 | 1055,43714685954 | 7,47557394424575 | 0.000828089724498681 |
| 1894 | 24,8208444696334 | + | NA | + | NA | NA | + | + | NA | + | + | + | NA | 0,161884941328286 | 9 | -518,011230509247 | 1055,43978385314 | 7,47821093784887 | 0.000826998610318873 |
| 2721 | 24,0548180583445 | NA | NA | NA | NA | NA | + | NA | + | NA | + | NA | + | 0,119387912741473 | 6 | -521,397647560291 | 1055,44144896674 | 7,47987605144476 | 0.000826310373541818 |
| 876 | 24,2283322086802 | + | + | NA | + | NA | + | + | NA | + | + | NA | NA | 0,161811255932246 | 9 | -518,017249497506 | 1055,45182182966 | 7,49024891436693 | 0.000822035865726455 |
| 2863 | 24,2085825040877 | NA | + | + | + | NA | + | NA | NA | + | + | NA | + | 0,161811197583444 | 9 | -518,017254263516 | 1055,45183136168 | 7,49025844638641 | 0.00082203194790485 |
| 2597 | 23,8674575666915 | NA | NA | + | NA | NA | + | NA | NA | NA | + | NA | + | 0,119210633925439 | 6 | -521,411429305043 | 1055,46901245624 | 7,50743954095014 | 0.000815000488871765 |
| 1570 | 24,0382119903135 | + | NA | NA | NA | NA | + | NA | NA | NA | + | + | NA | 0,119195765826708 | 6 | -521,412585032536 | 1055,47132391123 | 7,50975099593529 | 0.000814059114491224 |
| 1633 | 23,9703804586538 | NA | NA | NA | NA | NA | + | + | NA | NA | + | + | NA | 0,119159014693466 | 6 | -521,415441689052 | 1055,47703722426 | 7,51546430896724 | 0.000811736945616018 |
| 996 | 24,5529133552122 | + | + | NA | NA | NA | + | + | + | + | + | NA | NA | 0,16158119618825 | 9 | -518,036038514657 | 1055,48939986396 | 7,52782694866937 | 0.00080673481572969 |
| 1898 | 24,4538789212354 | + | NA | NA | + | NA | + | + | NA | + | + | + | NA | 0,161494924960239 | 9 | -518,043082971598 | 1055,50348877784 | 7,54191586255092 | 0.000801071776992636 |
| 3939 | 24,5027389745182 | NA | + | NA | NA | NA | + | + | NA | + | + | + | + | 0,161459491045287 | 9 | -518,045976109767 | 1055,50927505418 | 7,54770213888946 | 0.000798757515012373 |
| 613 | 23,8224565796424 | NA | NA | + | NA | NA | + | + | NA | NA | + | NA | NA | 0,118905661492407 | 6 | -521,435131530089 | 1055,51641690633 | 7,5548439910408 | 0.000795910297600856 |
| 737 | 23,9898869001813 | NA | NA | NA | NA | NA | + | + | NA | + | + | NA | NA | 0,118793685996067 | 6 | -521,443832122274 | 1055,5338180907 | 7,57224517541044 | 0.000789015444831938 |
| 1963 | 24,3778231363761 | NA | + | NA | + | NA | + | NA | + | + | + | + | NA | 0,161274344686312 | 9 | -518,061091108848 | 1055,53950505234 | 7,57793213705054 | 0.000786775081272758 |
| 1970 | 24,8232606729876 | + | NA | NA | NA | + | + | NA | + | + | + | + | NA | 0,161103604896927 | 9 | -518,075027027368 | 1055,56737688938 | 7,60580397409012 | 0.000775886693905069 |
| 940 | 24,3560027318008 | + | + | NA | + | NA | + | NA | + | + | + | NA | NA | 0,161036440957547 | 9 | -518,080508225055 | 1055,57833928475 | 7,61676636946436 | 0.000771645539474165 |
| 2987 | 24,3752166836074 | NA | + | NA | + | NA | + | NA | + | + | + | NA | + | 0,161019028270868 | 9 | -518,081929189497 | 1055,58118121364 | 7,61960829834834 | 0.000770549837262057 |
| 557 | 23,683118018145 | NA | NA | + | + | NA | + | NA | NA | NA | + | NA | NA | 0,118452077023888 | 6 | -521,470368606307 | 1055,58689105877 | 7,62531814347653 | 0.000768353114373814 |
| 3042 | 24,7878102346693 | + | NA | NA | NA | NA | + | + | + | + | + | NA | + | 0,16094139382552 | 9 | -518,088264199319 | 1055,59385123328 | 7,63227831799304 | 0.000765683825863O9 |
| 1892 | 24,4628113398085 | + | + | NA | NA | NA | + | + | NA | + | + | + | NA | 0,160928987905599 | 9 | -518,089276474341 | 1055,59587578333 | 7,63430286803623 | 0.000764909135417776 |
| 4065 | 24,8316260854833 | NA | NA | NA | NA | NA | + | + | + | + | + | + | + | 0,160901202706733 | 9 | -518,091543584697 | 1055,60041000404 | 7,63883708874801 | 0.000763176966246806 |
| 2916 | 24,4445744557042 | + | + | NA | NA | NA | + | + | NA | + | + | NA | + | 0,160731827399072 | 9 | -518,105362000477 | 1055,6280468356 | 7,66647392030973 | 0.00075270359894058 |
| 777 | 25,3780571693339 | NA | NA | NA | + | NA | NA | NA | NA | + | + | NA | NA | 0,103974622835758 | 5 | -522,585643529523 | 1055,62930232622 | 7,66772941093132 | 0.000752231241061661 |
| 3617 | 23,9109431248794 | NA | NA | NA | NA | NA | + | NA | NA | NA | + | + | + | 0,118156406775727 | 6 | -521,493328233597 | 1055,63281031335 | 7,67123739805652 | 0.000750912988748253 |
| 4005 | 24,982713065042 | NA | NA | + | NA | NA | + | NA | + | + | + | + | + | 0,160659091514459 | 9 | -518,111295272653 | 1055,63991337995 | 7,67834046466101 | 0.000748250826438915 |
| 2982 | 24,9958801026887 | + | NA | + | NA | NA | + | NA | + | + | + | NA | + | 0,160654320042334 | 9 | -518,11168447719 | 1055,64069178903 | 7,67911887373589 | 0.000747959660487351 |
| 2939 | 24,7143002440152 | NA | + | NA | + | + | + | + | NA | + | + | NA | + | 0,174755192897054 | 10 | -516,951726698204 | 1055,64948514244 | 7,68791222715004 | 0.000744678342403514 |
| 1577 | 23,8134101638605 | NA | NA | NA | + | NA | + | NA | NA | NA | + | + | NA | 0,117959124817501 | 6 | -521,50864345384 | 1055,66344075383 | 7,70186783854388 | 0.00073950020863932 |
| 895 | 24,5872797292377 | NA | + | + | + | + | + | + | NA | + | + | NA | NA | 0,174664289951974 | 10 | -516,959267760461 | 1055,66456726695 | 7,70299435166385 | 0.000739083797579833 |
| 4003 | 24,5338992520422 | NA | + | NA | NA | NA | + | NA | + | + | + | + | + | 0,160272966396871 | 9 | -518,142783989738 | 1055,70289081412 | 7,74131789883086 | 0.000725056464682709 |
| 770 | 25,4397433798011 | + | NA | NA | NA | NA | NA | NA | NA | + | + | NA | NA | 0,103398108942391 | 5 | -522,629681697598 | 1055,71737866237 | 7,75580574708079 | 0.000719823188279245 |
| 1956 | 24,5254892056862 | + | + | NA | NA | NA | + | NA | + | + | + | + | NA | 0,160122464018905 | 9 | -518,15505362247 | 1055,72743007959 | 7,76585716429463 | 0.000716214642032722 |
| 2018 | 24,7609292354261 | + | NA | NA | NA | NA | + | + | + | + | + | + | NA | 0,160100751352638 | 9 | -518,156823555577 | 1055,7309699458 | 7,76939703050971 | 0.00071494811119375 |
| 1958 | 24,9430353325543 | + | NA | + | NA | NA | + | NA | + | + | + | + | NA | 0,160009959679397 | 9 | -518,164224048458 | 1055,74577093156 | 7,78419801627024 | 0.000709676672467996 |
| 3938 | 24,6676941061572 | + | NA | NA | NA | NA | + | + | NA | + | + | + | + | 0,159922317767985 | 9 | -518,171367043727 | 1055,7600569221 | 7,79848400680908 | 0.000704625517020287 |
| 2980 | 24,5065170812078 | + | + | NA | NA | NA | + | NA | + | + | + | NA | + | 0,15980446385842 | 9 | -518,180971204612 | 1055,77926524387 | 7,81769232857982 | 0.00069789057369962 |
| 2935 | 24,9630501415507 | NA | + | + | NA | + | + | + | NA | + | + | NA | + | 0,173901764377822 | 10 | -517,022492149667 | 1055,79101604537 | 7,82944313007465 | 0.000693802209052613 |
| 833 | 25,4846354191825 | NA | NA | NA | NA | NA | NA | + | NA | + | + | NA | NA | 0,102814498332556 | 5 | -522,674233144509 | 1055,80648155619 | 7,84490864090208 | 0.000688457896027719 |
| 2819 | 24,7713956849939 | NA | + | NA | NA | NA | NA | NA | NA | + | + | NA | + | 0,116726188311155 | 6 | -521,604280209024 | 1055,8547142642 | 7,8931413489106 | 0.000672053405007715 |
| 2941 | 25,0075544047236 | NA | NA | + | + | + | + | + | NA | + | + | NA | + | 0,173461315670675 | 10 | -517,058985141584 | 1055,8640020292 | 7,90242911390965 | 0.000668939703389429 |
| 832 | 24,5786038343073 | + | + | + | + | + | + | NA | NA | + | + | NA | NA | 0,173203199182928 | 10 | -517,080362125574 | 1055,90675599718 | 7,94518308188913 | 0.000654791551246632 |
| 2879 | 24,6104953062541 | NA | + | + | + | + | + | NA | NA | + | + | NA | + | 0,173054829212574 | 10 | -517,092646979222 | 1055,93132570448 | 7,96975278918558 | 0.000646796740952906 |
| 892 | 24,5936115139099 | + | + | NA | + | + | + | + | NA | + | + | NA | NA | 0,173047928136047 | 10 | -517,093218326374 | 1055,93246839878 | 7,97089548348845 | 0.00064642730102668 |
| 787 | 25,7724979009301 | NA | + | NA | NA | + | NA | NA | NA | + | + | NA | NA | 0,116089489771465 | 6 | -521,653615584073 | 1055,9533850143 | 7,99181209900848 | 0.000639701994330599 |
| 548 | 23,543134778816 | + | + | NA | NA | NA | + | NA | NA | NA | + | NA | NA | 0,116001896606978 | 6 | -521,660400069346 | 1055,96695398485 | 8,00538106955582 | 0.000635376634830513 |
| 2862 | 24,5257600914377 | + | NA | + | + | NA | + | NA | NA | + | + | NA | + | 0,158570306328395 | 9 | -518,281464474789 | 1055,98025178422 | 8,01867886893388 | 0.000631166092568921 |
| 2986 | 24,609208400548 | + | NA | NA | + | NA | + | NA | + | + | + | NA | + | 0,158502637749684 | 9 | -518,286970237053 | 1055,99126330875 | 8,02969039346135 | 0.000627700590982496 |
| 1587 | 24,0235501943615 | NA | + | NA | NA | + | + | NA | NA | NA | + | + | NA | 0,129994037813902 | 7 | -520,568035539589 | 1056,00428813344 | 8,04271521815031 | 0.000623626027923218 |
| 4009 | 24,601193439574 | NA | NA | NA | + | NA | + | NA | + | + | + | + | + | 0,158414751279125 | 9 | -518,294120340158 | 1056,00556351496 | 8,04399059967091 | 0.000623228474138988 |
| 610 | 23,7332599256963 | + | NA | NA | NA | NA | + | + | NA | NA | + | NA | NA | 0,115645592135155 | 6 | -521,687990532331 | 1056,02213491082 | 8,06056199552495 | 0.000618085925485936 |

| | | | | | | | | | | | | | | | | | |
|---|---|---|---|---|---|---|---|---|---|---|---|---|---|---|---|---|---|
| **1838** | 24,5340700342639 | + | NA | + | + | NA | + | NA | NA | + | + | + | NA | 0,158235465294913 | 9 | -518,308704040182 | 1056,03473091501 | 8,07315799971843 | 0.00061420545147451 |
| **773** | 25,9004341371107 | NA | NA | + | NA | NA | NA | NA | NA | + | + | NA | NA | 0,101248961579561 | 5 | -522,793599581332 | 1056,04521442984 | 8,0836415145493 | 0.000610994358731576 |
| **567** | 23,9637045856201 | NA | + | + | NA | + | + | NA | NA | NA | + | NA | NA | 0,129683516339427 | 7 | -520,592467877683 | 1056,05315280963 | 8,09157989433857 | 0.000608574012681368 |
| **1962** | 24,5871570264312 | + | NA | NA | + | NA | + | NA | + | + | + | + | NA | 0,15812150224328 | 9 | -518,317972548438 | 1056,05326793152 | 8,09169501623182 | 0.000608538983593267 |
| **3059** | 24,8216062728699 | NA | + | NA | NA | + | + | + | + | + | + | NA | + | 0,172304167380749 | 10 | -517,154767119936 | 1056,0555659859 | 8,09399307061244 | 0.000607840157316838 |
| **897** | 25,9560284978021 | NA | NA | NA | NA | NA | NA | NA | + | + | + | NA | NA | 0,101129113718348 | 5 | -522,802728975723 | 1056,06347321862 | 8,10190030333183 | 0.000605441734872059 |
| **3885** | 24,5016521337383 | NA | NA | + | + | NA | + | NA | NA | + | + | + | + | 0,158043502558219 | 9 | -518,324315466946 | 1056,06595376854 | 8,1043808532479 | 0.000604691286127341 |
| **2594** | 23,7274721710085 | + | NA | NA | NA | NA | + | NA | NA | NA | + | NA | + | 0,115343896079616 | 6 | -521,711343703106 | 1056,06884125237 | 8,10726833707463 | 0.000603818897876188 |
| **554** | 23,6072808374862 | + | NA | NA | + | NA | + | NA | NA | NA | + | NA | NA | 0,115315123826763 | 6 | -521,71357044051 | 1056,07329472717 | 8,11172181188226 | 0.000602475847616224 |
| **1915** | 24,543589506282 | NA | + | NA | + | + | + | + | NA | + | + | + | NA | 0,172103434938283 | 10 | -517,17136895627 | 1056,08876965857 | 8,12719674328059 | 0.000597832199611577 |
| **888** | 24,846802858227 | + | + | + | NA | + | + | + | NA | + | + | NA | NA | 0,172063555159408 | 10 | -517,174666785926 | 1056,09536531788 | 8,13379240259269 | 0.000595863898199626 |
| **2872** | 24,8558662311058 | + | + | + | NA | + | + | NA | NA | + | + | NA | + | 0,172022949302686 | 10 | -517,178024494944 | 1056,10208073592 | 8,14050782062941 | 0.00059386651580594 |
| **1019** | 24,5135294854927 | NA | + | NA | + | + | + | + | + | + | + | NA | NA | 0,172016905765991 | 10 | -517,178524222519 | 1056,10308019107 | 8,14150727577953 | 0.00059356981847218 |
| **3961** | 24,8817494234576 | NA | NA | NA | + | + | + | + | NA | + | + | + | + | 0,171943384048217 | 10 | -517,184603289679 | 1056,11523832539 | 8,15366541009985 | 0.000589972413194109 |
| **564** | 24,0639897472255 | + | + | NA | NA | + | + | NA | NA | NA | + | NA | NA | 0,12925929161838 | 7 | -520,625832494568 | 1056,1198820434 | 8,15830912810884 | 0.000588604169483306 |
| **1836** | 24,150909231843 | + | + | NA | + | NA | + | NA | NA | + | + | + | NA | 0,157659437744328 | 9 | -518,355538976261 | 1056,12840078717 | 8,1668278718771 | 0.000586102417161959 |
| **2657** | 23,6283773820723 | NA | NA | NA | NA | NA | + | + | NA | NA | + | NA | + | 0,114915170123064 | 6 | -521,744516096804 | 1056,13518603976 | 8,17361312447019 | 0.000584117359871794 |
| **2932** | 24,9006267172719 | + | + | NA | NA | + | + | + | NA | + | + | NA | + | 0,171801646348372 | 10 | -517,196321202409 | 1056,13867415085 | 8,17710123555844 | 0.000583099514599238 |
| **2833** | 25,5602994341546 | NA | NA | NA | NA | + | NA | NA | NA | + | + | NA | + | 0,114891950239343 | 6 | -521,746312261791 | 1056,13877836974 | 8,17720545444445 | 0.000583069130399975 |
| **1793** | 25,7166428764032 | NA | NA | NA | NA | NA | NA | NA | NA | + | + | + | NA | 0,100607989222766 | 5 | -522,84241142117 | 1056,14283810951 | 8,18126519422412 | 0.000581886776346905 |
| **529** | 25,4482186078033 | NA | NA | NA | NA | + | NA | NA | NA | NA | + | NA | NA | 0,0863400849059778 | 4 | -523,920044045006 | 1056,14311839304 | 8,18154547775157 | 0.000581805235421559 |
| **3955** | 24,8823279657412 | NA | + | NA | NA | + | + | + | NA | + | + | + | + | 0,171758652348737 | 10 | -517,199875258804 | 1056,14578226364 | 8,18420934834899 | 0.000581030824338744 |
| **3065** | 24,9007934855014 | NA | NA | NA | + | + | + | + | + | + | + | NA | + | 0,171748376726925 | 10 | -517,200724655602 | 1056,14748105723 | 8,18590814194431 | 0.000580537508157459 |
| **2938** | 24,9483201802958 | + | NA | NA | + | + | + | + | NA | + | + | NA | + | 0,171732988105347 | 10 | -517,201996680178 | 1056,15002510639 | 8,18845219109812 | 0.000579799519648536 |
| **1911** | 24,8015499417685 | NA | + | + | NA | + | + | + | NA | + | + | + | NA | 0,171707076478144 | 10 | -517,204138483793 | 1056,15430871362 | 8,19273579832748 | 0.000578559031856393 |
| **959** | 24,5577970543484 | NA | + | + | + | + | + | NA | + | + | + | NA | NA | 0,171646989743873 | 10 | -517,20910487616 | 1056,16424149835 | 8,20266858306195 | 0.000575692803993978 |
| **2611** | 24,0296214093515 | NA | + | NA | NA | + | + | NA | NA | NA | + | NA | + | 0,128927627344886 | 7 | -520,651906061184 | 1056,17202917663 | 8,21045626134196 | 0.000573455507484345 |
| **1015** | 24,7616150372555 | NA | + | + | NA | + | + | + | + | + | + | NA | NA | 0,171536917506252 | 10 | -517,218201822666 | 1056,18243539136 | 8,22086247607331 | 0.000570479505851413 |
| **2561** | 24,3697409126064 | NA | NA | NA | NA | NA | NA | NA | NA | NA | + | NA | + | 0,0860766736336875 | 4 | -523,939780569561 | 1056,18259144215 | 8,2210185268616 | 0.000570434995699601 |
| **3878** | 24,6917193135393 | + | NA | + | NA | NA | + | NA | NA | + | + | + | + | 0,15729626073559 | 9 | -518,385051272758 | 1056,18742538016 | 8,22585246487051 | 0.000569057936820619 |
| **2999** | 24,8086474508613 | NA | + | + | NA | + | + | NA | + | + | + | NA | + | 0,171482754122777 | 10 | -517,222677725042 | 1056,19138719612 | 8,22981428082517 | 0.000567931801167805 |
| **3895** | 24,7841566614705 | NA | + | + | NA | + | + | NA | NA | + | + | + | + | 0,171456629636484 | 10 | -517,224836471047 | 1056,19570468813 | 8,2341317728351 | 0.000566707103042644 |
| **3876** | 24,267557344936 | + | + | NA | NA | NA | + | NA | NA | + | + | + | + | 0,157235486724108 | 9 | -518,389988616434 | 1056,19730006751 | 8,23572715222326 | 0.000566255226879424 |
| **894** | 24,8937680048758 | + | NA | + | + | + | + | + | NA | + | + | NA | NA | 0,171332541997687 | 10 | -517,235089282309 | 1056,21621031065 | 8,2546373953594 | 0.000560926446701972 |
| **3883** | 24,1245037837879 | NA | + | NA | + | NA | + | NA | NA | + | + | + | + | 0,157049166524754 | 9 | -518,405123244948 | 1056,22756932454 | 8,26599640925178 | 0.000557749690794572 |
| **2595** | 23,4201113616466 | NA | + | NA | NA | NA | + | NA | NA | NA | + | NA | + | 0,114273308684367 | 6 | -521,794149701252 | 1056,23445324866 | 8,27288033336754 | 0.000555833237592702 |
| **611** | 23,4369754282682 | NA | + | NA | NA | NA | + | + | NA | NA | + | NA | NA | 0,114216300258256 | 6 | -521,798556287568 | 1056,24326642129 | 8,28169350599978 | 0.000553389299121492 |
| **1855** | 24,4875171604035 | NA | + | + | + | + | + | NA | NA | + | + | + | NA | 0,171129707116774 | 10 | -517,251845325221 | 1056,24972239647 | 8,28814948118293 | 0.000551605845362539 |
| **617** | 23,4663165926777 | NA | NA | NA | + | NA | + | + | NA | NA | + | NA | NA | 0,114120892355441 | 6 | -521,805930408144 | 1056,25801466244 | 8,2964417471519 | 0.000549323548794654 |
| **4002** | 24,7311668591669 | + | NA | NA | NA | NA | + | NA | + | + | + | + | + | 0,15685625560524 | 9 | -518,420789709337 | 1056,25890225332 | 8,29732933802984 | 0.000549079815597022 |
| **2876** | 24,584585831945 | + | + | NA | + | + | + | NA | NA | + | + | NA | + | 0,170848965904181 | 10 | -517,27503039345 | 1056,29609253293 | 8,33451961764126 | 0.000538963944169726 |
| **2601** | 23,5244973050655 | NA | NA | NA | + | NA | + | NA | NA | NA | + | NA | + | 0,113768031967951 | 6 | -521,833196254253 | 1056,31254635466 | 8,35097343936968 | 0.000534548125023225 |
| **1021** | 24,8442369500072 | NA | NA | + | + | + | + | + | + | + | + | NA | NA | 0,170730454628927 | 10 | -517,28481531536 | 1056,31566237675 | 8,35408946146094 | 0.000533715941583849 |
| **2825** | 24,8819865259146 | NA | NA | NA | + | NA | NA | NA | NA | + | + | NA | + | 0,113729395297216 | 6 | -521,836181086146 | 1056,31851601845 | 8,3569431031558 | 0.000532954967567689 |
| **2860** | 24,0794131300873 | + | + | NA | + | NA | + | NA | NA | + | + | NA | + | 0,156485476643233 | 9 | -518,450890935129 | 1056,3191047049 | 8,3575317896134 | 0.000532798118966523 |
| **1848** | 24,7762214619537 | + | + | + | NA | + | + | NA | NA | + | + | + | NA | 0,170693519366726 | 10 | -517,287864601603 | 1056,32176094924 | 8,36018803394791 | 0.000532090967669656 |
| **1917** | 24,7952097233202 | NA | NA | + | + | + | + | + | NA | + | + | + | NA | 0,170648818165642 | 10 | -517,291554843506 | 1056,32914143304 | 8,36756851775363 | 0.000530131041809067 |
| **555** | 23,3402031327481 | NA | + | NA | + | NA | + | NA | NA | NA | + | NA | NA | 0,113531933955904 | 6 | -521,851433706068 | 1056,34902125829 | 8,38744834299837 | 0.000524887687985528 |
| **952** | 24,7686650237798 | + | + | + | NA | + | + | NA | + | + | + | NA | NA | 0,17051372127785 | 10 | -517,302706360194 | 1056,35144446642 | 8,38987155112932 | 0.00052425211703677 |
| **1575** | 23,942958816188 | NA | + | + | NA | NA | + | NA | NA | NA | + | + | NA | 0,127712847508417 | 7 | -520,747320515629 | 1056,36285808552 | 8,40128517023049 | 0.000521268830668472 |
| **2613** | 24,3747680460229 | NA | NA | + | NA | + | + | NA | NA | NA | + | NA | + | 0,127615230651734 | 7 | -520,75498203134 | 1056,37818111694 | 8,41660820165225 | 0.00051729038126984 |
| **566** | 24,4080463602397 | + | NA | + | NA | + | + | NA | NA | NA | + | NA | NA | 0,127290528637918 | 7 | -520,780460293568 | 1056,4291376414 | 8,467564726109 | 0.000504277201829067 |
| **1012** | 24,6994126275063 | + | + | NA | NA | + | + | + | + | + | + | NA | NA | 0,170033218127102 | 10 | -517,342354589057 | 1056,43074092415 | 8,46916800885492 | 0.000503873114348901 |
| **1007** | 24,5837442081543 | NA | + | + | + | NA | + | + | + | + | + | NA | NA | 0,169970498110271 | 10 | -517,347528174636 | 1056,4410880953 | 8,47951518001332 | 0.00050127301538752 |
| **3003** | 24,5493509949703 | NA | + | NA | + | + | + | NA | + | + | + | NA | + | 0,169929457662595 | 10 | -517,35091326627 | 1056,44785827857 | 8,48628536328079 | 0.000499579029063976 |
| **3899** | 24,5303451115463 | NA | + | NA | + | + | + | NA | NA | + | + | + | + | 0,169876880267073 | 10 | -517,355249702173 | 1056,45653115038 | 8,49495823508732 | 0.000497417327054433 |
| **2945** | 25,4131128959416 | NA | NA | NA | NA | NA | NA | NA | + | + | + | NA | + | 0,112827799845711 | 6 | -521,905795910194 | 1056,45774566654 | 8,49617275125024 | 0.000497115358058558 |
| **1975** | 24,7181263537101 | NA | + | + | NA | + | + | NA | + | + | + | + | NA | 0,169844252900342 | 10 | -517,357940577708 | 1056,46191290145 | 8,5003399861564 | 0.000496080638176865 |
| **1908** | 24,7658042034951 | + | + | NA | NA | + | + | + | NA | + | + | + | NA | 0,169809128931052 | 10 | -517,360837237304 | 1056,46770622064 | 8,50613330534998 | 0.000494645740643673 |
| **2035** | 24,6689678113507 | NA | + | NA | NA | + | + | + | + | + | + | + | NA | 0,169711140772617 | 10 | -517,368917633786 | 1056,4838670136 | 8,52229409831307 | 0.00049066491193843 |
| **571** | 23,756818804124 | NA | + | NA | + | + | + | NA | NA | NA | + | NA | NA | 0,126888247959016 | 7 | -520,812012752113 | 1056,49224255849 | 8,53066964319919 | 0.000488614415444497 |
| **1589** | 24,3509127744743 | NA | NA | + | NA | + | + | NA | NA | NA | + | + | NA | 0,126870878173085 | 7 | -520,813374805594 | 1056,49496666545 | 8,53339375016026 | 0.000487949349509805 |
| **691** | 23,9319111472383 | NA | + | NA | NA | + | + | NA | + | NA | + | NA | NA | 0,126830438352462 | 7 | -520,816545793201 | 1056,50130864067 | 8,53973572537575 | 0.000486404518783307 |
| **2821** | 25,3014240067685 | NA | NA | + | NA | NA | NA | NA | NA | + | + | NA | + | 0,112389866997751 | 6 | -521,939584476258 | 1056,52532279867 | 8,5637498833787 | 0.000480599143843914 |
| **1852** | 24,4914498461632 | + | + | NA | + | + | + | NA | NA | + | + | + | NA | 0,169459360101141 | 10 | -517,389675850509 | 1056,52538344705 | 8,56381053175824 | 0.000480584570285242 |
| **1914** | 24,7691304381491 | + | NA | NA | + | + | + | + | NA | + | + | + | NA | 0,169392654461299 | 10 | -517,395174385276 | 1056,53638051658 | 8,57480760129306 | 0.000477949310988785 |
| **956** | 24,4901679149384 | + | + | NA | + | + | + | NA | + | + | + | NA | NA | 0,169386039745491 | 10 | -517,395719611144 | 1056,53747096832 | 8,57589805302882 | 0.000477688791688283 |
| **2881** | 25,0302362681335 | NA | NA | NA | NA | NA | NA | + | NA | + | + | NA | + | 0,112288791744511 | 6 | -521,947380537778 | 1056,54091492171 | 8,57934200641853 | 0.000476866930537559 |
| **627** | 23,8716600558351 | NA | + | NA | NA | + | + | + | NA | NA | + | NA | NA | 0,126576384058358 | 7 | -520,83646346954 | 1056,54114399334 | 8,57957107805396 | 0.000476812315321487 |
| **1018** | 24,7825370344373 | + | NA | NA | + | + | + | + | + | + | + | NA | NA | 0,169345374177254 | 10 | -517,399071423934 | 1056,5441745939 | 8,58260167860885 | 0.000476090348624103 |
| **1701** | 24,5113121579487 | NA | NA | + | NA | NA | + | NA | + | NA | + | + | NA | 0,126548731869958 | 7 | -520,83863103204 | 1056,54547911834 | 8,58390620305204 | 0.000475779914128984 |
| **693** | 24,3548944560572 | NA | NA | + | NA | + | + | NA | + | NA | + | NA | NA | 0,126478621601409 | 7 | -520,844126433948 | 1056,55646992216 | 8,59489700686868 | 0.000473172483283072 |
| **679** | 24,105107714414 | NA | + | + | NA | NA | + | NA | + | NA | + | NA | NA | 0,126460647666964 | 7 | -520,84553520071 | 1056,55928745568 | 8,59771454039378 | 0.000472506362929942 |
| **2610** | 24,3783682551531 | + | NA | NA | NA | + | + | NA | NA | NA | + | NA | + | 0,126449462582783 | 7 | -520,846411854179 | 1056,56104076262 | 8,59946784733074 | 0.000472092320100558 |
| **3957** | 25,1829678116413 | NA | NA | + | NA | + | + | + | NA | + | + | + | + | 0,169209428713946 | 10 | -517,410275382266 | 1056,56658251056 | 8,60500959527394 | 0.000470786022407523 |
| **2996** | 24,7051392068318 | + | + | NA | NA | + | + | NA | + | + | + | NA | + | 0,16919793863837 | 10 | -517,411222253959 | 1056,56847625395 | 8,60690333865864 | 0.000470340459428527 |

| | | | | | | | | | | | | | | | | | | | |
|---|---|---|---|---|---|---|---|---|---|---|---|---|---|---|---|---|---|---|---|
| 3892 | 24,7295290823454 | + | + | NA | NA | + | + | NA | NA | + | + | + | + | 0,169154319566343 | 10 | -517,414816685973 | 1056,57566511798 | 8,61409220268638 | 0.000468652887372196 |
| 570 | 24,1297187442257 | + | NA | NA | + | + | + | NA | NA | NA | + | NA | NA | 0,126339245699682 | 7 | -520,855049724145 | 1056,57831650255 | 8,61674358726282 | 0.000468032009490995 |
| 2818 | 25,0626647980979 | + | NA | NA | NA | NA | NA | NA | NA | + | + | NA | + | 0,112029482273952 | 6 | -521,967377346493 | 1056,58090853914 | 8,61933562384979 | 0.000467425824343126 |
| 2673 | 24,3061797463395 | NA | NA | NA | NA | + | + | + | NA | NA | + | NA | + | 0,126225954320418 | 7 | -520,863927412104 | 1056,59607187847 | 8,63449896318139 | 0.000463895356481669 |
| 573 | 24,057481522004 | NA | NA | + | + | + | + | NA | NA | NA | + | NA | NA | 0,126225624060826 | 7 | -520,863953290076 | 1056,59612363441 | 8,63455071912404 | 0.000463883351966263 |
| 3061 | 25,187658337477 | NA | NA | + | NA | + | + | + | + | + | + | NA | + | 0,16901820743232 | 10 | -517,426031801396 | 1056,59809534882 | 8,63652243353295 | 0.0004634262545751 |
| 3633 | 24,3396476543413 | NA | NA | NA | NA | + | + | NA | NA | NA | + | + | + | 0,126203483085473 | 7 | -520,86568815615 | 1056,59959336656 | 8,63802045127295 | 0.000463079274161757 |
| 2041 | 24,7081797643555 | NA | NA | NA | + | + | + | + | + | + | + | + | NA | 0,168947274686876 | 10 | -517,431875658621 | 1056,60978306327 | 8,64821014798258 | 0.000460725955475081 |
| 685 | 24,1624807299606 | NA | NA | + | + | NA | + | NA | + | NA | + | NA | NA | 0,12613778921052 | 7 | -520,870835370673 | 1056,60988779561 | 8,64831488031837 | 0.000460701829654037 |
| 2617 | 24,1310895825347 | NA | NA | NA | + | + | + | NA | NA | NA | + | NA | + | 0,126049378577109 | 7 | -520,877761867144 | 1056,62374078855 | 8,66216787326198 | 0.000457521805981576 |
| 2934 | 25,2198676222425 | + | NA | + | NA | + | + | + | NA | + | + | NA | + | 0,16882445277995 | 10 | -517,441993271646 | 1056,63001828932 | 8,66844537403267 | 0.000456088010576803 |
| 1586 | 24,3807705246661 | + | NA | NA | NA | + | + | NA | NA | NA | + | + | NA | 0,125999974331454 | 7 | -520,881632118415 | 1056,63148129109 | 8,66990837580306 | 0.000455754503788466 |
| 3053 | 24,8648208513555 | NA | NA | + | + | NA | + | + | + | + | + | NA | + | 0,168806792734483 | 10 | -517,443447917796 | 1056,63292758162 | 8,67135466633385 | 0.000455425046214068 |
| 4019 | 24,6613867741963 | NA | + | NA | NA | + | + | NA | + | + | + | + | + | 0,168800488233919 | 10 | -517,443967207815 | 1056,63396616166 | 8,67239324637148 | 0.000455188609928066 |
| 944 | 24,6027136482805 | + | + | + | + | NA | + | NA | + | + | + | NA | NA | 0,168780435104602 | 10 | -517,445618920711 | 1056,63726958745 | 8,67569667216276 | 0.000454437389602038 |
| 577 | 24,5839812338395 | NA | NA | NA | NA | NA | NA | + | NA | NA | + | NA | NA | 0,0830009290016545 | 4 | -524,169815935603 | 1056,64266217424 | 8,68108925894603 | 0.00045321374346736 |
| 3841 | 25,2043914489883 | NA | NA | NA | NA | NA | NA | NA | NA | + | + | + | + | 0,111605153656593 | 6 | -522,000087121124 | 1056,6463280884 | 8,68475517311163 | 0.00045238378299958 |
| 629 | 24,2258089735477 | NA | NA | + | NA | + | + | + | NA | NA | + | NA | NA | 0,125859510781422 | 7 | -520,892634618543 | 1056,65348629135 | 8,69191337605935 | 0.000450767549592105 |
| 4081 | 25,0380639803942 | NA | NA | NA | NA | + | + | + | + | + | + | + | + | 0,168642620869099 | 10 | -517,456969166551 | 1056,65997007913 | 8,69839716384377 | 0.000449308575224125 |
| 2737 | 24,3073966558822 | NA | NA | NA | NA | + | + | NA | + | NA | + | NA | + | 0,12568371067581 | 7 | -520,906402539062 | 1056,68102213239 | 8,71944921709746 | 0.000444603945166527 |
| 3882 | 24,3476834480776 | + | NA | NA | + | NA | + | NA | NA | + | + | + | + | 0,154143877066648 | 9 | -518,640685824834 | 1056,69869448431 | 8,73712156902343 | 0.000440692652357745 |
| 1967 | 24,5963199714233 | NA | + | + | + | NA | + | NA | + | + | + | + | NA | 0,168384698826261 | 10 | -517,47820632188 | 1056,70244438979 | 8,74087147450064 | 0.000439867148594429 |
| 515 | 24,5496713070379 | NA | + | NA | NA | NA | NA | NA | NA | NA | + | NA | NA | 0,0824973353280244 | 4 | -524,207406289777 | 1056,71784288258 | 8,75626996729352 | 0.000436493506950159 |
| 1649 | 24,2794701289435 | NA | NA | NA | NA | + | + | + | NA | NA | + | + | NA | 0,125421137537034 | 7 | -520,926961004664 | 1056,72213906359 | 8,76056614830009 | 0.000435556885726866 |
| 2991 | 24,6006430423335 | NA | + | + | + | NA | + | NA | + | + | + | NA | + | 0,168178277090968 | 10 | -517,495198228842 | 1056,73642820372 | 8,77485528842544 | 0.000432456109088366 |
| 3058 | 25,0829302974328 | + | NA | NA | NA | + | + | + | + | + | + | NA | + | 0,168175728929019 | 10 | -517,495407958184 | 1056,7368476624 | 8,77527474710951 | 0.000432365419863578 |
| 3954 | 25,0767094765034 | + | NA | NA | NA | + | + | + | NA | + | + | + | + | 0,168173938049934 | 10 | -517,495555358122 | 1056,73714246228 | 8,77556954698525 | 0.000432301693924266 |
| 1979 | 24,4432465177985 | NA | + | NA | + | + | + | NA | + | + | + | + | NA | 0,16813000236176 | 10 | -517,499171425707 | 1056,74437459745 | 8,78280168215565 | 0.000430741284754667 |
| 626 | 24,2843432141235 | + | NA | NA | NA | + | + | + | NA | NA | + | NA | NA | 0,125272835343018 | 7 | -520,938569769349 | 1056,74535659296 | 8,7837836776705 | 0.000430529843662622 |
| 1593 | 24,1039405126963 | NA | NA | NA | + | + | + | NA | NA | NA | + | + | NA | 0,125248085214575 | 7 | -520,940506962604 | 1056,74923097947 | 8,78765806418096 | 0.0004296966314604 |
| 1713 | 24,3125428323922 | NA | NA | NA | NA | + | + | NA | + | NA | + | + | NA | 0,125132247073702 | 7 | -520,949572889012 | 1056,76736283229 | 8,80578991699781 | 0.000425818638781065 |
| 1006 | 24,8248230765252 | + | NA | + | + | NA | + | + | + | + | + | NA | NA | 0,167957556004762 | 10 | -517,513362545946 | 1056,77275683792 | 8,81118392263306 | 0.000424671751986121 |
| 690 | 24,306521189643 | + | NA | NA | NA | + | + | NA | + | NA | + | NA | NA | 0,125086555527487 | 7 | -520,953148550457 | 1056,77451415518 | 8,81294123988664 | 0.00042429877437168 |
| 678 | 24,4235446028719 | + | NA | + | NA | NA | + | NA | + | NA | + | NA | NA | 0,125085680734025 | 7 | -520,953217006929 | 1056,77465106812 | 8,81307815283049 | 0.000424269729368717 |
| 793 | 25,7839231447308 | NA | NA | NA | + | + | NA | NA | NA | + | + | NA | NA | 0,110671539835297 | 6 | -522,072000687462 | 1056,79015522108 | 8,82858230578768 | 0.000420993473280095 |
| 2725 | 24,3839388704818 | NA | NA | + | NA | NA | + | NA | + | NA | + | NA | + | 0,124937660598389 | 7 | -520,964799254638 | 1056,79781556354 | 8,8362426482488 | 0.000419384080283372 |
| 1699 | 24,0492288845849 | NA | + | NA | NA | NA | + | NA | + | NA | + | + | NA | 0,124907084925067 | 7 | -520,967191489316 | 1056,8026000329 | 8,84102711760488 | 0.000418382014209589 |
| 1903 | 24,4253702042006 | NA | + | + | + | NA | + | + | NA | + | + | + | NA | 0,167725831219449 | 10 | -517,532427234787 | 1056,81088621561 | 8,84931330031554 | 0.000416652205156136 |
| 1972 | 24,6236467870719 | + | + | NA | NA | + | + | NA | + | + | + | + | NA | 0,167703318133405 | 10 | -517,534279170724 | 1056,81459008748 | 8,85301717218863 | 0.000415881306012608 |
| 633 | 23,9692411571325 | NA | NA | NA | + | + | + | + | NA | NA | + | NA | NA | 0,124749336559339 | 7 | -520,979532363245 | 1056,82728178075 | 8,86570886546201 | 0.000413250543063039 |
| 2029 | 24,804110519773 | NA | NA | + | + | NA | + | + | + | + | + | + | NA | 0,167612301985053 | 10 | -517,541765687076 | 1056,82956312018 | 8,86799020489411 | 0.000412779429427036 |
| 2927 | 24,4304598343509 | NA | + | + | + | NA | + | + | NA | + | + | NA | + | 0,167600791151656 | 10 | -517,542712450255 | 1056,83145664654 | 8,869883731252 | 0.00041238881000345 |
| 697 | 24,053385295148 | NA | NA | NA | + | + | + | NA | + | NA | + | NA | NA | 0,124696215498564 | 7 | -520,983687597315 | 1056,83559224889 | 8,87401933360229 | 0.00041153695297652 |
| 880 | 24,3942914387666 | + | + | + | + | NA | + | + | NA | + | + | NA | NA | 0,167496236582349 | 10 | -517,551311438352 | 1056,84865462274 | 8,88708170744553 | 0.00040885788647679 |
| 641 | 25,2482988505747 | NA | NA | NA | NA | NA | NA | NA | + | NA | + | NA | NA | 0,081488258095056 | 4 | -524,282666045551 | 1056,86836239413 | 8,90678947884248 | 0.000404848832416327 |
| 517 | 25,1286619953287 | NA | NA | + | NA | NA | NA | NA | NA | NA | + | NA | NA | 0,0814520765063949 | 4 | -524,285363033 | 1056,87375636903 | 8,91218345374023 | 0.000403758431256672 |
| 521 | 24,6023384087696 | NA | NA | NA | + | NA | NA | NA | NA | NA | + | NA | NA | 0,0813667503219923 | 4 | -524,291722854233 | 1056,8864760115 | 8,9249030962053 | 0.000401198748004388 |
| 835 | 24,9291597734184 | NA | + | NA | NA | NA | NA | + | NA | + | + | NA | NA | 0,109942023663431 | 6 | -522,128140706024 | 1056,9024352582 | 8,9408623429108 | 0.000398010072241239 |
| 3047 | 24,8223536607156 | NA | + | + | NA | NA | + | + | + | + | + | NA | + | 0,16713735014398 | 10 | -517,580819492671 | 1056,90767073137 | 8,94609781608392 | 0.000396969549215098 |
| 1574 | 24,3272074260282 | + | NA | + | NA | NA | + | NA | NA | NA | + | + | NA | 0,124134940085696 | 7 | -521,027576271764 | 1056,92336959779 | 8,96179668250147 | 0.000393865760679302 |
| 2023 | 24,7968946099422 | NA | + | + | NA | NA | + | + | + | + | + | + | NA | 0,167020710285476 | 10 | -517,590407019222 | 1056,92684578447 | 8,96527286918422 | 0.000393181779806761 |
| 753 | 24,1722499512714 | NA | NA | NA | NA | + | + | + | + | NA | + | NA | NA | 0,12407820833984 | 7 | -521,032010820336 | 1056,93223869494 | 8,97066577964461 | 0.000392123010843253 |
| 741 | 24,2978911842113 | NA | NA | + | NA | NA | + | + | + | NA | + | NA | NA | 0,124033593386927 | 7 | -521,035498034292 | 1056,93921312285 | 8,97764020755744 | 0.000390757975475711 |
| 2835 | 25,2285712001238 | NA | + | NA | NA | + | NA | NA | NA | + | + | NA | + | 0,124002749566341 | 7 | -521,037908759043 | 1056,94403457235 | 8,98246165705882 | 0.00038981710010414 |
| 1537 | 25,1548640464324 | NA | NA | NA | NA | NA | NA | NA | NA | NA | + | + | NA | 0,0809277859657419 | 4 | -524,324431908432 | 1056,95189411989 | 8,99032120460402 | 0.000388288213145542 |
| 1910 | 25,0344777838268 | + | NA | + | NA | + | + | + | NA | + | + | + | NA | 0,166696460997919 | 10 | -517,61705251245 | 1056,98013677093 | 9,0185638556411 | 0.000382843601932687 |
| 2037 | 24,993725635481 | NA | NA | + | NA | + | + | + | + | + | + | + | NA | 0,166686122218578 | 10 | -517,617901940978 | 1056,98183562799 | 9,02026271269665 | 0.000382518541732779 |
| 1014 | 25,047606578977 | + | NA | + | NA | + | + | + | + | + | + | NA | NA | 0,166622838417116 | 10 | -517,623101074384 | 1056,9922338948 | 9,03066097951 | 0.000380534937782793 |
| 2926 | 24,6591578013572 | + | NA | + | + | NA | + | + | NA | + | + | NA | + | 0,166479719584306 | 10 | -517,634857666122 | 1057,01574707828 | 9,05417416298565 | 0.000376087339404336 |
| 1637 | 24,2118186277796 | NA | NA | + | NA | NA | + | + | NA | NA | + | + | NA | 0,123476982794517 | 7 | -521,078989164295 | 1057,02619538285 | 9,06462246756382 | 0.000374127724988135 |
| 1000 | 24,7672341610203 | + | + | + | NA | NA | + | + | + | + | + | NA | NA | 0,166415763946645 | 10 | -517,640110692288 | 1057,02625313061 | 9,06468021531714 | 0.000374116922626303 |
| 3949 | 24,6722474584384 | NA | NA | + | + | NA | + | + | NA | + | + | + | + | 0,166363221189482 | 10 | -517,644426014784 | 1057,0348837756 | 9,07331086030831 | 0.000372505965853052 |
| 779 | 25,0201158338538 | NA | + | NA | + | NA | NA | NA | NA | + | + | NA | NA | 0,109053337313543 | 6 | -522,196467582503 | 1057,03908901116 | 9,07751609586899 | 0.000371723551033878 |
| 3005 | 24,9494333662867 | NA | NA | + | + | + | + | NA | + | + | + | NA | + | 0,166268527844716 | 10 | -517,652202467296 | 1057,05043668062 | 9,08886376533269 | 0.00036962042507598 |
| 552 | 23,8356623103983 | + | + | + | NA | NA | + | NA | NA | NA | + | NA | NA | 0,123310525825189 | 7 | -521,091990025846 | 1057,05219710596 | 9,09062419066527 | 0.000369295223640317 |
| 4007 | 24,7878751107364 | NA | + | + | NA | NA | + | NA | + | + | + | + | + | 0,166191735513705 | 10 | -517,65850819593 | 1057,06304813789 | 9,10147522260172 | 0.000367297032019339 |
| 1960 | 24,7943172579993 | + | + | + | NA | NA | + | NA | + | + | + | + | NA | 0,166147395952444 | 10 | -517,662148832168 | 1057,07032941037 | 9,10875649507784 | 0.000365962268303287 |
| 514 | 24,7562492757654 | + | NA | NA | NA | NA | NA | NA | NA | NA | + | NA | NA | 0,0800926868153584 | 4 | -524,386615520809 | 1057,07626134465 | 9,11468842935733 | 0.000364878444328789 |
| 775 | 25,4876559181001 | NA | + | + | NA | NA | NA | NA | NA | + | + | NA | NA | 0,108800811981161 | 6 | -522,215870628776 | 1057,07789510371 | 9,11632218841532 | 0.000364580504304166 |
| 1572 | 23,8676638669657 | + | + | NA | NA | NA | + | NA | NA | NA | + | + | NA | 0,123064459584255 | 7 | -521,111204126511 | 1057,09062530729 | 9,12905239199586 | 0.000362267282041027 |
| 1581 | 24,0217163366811 | NA | NA | + | + | NA | + | NA | NA | NA | + | + | NA | 0,123019950120362 | 7 | -521,114679075764 | 1057,09757520579 | 9,13600229050121 | 0.000361010606326754 |
| 3943 | 24,6771896584525 | NA | + | + | NA | NA | + | + | NA | + | + | + | + | 0,165966552173141 | 10 | -517,676995564002 | 1057,10002287403 | 9,13844995874433 | 0.00036056905947365 |
| 2878 | 24,9190268189825 | + | NA | + | + | + | + | NA | NA | + | + | NA | + | 0,165929647116658 | 10 | -517,680024963136 | 1057,1060816723 | 9,14450875701414 | 0.000359478404725185 |
| 3621 | 24,1866293454177 | NA | NA | + | NA | NA | + | NA | NA | NA | + | + | + | 0,12295986638642 | 7 | -521,119369662084 | 1057,10695637843 | 9,14538346314066 | 0.000359321220118792 |
| 1705 | 24,119506855946 | NA | NA | NA | + | NA | + | NA | + | NA | + | + | NA | 0,122881490430795 | 7 | -521,125487793529 | 1057,11919264132 | 9,15761972603082 | 0.00035712955694124 |
| 786 | 25,8594091268795 | + | NA | NA | NA | + | NA | NA | NA | + | + | NA | NA | 0,108531683464971 | 6 | -522,236543349769 | 1057,11924054569 | 9,15766763040142 | 0.000357121003010358 |

| | | | | | | | | | | | | | | | | | | | |
|---|---|---|---|---|---|---|---|---|---|---|---|---|---|---|---|---|---|---|---|
| 3901 | 24,8105998178306 | NA | NA | + | + | + | + | NA | NA | + | + | + | + | 0,165810645812879 | 10 | -517,689792426152 | 1057,12561659834 | 9,16404368304552 | 0.000355984304727365 |
| 2984 | 24,7735374304282 | + | + | + | NA | NA | + | NA | + | + | + | NA | + | 0,165778444212014 | 10 | -517,692435249762 | 1057,13090224556 | 9,16932933026465 | 0.000355044743096893 |
| 2034 | 24,9057514830038 | + | NA | NA | NA | + | + | + | + | + | + | + | NA | 0,165769902587273 | 10 | -517,693136253934 | 1057,1323042539 | 9,17073133860936 | 0.000354795942465937 |
| 849 | 25,8970617117125 | NA | NA | NA | NA | + | NA | + | NA | + | + | NA | NA | 0,108436024981675 | 6 | -522,243889716372 | 1057,1339332789 | 9,17236036360759 | 0.000354507074395459 |
| 1902 | 24,6606981158909 | + | NA | + | + | NA | + | + | NA | + | + | + | NA | 0,165694454914748 | 10 | -517,699327871334 | 1057,1446874887 | 9,18311457340951 | 0.000352605968471887 |
| 1896 | 24,661974263136 | + | + | + | NA | NA | + | + | NA | + | + | + | NA | 0,165591712966251 | 10 | -517,70775849513 | 1057,16154873629 | 9,19997582100041 | 0.000349645775871814 |
| 958 | 24,9042706920097 | + | NA | + | + | + | + | NA | + | + | + | NA | NA | 0,165465651001381 | 10 | -517,718101255715 | 1057,18223425746 | 9,22066134217084 | 0.00034604811029017 |
| 772 | 25,1266758156102 | + | + | NA | NA | NA | NA | NA | NA | + | + | NA | NA | 0,108031211249578 | 6 | -522,274969824843 | 1057,19609349584 | 9,23452058054977 | 0.000343658418055876 |
| 1698 | 24,3197169370905 | + | NA | NA | NA | NA | + | NA | + | NA | + | + | NA | 0,122376601742479 | 7 | -521,164886983924 | 1057,19799102211 | 9,23641810682011 | 0.000343332522241147 |
| 2920 | 24,6314433590754 | + | + | + | NA | NA | + | + | NA | + | + | NA | + | 0,165207886273485 | 10 | -517,73924471652 | 1057,22452117907 | 9,26294826378194 | 0.00033880826305654 |
| 3745 | 24,2732063732244 | NA | NA | NA | NA | NA | + | NA | + | NA | + | + | + | 0,122090336576334 | 7 | -521,187215737769 | 1057,2426485298 | 9,2810756145102 | 0.00033575128958464 |
| 531 | 25,0484969477863 | NA | + | NA | NA | + | NA | NA | NA | NA | + | NA | NA | 0,093289495730688 | 5 | -523,397283272046 | 1057,25258181127 | 9,29100889597612 | 0.000334087867782258 |
| 1761 | 24,2480136008398 | NA | NA | NA | NA | NA | + | + | + | NA | + | + | NA | 0,122012980035738 | 7 | -521,193248318139 | 1057,25471369054 | 9,29314077525078 | 0.000333731940014097 |
| 899 | 25,5509576983361 | NA | + | NA | NA | NA | NA | NA | + | + | + | NA | NA | 0,107641412519484 | 6 | -522,304883815696 | 1057,25592147755 | 9,29434856225521 | 0.000333530462305703 |
| 3051 | 24,4838417494889 | NA | + | NA | + | NA | + | + | + | + | + | NA | + | 0,164916450128934 | 10 | -517,763142257614 | 1057,27231626126 | 9,3107433459686 | 0.000330807558014389 |
| 1981 | 24,7987899976203 | NA | NA | + | + | + | + | NA | + | + | + | + | NA | 0,16486436936693 | 10 | -517,767411962368 | 1057,28085567077 | 9,31928275547716 | 0.000329398118497704 |
| 558 | 23,8959814391707 | + | NA | + | + | NA | + | NA | NA | NA | + | NA | NA | 0,121838138024182 | 7 | -521,206881258358 | 1057,28197957098 | 9,32040665568957 | 0.000329213065190264 |
| 1840 | 24,3733537278716 | + | + | + | + | NA | + | NA | NA | + | + | + | NA | 0,164849114028812 | 10 | -517,768662580881 | 1057,28335690779 | 9,32178399250392 | 0.0003289864246021 |
| 851 | 25,3134055986671 | NA | + | NA | NA | + | NA | + | NA | + | + | NA | NA | 0,121808063563604 | 7 | -521,209225978064 | 1057,28666901039 | 9,32509609510112 | 0.000328442057081098 |
| 2027 | 24,4589896673339 | NA | + | NA | + | NA | + | + | + | + | + | + | NA | 0,164744114211296 | 10 | -517,777269749481 | 1057,30057124499 | 9,33899832970383 | 0.000326166934299076 |
| 1854 | 24,7673778700249 | + | NA | + | + | + | + | NA | NA | + | + | + | NA | 0,164589636576062 | 10 | -517,789930806282 | 1057,3258933586 | 9,36432044330468 | 0.000322063348899134 |
| 1635 | 23,7710921181956 | NA | + | NA | NA | NA | + | + | NA | NA | + | + | NA | 0,121466471835692 | 7 | -521,235852141331 | 1057,33992133693 | 9,37834842163488 | 0.000319812303691363 |
| 1795 | 25,4027988233415 | NA | + | NA | NA | NA | NA | NA | NA | + | + | + | NA | 0,107032983774525 | 6 | -522,351549849832 | 1057,34925354582 | 9,38768063052748 | 0.000318323502231929 |
| 3898 | 24,7643813544697 | + | NA | NA | + | + | + | NA | NA | + | + | + | + | 0,16436216314273 | 10 | -517,808570372554 | 1057,36317249114 | 9,4015995758491 | 0.000316115829550264 |
| 1004 | 24,4292698387458 | + | + | NA | + | NA | + | + | + | + | + | NA | NA | 0,164349666487868 | 10 | -517,809594223092 | 1057,36522019222 | 9,40364727692577 | 0.000315792339818831 |
| 4021 | 25,0330658844289 | NA | NA | + | NA | + | + | NA | + | + | + | + | + | 0,164334435353866 | 10 | -517,810842088728 | 1057,36771592349 | 9,4061430081972 | 0.000315398519178673 |
| 4025 | 24,7663760819263 | NA | NA | NA | + | + | + | NA | + | + | + | + | + | 0,164330723432879 | 10 | -517,811146197809 | 1057,36832414165 | 9,40675122636003 | 0.000315302618207644 |
| 3894 | 25,0027432304904 | + | NA | + | NA | + | + | NA | NA | + | + | + | + | 0,164199457466481 | 10 | -517,821899646154 | 1057,38983103834 | 9,42825812304886 | 0.000311930192888556 |
| 682 | 23,9777597068097 | + | NA | NA | + | NA | + | NA | + | NA | + | NA | NA | 0,121142529153217 | 7 | -521,261093050829 | 1057,39040315592 | 9,42883024063076 | 0.000311840975276067 |
| 3046 | 25,0309230807441 | + | NA | + | NA | NA | + | + | + | + | + | NA | + | 0,164157697775373 | 10 | -517,825320290791 | 1057,39667232761 | 9,43509941232264 | 0.000310865013386065 |
| 2598 | 24,0740936942494 | + | NA | + | NA | NA | + | NA | NA | NA | + | NA | + | 0,121084711559432 | 7 | -521,265597094045 | 1057,39941124235 | 9,4378383270639 | 0.000310439588369413 |
| 4069 | 25,0530272847979 | NA | NA | + | NA | NA | + | + | + | + | + | + | + | 0,164136555112271 | 10 | -517,827052076045 | 1057,40013589812 | 9,43856298283049 | 0.000310327127825494 |
| 3619 | 23,7171396268792 | NA | + | NA | NA | NA | + | NA | NA | NA | + | + | + | 0,121069303750552 | 7 | -521,266797326416 | 1057,4018117071 | 9,44023879180463 | 0.000310067212240054 |
| 4073 | 24,6534315572344 | NA | NA | NA | + | NA | + | + | + | + | + | + | + | 0,164119447478672 | 10 | -517,828453321949 | 1057,40293838993 | 9,44136547463881 | 0.000309892587728545 |
| 3002 | 24,8487724258147 | + | NA | NA | + | + | + | NA | + | + | + | NA | + | 0,164072741425763 | 10 | -517,832278758596 | 1057,41058926322 | 9,44901634793405 | 0.000308709377856747 |
| 3050 | 24,6321435323094 | + | NA | NA | + | NA | + | + | + | + | + | NA | + | 0,164061455570472 | 10 | -517,833203089062 | 1057,41243792416 | 9,450865008866 | 0.000308424160211647 |
| 614 | 24,0533171271653 | + | NA | + | NA | NA | + | + | NA | NA | + | NA | NA | 0,120990333731244 | 7 | -521,272948576619 | 1057,4141142075 | 9,45254129221053 | 0.00030816576537111 |
| 1579 | 23,6423613512308 | NA | + | NA | + | NA | + | NA | NA | NA | + | + | NA | 0,120940913679082 | 7 | -521,276797795841 | 1057,42181264595 | 9,46023973065485 | 0.000306981847824312 |
| 2599 | 23,677138174757 | NA | + | + | NA | NA | + | NA | NA | NA | + | NA | + | 0,120919082441013 | 7 | -521,278498114105 | 1057,42521328247 | 9,46364036718364 | 0.000306460324485095 |
| 1809 | 26,0644437384435 | NA | NA | NA | NA | + | NA | NA | NA | + | + | + | NA | 0,106499128166212 | 6 | -522,392470010551 | 1057,43109386725 | 9,46952095196434 | 0.000305560564951348 |
| 676 | 23,938045733372 | + | + | NA | NA | NA | + | NA | + | NA | + | NA | NA | 0,120860508301037 | 7 | -521,283059931782 | 1057,43433691783 | 9,47276400253736 | 0.000305065492263637 |
| 2998 | 25,1071492197974 | + | NA | + | NA | + | + | NA | + | + | + | NA | + | 0,163868123749386 | 10 | -517,849035352665 | 1057,44410245136 | 9,4825295360713 | 0.000303579559300535 |
| 2990 | 24,8845144612503 | + | NA | + | + | NA | + | NA | + | + | + | NA | + | 0,163807768204692 | 10 | -517,85397721895 | 1057,45398618393 | 9,49241326864058 | 0.000302083010629163 |
| 905 | 25,6430227123549 | NA | NA | NA | + | NA | NA | NA | + | + | + | NA | NA | 0,106321831925861 | 6 | -522,406054404638 | 1057,45826265543 | 9,49668974014003 | 0.000301437776014149 |
| 795 | 25,4119491904713 | NA | + | NA | + | + | NA | NA | NA | + | + | NA | NA | 0,12063833245774 | 7 | -521,300360466839 | 1057,46893798794 | 9,50736507265174 | 0.000299833088222701 |
| 683 | 23,7474009388565 | NA | + | NA | + | NA | + | NA | + | NA | + | NA | NA | 0,12045420447645 | 7 | -521,314694955507 | 1057,49760696528 | 9,53603404998694 | 0.000295565792040999 |
| 2577 | 24,9156698447226 | NA | NA | NA | NA | + | NA | NA | NA | NA | + | NA | + | 0,0916499430727498 | 5 | -523,520976643267 | 1057,49996855371 | 9,53839563841916 | 0.000295216995632194 |
| 4013 | 24,8415652994658 | NA | NA | + | + | NA | + | NA | + | + | + | + | + | 0,163466353504043 | 10 | -517,881925285486 | 1057,509882317 | 9,54830940171382 | 0.000293757260779903 |
| 2729 | 23,9409106463975 | NA | NA | NA | + | NA | + | NA | + | NA | + | NA | + | 0,120317147132886 | 7 | -521,325363015896 | 1057,51894308606 | 9,55737017076444 | 0.000292429437477601 |
| 2723 | 23,9079896808206 | NA | + | NA | NA | NA | + | NA | + | NA | + | NA | + | 0,120296932331842 | 7 | -521,326936324151 | 1057,52208970257 | 9,5605167872759 | 0.000291969717564795 |
| 2026 | 24,6175127841707 | + | NA | NA | + | NA | + | + | + | + | + | + | NA | 0,163352373194874 | 10 | -517,891253128298 | 1057,52853800263 | 9,56696508733671 | 0.00029102987929497 |
| 1634 | 24,1060736565208 | + | NA | NA | NA | NA | + | + | NA | NA | + | + | NA | 0,120244947490796 | 7 | -521,330982113417 | 1057,5301812811 | 9,5686083658079 | 0.000290790855936342 |
| 2864 | 24,28664963212 | + | + | + | + | NA | + | NA | NA | + | + | NA | + | 0,163326052719897 | 10 | -517,893406944867 | 1057,53284563577 | 9,57127272047501 | 0.000290403728867465 |
| 3887 | 24,308672231954 | NA | + | + | + | NA | + | NA | NA | + | + | + | + | 0,163309163476678 | 10 | -517,894788963736 | 1057,5356096735 | 9,57403675821297 | 0.000290002662638931 |
| 3880 | 24,5390640911698 | + | + | + | NA | NA | + | NA | NA | + | + | + | + | 0,163285167808854 | 10 | -517,896752441641 | 1057,53953662931 | 9,57796371402401 | 0.000289433807468334 |
| 3947 | 24,353227666459 | NA | + | NA | + | NA | + | + | NA | + | + | + | + | 0,163276724103901 | 10 | -517,897443345809 | 1057,54091843765 | 9,57934552235884 | 0.000289233905509064 |
| 615 | 23,6347738964474 | NA | + | + | NA | NA | + | + | NA | NA | + | NA | NA | 0,120161007832823 | 7 | -521,337514324348 | 1057,54324570296 | 9,58167278766882 | 0.000288897539233028 |
| 1966 | 24,8404074944693 | + | NA | + | + | NA | + | NA | + | + | + | + | NA | 0,163253697985902 | 10 | -517,899327417074 | 1057,54468658018 | 9,5831136648892 | 0.000288689481246879 |
| 2661 | 23,9357320255089 | NA | NA | + | NA | NA | + | + | NA | NA | + | NA | + | 0,120146109633131 | 7 | -521,338673641907 | 1057,54556433808 | 9,58399142278654 | 0.000288562809309713 |
| 2943 | 24,7849670196862 | NA | + | + | + | + | + | + | NA | + | + | NA | + | 0,177565821011731 | 11 | -516,71815399499 | 1057,54830798998 | 9,58673507468893 | 0.000288167222759338 |
| 559 | 23,5098775383373 | NA | + | + | + | NA | + | NA | NA | NA | + | NA | NA | 0,120082353556866 | 7 | -521,3436346599 | 1057,55548637406 | 9,59391345877361 | 0.000287134789166108 |
| 2022 | 24,9790523535591 | + | NA | + | NA | NA | + | + | + | + | + | + | NA | 0,163108615923821 | 10 | -517,911197307741 | 1057,56842636151 | 9,60685344622402 | 0.000285283025790317 |
| 781 | 25,5173585799395 | NA | NA | + | + | NA | NA | NA | NA | + | + | NA | NA | 0,105581330489498 | 6 | -522,462762336706 | 1057,57167851957 | 9,61010560427553 | 0.000284819510004457 |
| 2722 | 24,1314975380155 | + | NA | NA | NA | NA | + | NA | + | NA | + | NA | + | 0,119972040597461 | 7 | -521,352217535575 | 1057,57265212541 | 9,61107921012308 | 0.000284680892776612 |
| 1900 | 24,3254080040946 | + | + | NA | + | NA | + | + | NA | + | + | + | NA | 0,16303388393761 | 10 | -517,917310703578 | 1057,58065315319 | 9,61908023789692 | 0.000283544297909342 |
| 913 | 26,1153633192252 | NA | NA | NA | NA | + | NA | NA | + | + | + | NA | NA | 0,105518242213311 | 6 | -522,467591495696 | 1057,58133683755 | 9,61976392225529 | 0.000283447387073699 |
| 745 | 23,8642166299369 | NA | NA | NA | + | NA | + | + | + | NA | + | NA | NA | 0,119849829844849 | 7 | -521,361724860966 | 1057,5916667762 | 9,63009386090494 | 0.000281987164263458 |
| 1578 | 23,9382644617248 | + | NA | NA | + | NA | + | NA | NA | NA | + | + | NA | 0,11984187663768 | 7 | -521,362343531077 | 1057,59290411642 | 9,63133120112775 | 0.000281812761187675 |
| 1978 | 24,7094395340959 | + | NA | NA | + | + | + | NA | + | + | + | + | NA | 0,162953683907615 | 10 | -517,923870801878 | 1057,59377334979 | 9,63220043449769 | 0.000281690307271811 |
| 3942 | 24,8814923899207 | + | NA | + | NA | NA | + | + | NA | + | + | + | + | 0,162952872109345 | 10 | -517,923937201091 | 1057,59390614821 | 9,63233323292366 | 0.000281671603878055 |
| 1641 | 23,8582632497147 | NA | NA | NA | + | NA | + | + | NA | NA | + | + | NA | 0,119761507784776 | 7 | -521,36859501086 | 1057,60540707598 | 9,64383416069245 | 0.000280056509713662 |
| 4018 | 24,9097778377762 | + | NA | NA | NA | + | + | NA | + | + | + | + | + | 0,162875284137672 | 10 | -517,930283037642 | 1057,60659782132 | 9,64502490602467 | 0.000279889821348584 |
| 789 | 26,2217593769328 | NA | NA | + | NA | + | NA | NA | NA | + | + | NA | NA | 0,105315984632533 | 6 | -522,483071221745 | 1057,61229628964 | 9,65072337435276 | 0.000279093484720922 |
| 739 | 23,8402918125305 | NA | + | NA | NA | NA | + | + | + | NA | + | NA | NA | 0,119678875267965 | 7 | -521,375021974616 | 1057,6182610035 | 9,65668808820556 | 0.000278262368292598 |
| 593 | 25,1239997715694 | NA | NA | NA | NA | + | NA | + | NA | NA | + | NA | NA | 0,0908279821416793 | 5 | -523,582904186605 | 1057,62382364039 | 9,66225072509587 | 0.000277489507319533 |
| 3681 | 24,0190917418403 | NA | NA | NA | NA | NA | + | + | NA | NA | + | + | + | 0,11964076241956 | 7 | -521,377986099571 | 1057,62418925341 | 9,66261633811496 | 0.000277438785067585 |

| | | | | | | | | | | | | | | | | | |
|---|---|---|---|---|---|---|---|---|---|---|---|---|---|---|---|---|---|
| 4067 | 24,6590391051374 | NA | + | NA | NA | NA | + | + | + | + | + | + | + | 0,162765838442093 | 10 | -517,939233483691 | 1057,62449871341 | 9,66292579812352 | 0.000277395860284145 |
| 738 | 24,0909568070185 | + | NA | NA | NA | NA | + | + | + | NA | + | NA | NA | 0,119585667601346 | 7 | -521,382270725147 | 1057,63275850456 | 9,67118558926654 | 0.000276252606733039 |
| 621 | 23,697386238415 | NA | NA | + | + | NA | + | + | NA | NA | + | NA | NA | 0,119583815625835 | 7 | -521,382414745318 | 1057,6330465449 | 9,6714736296085 | 0.000276212823650236 |
| 778 | 25,2389299590389 | + | NA | NA | + | NA | NA | NA | NA | + | + | NA | NA | 0,105177473133222 | 6 | -522,493670141957 | 1057,63349413007 | 9,67192121477774 | 0.00027615101618481 |
| 1974 | 24,9690981241428 | + | NA | + | NA | + | + | NA | + | + | + | + | NA | 0,16268947132728 | 10 | -517,945478077697 | 1057,63698790142 | 9,67541498613423 | 0.000275669033036902 |
| 2605 | 23,7763197970537 | NA | NA | + | + | NA | + | NA | NA | NA | + | NA | + | 0,119535846537719 | 7 | -521,386144989516 | 1057,6405070333 | 9,67893411800583 | 0.000275184401692256 |
| 2924 | 24,3056322753346 | + | + | NA | + | NA | + | + | NA | + | + | NA | + | 0,162652558155546 | 10 | -517,948496290275 | 1057,64302432658 | 9,68145141129048 | 0.000274838259650001 |
| 2785 | 24,0576337617549 | NA | NA | NA | NA | NA | + | + | + | NA | + | NA | + | 0,119497834655276 | 7 | -521,389100782293 | 1057,64641861885 | 9,68484570355918 | 0.000274372214545252 |
| 1591 | 24,170074018838 | NA | + | + | NA | + | + | NA | NA | NA | + | + | NA | 0,133866218703488 | 8 | -520,262631863252 | 1057,6502637265 | 9,68869081121352 | 0.000273845225938412 |
| 3618 | 24,0529717493793 | + | NA | NA | NA | NA | + | NA | NA | NA | + | + | + | 0,119342983431357 | 7 | -521,401140649891 | 1057,67049835405 | 9,70892543875584 | 0.000271088616119948 |
| 898 | 25,6794315284132 | + | NA | NA | NA | NA | NA | NA | + | + | + | NA | NA | 0,104917608778923 | 6 | -522,513550576923 | 1057,673255 | 9,71168208470999 | 0.000270715225836843 |
| 961 | 25,74359514024 | NA | NA | NA | NA | NA | NA | + | + | + | + | NA | NA | 0,104914004338893 | 6 | -522,513826287281 | 1057,67380642072 | 9,71223350542459 | 0.000270640597133627 |
| 841 | 25,2706999768142 | NA | NA | NA | + | NA | NA | + | NA | + | + | NA | NA | 0,104779870413405 | 6 | -522,524085653068 | 1057,69432515229 | 9,73275223700011 | 0.000267878190749011 |
| 3946 | 24,5037278712294 | + | NA | NA | + | NA | + | + | NA | + | + | + | + | 0,162334634793262 | 10 | -517,974485851416 | 1057,69500344886 | 9,73343053357416 | 0.000267787355723629 |
| 3044 | 24,6194071336079 | + | + | NA | NA | NA | + | + | + | + | + | NA | + | 0,162302164179891 | 10 | -517,977139700351 | 1057,70031114673 | 9,7387382314439 | 0.00026707763070291 |
| 2020 | 24,6112280604621 | + | + | NA | NA | NA | + | + | + | + | + | + | NA | 0,162238908569287 | 10 | -517,982309336255 | 1057,71065041854 | 9,74907750325042 | 0.000265700499295256 |
| 2823 | 24,9369458741402 | NA | + | + | NA | NA | NA | NA | NA | + | + | NA | + | 0,119050862522542 | 7 | -521,423847639527 | 1057,71591233332 | 9,75433941802794 | 0.000265002371378538 |
| 834 | 25,2853048455257 | + | NA | NA | NA | NA | NA | + | NA | + | + | NA | NA | 0,104598783708881 | 6 | -522,537933811492 | 1057,72202146914 | 9,76044855384703 | 0.000264194138668249 |
| 896 | 24,6883360204698 | + | + | + | + | + | + | + | NA | + | + | NA | NA | 0,176414753620537 | 11 | -516,813907904968 | 1057,73981580994 | 9,77824289464502 | 0.000261853984211711 |
| 568 | 24,2199691431977 | + | + | + | NA | + | + | NA | NA | NA | + | NA | NA | 0,133127319459111 | 8 | -520,321014977286 | 1057,76702995457 | 9,80545703928055 | 0.000258315049982191 |
| 537 | 25,0884300057429 | NA | NA | NA | + | + | NA | NA | NA | NA | + | NA | NA | 0,0897909919993831 | 5 | -523,660952508123 | 1057,77992028342 | 9,81834736813084 | 0.000256655520725782 |
| 1964 | 24,4168237246179 | + | + | NA | + | NA | + | NA | + | + | + | + | NA | 0,161804975964518 | 10 | -518,01776245193 | 1057,78155664989 | 9,81998373460101 | 0.000256445615363767 |
| 2947 | 25,0480517898367 | NA | + | NA | NA | NA | NA | NA | + | + | + | NA | + | 0,118604801100689 | 7 | -521,458506136906 | 1057,78522932808 | 9,82365641278466 | 0.00025597512637622 |
| 2841 | 25,2755821962779 | NA | NA | NA | + | + | NA | NA | NA | + | + | NA | + | 0,118536757693673 | 7 | -521,463791495984 | 1057,79580004623 | 9,83422713094069 | 0.000254625774962831 |
| 774 | 25,5670838370966 | + | NA | + | NA | NA | NA | NA | NA | + | + | NA | NA | 0,104068518241169 | 6 | -522,578468459179 | 1057,80309076451 | 9,8415178492221 | 0.00025369926232827 |
| 1801 | 25,367973423778 | NA | NA | NA | + | NA | NA | NA | NA | + | + | + | NA | 0,103979123782563 | 6 | -522,585299604285 | 1057,81675305472 | 9,85518013943238 | 0.000251972111778636 |
| 4011 | 24,4172243707768 | NA | + | NA | + | NA | + | NA | + | + | + | + | + | 0,161585864530771 | 10 | -518,035657301579 | 1057,81734634919 | 9,85577343389855 | 0.000251897376034486 |
| 837 | 25,6203021040474 | NA | NA | + | NA | NA | NA | + | NA | + | + | NA | NA | 0,103943374660577 | 6 | -522,58803120978 | 1057,82221626571 | 9,86064335042329 | 0.000251284762584216 |
| 3625 | 23,8492482420607 | NA | NA | NA | + | NA | + | NA | NA | NA | + | + | + | 0,118300216935636 | 7 | -521,482161930979 | 1057,83254091622 | 9,8709680009315 | 0.000249990891473539 |
| 3940 | 24,5152030117911 | + | + | NA | NA | NA | + | + | NA | + | + | + | + | 0,161489317709572 | 10 | -518,043540805267 | 1057,83311335657 | 9,87154044127601 | 0.000249919349276417 |
| 3843 | 24,8797393476333 | NA | + | NA | NA | NA | NA | NA | NA | + | + | + | + | 0,118256685484793 | 7 | -521,485542171851 | 1057,83930139796 | 9,87772848267423 | 0.000249147288638354 |
| 2827 | 24,557926869285 | NA | + | NA | + | NA | NA | NA | NA | + | + | NA | + | 0,118250911485286 | 7 | -521,485990513528 | 1057,84019808132 | 9,87862516602945 | 0.000249035610561833 |
| 2883 | 24,5535123428346 | NA | + | NA | NA | NA | NA | + | NA | + | + | NA | + | 0,118169049740987 | 7 | -521,492346629226 | 1057,85291031272 | 9,89133739742465 | 0.000247457731312031 |
| 2988 | 24,3956834435492 | + | + | NA | + | NA | + | NA | + | + | + | NA | + | 0,161345058064955 | 10 | -518,055318595814 | 1057,85666893766 | 9,89509602237013 | 0.000246993117624993 |
| 1588 | 24,224449491676 | + | + | NA | NA | + | + | NA | NA | NA | + | + | NA | 0,132504740968421 | 8 | -520,370168568621 | 1057,86533713724 | 9,90376422195209 | 0.000245924941269071 |
| 1794 | 25,4172669764237 | + | NA | NA | NA | NA | NA | NA | NA | + | + | + | NA | 0,10341832319875 | 6 | -522,628138070001 | 1057,90242998615 | 9,94085707086424 | 0.000241405948039612 |
| 2615 | 24,1532983657042 | NA | + | + | NA | + | + | NA | NA | NA | + | NA | + | 0,132237529168811 | 8 | -520,391254564625 | 1057,90750912925 | 9,94593621396029 | 0.000240793658168491 |
| 3585 | 24,6827852718897 | NA | NA | NA | NA | NA | NA | NA | NA | NA | + | + | + | 0,088913537406481 | 5 | -523,72692411314 | 1057,91186349345 | 9,95029057816419 | 0.000240269976807367 |
| 4066 | 24,8090567578201 | + | NA | NA | NA | NA | + | + | + | + | + | + | + | 0,160978902203411 | 10 | -518,085203569952 | 1057,91643888594 | 9,95486597064496 | 0.000239720940338531 |
| 2689 | 24,7574859828693 | NA | NA | NA | NA | NA | NA | NA | + | NA | + | NA | + | 0,088837032906538 | 5 | -523,732673109575 | 1057,92336148633 | 9,96178857103519 | 0.000238892628545822 |
| 2940 | 24,747689763365 | + | + | NA | + | + | + | + | NA | + | + | NA | + | 0,17526112954676 | 11 | -516,909740388028 | 1057,93148077606 | 9,9699078607664 | 0.000237924775212565 |
| 1703 | 24,3261150130822 | NA | + | + | NA | NA | + | NA | + | NA | + | + | NA | 0,13205634122756 | 8 | -520,405548625368 | 1057,93609725074 | 9,97452433544527 | 0.000237376221701846 |
| 788 | 25,5807738519358 | + | + | NA | NA | + | NA | NA | NA | + | + | NA | NA | 0,117475846290914 | 7 | -521,546146447392 | 1057,96050994905 | 9,99893703375733 | 0.000234496336855667 |
| 4006 | 24,9874074329767 | + | NA | + | NA | NA | + | NA | + | + | + | + | + | 0,16066165122489 | 10 | -518,111086478546 | 1057,96820470312 | 10,0066317878327 | 0.000233595874353946 |
| 3067 | 24,6843216603962 | NA | + | NA | + | + | + | + | + | + | + | NA | + | 0,17494476058147 | 11 | -516,93599800495 | 1057,9839960099 | 10,0224230946085 | 0.000231758744511524 |
| 1857 | 25,4423588399721 | NA | NA | NA | NA | NA | NA | + | NA | + | + | + | NA | 0,102876857237223 | 6 | -522,669474196316 | 1057,98510223879 | 10,0235293234953 | 0.00023163059084764 |
| 1919 | 24,6216930213254 | NA | + | + | + | + | + | + | NA | + | + | + | NA | 0,174918628188766 | 11 | -516,938166460383 | 1057,98833292077 | 10,0267600054754 | 0.000231256730496964 |
| 2563 | 24,081726422273 | NA | + | NA | NA | NA | NA | NA | NA | NA | + | NA | + | 0,08825954646324 | 5 | -523,776053265018 | 1058,01012179721 | 10,0485488819206 | 0.000228750992420868 |
| 1023 | 24,6226831379334 | NA | + | + | + | + | + | + | + | + | + | NA | NA | 0,174780547909355 | 11 | -516,949623166937 | 1058,01124633387 | 10,0496734185829 | 0.000228622409134659 |
| 3963 | 24,7128773825924 | NA | + | NA | + | + | + | + | NA | + | + | + | + | 0,174756458628155 | 11 | -516,951621690718 | 1058,01524338144 | 10,0536704661456 | 0.00022816595807912 |
| 4004 | 24,5528719091164 | + | + | NA | NA | NA | + | NA | + | + | + | + | + | 0,160353278481052 | 10 | -518,136235686966 | 1058,01850311996 | 10,0569302046733 | 0.000227794380290852 |
| 2880 | 24,6956146722581 | + | + | + | + | + | + | NA | NA | + | + | NA | + | 0,17473192558986 | 11 | -516,953656970079 | 1058,01931394016 | 10,0577410248684 | 0.000227702048866232 |
| 2565 | 24,6001557047133 | NA | NA | + | NA | NA | NA | NA | NA | NA | + | NA | + | 0,088094202996181 | 5 | -523,788468629776 | 1058,03495252673 | 10,0733796114378 | 0.000225928522661701 |
| 3635 | 24,1419490002912 | NA | + | NA | NA | + | + | NA | NA | NA | + | + | + | 0,13141038781823 | 8 | -520,456484144709 | 1058,03796828942 | 10,0763953741277 | 0.000225588105975563 |
| 2625 | 24,1795645291528 | NA | NA | NA | NA | NA | NA | + | NA | NA | + | NA | + | 0,087958294703530 | 5 | -523,798672071725 | 1058,05535941063 | 10,0937864953355 | 0.00022363499493292 |
| 791 | 25,8399939516815 | NA | + | + | NA | + | NA | NA | NA | + | + | NA | NA | 0,116803916743257 | 7 | -521,598254890754 | 1058,06472683577 | 10,1031539204812 | 0.000222590002028974 |
| 2820 | 24,7424399957154 | + | + | NA | NA | NA | NA | NA | NA | + | + | NA | + | 0,116791330845509 | 7 | -521,599230554778 | 1058,06667816382 | 10,1051052485298 | 0.000222372934881352 |
| 2953 | 25,1661905991614 | NA | NA | NA | + | NA | NA | NA | + | + | + | NA | + | 0,116788068598573 | 7 | -521,599483443249 | 1058,06718394076 | 10,1056110254708 | 0.000222316706440025 |
| 915 | 25,6384173466092 | NA | + | NA | NA | + | NA | NA | + | + | + | NA | NA | 0,11675100171886 | 7 | -521,602356791834 | 1058,07293063793 | 10,1113577226402 | 0.000221678829905911 |
| 705 | 25,02846281942 | NA | NA | NA | NA | NA | NA | + | + | NA | + | NA | NA | 0,087550078101702 | 5 | -523,829310178979 | 1058,11663562513 | 10,1550627098427 | 0.000216887140554655 |
| 1576 | 24,1807124976004 | + | + | + | NA | NA | + | NA | NA | NA | + | + | NA | 0,130850476861037 | 8 | -520,500604315323 | 1058,12620863065 | 10,1646357153554 | 0.000215851490207473 |
| 2612 | 24,1823120012781 | + | + | NA | NA | + | + | NA | NA | NA | + | NA | + | 0,130850170628011 | 8 | -520,500628438267 | 1058,12625687653 | 10,1646839612438 | 0.000215846283296826 |
| 618 | 23,6196455554747 | + | NA | NA | + | NA | + | + | NA | NA | + | NA | NA | 0,11639467075585 | 7 | -521,629972691383 | 1058,12816243703 | 10,1665895217398 | 0.000215640727161963 |
| 2936 | 24,9960120491938 | + | + | + | NA | + | + | + | NA | + | + | NA | + | 0,174073517325841 | 11 | -517,008256439166 | 1058,12851287833 | 10,1669399630416 | 0.00021560294576352 |
| 3063 | 24,9227538565922 | NA | + | + | NA | + | + | + | + | + | + | NA | + | 0,174059241292859 | 11 | -517,009439818191 | 1058,13087963638 | 10,1693067210908 | 0.000215347956663992 |
| 572 | 23,9279177098454 | + | + | NA | + | + | + | NA | NA | NA | + | NA | NA | 0,130789449323107 | 8 | -520,505411479694 | 1058,13582295939 | 10,1742500440971 | 0.000214816346660355 |
| 612 | 23,5967547934361 | + | + | NA | NA | NA | + | + | NA | NA | + | NA | NA | 0,116332045872104 | 7 | -521,634825013719 | 1058,1378670817 | 10,176294166411 | 0.000214596903377884 |
| 2596 | 23,5806270462177 | + | + | NA | NA | NA | + | NA | NA | NA | + | NA | + | 0,116325933852383 | 7 | -521,63529856888 | 1058,13881419202 | 10,177241276733 | 0.000214495303965124 |
| 901 | 26,0837709401217 | NA | NA | + | NA | NA | NA | NA | + | + | + | NA | NA | 0,101857388005393 | 6 | -522,747234005008 | 1058,14062185617 | 10,1790489408797 | 0.000214301523815358 |
| 556 | 23,4684813688311 | + | + | NA | + | NA | + | NA | NA | NA | + | NA | NA | 0,116311792003569 | 7 | -521,636394257227 | 1058,14100556872 | 10,1794326534261 | 0.000214260412667498 |
| 3959 | 24,9675409538833 | NA | + | + | NA | + | + | + | NA | + | + | + | + | 0,173910632606363 | 11 | -517,021757180657 | 1058,15551436131 | 10,1939414460242 | 0.000212711706977112 |
| 2897 | 25,4166443973771 | NA | NA | NA | NA | + | NA | + | NA | + | + | NA | + | 0,116197265506769 | 7 | -521,645266945605 | 1058,15875094547 | 10,1971780301824 | 0.000212367755688462 |
| 2658 | 23,7809757251323 | + | NA | NA | NA | NA | + | + | NA | NA | + | NA | + | 0,116185295198785 | 7 | -521,646194252546 | 1058,16060555936 | 10,1990326440653 | 0.00021217091687358 |
| 1811 | 25,7655036917599 | NA | + | NA | NA | + | NA | NA | NA | + | + | + | NA | 0,116097049417467 | 7 | -521,653030023927 | 1058,17427710212 | 10,2127041868266 | 0.000210725510853136 |
| 695 | 24,1153309093482 | NA | + | + | NA | + | + | NA | + | NA | + | NA | NA | 0,13053831634026 | 8 | -520,525189778769 | 1058,17537955754 | 10,2138066422481 | 0.000210609385120997 |
| 1595 | 23,9211045386784 | NA | + | NA | + | + | + | NA | NA | NA | + | + | NA | 0,130514055438191 | 8 | -520,527100174668 | 1058,17920034934 | 10,2176274340454 | 0.000210207421892142 |

| | | | | | | | | | | | | | | | | | | |
|---|---|---|---|---|---|---|---|---|---|---|---|---|---|---|---|---|---|---|
| **2942** | 25,048978527871 | + | NA | + | + | + | + | + | NA | + | + | NA | + | 0,173741040139109 | 11 | -517,035811068607 | 1058,18362213721 | 10,2220492219226 | 0.000209743188951233 |
| **3965** | 24,9686168974405 | NA | NA | + | + | + | + | + | NA | + | + | + | + | 0,173713170702286 | 11 | -517,038120293604 | 1058,18824058721 | 10,2266676719171 | 0.000209259403535975 |
| **575** | 23,8418411136316 | NA | + | + | + | + | + | NA | NA | NA | + | NA | NA | 0,130448771411135 | 8 | -520,532240623221 | 1058,18948124644 | 10,2279083311505 | 0.000209129633984496 |
| **960** | 24,6537863658175 | + | + | + | + | + | + | NA | + | + | + | NA | NA | 0,173678672089212 | 11 | -517,040978695633 | 1058,19395739127 | 10,232384475976 | 0.000208662110091301 |
| **530** | 25,2702730671652 | + | NA | NA | NA | + | NA | NA | NA | NA | + | NA | NA | 0,0870040092176025 | 5 | -523,870273175515 | 1058,19856161821 | 10,2369887029147 | 0.000208182298739113 |
| **2569** | 24,171394283361 | NA | NA | NA | + | NA | NA | NA | NA | NA | + | NA | + | 0,0870028204486344 | 5 | -523,870362323518 | 1058,19873991421 | 10,2371669989202 | 0.000208163740530197 |
| **1797** | 25,8400794018669 | NA | NA | + | NA | NA | NA | NA | NA | + | + | + | NA | 0,101378615409358 | 6 | -522,783721848307 | 1058,21359754277 | 10,2520246274764 | 0.000206623060560527 |
| **1856** | 24,6210437971772 | + | + | + | + | + | + | NA | NA | + | + | + | NA | 0,17355058837454 | 11 | -517,051590100628 | 1058,21518020126 | 10,2536072859652 | 0.000206459618366959 |
| **1921** | 25,8933533711233 | NA | NA | NA | NA | NA | NA | NA | + | + | + | + | NA | 0,101360198607083 | 6 | -522,785125027139 | 1058,21640390043 | 10,2548309851418 | 0.00020633333477169 |
| **3007** | 24,6830844169424 | NA | + | + | + | + | + | NA | + | + | + | NA | + | 0,17353191892044 | 11 | -517,053136679433 | 1058,21827335887 | 10,2567004435753 | 0.000206140559085861 |
| **1715** | 24,0895219649759 | NA | + | NA | NA | + | + | NA | + | NA | + | + | NA | 0,130205885914415 | 8 | -520,551361982055 | 1058,22772396411 | 10,2661510488194 | 0.000205168780344007 |
| **3069** | 25,0079343541122 | NA | NA | + | + | + | + | + | + | + | + | NA | + | 0,173461336072064 | 11 | -517,058983451695 | 1058,22996690339 | 10,2683939880999 | 0.0002049388187573 |
| **579** | 24,1239953655034 | NA | + | NA | NA | NA | NA | + | NA | NA | + | NA | NA | 0,0867615508958795 | 5 | -523,888453176523 | 1058,23492162022 | 10,2733487049311 | 0.000204431740216953 |
| **2602** | 23,6456565746023 | + | NA | NA | + | NA | + | NA | NA | NA | + | NA | + | 0,115672013309955 | 7 | -521,685944988149 | 1058,24010703056 | 10,2785341152714 | 0.000203902395501063 |
| **533** | 25,5190670432935 | NA | NA | + | NA | + | NA | NA | NA | NA | + | NA | NA | 0,0867228197102401 | 5 | -523,891356869976 | 1058,24072900713 | 10,2791560918367 | 0.000203838994104315 |
| **2834** | 25,44983417059 | + | NA | NA | NA | + | NA | NA | NA | + | + | NA | + | 0,115650348098443 | 7 | -521,687622327132 | 1058,24346170853 | 10,2818887932369 | 0.000203560668740342 |
| **1651** | 24,0518976857912 | NA | + | NA | NA | + | + | + | NA | NA | + | + | NA | 0,13008465639548 | 8 | -520,560903878263 | 1058,24680775653 | 10,2852348412348 | 0.000203220391581597 |
| **1916** | 24,6142103531835 | + | + | NA | + | + | + | + | NA | + | + | + | NA | 0,173152012146275 | 11 | -517,08460059908 | 1058,28120119816 | 10,319628282869 | 0.000199755544633092 |
| **1539** | 24,8522189703184 | NA | + | NA | NA | NA | NA | NA | NA | NA | + | + | NA | 0,0864271601943126 | 5 | -523,913518534438 | 1058,28505233605 | 10,3234794207613 | 0.000199371271650321 |
| **1020** | 24,5715363229949 | + | + | NA | + | + | + | + | + | + | + | NA | NA | 0,173124815148274 | 11 | -517,086852503117 | 1058,28570500623 | 10,3241320909442 | 0.000199306220422977 |
| **3886** | 24,5707940097728 | + | NA | + | + | NA | + | NA | NA | + | + | + | + | 0,158707176303243 | 10 | -518,270326881696 | 1058,28668550942 | 10,3251125941338 | 0.000199208534177939 |
| **3903** | 24,6226485504904 | NA | + | + | + | + | + | NA | NA | + | + | + | + | 0,173117093478418 | 11 | -517,087491841789 | 1058,28698368358 | 10,3254107682867 | 0.000199178836973761 |
| **1553** | 25,4836160760359 | NA | NA | NA | NA | + | NA | NA | NA | NA | + | + | NA | 0,0863876049359884 | 5 | -523,916482922736 | 1058,29098111265 | 10,3294081973575 | 0.000198781132917999 |
| **657** | 25,4250583876943 | NA | NA | NA | NA | + | NA | NA | + | NA | + | NA | NA | 0,086359217887645 | 5 | -523,918610253211 | 1058,2952357736 | 10,3336628583072 | 0.000198358709232809 |
| **2829** | 25,0207174897797 | NA | NA | + | + | NA | NA | NA | NA | + | + | NA | + | 0,115304770366638 | 7 | -521,714371696063 | 1058,29696044639 | 10,3353875310984 | 0.000198187731029416 |
| **631** | 23,9730045513159 | NA | + | + | NA | + | + | + | NA | NA | + | NA | NA | 0,129694988740003 | 8 | -520,591565365435 | 1058,30813073087 | 10,346557815579 | 0.00019708390973018 |
| **4010** | 24,6190619058553 | + | NA | NA | + | NA | + | NA | + | + | + | + | + | 0,15851190023472 | 10 | -518,286216633643 | 1058,31846501332 | 10,3568920980274 | 0.000196068175815828 |
| **2659** | 23,5349788722909 | NA | + | NA | NA | NA | + | + | NA | NA | + | NA | + | 0,115135199518604 | 7 | -521,727493471231 | 1058,32320399673 | 10,3616310814352 | 0.000195604143876853 |
| **2665** | 23,5510588315205 | NA | NA | NA | + | NA | + | + | NA | NA | + | NA | + | 0,115127604478549 | 7 | -521,728081133774 | 1058,32437932181 | 10,3628064065208 | 0.000195489228417346 |
| **574** | 24,267863265482 | + | NA | + | + | + | + | NA | NA | NA | + | NA | NA | 0,129566990650726 | 8 | -520,60163406149 | 1058,32826812298 | 10,3666952076899 | 0.000195109488350386 |
| **2562** | 24,4087375203695 | + | NA | NA | NA | NA | NA | NA | NA | NA | + | NA | + | 0,0861281595832692 | 5 | -523,935923346862 | 1058,3298619609 | 10,3682890456084 | 0.000194954063838579 |
| **3857** | 25,5282912713294 | NA | NA | NA | NA | + | NA | NA | NA | + | + | + | + | 0,115083962545101 | 7 | -521,731457809509 | 1058,33113267328 | 10,3695597579901 | 0.000194830237908138 |
| **2837** | 25,5988097104401 | NA | NA | + | NA | + | NA | NA | NA | + | + | NA | + | 0,115073912618066 | 7 | -521,732235371662 | 1058,33268779759 | 10,3711148822972 | 0.000194678804170979 |
| **2619** | 23,9304922809999 | NA | + | NA | + | + | + | NA | NA | NA | + | NA | + | 0,129423401543668 | 8 | -520,612927430542 | 1058,35085486108 | 10,3892819457933 | 0.000192918440337039 |
| **2961** | 25,5715524601285 | NA | NA | NA | NA | + | NA | NA | + | + | + | NA | + | 0,114901592610119 | 7 | -521,745566385704 | 1058,35934982567 | 10,3977769103806 | 0.000192100760447339 |
| **692** | 24,0803110500888 | + | + | NA | NA | + | + | NA | + | NA | + | NA | NA | 0,129278265323278 | 8 | -520,624340589195 | 1058,37368117839 | 10,4121082630998 | 0.00019072914870147 |
| **628** | 24,0623079121175 | + | + | NA | NA | + | + | + | NA | NA | + | NA | NA | 0,129259856135572 | 8 | -520,625788106962 | 1058,37657621392 | 10,4150032986338 | 0.000190453264591784 |
| **519** | 24,7671938983505 | NA | + | + | NA | NA | NA | NA | NA | NA | + | NA | NA | 0,0857947643315368 | 5 | -523,960896791843 | 1058,37980885086 | 10,4182359355716 | 0.000190145680106966 |
| **595** | 24,5633596091042 | NA | + | NA | NA | + | NA | + | NA | NA | + | NA | NA | 0,100267147956108 | 6 | -522,868353279481 | 1058,38286040512 | 10,4212874898258 | 0.000189855781394097 |
| **3009** | 25,3001265961706 | NA | NA | NA | NA | NA | NA | + | + | + | + | NA | + | 0,114696700952746 | 7 | -521,761413830895 | 1058,39104471605 | 10,429471800762 | 0.000189080449491411 |
| **3060** | 24,8417915742619 | + | + | NA | NA | + | + | + | + | + | + | NA | + | 0,17242394747013 | 11 | -517,144858635507 | 1058,40171727101 | 10,440144355724 | 0.00018807415107841 |
| **2675** | 24,0698981104612 | NA | + | NA | NA | + | + | + | NA | NA | + | NA | + | 0,129082515422013 | 8 | -520,63973087301 | 1058,40446174602 | 10,4428888307286 | 0.00018781624566929 |
| **2739** | 24,0714375120622 | NA | + | NA | NA | + | + | NA | + | NA | + | NA | + | 0,129029149951432 | 8 | -520,643925982837 | 1058,41285196567 | 10,4512790503827 | 0.000187029986266789 |
| **4083** | 24,8185928642626 | NA | + | NA | NA | + | + | + | + | + | + | + | + | 0,172309223454056 | 11 | -517,154348898897 | 1058,42069779779 | 10,4591248825031 | 0.000186297720577423 |
| **1912** | 24,8786226327105 | + | + | + | NA | + | + | + | NA | + | + | + | NA | 0,172246646904334 | 11 | -517,159524836663 | 1058,43104967333 | 10,4694767580363 | 0.000185335946358597 |
| **1709** | 24,3774823768964 | NA | NA | + | + | NA | + | NA | + | NA | + | + | NA | 0,128903764902585 | 8 | -520,653781608585 | 1058,43256321717 | 10,47099030188 | 0.000185195742376197 |
| **581** | 24,8172712500451 | NA | NA | + | NA | NA | NA | + | NA | NA | + | NA | NA | 0,0854290006008188 | 5 | -523,988284374238 | 1058,43458401565 | 10,473011100361 | 0.000185008715240705 |
| **3884** | 24,1723213432219 | + | + | NA | + | NA | + | NA | NA | + | + | + | + | 0,157796431164623 | 10 | -518,344403386452 | 1058,43483851894 | 10,4732656036447 | 0.000184985174075791 |
| **2614** | 24,5266089638878 | + | NA | + | NA | + | + | NA | NA | NA | + | NA | + | 0,128862101169192 | 8 | -520,657056184104 | 1058,43911236821 | 10,4775394529181 | 0.000184590296761677 |
| **619** | 23,3935533820964 | NA | + | NA | + | NA | + | + | NA | NA | + | NA | NA | 0,114315244352772 | 7 | -521,790908014507 | 1058,45003308328 | 10,4884601679878 | 0.000183585114568477 |
| **2043** | 24,5328930799285 | NA | + | NA | + | + | + | + | + | + | + | + | NA | 0,172118338048397 | 11 | -517,170136513535 | 1058,45227302707 | 10,4907001117795 | 0.0001833796194955 |
| **1016** | 24,8244098136708 | + | + | + | NA | + | + | + | + | + | + | NA | NA | 0,172106446396603 | 11 | -517,171119919441 | 1058,45423983888 | 10,4926669235908 | 0.000183199371537685 |
| **2603** | 23,4136334366466 | NA | + | NA | + | NA | + | NA | NA | NA | + | NA | + | 0,114274832586996 | 7 | -521,7940319041 | 1058,45628086246 | 10,4947079471731 | 0.000183012509782434 |
| **4089** | 24,8567062565517 | NA | NA | NA | + | + | + | + | + | + | + | + | + | 0,172087625206629 | 11 | -517,172676349663 | 1058,45735269933 | 10,4957797840345 | 0.000182914456282073 |
| **3896** | 24,8690518820368 | + | + | + | NA | + | + | NA | NA | + | + | + | + | 0,172072191108413 | 11 | -517,173952655797 | 1058,45990531159 | 10,4983323963027 | 0.000182681150356075 |
| **3962** | 24,9059254065267 | + | NA | NA | + | + | + | + | NA | + | + | + | + | 0,172067662046145 | 11 | -517,174327177211 | 1058,46065435442 | 10,4990814391324 | 0.000182612745163578 |
| **3000** | 24,8655756454705 | + | + | + | NA | + | + | NA | + | + | + | NA | + | 0,172029478848432 | 11 | -517,177484576187 | 1058,46696915237 | 10,5053962370828 | 0.000182037073160486 |
| **2621** | 24,2370474921621 | NA | NA | + | + | + | + | NA | NA | NA | + | NA | + | 0,128658040076643 | 8 | -520,673092175652 | 1058,4711843513 | 10,5096114360128 | 0.0001816538159407 |
| **1983** | 24,5956894327098 | NA | + | + | + | + | + | NA | + | + | + | + | NA | 0,171945871649882 | 11 | -517,184397613727 | 1058,48079522745 | 10,5192223121642 | 0.000180782983817697 |
| **649** | 24,9626392698791 | NA | NA | NA | + | NA | NA | NA | + | NA | + | NA | NA | 0,0851071924886072 | 5 | -524,012371604827 | 1058,48275847683 | 10,5211855615382 | 0.000180605609849363 |
| **3066** | 24,9251777388184 | + | NA | NA | + | + | + | + | + | + | + | NA | + | 0,171900104139796 | 11 | -517,188181591876 | 1058,48836318375 | 10,5267902684611 | 0.000180100197596092 |
| **686** | 24,3126818680046 | + | NA | + | + | NA | + | NA | + | NA | + | NA | NA | 0,12852334474126 | 8 | -520,683675053539 | 1058,49235010708 | 10,530777191788 | 0.000179741532368068 |
| **1590** | 24,5361557059138 | + | NA | + | NA | + | + | NA | NA | NA | + | + | NA | 0,128495227071645 | 8 | -520,685884024479 | 1058,49676804896 | 10,5351951336668 | 0.000179344926752881 |
| **3637** | 24,4751443134466 | NA | NA | + | NA | + | + | NA | NA | NA | + | + | + | 0,128482072870424 | 8 | -520,68691741597 | 1058,49883483194 | 10,5372619166499 | 0.000179159688959587 |
| **3956** | 24,8981471725552 | + | + | NA | NA | + | + | + | NA | + | + | + | + | 0,171804700057888 | 11 | -517,196068763522 | 1058,50413752704 | 10,542564611754 | 0.000178685303515079 |
| **687** | 23,9781344230175 | NA | + | + | + | NA | + | NA | + | NA | + | NA | NA | 0,128444574332472 | 8 | -520,689863208262 | 1058,50472641652 | 10,5431535012333 | 0.000178632698312446 |
| **643** | 24,9129079283278 | NA | + | NA | NA | NA | NA | NA | + | NA | + | NA | NA | 0,0849197712382892 | 5 | -524,026396121377 | 1058,51080750993 | 10,5492345946384 | 0.000178090382137259 |
| **3849** | 24,9371049886996 | NA | NA | NA | + | NA | NA | NA | NA | + | + | + | + | 0,113918782557957 | 7 | -521,821548945608 | 1058,51131494548 | 10,5497420301897 | 0.000178045203173234 |
| **2039** | 24,7918496031997 | NA | + | + | NA | + | + | + | + | + | + | + | NA | 0,171713173553121 | 11 | -517,203634517716 | 1058,51926903543 | 10,5576961201418 | 0.000177338515588062 |
| **2889** | 24,8472393095009 | NA | NA | NA | + | NA | NA | + | NA | + | + | NA | + | 0,113866274584506 | 7 | -521,825606047195 | 1058,51942914865 | 10,5578562333624 | 0.000177324319035905 |
| **2677** | 24,4282739267226 | NA | NA | + | NA | + | + | + | NA | NA | + | NA | + | 0,128330455990765 | 8 | -520,698827283303 | 1058,52265456661 | 10,5610816513145 | 0.000177038576986241 |
| **680** | 24,239624982182 | + | + | + | NA | NA | + | NA | + | NA | + | NA | NA | 0,128318503413177 | 8 | -520,699766098715 | 1058,52453219743 | 10,562959282139 | 0.000176872448435823 |
| **2741** | 24,489961037838 | NA | NA | + | NA | + | + | NA | + | NA | + | NA | + | 0,128302895776602 | 8 | -520,700991981428 | 1058,52698396286 | 10,5654110475655 | 0.000176655756405481 |
| **701** | 24,2252201663578 | NA | NA | + | + | + | + | NA | + | NA | + | NA | NA | 0,128278105838779 | 8 | -520,702939031661 | 1058,53087806332 | 10,5693051480305 | 0.000176312133408073 |
| **2946** | 25,3153429574846 | + | NA | NA | NA | NA | NA | NA | + | + | + | NA | + | 0,113789444372659 | 7 | -521,831542007355 | 1058,53130106897 | 10,5697281536834 | 0.000176274846836765 |
| **1583** | 23,8377744884482 | NA | + | + | + | NA | + | NA | NA | NA | + | + | NA | 0,128262748159147 | 8 | -520,704145226113 | 1058,53329045223 | 10,5717175369346 | 0.000176099594898084 |

| | | | | | | | | | | | | | | | | | | |
|---|---|---|---|---|---|---|---|---|---|---|---|---|---|---|---|---|---|---|
| **2826** | 24,8648667866732 | + | NA | NA | + | NA | NA | NA | NA | + | + | NA | + | 0,113770395494568 | 7 | -521,833013658481 | 1058,53424437123 | 10,5726714559348 | 0.000176015622550624 |
| **2949** | 25,5481400082694 | NA | NA | + | NA | NA | NA | NA | + | + | + | NA | + | 0,113676394580427 | 7 | -521,840275384452 | 1058,54876782317 | 10,5871949078773 | 0.000174742075006098 |
| **4023** | 24,8193462077393 | NA | + | + | NA | + | + | NA | + | + | + | + | + | 0,171519036097171 | 11 | -517,21967952195 | 1058,5513590439 | 10,5897861286094 | 0.00017451582396059 |
| **963** | 25,2095387761987 | NA | + | NA | NA | NA | NA | + | + | + | + | NA | NA | 0,11360181795061 | 7 | -521,846036004764 | 1058,56028906379 | 10,5987161485016 | 0.000173738346085746 |
| **3623** | 23,989091670465 | NA | + | + | NA | NA | + | NA | NA | NA | + | + | + | 0,128077414985019 | 8 | -520,71869964681 | 1058,56239929362 | 10,6008263783292 | 0.000173555128840312 |
| **1022** | 24,915160836655 | + | NA | + | + | + | + | + | + | + | + | NA | NA | 0,171384411070999 | 11 | -517,230803757487 | 1058,57360751497 | 10,6120345996826 | 0.000172585226946958 |
| **1918** | 24,8757152271593 | + | NA | + | + | + | + | + | NA | + | + | + | NA | 0,171368429691298 | 11 | -517,232124199078 | 1058,57624839816 | 10,6146754828662 | 0.000172357488625829 |
| **1717** | 24,5108613837741 | NA | NA | + | NA | + | + | NA | + | NA | + | + | NA | 0,127942901145829 | 8 | -520,729261232094 | 1058,58352246419 | 10,6219495488979 | 0.000171731757341608 |
| **1639** | 23,9895336981066 | NA | + | + | NA | NA | + | + | NA | NA | + | + | NA | 0,127931132625927 | 8 | -520,730185180199 | 1058,5853703604 | 10,6237974451083 | 0.000171573159389232 |
| **1597** | 24,2152342210633 | NA | NA | + | + | + | + | NA | NA | NA | + | + | NA | 0,127916441267798 | 8 | -520,731338583244 | 1058,58767716649 | 10,6261042511969 | 0.000171375380466199 |
| **2618** | 24,2477919718257 | + | NA | NA | + | + | + | NA | NA | NA | + | NA | + | 0,127874584153165 | 8 | -520,734624634717 | 1058,59424926943 | 10,6326763541435 | 0.000170813156399255 |
| **2885** | 25,1645156486707 | NA | NA | + | NA | NA | NA | + | NA | + | + | NA | + | 0,113379907741465 | 7 | -521,863174440844 | 1058,59456593595 | 10,6329930206612 | 0.000170786113136535 |
| **694** | 24,501969009661 | + | NA | + | NA | + | + | NA | + | NA | + | NA | NA | 0,127865215475388 | 8 | -520,735360114312 | 1058,59572022862 | 10,6341473133341 | 0.000170687572995795 |
| **1601** | 24,838925928166 | NA | NA | NA | NA | NA | NA | + | NA | NA | + | + | NA | 0,0843008669357579 | 5 | -524,072687632616 | 1058,60339053241 | 10,6418176171164 | 0.00017003421389394 |
| **839** | 25,0673320343589 | NA | + | + | NA | NA | NA | + | NA | + | + | NA | NA | 0,113304755442043 | 7 | -521,868977586282 | 1058,60617222683 | 10,6445993115362 | 0.000169797886667586 |
| **3055** | 24,6809589167361 | NA | + | + | + | NA | + | + | + | + | + | NA | + | 0,171119850870509 | 11 | -517,252659438187 | 1058,61731887637 | 10,6557459610835 | 0.00016885418013359 |
| **1008** | 24,6470674178372 | + | + | + | + | NA | + | + | + | + | + | NA | NA | 0,171010257960444 | 11 | -517,261711016429 | 1058,63542203286 | 10,6738491175672 | 0.000167332679688975 |
| **630** | 24,4270086257134 | + | NA | + | NA | + | + | + | NA | NA | + | NA | NA | 0,127596206579472 | 8 | -520,756475046936 | 1058,63795009387 | 10,6763771785811 | 0.000167121299700561 |
| **525** | 24,8038951497167 | NA | NA | + | + | NA | NA | NA | NA | NA | + | NA | NA | 0,084069507934384 | 5 | -524,089984310145 | 1058,63798388747 | 10,6764109721751 | 0.000167118475909746 |
| **1653** | 24,4068662841552 | NA | NA | + | NA | + | + | + | NA | NA | + | + | NA | 0,127581575592452 | 8 | -520,757623269586 | 1058,64024653917 | 10,6786736238821 | 0.000166929517364346 |
| **1594** | 24,250845016395 | + | NA | NA | + | + | + | NA | NA | NA | + | + | NA | 0,127549984451586 | 8 | -520,760102439517 | 1058,64520487903 | 10,6836319637437 | 0.000166516183298735 |
| **699** | 23,843190485862 | NA | + | NA | + | + | + | NA | + | NA | + | NA | NA | 0,127496697672364 | 8 | -520,764284009237 | 1058,65356801847 | 10,6919951031828 | 0.000165821338053235 |
| **2031** | 24,651627042812 | NA | + | + | + | NA | + | + | + | + | + | + | NA | 0,170893032409789 | 11 | -517,271391671593 | 1058,65478334319 | 10,693210427896 | 0.000165720605277032 |
| **2681** | 24,1592987456153 | NA | NA | NA | + | + | + | + | NA | NA | + | NA | + | 0,127456600788174 | 8 | -520,767430360943 | 1058,65986072189 | 10,6982878065953 | 0.000165300425719751 |
| **3900** | 24,5891387188627 | + | + | NA | + | + | + | NA | NA | + | + | + | + | 0,170858753632905 | 11 | -517,274222203962 | 1058,66044440792 | 10,6988714926331 | 0.000165252190983314 |
| **3004** | 24,5914109702388 | + | + | NA | + | + | + | NA | + | + | + | NA | + | 0,170855772013641 | 11 | -517,27446840237 | 1058,66093680474 | 10,6993638894496 | 0.000165211511164791 |
| **1702** | 24,6151857052844 | + | NA | + | NA | NA | + | NA | + | NA | + | + | NA | 0,127403885175676 | 8 | -520,771566668389 | 1058,66813333678 | 10,7065604214881 | 0.000164618104456037 |
| **2727** | 24,1902088367931 | NA | + | + | NA | NA | + | NA | + | NA | + | NA | + | 0,127363967059282 | 8 | -520,774698659742 | 1058,67439731948 | 10,7128244041942 | 0.000164103328533999 |
| **2822** | 25,1999427916533 | + | NA | + | NA | NA | NA | NA | NA | + | + | NA | + | 0,112846347621656 | 7 | -521,904364494634 | 1058,67694604353 | 10,7153731282403 | 0.000163894334679453 |
| **2045** | 24,8291036431541 | NA | NA | + | + | + | + | + | + | + | + | + | NA | 0,170753561534279 | 11 | -517,28290759591 | 1058,67781519182 | 10,7162422765302 | 0.000163823125912963 |
| **3969** | 25,4182381104945 | NA | NA | NA | NA | NA | NA | NA | + | + | + | + | + | 0,112830087250761 | 7 | -521,905619382442 | 1058,67945581915 | 10,7178829038564 | 0.000163688794668856 |
| **1976** | 24,804325494667 | + | + | + | NA | + | + | NA | + | + | + | + | NA | 0,170741240980635 | 11 | -517,283924794546 | 1058,67984958909 | 10,7182766738015 | 0.000163656569977397 |
| **634** | 24,1388950996223 | + | NA | NA | + | + | + | + | NA | NA | + | NA | NA | 0,127308789885606 | 8 | -520,779027647061 | 1058,68305529412 | 10,7214823788315 | 0.000163394462748341 |
| **637** | 24,0687545822175 | NA | NA | + | + | + | + | + | NA | NA | + | NA | NA | 0,127257625242124 | 8 | -520,783041582358 | 1058,69108316472 | 10,7295102494261 | 0.000162739922466916 |
| **523** | 24,3274693383782 | NA | + | NA | + | NA | NA | NA | NA | NA | + | NA | NA | 0,0836596560848473 | 5 | -524,120614610426 | 1058,69924448803 | 10,7376715727362 | 0.000162077189020682 |
| **3697** | 24,4146444984409 | NA | NA | NA | NA | + | + | + | NA | NA | + | + | + | 0,127169461575039 | 8 | -520,789957589178 | 1058,70491517836 | 10,7433422630656 | 0.000161618295115215 |
| **2899** | 24,9611297231961 | NA | + | NA | NA | + | NA | + | NA | + | + | NA | + | 0,127159189980987 | 8 | -520,790763299887 | 1058,70652659977 | 10,744953684484 | 0.000161488129968808 |
| **3634** | 24,4585790164116 | + | NA | NA | NA | + | + | NA | NA | NA | + | + | + | 0,127128415682489 | 8 | -520,79317719972 | 1058,71135439944 | 10,7497814841495 | 0.00016109878390914 |
| **2733** | 24,2486847084858 | NA | NA | + | + | NA | + | NA | + | NA | + | NA | + | 0,127127522345763 | 8 | -520,793247270733 | 1058,71149454147 | 10,7499216261758 | 0.000161087495949615 |
| **635** | 23,7799636962081 | NA | + | NA | + | + | + | + | NA | NA | + | NA | NA | 0,127121371975294 | 8 | -520,793729687877 | 1058,71245937575 | 10,7508864604642 | 0.000161009803321485 |
| **2843** | 24,9744554097693 | NA | + | NA | + | + | NA | NA | NA | + | + | NA | + | 0,12711784051304 | 8 | -520,794006683954 | 1058,71301336791 | 10,7514404526182 | 0.000160965210413923 |
| **3749** | 24,5646559847535 | NA | NA | + | NA | NA | + | NA | + | NA | + | + | + | 0,127104819078022 | 8 | -520,795028032046 | 1058,71505606409 | 10,7534831488019 | 0.000160800892830421 |
| **2882** | 24,9710925751095 | + | NA | NA | NA | NA | NA | + | NA | + | + | NA | + | 0,112560477032937 | 7 | -521,926423096291 | 1058,72106324685 | 10,7594903315551 | 0.000160318637267255 |
| **2674** | 24,4167695893852 | + | NA | NA | NA | + | + | + | NA | NA | + | NA | + | 0,127044498196941 | 8 | -520,799759156525 | 1058,72451831305 | 10,762945397759 | 0.000160041920601929 |
| **585** | 24,4576859034075 | NA | NA | NA | + | NA | NA | + | NA | NA | + | NA | NA | 0,0834814658965907 | 5 | -524,13392739314 | 1058,72587005346 | 10,7642971381656 | 0.000159933789581902 |
| **3845** | 25,3311129452009 | NA | NA | + | NA | NA | NA | NA | NA | + | + | + | + | 0,112435496244372 | 7 | -521,936064744317 | 1058,7403465429 | 10,7787736276071 | 0.00015878032922399 |
| **3641** | 24,2211963517712 | NA | NA | NA | + | + | + | NA | NA | NA | + | + | + | 0,126916246822002 | 8 | -520,80981716149 | 1058,74463432298 | 10,7830614076895 | 0.000158440286294992 |
| **783** | 25,1586838019937 | NA | + | + | + | NA | NA | NA | NA | + | + | NA | NA | 0,112380726215011 | 7 | -521,940289552831 | 1058,74879615993 | 10,7872232446346 | 0.000158110927779407 |
| **698** | 24,1891280052943 | + | NA | NA | + | + | + | NA | + | NA | + | NA | NA | 0,126874954541574 | 8 | -520,813055159188 | 1058,75111031838 | 10,789537403085 | 0.000157928086710796 |
| **3905** | 25,0698408035174 | NA | NA | NA | NA | NA | NA | + | NA | + | + | + | + | 0,112361750569467 | 7 | -521,941753220847 | 1058,75172349596 | 10,790150580666 | 0.000157879675151302 |
| **755** | 23,9358029822856 | NA | + | NA | NA | + | + | + | + | NA | + | NA | NA | 0,126837360355532 | 8 | -520,816003032018 | 1058,75700606404 | 10,7954331487463 | 0.000157463220314933 |
| **645** | 25,4634534242129 | NA | NA | + | NA | NA | NA | NA | + | NA | + | NA | NA | 0,0831837745168243 | 5 | -524,156162469586 | 1058,77034020635 | 10,8087672910576 | 0.000156416893663561 |
| **1545** | 24,8581995222096 | NA | NA | NA | + | NA | NA | NA | NA | NA | + | + | NA | 0,0831489832908063 | 5 | -524,158760614562 | 1058,7755364963 | 10,8139635810094 | 0.000156011027375188 |
| **1765** | 24,5157572644843 | NA | NA | + | NA | NA | + | + | + | NA | + | + | NA | 0,126675238217209 | 8 | -520,828714061955 | 1058,78242812391 | 10,8208552086203 | 0.000155474367569148 |
| **2036** | 24,7170518291319 | + | + | NA | NA | + | + | + | + | + | + | + | NA | 0,170109970420404 | 11 | -517,336022991488 | 1058,78404598298 | 10,8224730676855 | 0.000155348650616595 |
| **1657** | 24,1359069176545 | NA | NA | NA | + | + | + | + | NA | NA | + | + | NA | 0,126647830442504 | 8 | -520,830862709042 | 1058,78672541808 | 10,8251525027943 | 0.000155140666653303 |
| **743** | 24,0920875827283 | NA | + | + | NA | NA | + | + | + | NA | + | NA | NA | 0,126633099043906 | 8 | -520,83201755703 | 1058,78903511406 | 10,8274621987687 | 0.000154961606180398 |
| **3842** | 25,104871912391 | + | NA | NA | NA | NA | NA | NA | NA | + | + | + | + | 0,112119756052038 | 7 | -521,960416491136 | 1058,78905003654 | 10,8274771212452 | 0.000154960449979243 |
| **2745** | 24,1928168094618 | NA | NA | NA | + | + | + | NA | + | NA | + | NA | + | 0,126629374654107 | 8 | -520,832309522427 | 1058,78961904485 | 10,8280461295628 | 0.000154916369357632 |
| **578** | 24,5481924591549 | + | NA | NA | NA | NA | NA | + | NA | NA | + | NA | NA | 0,0830346241791257 | 5 | -524,16730004911 | 1058,7926153654 | 10,8310424501046 | 0.000154684453574632 |
| **757** | 24,3537012005499 | NA | NA | + | NA | + | + | + | + | NA | + | NA | NA | 0,126606577043892 | 8 | -520,834096664124 | 1058,79319332825 | 10,8316204129569 | 0.000154639759098885 |
| **1650** | 24,4206063835359 | + | NA | NA | NA | + | + | + | NA | NA | + | + | NA | 0,126569835988793 | 8 | -520,836976756996 | 1058,79895351399 | 10,837380598701 | 0.000154195022979057 |
| **2738** | 24,4063819668574 | + | NA | NA | NA | + | + | NA | + | NA | + | NA | + | 0,126540447893466 | 8 | -520,83928037188 | 1058,80356074376 | 10,8419878284685 | 0.000153840225843998 |
| **4027** | 24,5516990521404 | NA | + | NA | + | + | + | NA | + | + | + | + | + | 0,169932395516023 | 11 | -517,350670952272 | 1058,81334190455 | 10,8517689892544 | 0.000153089694614415 |
| **3761** | 24,4018950552643 | NA | NA | NA | NA | + | + | NA | + | NA | + | + | + | 0,126474480585392 | 8 | -520,84445100239 | 1058,81390200478 | 10,8523290894886 | 0.000153046827830213 |
| **907** | 25,2946693564863 | NA | + | NA | + | NA | NA | NA | + | + | + | NA | NA | 0,11190781423937 | 7 | -521,976757839857 | 1058,82173273398 | 10,8601598186874 | 0.000152448765278979 |
| **2579** | 24,6152223266009 | NA | + | NA | NA | + | NA | NA | NA | NA | + | NA | + | 0,0973471935256524 | 6 | -523,090193007066 | 1058,82653986028 | 10,8649669449942 | 0.000152082785048465 |
| **1968** | 24,6791830070692 | + | + | + | + | NA | + | NA | + | + | + | + | NA | 0,169839707663145 | 11 | -517,358315428488 | 1058,82863085698 | 10,8670579416855 | 0.000151923865837655 |
| **749** | 24,1506121372364 | NA | NA | + | + | NA | + | + | + | NA | + | NA | NA | 0,126302475673708 | 8 | -520,85793120645 | 1058,8408624129 | 10,8792894976091 | 0.000150997568608356 |
| **794** | 25,6482680575701 | + | NA | NA | + | + | NA | NA | NA | + | + | NA | NA | 0,111756345519836 | 7 | -521,988434143791 | 1058,84508534185 | 10,8835124265552 | 0.000150679078964834 |
| **2801** | 24,32712662689 | NA | NA | NA | NA | + | + | + | + | NA | + | NA | + | 0,126268481646952 | 8 | -520,86059503981 | 1058,84619007962 | 10,8846171643286 | 0.000150595871512456 |
| **857** | 25,6687216814772 | NA | NA | NA | + | + | NA | + | NA | + | + | NA | NA | 0,111733372951961 | 7 | -521,990204861593 | 1058,84862677745 | 10,8870538621588 | 0.000150412504920702 |
| **1714** | 24,4268334731566 | + | NA | NA | NA | + | + | NA | + | NA | + | + | NA | 0,126198848801757 | 8 | -520,866051271973 | 1058,85710254395 | 10,8955296286547 | 0.000149776423058139 |
| **1721** | 24,1977822401373 | NA | NA | NA | + | + | + | NA | + | NA | + | + | NA | 0,126194622775307 | 8 | -520,866382397426 | 1058,85776479485 | 10,8961918795605 | 0.000149726836482357 |
| **516** | 24,5033402706385 | + | + | NA | NA | NA | NA | NA | NA | NA | + | NA | NA | 0,0825578452553764 | 5 | -524,202890664027 | 1058,86379659523 | 10,9022236799394 | 0.000149275955535567 |
| **642** | 25,0739105960532 | + | NA | NA | NA | NA | NA | NA | + | NA | + | NA | NA | 0,0825254056276052 | 5 | -524,20531154709 | 1058,86863836136 | 10,9070654460659 | 0.000148915012979206 |

| | | | | | | | | | | | | | | | | | | | |
|---|---|---|---|---|---|---|---|---|---|---|---|---|---|---|---|---|---|---|---|
| **1817** | 25,6829223803975 | NA | NA | NA | + | + | NA | NA | NA | + | + | + | NA | 0,111597781522078 | 7 | -522,0006552711 | 1058,86952759646 | 10,9079546811734 | 0.000148848817467353 |
| **1700** | 24,1462901607506 | + | + | NA | NA | NA | + | NA | + | NA | + | + | NA | 0,126116393262513 | 8 | -520,872511692085 | 1058,87002338417 | 10,9084504688801 | 0.000148811923333523 |
| **1541** | 25,3401530280252 | NA | NA | + | NA | NA | NA | NA | NA | NA | + | + | NA | 0,0824464497709314 | 5 | -524,211203454969 | 1058,88042217711 | 10,9188492618223 | 0.000148040199133077 |
| **1980** | 24,5118669235137 | + | + | NA | + | + | + | NA | + | + | + | + | NA | 0,16950897590628 | 11 | -517,385585741708 | 1058,88317148342 | 10,9215985681255 | 0.000147836835016358 |
| **4085** | 25,1310272560774 | NA | NA | + | NA | + | + | + | + | + | + | + | + | 0,169443390584134 | 11 | -517,390992255317 | 1058,89398451063 | 10,9324115953436 | 0.000147039709931422 |
| **2042** | 24,7588600135923 | + | NA | NA | + | + | + | + | + | + | + | + | NA | 0,169413223809278 | 11 | -517,393478904418 | 1058,89895780884 | 10,9373848935463 | 0.000146674527996697 |
| **1582** | 24,1975642242816 | + | NA | + | + | NA | + | NA | NA | NA | + | + | NA | 0,125865067671283 | 8 | -520,892199381318 | 1058,90939876264 | 10,9478258473455 | 0.000145910812226288 |
| **1707** | 23,9483898323612 | NA | + | NA | + | NA | + | NA | + | NA | + | + | NA | 0,125834283587048 | 8 | -520,894610475743 | 1058,91422095149 | 10,9526480361963 | 0.0001455594312569 |
| **1665** | 25,3886927712309 | NA | NA | NA | NA | NA | NA | NA | + | NA | + | + | NA | 0,0821624757480253 | 5 | -524,232390206332 | 1058,92279567984 | 10,9612227645482 | 0.000144936700855752 |
| **3958** | 25,167928074079 | + | NA | + | NA | + | + | + | NA | + | + | + | + | 0,16922993622504 | 11 | -517,408585371147 | 1058,92917074229 | 10,9675978270041 | 0.000144475446118185 |
| **2992** | 24,6513646243161 | + | + | + | + | NA | + | NA | + | + | + | NA | + | 0,169206290372435 | 11 | -517,410534007342 | 1058,93306801468 | 10,9714950993937 | 0.000144194190156389 |
| **4020** | 24,707263214172 | + | + | NA | NA | + | + | NA | + | + | + | + | + | 0,169199810626579 | 11 | -517,411067988447 | 1058,93413597689 | 10,9725630616035 | 0.000144117213737212 |
| **2726** | 24,4769418076186 | + | NA | + | NA | NA | + | NA | + | NA | + | NA | + | 0,125696912819525 | 8 | -520,905368699118 | 1058,93573739824 | 10,9741644829455 | 0.00014400186373345 |
| **1904** | 24,5132429080724 | + | + | + | + | NA | + | + | NA | + | + | + | NA | 0,169132316975054 | 11 | -517,416629739146 | 1058,94525947829 | 10,983686563002 | 0.000143317894590295 |
| **797** | 25,8314078461452 | NA | NA | + | + | + | NA | NA | NA | + | + | NA | NA | 0,111090459371606 | 7 | -522,039741867431 | 1058,94770078913 | 10,9861278738347 | 0.000143143059554383 |
| **1777** | 24,3301360549378 | NA | NA | NA | NA | + | + | + | + | NA | + | + | NA | 0,12560951589926 | 8 | -520,91221232978 | 1058,94942465956 | 10,9878517442696 | 0.000143019732667745 |
| **3054** | 24,8981104649455 | + | NA | + | + | NA | + | + | + | + | + | NA | + | 0,169033382956711 | 11 | -517,4247814876 | 1058,9615629752 | 10,9999900599087 | 0.00014215435206095 |
| **3062** | 25,1789832616236 | + | NA | + | NA | + | + | + | + | + | + | NA | + | 0,169026913100249 | 11 | -517,425314542863 | 1058,96262908573 | 11,0010561704348 | 0.000142078596128257 |
| **518** | 24,9813941634093 | + | NA | + | NA | NA | NA | NA | NA | NA | + | NA | NA | 0,0818532779385449 | 5 | -524,25545140604 | 1058,96891807926 | 11,0073451639653 | 0.00014163253213327 |
| **4015** | 24,6508179055027 | NA | + | + | + | NA | + | NA | + | + | + | + | + | 0,168903379462403 | 11 | -517,435491755968 | 1058,98298351194 | 11,0214105966447 | 0.000140639965022721 |
| **843** | 24,8182232342673 | NA | + | NA | + | NA | NA | + | NA | + | + | NA | NA | 0,110832274073438 | 7 | -522,059625173849 | 1058,98746740196 | 11,02589448667 | 0.000140325011141122 |
| **754** | 24,3093967199902 | + | NA | NA | NA | + | + | + | + | NA | + | NA | NA | 0,125340920881692 | 8 | -520,933240429644 | 1058,99148085929 | 11,0299079439976 | 0.000140043699271717 |
| **4077** | 24,8846225506371 | NA | NA | + | + | NA | + | + | + | + | + | + | + | 0,168847866277489 | 11 | -517,440064669864 | 1058,99212933973 | 11,0305564244372 | 0.000139998298832604 |
| **761** | 24,040633168202 | NA | NA | NA | + | + | + | + | + | NA | + | NA | NA | 0,125282061818518 | 8 | -520,937847598831 | 1059,00069519766 | 11,0391222823723 | 0.00013939997825999 |
| **1859** | 25,0123353368177 | NA | + | NA | NA | NA | NA | + | NA | + | + | + | NA | 0,11071341862066 | 7 | -522,068776502829 | 1059,00577005992 | 11,0441971446305 | 0.000139046708803761 |
| **1538** | 24,9915939764765 | + | NA | NA | NA | NA | NA | NA | NA | NA | + | + | NA | 0,081600559762724 | 5 | -524,274294366182 | 1059,00660399954 | 11,0450310842493 | 0.000138988742610008 |
| **921** | 25,8024595283454 | NA | NA | NA | + | + | NA | NA | + | + | + | NA | NA | 0,110702449673777 | 7 | -522,069621000204 | 1059,00745905467 | 11,0458861393802 | 0.000138929333791618 |
| **3951** | 24,5104565051277 | NA | + | + | + | NA | + | + | NA | + | + | + | + | 0,168722208334095 | 11 | -517,450414650105 | 1059,01282930021 | 11,0512563849193 | 0.000138556791858342 |
| **539** | 24,7441614200063 | NA | + | NA | + | + | NA | NA | NA | NA | + | NA | NA | 0,0960605738475621 | 6 | -523,187714694162 | 1059,02158323448 | 11,060010319188 | 0.00013795165862436 |
| **4082** | 25,0281457611081 | + | NA | NA | NA | + | + | + | + | + | + | + | + | 0,168656090870123 | 11 | -517,455859873346 | 1059,02371974669 | 11,0621468314005 | 0.000137804369607872 |
| **3747** | 24,080761830335 | NA | + | NA | NA | NA | + | NA | + | NA | + | + | + | 0,125108118341691 | 8 | -520,951461142892 | 1059,02792228578 | 11,066349370494 | 0.000137515109495769 |
| **742** | 24,4228597547597 | + | NA | + | NA | NA | + | + | + | NA | + | NA | NA | 0,125087735184966 | 8 | -520,953056236914 | 1059,03111247383 | 11,0695395585369 | 0.000137295934815333 |
| **522** | 24,5784331518452 | + | NA | NA | + | NA | NA | NA | NA | NA | + | NA | NA | 0,081390201069498 | 5 | -524,289975002242 | 1059,03796527166 | 11,0763923563684 | 0.000136826309194963 |
| **2789** | 24,3841630900382 | NA | NA | + | NA | NA | + | + | + | NA | + | NA | + | 0,124980459710013 | 8 | -520,961450520222 | 1059,04790104044 | 11,0863281251527 | 0.000136148257544701 |
| **836** | 24,8605970598402 | + | + | NA | NA | NA | NA | + | NA | + | + | NA | NA | 0,110411738367258 | 7 | -522,091999021114 | 1059,05221509649 | 11,0906421812001 | 0.000135854898443444 |
| **1763** | 24,0524900047562 | NA | + | NA | NA | NA | + | + | + | NA | + | + | NA | 0,124911397093427 | 8 | -520,96685411117 | 1059,05870822234 | 11,0971353070502 | 0.000135414552159711 |
| **2839** | 25,2976782648908 | NA | + | + | NA | + | NA | NA | NA | + | + | NA | + | 0,124904490063828 | 8 | -520,967394506904 | 1059,05978901381 | 11,0982160985175 | 0.000135341394482278 |
| **1638** | 24,3777170080686 | + | NA | + | NA | NA | + | + | NA | NA | + | + | NA | 0,124903300785307 | 8 | -520,967487553858 | 1059,05997510772 | 11,098402192426 | 0.000135328801963595 |
| **2928** | 24,477410759193 | + | + | + | + | NA | + | + | NA | + | + | NA | + | 0,16843483025321 | 11 | -517,47407904397 | 1059,06015808794 | 11,09858517265 | 0.000135316421282706 |
| **850** | 25,6953726962354 | + | NA | NA | NA | + | NA | + | NA | + | + | NA | NA | 0,110350809795608 | 7 | -522,096688179988 | 1059,06159341424 | 11,1000204989496 | 0.00013521934451197 |
| **1645** | 24,066146627982 | NA | NA | + | + | NA | + | + | NA | NA | + | + | NA | 0,124809526418327 | 8 | -520,974823889225 | 1059,07464777845 | 11,1130748631585 | 0.000134339617419089 |
| **560** | 23,7027358572381 | + | + | + | + | NA | + | NA | NA | NA | + | NA | NA | 0,124688069878979 | 8 | -520,984324741445 | 1059,09364948289 | 11,1320765675987 | 0.000133069320573383 |
| **776** | 25,2997688672987 | + | + | + | NA | NA | NA | NA | NA | + | + | NA | NA | 0,110024580687749 | 7 | -522,121789827033 | 1059,11179670833 | 11,1502237930388 | 0.000131867362391573 |
| **2030** | 24,8591507646927 | + | NA | + | + | NA | + | + | + | + | + | + | NA | 0,168063574727395 | 11 | -517,504638300142 | 1059,12127660028 | 11,159703684993 | 0.000131243797217784 |
| **903** | 25,7251809461835 | NA | + | + | NA | NA | NA | NA | + | + | + | NA | NA | 0,109962954660336 | 7 | -522,126530599194 | 1059,12127825265 | 11,1597053373614 | 0.000131243688786278 |
| **900** | 25,3799009968074 | + | + | NA | NA | NA | NA | NA | + | + | + | NA | NA | 0,109876692438762 | 7 | -522,133166035531 | 1059,13454912533 | 11,1729762100349 | 0.000130375712530008 |
| **3622** | 24,3518760732027 | + | NA | + | NA | NA | + | NA | NA | NA | + | + | + | 0,124412123856787 | 8 | -521,005905527886 | 1059,13681105577 | 11,1752381404813 | 0.000130228345482399 |
| **1803** | 25,1027483830874 | NA | + | NA | + | NA | NA | NA | NA | + | + | + | NA | 0,10977145629841 | 7 | -522,14126010759 | 1059,15073726944 | 11,1891643541524 | 0.00012932470133624 |
| **3859** | 25,2494303373613 | NA | + | NA | NA | + | NA | NA | NA | + | + | + | + | 0,12425304036915 | 8 | -521,018343809697 | 1059,16168761939 | 11,2001147041035 | 0.00012861856085314 |
| **1873** | 25,7655287330227 | NA | NA | NA | NA | + | NA | + | NA | + | + | + | NA | 0,109697372479967 | 7 | -522,146957574272 | 1059,16213220281 | 11,2005592875173 | 0.000128589973191227 |
| **3685** | 24,277536561528 | NA | NA | + | NA | NA | + | + | NA | NA | + | + | + | 0,124180571556902 | 8 | -521,024009189674 | 1059,17301837935 | 11,2114454640582 | 0.000127891948046069 |
| **1706** | 24,2064003918973 | + | NA | NA | + | NA | + | NA | + | NA | + | + | NA | 0,124174585693867 | 8 | -521,024477124354 | 1059,17395424871 | 11,2123813334169 | 0.000127832116967977 |
| **4071** | 24,8675812381428 | NA | + | + | NA | NA | + | + | + | + | + | + | + | 0,167714630726725 | 11 | -517,533348598293 | 1059,17869719659 | 11,2171242812949 | 0.000127529325606839 |
| **780** | 24,948825989626 | + | + | NA | + | NA | NA | NA | NA | + | + | NA | NA | 0,109557573653166 | 7 | -522,157707607901 | 1059,18363227006 | 11,2220593547743 | 0.000127215030237674 |
| **2836** | 25,1926183072279 | + | + | NA | NA | + | NA | NA | NA | + | + | NA | + | 0,124112011714586 | 8 | -521,029368547586 | 1059,18373709517 | 11,2221641798806 | 0.000127208362747872 |
| **2963** | 25,1991272875183 | NA | + | NA | NA | + | NA | NA | + | + | + | NA | + | 0,124056247554291 | 8 | -521,033727351289 | 1059,19245470258 | 11,2308817872884 | 0.000126655093137944 |
| **1810** | 25,7613911372477 | + | NA | NA | NA | + | NA | NA | NA | + | + | + | NA | 0,109468231898499 | 7 | -522,164576788276 | 1059,19737063082 | 11,2357977155259 | 0.000126344161749193 |
| **1799** | 25,5775928375557 | NA | + | + | NA | NA | NA | NA | NA | + | + | + | NA | 0,109372344595307 | 7 | -522,171948467065 | 1059,21211398839 | 11,2505410731037 | 0.00012541621762709 |
| **1555** | 25,1881422102835 | NA | + | NA | NA | + | NA | NA | NA | NA | + | + | NA | 0,0946937678823378 | 6 | -523,291162384316 | 1059,22847861478 | 11,2669056994944 | 0.000124394209750924 |
| **2600** | 23,8841132920861 | + | + | + | NA | NA | + | NA | NA | NA | + | NA | + | 0,123781617192821 | 8 | -521,055189778587 | 1059,23537955717 | 11,2738066418831 | 0.000123965730765745 |
| **1580** | 23,7694432454207 | + | + | NA | + | NA | + | NA | NA | NA | + | + | NA | 0,123693249576851 | 8 | -521,062094298786 | 1059,24918859757 | 11,2876156822808 | 0.00012311275495667 |
| **535** | 25,1472478360173 | NA | + | + | NA | + | NA | NA | NA | NA | + | NA | NA | 0,0945373851126342 | 6 | -523,302988372818 | 1059,25213059179 | 11,2905576764992 | 0.000122931789582425 |
| **3587** | 24,373490390011 | NA | + | NA | NA | NA | NA | NA | NA | NA | + | + | + | 0,0945294171132513 | 6 | -523,303590874752 | 1059,25333559566 | 11,2917626803674 | 0.000122857745249612 |
| **2641** | 24,7303357219392 | NA | NA | NA | NA | + | NA | + | NA | NA | + | NA | + | 0,094522450923164 | 6 | -523,304117620326 | 1059,25438908681 | 11,2928161715145 | 0.000122793047517243 |
| **2024** | 24,8372465850435 | + | + | + | NA | NA | + | + | + | + | + | + | NA | 0,167252434404988 | 11 | -517,571358517273 | 1059,25471703455 | 11,2931441192545 | 0.000122772914316733 |
| **859** | 25,1715014721824 | NA | + | NA | + | + | NA | + | NA | + | + | NA | NA | 0,12361483019234 | 8 | -521,068220940536 | 1059,26144188107 | 11,2998689657816 | 0.000122360793064829 |
| **3048** | 24,8380574242614 | + | + | + | NA | NA | + | + | + | + | + | NA | + | 0,167200541118539 | 11 | -517,57562478026 | 1059,26324956052 | 11,3016766452297 | 0.000122250248484207 |
| **3629** | 24,0736467203024 | NA | NA | + | + | NA | + | NA | NA | NA | + | + | + | 0,123593106115926 | 8 | -521,069918072359 | 1059,26483614472 | 11,3032632294278 | 0.000122153306784604 |
| **1769** | 24,1165007694754 | NA | NA | NA | + | NA | + | + | + | NA | + | + | NA | 0,123480059249786 | 8 | -521,078748859338 | 1059,28249771868 | 11,320924803385 | 0.000121079345892785 |
| **1636** | 23,9168572700385 | + | + | NA | NA | NA | + | + | NA | NA | + | + | NA | 0,123352872602791 | 8 | -521,088682830945 | 1059,30236566189 | 11,3407927466001 | 0.000119882501659154 |
| **616** | 23,8516139737651 | + | + | + | NA | NA | + | + | NA | NA | + | NA | NA | 0,12335121402677 | 8 | -521,088812365274 | 1059,30262473055 | 11,3410518152566 | 0.00011986697376555 |
| **3753** | 24,1556504722317 | NA | NA | NA | + | NA | + | NA | + | NA | + | + | + | 0,123226238466159 | 8 | -521,098572217418 | 1059,32214443484 | 11,3605715195458 | 0.000118702780244124 |
| **853** | 25,9347633007059 | NA | NA | + | NA | + | NA | + | NA | + | + | NA | NA | 0,108635645977083 | 7 | -522,22855835801 | 1059,32533377028 | 11,3637608549925 | 0.000118513639600119 |
| **855** | 25,3775847703253 | NA | + | + | NA | + | NA | + | NA | + | + | NA | NA | 0,123197230639578 | 8 | -521,100837358287 | 1059,32667471657 | 11,3651018012843 | 0.000118434206019365 |
| **790** | 25,8803509810674 | + | NA | + | NA | + | NA | NA | NA | + | + | NA | NA | 0,10856921564998 | 7 | -522,233660743139 | 1059,33553854054 | 11,3739656252519 | 0.000117910477459103 |
| **1796** | 25,2072407582209 | + | + | NA | NA | NA | NA | NA | NA | + | + | + | NA | 0,108553515155257 | 7 | -522,234866612291 | 1059,33795027885 | 11,3763773635553 | 0.00011776837854524 |

| | | | | | | | | | | | | | | | | | | |
|---|---|---|---|---|---|---|---|---|---|---|---|---|---|---|---|---|---|---|
| 2038 | 25,0151708761153 | + | NA | + | NA | + | + | + | + | + | + | + | NA | 0,166729362525122 | 11 | -517,614349270642 | 1059,34069854129 | 11,3791256259944 | 0.000117606660475974 |
| 3950 | 24,7210907846836 | + | NA | + | + | NA | + | + | NA | + | + | + | + | 0,166729058304481 | 11 | -517,614374266383 | 1059,34074853277 | 11,3791756174753 | 0.000117603720847155 |
| 914 | 25,8532966973317 | + | NA | NA | NA | + | NA | NA | + | + | + | NA | NA | 0,108534834126466 | 7 | -522,236301372313 | 1059,34081979889 | 11,3792468835984 | 0.000117599530341193 |
| 2585 | 24,6741567651686 | NA | NA | NA | + | + | NA | NA | NA | NA | + | NA | + | 0,0939453321933072 | 6 | -523,347742151696 | 1059,34163814955 | 11,3800652342557 | 0.000117551421357895 |
| 3620 | 23,8759686509443 | + | + | NA | NA | NA | + | NA | NA | NA | + | + | + | 0,123094276109222 | 8 | -521,108876187893 | 1059,34275237579 | 11,3811794604956 | 0.000117485950157913 |
| 3006 | 24,995908842235 | + | NA | + | + | + | + | NA | + | + | + | NA | + | 0,166679846130612 | 11 | -517,618417575838 | 1059,34883515168 | 11,3872622363858 | 0.000117129172629834 |
| 977 | 25,9116842544058 | NA | NA | NA | NA | + | NA | + | + | + | + | NA | NA | 0,108458170898157 | 7 | -522,242189027511 | 1059,35259510929 | 11,391022193995 | 0.000116909179124248 |
| 4029 | 24,9015741765206 | NA | NA | + | + | + | + | NA | + | + | + | + | + | 0,166625450758299 | 11 | -517,622886463128 | 1059,35777292626 | 11,3962000109645 | 0.00011660690340938 |
| 622 | 23,9085096748413 | + | NA | + | + | NA | + | + | NA | NA | + | NA | NA | 0,122923551114992 | 8 | -521,12220454846 | 1059,36940909692 | 11,4078361816296 | 0.000115930444251718 |
| 4008 | 24,8291573434036 | + | + | + | NA | NA | + | NA | + | + | + | + | + | 0,166473097060707 | 11 | -517,635401628888 | 1059,38280325778 | 11,4212303424854 | 0.00011515664274014 |
| 909 | 25,8126827429468 | NA | NA | + | + | NA | NA | NA | + | + | + | NA | NA | 0,108232244083969 | 7 | -522,259537026343 | 1059,38729110695 | 11,4257181916591 | 0.000114898529619646 |
| 969 | 25,5403654420667 | NA | NA | NA | + | NA | NA | + | + | + | + | NA | NA | 0,108216579621153 | 7 | -522,260739673716 | 1059,3896964017 | 11,4281234864056 | 0.000114760430263868 |
| 3902 | 24,8706360443643 | + | NA | + | + | + | + | NA | NA | + | + | + | + | 0,166350451516025 | 11 | -517,645474743563 | 1059,40294948713 | 11,4413765718361 | 0.000114002479440989 |
| 906 | 25,5054899695695 | + | NA | NA | + | NA | NA | NA | + | + | + | NA | NA | 0,108037615612738 | 7 | -522,274478231179 | 1059,41717351662 | 11,4556006013302 | 0.000113194568468776 |
| 1762 | 24,3269984159016 | + | NA | NA | NA | NA | + | + | + | NA | + | + | NA | 0,122595269390835 | 8 | -521,147825948546 | 1059,42065189709 | 11,4590789818008 | 0.000112997872675888 |
| 3746 | 24,3395519671946 | + | NA | NA | NA | NA | + | NA | + | NA | + | + | + | 0,122577843014927 | 8 | -521,14918575689 | 1059,42337151378 | 11,4617985984885 | 0.000112844321649428 |
| 659 | 24,9886726110798 | NA | + | NA | NA | + | NA | NA | + | NA | + | NA | NA | 0,0934010356502153 | 6 | -523,388860205038 | 1059,42387425623 | 11,4623013409384 | 0.00011281595939894 |
| 3944 | 24,7166894657305 | + | + | + | NA | NA | + | + | NA | + | + | + | + | 0,166186324660634 | 11 | -517,658952481052 | 1059,4299049621 | 11,4683320468134 | 0.000112476291830535 |
| 532 | 25,0052812096626 | + | + | NA | NA | + | NA | NA | NA | NA | + | NA | NA | 0,0933416903350118 | 6 | -523,393341864609 | 1059,43283757537 | 11,4712646600813 | 0.000112311487953703 |
| 1592 | 24,4122831925458 | + | + | + | NA | + | + | NA | NA | NA | + | + | NA | 0,137083010288502 | 9 | -520,007880000547 | 1059,43308283574 | 11,4715099204486 | 0.00011229771601975 |
| 1923 | 25,5903619643323 | NA | + | NA | NA | NA | NA | NA | + | + | + | + | NA | 0,107908036973025 | 7 | -522,28442389495 | 1059,43706484416 | 11,4754919288739 | 0.000112074353226002 |
| 3809 | 24,2890890800942 | NA | NA | NA | NA | NA | + | + | + | NA | + | + | + | 0,122417805866331 | 8 | -521,161672452761 | 1059,44834490552 | 11,4867719902318 | 0.000111444029628579 |
| 684 | 23,8310120469923 | + | + | NA | + | NA | + | NA | + | NA | + | NA | NA | 0,122376902642554 | 8 | -521,164863509813 | 1059,45472701963 | 11,4931541043363 | 0.000111088972177481 |
| 707 | 24,5484556383824 | NA | + | NA | NA | NA | NA | + | + | NA | + | NA | NA | 0,0931869979728326 | 6 | -523,405022596588 | 1059,45619903933 | 11,4946261240389 | 0.000111007239681242 |
| 2606 | 23,9458864692631 | + | NA | + | + | NA | + | NA | NA | NA | + | NA | + | 0,122352536641705 | 8 | -521,166764347984 | 1059,45852869597 | 11,4969557806767 | 0.000110878010584325 |
| 2753 | 24,6288484341296 | NA | NA | NA | NA | NA | NA | + | + | NA | + | NA | + | 0,0931196869333799 | 6 | -523,410104592916 | 1059,46636303198 | 11,5047901166943 | 0.00011044453234621 |
| 3011 | 24,8372445689045 | NA | + | NA | NA | NA | NA | + | + | + | + | NA | + | 0,122052718903067 | 8 | -521,190149384605 | 1059,50529876921 | 11,5437258539203 | 0.000108315206668145 |
| 799 | 25,4767201872051 | NA | + | + | + | + | NA | NA | NA | + | + | NA | NA | 0,122043557939791 | 8 | -521,190863791239 | 1059,50672758248 | 11,545154667187 | 0.000108237853200209 |
| 962 | 25,5494433473435 | + | NA | NA | NA | NA | NA | + | + | + | + | NA | NA | 0,107425609563126 | 7 | -522,321439381161 | 1059,51109581659 | 11,5495229012945 | 0.000108001707038481 |
| 852 | 25,2649289094986 | + | + | NA | NA | + | NA | + | NA | + | + | NA | NA | 0,122009583530258 | 8 | -521,193513179394 | 1059,51202635879 | 11,5504534434983 | 0.000107951468653379 |
| 1982 | 24,8701403228843 | + | NA | + | + | + | + | NA | + | + | + | + | NA | 0,165594251141683 | 11 | -517,707550234353 | 1059,52710046871 | 11,5655275534157 | 0.000107140891022561 |
| 2955 | 24,847158549176 | NA | + | NA | + | NA | NA | NA | + | + | + | NA | + | 0,121831699750731 | 8 | -521,207383217476 | 1059,53976643495 | 11,5781935196605 | 0.000106464513572084 |
| 979 | 25,3189147348378 | NA | + | NA | NA | + | NA | + | + | + | + | NA | NA | 0,121810955568007 | 8 | -521,209000509853 | 1059,54300101971 | 11,5814281044156 | 0.000106292468486908 |
| 1875 | 25,3166846477674 | NA | + | NA | NA | + | NA | + | NA | + | + | + | NA | 0,121810270025672 | 8 | -521,20905395659 | 1059,54310791318 | 11,5815349978891 | 0.000106286787653141 |
| 1643 | 23,6991206974963 | NA | + | NA | + | NA | + | + | NA | NA | + | + | NA | 0,121787109989028 | 8 | -521,210859551256 | 1059,54671910251 | 11,5851461872212 | 0.000106095049948662 |
| 3683 | 23,8166718951428 | NA | + | NA | NA | NA | + | + | NA | NA | + | + | + | 0,121762029381603 | 8 | -521,212814823113 | 1059,55062964623 | 11,5890567309345 | 0.000105887807956608 |
| 4075 | 24,5227566181463 | NA | + | NA | + | NA | + | + | + | + | + | + | + | 0,165423286296126 | 11 | -517,721576719641 | 1059,55515343928 | 11,5935805239919 | 0.000105648571357575 |
| 2662 | 24,1139741570155 | + | NA | + | NA | NA | + | + | NA | NA | + | NA | + | 0,12170669864095 | 8 | -521,217128183154 | 1059,55925636631 | 11,5976834510166 | 0.000105432059327504 |
| 3601 | 25,0054135730259 | NA | NA | NA | NA | + | NA | NA | NA | NA | + | + | + | 0,0923704651593722 | 6 | -523,466645540299 | 1059,57944492675 | 11,6178720114608 | 0.000104373152021183 |
| 1642 | 23,9796845248631 | + | NA | NA | + | NA | + | + | NA | NA | + | + | NA | 0,121568958306947 | 8 | -521,227864681211 | 1059,58072936242 | 11,6191564471319 | 0.000104306143240823 |
| 2730 | 24,0162354159177 | + | NA | NA | + | NA | + | NA | + | NA | + | NA | + | 0,121500587421728 | 8 | -521,233193387634 | 1059,59138677527 | 11,6298138599777 | 0.00010375180469079 |
| 601 | 24,942472923924 | NA | NA | NA | + | + | NA | + | NA | NA | + | NA | NA | 0,092283799374173 | 6 | -523,473182870455 | 1059,59251958706 | 11,6309466717728 | 0.000103693055696193 |
| 2581 | 24,9924742172416 | NA | NA | + | NA | + | NA | NA | NA | NA | + | NA | + | 0,0922442510369587 | 6 | -523,476165853546 | 1059,59848555325 | 11,6369126379559 | 0.000103384201946116 |
| 2831 | 24,6970858194056 | NA | + | + | + | NA | NA | NA | NA | + | + | NA | + | 0,121429230326039 | 8 | -521,238754391999 | 1059,602508784 | 11,6409358687072 | 0.000103176441732764 |
| 2028 | 24,4896459521489 | + | + | NA | + | NA | + | + | + | + | + | + | NA | 0,165109424877119 | 11 | -517,747319408334 | 1059,60663881667 | 11,6450659013778 | 0.000102963600531077 |
| 746 | 23,9655753379141 | + | NA | NA | + | NA | + | + | + | NA | + | NA | NA | 0,121363556540526 | 8 | -521,243872085459 | 1059,61274417092 | 11,6511712556269 | 0.000102649765165814 |
| 3052 | 24,4955724738738 | + | + | NA | + | NA | + | + | + | + | + | NA | + | 0,165062645144918 | 11 | -517,751155420176 | 1059,61431084035 | 11,652737925062 | 0.00010256938752642 |
| 1937 | 25,9892972779665 | NA | NA | NA | NA | + | NA | NA | + | + | + | + | NA | 0,106748023174684 | 7 | -522,373395189912 | 1059,61500743409 | 11,6534345187979 | 0.000102533669150655 |
| 2663 | 23,7500326923621 | NA | + | + | NA | NA | + | + | NA | NA | + | NA | + | 0,121333237450162 | 8 | -521,246234600998 | 1059,617469202 | 11,6558962867045 | 0.000102407539743878 |
| 3888 | 24,3994764375147 | + | + | + | + | NA | + | NA | NA | + | + | + | + | 0,165039269897444 | 11 | -517,753072146861 | 1059,61814429372 | 11,6565713784305 | 0.000102372978335858 |
| 2887 | 24,6921792091081 | NA | + | + | NA | NA | NA | + | NA | + | + | NA | + | 0,121239361716557 | 8 | -521,253549042545 | 1059,63209808509 | 11,6705251698002 | 0.000101661218569083 |
| 2724 | 23,9780353174403 | + | + | NA | NA | NA | + | NA | + | NA | + | NA | + | 0,121209979199688 | 8 | -521,255838256477 | 1059,63667651295 | 11,6751035976631 | 0.000101428760465924 |
| 576 | 24,0618032819436 | + | + | + | + | + | + | NA | NA | NA | + | NA | NA | 0,135789999682218 | 9 | -520,11039309496 | 1059,63810902456 | 11,6765361092741 | 0.000101356137538795 |
| 2607 | 23,6031507872458 | NA | + | + | + | NA | + | NA | NA | NA | + | NA | + | 0,121159033039865 | 8 | -521,259807328692 | 1059,64461465738 | 11,6830417420936 | 0.000101026980265703 |
| 1813 | 26,0877543214965 | NA | NA | + | NA | + | NA | NA | NA | + | + | + | NA | 0,10654806108773 | 7 | -522,388720307794 | 1059,64565766985 | 11,6840847545614 | 0.000100974307801406 |
| 3627 | 23,6737124748717 | NA | + | NA | + | NA | + | NA | NA | NA | + | + | + | 0,12114660122312 | 8 | -521,260775821711 | 1059,64655164342 | 11,684978728131 | 0.000100929183705901 |
| 845 | 25,4069740824458 | NA | NA | + | + | NA | NA | + | NA | + | + | NA | NA | 0,10652964855077 | 7 | -522,390131274421 | 1059,64847960311 | 11,6869066878153 | 0.000100831936886842 |
| 782 | 25,3826082087259 | + | NA | + | + | NA | NA | NA | NA | + | + | NA | NA | 0,106514883959752 | 7 | -522,391262675197 | 1059,65074240466 | 11,6891694893663 | 0.000100717920066771 |
| 2731 | 23,8127152813261 | NA | + | NA | + | NA | + | NA | + | NA | + | NA | + | 0,121095013686203 | 8 | -521,26479457067 | 1059,65458914134 | 11,6930162260489 | 0.000100524388583739 |
| 1929 | 25,6016324411218 | NA | NA | NA | + | NA | NA | NA | + | + | + | + | NA | 0,106436849246495 | 7 | -522,397242112931 | 1059,66270128013 | 11,7011283648353 | 0.00010011748046957 |
| 2951 | 25,2236136243898 | NA | + | + | NA | NA | NA | NA | + | + | + | NA | + | 0,121021194868523 | 8 | -521,270544760764 | 1059,66608952153 | 11,7045166062383 | 9.99480129630644e-05 |
| 2705 | 24,958733529753 | NA | NA | NA | NA | + | NA | NA | + | NA | + | NA | + | 0,0917754766831882 | 6 | -523,511513857233 | 1059,66918156062 | 11,7076086453285 | 9.97936107666887e-05 |
| 2669 | 23,8048340601049 | NA | NA | + | + | NA | + | + | NA | NA | + | NA | + | 0,12095701977413 | 8 | -521,275543351263 | 1059,67608670253 | 11,7145137872355 | 9.94496603430934e-05 |
| 2567 | 24,298045732135 | NA | + | + | NA | NA | NA | NA | NA | NA | + | NA | + | 0,0917079743789244 | 6 | -523,516602376996 | 1059,67935860015 | 11,7177856848555 | 9.92870987970435e-05 |
| 740 | 23,9321908454312 | + | + | NA | NA | NA | + | + | + | NA | + | NA | NA | 0,120886590197189 | 8 | -521,281028682885 | 1059,68705736577 | 11,7254844504798 | 9.89056394081015e-05 |
| 2691 | 24,4731918703555 | NA | + | NA | NA | NA | NA | NA | + | NA | + | NA | + | 0,0916506261142928 | 6 | -523,520925158792 | 1059,68800416374 | 11,7264312484481 | 9.88588286598359e-05 |
| 2578 | 24,9119490331936 | + | NA | NA | NA | + | NA | NA | NA | NA | + | NA | + | 0,0916505748363103 | 6 | -523,520929023888 | 1059,68801189393 | 11,7264389786394 | 9.88584465617443e-05 |
| 796 | 25,3630602422812 | + | + | NA | + | + | NA | NA | NA | + | + | NA | NA | 0,120836862219906 | 8 | -521,284901428206 | 1059,69480285641 | 11,733229941121 | 9.85233438004798e-05 |
| 4026 | 24,7969315890428 | + | NA | NA | + | + | + | NA | + | + | + | + | + | 0,164570335730394 | 11 | -517,791512547868 | 1059,69502509574 | 11,7334521804462 | 9.85123965279964e-05 |
| 965 | 25,8973288383493 | NA | NA | + | NA | NA | NA | + | + | + | + | NA | NA | 0,106191235210249 | 7 | -522,416058969942 | 1059,70033499415 | 11,7387620788561 | 9.82511980069863e-05 |
| 2697 | 24,538581643754 | NA | NA | NA | + | NA | NA | NA | + | NA | + | NA | + | 0,0915427815301024 | 6 | -523,529053499571 | 1059,7042608453 | 11,7426879300044 | 9.80585273786428e-05 |
| 583 | 24,3146102638472 | NA | + | + | NA | NA | NA | + | NA | NA | + | NA | NA | 0,0915194758694331 | 6 | -523,530809940896 | 1059,70777372795 | 11,7462008126558 | 9.78864444997974e-05 |
| 3847 | 25,0564329746812 | NA | + | + | NA | NA | NA | NA | NA | + | + | + | + | 0,120738301086055 | 8 | -521,292576584482 | 1059,71015316896 | 11,7485802536739 | 9.77700562376979e-05 |
| 597 | 25,206622250705 | NA | NA | + | NA | + | NA | + | NA | NA | + | NA | NA | 0,0915028626394019 | 6 | -523,532061976761 | 1059,71027779968 | 11,7487048843846 | 9.77639638517316e-05 |
| 3639 | 24,2895692997213 | NA | + | + | NA | + | + | NA | NA | NA | + | + | + | 0,135300641079498 | 9 | -520,14915067792 | 1059,71562419049 | 11,7540512751953 | 9.75029706710609e-05 |
| 923 | 25,391060487005 | NA | + | NA | + | + | NA | NA | + | + | + | NA | NA | 0,120669027554667 | 8 | -521,297970540912 | 1059,72094108182 | 11,7593681665342 | 9.72441085588076e-05 |

| | | | | | | | | | | | | | | | | | | | |
|---|---|---|---|---|---|---|---|---|---|---|---|---|---|---|---|---|---|---|---|
| **1603** | 24,4182424127692 | NA | + | NA | NA | NA | NA | + | NA | NA | + | + | NA | 0,0914016977983868 | 6 | -523,539685647966 | 1059,72552514209 | 11,7639522267953 | 9.70214773681353e-05 |
| **1819** | 25,413655684516 | NA | + | NA | + | + | NA | NA | NA | + | + | + | NA | 0,120638897879533 | 8 | -521,300316443748 | 1059,7256328875 | 11,7640599722049 | 9.7016250699512e-05 |
| **2793** | 23,9310930249145 | NA | NA | NA | + | NA | + | + | + | NA | + | NA | + | 0,120625179438048 | 8 | -521,301384537867 | 1059,72776907573 | 11,7661961604431 | 9.69126835322707e-05 |
| **4022** | 25,0485909637923 | + | NA | + | NA | + | + | NA | + | + | + | + | + | 0,16436142264822 | 11 | -517,808631041672 | 1059,72926208334 | 11,7676891680542 | 9.68403648416556e-05 |
| **623** | 23,550137400627 | NA | + | + | + | NA | + | + | NA | NA | + | NA | NA | 0,120598413880999 | 8 | -521,303468409886 | 1059,73193681977 | 11,7703639044812 | 9.67109401794363e-05 |
| **721** | 25,2046007373137 | NA | NA | NA | NA | + | NA | + | + | NA | + | NA | NA | 0,0912761027465025 | 6 | -523,5491491726 | 1059,74445219135 | 11,7828792760636 | 9.61076430937135e-05 |
| **3682** | 24,1297136054682 | + | NA | NA | NA | NA | + | + | NA | NA | + | + | + | 0,120502567576996 | 8 | -521,310930145675 | 1059,74686029135 | 11,7852873760598 | 9.59919943236479e-05 |
| **747** | 23,750714035558 | NA | + | NA | + | NA | + | + | + | NA | + | NA | NA | 0,120485036122779 | 8 | -521,312294899935 | 1059,74958979987 | 11,7880168845797 | 9.58610781949654e-05 |
| **4070** | 25,0384665367109 | + | NA | + | NA | NA | + | + | + | + | + | + | + | 0,164163284301924 | 11 | -517,824862693805 | 1059,76172538761 | 11,8001524723188 | 9.52811740809284e-05 |
| **4074** | 24,6652308577424 | + | NA | NA | + | NA | + | + | + | + | + | + | + | 0,164163154777015 | 11 | -517,824873303331 | 1059,76174660666 | 11,8001736913718 | 9.52801631981497e-05 |
| **842** | 25,1654068012577 | + | NA | NA | + | NA | NA | + | NA | + | + | NA | NA | 0,105711691347697 | 7 | -522,452782647923 | 1059,77378235011 | 11,8122094348198 | 9.47085012176325e-05 |
| **1599** | 24,0348813346168 | NA | + | + | + | + | + | NA | NA | NA | + | + | NA | 0,134905542692739 | 9 | -520,180426786142 | 1059,77817640693 | 11,8166034916383 | 9.45006523583923e-05 |
| **2787** | 23,9109935905927 | NA | + | NA | NA | NA | + | + | + | NA | + | NA | + | 0,120299778996689 | 8 | -521,32671477178 | 1059,77842954356 | 11,8168566282689 | 9.44886923269255e-05 |
| **3689** | 23,905249275182 | NA | NA | NA | + | NA | + | + | NA | NA | + | + | + | 0,120283336333127 | 8 | -521,327994473876 | 1059,78098894775 | 11,8194160324617 | 9.43678522853832e-05 |
| **2944** | 24,8405846282529 | + | + | + | + | + | + | + | NA | + | + | NA | + | 0,17859054417241 | 12 | -516,632797523986 | 1059,78172408023 | 11,8201511649402 | 9.43331722228159e-05 |
| **1805** | 25,5010807834193 | NA | NA | + | + | NA | NA | NA | NA | + | + | + | NA | 0,105593661642785 | 7 | -522,461818395727 | 1059,79185384572 | 11,8302809304275 | 9.38565936926613e-05 |
| **902** | 25,802802818675 | + | NA | + | NA | NA | NA | NA | + | + | + | NA | NA | 0,105565196465284 | 7 | -522,463997365096 | 1059,79621178446 | 11,8346388691648 | 9.36523056994294e-05 |
| **838** | 25,4199979000979 | + | NA | + | NA | NA | NA | + | NA | + | + | NA | NA | 0,105564701782914 | 7 | -522,464035231724 | 1059,79628751771 | 11,8347146024209 | 9.36487594695432e-05 |
| **594** | 25,0667318616105 | + | NA | NA | NA | + | NA | + | NA | NA | + | NA | NA | 0,0909221039316336 | 6 | -523,575815773383 | 1059,79778539292 | 11,8362124776297 | 9.35786486495644e-05 |
| **917** | 26,1376579118821 | NA | NA | + | NA | + | NA | NA | + | + | + | NA | NA | 0,105546792745977 | 7 | -522,465406107075 | 1059,79902926841 | 11,8374563531236 | 9.35204666503289e-05 |
| **2627** | 23,8056286321998 | NA | + | NA | NA | NA | NA | + | NA | NA | + | NA | + | 0,0908776820852689 | 6 | -523,579161321806 | 1059,80447648977 | 11,8429035744755 | 9.32660998642289e-05 |
| **2616** | 24,339332945679 | + | + | + | NA | + | + | NA | NA | NA | + | NA | + | 0,134736368615908 | 9 | -520,193814292312 | 1059,80495141927 | 11,8433785039797 | 9.32439550823516e-05 |
| **1617** | 25,1514403915401 | NA | NA | NA | NA | + | NA | + | NA | NA | + | + | NA | 0,0908552220716585 | 6 | -523,580852793608 | 1059,80785943337 | 11,8462865180782 | 9.31084762318337e-05 |
| **2786** | 24,1319435101679 | + | NA | NA | NA | NA | + | + | + | NA | + | NA | + | 0,120050234758693 | 8 | -521,346133766943 | 1059,81726753389 | 11,8556946185954 | 9.26715178230474e-05 |
| **2693** | 24,9797190312695 | NA | NA | + | NA | NA | NA | NA | + | NA | + | NA | + | 0,0907902953536236 | 6 | -523,585742213505 | 1059,81763827316 | 11,8560653578738 | 9.26543409293099e-05 |
| **4014** | 24,8828467214617 | + | NA | + | + | NA | + | NA | + | + | + | + | + | 0,163808066195893 | 11 | -517,853952820532 | 1059,81990564106 | 11,8583327257727 | 9.25493597091589e-05 |
| **3851** | 24,6576745149047 | NA | + | NA | + | NA | NA | NA | NA | + | + | + | + | 0,120029894240896 | 8 | -521,347716379609 | 1059,82043275922 | 11,8588598439271 | 9.25249706994311e-05 |
| **1719** | 24,3046998365915 | NA | + | + | NA | + | + | NA | + | NA | + | + | NA | 0,134574213769706 | 9 | -520,206643879794 | 1059,83061059423 | 11,869037678942 | 9.2055314789811e-05 |
| **3626** | 23,9536998218442 | + | NA | NA | + | NA | + | NA | NA | NA | + | + | + | 0,119956901636391 | 8 | -521,353395335214 | 1059,83179067043 | 11,870217755137 | 9.20010146681559e-05 |
| **3713** | 24,9467720674528 | NA | NA | NA | NA | NA | NA | NA | + | NA | + | + | + | 0,0906874994151501 | 6 | -523,59348272756 | 1059,83311930127 | 11,871546385983 | 9.19399172713965e-05 |
| **3649** | 24,4921553536538 | NA | NA | NA | NA | NA | NA | + | NA | NA | + | + | + | 0,0906841955957352 | 6 | -523,593731490008 | 1059,83361682617 | 11,8720439108799 | 9.19170489169776e-05 |
| **541** | 25,1669502335132 | NA | NA | + | + | + | NA | NA | NA | NA | + | NA | NA | 0,0906827406591513 | 6 | -523,593841039773 | 1059,8338359257 | 11,8722630104098 | 9.19069799774095e-05 |
| **3907** | 24,6538054897696 | NA | + | NA | NA | NA | NA | + | NA | + | + | + | + | 0,119901245516369 | 8 | -521,357725164884 | 1059,84045032977 | 11,8788774144766 | 9.16035270929299e-05 |
| **1985** | 25,667507885599 | NA | NA | NA | NA | NA | NA | + | + | + | + | + | NA | 0,105231186875308 | 7 | -522,489560153215 | 1059,84733736069 | 11,8857644454029 | 9.12886314161394e-05 |
| **3589** | 24,8720342172667 | NA | NA | + | NA | NA | NA | NA | NA | NA | + | + | + | 0,0905645210823067 | 6 | -523,602741822976 | 1059,85163749211 | 11,8900645768151 | 9.10925657129748e-05 |
| **1802** | 25,2379404106192 | + | NA | NA | + | NA | NA | NA | NA | + | + | + | NA | 0,105177520647956 | 7 | -522,493666506404 | 1059,85555006707 | 11,8939771517812 | 9.09145366616522e-05 |
| **3948** | 24,3776428349038 | + | + | NA | + | NA | + | + | NA | + | + | + | + | 0,163583320798586 | 11 | -517,872351675054 | 1059,85670335011 | 11,895130434817 | 9.08621266775702e-05 |
| **2629** | 24,4135539378148 | NA | NA | + | NA | NA | NA | + | NA | NA | + | NA | + | 0,0904894548263928 | 6 | -523,608392980947 | 1059,86293980805 | 11,9013668927571 | 9.0579239045308e-05 |
| **1711** | 24,1958970996177 | NA | + | + | + | NA | + | NA | + | NA | + | + | NA | 0,134331613962829 | 9 | -520,225833735409 | 1059,86899030546 | 11,9074173901734 | 9.0305628398208e-05 |
| **1922** | 25,6321280897027 | + | NA | NA | NA | NA | NA | NA | + | + | + | + | NA | 0,105064572216165 | 7 | -522,502308123678 | 1059,87283330162 | 11,9112603863282 | 9.01322729113158e-05 |
| **3971** | 25,1034785904628 | NA | + | NA | NA | NA | NA | NA | + | + | + | + | + | 0,119646372135709 | 8 | -521,377549826935 | 1059,88009965387 | 11,9185267385803 | 8.98054006410087e-05 |
| **3593** | 24,4633538256009 | NA | NA | NA | + | NA | NA | NA | NA | NA | + | + | + | 0,0903576253896996 | 6 | -523,618316271717 | 1059,88278638959 | 11,9212134742968 | 8.96848399491024e-05 |
| **1596** | 24,0845096555999 | + | + | NA | + | + | + | NA | NA | NA | + | + | NA | 0,134242691113363 | 9 | -520,232866263938 | 1059,88305536252 | 11,9214824472301 | 8.96727793628753e-05 |
| **709** | 25,2797890681467 | NA | NA | + | NA | NA | NA | + | + | NA | + | NA | NA | 0,090164281529081 | 6 | -523,632867381652 | 1059,91188860946 | 11,9503156941678 | 8.8389274798528e-05 |
| **1865** | 25,2591837413812 | NA | NA | NA | + | NA | NA | + | NA | + | + | + | NA | 0,104785648601695 | 7 | -522,523643734298 | 1059,91550452286 | 11,9539316075693 | 8.8229615189201e-05 |
| **1704** | 24,4635951736652 | + | + | + | NA | NA | + | NA | + | NA | + | + | NA | 0,133967225749365 | 9 | -520,25464706212 | 1059,92661695889 | 11,9650440435944 | 8.77407515860901e-05 |
| **665** | 25,1391792434149 | NA | NA | NA | + | + | NA | NA | + | NA | + | NA | NA | 0,089988454323042 | 6 | -523,646097500438 | 1059,93834884703 | 11,9767759317394 | 8.72275758460008e-05 |
| **1655** | 24,1818213656949 | NA | + | + | NA | + | + | + | NA | NA | + | + | NA | 0,133883988101259 | 9 | -520,261227225668 | 1059,93977728598 | 11,9782043706912 | 8.71652984550091e-05 |
| **1858** | 25,2651639961254 | + | NA | NA | NA | NA | NA | + | NA | + | + | + | NA | 0,104615372302102 | 7 | -522,536665356277 | 1059,94154776682 | 11,9799748515275 | 8.70881703532506e-05 |
| **1561** | 25,1459715596834 | NA | NA | NA | + | + | NA | NA | NA | NA | + | + | NA | 0,0899497124155542 | 6 | -523,649012291901 | 1059,94417842996 | 11,9826055146661 | 8.69736958359461e-05 |
| **3967** | 24,7917596899326 | NA | + | + | + | + | + | + | NA | + | + | + | + | 0,177588665425195 | 12 | -516,716252280197 | 1059,94863359265 | 11,9870606773613 | 8.67801704801314e-05 |
| **3071** | 24,7865189105283 | NA | + | + | + | + | + | + | + | + | + | NA | + | 0,177566167621829 | 12 | -516,718125141347 | 1059,95237931495 | 11,9908063996618 | 8.66177953707093e-05 |
| **1543** | 25,0799497998217 | NA | + | + | NA | NA | NA | NA | NA | NA | + | + | NA | 0,0898673883576495 | 6 | -523,655205624244 | 1059,95656509464 | 11,9949921793518 | 8.6436703435218e-05 |
| **538** | 25,0679255165377 | + | NA | NA | + | + | NA | NA | NA | NA | + | NA | NA | 0,0898079557078987 | 6 | -523,659676461936 | 1059,96550677003 | 12,0039338547356 | 8.60511215428298e-05 |
| **2845** | 25,3253115957935 | NA | NA | + | + | + | NA | NA | NA | + | + | NA | + | 0,119072450345367 | 8 | -521,422169843729 | 1059,96933968746 | 12,0077667721678 | 8.58863660449006e-05 |
| **2824** | 24,9211413185984 | + | + | + | NA | NA | NA | NA | NA | + | + | NA | + | 0,119067391776676 | 8 | -521,42256299708 | 1059,97012599416 | 12,0085530788695 | 8.58526061691374e-05 |
| **2573** | 24,3728763898824 | NA | NA | + | NA | + | NA | NA | NA | NA | + | NA | + | 0,0897692951107713 | 6 | -523,662584559526 | 1059,97132296521 | 12,0097500499155 | 8.58012399997185e-05 |
| **2969** | 25,325282929522 | NA | NA | NA | + | + | NA | NA | + | + | + | NA | + | 0,118960052438829 | 8 | -521,430904908059 | 1059,98680981612 | 12,0252369008269 | 8.51394102155466e-05 |
| **4068** | 24,6602172149464 | + | + | NA | NA | NA | + | + | + | + | + | + | + | 0,162766304903776 | 11 | -517,939195339034 | 1059,99039067807 | 12,0288177627774 | 8.49871103601577e-05 |
| **2948** | 25,0076963107994 | + | + | NA | NA | NA | NA | NA | + | + | + | NA | + | 0,118908653671918 | 8 | -521,434899020012 | 1059,99479804002 | 12,033225124733 | 8.48000320878215e-05 |
| **2891** | 24,4576801843279 | NA | + | NA | + | NA | NA | + | NA | NA | + | + | + | 0,118885813732031 | 8 | -521,436673798691 | 1059,99834759738 | 12,0367746820905 | 8.46496642731811e-05 |
| **2905** | 25,2323141217278 | NA | NA | NA | + | + | NA | + | NA | + | + | NA | + | 0,118824756630103 | 8 | -521,441418018115 | 1060,00783603623 | 12,0462631209386 | 8.42490188181335e-05 |
| **1798** | 25,5382515675216 | + | NA | + | NA | NA | NA | NA | NA | + | + | + | NA | 0,104106160457156 | 7 | -522,575591796967 | 1060,0194006482 | 12,0578277329066 | 8.37632709373323e-05 |
| **696** | 24,2841045468615 | + | + | + | NA | + | + | NA | + | NA | + | NA | NA | 0,133356691909901 | 9 | -520,302896751852 | 1060,02311633835 | 12,0615434230592 | 8.36077962255299e-05 |
| **2957** | 25,3360360425145 | NA | NA | + | + | NA | NA | NA | + | + | + | NA | + | 0,118707079464306 | 8 | -521,450560766299 | 1060,0261215326 | 12,0645486173069 | 8.34822617288988e-05 |
| **3636** | 24,2945669604531 | + | + | NA | NA | + | + | NA | NA | NA | + | + | + | 0,133331765834742 | 9 | -520,304865901961 | 1060,02705463857 | 12,0654817232771 | 8.34433219149332e-05 |
| **1861** | 25,5700301191809 | NA | NA | + | NA | NA | NA | + | NA | + | + | + | NA | 0,104039173200116 | 7 | -522,580710957775 | 1060,02963896981 | 12,0680660545238 | 8.33355689551855e-05 |
| **3865** | 25,2509352450146 | NA | NA | NA | + | + | NA | NA | NA | + | + | + | + | 0,11867045292807 | 8 | -521,453406160258 | 1060,03181232052 | 12,0702394052257 | 8.32450594326837e-05 |
| **2842** | 25,2559183359935 | + | NA | NA | + | + | NA | NA | NA | + | + | NA | + | 0,11859518194291 | 8 | -521,459253342098 | 1060,0435066842 | 12,0819337689061 | 8.27597307182697e-05 |
| **713** | 24,8565073508766 | NA | NA | NA | + | NA | NA | + | + | NA | + | NA | NA | 0,0892604317316881 | 6 | -523,700850385151 | 1060,04785461646 | 12,0862817011653 | 8.25800092917234e-05 |
| **632** | 24,2051804172587 | + | + | + | NA | + | + | + | NA | NA | + | NA | NA | 0,133183059084215 | 9 | -520,316612521842 | 1060,05054787833 | 12,0889749630398 | 8.24688793387195e-05 |
| **2623** | 24,0245963525707 | NA | + | + | + | + | + | NA | NA | NA | + | NA | + | 0,133181031021136 | 9 | -520,316772708313 | 1060,05086825127 | 12,0892953359812 | 8.24556699980044e-05 |
| **1584** | 24,0337725772432 | + | + | + | + | NA | + | NA | NA | NA | + | + | NA | 0,132945087459651 | 9 | -520,335406143493 | 1060,08813512163 | 12,1265622063409 | 8.09334636256445e-05 |
| **3844** | 24,8546368082036 | + | + | NA | NA | NA | NA | NA | NA | + | + | + | + | 0,118303269459387 | 8 | -521,481924894589 | 1060,08884978918 | 12,1272768738866 | 8.09045485322144e-05 |
| **2828** | 24,5677471724338 | + | + | NA | + | NA | NA | NA | NA | + | + | NA | + | 0,118278420338959 | 8 | -521,483854469435 | 1060,09270893887 | 12,1311360235795 | 8.0748587667871e-05 |
| **1920** | 24,7264123835105 | + | + | + | + | + | + | + | NA | + | + | + | NA | 0,176707502860964 | 12 | -516,7895567646127 | 1060,09526432451 | 12,1336914092221 | 8.06454816600071e-05 |

| | | | | | | | | | | | | | | | | | | | |
|---|---|---|---|---|---|---|---|---|---|---|---|---|---|---|---|---|---|---|---|
| **3586** | 24,6927923274018 | + | NA | NA | NA | NA | NA | NA | NA | NA | + | + | + | 0,0889177243508135 | 6 | -523,726609467681 | 1060,09937278152 | 12,1377998662258 | 8.04799874526925e-05 |
| **2571** | 23,9383060418185 | NA | + | NA | + | NA | NA | NA | NA | NA | + | NA | + | 0,0888730675242154 | 6 | -523,729965317352 | 1060,10608448086 | 12,1445115655677 | 8.02103613786073e-05 |
| **2690** | 24,73704125272 | + | NA | NA | NA | NA | NA | NA | + | NA | + | NA | + | 0,0888659813824602 | 6 | -523,730497808202 | 1060,10714946256 | 12,1455765472681 | 8.01676614647461e-05 |
| **2884** | 24,5576830208964 | + | + | NA | NA | NA | NA | + | NA | + | + | NA | + | 0,118172767698763 | 8 | -521,49205796297 | 1060,10911592594 | 12,1475430106498 | 8.00888768175415e-05 |
| **4012** | 24,442426413049 | + | + | NA | + | NA | + | NA | + | + | + | + | + | 0,161980970398233 | 11 | -518,003385585154 | 1060,11877117031 | 12,1571982550174 | 7.9703169752237e-05 |
| **2743** | 24,2523978408572 | NA | + | + | NA | + | + | NA | + | NA | + | NA | + | 0,132693480599612 | 9 | -520,355270990894 | 1060,12786481643 | 12,1662919011435 | 7.93415961702798e-05 |
| **792** | 25,6508838327182 | + | + | + | NA | + | NA | NA | NA | + | + | NA | NA | 0,11805017961305 | 8 | -521,501575193166 | 1060,12815038633 | 12,1665774710423 | 7.93302681932377e-05 |
| **1024** | 24,7062766507134 | + | + | + | + | + | + | + | + | + | + | NA | NA | 0,176449996305525 | 12 | -516,810978155165 | 1060,13808534259 | 12,1765124272979 | 7.89371739729125e-05 |
| **2564** | 24,1567078693306 | + | + | NA | NA | NA | NA | NA | NA | NA | + | NA | + | 0,0885823399134084 | 6 | -523,751808757391 | 1060,14977136094 | 12,1881984456459 | 7.84772882079951e-05 |
| **1716** | 24,2300528125491 | + | + | NA | NA | + | + | NA | + | NA | + | + | NA | 0,132507146753436 | 9 | -520,369978695823 | 1060,15728022629 | 12,1957073110016 | 7.81832029198322e-05 |
| **1652** | 24,2264529730561 | + | + | NA | NA | + | + | + | NA | NA | + | + | NA | 0,132505487229112 | 9 | -520,370109671226 | 1060,1575421771 | 12,1959692618079 | 7.81729635138776e-05 |
| **3751** | 24,3723325067807 | NA | + | + | NA | NA | + | NA | + | NA | + | + | + | 0,132421697014855 | 9 | -520,376722359416 | 1060,17076755348 | 12,2091946381863 | 7.76577354799767e-05 |
| **916** | 25,5186090771581 | + | + | NA | NA | + | NA | NA | + | + | + | NA | NA | 0,117727378965541 | 8 | -521,526629772302 | 1060,1782595446 | 12,2166866293128 | 7.73673741331171e-05 |
| **2620** | 24,0464365405525 | + | + | NA | + | + | + | NA | NA | NA | + | NA | + | 0,132353361894047 | 9 | -520,382114865621 | 1060,18155256589 | 12,2199796505965 | 7.72400927421103e-05 |
| **967** | 25,3905223034954 | NA | + | + | NA | NA | NA | + | + | + | + | NA | NA | 0,117691218098212 | 8 | -521,5294358729 | 1060,1838717458 | 12,2222988305102 | 7.71505778166358e-05 |
| **2679** | 24,1764587962656 | NA | + | + | NA | + | + | + | NA | NA | + | NA | + | 0,132294671192475 | 9 | -520,386745966196 | 1060,19081476704 | 12,229241851747 | 7.68832131147403e-05 |
| **647** | 25,1388961783302 | NA | + | + | NA | NA | NA | NA | + | NA | + | NA | NA | 0,0882956809399784 | 6 | -523,773339687869 | 1060,19283322189 | 12,2312603066014 | 7.68056596085209e-05 |
| **653** | 25,2021018791372 | NA | NA | + | + | NA | NA | NA | + | NA | + | NA | NA | 0,0882529906246166 | 6 | -523,776545574834 | 1060,19924499582 | 12,2376720805307 | 7.65598236173207e-05 |
| **2633** | 24,0901283246497 | NA | NA | NA | + | NA | NA | + | NA | NA | + | NA | + | 0,0882379316720603 | 6 | -523,777676411444 | 1060,20150666904 | 12,2399337537513 | 7.64732958994736e-05 |
| **2626** | 24,2552292475319 | + | NA | NA | NA | NA | NA | + | NA | NA | + | NA | + | 0,0882238378483103 | 6 | -523,778734755784 | 1060,20362335772 | 12,2420504424308 | 7.63924036331028e-05 |
| **3017** | 25,1207435548829 | NA | NA | NA | + | NA | NA | + | + | + | + | NA | + | 0,117554649419349 | 8 | -521,540032630941 | 1060,20506526188 | 12,2434923465921 | 7.63373482192995e-05 |
| **1547** | 24,602581032218 | NA | + | NA | + | NA | NA | NA | NA | NA | + | + | NA | 0,0881576230988369 | 6 | -523,783706786225 | 1060,2135674186 | 12,2519945033134 | 7.60135209653654e-05 |
| **1767** | 24,3151910599929 | NA | + | + | NA | NA | + | + | + | NA | + | + | NA | 0,132149807416362 | 9 | -520,398175375894 | 1060,21367358643 | 12,2521006711434 | 7.60094859771799e-05 |
| **2566** | 24,6426022653678 | + | NA | + | NA | NA | NA | NA | NA | NA | + | NA | + | 0,0881538188269756 | 6 | -523,78399243602 | 1060,21413871819 | 12,2525658029035 | 7.59918108195779e-05 |
| **1812** | 25,576306386604 | + | + | NA | NA | + | NA | NA | NA | + | + | + | NA | 0,1174792554287 | 8 | -521,545881966106 | 1060,21676393221 | 12,2551910169204 | 7.58921288720343e-05 |
| **703** | 24,0023285629614 | NA | + | + | + | + | + | NA | + | NA | + | NA | NA | 0,132083960002747 | 9 | -520,403369950907 | 1060,22406273646 | 12,2624898211689 | 7.56156727323529e-05 |
| **3643** | 24,0351568239719 | NA | + | NA | + | + | + | NA | NA | NA | + | + | + | 0,132011536538374 | 9 | -520,409082842977 | 1060,2354885206 | 12,273915605309 | 7.51849201482831e-05 |
| **688** | 24,1188221868753 | + | + | + | + | NA | + | NA | + | NA | + | NA | NA | 0,13197924620495 | 9 | -520,411629808519 | 1060,24058245168 | 12,279009536393 | 7.49936704040268e-05 |
| **1729** | 25,1423098465129 | NA | NA | NA | NA | NA | NA | + | + | NA | + | + | NA | 0,0879584890143346 | 6 | -523,798657484745 | 1060,24346881564 | 12,2818959003534 | 7.4885518950155e-05 |
| **706** | 24,9334172506109 | + | NA | NA | NA | NA | NA | + | + | NA | + | NA | NA | 0,0879112164245416 | 6 | -523,802206163078 | 1060,25056617231 | 12,2889932570199 | 7.46202452954044e-05 |
| **651** | 24,682768238937 | NA | + | NA | + | NA | NA | NA | + | NA | + | NA | NA | 0,0879103948025412 | 6 | -523,802267839313 | 1060,25068952478 | 12,2891166094898 | 7.46156431415458e-05 |
| **919** | 25,7173132923799 | NA | + | + | NA | + | NA | NA | + | + | + | NA | NA | 0,11725939486143 | 8 | -521,562936682904 | 1060,25087336581 | 12,2893004505181 | 7.46087847484838e-05 |
| **527** | 24,4967309068911 | NA | + | + | + | NA | NA | NA | NA | NA | + | NA | NA | 0,0878907370091996 | 6 | -523,803743463233 | 1060,25364077262 | 12,2920678573284 | 7.45056197103395e-05 |
| **1667** | 25,096158246636 | NA | + | NA | NA | NA | NA | NA | + | NA | + | + | NA | 0,0878189007871919 | 6 | -523,809135621539 | 1060,26442508923 | 12,3028521739413 | 7.41049548092182e-05 |
| **599** | 24,6591078919392 | NA | + | + | NA | + | NA | + | NA | NA | + | NA | NA | 0,102449161291472 | 7 | -522,702107366513 | 1060,27243178729 | 12,3108588719988 | 7.38088798514843e-05 |
| **2643** | 24,3031812763768 | NA | + | NA | NA | + | NA | + | NA | NA | + | NA | + | 0,102394024309781 | 7 | -522,706313183241 | 1060,28084342075 | 12,3192705054553 | 7.34991051144284e-05 |
| **3068** | 24,7141407394587 | + | + | NA | + | + | + | + | + | + | + | NA | + | 0,175540412770422 | 12 | -516,886552402141 | 1060,28923383654 | 12,3276609212487 | 7.31914069683815e-05 |
| **2954** | 25,139126949332 | + | NA | NA | + | NA | NA | NA | + | + | + | NA | + | 0,116956466543419 | 8 | -521,586428061706 | 1060,29785612341 | 12,3362832081223 | 7.28765475044205e-05 |
| **2666** | 23,6677182447789 | + | NA | NA | + | NA | + | + | NA | NA | + | NA | + | 0,116923026279762 | 8 | -521,589020781852 | 1060,3030415637 | 12,3414686484125 | 7.26878437453267e-05 |
| **620** | 23,5116549221305 | + | + | NA | + | NA | + | + | NA | NA | + | NA | NA | 0,116918569884453 | 8 | -521,58936629163 | 1060,30373258326 | 12,3421596679698 | 7.26627337226534e-05 |
| **3699** | 24,1828796686532 | NA | + | NA | NA | + | + | + | NA | NA | + | + | + | 0,131569579702371 | 9 | -520,443934865255 | 1060,30519256515 | 12,3436196498642 | 7.26097099406316e-05 |
| **3763** | 24,1781738799531 | NA | + | NA | NA | + | + | NA | + | NA | + | + | + | 0,131488619765276 | 9 | -520,450317317575 | 1060,3179574698 | 12,3563845545054 | 7.21477576911858e-05 |
| **1925** | 26,0189499854556 | NA | NA | + | NA | NA | NA | NA | + | + | + | + | NA | 0,102141759392847 | 7 | -522,725552510806 | 1060,31932207588 | 12,3577491605859 | 7.20985478466913e-05 |
| **3977** | 25,1810708585004 | NA | NA | NA | + | NA | NA | NA | + | + | + | + | + | 0,116809729691878 | 8 | -521,59780426387 | 1060,32060852774 | 12,3590356124487 | 7.20521871029138e-05 |
| **1815** | 25,8397210863791 | NA | + | + | NA | + | NA | NA | NA | + | + | + | NA | 0,116803926183942 | 8 | -521,598254158903 | 1060,32150831781 | 12,3599354025155 | 7.20197784725783e-05 |
| **2660** | 23,649960559226 | + | + | NA | NA | NA | + | + | NA | NA | + | NA | + | 0,11676906036524 | 8 | -521,600956937615 | 1060,32691387523 | 12,3653409599394 | 7.18253877643508e-05 |
| **1939** | 25,6351120532197 | NA | + | NA | NA | + | NA | NA | + | + | + | + | NA | 0,116753098539386 | 8 | -521,602194253818 | 1060,32938850764 | 12,3678155923451 | 7.17365720062146e-05 |
| **2570** | 24,2356722090842 | + | NA | NA | + | NA | NA | NA | NA | NA | + | NA | + | 0,0873770026078997 | 6 | -523,842295974564 | 1060,33074579528 | 12,3691728799902 | 7.16879049404421e-05 |
| **534** | 25,341910218031 | + | NA | + | NA | + | NA | NA | NA | NA | + | NA | NA | 0,0873671107666114 | 6 | -523,843038081695 | 1060,33223000954 | 12,3706570942534 | 7.1634724570143e-05 |
| **3964** | 24,7477090954445 | + | + | NA | + | + | + | + | NA | + | + | + | + | 0,17526112975439 | 12 | -516,909740370793 | 1060,33560977384 | 12,3740368585525 | 7.15137725539142e-05 |
| **1619** | 24,7061878454739 | NA | + | NA | NA | + | NA | + | NA | NA | + | + | NA | 0,101934089422827 | 7 | -522,741386689765 | 1060,35099043379 | 12,3894175185028 | 7.09659173389608e-05 |
| **2604** | 23,507569708359 | + | + | NA | + | NA | + | NA | NA | NA | + | NA | + | 0,11661040239841 | 8 | -521,613254667621 | 1060,35150933524 | 12,3899364199522 | 7.09475075685968e-05 |
| **2898** | 25,3513865072651 | + | NA | NA | NA | + | NA | + | NA | + | + | NA | + | 0,116605806619909 | 8 | -521,613610857887 | 1060,35222171577 | 12,3906488004825 | 7.09222412571373e-05 |
| **1554** | 25,3151395282511 | + | NA | NA | NA | + | NA | NA | NA | NA | + | + | NA | 0,0871192318785864 | 6 | -523,861631862549 | 1060,36941757125 | 12,4078446559611 | 7.03150708955328e-05 |
| **2901** | 25,4588529722068 | NA | NA | + | NA | + | NA | + | NA | + | + | NA | + | 0,116482453768794 | 8 | -521,623170479772 | 1060,37134095954 | 12,4097680442535 | 7.02474818086858e-05 |
| **2047** | 24,657262717751 | NA | + | + | + | + | + | + | + | + | + | + | NA | 0,175035642547677 | 12 | -516,928456121422 | 1060,3730412751 | 12,4114683598104 | 7.01877857447401e-05 |
| **3025** | 25,4562436791233 | NA | NA | NA | NA | + | NA | + | + | + | + | NA | + | 0,116466806103334 | 8 | -521,624383050074 | 1060,37376610015 | 12,4121931848581 | 7.01623534209412e-05 |
| **3008** | 24,7481487766162 | + | + | + | + | + | + | NA | + | + | + | NA | + | 0,17502035625804 | 12 | -516,929724719583 | 1060,37557847142 | 12,4140055561336 | 7.00988021029554e-05 |
| **3921** | 25,3768150905718 | NA | NA | NA | NA | + | NA | + | NA | + | + | + | + | 0,116446434809837 | 8 | -521,625961632025 | 1060,37692326405 | 12,4153503487594 | 7.00516837697551e-05 |
| **2622** | 24,3862019988316 | + | NA | + | + | + | + | NA | NA | NA | + | NA | + | 0,131106765664689 | 9 | -520,480412656366 | 1060,37814814738 | 12,4165752320869 | 7.00087943349779e-05 |
| **658** | 25,279812064362 | + | NA | NA | NA | + | NA | NA | + | NA | + | NA | NA | 0,0870111735179039 | 6 | -523,869735908813 | 1060,38562566378 | 12,4240527484883 | 6.9747537074227e-05 |
| **4091** | 24,6826613829514 | NA | + | NA | + | + | + | + | + | + | + | + | + | 0,174946416147425 | 12 | -516,93586062444 | 1060,38785028114 | 12,4262773658481 | 6.96699994142366e-05 |
| **1710** | 24,5087692085739 | + | NA | + | + | NA | + | NA | + | NA | + | + | NA | 0,131020325813643 | 9 | -520,487223466874 | 1060,39176976839 | 12,4301968531029 | 6.95335977766412e-05 |
| **702** | 24,3916398685714 | + | NA | + | + | + | + | NA | + | NA | + | NA | NA | 0,131014717644882 | 9 | -520,487665325025 | 1060,39265348469 | 12,431080569404 | 6.95028805765478e-05 |
| **1723** | 23,9956857164081 | NA | + | NA | + | + | + | NA | + | NA | + | + | NA | 0,130999609584263 | 9 | -520,488855649438 | 1060,39503413352 | 12,4334612182317 | 6.94201988197826e-05 |
| **700** | 23,9687681984401 | + | + | NA | + | + | + | NA | + | NA | + | NA | NA | 0,130968916670656 | 9 | -520,491273799791 | 1060,39987043423 | 12,4382975189378 | 6.9252533143607e-05 |
| **580** | 24,1741320704802 | + | + | NA | NA | NA | NA | + | NA | NA | + | NA | NA | 0,0869017380867257 | 6 | -523,877942254092 | 1060,40203835434 | 12,4404654390466 | 6.91775068341196e-05 |
| **3904** | 24,7132665848975 | + | + | + | + | + | + | NA | NA | + | + | + | + | 0,17484279389839 | 12 | -516,944458770679 | 1060,40504657362 | 12,443473658326 | 6.90735344917462e-05 |
| **3624** | 24,1954554277482 | + | + | + | NA | NA | + | NA | NA | NA | + | + | + | 0,130929399936916 | 9 | -520,494387011747 | 1060,40609685814 | 12,4445239428492 | 6.9037270582315e-05 |
| **1598** | 24,3991453726135 | + | NA | + | + | + | + | NA | NA | NA | + | + | NA | 0,13092090641782 | 9 | -520,495056130692 | 1060,40743509603 | 12,4458621807391 | 6.8991091887901e-05 |
| **587** | 24,0802498405788 | NA | + | NA | + | NA | NA | + | NA | NA | + | NA | NA | 0,0868618940589606 | 6 | -523,880929834354 | 1060,40801351486 | 12,4464405995716 | 6.89711418994927e-05 |
| **636** | 23,9419255286599 | + | + | NA | + | + | + | + | NA | NA | + | NA | NA | 0,13090282319879 | 9 | -520,496480703874 | 1060,41028424239 | 12,448711327103 | 6.88928790009389e-05 |
| **1640** | 24,1922318976043 | + | + | + | NA | NA | + | + | NA | NA | + | + | NA | 0,13087301416932 | 9 | -520,49882895694 | 1060,41498074853 | 12,4534078332356 | 6.87312908857248e-05 |
| **2676** | 24,1930173624361 | + | + | NA | NA | + | + | + | NA | NA | + | NA | + | 0,130866922071166 | 9 | -520,499308861624 | 1060,41594055789 | 12,4543676426026 | 6.86983143307591e-05 |
| **2740** | 24,1809503101571 | + | + | NA | NA | + | + | NA | + | NA | + | NA | + | 0,130850329350022 | 9 | -520,500615935233 | 1060,41855470511 | 12,4569817898216 | 6.86085792350439e-05 |
| **1557** | 25,556034164746 | NA | NA | + | NA | + | NA | NA | NA | NA | + | + | NA | 0,0867737462587916 | 6 | -523,887538859501 | 1060,42123156515 | 12,4596586498642 | 6.85168128780002e-05 |

| | | | | | | | | | | | | | | | | | |
|---|---|---|---|---|---|---|---|---|---|---|---|---|---|---|---|---|---|
| **1659** | 23,9499760961944 | NA | + | NA | + | + | + | + | NA | NA | + | + | NA | 0,130828489683826 | 9 | -520,502336293231 | 1060,42199542111 | 12,4604225058167 | 6.84906493869203e-05 |
| **661** | 25,5137804187889 | NA | NA | + | NA | + | NA | NA | + | NA | + | NA | NA | 0,0867235849208459 | 6 | -523,891299503003 | 1060,42875285216 | 12,4671799368698 | 6.82596294614901e-05 |
| **1540** | 24,7833116524455 | + | + | NA | NA | NA | NA | NA | NA | NA | + | + | NA | 0,086577018625472 | 6 | -523,902286535612 | 1060,45072691738 | 12,4891540020867 | 6.7513763616016e-05 |
| **3013** | 25,4527102171816 | NA | NA | + | NA | NA | NA | + | + | + | + | NA | + | 0,115943626410528 | 8 | -521,664912986458 | 1060,45482597292 | 12,493253057625 | 6.73755339838072e-05 |
| **639** | 23,8601067788801 | NA | + | + | + | + | + | + | NA | NA | + | NA | NA | 0,13060686558232 | 9 | -520,519791662212 | 1060,45690615907 | 12,4953332437799 | 6.73054935879007e-05 |
| **759** | 24,1080605365308 | NA | + | + | NA | + | + | + | + | NA | + | NA | NA | 0,13059004333898 | 9 | -520,521116419826 | 1060,4595556743 | 12,4979827590071 | 6.7216389156752e-05 |
| **638** | 24,2775317195863 | + | NA | + | + | + | + | + | NA | NA | + | NA | NA | 0,130555326092602 | 9 | -520,523850334264 | 1060,46502350317 | 12,5034505878839 | 6.70328762685084e-05 |
| **911** | 25,4753482826737 | NA | + | + | + | NA | NA | NA | + | + | + | NA | NA | 0,115847809199049 | 8 | -521,672333202904 | 1060,46966640581 | 12,5080934905179 | 6.68774431945728e-05 |
| **1605** | 25,0301265319148 | NA | NA | + | NA | NA | NA | + | NA | NA | + | + | NA | 0,0864438458825783 | 6 | -523,912268021032 | 1060,47068988822 | 12,5091169729267 | 6.68432280066541e-05 |
| **3858** | 25,4196938819892 | + | NA | NA | NA | + | NA | NA | NA | + | + | + | + | 0,115830410179423 | 8 | -521,673680520653 | 1060,47236104131 | 12,5107881260153 | 6.67873987012626e-05 |
| **589** | 24,6601023107208 | NA | NA | + | + | NA | NA | + | NA | NA | + | NA | NA | 0,0864301528011079 | 6 | -523,913294254395 | 1060,47274235494 | 12,511169439653 | 6.67746664420709e-05 |
| **1681** | 25,4598440773388 | NA | NA | NA | NA | + | NA | NA | + | NA | + | + | NA | 0,0864096107613236 | 6 | -523,914833756794 | 1060,47582135974 | 12,5142484444511 | 6.66719457725338e-05 |
| **2838** | 25,4866565896707 | + | NA | + | NA | + | NA | NA | NA | + | + | NA | + | 0,115791889905394 | 8 | -521,6766632984 | 1060,4783265968 | 12,516753681509 | 6.65884835419791e-05 |
| **2962** | 25,4756670407599 | + | NA | NA | NA | + | NA | NA | + | + | + | NA | + | 0,115732680389595 | 8 | -521,681247873033 | 1060,48749574607 | 12,5259228307759 | 6.62839023904458e-05 |
| **3064** | 24,9540602082163 | + | + | + | NA | + | + | + | + | + | + | NA | + | 0,174284764971259 | 12 | -516,990743160048 | 1060,49761535235 | 12,5360424370638 | 6.59493659507261e-05 |
| **971** | 25,0871014649632 | NA | + | NA | + | NA | NA | + | + | + | + | NA | NA | 0,115590847291762 | 8 | -521,692228718579 | 1060,50945743716 | 12,5478845218674 | 6.55600307311956e-05 |
| **33** | 25,523595505618 | NA | NA | NA | NA | NA | + | NA | NA | NA | NA | NA | NA | 0,0419917217098653 | 3 | -527,165295242648 | 1060,51104161312 | 12,5494686978248 | 6.55081219798362e-05 |
| **1779** | 24,0976139598593 | NA | + | NA | NA | + | + | + | + | NA | + | + | NA | 0,130231639433951 | 9 | -520,549334768179 | 1060,515992371 | 12,5544194557122 | 6.53461650890839e-05 |
| **603** | 24,4686268304164 | NA | + | NA | + | + | NA | + | NA | NA | + | NA | NA | 0,100843682179278 | 7 | -522,824466785736 | 1060,51715062573 | 12,5555777104444 | 6.53083322926718e-05 |
| **2893** | 24,9816741322696 | NA | NA | + | + | NA | NA | + | NA | + | + | NA | + | 0,115504392483617 | 8 | -521,698921263898 | 1060,5228425278 | 12,5612696125058 | 6.51227322067171e-05 |
| **1673** | 25,1190112688959 | NA | NA | NA | + | NA | NA | NA | + | NA | + | + | NA | 0,0860745754179831 | 6 | -523,939937759011 | 1060,52602936418 | 12,564456448885 | 6.50190470894762e-05 |
| **3853** | 25,0686788573946 | NA | NA | + | + | NA | NA | NA | NA | + | + | + | + | 0,115457346759479 | 8 | -521,702562841487 | 1060,53012568297 | 12,568552767684 | 6.48860140000277e-05 |
| **3960** | 25,0022587256319 | + | + | + | NA | + | + | + | NA | + | + | + | + | 0,174087673838013 | 12 | -517,007082947357 | 1060,53029492697 | 12,5687220116818 | 6.48805234481367e-05 |
| **2749** | 24,3603233708681 | NA | NA | + | + | + | + | NA | + | NA | + | NA | + | 0,130139250171572 | 9 | -520,556607002331 | 1060,53053683931 | 12,5689639240168 | 6.48726762232693e-05 |
| **4087** | 24,9268597928118 | NA | + | + | NA | + | + | + | + | + | + | + | + | 0,174065626180453 | 12 | -517,008910560099 | 1060,53395015246 | 12,5723772371648 | 6.47620552667605e-05 |
| **2685** | 24,2664033837475 | NA | NA | + | + | + | + | + | NA | NA | + | NA | + | 0,13010302958419 | 9 | -520,559457822041 | 1060,53623847873 | 12,5746655634371 | 6.46879992846454e-05 |
| **1984** | 24,6952445693627 | + | + | + | + | + | + | NA | + | + | + | + | NA | 0,174019054725642 | 12 | -517,01277088178 | 1060,54167079582 | 12,5800978805273 | 6.45125348251411e-05 |
| **723** | 24,6416420359307 | NA | + | NA | NA | + | NA | + | + | NA | + | NA | NA | 0,100678286107271 | 7 | -522,837059814404 | 1060,54233668307 | 12,5807637677813 | 6.44910593630688e-05 |
| **3966** | 25,0100302898182 | + | NA | + | + | + | + | + | NA | + | + | + | + | 0,173996240731322 | 12 | -517,01466186114 | 1060,54545275454 | 12,5838798392476 | 6.43906582227026e-05 |
| **1725** | 24,3854105970197 | NA | NA | + | + | + | + | NA | + | NA | + | + | NA | 0,129996780258092 | 9 | -520,567819720763 | 1060,55296227617 | 12,5913893608802 | 6.41493400318314e-05 |
| **596** | 24,64560297049 | + | + | NA | NA | + | NA | + | NA | NA | + | NA | NA | 0,10060780725278 | 7 | -522,842425273756 | 1060,55306760178 | 12,591494686485 | 6.41459618367645e-05 |
| **2830** | 25,0145237866641 | + | NA | + | + | NA | NA | NA | NA | + | + | NA | + | 0,115308812700672 | 8 | -521,71405886045 | 1060,5531177209 | 12,5915448056087 | 6.41443543872085e-05 |
| **3010** | 25,2326046717439 | + | NA | NA | NA | NA | NA | + | + | + | + | NA | + | 0,115291307363091 | 8 | -521,715413585547 | 1060,55582717109 | 12,5942542558025 | 6.40575152553514e-05 |
| **2667** | 23,4852873543552 | NA | + | NA | + | NA | + | + | NA | NA | + | NA | + | 0,115277531524564 | 8 | -521,716479668639 | 1060,55795933728 | 12,5963864219868 | 6.3989261010225e-05 |
| **3861** | 25,5663857754977 | NA | NA | + | NA | + | NA | NA | NA | + | + | + | + | 0,115247645776233 | 8 | -521,718792406637 | 1060,56258481327 | 12,6010118979839 | 6.38414416144435e-05 |
| **520** | 24,7413217303713 | + | + | + | NA | NA | NA | NA | NA | NA | + | NA | NA | 0,0858121377126792 | 6 | -523,959595638886 | 1060,56534512393 | 12,6037722086346 | 6.37533912843631e-05 |
| **2683** | 23,969894517985 | NA | + | NA | + | + | + | + | NA | NA | + | NA | + | 0,129855209029714 | 9 | -520,57895989278 | 1060,5752426202 | 12,6136697049142 | 6.34386711853859e-05 |
| **2965** | 25,6244119619797 | NA | NA | + | NA | + | NA | NA | + | + | + | NA | + | 0,115109493397455 | 8 | -521,72948244955 | 1060,5839648991 | 12,6223919838098 | 6.31626087049958e-05 |
| **3985** | 25,5384482722458 | NA | NA | NA | NA | + | NA | NA | + | + | + | + | + | 0,115091567416107 | 8 | -521,730869415303 | 1060,58673883061 | 12,6251659153152 | 6.30750650539302e-05 |
| **3070** | 25,0459171616628 | + | NA | + | + | + | + | + | + | + | + | NA | + | 0,173742583808234 | 12 | -517,035683159931 | 1060,58749535212 | 12,62592243683 | 6.30512107439071e-05 |
| **4093** | 24,9672986002581 | NA | NA | + | + | + | + | + | + | + | + | + | + | 0,173713399157545 | 12 | -517,038101364417 | 1060,59233176109 | 12,6307588458014 | 6.28989242274362e-05 |
| **1549** | 25,03406966598 | NA | NA | + | + | NA | NA | NA | NA | NA | + | + | NA | 0,0856171650787 | 6 | -523,974196400274 | 1060,5945466467 | 12,6329737314113 | 6.28293058226781e-05 |
| **2747** | 23,9819862003922 | NA | + | NA | + | + | + | NA | + | NA | + | NA | + | 0,129716701235621 | 9 | -520,589857251556 | 1060,59703733776 | 12,6354644224675 | 6.27511103279488e-05 |
| **3603** | 24,7329059820082 | NA | + | NA | NA | + | NA | NA | NA | NA | + | + | + | 0,100279699161931 | 7 | -522,867398165599 | 1060,60301338546 | 12,64144047017 | 6.2563888364741e-05 |
| **3645** | 24,3362974545396 | NA | NA | + | + | + | + | NA | NA | NA | + | + | + | 0,129652656826845 | 9 | -520,594895478681 | 1060,60711379201 | 12,6455408767174 | 6.2435751074629e-05 |
| **4031** | 24,6968903319489 | NA | + | + | + | + | + | NA | + | + | + | + | + | 0,173603735757064 | 12 | -517,047187176765 | 1060,61050338579 | 12,6489304704965 | 6.23300247755208e-05 |
| **3638** | 24,6133228290901 | + | NA | + | NA | + | + | NA | NA | NA | + | + | + | 0,129613893882272 | 9 | -520,597944690846 | 1060,61321221634 | 12,6516393010472 | 6.22456611824159e-05 |
| **582** | 24,7901191310338 | + | NA | + | NA | NA | NA | + | NA | NA | + | NA | NA | 0,0854474987710592 | 6 | -523,986899534861 | 1060,61995291588 | 12,6583800005847 | 6.20362246680537e-05 |
| **3757** | 24,431696390732 | NA | NA | + | + | NA | + | NA | + | NA | + | + | + | 0,129507223845716 | 9 | -520,606334982617 | 1060,62999279988 | 12,6684198845892 | 6.17255867626497e-05 |
| **644** | 24,8257427024826 | + | + | NA | NA | NA | NA | NA | + | NA | + | NA | NA | 0,0853305361167004 | 6 | -523,995655307611 | 1060,63746446138 | 12,6758915460859 | 6.14954206162347e-05 |
| **650** | 24,9054046879876 | + | NA | NA | + | NA | NA | NA | + | NA | + | NA | NA | 0,0852997961584565 | 6 | -523,997956301625 | 1060,6420664494 | 12,6804935341136 | 6.13540826927384e-05 |
| **847** | 24,9375404222586 | NA | + | + | + | NA | NA | + | NA | + | + | NA | NA | 0,114732777207882 | 8 | -521,758623761268 | 1060,64224752254 | 12,6806746072461 | 6.13485281562117e-05 |
| **2735** | 24,0642920730218 | NA | + | + | + | NA | + | NA | + | NA | + | NA | + | 0,129421927232512 | 9 | -520,613043376339 | 1060,64340958732 | 12,6818366720329 | 6.13128930276519e-05 |
| **1773** | 24,3727296578818 | NA | NA | + | + | NA | + | + | + | NA | + | + | NA | 0,129404993837446 | 9 | -520,614375072998 | 1060,64607298064 | 12,6845000653504 | 6.1231297195374e-05 |
| **4033** | 25,2945113996461 | NA | NA | NA | NA | NA | NA | + | + | + | + | + | + | 0,114699155960431 | 8 | -521,761223968827 | 1060,64744793765 | 12,6858750223644 | 6.11892164610564e-05 |
| **1661** | 24,248856830592 | NA | NA | + | + | + | + | + | NA | NA | + | + | NA | 0,129367441610562 | 9 | -520,617328208552 | 1060,65197925175 | 12,6904063364598 | 6.10507396112992e-05 |
| **2678** | 24,5587248970911 | + | NA | + | NA | + | + | + | NA | NA | + | NA | + | 0,129357438702362 | 9 | -520,618114823354 | 1060,65355248135 | 12,6919795660631 | 6.10027350788821e-05 |
| **3701** | 24,5438633913794 | NA | NA | + | NA | + | + | + | NA | NA | + | + | + | 0,129356651274971 | 9 | -520,618176745166 | 1060,65367632498 | 12,6921034096879 | 6.09989577959138e-05 |
| **756** | 24,0777795545193 | + | + | NA | NA | + | + | + | + | NA | + | NA | NA | 0,129282732900379 | 9 | -520,623989298185 | 1060,66530143102 | 12,7037285157257 | 6.06454265744699e-05 |
| **2950** | 25,4487251956044 | + | NA | + | NA | NA | NA | NA | + | + | + | NA | + | 0,114581315099739 | 8 | -521,770336792527 | 1060,66567358505 | 12,7041006697636 | 6.06341429041283e-05 |
| **2044** | 24,5918919853417 | + | + | NA | + | + | + | + | + | + | + | + | NA | 0,173234486837273 | 12 | -517,077771264422 | 1060,6716715611 | 12,7100986458106 | 6.04525742329647e-05 |
| **1609** | 24,7072908070393 | NA | NA | NA | + | NA | NA | + | NA | NA | + | + | NA | 0,0850990629076729 | 6 | -524,012979991592 | 1060,67211382934 | 12,7105409140468 | 6.04392075842416e-05 |
| **2742** | 24,5962344472088 | + | NA | + | NA | + | + | NA | + | NA | + | NA | + | 0,129238606657467 | 9 | -520,627458919247 | 1060,67224067314 | 12,7106677578499 | 6.04353745363171e-05 |
| **3765** | 24,5941484087763 | NA | NA | + | NA | + | + | NA | + | NA | + | + | + | 0,129196651858061 | 9 | -520,630757637984 | 1060,67883811061 | 12,7172651953226 | 6.02363436878708e-05 |
| **2682** | 24,2714642866284 | + | NA | NA | + | + | + | + | NA | NA | + | NA | + | 0,129173499596382 | 9 | -520,632577929032 | 1060,68247869271 | 12,7209057774185 | 6.01267957455142e-05 |
| **964** | 25,1329013638778 | + | + | NA | NA | NA | NA | + | + | + | + | NA | NA | 0,114459881491241 | 8 | -521,779726181058 | 1060,68445236212 | 12,7228794468251 | 6.00674898042661e-05 |
| **2803** | 24,0899654851408 | NA | + | NA | NA | + | + | + | + | NA | + | NA | + | 0,129120897772589 | 9 | -520,636713441484 | 1060,69074971761 | 12,7291768023222 | 5.98786540831386e-05 |
| **1718** | 24,6313371947735 | + | NA | + | NA | + | + | NA | + | NA | + | + | NA | 0,129079250598665 | 9 | -520,639987531222 | 1060,69729789709 | 12,7357249817981 | 5.96829265858228e-05 |
| **2734** | 24,3636110287255 | + | NA | + | + | NA | + | NA | + | NA | + | NA | + | 0,129050613332716 | 9 | -520,642238757136 | 1060,70180034892 | 12,7402274336266 | 5.95487179585036e-05 |
| **1654** | 24,569859589202 | + | NA | + | NA | + | + | + | NA | NA | + | + | NA | 0,12897249065969 | 9 | -520,648379741866 | 1060,71408231838 | 12,7525094030871 | 5.91841507373883e-05 |
| **1863** | 25,1644391278374 | NA | + | + | NA | NA | NA | + | NA | + | + | + | NA | 0,114266796058479 | 8 | -521,794653122724 | 1060,71430624545 | 12,7527333301568 | 5.91775246416097e-05 |
| **1658** | 24,27916298841 | + | NA | NA | + | + | + | + | NA | NA | + | + | NA | 0,128865276680932 | 9 | -520,656806609173 | 1060,73093605299 | 12,7693631377008 | 5.86875092573396e-05 |
| **2728** | 24,2910782522734 | + | + | + | NA | NA | + | NA | + | NA | + | NA | + | 0,128822375269088 | 9 | -520,660178309167 | 1060,73767945298 | 12,7761065376899 | 5.84899657984817e-05 |
| **1647** | 23,8803373897018 | NA | + | + | + | NA | + | + | NA | NA | + | + | NA | 0,128818026506596 | 9 | -520,660520077058 | 1060,73836298876 | 12,7767900734714 | 5.84699792218206e-05 |
| **3913** | 24,9021589157971 | NA | NA | NA | + | NA | NA | + | NA | + | + | + | + | 0,114043846469126 | 8 | -521,81188474073 | 1060,74876948146 | 12,7871965661684 | 5.81665356463881e-05 |
| **750** | 24,3007585843889 | + | NA | + | + | NA | + | + | + | NA | + | NA | NA | 0,128707584807368 | 9 | -520,669199084332 | 1060,75572100331 | 12,794148088019 | 5.79647136203664e-05 |

| | | | | | | | | | | | | | | | | | | |
|---|---|---|---|---|---|---|---|---|---|---|---|---|---|---|---|---|---|---|
| **765** | 24,2105986597514 | NA | NA | + | + | + | + | + | + | NA | + | NA | NA | 0,12868324774478 | 9 | -520,671111452723 | 1060,75954574009 | 12,7979728248001 | 5.78539696596472e-05 |
| **2587** | 24,4051460806384 | NA | + | NA | + | + | NA | NA | NA | NA | + | NA | + | 0,0992483490420137 | 7 | -522,945836828249 | 1060,75989071076 | 12,7983177954704 | 5.78439915588631e-05 |
| **3850** | 24,9198834345471 | + | NA | NA | + | NA | NA | NA | NA | + | + | + | + | 0,113961466254643 | 8 | -521,818250753358 | 1060,76150150672 | 12,7999285914259 | 5.77974228807147e-05 |
| **744** | 24,228128722242 | + | + | + | NA | NA | + | + | + | NA | + | NA | NA | 0,128656176572844 | 9 | -520,673238600437 | 1060,76380003552 | 12,802227120229 | 5.77310365151479e-05 |
| **1987** | 25,2518581319777 | NA | + | NA | NA | NA | NA | + | + | + | + | + | NA | 0,113942648260266 | 8 | -521,8197048495 | 1060,764409699 | 12,8028367837092 | 5.77134409448099e-05 |
| **2907** | 24,842318578361 | NA | + | NA | + | + | NA | + | NA | + | + | NA | + | 0,128646979977491 | 9 | -520,673961218206 | 1060,76524527106 | 12,8036723557668 | 5.76893341116069e-05 |
| **3631** | 23,8833152557875 | NA | + | + | + | NA | + | NA | NA | NA | + | + | + | 0,128645256712364 | 9 | -520,674096622032 | 1060,76551607871 | 12,8039431634199 | 5.76815232838363e-05 |
| **2805** | 24,4973389077136 | NA | NA | + | NA | + | + | + | + | NA | + | NA | + | 0,128639082243725 | 9 | -520,674581772507 | 1060,76648637966 | 12,8049134643688 | 5.76535458526291e-05 |
| **3642** | 24,3289641700567 | + | NA | NA | + | + | + | NA | NA | NA | + | + | + | 0,128635460952445 | 9 | -520,674866308945 | 1060,76705545254 | 12,8054825372446 | 5.76371436516823e-05 |
| **2847** | 25,0380723318145 | NA | + | + | + | + | NA | NA | NA | + | + | NA | + | 0,128623628847433 | 9 | -520,675795987039 | 1060,76891480872 | 12,8073418934321 | 5.7583584562059e-05 |
| **3705** | 24,2670239121474 | NA | NA | NA | + | + | + | + | NA | NA | + | + | + | 0,12861259426323 | 9 | -520,676662990557 | 1060,77064881576 | 12,8090759004683 | 5.75336810280533e-05 |
| **1602** | 24,7790232731135 | + | NA | NA | NA | NA | NA | + | NA | NA | + | + | NA | 0,0844391504322925 | 6 | -524,062347306142 | 1060,77084845844 | 12,809275543148 | 5.75279382255579e-05 |
| **2890** | 24,835396653202 | + | NA | NA | + | NA | NA | + | NA | + | + | NA | + | 0,113891884027227 | 8 | -521,823627328009 | 1060,77225465602 | 12,810681740727 | 5.74875046179067e-05 |
| **2903** | 25,0250246974285 | NA | + | + | NA | + | NA | + | NA | + | + | NA | + | 0,128576705697855 | 9 | -520,679482732066 | 1060,77628829878 | 12,8147153834868 | 5.73716794281349e-05 |
| **3970** | 25,3209663766244 | + | NA | NA | NA | NA | NA | NA | + | + | + | + | + | 0,113792223767286 | 8 | -521,831327278196 | 1060,78765455639 | 12,8260816411009 | 5.70465535253347e-05 |
| **751** | 23,9770660963441 | NA | + | + | + | NA | + | + | + | NA | + | NA | NA | 0,128453875711411 | 9 | -520,689132526778 | 1060,7955878882 | 12,8340149729106 | 5.68207171113961e-05 |
| **646** | 25,2892840917243 | + | NA | + | NA | NA | NA | NA | + | NA | + | NA | NA | 0,0842103816161699 | 6 | -524,079452944118 | 1060,80505973439 | 12,8434868190986 | 5.65522547752759e-05 |
| **4084** | 24,8392680762161 | + | + | NA | NA | + | + | + | + | + | + | + | + | 0,172427098755134 | 12 | -517,144597934615 | 1060,80532490149 | 12,8437519861966 | 5.65447573736623e-05 |
| **3973** | 25,5475769520997 | NA | NA | + | NA | NA | NA | NA | + | + | + | + | + | 0,113676427049557 | 8 | -521,840272876291 | 1060,80554575258 | 12,8439728372916 | 5.65385137326051e-05 |
| **711** | 24,7924146352577 | NA | + | + | NA | NA | NA | + | + | NA | + | NA | NA | 0,0989474221720312 | 7 | -522,968706708749 | 1060,80563047176 | 12,8440575564719 | 5.65361188350617e-05 |
| **3687** | 24,047355966897 | NA | + | + | NA | NA | + | + | NA | NA | + | + | + | 0,128364928966462 | 9 | -520,696119530762 | 1060,80956189617 | 12,8479889808782 | 5.64250942532853e-05 |
| **2886** | 25,1088530907696 | + | NA | + | NA | NA | NA | + | NA | + | + | NA | + | 0,113589310987059 | 8 | -521,847002049101 | 1060,8190040982 | 12,8574311829104 | 5.61593335187323e-05 |
| **526** | 24,8028802258411 | + | NA | + | + | NA | NA | NA | NA | NA | + | NA | NA | 0,084069544540382 | 6 | -524,089981573782 | 1060,82611699372 | 12,8645440784271 | 5.59599605228961e-05 |
| **2583** | 24,7104452616328 | NA | + | + | NA | + | NA | NA | NA | NA | + | NA | + | 0,0988111639394686 | 7 | -522,979059569273 | 1060,82633619281 | 12,8647632775194 | 5.5953827672706e-05 |
| **2040** | 24,855887171222 | + | + | + | NA | + | + | + | + | + | + | + | NA | 0,172291539079812 | 12 | -517,155811678514 | 1060,82775238929 | 12,8661794739953 | 5.59142208903249e-05 |
| **2746** | 24,2887894210289 | + | NA | NA | + | + | + | NA | + | NA | + | NA | + | 0,128216950288582 | 9 | -520,707742071603 | 1060,83280697785 | 12,8712340625618 | 5.57730876172939e-05 |
| **840** | 25,018464939463 | + | + | + | NA | NA | NA | + | NA | + | + | NA | NA | 0,113497215129084 | 8 | -521,854115161335 | 1060,83323032267 | 12,8716574073787 | 5.57612832428853e-05 |
| **1781** | 24,5172599157297 | NA | NA | + | NA | + | + | + | + | NA | + | + | NA | 0,128202809103266 | 9 | -520,708852645312 | 1060,83502812527 | 12,8734552099793 | 5.57111818743395e-05 |
| **4090** | 24,8811971830311 | + | NA | NA | + | + | + | + | + | + | + | + | + | 0,172246743731411 | 12 | -517,15951682804 | 1060,83516268834 | 12,8735897730476 | 5.57074336666481e-05 |
| **3909** | 25,1938379393873 | NA | NA | + | NA | NA | NA | + | NA | + | + | + | + | 0,113424122180764 | 8 | -521,859760039854 | 1060,84452007971 | 12,8829471644169 | 5.54474043106016e-05 |
| **1722** | 24,3117481204657 | + | NA | NA | + | + | + | NA | + | NA | + | + | NA | 0,128099859493544 | 9 | -520,71693721911 | 1060,85119727286 | 12,8896243575743 | 5.52625964673473e-05 |
| **4024** | 24,8787760691179 | + | + | + | NA | + | + | NA | + | + | + | + | + | 0,172078737497903 | 12 | -517,17341131207 | 1060,8629516564 | 12,9013787411081 | 5.49387601465967e-05 |
| **1807** | 25,2557053444282 | NA | + | + | + | NA | NA | NA | NA | + | + | + | NA | 0,113286369780509 | 8 | -521,870397224054 | 1060,86579444811 | 12,9042215328184 | 5.48607258925598e-05 |
| **2032** | 24,7222120579191 | + | + | + | + | NA | + | + | + | + | + | + | NA | 0,172040817009487 | 12 | -517,176547029826 | 1060,86922309191 | 12,9076501766201 | 5.47667575177e-05 |
| **758** | 24,4995411940347 | + | NA | + | NA | + | + | + | + | NA | + | NA | NA | 0,127965856613779 | 9 | -520,727458959306 | 1060,87224075326 | 12,9106678379665 | 5.46841860629239e-05 |
| **1669** | 25,5751119516217 | NA | NA | + | NA | NA | NA | NA | + | NA | + | + | NA | 0,0836883990281526 | 6 | -524,118466951526 | 1060,88308774921 | 12,9215148339144 | 5.43884092879779e-05 |
| **524** | 24,3462081748544 | + | + | NA | + | NA | NA | NA | NA | NA | + | NA | NA | 0,0836831645635861 | 6 | -524,118858073215 | 1060,88386999258 | 12,9222970772937 | 5.4367140960964e-05 |
| **3698** | 24,5101624092353 | + | NA | NA | NA | + | + | + | NA | NA | + | + | + | 0,127867516037478 | 9 | -520,735179511493 | 1060,88768185763 | 12,9261089423405 | 5.4263619542558e-05 |
| **1708** | 24,0360722643431 | + | + | NA | + | NA | + | NA | + | NA | + | + | NA | 0,127857763976243 | 9 | -520,735945081891 | 1060,88921299843 | 12,9276400831368 | 5.4222092819584e-05 |
| **3750** | 24,6465302499818 | + | NA | + | NA | NA | + | NA | + | NA | + | + | + | 0,127756484271721 | 9 | -520,743895381741 | 1060,90511359813 | 12,9435406828372 | 5.37927200060903e-05 |
| **3056** | 24,7179496453825 | + | + | + | + | NA | + | + | + | + | + | NA | + | 0,171796876751738 | 12 | -517,196715485484 | 1060,90956000323 | 12,9479870879352 | 5.36732607341389e-05 |
| **4079** | 24,7277391633733 | NA | + | + | + | NA | + | + | + | + | + | + | + | 0,171775183210172 | 12 | -517,198508773336 | 1060,91314657893 | 12,9515736636404 | 5.35770953795079e-05 |
| **586** | 24,4635970217431 | + | NA | NA | + | NA | NA | + | NA | NA | + | NA | NA | 0,0834831478068057 | 6 | -524,133801747894 | 1060,91375734194 | 12,952184426651 | 5.3560736423459e-05 |
| **3591** | 24,6019062901492 | NA | + | + | NA | NA | NA | NA | NA | NA | + | + | + | 0,0981994905803421 | 7 | -523,025515063544 | 1060,91924718135 | 12,9576742660611 | 5.34139180970771e-05 |
| **2809** | 24,1924854290105 | NA | NA | NA | + | + | + | + | + | NA | + | NA | + | 0,127640035685078 | 9 | -520,753035275093 | 1060,92339338483 | 12,9618204695412 | 5.33033003111066e-05 |
| **3846** | 25,2325232874238 | + | NA | + | NA | NA | NA | NA | NA | + | + | + | + | 0,112907535138063 | 8 | -521,899642165794 | 1060,92428433159 | 12,9627114162984 | 5.32795603979693e-05 |
| **1666** | 25,2174876853314 | + | NA | NA | NA | NA | NA | NA | + | NA | + | + | NA | 0,0833964840093922 | 6 | -524,140275571247 | 1060,92670498865 | 12,9651320733565 | 5.32151136347495e-05 |
| **763** | 23,8492280135748 | NA | + | NA | + | + | + | + | + | NA | + | NA | NA | 0,127583418371348 | 9 | -520,757478651526 | 1060,9322801377 | 12,9707072224064 | 5.30669791035845e-05 |
| **1646** | 24,2379216195603 | + | NA | + | + | NA | + | + | NA | NA | + | + | NA | 0,127560413557289 | 9 | -520,759284007161 | 1060,93589084897 | 12,9743179336776 | 5.29712607626161e-05 |
| **762** | 24,1779999405981 | + | NA | NA | + | + | + | + | + | NA | + | NA | NA | 0,127537420408467 | 9 | -520,761088399797 | 1060,93949963424 | 12,9779267189488 | 5.28757659907407e-05 |
| **3769** | 24,2871444735841 | NA | NA | NA | + | + | + | NA | + | NA | + | + | + | 0,127536373348573 | 9 | -520,761170566931 | 1060,93966396851 | 12,9780910532172 | 5.28714215190682e-05 |
| **1766** | 24,6168946337528 | + | NA | + | NA | NA | + | + | + | NA | + | + | NA | 0,127493964221672 | 9 | -520,76449850424 | 1060,94631984313 | 12,9847469278347 | 5.26957611982173e-05 |
| **908** | 25,2197360619521 | + | + | NA | + | NA | NA | NA | + | + | + | NA | NA | 0,112759408156214 | 8 | -521,911073746988 | 1060,94714749398 | 12,9855745786854 | 5.26739588639186e-05 |
| **543** | 24,8284655588088 | NA | + | + | + | + | NA | NA | NA | NA | + | NA | NA | 0,0980113444244199 | 7 | -523,039798093056 | 1060,94781324037 | 12,9862403250841 | 5.26564280326434e-05 |
| **1546** | 24,8173891632498 | + | NA | NA | + | NA | NA | NA | NA | NA | + | + | NA | 0,0832388965437948 | 6 | -524,152045855906 | 1060,95024555797 | 12,988672642676 | 5.25924283792868e-05 |
| **2791** | 24,1773656999865 | NA | + | + | NA | NA | + | + | + | NA | + | NA | + | 0,127454784354797 | 9 | -520,767572890747 | 1060,95246861614 | 12,9908957008495 | 5.25340028422573e-05 |
| **1881** | 25,5614851384473 | NA | NA | NA | + | + | NA | + | NA | + | + | + | NA | 0,112716945745087 | 8 | -521,914350397765 | 1060,95370079553 | 12,9921278802399 | 5.25016471524856e-05 |
| **3923** | 24,9819637425669 | NA | + | NA | NA | + | NA | + | NA | + | + | + | + | 0,127423727997125 | 9 | -520,770009739408 | 1060,95734231346 | 12,9957693981717 | 5.24061412806796e-05 |
| **3867** | 24,9952112448461 | NA | + | NA | + | + | NA | NA | NA | + | + | + | + | 0,127413716019531 | 9 | -520,770795314478 | 1060,9589134636 | 12,9973405483106 | 5.23649884890223e-05 |
| **2797** | 24,2387228594792 | NA | NA | + | + | NA | + | + | + | NA | + | NA | + | 0,127411100538886 | 9 | -520,771000532829 | 1060,9593239003 | 12,9977509850123 | 5.2354243335031e-05 |
| **3906** | 25,0128519960153 | + | NA | NA | NA | NA | NA | + | NA | + | + | + | + | 0,112647761958519 | 8 | -521,919688692211 | 1060,96437738442 | 13,0028044691312 | 5.22221246516149e-05 |
| **2046** | 24,8971259400512 | + | NA | + | + | + | + | + | + | + | + | + | NA | 0,171419907375166 | 12 | -517,227870828513 | 1060,97187068928 | 13,0102977739925 | 5.20268325755697e-05 |
| **1542** | 25,1785365528774 | + | NA | + | NA | NA | NA | NA | NA | NA | + | + | NA | 0,0830922850409916 | 6 | -524,162994524766 | 1060,97214289569 | 13,0105699803955 | 5.20197520389451e-05 |
| **784** | 25,1093447700148 | + | + | + | + | NA | NA | NA | NA | + | + | NA | NA | 0,112569994976882 | 8 | -521,925688778504 | 1060,97637755701 | 13,0148046417175 | 5.19097255453141e-05 |
| **3027** | 24,9918242830927 | NA | + | NA | NA | + | NA | + | + | + | + | NA | + | 0,127297680173663 | 9 | -520,77989923894 | 1060,97712131253 | 13,0155483972358 | 5.18904250618351e-05 |
| **1818** | 25,5616747505556 | + | NA | NA | + | + | NA | NA | NA | + | + | + | NA | 0,112559449215447 | 8 | -521,926502392848 | 1060,9780047857 | 13,0164318704051 | 5.1867508224667e-05 |
| **3813** | 24,5729683652003 | NA | NA | + | NA | NA | + | + | + | NA | + | + | + | 0,127288517191329 | 9 | -520,780618095407 | 1060,97855902546 | 13,0169861101699 | 5.18531366982966e-05 |
| **1563** | 24,8813161618084 | NA | + | NA | + | + | NA | NA | NA | NA | + | + | NA | 0,0977874680713999 | 7 | -523,056789684835 | 1060,98179642393 | 13,0202235086435 | 5.17692699613204e-05 |
| **858** | 25,5679568590132 | + | NA | NA | + | + | NA | + | NA | + | + | NA | NA | 0,11251934412614 | 8 | -521,929596445887 | 1060,98419289177 | 13,0226199764827 | 5.17072754154621e-05 |
| **3762** | 24,4891892732957 | + | NA | NA | NA | + | + | NA | + | NA | + | + | + | 0,127231740400524 | 9 | -520,785072193504 | 1060,98746722165 | 13,0258943063632 | 5.1622691334935e-05 |
| **2755** | 24,2278568628528 | NA | + | NA | NA | NA | NA | + | + | NA | + | NA | + | 0,0977493612422351 | 7 | -523,059681467336 | 1060,98757998893 | 13,0260070736442 | 5.16197807417195e-05 |
| **289** | 26,1945672410541 | NA | NA | NA | NA | NA | + | NA | NA | + | NA | NA | NA | 0,0534170806611641 | 4 | -526,343818743911 | 1060,99066779085 | 13,0290948755619 | 5.15401464023222e-05 |
| **3825** | 24,4327067648517 | NA | NA | NA | NA | + | + | + | + | NA | + | + | + | 0,127202823780316 | 9 | -520,787340570096 | 1060,99200397484 | 13,0304310595466 | 5.15057243430583e-05 |
| **2971** | 24,9997425026794 | NA | + | NA | + | + | NA | NA | + | + | + | NA | + | 0,127196440671721 | 9 | -520,787841285648 | 1060,99300540594 | 13,0314324906515 | 5.14799410814183e-05 |
| **2844** | 24,9921485585296 | + | + | NA | + | + | NA | NA | NA | + | + | NA | + | 0,127193403305892 | 9 | -520,788079547004 | 1060,99348192865 | 13,0319090133623 | 5.14676768619814e-05 |
| **2900** | 24,9710900926258 | + | + | NA | NA | + | NA | + | NA | + | + | NA | + | 0,127178362110259 | 9 | -520,789259417577 | 1060,9958416698 | 13,0342687545092 | 5.14069874744211e-05 |

| | | | | | | | | | | | | | | | | | | | |
|---|---|---|---|---|---|---|---|---|---|---|---|---|---|---|---|---|---|---|---|
| **1785** | 24,1979503256352 | NA | NA | NA | + | + | + | + | + | NA | + | + | NA | 0,1270968078181 | 9 | -520,795656394934 | 1061,00863562451 | 13,0470627092225 | 5.10791877212058e-05 |
| **2802** | 24,4202657444282 | + | NA | NA | NA | + | + | + | + | NA | + | NA | + | 0,127046370623498 | 9 | -520,799612302127 | 1061,0165474389 | 13,0549745236081 | 5.08775223428498e-05 |
| **2580** | 24,6726764608098 | + | + | NA | NA | + | NA | NA | NA | NA | + | NA | + | 0,0975566010196064 | 7 | -523,074307438027 | 1061,01683193032 | 13,0552590150266 | 5.08702857482963e-05 |
| **3651** | 24,0873385755759 | NA | + | NA | NA | NA | NA | + | NA | NA | + | + | + | 0,0975055914825504 | 7 | -523,078177340585 | 1061,02457173543 | 13,0629988201431 | 5.06738031288499e-05 |
| **1931** | 25,3368762939492 | NA | + | NA | + | NA | NA | NA | + | + | + | + | NA | 0,112248271318955 | 8 | -521,950505680075 | 1061,02601136015 | 13,0644384448588 | 5.06373406237948e-05 |
| **861** | 25,7162060811368 | NA | NA | + | + | + | NA | + | NA | + | + | NA | NA | 0,112219788924475 | 8 | -521,952702302566 | 1061,03040460513 | 13,0688316898409 | 5.05262315794827e-05 |
| **2707** | 24,6305840006088 | NA | + | NA | NA | + | NA | NA | + | NA | + | NA | + | 0,0973601522258811 | 7 | -523,089210072156 | 1061,04663719858 | 13,0850642832852 | 5.01178053878733e-05 |
| **798** | 25,6956029689016 | + | NA | + | + | + | NA | NA | NA | + | + | NA | NA | 0,112048462986821 | 8 | -521,965913835181 | 1061,05682767036 | 13,0952547550719 | 4.98630928083622e-05 |
| **4028** | 24,5958694661611 | + | + | NA | + | + | + | NA | + | + | + | + | + | 0,170865431209457 | 12 | -517,273670819582 | 1061,06347067142 | 13,1018977561312 | 4.9697747268851e-05 |
| **985** | 25,7068691352444 | NA | NA | NA | + | + | NA | + | + | + | + | NA | NA | 0,111987392847693 | 8 | -521,970622547807 | 1061,06624509561 | 13,1046721803232 | 4.96288537487057e-05 |
| **1821** | 25,7301369237262 | NA | NA | + | + | + | NA | NA | NA | + | + | + | NA | 0,111957409191187 | 8 | -521,972934269858 | 1061,07086853972 | 13,1092956244258 | 4.95142581407433e-05 |
| **922** | 25,6785024333581 | + | NA | NA | + | + | NA | NA | + | + | + | NA | NA | 0,111884089451858 | 8 | -521,978586849458 | 1061,08217369892 | 13,1206007836261 | 4.92351643981854e-05 |
| **904** | 25,5543942003926 | + | + | + | NA | NA | NA | NA | + | + | + | NA | NA | 0,111882560903357 | 8 | -521,978704687823 | 1061,08240937565 | 13,1208364603549 | 4.92293629487642e-05 |
| **1778** | 24,4401126355674 | + | NA | NA | NA | + | + | + | + | NA | + | + | NA | 0,126615637805705 | 9 | -520,83338638187 | 1061,08409559839 | 13,122522683095 | 4.91878746052268e-05 |
| **1867** | 24,9004618460192 | NA | + | NA | + | NA | NA | + | NA | + | + | + | NA | 0,111709358281865 | 8 | -521,99205585535 | 1061,1091117107 | 13,1475387954094 | 4.85764616724094e-05 |
| **1945** | 25,7014659503456 | NA | NA | NA | + | + | NA | NA | + | + | + | + | NA | 0,111628720663462 | 8 | -521,998270845607 | 1061,12154169121 | 13,1599687759242 | 4.82754956555702e-05 |
| **3748** | 24,159319441256 | + | + | NA | NA | NA | + | NA | + | NA | + | + | + | 0,126174028423474 | 9 | -520,867996021325 | 1061,1533148773 | 13,1917419620045 | 4.75146223471918e-05 |
| **1764** | 24,1443606940114 | + | + | NA | NA | NA | + | + | + | NA | + | + | NA | 0,126118626061245 | 9 | -520,872336759556 | 1061,16199635376 | 13,2004234384667 | 4.73088207979014e-05 |
| **1874** | 25,590520478902 | + | NA | NA | NA | + | NA | + | NA | + | + | + | NA | 0,11135755823891 | 8 | -522,019166035693 | 1061,16333207139 | 13,2017591560955 | 4.72772357332894e-05 |
| **3630** | 24,2216113168957 | + | NA | + | + | NA | + | NA | NA | NA | + | + | + | 0,126085819994407 | 9 | -520,874906963416 | 1061,16713676148 | 13,205563846187 | 4.71873836100821e-05 |
| **3755** | 23,9801089234545 | NA | + | NA | + | NA | + | NA | + | NA | + | + | + | 0,126043400388803 | 9 | -520,878230200688 | 1061,17378323602 | 13,2122103207307 | 4.70308290162202e-05 |
| **3715** | 24,6462552818684 | NA | + | NA | NA | NA | NA | NA | + | NA | + | + | + | 0,0964980883114456 | 7 | -523,154568063217 | 1061,1773531807 | 13,2157802654071 | 4.69469551659037e-05 |
| **1607** | 24,6254433492413 | NA | + | + | NA | NA | NA | + | NA | NA | + | + | NA | 0,0964802294825395 | 7 | -523,155921383979 | 1061,18005982222 | 13,2184869069306 | 4.68834638485272e-05 |
| **925** | 25,8759905665844 | NA | NA | + | + | + | NA | NA | + | + | + | NA | NA | 0,111212078078825 | 8 | -522,030373795679 | 1061,18574759136 | 13,2241746760674 | 4.67503220987933e-05 |
| **1771** | 23,9560596278603 | NA | + | NA | + | NA | + | + | + | NA | + | + | NA | 0,125934889153482 | 9 | -520,886730455608 | 1061,19078374586 | 13,2292108305703 | 4.66327492670788e-05 |
| **1860** | 24,9439999759574 | + | + | NA | NA | NA | NA | + | NA | + | + | + | NA | 0,111171406268108 | 8 | -522,033506815187 | 1061,19201363037 | 13,2304407150841 | 4.66040816343717e-05 |
| **4016** | 24,7105719014386 | + | + | + | + | NA | + | NA | + | + | + | + | + | 0,1700841284147 | 12 | -517,3381548652 | 1061,19243876266 | 13,2308658473671 | 4.65941762373738e-05 |
| **3015** | 25,0180570730783 | NA | + | + | NA | NA | NA | + | + | + | + | NA | + | 0,125834338856465 | 9 | -520,894606146966 | 1061,20653512858 | 13,2449622132874 | 4.62669265708646e-05 |
| **844** | 24,7857279507293 | + | + | NA | + | NA | NA | + | NA | + | + | NA | NA | 0,111037855082496 | 8 | -522,043793484958 | 1061,21258696992 | 13,2510140546244 | 4.61271381225239e-05 |
| **540** | 24,7981403315593 | + | + | NA | + | + | NA | NA | NA | NA | + | NA | NA | 0,0962298388527468 | 7 | -523,174892872559 | 1061,21800279938 | 13,2564298840919 | 4.60023987336163e-05 |
| **2790** | 24,4759917471814 | + | NA | + | NA | NA | + | + | + | NA | + | NA | + | 0,125717959647101 | 9 | -520,903720521557 | 1061,22476387776 | 13,2631909624699 | 4.58471483847025e-05 |
| **1559** | 25,3075510898273 | NA | + | + | NA | + | NA | NA | NA | NA | + | + | NA | 0,0961729034400284 | 7 | -523,179205997092 | 1061,22662904845 | 13,2650561331575 | 4.58044119368548e-05 |
| **2959** | 25,027783924687 | NA | + | + | + | NA | NA | NA | + | + | + | NA | + | 0,125674871808269 | 9 | -520,907094688716 | 1061,23151221208 | 13,269939296787 | 4.56927131340061e-05 |
| **1731** | 24,7319845834907 | NA | + | NA | NA | NA | NA | + | + | NA | + | + | NA | 0,0961155351538997 | 7 | -523,183551639151 | 1061,23532033257 | 13,2737474172752 | 4.56057942311562e-05 |
| **667** | 24,7687687680669 | NA | + | NA | + | + | NA | NA | + | NA | + | NA | NA | 0,096097051932728 | 7 | -523,184951682628 | 1061,23812041952 | 13,2765475042286 | 4.55419888120467e-05 |
| **3693** | 24,1309238841213 | NA | NA | + | + | NA | + | + | NA | NA | + | + | + | 0,125612338221046 | 9 | -520,911991338077 | 1061,2413055108 | 13,2797325955091 | 4.5469518837502e-05 |
| **3952** | 24,5677266597205 | + | + | + | + | NA | + | + | NA | + | + | + | + | 0,169717752337828 | 12 | -517,368372454339 | 1061,25287394094 | 13,2913010256464 | 4.52072725371478e-05 |
| **1600** | 24,252048152838 | + | + | + | + | + | + | NA | NA | NA | + | + | NA | 0,140180606631363 | 10 | -519,761669057244 | 1061,26936986052 | 13,3077969452283 | 4.48359382512482e-05 |
| **2757** | 24,8816534572222 | NA | NA | + | NA | NA | NA | + | + | NA | + | NA | + | 0,0958605654796619 | 7 | -523,202862232246 | 1061,27394151876 | 13,3123686034648 | 4.47335680030704e-05 |
| **863** | 25,2288833765612 | NA | + | + | + | + | NA | + | NA | + | + | NA | NA | 0,12540389811766 | 9 | -520,928310568987 | 1061,27394397262 | 13,3123710573295 | 4.47335131180427e-05 |
| **1800** | 25,3897319036632 | + | + | + | NA | NA | NA | NA | NA | + | + | + | NA | 0,110625406644811 | 8 | -522,075552236684 | 1061,27610447337 | 13,3145315580778 | 4.46852158150479e-05 |
| **3686** | 24,4115954004 | + | NA | + | NA | NA | + | + | NA | NA | + | + | + | 0,125300491034297 | 9 | -520,936405092797 | 1061,29013302024 | 13,3285601049492 | 4.43728781841438e-05 |
| **978** | 25,7248356259654 | + | NA | NA | NA | + | NA | + | + | + | + | NA | NA | 0,110511054500233 | 8 | -522,084354806901 | 1061,2937096138 | 13,332136698511 | 4.42935972190041e-05 |
| **3863** | 25,3280224264276 | NA | + | + | NA | + | NA | NA | NA | + | + | + | + | 0,125273690060035 | 9 | -520,938502869645 | 1061,29432857394 | 13,3327556586448 | 4.42798913545287e-05 |
| **4086** | 25,124330806893 | + | NA | + | NA | + | + | + | + | + | + | + | + | 0,169448872277446 | 12 | -517,390540389309 | 1061,29720981088 | 13,3356368955858 | 4.42161468519502e-05 |
| **854** | 25,7287179828528 | + | NA | + | NA | + | NA | + | NA | + | + | NA | NA | 0,110474930052911 | 8 | -522,087135350294 | 1061,29927070059 | 13,337697785297 | 4.41706080175511e-05 |
| **973** | 25,7159597172278 | NA | NA | + | + | NA | NA | + | + | + | + | NA | NA | 0,110469289120472 | 8 | -522,087569529653 | 1061,30013905931 | 13,338566144015 | 4.41514342140119e-05 |
| **2649** | 24,5957017326155 | NA | NA | NA | + | + | NA | + | NA | NA | + | NA | + | 0,0955743557897911 | 7 | -523,224532361716 | 1061,3172817777 | 13,3557088624048 | 4.37746136516147e-05 |
| **3595** | 24,196857291796 | NA | + | NA | + | NA | NA | NA | NA | NA | + | + | + | 0,095571858953498 | 7 | -523,224721377446 | 1061,31765980916 | 13,3560868938657 | 4.37663403429561e-05 |
| **3811** | 24,0879305556779 | NA | + | NA | NA | NA | + | + | + | NA | + | + | + | 0,125122986814603 | 9 | -520,950297579754 | 1061,31791799415 | 13,356345078862 | 4.37606908014119e-05 |
| **2608** | 23,7509230828694 | + | + | + | + | NA | + | NA | NA | NA | + | NA | + | 0,125107591104711 | 9 | -520,951502402551 | 1061,32032763975 | 13,3587547244572 | 4.37079986722087e-05 |
| **1927** | 25,7709962318268 | NA | + | + | NA | NA | NA | NA | + | + | + | + | NA | 0,110287943516845 | 8 | -522,101526131408 | 1061,32805226282 | 13,3664793475255 | 4.35395103514467e-05 |
| **2631** | 23,9988595286296 | NA | + | + | NA | NA | NA | + | NA | NA | + | NA | + | 0,0954957977735901 | 7 | -523,230479117943 | 1061,32917529015 | 13,3676023748599 | 4.35150691839878e-05 |
| **1804** | 25,0316657294879 | + | + | NA | + | NA | NA | NA | NA | + | + | + | NA | 0,110260295233244 | 8 | -522,103653730732 | 1061,33230746146 | 13,3707345461742 | 4.34469741934806e-05 |
| **2769** | 24,8182091651673 | NA | NA | NA | NA | + | NA | + | + | NA | + | NA | + | 0,0954684897697509 | 7 | -523,232546183163 | 1061,33330942059 | 13,371736505298 | 4.34252135986435e-05 |
| **624** | 23,7300074031998 | + | + | + | + | NA | + | + | NA | NA | + | NA | NA | 0,124994886870471 | 9 | -520,960321657823 | 1061,33796615029 | 13,3763932350014 | 4.33242214765032e-05 |
| **2840** | 25,2693495601466 | + | + | + | NA | + | NA | NA | NA | + | + | NA | + | 0,12496418691292 | 9 | -520,962723773314 | 1061,34277038127 | 13,3811974659832 | 4.32202765867599e-05 |
| **2645** | 24,8134462295056 | NA | NA | + | NA | + | NA | + | NA | NA | + | NA | + | 0,0953588407802951 | 7 | -523,240845378681 | 1061,34990781163 | 13,3883348963348 | 4.30663106235434e-05 |
| **2967** | 25,290000778763 | NA | + | + | NA | + | NA | NA | + | + | + | NA | + | 0,124907349790893 | 9 | -520,967170766624 | 1061,35166436789 | 13,3900914526039 | 4.30285030298082e-05 |
| **1924** | 25,4180254952891 | + | + | NA | NA | NA | NA | NA | + | + | + | + | NA | 0,110110423951325 | 8 | -522,11518552243 | 1061,35537104486 | 13,3937981295703 | 4.2948830502154e-05 |
| **4078** | 24,9178429165661 | + | NA | + | + | NA | + | + | + | + | + | + | + | 0,169074236452014 | 12 | -517,421415448398 | 1061,35895992905 | 13,3973870137629 | 4.28718304194709e-05 |
| **1770** | 24,2038853353757 | + | NA | NA | + | NA | + | + | + | NA | + | + | NA | 0,124788886088966 | 9 | -520,976438557391 | 1061,37019994943 | 13,4086270341377 | 4.26315660721871e-05 |
| **2695** | 24,6992491923593 | NA | + | + | NA | NA | NA | NA | + | NA | + | NA | + | 0,0952083699426076 | 7 | -523,252232692628 | 1061,37268243952 | 13,4111095242283 | 4.257868267944e-05 |
| **1877** | 25,8014588593451 | NA | NA | + | NA | + | NA | + | NA | + | + | + | NA | 0,109860545388932 | 8 | -522,134408022565 | 1061,39381604513 | 13,4322431298399 | 4.2131130895919e-05 |
| **3665** | 24,8123864386628 | NA | NA | NA | NA | + | NA | + | NA | NA | + | + | + | 0,0950316567861723 | 7 | -523,265603553054 | 1061,39942416037 | 13,437851245081 | 4.20131582556074e-05 |
| **2589** | 24,7512361038337 | NA | NA | + | + | + | NA | NA | NA | NA | + | NA | + | 0,094999654211987 | 7 | -523,268024723274 | 1061,40426650081 | 13,4426935855208 | 4.1911560290609e-05 |
| **2001** | 25,7822010678587 | NA | NA | NA | NA | + | NA | + | + | + | + | + | NA | 0,109727769843917 | 8 | -522,144619901555 | 1061,41423980311 | 13,4526668878193 | 4.17030821960974e-05 |
| **1683** | 25,1157176541551 | NA | + | NA | NA | + | NA | NA | + | NA | + | + | NA | 0,0948736459861104 | 7 | -523,27755710475 | 1061,42333126376 | 13,4617583484726 | 4.1513941447757e-05 |
| **3588** | 24,4457161718448 | + | + | NA | NA | NA | NA | NA | NA | NA | + | + | + | 0,0948326378922009 | 7 | -523,280659035153 | 1061,42953512457 | 13,4679622092799 | 4.13853676072515e-05 |
| **910** | 25,6739023649667 | + | NA | + | + | NA | NA | NA | + | + | + | NA | NA | 0,109621972615365 | 8 | -522,152755765838 | 1061,43051153168 | 13,4689386163845 | 4.13651680549193e-05 |
| **3609** | 24,7648346327438 | NA | NA | NA | + | + | NA | NA | NA | NA | + | + | + | 0,0948253958045201 | 7 | -523,281206825879 | 1061,43063070602 | 13,4690577907309 | 4.13627032949226e-05 |
| **2701** | 24,7786266411118 | NA | NA | + | + | NA | NA | NA | + | NA | + | NA | + | 0,094810615874538 | 7 | -523,282324764585 | 1061,43286658343 | 13,4712936681422 | 4.13164881655932e-05 |
| **1556** | 25,1306852737062 | + | + | NA | NA | + | NA | NA | NA | NA | + | + | NA | 0,0947946691552895 | 7 | -523,283530937568 | 1061,4352789294 | 13,4737060141099 | 4.12666833764132e-05 |
| **1644** | 23,8157538976266 | + | + | NA | + | NA | + | + | NA | NA | + | + | NA | 0,1243563953348 | 9 | -521,010263038268 | 1061,43784891118 | 13,4762759958912 | 4.12136901194046e-05 |
| **3860** | 25,2151380819442 | + | + | NA | NA | + | NA | NA | NA | + | + | + | + | 0,124349192677081 | 9 | -521,010826206416 | 1061,43897524748 | 13,4774023321875 | 4.11904864162695e-05 |
| **3754** | 24,2249897525402 | + | NA | NA | + | NA | + | NA | + | NA | + | + | + | 0,124322521158596 | 9 | -521,01291158377 | 1061,44314600218 | 13,4815730868943 | 4.11046782110618e-05 |

| | | | | | | | | | | | | | | | | | | |
|---|---|---|---|---|---|---|---|---|---|---|---|---|---|---|---|---|---|---|
| **3987** | 25,2195403211462 | NA | + | NA | NA | + | NA | NA | + | + | + | + | + | 0,124308466869426 | 9 | -521,014010426911 | 1061,44534368847 | 13,4837707731772 | 4.10595354242913e-05 |
| **970** | 25,4307444402623 | + | NA | NA | + | NA | NA | + | + | + | + | NA | NA | 0,109517411646295 | 8 | -522,160795611768 | 1061,44659122354 | 13,4850183082447 | 4.10339318053265e-05 |
| **1814** | 25,7792367435924 | + | NA | + | NA | + | NA | NA | NA | + | + | + | NA | 0,109493739224439 | 8 | -522,162615687932 | 1061,45023137586 | 13,4886584605738 | 4.09593148489797e-05 |
| **1938** | 25,7550759874264 | + | NA | NA | NA | + | NA | NA | + | + | + | + | NA | 0,109471583629531 | 8 | -522,164319097636 | 1061,45363819527 | 13,4920652799817 | 4.08896037447401e-05 |
| **2642** | 24,769787233443 | + | NA | NA | NA | + | NA | + | NA | NA | + | NA | + | 0,0946152433974887 | 7 | -523,297100823481 | 1061,46241870123 | 13,5008457859349 | 4.07104815238253e-05 |
| **2713** | 24,7390972755304 | NA | NA | NA | + | + | NA | NA | + | NA | + | NA | + | 0,0946003257099201 | 7 | -523,298228920163 | 1061,46467489459 | 13,5031019792998 | 4.06645820590614e-05 |
| **536** | 25,1164243377091 | + | + | + | NA | + | NA | NA | NA | NA | + | NA | NA | 0,0945623304338082 | 7 | -523,301102092892 | 1061,47042124005 | 13,5088483247569 | 4.05479133757714e-05 |
| **663** | 25,1183308905584 | NA | + | + | NA | + | NA | NA | + | NA | + | NA | NA | 0,0945589414896573 | 7 | -523,301358356314 | 1061,47093376689 | 13,509360851602 | 4.05375237600082e-05 |
| **2964** | 25,1760445169465 | + | + | NA | NA | + | NA | NA | + | + | + | NA | + | 0,124137055534767 | 9 | -521,027410907972 | 1061,47214465059 | 13,5105717352988 | 4.05129880749085e-05 |
| **2761** | 24,4871669840723 | NA | NA | NA | + | NA | NA | + | + | NA | + | NA | + | 0,094480986941717 | 7 | -523,307252818278 | 1061,48272269082 | 13,5211497755288 | 4.02992797181446e-05 |
| **3777** | 24,7968060427187 | NA | NA | NA | NA | NA | NA | + | + | NA | + | + | + | 0,0944314403236908 | 7 | -523,310998976936 | 1061,49021500814 | 13,5286420928446 | 4.01485946438409e-05 |
| **3817** | 24,1577626712411 | NA | NA | NA | + | NA | + | + | + | NA | + | + | + | 0,1239304011552 | 9 | -521,043563111677 | 1061,504449058 | 13,5428761427088 | 3.98638704910403e-05 |
| **2664** | 23,9113942293946 | + | + | + | NA | NA | + | + | NA | NA | + | NA | + | 0,12387332609923 | 9 | -521,048023453823 | 1061,51336974229 | 13,5517968270008 | 3.96864599393938e-05 |
| **2586** | 24,7266700975331 | + | NA | NA | + | + | NA | NA | NA | NA | + | NA | + | 0,0942257782551539 | 7 | -523,326546642414 | 1061,52131033909 | 13,5597374238009 | 3.95292052316197e-05 |
| **3019** | 24,7274628074286 | NA | + | NA | + | NA | NA | + | + | + | + | NA | + | 0,123803063373245 | 9 | -521,053513996587 | 1061,52435082782 | 13,5627779125284 | 3.94691568358456e-05 |
| **987** | 25,1984483807905 | NA | + | NA | + | + | NA | + | + | + | + | NA | NA | 0,123719246560395 | 9 | -521,060063121287 | 1061,53744907722 | 13,5758761619297 | 3.9211512997594e-05 |
| **3628** | 23,7765296455568 | + | + | NA | + | NA | + | NA | NA | NA | + | + | + | 0,123709794489446 | 9 | -521,06080163062 | 1061,53892609589 | 13,5773531805958 | 3.91825656195536e-05 |
| **715** | 24,4525909218335 | NA | + | NA | + | NA | NA | + | + | NA | + | NA | NA | 0,0940859717144685 | 7 | -523,337113739412 | 1061,54244453309 | 13,5808716177962 | 3.91136955178549e-05 |
| **2670** | 23,9691808396051 | + | NA | + | + | NA | + | + | NA | NA | + | NA | + | 0,123642840619556 | 9 | -521,066032643335 | 1061,54938812131 | 13,5878152060241 | 3.89781362730384e-05 |
| **1883** | 25,1776169728846 | NA | + | NA | + | + | NA | + | NA | + | + | + | NA | 0,12362334025896 | 9 | -521,067556104268 | 1061,55243504318 | 13,5908621278918 | 3.89187998150156e-05 |
| **860** | 25,1644873325398 | + | + | NA | + | + | NA | + | NA | + | + | NA | NA | 0,123622094574546 | 9 | -521,067653421909 | 1061,55262967846 | 13,5910567631736 | 3.89150125135199e-05 |
| **4072** | 24,8883148216984 | + | + | + | NA | NA | + | + | + | + | + | + | + | 0,167810099356202 | 12 | -517,525494857898 | 1061,56711874805 | 13,6055458327637 | 3.86341100835849e-05 |
| **981** | 25,9673320856077 | NA | NA | + | NA | + | NA | + | + | + | + | NA | NA | 0,108706290021577 | 8 | -522,223131908363 | 1061,57126381673 | 13,6096909014361 | 3.85541224810458e-05 |
| **2699** | 24,2965997918233 | NA | + | NA | + | NA | NA | NA | + | NA | + | NA | + | 0,0938911660828629 | 7 | -523,35183515408 | 1061,57188736242 | 13,6103144471324 | 3.85421042260527e-05 |
| **3855** | 24,8106201842398 | NA | + | + | + | NA | NA | NA | NA | + | + | + | + | 0,123484870766832 | 9 | -521,078373025302 | 1061,57406888525 | 13,6124959699587 | 3.85000869055835e-05 |
| **3684** | 23,9329462556 | + | + | NA | NA | NA | + | + | NA | NA | + | + | + | 0,123418808103395 | 9 | -521,083533088157 | 1061,58438901096 | 13,622816095668 | 3.83019357132131e-05 |
| **918** | 25,8776309135253 | + | NA | + | NA | + | NA | NA | + | + | + | NA | NA | 0,108569646224751 | 8 | -522,233627672747 | 1061,59225534549 | 13,6306824302026 | 3.81515836670915e-05 |
| **3721** | 24,7309341984079 | NA | NA | NA | + | NA | NA | NA | + | NA | + | + | + | 0,0937125721753497 | 7 | -523,365328673411 | 1061,59887440109 | 13,6373014857952 | 3.80255286468248e-05 |
| **983** | 25,4195014898504 | NA | + | + | NA | + | NA | + | + | + | + | NA | NA | 0,123320692399274 | 9 | -521,091196052206 | 1061,59971493906 | 13,638142023766 | 3.80095510541593e-05 |
| **966** | 25,6988400508356 | + | NA | + | NA | NA | NA | + | + | + | + | NA | NA | 0,108518298563823 | 8 | -522,237571328914 | 1061,60014265783 | 13,638569742538 | 3.80014232240458e-05 |
| **856** | 25,3425204287026 | + | + | + | NA | + | NA | + | NA | + | + | NA | NA | 0,123288971393243 | 9 | -521,093673320652 | 1061,60466947595 | 13,6430965606594 | 3.79155077260392e-05 |
| **1712** | 24,3444187512111 | + | + | + | + | NA | + | NA | + | NA | + | + | NA | 0,138070354722798 | 10 | -519,929497448386 | 1061,6050266428 | 13,6434537275138 | 3.79087372492904e-05 |
| **1879** | 25,3909255020882 | NA | + | + | NA | + | NA | + | NA | + | + | + | NA | 0,123227938038948 | 9 | -521,098439500161 | 1061,61420183497 | 13,6526289196779 | 3.77352255798486e-05 |
| **3911** | 24,8053699616124 | NA | + | + | NA | NA | NA | + | NA | + | + | + | + | 0,123222281875562 | 9 | -521,098881181036 | 1061,61508519672 | 13,6535122814262 | 3.77185623326186e-05 |
| **1933** | 25,7654638278534 | NA | NA | + | + | NA | NA | NA | + | + | + | + | NA | 0,108397161690952 | 8 | -522,246874108798 | 1061,6187482176 | 13,6571753023047 | 3.76495436154957e-05 |
| **1993** | 25,4864753766558 | NA | NA | NA | + | NA | NA | + | + | + | + | + | NA | 0,108396980777918 | 8 | -522,24688800118 | 1061,61877600236 | 13,6572030870684 | 3.76490205772926e-05 |
| **4035** | 24,8926744971242 | NA | + | NA | NA | NA | NA | + | + | + | + | + | + | 0,123093499210614 | 9 | -521,108936845606 | 1061,63519652586 | 13,6736236105664 | 3.73411777282075e-05 |
| **3640** | 24,4786610794604 | + | + | + | NA | + | + | NA | NA | NA | + | + | + | 0,137850901676131 | 10 | -519,946926974441 | 1061,63988569491 | 13,6783127796241 | 3.7253730734111e-05 |
| **660** | 24,9692523431025 | + | + | NA | NA | + | NA | NA | + | NA | + | NA | NA | 0,0934195542095892 | 7 | -523,387461654185 | 1061,64314036263 | 13,6815674473437 | 3.71931557778901e-05 |
| **3979** | 24,9021894918229 | NA | + | NA | + | NA | NA | NA | + | + | + | + | + | 0,122949368867615 | 9 | -521,120189144072 | 1061,65770112279 | 13,6961282074994 | 3.69233587715972e-05 |
| **1930** | 25,4703144895471 | + | NA | NA | + | NA | NA | NA | + | + | + | + | NA | 0,108125477028186 | 8 | -522,26773370475 | 1061,6604674095 | 13,69889449421 | 3.6872323775752e-05 |
| **605** | 25,0226289375952 | NA | NA | + | + | + | NA | + | NA | NA | + | NA | NA | 0,0932988970736526 | 7 | -523,396573358645 | 1061,66136377155 | 13,6997908562623 | 3.6855802002494e-05 |
| **3810** | 24,3503109344817 | + | NA | NA | NA | NA | + | + | + | NA | + | + | + | 0,122849364050792 | 9 | -521,127995463632 | 1061,67331376191 | 13,7117408466197 | 3.66362453414091e-05 |
| **291** | 25,599478377895 | NA | + | NA | NA | NA | + | NA | NA | + | NA | NA | NA | 0,0635278668145052 | 5 | -525,608550746417 | 1061,67511676001 | 13,7135438447199 | 3.66032326837106e-05 |
| **708** | 24,5486254359183 | + | + | NA | NA | NA | NA | + | + | NA | + | NA | NA | 0,0931870003351229 | 7 | -523,405022418228 | 1061,67826189072 | 13,7166889754287 | 3.65457169436153e-05 |
| **729** | 25,0281809764845 | NA | NA | NA | + | + | NA | + | + | NA | + | NA | NA | 0,0931602514452904 | 7 | -523,407042009666 | 1061,6823010736 | 13,7207281583051 | 3.64719840068396e-05 |
| **2754** | 24,6407970745405 | + | NA | NA | NA | NA | NA | + | + | NA | + | NA | + | 0,0931308871932209 | 7 | -523,409258997308 | 1061,68673504888 | 13,7251621335886 | 3.63912156333506e-05 |
| **4030** | 24,9479846211093 | + | NA | + | + | + | + | NA | + | + | + | + | + | 0,167042975207742 | 12 | -517,588576997741 | 1061,69328302774 | 13,731710112449 | 3.627226600418e-05 |
| **2575** | 24,1086772890703 | NA | + | + | + | NA | NA | NA | NA | NA | + | NA | + | 0,0930399293036999 | 7 | -523,416125821168 | 1061,7004686966 | 13,7388957813089 | 3.61421795874419e-05 |
| **2732** | 23,8695910363886 | + | + | NA | + | NA | + | NA | + | NA | + | NA | + | 0,122670178936067 | 9 | -521,141980327936 | 1061,70128349052 | 13,7397105752273 | 3.61274583722722e-05 |
| **3605** | 25,087454364544 | NA | NA | + | NA | + | NA | NA | NA | NA | + | + | + | 0,0930060415441835 | 7 | -523,418683985926 | 1061,70558502612 | 13,744012110825 | 3.60498400975138e-05 |
| **655** | 24,9016925853368 | NA | + | + | + | NA | NA | NA | + | NA | + | NA | NA | 0,092945096211239 | 7 | -523,423284468775 | 1061,71478599181 | 13,7532130765228 | 3.5884374329486e-05 |
| **717** | 25,1092167549245 | NA | NA | + | + | NA | NA | + | + | NA | + | NA | NA | 0,0929397590800523 | 7 | -523,423687329546 | 1061,71559171335 | 13,7540187980642 | 3.58699208343564e-05 |
| **2624** | 24,1809141574675 | + | + | + | + | + | + | NA | NA | NA | + | NA | + | 0,137365464819872 | 10 | -519,985465871695 | 1061,71696348942 | 13,7553905741315 | 3.58453265203243e-05 |
| **35** | 24,9714207410656 | NA | + | NA | NA | NA | + | NA | NA | NA | NA | NA | NA | 0,0482679404356094 | 4 | -526,7152587775 | 1061,73354785803 | 13,7719749427392 | 3.55493194351652e-05 |
| **1986** | 25,4894559825174 | + | NA | NA | NA | NA | NA | + | + | + | + | + | NA | 0,107645354629908 | 8 | -522,304581355064 | 1061,73416271013 | 13,7725897948367 | 3.55383923280813e-05 |
| **1720** | 24,4659447651144 | + | + | + | NA | + | + | NA | + | NA | + | + | NA | 0,137251675497929 | 10 | -519,99449648519 | 1061,73502471641 | 13,7734518011209 | 3.55230784697194e-05 |
| **3597** | 24,6390160930275 | NA | NA | + | + | NA | NA | NA | NA | NA | + | + | + | 0,092800899164173 | 7 | -523,434168010424 | 1061,73655307511 | 13,7749801598209 | 3.54959428362713e-05 |
| **3653** | 24,6828217869401 | NA | NA | + | NA | NA | NA | + | NA | NA | + | + | + | 0,0927968566314783 | 7 | -523,434473103218 | 1061,7371632607 | 13,7755903454085 | 3.54851149317433e-05 |
| **2895** | 24,5793885455155 | NA | + | + | + | NA | NA | + | NA | + | + | NA | + | 0,12242427684152 | 9 | -521,16116760727 | 1061,73965804919 | 13,7780851338957 | 3.54408785994533e-05 |
| **1551** | 24,79134162774 | NA | + | + | + | NA | NA | NA | NA | NA | + | + | NA | 0,0927740825144424 | 7 | -523,436191856503 | 1061,74060076727 | 13,7790278519797 | 3.54241771573673e-05 |
| **748** | 23,8331045296463 | + | + | NA | + | NA | + | + | + | NA | + | NA | NA | 0,122392618130249 | 9 | -521,163637486681 | 1061,74459780801 | 13,7830248927169 | 3.53534519142371e-05 |
| **1656** | 24,3994668712583 | + | + | + | NA | + | + | + | NA | NA | + | + | NA | 0,137122535700966 | 10 | -520,00474390982 | 1061,75551956567 | 13,7939466503806 | 3.51609171799409e-05 |
| **3975** | 25,2867169105515 | NA | + | + | NA | NA | NA | NA | + | + | + | + | + | 0,12218445331369 | 9 | -521,17987541203 | 1061,7770736587 | 13,8155007434143 | 3.47840209021453e-05 |
| **2709** | 25,0670511138426 | NA | NA | + | NA | + | NA | NA | + | NA | + | NA | + | 0,0925119138229619 | 7 | -523,455974515184 | 1061,78016608463 | 13,818593169342 | 3.47302789570454e-05 |
| **3729** | 25,0499255235805 | NA | NA | NA | NA | + | NA | NA | + | NA | + | + | + | 0,0925022221716776 | 7 | -523,456705715858 | 1061,78162848598 | 13,8200555706892 | 3.47048934357634e-05 |
| **800** | 25,4434279589337 | + | + | + | + | + | NA | NA | NA | + | + | NA | NA | 0,122122286303259 | 9 | -521,184724011964 | 1061,78677085857 | 13,8251979432823 | 3.46157753080887e-05 |
| **3691** | 23,7435271135808 | NA | + | NA | + | NA | + | + | NA | NA | + | + | + | 0,122115722452637 | 9 | -521,18523592721 | 1061,78779468907 | 13,8262217737752 | 3.45980594998249e-05 |
| **3012** | 24,8280375122999 | + | + | NA | NA | NA | NA | + | + | + | + | NA | + | 0,122084915187447 | 9 | -521,187638537226 | 1061,7925999091 | 13,8310269938072 | 3.45150336348622e-05 |
| **1823** | 25,4887131057093 | NA | + | + | + | + | NA | NA | NA | + | + | + | NA | 0,12206710030019 | 9 | -521,189027853727 | 1061,7953785421 | 13,8338056268094 | 3.44671146241411e-05 |
| **927** | 25,4914258103758 | NA | + | + | + | + | NA | NA | + | + | + | NA | NA | 0,122055249693078 | 9 | -521,189952022696 | 1061,79722688004 | 13,8356539647468 | 3.44352759008107e-05 |
| **3602** | 24,9945142717153 | + | NA | NA | NA | + | NA | NA | NA | NA | + | + | + | 0,0923763669611643 | 7 | -523,466200335848 | 1061,80061772596 | 13,8390448106686 | 3.43769430068207e-05 |
| **1625** | 24,987804994736 | NA | NA | NA | + | + | NA | + | NA | NA | + | + | NA | 0,0923746022082511 | 7 | -523,466333460895 | 1061,80088397605 | 13,8393110607628 | 3.43723668792686e-05 |
| **980** | 25,2767635134587 | + | + | NA | NA | + | NA | + | + | + | + | NA | NA | 0,122029721507918 | 9 | -521,191942794507 | 1061,80120842366 | 13,8396355083692 | 3.43667913154467e-05 |
| **846** | 25,3075164788233 | + | NA | + | + | NA | NA | + | NA | + | + | NA | NA | 0,107192537338576 | 8 | -522,339315284648 | 1061,8036305693 | 13,8420576540052 | 3.4325195821312e-05 |
| **1876** | 25,2683909770441 | + | + | NA | NA | + | NA | + | NA | + | + | + | NA | 0,122012097613781 | 9 | -521,193317129958 | 1061,80395709456 | 13,8423841792717 | 3.43195922568947e-05 |

| | | | | | | | | | | | | | | | | | | |
|---|---|---|---|---|---|---|---|---|---|---|---|---|---|---|---|---|---|---|
| **3717** | 25,1378669102695 | NA | NA | + | NA | NA | NA | NA | + | NA | + | + | + | 0,0923524966249349 | 7 | -523,468000985068 | 1061,8042190244 | 13,8426461091087 | 3.43150978885996e-05 |
| **602** | 24,9544188178408 | + | NA | NA | + | + | NA | + | NA | NA | + | NA | NA | 0,092290459033372 | 7 | -523,472680544502 | 1061,81357814327 | 13,8520052279766 | 3.41548934839146e-05 |
| **725** | 25,3366625456887 | NA | NA | + | NA | + | NA | + | + | NA | + | NA | NA | 0,0922804849578235 | 7 | -523,473432869465 | 1061,81508279319 | 13,8535098779032 | 3.41292075682175e-05 |
| **2582** | 24,9939530021089 | + | NA | + | NA | + | NA | NA | NA | NA | + | NA | + | 0,0922443448364071 | 7 | -523,476158778759 | 1061,82053461178 | 13,8589616964903 | 3.40363011289187e-05 |
| **2671** | 23,66059573058 | NA | + | + | + | NA | + | + | NA | NA | + | NA | + | 0,121883405488754 | 9 | -521,203351882763 | 1061,82402660017 | 13,8624536848808 | 3.3976925794297e-05 |
| **2568** | 24,3972705407352 | + | + | + | NA | NA | NA | NA | NA | NA | + | NA | + | 0,092204624951551 | 7 | -523,479154570615 | 1061,82652619549 | 13,8649532802031 | 3.39344880367369e-05 |
| **704** | 24,1675009962169 | + | + | + | + | + | + | NA | + | NA | + | NA | NA | 0,136674117999241 | 10 | -520,040314586951 | 1061,82666091993 | 13,865088004643 | 3.39322022112794e-05 |
| **2956** | 24,8455095004052 | + | + | NA | + | NA | NA | NA | + | + | + | NA | + | 0,121833171713061 | 9 | -521,207268456456 | 1061,83185974756 | 13,8702868322675 | 3.3844112916037e-05 |
| **2003** | 25,3219226352857 | NA | + | NA | NA | + | NA | + | + | + | + | + | NA | 0,12181300201992 | 9 | -521,208840962673 | 1061,83500475999 | 13,8734318447005 | 3.37909346606127e-05 |
| **2794** | 24,0062965865392 | + | NA | NA | + | NA | + | + | + | NA | + | NA | + | 0,121800881243348 | 9 | -521,209785927286 | 1061,83689468922 | 13,8753217739261 | 3.37590185053182e-05 |
| **3690** | 24,0033194806066 | + | NA | NA | + | NA | + | + | NA | NA | + | + | + | 0,121800412712511 | 9 | -521,209822454804 | 1061,83696774425 | 13,875394828963 | 3.37577853946688e-05 |
| **3647** | 24,151713577212 | NA | + | + | + | + | + | NA | NA | NA | + | + | + | 0,136457652510708 | 10 | -520,057479076065 | 1061,86098989816 | 13,8994169828711 | 3.33547433663289e-05 |
| **2832** | 24,7296155582976 | + | + | + | + | NA | NA | NA | NA | + | + | NA | + | 0,121631696439476 | 9 | -521,222974616882 | 1061,86327206841 | 13,9016991531184 | 3.33167044718058e-05 |
| **1727** | 24,1880639796967 | NA | + | + | + | + | + | NA | + | NA | + | + | NA | 0,136437868865454 | 10 | -520,059047592771 | 1061,86412693157 | 13,9025540162822 | 3.33024669031198e-05 |
| **584** | 24,4067296628501 | + | + | + | NA | NA | NA | + | NA | NA | + | NA | NA | 0,0919222161640891 | 7 | -523,500450904772 | 1061,86911886381 | 13,9075459485171 | 3.32194487224857e-05 |
| **1941** | 26,0069770405967 | NA | NA | + | NA | + | NA | NA | + | + | + | + | NA | 0,106764913748351 | 8 | -522,372100537308 | 1061,86920107462 | 13,9076281593257 | 3.32180832516794e-05 |
| **591** | 24,2348762052097 | NA | + | + | + | NA | NA | + | NA | NA | + | NA | NA | 0,0919126795950671 | 7 | -523,501169938091 | 1061,87055693044 | 13,9089840151541 | 3.31955714173482e-05 |
| **2628** | 23,8981704621493 | + | + | NA | NA | NA | NA | + | NA | NA | + | NA | + | 0,0918692007691909 | 7 | -523,504448036563 | 1061,87711312739 | 13,9155402120984 | 3.30869312293067e-05 |
| **2706** | 24,9457787321674 | + | NA | NA | NA | + | NA | NA | + | NA | + | NA | + | 0,0917862511733819 | 7 | -523,510701609882 | 1061,88962027403 | 13,9280473587373 | 3.28806653007089e-05 |
| **161** | 26,0257008174899 | NA | NA | NA | NA | NA | + | NA | + | NA | NA | NA | NA | 0,047181330168077 | 4 | -526,793386139509 | 1061,88980258205 | 13,9282296667573 | 3.28776682328159e-05 |
| **1989** | 25,8164791942709 | NA | NA | + | NA | NA | NA | + | + | + | + | + | NA | 0,106601115096939 | 8 | -522,384654576433 | 1061,89430915287 | 13,9327362375745 | 3.28036688648744e-05 |
| **1869** | 25,3888117724778 | NA | NA | + | + | NA | NA | + | NA | + | + | + | NA | 0,106544716733047 | 8 | -522,388976590382 | 1061,90295318076 | 13,9413802654731 | 3.26621968925946e-05 |
| **2692** | 24,4882434679349 | + | + | NA | NA | NA | NA | NA | + | NA | + | NA | + | 0,0916800097185995 | 7 | -523,518710323587 | 1061,90563770144 | 13,9440647861475 | 3.26183851311526e-05 |
| **1806** | 25,3751039263304 | + | NA | + | + | NA | NA | NA | NA | + | + | + | NA | 0,106517815439662 | 8 | -522,391038039331 | 1061,90707607866 | 13,9455031633704 | 3.25949347936604e-05 |
| **1611** | 24,3530164166719 | NA | + | NA | + | NA | NA | + | NA | NA | + | + | NA | 0,091669474472315 | 7 | -523,519504442515 | 1061,90722593929 | 13,9456530240029 | 3.25924925363885e-05 |
| **2698** | 24,5673759590261 | + | NA | NA | + | NA | NA | NA | + | NA | + | NA | + | 0,0916607374654704 | 7 | -523,520163007939 | 1061,90854307014 | 13,9469701548512 | 3.25710353139852e-05 |
| **2888** | 24,7159658540265 | + | + | + | NA | NA | NA | + | NA | + | + | NA | + | 0,121328393209249 | 9 | -521,246612065011 | 1061,91054696467 | 13,9489740493777 | 3.25384171978357e-05 |
| **598** | 25,1548505391593 | + | NA | + | NA | + | NA | + | NA | NA | + | NA | NA | 0,0915760902331975 | 7 | -523,526543096386 | 1061,92130324703 | 13,9597303317444 | 3.23638907309264e-05 |
| **4076** | 24,5380779168254 | + | + | NA | + | NA | + | + | + | + | + | + | + | 0,165615262087286 | 12 | -517,705826233125 | 1061,92778149851 | 13,9662085832169 | 3.22592296162966e-05 |
| **1621** | 25,2356180719258 | NA | NA | + | NA | + | NA | + | NA | NA | + | + | NA | 0,0915329071752246 | 7 | -523,52979769015 | 1061,92781243456 | 13,9662395192734 | 3.2258730633482e-05 |
| **2788** | 23,9746344937366 | + | + | NA | NA | NA | + | + | + | NA | + | NA | + | 0,121215187456253 | 9 | -521,255432482879 | 1061,9281878004 | 13,9666148851134 | 3.22526767888406e-05 |
| **2952** | 25,1883308309347 | + | + | + | NA | NA | NA | NA | + | + | + | NA | + | 0,12120368748703 | 9 | -521,256328438585 | 1061,92997971182 | 13,9684067965254 | 3.22237927603742e-05 |
| **2795** | 23,8214659833656 | NA | + | NA | + | NA | + | + | + | NA | + | NA | + | 0,121178094529916 | 9 | -521,258322328468 | 1061,93396749158 | 13,9723945762903 | 3.21596060778688e-05 |
| **722** | 25,1298770400931 | + | NA | NA | NA | + | NA | + | + | NA | + | NA | NA | 0,0914791018674577 | 7 | -523,533852638653 | 1061,93592233157 | 13,9743494162794 | 3.21281879927044e-05 |
| **1604** | 24,4523196715974 | + | + | NA | NA | NA | NA | + | NA | NA | + | + | NA | 0,0914733819362475 | 7 | -523,534283697744 | 1061,93678444975 | 13,9752115344606 | 3.21143418296773e-05 |
| **640** | 24,0696879890147 | + | + | + | + | + | + | + | NA | NA | + | NA | NA | 0,135832663895959 | 10 | -520,10701301645 | 1061,96005777893 | 13,9984848636411 | 3.17428039287835e-05 |
| **3657** | 24,3800121047027 | NA | NA | NA | + | NA | NA | + | NA | NA | + | + | + | 0,0913093089731093 | 7 | -523,546647227093 | 1061,96151150845 | 13,9999385931587 | 3.17197395866063e-05 |
| **1671** | 25,32920167382 | NA | + | + | NA | NA | NA | NA | + | NA | + | + | NA | 0,0913032224673885 | 7 | -523,547105825805 | 1061,96242870587 | 14,000855790584 | 3.17051962898909e-05 |
| **1745** | 25,2247479701674 | NA | NA | NA | NA | + | NA | + | + | NA | + | + | NA | 0,0912920534947035 | 7 | -523,54794736414 | 1061,96411178254 | 14,0025388672525 | 3.16785263752691e-05 |
| **3767** | 24,3863490965587 | NA | + | + | NA | + | + | NA | + | NA | + | + | + | 0,135738971678221 | 10 | -520,114435576938 | 1061,97490289991 | 14,0133299846173 | 3.15080633125131e-05 |
| **2637** | 24,2925193658803 | NA | NA | + | + | NA | NA | + | NA | NA | + | NA | + | 0,0911924791086775 | 7 | -523,555449447105 | 1061,97911594847 | 14,0175430331822 | 3.14417606706326e-05 |
| **669** | 25,2637558763963 | NA | NA | + | + | + | NA | NA | + | NA | + | NA | NA | 0,0911689064366453 | 7 | -523,557225327153 | 1061,98266770857 | 14,021094793279 | 3.13859734255604e-05 |
| **924** | 25,3540997693399 | + | + | NA | + | + | NA | NA | + | + | + | NA | NA | 0,120844938123032 | 9 | -521,284272503057 | 1061,98586784076 | 14,0242949254687 | 3.13357939495732e-05 |
| **1820** | 25,3649458310883 | + | + | NA | + | + | NA | NA | NA | + | + | + | NA | 0,120837565710804 | 9 | -521,284846642845 | 1061,98701612034 | 14,0254432050456 | 3.13178079871863e-05 |
| **1675** | 24,8639284405625 | NA | + | NA | + | NA | NA | NA | + | NA | + | + | NA | 0,0911114277216222 | 7 | -523,56155537314 | 1061,99132780054 | 14,0297548852525 | 3.1250364525556e-05 |
| **975** | 25,261612163777 | NA | + | + | + | NA | NA | + | + | + | + | NA | NA | 0,120754556032556 | 9 | -521,291310837902 | 1061,99994451045 | 14,0383715951589 | 3.11160164801097e-05 |
| **3915** | 24,5532794116579 | NA | + | NA | + | NA | NA | + | NA | + | + | + | + | 0,120749805955701 | 9 | -521,291680721177 | 1062,000684277 | 14,0391113617095 | 3.11045093143047e-05 |
| **3848** | 25,0460013876731 | + | + | + | NA | NA | NA | NA | NA | + | + | + | + | 0,120745126487726 | 9 | -521,292045104264 | 1062,00141304317 | 14,0398401278821 | 3.10931774219026e-05 |
| **1618** | 25,0966526686653 | + | NA | NA | NA | + | NA | + | NA | NA | + | + | NA | 0,0909684944717139 | 7 | -523,57232178204 | 1062,01286061834 | 14,0512877030537 | 3.09157150419537e-05 |
| **1947** | 25,3930991385816 | NA | + | NA | + | + | NA | NA | + | + | + | + | NA | 0,120669941526289 | 9 | -521,297899377634 | 1062,01312158991 | 14,0515486746237 | 3.09116812437898e-05 |
| **2635** | 23,7788538418748 | NA | + | NA | + | NA | NA | + | NA | NA | + | NA | + | 0,0909202779624601 | 7 | -523,575953296063 | 1062,02012364639 | 14,0585507310984 | 3.08036477993105e-05 |
| **1926** | 25,7532769801645 | + | NA | + | NA | NA | NA | NA | + | + | + | + | NA | 0,105753481936708 | 8 | -522,449583089399 | 1062,0241661788 | 14,0625932635078 | 3.07414483090731e-05 |
| **1565** | 25,2295417244234 | NA | NA | + | + | + | NA | NA | NA | NA | + | + | NA | 0,0908642541318409 | 7 | -523,580172590888 | 1062,02856223604 | 14,0669893207491 | 3.0673951932507e-05 |
| **1866** | 25,1623804843783 | + | NA | NA | + | NA | NA | + | NA | + | + | + | NA | 0,105712123986015 | 8 | -522,452749525164 | 1062,03049905033 | 14,0689261350374 | 3.06442614368818e-05 |
| **2630** | 24,50101337165 | + | NA | + | NA | NA | NA | + | NA | NA | + | NA | + | 0,0908190656996063 | 7 | -523,583575655413 | 1062,03536836509 | 14,0737954497993 | 3.05697439087247e-05 |
| **3703** | 24,3118983114766 | NA | + | + | NA | + | + | + | NA | NA | + | + | + | 0,13535413369968 | 10 | -520,144915088071 | 1062,03586192217 | 14,074289006883 | 3.0562200882665e-05 |
| **2694** | 24,961558085795 | + | NA | + | NA | NA | NA | NA | + | NA | + | NA | + | 0,0908122034571418 | 7 | -523,584092424519 | 1062,0364019033 | 14,0748289880119 | 3.0553950490612e-05 |
| **3650** | 24,5364161256216 | + | NA | NA | NA | NA | NA | + | NA | NA | + | + | + | 0,0908039514124582 | 7 | -523,584713849127 | 1062,03764475252 | 14,0760718372262 | 3.05349694121961e-05 |
| **3714** | 24,9199392244686 | + | NA | NA | NA | NA | NA | NA | + | NA | + | + | + | 0,0907433339709614 | 7 | -523,589278504952 | 1062,04677406417 | 14,0852011488776 | 3.03959054172816e-05 |
| **1862** | 25,3918638791872 | + | NA | + | NA | NA | NA | + | NA | + | + | + | NA | 0,10560074130963 | 8 | -522,461276446408 | 1062,04755289282 | 14,0859799775249 | 3.03840711207036e-05 |
| **97** | 25,2238898756661 | NA | NA | NA | NA | NA | + | + | NA | NA | NA | NA | NA | 0,046067354606595 | 4 | -526,87338867319 | 1062,04980764941 | 14,0882347341189 | 3.03498360798934e-05 |
| **542** | 25,1594879777179 | + | NA | + | + | + | NA | NA | NA | NA | + | NA | NA | 0,0906847705982562 | 7 | -523,593688195021 | 1062,05559344431 | 14,0940205290146 | 3.02621639908794e-05 |
| **3594** | 24,5135099289927 | + | NA | NA | + | NA | NA | NA | NA | NA | + | + | + | 0,0906253124017548 | 7 | -523,598164972911 | 1062,06454700009 | 14,102974084796 | 3.01269898023689e-05 |
| **1663** | 24,0582579614408 | NA | + | + | + | + | + | + | NA | NA | + | + | NA | 0,135130551172672 | 10 | -520,162616797148 | 1062,07126534033 | 14,109692425038 | 3.00259579050574e-05 |
| **3590** | 24,8863564409612 | + | NA | + | NA | NA | NA | NA | NA | NA | + | + | + | 0,0905727715634602 | 7 | -523,602120679608 | 1062,07245841348 | 14,1108854981887 | 3.00080516643517e-05 |
| **710** | 25,1890479670428 | + | NA | + | NA | NA | NA | + | + | NA | + | NA | NA | 0,0904771259894331 | 7 | -523,609321079179 | 1062,08685921262 | 14,125286297332 | 2.97927577333392e-05 |
| **3644** | 24,1545102920002 | + | + | NA | + | + | + | NA | NA | NA | + | + | + | 0,13502671507741 | 10 | -520,17083626151 | 1062,08770426905 | 14,1261313537607 | 2.97801721116907e-05 |
| **1733** | 25,3590754939153 | NA | NA | + | NA | NA | NA | + | + | NA | + | + | NA | 0,0903985867104961 | 7 | -523,615233115077 | 1062,09868328442 | 14,137110369126 | 2.9617141516351e-05 |
| **2574** | 24,4639690066674 | + | NA | + | + | NA | NA | NA | NA | NA | + | NA | + | 0,09037186215745 | 7 | -523,617244686629 | 1062,10270642752 | 14,1411335122314 | 2.95576243985455e-05 |
| **3852** | 24,6734602775312 | + | + | NA | + | NA | NA | NA | NA | + | + | + | + | 0,120090996228952 | 9 | -521,342962173259 | 1062,10324718116 | 14,1416742658728 | 2.95496337823203e-05 |
| **3667** | 24,4220218306787 | NA | + | NA | NA | + | NA | + | NA | NA | + | + | + | 0,105217853276465 | 8 | -522,490580416837 | 1062,10616083367 | 14,1445879183841 | 2.95066164419845e-05 |
| **2744** | 24,3799344251868 | + | + | + | NA | + | + | NA | + | NA | + | NA | + | 0,134844459305369 | 10 | -520,185260891335 | 1062,1165535287 | 14,1549806134112 | 2.93536874891105e-05 |
| **3908** | 24,6630260398696 | + | + | NA | NA | NA | NA | + | NA | + | + | + | + | 0,119917495236334 | 9 | -521,35646102816 | 1062,13024489097 | 14,1686719756747 | 2.91534277437695e-05 |
| **2680** | 24,3331279090725 | + | + | + | NA | + | + | + | NA | NA | + | NA | + | 0,13474329183436 | 10 | -520,193266478217 | 1062,13256470247 | 14,1709917871751 | 2.91196321189494e-05 |
| **3759** | 24,243397892655 | NA | + | + | + | NA | + | NA | + | NA | + | + | + | 0,134736234850614 | 10 | -520,193824876727 | 1062,13368149949 | 14,1721085841953 | 2.91033762987958e-05 |
| **3972** | 25,0663171319975 | + | + | NA | NA | NA | NA | NA | + | + | + | + | + | 0,119878306724906 | 9 | -521,359509633999 | 1062,13634210264 | 14,1747691873538 | 2.90646857720832e-05 |

| | | | | | | | | | | | | | | | | | | |
|---|---|---|---|---|---|---|---|---|---|---|---|---|---|---|---|---|---|---|
| 1689 | 25,194878877134 | NA | NA | NA | + | + | NA | NA | + | NA | + | + | NA | 0,0901412305861856 | 7 | -523,634601995484 | 1062,13742104523 | 14,1758481299407 | 2.90490104370328e-05 |
| 2973 | 25,4099131736581 | NA | NA | + | + | + | NA | NA | + | + | + | NA | + | 0,119828688294289 | 9 | -521,363369423621 | 1062,14406168189 | 14,1824887665966 | 2.89527184238024e-05 |
| 666 | 25,1095716754952 | + | NA | NA | + | + | NA | NA | + | NA | + | NA | NA | 0,09003357342441 | 7 | -523,6427027572 | 1062,15362256866 | 14,1920496533739 | 2.881464188891e-05 |
| 1783 | 24,2988147457574 | NA | + | + | NA | + | + | + | + | NA | + | + | NA | 0,134602981536266 | 10 | -520,204367967871 | 1062,15476768177 | 14,1931947664841 | 2.87981485989353e-05 |
| 1562 | 25,120743110208 | + | NA | NA | + | + | NA | NA | NA | NA | + | + | NA | 0,0899815453806506 | 7 | -523,646617311662 | 1062,16145167759 | 14,1998787622974 | 2.87020658901364e-05 |
| 3021 | 25,2944387144556 | NA | NA | + | + | NA | NA | + | + | + | + | NA | + | 0,119696289883093 | 9 | -521,373667556759 | 1062,16465794816 | 14,2030850328738 | 2.8656089458531e-05 |
| 1544 | 25,0291888576384 | + | + | + | NA | NA | NA | NA | NA | NA | + | + | NA | 0,0899419097070309 | 7 | -523,649599322545 | 1062,16741569935 | 14,2058427840623 | 2.8616603505515e-05 |
| 1737 | 24,9857940847498 | NA | NA | NA | + | NA | NA | + | + | NA | + | + | NA | 0,0898741598862726 | 7 | -523,654696215593 | 1062,17760948545 | 14,2160365701586 | 2.84711188129582e-05 |
| 1660 | 24,1033321673619 | + | + | NA | + | + | + | + | NA | NA | + | + | NA | 0,134415882527125 | 10 | -520,219168628062 | 1062,18436900216 | 14,2227960868647 | 2.8375055737762e-05 |
| 1724 | 24,1169799377845 | + | + | NA | + | + | + | NA | + | NA | + | + | NA | 0,134364337871212 | 10 | -520,223245559455 | 1062,19252286494 | 14,2309499496521 | 2.82596080782291e-05 |
| 3033 | 25,279132245806 | NA | NA | NA | + | + | NA | + | + | + | + | NA | + | 0,119513920718301 | 9 | -521,387849950339 | 1062,19302273532 | 14,2314498200337 | 2.82525458902729e-05 |
| 2647 | 24,3960932317887 | NA | + | + | NA | + | NA | + | NA | NA | + | NA | + | 0,104648380397099 | 8 | -522,53414130564 | 1062,19328261128 | 14,2317096959896 | 2.82488750500844e-05 |
| 1775 | 24,1965998380372 | NA | + | + | + | NA | + | + | + | NA | + | + | NA | 0,134333480233738 | 10 | -520,225686132275 | 1062,19740401058 | 14,2358310952909 | 2.81907225411434e-05 |
| 2751 | 24,1392705969755 | NA | + | + | + | + | + | NA | + | NA | + | NA | + | 0,13426585109409 | 10 | -520,231034709471 | 1062,20810116497 | 14,2465282496823 | 2.80403447974058e-05 |
| 2572 | 24,0040225164721 | + | + | NA | + | NA | NA | NA | NA | NA | + | NA | + | 0,0896627093522375 | 7 | -523,670601431838 | 1062,20941991794 | 14,2478470026485 | 2.80218617477822e-05 |
| 2909 | 25,2810782787756 | NA | NA | + | + | + | NA | + | NA | + | + | NA | + | 0,119396045426965 | 9 | -521,397015254687 | 1062,21135334402 | 14,2497804287284 | 2.79947857381019e-05 |
| 3968 | 24,8515354288169 | + | + | + | + | + | + | + | NA | + | + | + | + | 0,178640553341576 | 13 | -516,628629181807 | 1062,21660795711 | 14,2550350418185 | 2.79213313901029e-05 |
| 1623 | 24,825229723322 | NA | + | + | NA | + | NA | + | NA | NA | + | + | NA | 0,104476921841425 | 8 | -522,547251321148 | 1062,2195026423 | 14,2579297270047 | 2.78809488886323e-05 |
| 1768 | 24,4521904211126 | + | + | + | NA | NA | + | + | + | NA | + | + | NA | 0,134190687658088 | 10 | -520,236978660643 | 1062,21998906732 | 14,2584161520276 | 2.78741687175758e-05 |
| 3072 | 24,8353837726463 | + | + | + | + | + | + | + | + | + | + | NA | + | 0,178594871440239 | 13 | -516,632436849494 | 1062,22422329248 | 14,2626503771933 | 2.78152183885085e-05 |
| 3752 | 24,4841303430338 | + | + | + | NA | NA | + | NA | + | NA | + | + | + | 0,134089466975491 | 10 | -520,244982413227 | 1062,23599657249 | 14,2744236571946 | 2.76519612009006e-05 |
| 3869 | 25,3021768538486 | NA | NA | + | + | + | NA | NA | NA | + | + | + | + | 0,119177219818338 | 9 | -521,414026624079 | 1062,2453760828 | 14,2838031675135 | 2.75225838842834e-05 |
| 2846 | 25,3112958014577 | + | NA | + | + | + | NA | NA | NA | + | + | NA | + | 0,119096651271066 | 9 | -521,420288909902 | 1062,25790065445 | 14,2963277391591 | 2.7350768138569e-05 |
| 3993 | 25,3012628698516 | NA | NA | NA | + | + | NA | NA | + | + | + | + | + | 0,119076820727255 | 9 | -521,421830174629 | 1062,2609831839 | 14,2994102686134 | 2.73086458335326e-05 |
| 714 | 24,8266169215172 | + | NA | NA | + | NA | NA | + | + | NA | + | NA | NA | 0,0893168314640652 | 7 | -523,69661025608 | 1062,26143756642 | 14,2998646511339 | 2.73024422525897e-05 |
| 2970 | 25,3016170255829 | + | NA | NA | + | + | NA | NA | + | + | + | NA | + | 0,11906771255597 | 9 | -521,422538066092 | 1062,26239896683 | 14,3008260515389 | 2.72893211169891e-05 |
| 3929 | 25,2027520681633 | NA | NA | NA | + | + | NA | + | NA | + | + | + | + | 0,118987908204172 | 9 | -521,42874018595 | 1062,27480320655 | 14,3132302912552 | 2.71205932526544e-05 |
| 2892 | 24,4694515320164 | + | + | NA | + | NA | NA | + | NA | + | + | NA | + | 0,118972270842435 | 9 | -521,429955402149 | 1062,27723363894 | 14,3156607236533 | 2.70876558854702e-05 |
| 2906 | 25,2192557087314 | + | NA | NA | + | + | NA | + | NA | + | + | NA | + | 0,118857257716081 | 9 | -521,438892680664 | 1062,29510819597 | 14,3335352806835 | 2.68466445574117e-05 |
| 2958 | 25,3148116538574 | + | NA | + | + | NA | NA | NA | + | + | + | NA | + | 0,118779144669221 | 9 | -521,444961914782 | 1062,30724666421 | 14,3456737489184 | 2.66841994436836e-05 |
| 1677 | 25,3310939624377 | NA | NA | + | + | NA | NA | NA | + | NA | + | + | NA | 0,0889939681597003 | 7 | -523,720879554706 | 1062,30997616368 | 14,3484032483855 | 2.66478070285218e-05 |
| 3866 | 25,2314235870127 | + | NA | NA | + | + | NA | NA | NA | + | + | + | + | 0,11872829306188 | 9 | -521,448912698302 | 1062,31514823125 | 14,3535753159586 | 2.65789839269967e-05 |
| 3981 | 25,343664676042 | NA | NA | + | + | NA | NA | NA | + | + | + | + | + | 0,118713491385477 | 9 | -521,450062633281 | 1062,31744810121 | 14,3558751859164 | 2.65484373903067e-05 |
| 760 | 24,2753690254167 | + | + | + | NA | + | + | + | + | NA | + | NA | NA | 0,133523599432121 | 10 | -520,289709644138 | 1062,32545103431 | 14,3638781190175 | 2.64424169661743e-05 |
| 2687 | 24,0579683219709 | NA | + | + | + | + | + | + | NA | NA | + | NA | + | 0,133512515229956 | 10 | -520,290585468678 | 1062,32720268339 | 14,3656297680977 | 2.64192681871095e-05 |
| 2634 | 24,1609787359765 | + | NA | NA | + | NA | NA | + | NA | NA | + | NA | + | 0,0887527779549315 | 7 | -523,739003962123 | 1062,34622497851 | 14,3846520632198 | 2.61691818183765e-05 |
| 3700 | 24,3057709458516 | + | + | NA | NA | + | + | + | NA | NA | + | + | + | 0,13335000177848 | 10 | -520,303425275975 | 1062,35288229798 | 14,3913093826918 | 2.60822183327566e-05 |
| 3764 | 24,288967249479 | + | + | NA | NA | + | + | NA | + | NA | + | + | + | 0,133334540575065 | 10 | -520,304646704089 | 1062,35532515421 | 14,3937522389185 | 2.6050380225964e-05 |
| 37 | 25,2185469978388 | NA | NA | + | NA | NA | + | NA | NA | NA | NA | NA | NA | 0,0439072244197732 | 4 | -527,028257278399 | 1062,35954485983 | 14,3979719445379 | 2.59954756986541e-05 |
| 41 | 25,2194340329835 | NA | NA | NA | + | NA | + | NA | NA | NA | NA | NA | NA | 0,0439014092936202 | 4 | -527,028673716479 | 1062,36037773599 | 14,3988048206979 | 2.59846524464281e-05 |
| 1648 | 24,0629006559233 | + | + | + | + | NA | + | + | NA | NA | + | + | NA | 0,133280931544474 | 10 | -520,308881624715 | 1062,36379499546 | 14,4022220801719 | 2.5940292204959e-05 |
| 727 | 24,8019935730204 | NA | + | + | NA | + | NA | + | + | NA | + | NA | NA | 0,103475273583476 | 8 | -522,62378896285 | 1062,3725779257 | 14,4110050104102 | 2.58266260796427e-05 |
| 648 | 25,0628261792552 | + | + | + | NA | NA | NA | NA | + | NA | + | NA | NA | 0,0885753467666166 | 7 | -523,75233409258 | 1062,37288523942 | 14,4113123241327 | 2.58226579462161e-05 |
| 2644 | 24,401095850397 | + | + | NA | NA | + | NA | + | NA | NA | + | NA | + | 0,103464751794873 | 8 | -522,624592496849 | 1062,3741849937 | 14,4126120784072 | 2.5805881842988e-05 |
| 2759 | 24,4736151610086 | NA | + | + | NA | NA | NA | + | + | NA | + | NA | + | 0,10338166722426 | 8 | -522,630937216023 | 1062,38687443205 | 14,4253015167551 | 2.56426690867264e-05 |
| 4095 | 24,7939141074649 | NA | + | + | + | + | + | + | + | + | + | + | + | 0,177589301898372 | 13 | -516,716199295351 | 1062,3917481842 | 14,4301752689075 | 2.55802571558546e-05 |
| 607 | 24,5480966270645 | NA | + | + | + | + | NA | + | NA | NA | + | NA | NA | 0,10333583495755 | 8 | -522,634436926794 | 1062,39387385359 | 14,4323009382967 | 2.55530840138893e-05 |
| 1730 | 25,0482691796588 | + | NA | NA | NA | NA | NA | + | + | NA | + | + | NA | 0,0884245743182517 | 7 | -523,763659354836 | 1062,39553576393 | 14,4339628486441 | 2.55318593661111e-05 |
| 920 | 25,5968341453577 | + | + | + | NA | + | NA | NA | + | + | + | NA | NA | 0,118197581524613 | 9 | -521,490131359589 | 1062,39758555382 | 14,4360126385332 | 2.55057052973984e-05 |
| 968 | 25,3264652070301 | + | + | + | NA | NA | NA | + | + | + | + | NA | NA | 0,118147361077505 | 9 | -521,494030536228 | 1062,4053839071 | 14,4438109918117 | 2.54064476843736e-05 |
| 1991 | 25,4413827617294 | NA | + | + | NA | NA | NA | + | + | + | + | + | NA | 0,118136467391203 | 9 | -521,494876305999 | 1062,40707544664 | 14,4455025313525 | 2.53849687633398e-05 |
| 654 | 25,1601106593249 | + | NA | + | + | NA | NA | NA | + | NA | + | NA | NA | 0,0883406530818708 | 7 | -523,769962281764 | 1062,40814161779 | 14,4465687025008 | 2.53714400090041e-05 |
| 3632 | 24,045900514373 | + | + | + | + | NA | + | NA | NA | NA | + | + | + | 0,132989627350835 | 10 | -520,331889032856 | 1062,40980981174 | 14,4482368964536 | 2.53502865908331e-05 |
| 1668 | 25,0053067727 | + | + | NA | NA | NA | NA | NA | + | NA | + | + | NA | 0,0882815762001818 | 7 | -523,774398918976 | 1062,41701489222 | 14,4554419769247 | 2.52591254678071e-05 |
| 2771 | 24,387098354303 | NA | + | NA | NA | + | NA | + | + | NA | + | NA | + | 0,1031653868362 | 8 | -522,647450623705 | 1062,41990124741 | 14,4583283321194 | 2.52226983555215e-05 |
| 1816 | 25,6522369154142 | + | + | + | NA | + | NA | NA | NA | + | + | + | NA | 0,118050427359689 | 9 | -521,501555960479 | 1062,4204347556 | 14,4588618403129 | 2.52159709947197e-05 |
| 1548 | 24,6128352126344 | + | + | NA | + | NA | NA | NA | NA | NA | + | + | NA | 0,0881651490768388 | 7 | -523,783141682704 | 1062,43450041967 | 14,4729275043812 | 2.50392534469321e-05 |
| 600 | 24,7704238094818 | + | + | + | NA | + | NA | + | NA | NA | + | NA | NA | 0,103008755989084 | 8 | -522,659407193662 | 1062,44381438732 | 14,4822414720343 | 2.49229171469888e-05 |
| 528 | 24,5580514847006 | + | + | + | + | NA | NA | NA | NA | NA | + | NA | NA | 0,0880958104582406 | 7 | -523,788347938967 | 1062,4449129322 | 14,4833400169066 | 2.49092314345076e-05 |
| 1057 | 25,7645689572932 | NA | NA | NA | NA | NA | + | NA | NA | NA | NA | + | NA | 0,043269224920496 | 4 | -527,073931184738 | 1062,45089267251 | 14,4893197572153 | 2.48348672918362e-05 |
| 2807 | 24,2519983759234 | NA | + | + | NA | + | + | + | + | NA | + | NA | + | 0,132693554688154 | 10 | -520,355265142309 | 1062,45656203065 | 14,4949891153585 | 2.47645680981778e-05 |
| 1940 | 25,5160907570835 | + | + | NA | NA | + | NA | NA | + | + | + | + | NA | 0,117728639740993 | 9 | -521,52653193345 | 1062,47038670155 | 14,5088137862547 | 2.45939773656804e-05 |
| 652 | 24,6706616105325 | + | + | NA | + | NA | NA | NA | + | NA | + | NA | NA | 0,087924969577651 | 7 | -523,801173754652 | 1062,47056456357 | 14,5089916482762 | 2.45917902956659e-05 |
| 2684 | 24,0729880484049 | + | + | NA | + | + | + | + | NA | NA | + | NA | + | 0,132597947495697 | 10 | -520,362812005249 | 1062,47165575653 | 14,5100828412401 | 2.45783767609166e-05 |
| 2651 | 24,230416936728 | NA | + | NA | + | + | NA | + | NA | NA | + | NA | + | 0,102818566852398 | 8 | -522,673922663647 | 1062,47284532729 | 14,5112724120027 | 2.45637622484037e-05 |
| 3018 | 25,1002603632012 | + | NA | NA | + | NA | NA | + | + | + | + | NA | + | 0,117665476968783 | 9 | -521,531433326818 | 1062,48018948828 | 14,518616572991 | 2.44737275446971e-05 |
| 1780 | 24,2306156376398 | + | + | NA | NA | + | + | + | + | NA | + | + | NA | 0,132507329434089 | 10 | -520,369964278019 | 1062,48596030207 | 14,5243873867785 | 2.44032126635101e-05 |
| 1613 | 24,8825323563553 | NA | NA | + | + | NA | NA | + | NA | NA | + | + | NA | 0,0878057530022508 | 7 | -523,810122472391 | 1062,48846199904 | 14,5268890837542 | 2.43727070247942e-05 |
| 3707 | 24,0772530417647 | NA | + | NA | + | + | + | + | NA | NA | + | + | + | 0,132484557650367 | 10 | -520,371761484522 | 1062,48955471507 | 14,5279817997844 | 2.43593944380126e-05 |
| 3815 | 24,3616856729044 | NA | + | + | NA | NA | + | + | + | NA | + | + | + | 0,132477762246959 | 10 | -520,372297785732 | 1062,4906273175 | 14,5290544022057 | 2.43463339677703e-05 |
| 4041 | 25,128266765155 | NA | NA | NA | + | NA | NA | + | + | + | + | + | + | 0,117559682339254 | 9 | -521,539642141248 | 1062,49660711714 | 14,5350342018517 | 2.42736495818131e-05 |
| 2686 | 24,4105588890523 | + | NA | + | + | + | + | + | NA | NA | + | NA | + | 0,132428085859013 | 10 | -520,376218177024 | 1062,49846810008 | 14,5368951847886 | 2.42510736629475e-05 |
| 2748 | 24,0695093763703 | + | + | NA | + | + | + | NA | + | NA | + | NA | + | 0,132426493509038 | 10 | -520,376343839353 | 1062,49871942474 | 14,5371465094468 | 2.42480264080148e-05 |
| 1627 | 24,6065993969033 | NA | + | NA | + | + | NA | + | NA | NA | + | + | NA | 0,102649015231092 | 8 | -522,686860460105 | 1062,49872092021 | 14,5371480049191 | 2.4248008276895e-05 |
| 1726 | 24,5275211496283 | + | NA | + | + | + | + | NA | + | NA | + | + | NA | 0,132419047673033 | 10 | -520,376931433938 | 1062,49989461391 | 14,5383216986179 | 2.42337825841935e-05 |
| 2081 | 25,3927852274246 | NA | NA | NA | NA | NA | + | NA | NA | NA | NA | NA | + | 0,042907567190274 | 4 | -527,099808472616 | 1062,50264724826 | 14,5410743329708 | 2.42004521548554e-05 |

| | | | | | | | | | | | | | | | | | | |
|---|---|---|---|---|---|---|---|---|---|---|---|---|---|---|---|---|---|---|
| **2736** | 24,1672409763845 | + | + | + | + | NA | + | NA | + | NA | + | NA | + | 0,132401462504193 | 10 | -520,378319162731 | 1062,50267007149 | 14,5410971562039 | 2.420017599015e-05 |
| **49** | 25,7519581417473 | NA | NA | NA | NA | + | + | NA | NA | NA | NA | NA | NA | 0,0428792516920999 | 4 | -527,101834086948 | 1062,50669847693 | 14,5451255616358 | 2.4151480987211e-05 |
| **3771** | 24,0843115869936 | NA | + | NA | + | + | + | NA | + | NA | + | + | + | 0,132283404581481 | 10 | -520,38763494358 | 1062,52130163319 | 14,5597287179021 | 2.39757792921777e-05 |
| **1662** | 24,428884554955 | + | NA | + | + | + | + | + | NA | NA | + | + | NA | 0,132279505757112 | 10 | -520,387942572593 | 1062,52191689122 | 14,5603439759275 | 2.39684047812304e-05 |
| **3655** | 24,2952826255761 | NA | + | + | NA | NA | NA | + | NA | NA | + | + | + | 0,102472570549828 | 8 | -522,700321643876 | 1062,52564328775 | 14,5640703724621 | 2.39237884683976e-05 |
| **2668** | 23,5648271267576 | + | + | NA | + | NA | + | + | NA | NA | + | NA | + | 0,117358446209864 | 9 | -521,55525373564 | 1062,52783030592 | 14,5662573906341 | 2.38976418867038e-05 |
| **2750** | 24,4829446959308 | + | NA | + | + | + | + | NA | + | NA | + | NA | + | 0,132221542275157 | 10 | -520,392515903363 | 1062,53106355276 | 14,5694906374681 | 2.38590396102973e-05 |
| **2048** | 24,744285651371 | + | + | + | + | + | + | + | + | + | + | + | NA | 0,176742534914838 | 13 | -516,786654371595 | 1062,53265833669 | 14,5710854213951 | 2.38400221870334e-05 |
| **1558** | 25,3879408250092 | + | NA | + | NA | + | NA | NA | NA | NA | + | + | NA | 0,0874863095032694 | 7 | -523,834095001708 | 1062,53640705768 | 14,5748341423898 | 2.37953792427454e-05 |
| **3611** | 24,5157780477548 | NA | + | NA | + | + | NA | NA | NA | NA | + | + | + | 0,102385069502383 | 8 | -522,706996226339 | 1062,53899245268 | 14,5774195373881 | 2.37646388887773e-05 |
| **293** | 25,8129420031856 | NA | NA | + | NA | NA | + | NA | NA | + | NA | NA | NA | 0,0575751856968265 | 5 | -526,04239140481 | 1062,5427980768 | 14,5812251615052 | 2.37194622422969e-05 |
| **1943** | 25,7189505207413 | NA | + | + | NA | + | NA | NA | + | + | + | + | NA | 0,117259786815641 | 9 | -521,562906282558 | 1062,54313539976 | 14,5815624844704 | 2.37154620199804e-05 |
| **662** | 25,3687874029598 | + | NA | + | NA | + | NA | NA | + | NA | + | NA | NA | 0,0874058999788478 | 7 | -523,840127985817 | 1062,5484730259 | 14,5869001106073 | 2.36522542673719e-05 |
| **3779** | 24,4015591551615 | NA | + | NA | NA | NA | NA | + | + | NA | + | + | + | 0,102297642779999 | 8 | -522,713664489811 | 1062,55232897962 | 14,5907560643304 | 2.3606697199019e-05 |
| **767** | 24,002931396507 | NA | + | + | + | + | + | + | + | NA | + | NA | NA | 0,132086448486613 | 10 | -520,403173646393 | 1062,55237903882 | 14,5908061235277 | 2.36061063402573e-05 |
| **417** | 26,6069492289313 | NA | NA | NA | NA | NA | + | NA | + | + | NA | NA | NA | 0,0574625695710178 | 5 | -526,050572618132 | 1062,55916050344 | 14,5975875881491 | 2.35261998999923e-05 |
| **1747** | 24,7704357783105 | NA | + | NA | NA | + | NA | + | + | NA | + | + | NA | 0,102245597890693 | 8 | -522,71763378065 | 1062,5602675613 | 14,5986946460087 | 2.3513181071221e-05 |
| **752** | 24,1174240250228 | + | + | + | + | NA | + | + | + | NA | + | NA | NA | 0,132017916200017 | 10 | -520,408579623022 | 1062,56319099208 | 14,6016180767845 | 2.34788365996669e-05 |
| **3646** | 24,4759375554932 | + | NA | + | + | + | + | NA | NA | NA | + | + | + | 0,131983516513641 | 10 | -520,411292984694 | 1062,56861771542 | 14,6070448001283 | 2.34152163757078e-05 |
| **1620** | 24,7731393959354 | + | + | NA | NA | + | NA | + | NA | NA | + | + | NA | 0,102183989891879 | 8 | -522,722332120826 | 1062,56966424165 | 14,6080913263604 | 2.3402967261658e-05 |
| **3607** | 24,850667676029 | NA | + | + | NA | + | NA | NA | NA | NA | + | + | + | 0,102175359192159 | 8 | -522,722990288249 | 1062,5709805765 | 14,6094076612069 | 2.3387569258787e-05 |
| **3026** | 25,3897689513994 | + | NA | NA | NA | + | NA | + | + | + | + | NA | + | 0,116990847059432 | 9 | -521,583762338767 | 1062,58484751218 | 14,6232745968885 | 2.32259731575095e-05 |
| **3978** | 25,1536810756585 | + | NA | NA | + | NA | NA | NA | + | + | + | + | + | 0,116976912856218 | 9 | -521,584842751724 | 1062,58700833809 | 14,6254354228033 | 2.32008930660286e-05 |
| **1682** | 25,3243568486474 | + | NA | NA | NA | + | NA | NA | + | NA | + | + | NA | 0,0871260248616588 | 7 | -523,861122377581 | 1062,59046180943 | 14,6288888941356 | 2.31608658246637e-05 |
| **3029** | 25,5257456547066 | NA | NA | + | NA | + | NA | + | + | + | + | NA | + | 0,116937082917533 | 9 | -521,587930942115 | 1062,59318471888 | 14,6316118035854 | 2.31293548097306e-05 |
| **1735** | 24,9837920546174 | NA | + | + | NA | NA | NA | + | + | NA | + | + | NA | 0,102025600667202 | 8 | -522,734409697637 | 1062,59381939527 | 14,6322464799825 | 2.31220161464222e-05 |
| **588** | 24,1280914650738 | + | + | NA | + | NA | NA | + | NA | NA | + | NA | NA | 0,0870956244269016 | 7 | -523,86340243101 | 1062,59502191628 | 14,6334490009938 | 2.31081179699341e-05 |
| **2902** | 25,3937692568924 | + | NA | + | NA | + | NA | + | NA | + | + | NA | + | 0,116840098778782 | 9 | -521,59544996692 | 1062,60822276849 | 14,6466498531956 | 2.29560967991016e-05 |
| **3922** | 25,3143386094617 | + | NA | NA | NA | + | NA | + | NA | + | + | + | + | 0,116837569442381 | 9 | -521,595646051274 | 1062,60861493719 | 14,6470420219034 | 2.29515959089844e-05 |
| **4049** | 25,4164398570732 | NA | NA | NA | NA | + | NA | + | + | + | + | + | + | 0,11671206274133 | 9 | -521,6053751318 | 1062,62807309824 | 14,666500182954 | 2.27293807117881e-05 |
| **3925** | 25,4191770515693 | NA | NA | + | NA | + | NA | + | NA | + | + | + | + | 0,116707782230836 | 9 | -521,605706925821 | 1062,62873668629 | 14,6671637709978 | 2.27218404901154e-05 |
| **3827** | 24,1981992756658 | NA | + | NA | NA | + | + | + | + | NA | + | + | + | 0,131592820180115 | 10 | -520,44210259931 | 1062,63023694465 | 14,6686640293622 | 2.27048025656124e-05 |
| **34** | 25,5504493850521 | + | NA | NA | NA | NA | + | NA | NA | NA | NA | NA | NA | 0,0420112235397944 | 4 | -527,163901432672 | 1062,63083316838 | 14,6692602530845 | 2.26980350034571e-05 |
| **1774** | 24,5043205733549 | + | NA | + | + | NA | + | + | + | NA | + | + | NA | 0,131533061071475 | 10 | -520,446813874175 | 1062,63965949438 | 14,6780865790906 | 2.25980855838961e-05 |
| **1685** | 25,5491158899772 | NA | NA | + | NA | + | NA | NA | + | NA | + | + | NA | 0,0867751581388079 | 7 | -523,887433006517 | 1062,6430830673 | 14,6815101520076 | 2.25594355767231e-05 |
| **766** | 24,3778155464886 | + | NA | + | + | + | + | + | + | NA | + | NA | NA | 0,131477925623493 | 10 | -520,451160342568 | 1062,64835243117 | 14,6867795158764 | 2.25000768693524e-05 |
| **3709** | 24,3843235111669 | NA | NA | + | + | + | + | + | NA | NA | + | + | + | 0,131412608707088 | 10 | -520,456309085297 | 1062,65864991663 | 14,6970770013345 | 2.23845274844597e-05 |
| **604** | 24,560595922899 | + | + | NA | + | + | NA | + | NA | NA | + | NA | NA | 0,101551830674885 | 8 | -522,770523145476 | 1062,66604629095 | 14,7044733756611 | 2.23018981957212e-05 |
| **3014** | 25,3850285661366 | + | NA | + | NA | NA | NA | + | + | + | + | NA | + | 0,116448216410746 | 9 | -521,625823576311 | 1062,66896998727 | 14,7073970719771 | 2.22693200248953e-05 |
| **731** | 24,5489293515485 | NA | + | NA | + | + | NA | + | + | NA | + | NA | NA | 0,101511722434997 | 8 | -522,773579550063 | 1062,67215910013 | 14,7105861848347 | 2.22338386335426e-05 |
| **3758** | 24,5377128796873 | + | NA | + | + | NA | + | NA | + | NA | + | + | + | 0,131299870233159 | 10 | -520,465195021261 | 1062,67642178855 | 14,7148488732626 | 2.21865011343899e-05 |
| **1606** | 24,9790201042118 | + | NA | + | NA | NA | NA | + | NA | NA | + | + | NA | 0,0865372810851016 | 7 | -523,905265072712 | 1062,67874719969 | 14,7171742843973 | 2.21607197569806e-05 |
| **3773** | 24,4686495287284 | NA | NA | + | + | + | + | NA | + | NA | + | + | + | 0,131233120886257 | 10 | -520,470455596101 | 1062,68694293823 | 14,7253700229428 | 2.20700938378309e-05 |
| **297** | 25,8411866930379 | NA | NA | NA | + | NA | + | NA | NA | + | NA | NA | NA | 0,0565750973684153 | 5 | -526,115010561706 | 1062,68803639059 | 14,7264634752964 | 2.20580308376843e-05 |
| **590** | 24,6926952544459 | + | NA | + | + | NA | NA | + | NA | NA | + | NA | NA | 0,0864736350933891 | 7 | -523,910035403833 | 1062,68828786193 | 14,7267149466388 | 2.20552575307272e-05 |
| **1935** | 25,5263360980314 | NA | + | + | + | NA | NA | NA | + | + | + | + | NA | 0,116294672517171 | 9 | -521,637720624831 | 1062,69276408431 | 14,7311911690163 | 2.20059506098454e-05 |
| **912** | 25,4140331899432 | + | + | + | + | NA | NA | NA | + | + | + | NA | NA | 0,116261507103018 | 9 | -521,640290110794 | 1062,69790305623 | 14,7363301409428 | 2.19494792109487e-05 |
| **1787** | 24,0055376825271 | NA | + | NA | + | + | + | + | + | NA | + | + | NA | 0,131156025171993 | 10 | -520,476531074532 | 1062,6990938951 | 14,737520979806 | 2.19364139545518e-05 |
| **2591** | 24,4857866228888 | NA | + | + | + | + | NA | NA | NA | NA | + | NA | + | 0,101313702781076 | 8 | -522,788667422977 | 1062,70233484595 | 14,7407619306632 | 2.19008953210561e-05 |
| **1674** | 25,0575667656311 | + | NA | NA | + | NA | NA | NA | + | NA | + | + | NA | 0,0863475953679362 | 7 | -523,919481227853 | 1062,70717950997 | 14,7456065946787 | 2.18479082833292e-05 |
| **353** | 25,9070040277551 | NA | NA | NA | NA | NA | + | + | NA | + | NA | NA | NA | 0,0563847987851797 | 5 | -526,128819949859 | 1062,71565516689 | 14,7540822516032 | 2.17555165042946e-05 |
| **764** | 23,9732236537528 | + | + | NA | + | + | + | + | + | NA | + | NA | NA | 0,131026637827895 | 10 | -520,486726150516 | 1062,71948404706 | 14,757911131773 | 2.17139067138656e-05 |
| **3688** | 24,2119880108909 | + | + | + | NA | NA | + | + | NA | NA | + | + | + | 0,130966884049838 | 10 | -520,491433937415 | 1062,72889962086 | 14,7673267055713 | 2.16119225166002e-05 |
| **4092** | 24,7140370424295 | + | + | NA | + | + | + | + | + | + | + | + | + | 0,175540418647996 | 13 | -516,886551914062 | 1062,73245342162 | 14,7708805063294 | 2.15735543815613e-05 |
| **2813** | 24,3543932110257 | NA | NA | + | + | + | + | + | + | NA | + | NA | + | 0,130900403614078 | 10 | -520,496671313458 | 1062,73937437295 | 14,777801457657 | 2.14990286432636e-05 |
| **1995** | 25,130678844463 | NA | + | NA | + | NA | NA | + | + | + | + | + | NA | 0,115975018067593 | 9 | -521,662481798607 | 1062,74228643186 | 14,7807135165681 | 2.14677482024414e-05 |
| **2804** | 24,1882036596269 | + | + | NA | NA | + | + | + | + | NA | + | NA | + | 0,130870005709923 | 10 | -520,499065948577 | 1062,74416364319 | 14,7825907278959 | 2.14476079057639e-05 |
| **4037** | 25,4395410381742 | NA | NA | + | NA | NA | NA | + | + | + | + | + | + | 0,115959596546066 | 9 | -521,663676159076 | 1062,7446751528 | 14,7831022375069 | 2.14421232783654e-05 |
| **3862** | 25,4557826328428 | + | NA | + | NA | + | NA | NA | NA | + | + | + | + | 0,115956589176625 | 9 | -521,66390907032 | 1062,74514097528 | 14,7835680599942 | 2.14371297483135e-05 |
| **972** | 25,0488966580013 | + | + | NA | + | NA | NA | + | + | + | + | NA | NA | 0,115952513789963 | 9 | -521,664224694852 | 1062,74577222435 | 14,7841993090599 | 2.14303647319099e-05 |
| **2966** | 25,527544366234 | + | NA | + | NA | + | NA | NA | + | + | + | NA | + | 0,11592690511663 | 9 | -521,666207964319 | 1062,74973876328 | 14,7881658479926 | 2.13879046626816e-05 |
| **3986** | 25,4450033624526 | + | NA | NA | NA | + | NA | NA | + | + | + | + | + | 0,11590586408796 | 9 | -521,667837448546 | 1062,75299773174 | 14,7914248164468 | 2.13530817887497e-05 |
| **1871** | 25,0352994413849 | NA | + | + | + | NA | NA | + | NA | + | + | + | NA | 0,115865726947462 | 9 | -521,670945688959 | 1062,75921421256 | 14,7976412972728 | 2.12868143179592e-05 |
| **724** | 24,695539644112 | + | + | NA | NA | + | NA | + | + | NA | + | NA | NA | 0,100888055671884 | 8 | -522,821087855337 | 1062,76717571067 | 14,8056027953828 | 2.12022452876982e-05 |
| **1789** | 24,380512822975 | NA | NA | + | + | + | + | + | + | NA | + | + | NA | 0,130682895769862 | 10 | -520,51380393846 | 1062,77363962295 | 14,8120667076614 | 2.11338311761295e-05 |
| **719** | 24,6759313557974 | NA | + | + | + | NA | NA | + | + | NA | + | NA | NA | 0,100801314046451 | 8 | -522,827692857469 | 1062,78038571494 | 14,8188127996468 | 2.10626658809504e-05 |
| **3917** | 25,028811944891 | NA | NA | + | + | NA | NA | + | NA | + | + | + | + | 0,115643429094761 | 9 | -521,688157993591 | 1062,79363882183 | 14,8320659065366 | 2.09235544235973e-05 |
| **4032** | 24,766985392629 | + | + | + | + | + | + | NA | + | + | + | + | + | 0,175138161491007 | 13 | -516,919947540992 | 1062,79924467548 | 14,8376717601886 | 2.0864989346763e-05 |
| **1550** | 25,0120535765948 | + | NA | + | + | NA | NA | NA | NA | NA | + | + | NA | 0,0856394224371352 | 7 | -523,972529788431 | 1062,81327663113 | 14,8517037158344 | 2.07191133739083e-05 |
| **2911** | 24,8997954912468 | NA | + | + | + | + | NA | + | NA | + | + | NA | + | 0,130430340333351 | 10 | -520,533691812018 | 1062,81341537007 | 14,8518424547774 | 2.07176761498145e-05 |
| **2894** | 24,9810142525871 | + | NA | + | + | NA | NA | + | NA | + | + | NA | + | 0,115504449828645 | 9 | -521,698916824984 | 1062,81515648461 | 14,8535835693233 | 2.06996480745383e-05 |
| **3854** | 25,0620846519672 | + | NA | + | + | NA | NA | NA | NA | + | + | + | + | 0,115462087800344 | 9 | -521,702195869657 | 1062,82171457396 | 14,8601416586687 | 2.06318841649534e-05 |
| **3604** | 24,7912730118929 | + | + | NA | NA | + | NA | NA | NA | NA | + | + | + | 0,100494939790013 | 8 | -522,85101684489 | 1062,82703368978 | 14,8654607744891 | 2.0577085376796e-05 |
| **3702** | 24,6578416573438 | + | NA | + | NA | + | + | + | NA | NA | + | + | + | 0,130246167830138 | 10 | -520,548191124569 | 1062,84241399517 | 14,8808410798802 | 2.04194513399777e-05 |
| **4034** | 25,2284987292984 | + | NA | NA | NA | NA | NA | + | + | + | + | + | + | 0,115292654118904 | 9 | -521,715309362064 | 1062,84794155877 | 14,8863686434829 | 2.03630943470594e-05 |
| **3706** | 24,3688460411852 | + | NA | NA | + | + | + | + | NA | NA | + | + | + | 0,130199224893813 | 10 | -520,551886301144 | 1062,84980434832 | 14,8882314330297 | 2.0344137097127e-05 |

| | | | | | | | | | | | | | | | | | | | |
|---|---|---|---|---|---|---|---|---|---|---|---|---|---|---|---|---|---|---|---|
| 3989 | 25,5904529874166 | NA | NA | + | NA | + | NA | NA | + | + | + | + | + | 0,115277776902655 | 9 | -521,716460679491 | 1062,85024419363 | 14,8886712783367 | 2.03396634524591e-05 |
| 3731 | 24,7426430264553 | NA | + | NA | NA | + | NA | NA | + | NA | + | + | + | 0,100285031334494 | 8 | -522,866992397205 | 1062,85898479441 | 14,8974118791184 | 2.02509669696313e-05 |
| 3821 | 24,4320228423278 | NA | NA | + | + | NA | + | + | + | NA | + | + | + | 0,130133538167974 | 10 | -520,557056585815 | 1062,86014491766 | 14,8985720023704 | 2.02392235670805e-05 |
| 3719 | 24,8810220825118 | NA | + | + | NA | NA | NA | NA | + | NA | + | + | + | 0,100231508466269 | 8 | -522,8710652787 | 1062,8671305574 | 14,9055576421085 | 2.01686549184038e-05 |
| 2588 | 24,4824074386554 | + | + | NA | + | + | NA | NA | NA | NA | + | NA | + | 0,100220666093167 | 8 | -522,871890311503 | 1062,86878062301 | 14,9072077077162 | 2.01520219787906e-05 |
| 3599 | 24,3855397277097 | NA | + | + | + | NA | NA | NA | NA | NA | + | + | + | 0,100199600351157 | 8 | -522,873493246582 | 1062,87198649316 | 14,9104135778741 | 2.0119745471333e-05 |
| 3766 | 24,6878552266169 | + | NA | + | NA | + | + | NA | + | NA | + | + | + | 0,130023160293675 | 10 | -520,565743688158 | 1062,87751912235 | 14,9159462070581 | 2.00641648380508e-05 |
| 225 | 25,7817265453405 | NA | NA | NA | NA | NA | + | + | + | NA | NA | NA | NA | 0,0552393289202233 | 5 | -526,21188444521 | 1062,8817841576 | 14,9202112423045 | 2.00214232427396e-05 |
| 2811 | 24,0005680440634 | NA | + | NA | + | + | + | + | + | NA | + | NA | + | 0,129996028499914 | 10 | -520,567878881038 | 1062,88178950811 | 14,9202165928168 | 2.00213696803755e-05 |
| 1567 | 24,9888352130298 | NA | + | + | + | + | NA | NA | NA | NA | + | + | NA | 0,100117413592433 | 8 | -522,879746645299 | 1062,8844932906 | 14,9229203753082 | 1.99943212533632e-05 |
| 1610 | 24,6942385792579 | + | NA | NA | + | NA | NA | + | NA | NA | + | + | NA | 0,085110459442999 | 7 | -524,012127116924 | 1062,89247128811 | 14,930898372822 | 1.99147227953621e-05 |
| 2337 | 26,0444224596787 | NA | NA | NA | NA | NA | + | NA | NA | + | NA | NA | + | 0,0549753529902623 | 5 | -526,231012559696 | 1062,92004038657 | 14,9584674712769 | 1.9642090690945e-05 |
| 1670 | 25,4042113649368 | + | NA | + | NA | NA | NA | NA | + | NA | + | + | NA | 0,0848864657916337 | 7 | -524,028888029202 | 1062,92599311267 | 14,9644201973772 | 1.95837156137902e-05 |
| 1988 | 25,1746295592334 | + | + | NA | NA | NA | NA | + | + | + | + | + | NA | 0,114769589545799 | 9 | -521,755776647238 | 1062,92887612912 | 14,9673032138303 | 1.95555058638048e-05 |
| 848 | 24,9290417257223 | + | + | + | + | NA | NA | + | NA | + | + | NA | NA | 0,114743051876282 | 9 | -521,757829116973 | 1062,93298106859 | 14,9714081533018 | 1.95154099417609e-05 |
| 3829 | 24,6095311815283 | NA | NA | + | NA | + | + | + | + | NA | + | + | + | 0,129645160540241 | 10 | -520,595485170174 | 1062,93700208638 | 14,9754291710901 | 1.94762134521485e-05 |
| 4088 | 24,9599059644291 | + | + | + | NA | + | + | + | + | + | + | + | + | 0,174296026936892 | 13 | -516,98980937209 | 1062,93896833768 | 14,9773954223858 | 1.94570752963281e-05 |
| 3974 | 25,4488089466586 | + | NA | + | NA | NA | NA | NA | + | + | + | + | + | 0,114581315828655 | 9 | -521,770336736163 | 1062,95799630697 | 14,9964233916803 | 1.92728387820552e-05 |
| 2806 | 24,5992895240071 | + | NA | + | NA | + | + | + | + | NA | + | NA | + | 0,129509408830103 | 10 | -520,606163129701 | 1062,95835800543 | 14,9967850901437 | 1.92693536191228e-05 |
| 163 | 25,4779456279628 | NA | + | NA | NA | NA | + | NA | + | NA | NA | NA | NA | 0,0546815550324634 | 5 | -526,252295346882 | 1062,96260596094 | 15,0010330456485 | 1.92284693747159e-05 |
| 2799 | 24,0650716609911 | NA | + | + | + | NA | + | + | + | NA | + | NA | + | 0,129423099455865 | 10 | -520,612951187966 | 1062,97193412196 | 15,0103612066723 | 1.91389950653218e-05 |
| 2715 | 24,4557917579973 | NA | + | NA | + | + | NA | NA | + | NA | + | NA | + | 0,0995311505171684 | 8 | -522,924337482201 | 1062,9736749644 | 15,0121020491122 | 1.91223433259538e-05 |
| 1864 | 25,1167436178421 | + | + | + | NA | NA | NA | + | NA | + | + | + | NA | 0,114444488984966 | 9 | -521,780916255729 | 1062,9791553461 | 15,0175824308139 | 1.90700161816477e-05 |
| 3695 | 23,9383534385676 | NA | + | + | + | NA | + | + | NA | NA | + | + | + | 0,129328586229555 | 10 | -520,620383691122 | 1062,98679912828 | 15,0252262129852 | 1.89972717559834e-05 |
| 2798 | 24,3535192207593 | + | NA | + | + | NA | + | + | + | NA | + | NA | + | 0,129323287501675 | 10 | -520,620800358259 | 1062,98763246255 | 15,0260595472591 | 1.89893578659942e-05 |
| 4094 | 25,0052147724323 | + | NA | + | + | + | + | + | + | + | + | + | + | 0,17399977106073 | 13 | -517,014369246788 | 1062,98808808707 | 15,026515171782 | 1.89850323501574e-05 |
| 1782 | 24,6346787927755 | + | NA | + | NA | + | + | + | + | NA | + | + | NA | 0,129291155160665 | 10 | -520,623327040737 | 1062,99268582751 | 15,0311129122158 | 1.89414383523907e-05 |
| 2810 | 24,288864398811 | + | NA | NA | + | + | + | + | + | NA | + | NA | + | 0,129241112178812 | 10 | -520,627261916253 | 1063,00055557854 | 15,0389826632479 | 1.88670525957032e-05 |
| 3871 | 25,0686235660235 | NA | + | + | + | + | NA | NA | NA | + | + | + | + | 0,129097770107114 | 10 | -520,63853164065 | 1063,02309502733 | 15,0615221120406 | 1.86556197455304e-05 |
| 1786 | 24,3136323208763 | + | NA | NA | + | + | + | + | + | NA | + | + | NA | 0,12905879870984 | 10 | -520,641595297775 | 1063,02922234158 | 15,0676494262907 | 1.85985527846115e-05 |
| 2792 | 24,2762310050836 | + | + | + | NA | NA | + | + | + | NA | + | NA | + | 0,129054162985565 | 10 | -520,64195971669 | 1063,02995117941 | 15,068378264122 | 1.85917763549779e-05 |
| 3031 | 25,0928561573914 | NA | + | + | NA | + | NA | + | + | + | + | NA | + | 0,129048459686404 | 10 | -520,642408056074 | 1063,03084785818 | 15,0692749428899 | 1.85834427976904e-05 |
| 3035 | 24,8824117882015 | NA | + | NA | + | + | NA | + | + | + | + | NA | + | 0,12902594159101 | 10 | -520,644178186879 | 1063,03438811979 | 15,0728152045001 | 1.85505767702952e-05 |
| 2584 | 24,7846653631713 | + | + | + | NA | + | NA | NA | NA | NA | + | NA | + | 0,0991247339476775 | 8 | -522,955232268782 | 1063,03546453756 | 15,0738916222738 | 1.85405953713012e-05 |
| 3770 | 24,3740098472757 | + | NA | NA | + | + | + | NA | + | NA | + | + | + | 0,129013611698827 | 10 | -520,645147411194 | 1063,03632656842 | 15,0747536531292 | 1.85326058105932e-05 |
| 3914 | 24,8901798030743 | + | NA | NA | + | NA | NA | + | NA | + | + | + | + | 0,114071388658784 | 9 | -521,809756258787 | 1063,03683535222 | 15,0752624369297 | 1.8527891865404e-05 |
| 3927 | 25,0554544307616 | NA | + | + | NA | + | NA | + | NA | + | + | + | + | 0,129000147995108 | 10 | -520,646205746149 | 1063,03844323833 | 15,0768703230383 | 1.85130024813388e-05 |
| 2975 | 25,1013142705562 | NA | + | + | + | + | NA | NA | + | + | + | NA | + | 0,128979680185453 | 10 | -520,647814618204 | 1063,04166098244 | 15,0800880671495 | 1.84832413763058e-05 |
| 99 | 24,5280806435933 | NA | + | NA | NA | NA | + | + | NA | NA | NA | NA | NA | 0,0541177067847269 | 5 | -526,293122121454 | 1063,04425951008 | 15,0826865947934 | 1.84592423633744e-05 |
| 3931 | 24,863082982131 | NA | + | NA | + | + | NA | + | NA | + | + | + | + | 0,12894179424744 | 10 | -520,650792542432 | 1063,0476168309 | 15,0860439156047 | 1.842828155771e-05 |
| 712 | 24,823993204069 | + | + | + | NA | NA | NA | + | + | NA | + | NA | NA | 0,0990144158393264 | 8 | -522,963615975734 | 1063,05223195147 | 15,0906590361776 | 1.83858062130968e-05 |
| 2908 | 24,865302771103 | + | + | NA | + | + | NA | + | NA | + | + | NA | + | 0,128880651626606 | 10 | -520,655598223828 | 1063,05722819369 | 15,0956552783973 | 1.83399335640468e-05 |
| 3723 | 24,4649677589868 | NA | + | NA | + | NA | NA | NA | + | NA | + | + | + | 0,0989639800004846 | 8 | -522,967448543174 | 1063,05989708635 | 15,0983241710571 | 1.83154762291413e-05 |
| 2848 | 25,0722148091771 | + | + | + | + | + | NA | NA | NA | + | + | NA | + | 0,128841751382891 | 10 | -520,65865552569 | 1063,06334279741 | 15,1017698821204 | 1.82839484762858e-05 |
| 2711 | 24,7625359846573 | NA | + | + | NA | + | NA | NA | + | NA | + | NA | + | 0,0989285377553955 | 8 | -522,970141634588 | 1063,06528326918 | 15,1037103538845 | 1.82662173364935e-05 |
| 2703 | 24,5155665104303 | NA | + | + | + | NA | NA | NA | + | NA | + | NA | + | 0,0989218477083467 | 8 | -522,970649968278 | 1063,06629993656 | 15,1047270212657 | 1.82569343624453e-05 |
| 3833 | 24,2993116552405 | NA | NA | NA | + | + | + | + | + | NA | + | + | + | 0,1287894861433 | 10 | -520,66276301301 | 1063,07155777205 | 15,109984856761 | 1.82090014171738e-05 |
| 1059 | 25,2900125996137 | NA | + | NA | NA | NA | + | NA | NA | NA | NA | + | NA | 0,0538697291899907 | 5 | -526,311069828294 | 1063,08015492376 | 15,1185820084727 | 1.81308966332589e-05 |
| 2904 | 25,0487376754886 | + | + | + | NA | + | NA | + | NA | + | + | NA | + | 0,12866526621233 | 10 | -520,67252437918 | 1063,09108050439 | 15,129507589101 | 1.80321213866124e-05 |
| 419 | 26,0337854006909 | NA | + | NA | NA | NA | + | NA | + | + | NA | NA | NA | 0,06875081868073 | 6 | -525,225617550835 | 1063,09738894782 | 15,135816032534 | 1.79753336853228e-05 |
| 3910 | 25,1400864712634 | + | NA | + | NA | NA | NA | + | NA | + | + | + | + | 0,113644178730095 | 9 | -521,842763935018 | 1063,10285070468 | 15,1412777893904 | 1.79263122006446e-05 |
| 3592 | 24,6987867655535 | + | + | + | NA | NA | NA | NA | NA | NA | + | + | + | 0,0986772329314366 | 8 | -522,989234083349 | 1063,1034681667 | 15,1418952514082 | 1.79207786464271e-05 |
| 3023 | 24,9038747715552 | NA | + | + | + | NA | NA | + | + | + | + | NA | + | 0,128562014183818 | 10 | -520,680637001396 | 1063,10730574882 | 15,1457328335334 | 1.78864253854302e-05 |
| 290 | 26,1159461775965 | + | NA | NA | NA | NA | + | NA | NA | + | NA | NA | NA | 0,0536641703441815 | 5 | -526,325943858146 | 1063,10990298347 | 15,1483300681762 | 1.7863212838977e-05 |
| 305 | 26,2978365773702 | NA | NA | NA | NA | + | + | NA | NA | + | NA | NA | NA | 0,0536438122327522 | 5 | -526,327416774709 | 1063,11284881659 | 15,1512759013024 | 1.78369211843763e-05 |
| 3652 | 24,1810911919743 | + | + | NA | NA | NA | NA | + | NA | NA | + | + | + | 0,0985149625243276 | 8 | -523,001559469643 | 1063,12811893929 | 15,1665460239956 | 1.77012537704638e-05 |
| 1808 | 25,2077826054738 | + | + | + | + | NA | NA | NA | NA | + | + | + | NA | 0,11345801546694 | 9 | -521,85714256088 | 1063,13160795641 | 15,1700350411156 | 1.76704007012844e-05 |
| 2763 | 24,1476322616638 | NA | + | NA | + | NA | NA | + | + | NA | + | NA | + | 0,0984731174670451 | 8 | -523,00473748678 | 1063,13447497356 | 15,1729020582695 | 1.76450881775155e-05 |
| 1882 | 25,4745440150897 | + | NA | NA | + | + | NA | + | NA | + | + | + | NA | 0,113386456048596 | 9 | -521,862668767313 | 1063,14266036927 | 15,1810874539806 | 1.75730197407357e-05 |
| 1313 | 26,2159591334742 | NA | NA | NA | NA | NA | + | NA | NA | + | NA | + | NA | 0,0534346174888043 | 5 | -526,342550253891 | 1063,14311577496 | 15,1815428596672 | 1.75690187697091e-05 |
| 544 | 24,9128466972475 | + | + | + | + | + | NA | NA | NA | NA | + | NA | NA | 0,0983725683304302 | 8 | -523,012373314714 | 1063,14974662943 | 15,1881737141366 | 1.75108664196031e-05 |
| 671 | 24,9063323726862 | NA | + | + | + | + | NA | NA | + | NA | + | NA | NA | 0,0982923567038177 | 8 | -523,018464076264 | 1063,16192815253 | 15,2003552372366 | 1.74045360528764e-05 |
| 355 | 25,1779694401806 | NA | + | NA | NA | NA | + | + | NA | + | NA | NA | NA | 0,068248039471992 | 6 | -525,262573266992 | 1063,17130038014 | 15,2097274648472 | 1.73231672169197e-05 |
| 2765 | 24,739836028324 | NA | NA | + | + | NA | NA | + | + | NA | + | NA | + | 0,0981975631113696 | 8 | -523,025661401571 | 1063,17632280314 | 15,2147498878508 | 1.72797196559972e-05 |
| 1885 | 25,6092130940101 | NA | NA | + | + | + | NA | + | NA | + | + | + | NA | 0,113139049352249 | 9 | -521,881771419871 | 1063,18086567439 | 15,2192927590977 | 1.72405144281389e-05 |
| 2756 | 24,272651979005 | + | + | NA | NA | NA | NA | + | + | NA | + | NA | + | 0,0981485707984362 | 8 | -523,029380909481 | 1063,18376181896 | 15,2221889036718 | 1.72155669841911e-05 |
| 1932 | 25,2612625845473 | + | + | NA | + | NA | NA | NA | + | + | + | + | NA | 0,113065369399799 | 9 | -521,887459333303 | 1063,19224150125 | 15,2306685859603 | 1.71427302322102e-05 |
| 65 | 26,0391304347826 | NA | NA | NA | NA | NA | NA | + | NA | NA | NA | NA | NA | 0,0229764543599237 | 3 | -528,511010805313 | 1063,20247273845 | 15,2408998231558 | 1.70552584894131e-05 |
| 2009 | 25,5998085445118 | NA | NA | NA | + | + | NA | + | + | + | + | + | NA | 0,112979875067002 | 9 | -521,894058696299 | 1063,20544022724 | 15,2438673119532 | 1.70299716094337e-05 |
| 3694 | 24,2713899293163 | + | NA | + | + | NA | + | + | NA | NA | + | + | + | 0,127933831731835 | 10 | -520,729973274127 | 1063,20597829429 | 15,2444053789957 | 1.7025390592456e-05 |
| 1772 | 24,0419487357438 | + | + | NA | + | NA | + | + | + | NA | + | + | NA | 0,127929037589259 | 10 | -520,730349660508 | 1063,20673106705 | 15,2451581517566 | 1.70189836731289e-05 |
| 3814 | 24,65071847049 | + | NA | + | NA | NA | + | + | + | NA | + | + | + | 0,127890540160335 | 10 | -520,733372004594 | 1063,21277575522 | 15,2512028399292 | 1.69676241007156e-05 |
| 3756 | 24,046449775515 | + | + | NA | + | NA | + | NA | + | NA | + | + | + | 0,127889478188686 | 10 | -520,733455375641 | 1063,21294249731 | 15,2513695820242 | 1.69662095510881e-05 |
| 3826 | 24,5127984429643 | + | NA | NA | NA | + | + | + | + | NA | + | + | + | 0,127868611443229 | 10 | -520,735093517834 | 1063,2162187817 | 15,2546458664085 | 1.69384392394509e-05 |
| 986 | 25,6063334053982 | + | NA | NA | + | + | NA | + | + | + | + | NA | NA | 0,112908022659324 | 9 | -521,899604538582 | 1063,21653191181 | 15,2549589965183 | 1.69357874793732e-05 |
| 1564 | 24,92487425099 | + | + | NA | + | + | NA | NA | NA | NA | + | + | NA | 0,0979068071820148 | 8 | -523,047732703206 | 1063,22046540641 | 15,258892491122 | 1.69025117981981e-05 |

| | | | | | | | | | | | | | | | | | | |
|---|---|---|---|---|---|---|---|---|---|---|---|---|---|---|---|---|---|---|
| **862** | 25,6169294624472 | + | NA | + | + | + | NA | + | NA | + | + | NA | NA | 0,112878905193916 | 9 | -521,901851807541 | 1063,22102644973 | 15,2594535344372 | 1.68977709425583e-05 |
| **4080** | 24,7724848363327 | + | + | + | + | NA | + | + | + | + | + | + | + | 0,172575303608274 | 13 | -517,132336057564 | 1063,22402170862 | 15,2624487933329 | 1.68724832836896e-05 |
| **67** | 24,9715868373804 | NA | + | NA | NA | NA | NA | + | NA | NA | NA | NA | NA | 0,0378027756691876 | 4 | -527,464026879232 | 1063,23108406149 | 15,2695111462033 | 1.68130086378487e-05 |
| **1822** | 25,6077868565261 | + | NA | + | + | + | NA | NA | NA | + | + | + | NA | 0,112811309967272 | 9 | -521,9070684834 | 1063,23145980145 | 15,2698868861553 | 1.68098502750095e-05 |
| **1691** | 24,8960494039216 | NA | + | NA | + | + | NA | NA | + | NA | + | + | NA | 0,0978017396818506 | 8 | -523,055706635112 | 1063,23641327022 | 15,2748403549342 | 1.67682682558162e-05 |
| **989** | 25,7876079219788 | NA | NA | + | + | + | NA | + | + | + | + | NA | NA | 0,112746307237053 | 9 | -521,912084708112 | 1063,24149225087 | 15,2799193355791 | 1.67257394244903e-05 |
| **1946** | 25,5914402963923 | + | NA | NA | + | + | NA | NA | + | + | + | + | NA | 0,112680142355498 | 9 | -521,917190238182 | 1063,25170331101 | 15,2901303957181 | 1.66405632786307e-05 |
| **3659** | 24,036137938623 | NA | + | NA | + | NA | NA | + | NA | NA | + | + | + | 0,0976846290036438 | 8 | -523,064593471203 | 1063,25418694241 | 15,2926140271163 | 1.66199115913576e-05 |
| **4051** | 25,0123160333547 | NA | + | NA | NA | + | NA | + | + | + | + | + | + | 0,127559941530743 | 10 | -520,759321050031 | 1063,26467384609 | 15,3031009308029 | 1.65329939586406e-05 |
| **3868** | 25,0154166027199 | + | + | NA | + | + | NA | NA | NA | + | + | + | + | 0,127506120921928 | 10 | -520,763544557808 | 1063,27312086165 | 15,311547946357 | 1.64633139806221e-05 |
| **2708** | 24,6730605456421 | + | + | NA | NA | + | NA | NA | + | NA | + | NA | + | 0,0975566137034981 | 8 | -523,07430647572 | 1063,27361295144 | 15,3120400361499 | 1.64592637645261e-05 |
| **3995** | 25,0204198347163 | NA | + | NA | + | + | NA | NA | + | + | + | + | + | 0,127491915759448 | 10 | -520,764659247475 | 1063,27535024098 | 15,3137773256917 | 1.6444972718944e-05 |
| **3924** | 24,9939108206042 | + | + | NA | NA | + | NA | + | NA | + | + | + | + | 0,127449909404046 | 10 | -520,767955411308 | 1063,28194256865 | 15,3203696533569 | 1.63908566313812e-05 |
| **169** | 25,6724781139953 | NA | NA | NA | + | NA | + | NA | + | NA | NA | NA | NA | 0,0523389675767697 | 5 | -526,421756738607 | 1063,30152874439 | 15,3399558290994 | 1.62311229516831e-05 |
| **3028** | 24,9965005417906 | + | + | NA | NA | + | NA | + | + | + | + | NA | + | 0,12730390146749 | 10 | -520,779411160219 | 1063,30485406647 | 15,3432811511793 | 1.62041585184844e-05 |
| **926** | 25,7497849272962 | + | NA | + | + | + | NA | NA | + | + | + | NA | NA | 0,112298159182091 | 9 | -521,946658052829 | 1063,3106389403 | 15,3490660250129 | 1.61573567304191e-05 |
| **2972** | 25,0117069817957 | + | + | NA | + | + | NA | NA | + | + | + | NA | + | 0,12725138021143 | 10 | -520,783531497334 | 1063,3130947407 | 15,3515218254095 | 1.61375292844399e-05 |
| **1615** | 24,5286233737101 | NA | + | + | + | NA | NA | + | NA | NA | + | + | NA | 0,0972007914707919 | 8 | -523,101296820918 | 1063,32759364184 | 15,3660207265459 | 1.60209640909053e-05 |
| **1739** | 24,6327641623939 | NA | + | NA | + | NA | NA | + | + | NA | + | + | NA | 0,09718672822237 | 8 | -523,102363348444 | 1063,32972669689 | 15,3681537815983 | 1.60038864002389e-05 |
| **1928** | 25,5991144354457 | + | + | + | NA | NA | NA | NA | + | + | + | + | NA | 0,112169767183735 | 9 | -521,95655991451 | 1063,33044266367 | 15,3688697483744 | 1.59981583001063e-05 |
| **1949** | 25,773664672903 | NA | NA | + | + | + | NA | NA | + | + | + | + | NA | 0,112070819889863 | 9 | -521,964189962006 | 1063,34570275866 | 15,3841298433679 | 1.58765560976554e-05 |
| **2632** | 24,1299077131145 | + | + | + | NA | NA | NA | + | NA | NA | + | NA | + | 0,0970764189234853 | 8 | -523,110728400459 | 1063,34645680092 | 15,3848838856284 | 1.58705714287752e-05 |
| **4039** | 25,0813433069599 | NA | + | + | NA | NA | NA | + | + | + | + | + | + | 0,127030787035578 | 10 | -520,800834513196 | 1063,34770077242 | 15,3861278571328 | 1.5860703228725e-05 |
| **51** | 25,3125966946196 | NA | + | NA | NA | + | + | NA | NA | NA | NA | NA | NA | 0,051974773152642 | 5 | -526,448064734207 | 1063,35414473559 | 15,3925718202986 | 1.58096825729747e-05 |
| **3983** | 25,0915838219716 | NA | + | + | + | NA | NA | NA | + | + | + | + | + | 0,126974890020432 | 10 | -520,805218301801 | 1063,35646834963 | 15,3948954343425 | 1.57913254385352e-05 |
| **2777** | 24,6747381236891 | NA | NA | NA | + | + | NA | + | + | NA | + | NA | + | 0,0969980515101688 | 8 | -523,116670592447 | 1063,35834118489 | 15,3967682696043 | 1.57765450843515e-05 |
| **1868** | 24,8687275546918 | + | + | NA | + | NA | NA | + | NA | + | + | + | NA | 0,111893127162344 | 9 | -521,977890113076 | 1063,3731030608 | 15,411530145506 | 1.56605280679044e-05 |
| **2773** | 24,9553681363088 | NA | NA | + | NA | + | NA | + | + | NA | + | NA | + | 0,0968673674856295 | 8 | -523,126578533683 | 1063,37815706737 | 15,4165841520748 | 1.56210038220236e-05 |
| **3781** | 25,0130540805473 | NA | NA | + | NA | NA | NA | + | + | NA | + | + | + | 0,0968425593634497 | 8 | -523,128459224672 | 1063,38191844934 | 15,4203455340528 | 1.55916531493208e-05 |
| **1608** | 24,6993742601475 | + | + | + | NA | NA | NA | + | NA | NA | + | + | NA | 0,0967636749556923 | 8 | -523,134439067959 | 1063,39387813592 | 15,4323052206264 | 1.5498695719339e-05 |
| **2653** | 24,6738830139644 | NA | NA | + | + | + | NA | + | NA | NA | + | NA | + | 0,096720198760027 | 8 | -523,137734563777 | 1063,40046912755 | 15,4388962122632 | 1.54477039002173e-05 |
| **3725** | 24,9430666643102 | NA | NA | + | + | NA | NA | NA | + | NA | + | + | + | 0,0966365295896568 | 8 | -523,144076241132 | 1063,41315248226 | 15,4515795669743 | 1.53500495197643e-05 |
| **3716** | 24,668702637685 | + | + | NA | NA | NA | NA | NA | + | NA | + | + | + | 0,0965594972774111 | 8 | -523,149914360948 | 1063,4248287219 | 15,4632558066055 | 1.52606951759681e-05 |
| **2002** | 25,620203496523 | + | NA | NA | NA | + | NA | + | + | + | + | + | NA | 0,111523474363482 | 9 | -522,006381654769 | 1063,43008614418 | 15,4685132288921 | 1.52206318969687e-05 |
| **3596** | 24,2702265787571 | + | + | NA | + | NA | NA | NA | NA | NA | + | + | + | 0,0964886615792275 | 8 | -523,155282413077 | 1063,43556482615 | 15,4739919108633 | 1.51789944518036e-05 |
| **1878** | 25,6219008223855 | + | NA | + | NA | + | NA | + | NA | + | + | + | NA | 0,111459898808471 | 9 | -522,011280641522 | 1063,43988411769 | 15,4783112023988 | 1.51462485731539e-05 |
| **974** | 25,6083786556865 | + | NA | + | + | NA | NA | + | + | + | + | NA | NA | 0,111429606930537 | 9 | -522,013614741067 | 1063,44455231678 | 15,4829794014888 | 1.51109369477067e-05 |
| **227** | 25,0488748312064 | NA | + | NA | NA | NA | + | + | + | NA | NA | NA | NA | 0,0663313313005684 | 6 | -525,403274226289 | 1063,45270229873 | 15,4911293834402 | 1.50494853084844e-05 |
| **1679** | 25,0943209731123 | NA | + | + | + | NA | NA | NA | + | NA | + | + | NA | 0,0963755007202104 | 8 | -523,163857066637 | 1063,45271413327 | 15,4911412179824 | 1.50493962568634e-05 |
| **668** | 24,8095089966206 | + | + | NA | + | + | NA | NA | + | NA | + | NA | NA | 0,0962417436472561 | 8 | -523,173990994213 | 1063,47298198843 | 15,5114090731361 | 1.48976569214287e-05 |
| **1560** | 25,261246091211 | + | + | + | NA | + | NA | NA | NA | NA | + | + | NA | 0,0962343071588415 | 8 | -523,174554365924 | 1063,47410873185 | 15,5125358165567 | 1.48892663666917e-05 |
| **2717** | 24,8654166920883 | NA | NA | + | + | + | NA | NA | + | NA | + | NA | + | 0,0962306023329382 | 8 | -523,174835033509 | 1063,47467006702 | 15,5130971517274 | 1.48850880186439e-05 |
| **3673** | 24,6795073276591 | NA | NA | NA | + | + | NA | + | NA | NA | + | + | + | 0,0962271756073275 | 8 | -523,175094631945 | 1063,47518926389 | 15,5136163485993 | 1.4881224374595e-05 |
| **1687** | 25,2644878356574 | NA | + | + | NA | + | NA | NA | + | NA | + | + | NA | 0,0962245533048234 | 8 | -523,175293289039 | 1063,47558657808 | 15,5140136627872 | 1.48782684074278e-05 |
| **3819** | 23,9928223445724 | NA | + | NA | + | NA | + | + | + | NA | + | + | + | 0,126187647885952 | 10 | -520,866928903376 | 1063,47988955278 | 15,5183166374934 | 1.48462924114231e-05 |
| **3812** | 24,1587202497824 | + | + | NA | NA | NA | + | + | + | NA | + | + | + | 0,126174175941688 | 10 | -520,867984463004 | 1063,48200067204 | 15,5204277567484 | 1.48306295325411e-05 |
| **307** | 25,8643343868403 | NA | + | NA | NA | + | + | NA | NA | + | NA | NA | NA | 0,0661187584948906 | 6 | -525,41886089868 | 1063,48387564351 | 15,5223027282232 | 1.48167325440101e-05 |
| **1732** | 24,7279186888938 | + | + | NA | NA | NA | NA | + | + | NA | + | + | NA | 0,096116951163746 | 8 | -523,183444379862 | 1063,49188875972 | 15,5303158444337 | 1.47574872083349e-05 |
| **3785** | 24,658500208629 | NA | NA | NA | + | NA | NA | + | + | NA | + | + | + | 0,0960790171118884 | 8 | -523,186317733666 | 1063,49763546733 | 15,5360625520418 | 1.47151445881235e-05 |
| **1728** | 24,3518600856729 | + | + | + | + | + | + | NA | + | NA | + | + | NA | 0,140993025494518 | 11 | -519,696947581501 | 1063,505895163 | 15,5443222477124 | 1.46544985956566e-05 |
| **2650** | 24,6566013722585 | + | NA | NA | + | + | NA | + | NA | NA | + | NA | + | 0,0959943421100266 | 8 | -523,192731094085 | 1063,51046218817 | 15,5488892728799 | 1.46210730420657e-05 |
| **3613** | 24,8490586798883 | NA | NA | + | + | + | NA | NA | NA | NA | + | + | + | 0,0959608899874084 | 8 | -523,195264622331 | 1063,51552924466 | 15,5539563293714 | 1.45840770255459e-05 |
| **3793** | 24,8971669278262 | NA | NA | NA | NA | + | NA | + | + | NA | + | + | + | 0,0959533809961246 | 8 | -523,195833310033 | 1063,51666662007 | 15,5550937047753 | 1.4575785598137e-05 |
| **991** | 25,2994841438671 | NA | + | + | + | + | NA | + | + | + | + | NA | NA | 0,125914383747068 | 10 | -520,888336633288 | 1063,52270501261 | 15,5611320973167 | 1.45318448069323e-05 |
| **3648** | 24,3172220102397 | + | + | + | + | + | + | NA | NA | NA | + | + | + | 0,140883478682566 | 11 | -519,705678213133 | 1063,52335642627 | 15,5617835109756 | 1.45271124565552e-05 |
| **3669** | 24,8996739706514 | NA | NA | + | NA | + | NA | + | NA | NA | + | + | + | 0,0959001602792401 | 8 | -523,199863804943 | 1063,52472760989 | 15,5631546945945 | 1.4517156200594e-05 |
| **2758** | 24,9004162238151 | + | NA | + | NA | NA | NA | + | + | NA | + | NA | + | 0,0958859016945511 | 8 | -523,200943591401 | 1063,5268871828 | 15,5653142675105 | 1.4501489231932e-05 |
| **3016** | 25,0220062149559 | + | + | + | NA | NA | NA | + | + | + | + | NA | + | 0,125838226311803 | 10 | -520,894301675399 | 1063,53463509683 | 15,5730621815399 | 1.44454197614802e-05 |
| **2639** | 23,9381275317891 | NA | + | + | + | NA | NA | + | NA | NA | + | NA | + | 0,09575949700221 | 8 | -523,210515331263 | 1063,54603066253 | 15,5844577472346 | 1.43633469351501e-05 |
| **2960** | 25,0423646937081 | + | + | + | + | NA | NA | NA | + | + | + | NA | + | 0,125739729169286 | 10 | -520,902015708041 | 1063,55006316211 | 15,588490246824 | 1.43344160156225e-05 |
| **1997** | 25,6548713820467 | NA | NA | + | + | NA | NA | + | + | + | + | + | NA | 0,110725037267498 | 9 | -522,067881973809 | 1063,55308678226 | 15,5915138669729 | 1.43127614739798e-05 |
| **982** | 25,7778275291599 | + | NA | + | NA | + | NA | + | + | + | + | NA | NA | 0,110722406714949 | 9 | -522,068084503201 | 1063,55349184105 | 15,5919189257568 | 1.43098630126236e-05 |
| **1315** | 25,7410177594831 | NA | + | NA | NA | NA | + | NA | NA | + | NA | + | NA | 0,0656182880853253 | 6 | -525,455543354935 | 1063,55724055602 | 15,5956676407329 | 1.42830663348546e-05 |
| **299** | 25,3456090647757 | NA | + | NA | + | NA | + | NA | NA | + | NA | NA | NA | 0,0655491466392121 | 6 | -525,46060959917 | 1063,56737304449 | 15,6058001292024 | 1.42108878237468e-05 |
| **3737** | 24,8341163613887 | NA | NA | NA | + | + | NA | NA | + | NA | + | + | + | 0,0955233296327271 | 8 | -523,228395044966 | 1063,58179008993 | 15,6202171746422 | 1.41088166491814e-05 |
| **2672** | 23,7900568823148 | + | + | + | + | NA | + | + | NA | NA | + | NA | + | 0,125530564931534 | 10 | -520,918394010944 | 1063,58281976792 | 15,6212468526289 | 1.41015547497352e-05 |
| **195** | 25,495067769684 | NA | + | NA | NA | NA | NA | + | + | NA | NA | NA | NA | 0,0503860723204115 | 5 | -526,562708309634 | 1063,58343188644 | 15,6218589711532 | 1.40972394986886e-05 |
| **2770** | 24,835338801222 | + | NA | NA | NA | + | NA | + | + | NA | + | NA | + | 0,0954901893361767 | 8 | -523,230903650801 | 1063,5868073016 | 15,6252343863114 | 1.40734675464369e-05 |
| **2646** | 24,8625255352435 | + | NA | + | NA | + | NA | + | NA | NA | + | NA | + | 0,0954901472002705 | 8 | -523,230906840285 | 1063,58681368057 | 15,6252407652792 | 1.40734226594105e-05 |
| **1887** | 25,2463662457742 | NA | + | + | + | + | NA | + | NA | + | + | + | NA | 0,12546356417822 | 10 | -520,923639579952 | 1063,59331090593 | 15,6317379906443 | 1.40277777415442e-05 |
| **2590** | 24,8208034351552 | + | NA | + | + | + | NA | NA | NA | NA | + | NA | + | 0,0954151320429888 | 8 | -523,236584888678 | 1063,59816977736 | 15,6365968620648 | 1.39937395209391e-05 |
| **864** | 25,2411582348703 | + | + | + | + | + | NA | + | NA | + | + | NA | NA | 0,125422278303917 | 10 | -520,926871700353 | 1063,59977514674 | 15,6382022314478 | 1.39825114673449e-05 |
| **2696** | 24,7287700593332 | + | + | + | NA | NA | NA | NA | + | NA | + | NA | + | 0,0953018787037386 | 8 | -523,245156370525 | 1063,61531274105 | 15,6537398257603 | 1.38743050331649e-05 |
| **3864** | 25,3027362817147 | + | + | + | NA | + | NA | NA | NA | + | + | + | + | 0,125319068348501 | 10 | -520,934950963942 | 1063,61593367392 | 15,654360758625 | 1.38699981957791e-05 |
| **1664** | 24,2638512329569 | + | + | + | + | + | + | + | NA | NA | + | + | NA | 0,140261297643581 | 11 | -519,755243529662 | 1063,62248705932 | 15,6609141440342 | 1.38246248517274e-05 |
| **3991** | 25,3214154443628 | NA | + | + | NA | + | NA | NA | + | + | + | + | + | 0,125275789099996 | 10 | -520,938338575047 | 1063,62270889613 | 15,6611359808353 | 1.38230915314891e-05 |

| | | | | | | | | | | | | | | | | | | | |
|---|---|---|---|---|---|---|---|---|---|---|---|---|---|---|---|---|---|---|---|
| 2049 | 26,0888888888889 | NA | NA | NA | NA | NA | NA | NA | NA | NA | NA | NA | + | 0,0198128408868912 | 3 | -528,732357688581 | 1063,64516650498 | 15,6835935896906 | 1.36687429369507e-05 |
| 295 | 25,427329246988 | NA | + | + | NA | NA | + | NA | NA | + | NA | NA | NA | 0,0650095302776616 | 6 | -525,500136376382 | 1063,64642659892 | 15,6848536836264 | 1.36601336993051e-05 |
| 3666 | 24,8425863788684 | + | NA | NA | NA | + | NA | + | NA | NA | + | + | + | 0,0950940359505386 | 8 | -523,260883981233 | 1063,64676796247 | 15,6851950471746 | 1.36578023624151e-05 |
| 2702 | 24,8273228231814 | + | NA | + | + | NA | NA | NA | + | NA | + | NA | + | 0,0950675087890829 | 8 | -523,262891050802 | 1063,6507821016 | 15,6892091863133 | 1.363041769357e-05 |
| 3610 | 24,8113335025364 | + | NA | NA | + | + | NA | NA | NA | NA | + | + | + | 0,0950647333960005 | 8 | -523,263101036211 | 1063,65120207242 | 15,6896291571318 | 1.36275558052204e-05 |
| 2339 | 25,4585612807756 | NA | + | NA | NA | NA | + | NA | NA | + | NA | NA | + | 0,0649677145159903 | 6 | -525,503198419532 | 1063,65255068522 | 15,6909777699266 | 1.36183697546144e-05 |
| 3818 | 24,226500749622 | + | NA | NA | + | NA | + | + | + | NA | + | + | + | 0,125008495057493 | 10 | -520,959256859476 | 1063,66454546498 | 15,7029725496923 | 1.35369395101674e-05 |
| 2968 | 25,2694412093087 | + | + | + | NA | + | NA | NA | + | + | + | NA | + | 0,124964187488021 | 10 | -520,962723728316 | 1063,67147920266 | 15,7099062873735 | 1.34900899739052e-05 |
| 1684 | 25,0878579343835 | + | + | NA | NA | + | NA | NA | + | NA | + | + | NA | 0,0949149031870901 | 8 | -523,274436189657 | 1063,67387237931 | 15,7122994640229 | 1.34739575436076e-05 |
| 2005 | 25,8354336928516 | NA | NA | + | NA | + | NA | + | + | + | + | + | NA | 0,109939527160113 | 9 | -522,12833274581 | 1063,67398832627 | 15,7124154109747 | 1.34731764340964e-05 |
| 4043 | 24,7825974003662 | NA | + | NA | + | NA | NA | + | + | + | + | + | + | 0,124902357602131 | 10 | -520,967561346678 | 1063,68115443939 | 15,7195815240964 | 1.34249876639785e-05 |
| 733 | 25,1693178647249 | NA | NA | + | + | + | NA | + | + | NA | + | NA | NA | 0,0948665019913521 | 8 | -523,278097500237 | 1063,68119500047 | 15,7196220851833 | 1.34247154006932e-05 |
| 2714 | 24,7772066538944 | + | NA | NA | + | + | NA | NA | + | NA | + | NA | + | 0,0947822263121388 | 8 | -523,284472070712 | 1063,69394414142 | 15,7323712261336 | 1.33394107852678e-05 |
| 1934 | 25,6334038546925 | + | NA | + | + | NA | NA | NA | + | + | + | + | NA | 0,109752928753215 | 9 | -522,142685025552 | 1063,70269288575 | 15,7411199704582 | 1.32811866778689e-05 |
| 2762 | 24,5242371555434 | + | NA | NA | + | NA | NA | + | + | NA | + | NA | + | 0,0946847313910255 | 8 | -523,291845798062 | 1063,70869159612 | 15,7471186808332 | 1.32414113616794e-05 |
| 1994 | 25,3842297715417 | + | NA | NA | + | NA | NA | + | + | + | + | + | NA | 0,109660363999001 | 9 | -522,149803559307 | 1063,71692995326 | 15,7553570379696 | 1.31869798073961e-05 |
| 2113 | 25,3386959130609 | NA | NA | NA | NA | NA | NA | + | NA | NA | NA | NA | + | 0,0343291321779243 | 4 | -527,710762955801 | 1063,72455621463 | 15,7629832993412 | 1.31367918774185e-05 |
| 664 | 25,1008969765408 | + | + | + | NA | + | NA | NA | + | NA | + | NA | NA | 0,0945735082567537 | 8 | -523,300256847273 | 1063,72551369455 | 15,7639407792558 | 1.3130504275419e-05 |
| 3919 | 24,6888848018382 | NA | + | + | + | NA | NA | + | NA | + | + | + | + | 0,124613821271682 | 10 | -520,990132137494 | 1063,72629602102 | 15,7647231057283 | 1.31253691092825e-05 |
| 43 | 24,7676480755077 | NA | + | NA | + | NA | + | NA | NA | NA | NA | NA | NA | 0,0493882180929024 | 5 | -526,634617326898 | 1063,72724992097 | 15,7656770056813 | 1.31191104574452e-05 |
| 2576 | 24,2138259369128 | + | + | + | + | NA | NA | NA | NA | NA | + | NA | + | 0,0944908987229983 | 8 | -523,306503376132 | 1063,73800675226 | 15,7764338369736 | 1.30487398393497e-05 |
| 1942 | 25,7754222388106 | + | NA | + | NA | + | NA | NA | + | + | + | + | NA | 0,109494574534645 | 9 | -522,162551465153 | 1063,74242576495 | 15,7808528496619 | 1.30199403938871e-05 |
| 3778 | 24,8002390422147 | + | NA | NA | NA | NA | NA | + | + | NA | + | + | + | 0,0944325344004663 | 8 | -523,310916257353 | 1063,74683251471 | 15,7852595994148 | 1.299128416613e-05 |
| 3692 | 23,8314343250186 | + | + | NA | + | NA | + | + | NA | NA | + | + | + | 0,124416891163416 | 10 | -521,005532750991 | 1063,75709724801 | 15,7955243327224 | 1.29247789430657e-05 |
| 3988 | 25,1977586260853 | + | + | NA | NA | + | NA | NA | + | + | + | + | + | 0,124377175903437 | 10 | -521,008638201754 | 1063,76330814954 | 15,8017352342483 | 1.28847039361225e-05 |
| 2700 | 24,3320651905087 | + | + | NA | + | NA | NA | NA | + | NA | + | NA | + | 0,0942938759517261 | 8 | -523,321398975439 | 1063,76779795088 | 15,8062250355877 | 1.28558114980884e-05 |
| 165 | 25,7213522348652 | NA | NA | + | NA | NA | + | NA | + | NA | NA | NA | NA | 0,0490746034119763 | 5 | -526,657201963485 | 1063,77241919414 | 15,8108462788541 | 1.28261408739221e-05 |
| 2083 | 24,8568774332125 | NA | + | NA | NA | NA | + | NA | NA | NA | NA | NA | + | 0,0490440173510472 | 5 | -526,659404188533 | 1063,77682364424 | 15,8152507289512 | 1.27979259044724e-05 |
| 425 | 26,247593367919 | NA | NA | NA | + | NA | + | NA | + | + | NA | NA | NA | 0,064105295809056 | 6 | -525,566320258439 | 1063,77879436303 | 15,8172214477415 | 1.27853215588691e-05 |
| 716 | 24,4747933126115 | + | + | NA | + | NA | NA | + | + | NA | + | NA | NA | 0,0941504962881639 | 8 | -523,332236935318 | 1063,78947387064 | 15,8279009553446 | 1.27172330392209e-05 |
| 36 | 25,0752167794664 | + | + | NA | NA | NA | + | NA | NA | NA | NA | NA | NA | 0,0487608193139116 | 5 | -526,679791350677 | 1063,81759796853 | 15,8560250532387 | 1.25396541733167e-05 |
| 3661 | 24,5546680575322 | NA | NA | + | + | NA | NA | + | NA | NA | + | + | + | 0,0939232390675377 | 8 | -523,349411623906 | 1063,82382324781 | 15,8622503325221 | 1.25006834310464e-05 |
| 39 | 24,8629339062425 | NA | + | + | NA | NA | + | NA | NA | NA | NA | NA | NA | 0,0487154285364594 | 5 | -526,683058426218 | 1063,82413211961 | 15,8625592043209 | 1.24987530258235e-05 |
| 3722 | 24,7549102479597 | + | NA | NA | + | NA | NA | NA | + | NA | + | + | + | 0,0938024184907669 | 8 | -523,358540739407 | 1063,84208147881 | 15,8805085635242 | 1.23870825747947e-05 |
| 1990 | 25,6345740828319 | + | NA | + | NA | NA | NA | + | + | + | + | + | NA | 0,108813915894483 | 9 | -522,214863911169 | 1063,84705065698 | 15,8854777416936 | 1.23563439668559e-05 |
| 3020 | 24,732191793138 | + | + | NA | + | NA | NA | + | + | + | + | NA | + | 0,123824835107171 | 10 | -521,051812734243 | 1063,84965721452 | 15,8880842992262 | 1.23402506954225e-05 |
| 3856 | 24,8536953583189 | + | + | + | + | NA | NA | NA | NA | + | + | + | + | 0,123789504705269 | 10 | -521,05457346142 | 1063,85517866887 | 15,8936057535802 | 1.23062296130278e-05 |
| 481 | 26,3387991750729 | NA | NA | NA | NA | NA | + | + | + | + | NA | NA | NA | 0,0635716894120852 | 6 | -525,605346676969 | 1063,85684720009 | 15,8952742848003 | 1.22959672302443e-05 |
| 292 | 25,6131533918514 | + | + | NA | NA | NA | + | NA | NA | + | NA | NA | NA | 0,0635398624324327 | 6 | -525,607673707122 | 1063,8615012604 | 15,8999283451078 | 1.22673874096787e-05 |
| 988 | 25,1889715788614 | + | + | NA | + | + | NA | + | + | + | + | NA | NA | 0,123737097878925 | 10 | -521,058668340112 | 1063,86336842626 | 15,9017955109655 | 1.22559401305353e-05 |
| 2011 | 25,2038105794306 | NA | + | NA | + | + | NA | + | + | + | + | + | NA | 0,123726246240252 | 10 | -521,059516217076 | 1063,86506418018 | 15,903491264894 | 1.22455530053476e-05 |
| 1884 | 25,1706071313168 | + | + | NA | + | + | NA | + | NA | + | + | + | NA | 0,123630569871154 | 10 | -521,066991296525 | 1063,88001433908 | 15,9184414237902 | 1.21543577941123e-05 |
| 2209 | 25,8921680762519 | NA | NA | NA | NA | NA | + | NA | + | NA | NA | NA | + | 0,0482360590849629 | 5 | -526,717552318389 | 1063,89311990395 | 15,9315469886635 | 1.20749733105123e-05 |
| 1629 | 25,0727432857891 | NA | NA | + | + | + | NA | + | NA | NA | + | + | NA | 0,0934067216176478 | 8 | -523,388430794834 | 1063,90186158967 | 15,9402886743781 | 1.20223106735617e-05 |
| 984 | 25,3838552144326 | + | + | + | NA | + | NA | + | + | + | + | NA | NA | 0,123454911372033 | 10 | -521,080713160172 | 1063,90745806638 | 15,945885151086 | 1.19887164070345e-05 |
| 483 | 25,6026432748771 | NA | + | NA | NA | NA | + | + | + | + | NA | NA | NA | 0,0782921171520006 | 7 | -524,52049876944 | 1063,90921459314 | 15,9476416778537 | 1.19781917787697e-05 |
| 1741 | 25,207947602191 | NA | NA | + | + | NA | NA | + | + | NA | + | + | NA | 0,0933442216844958 | 8 | -523,39315070729 | 1063,91130141458 | 15,949728499289 | 1.19657001231864e-05 |
| 606 | 25,0504630400443 | + | NA | + | + | + | NA | + | NA | NA | + | NA | NA | 0,0933312291258607 | 8 | -523,394131847339 | 1063,91326369468 | 15,9516907793877 | 1.19539658529994e-05 |
| 3733 | 25,1651007223161 | NA | NA | + | NA | + | NA | NA | + | NA | + | + | + | 0,0932889779020178 | 8 | -523,397322374066 | 1063,91964474813 | 15,9580718328409 | 1.1915887183265e-05 |
| 3912 | 24,8375313829295 | + | + | + | NA | NA | NA | + | NA | + | + | + | + | 0,123367095683628 | 10 | -521,087572002211 | 1063,92117575045 | 15,9596028351637 | 1.19067690482042e-05 |
| 2007 | 25,4325269153015 | NA | + | + | NA | + | NA | + | + | + | + | + | NA | 0,12335058551589 | 10 | -521,088861451735 | 1063,9237546495 | 15,962181734212 | 1.18914257648413e-05 |
| 3598 | 24,7126163583791 | + | NA | + | + | NA | NA | NA | NA | NA | + | + | + | 0,0932592128705496 | 8 | -523,399569938758 | 1063,92413987752 | 15,9625669622249 | 1.18891355302552e-05 |
| 1753 | 25,0675487786921 | NA | NA | NA | + | + | NA | + | + | NA | + | + | NA | 0,0932317750628719 | 8 | -523,401641708901 | 1063,9282834178 | 15,9667105025119 | 1.1864529472081e-05 |
| 1880 | 25,3558418131028 | + | + | + | NA | + | NA | + | NA | + | + | + | NA | 0,123318980007089 | 10 | -521,09132978465 | 1063,92869131533 | 15,9671184000422 | 1.18621099626824e-05 |
| 730 | 25,0206442751676 | + | NA | NA | + | + | NA | + | + | NA | + | NA | NA | 0,0931633947029336 | 8 | -523,406804690896 | 1063,93860938179 | 15,977036466501 | 1.18034309809696e-05 |
| 3783 | 24,6539874679537 | NA | + | + | NA | NA | NA | + | + | NA | + | + | + | 0,108092962974773 | 9 | -522,270229666057 | 1063,95778216676 | 15,9962092514681 | 1.16908192912674e-05 |
| 3760 | 24,3609423433601 | + | + | + | + | NA | + | NA | + | NA | + | + | + | 0,138143525336576 | 11 | -519,923685063831 | 1063,95937012766 | 15,997797212372 | 1.16815406932832e-05 |
| 3606 | 25,081263513183 | + | NA | + | NA | + | NA | NA | NA | NA | + | + | + | 0,0930078278216671 | 8 | -523,418549143395 | 1063,96209828679 | 16,0005253714985 | 1.1665617005414e-05 |
| 4036 | 24,8867307546417 | + | + | NA | NA | NA | NA | + | + | + | + | + | + | 0,123105052968638 | 10 | -521,108034760274 | 1063,96210126658 | 16,0005283512892 | 1.16655996248786e-05 |
| 656 | 24,9223378506601 | + | + | + | + | NA | NA | NA | + | NA | + | NA | NA | 0,0929777000161557 | 8 | -523,420823395944 | 1063,96664679189 | 16,0050738765983 | 1.16391165919318e-05 |
| 3654 | 24,7382716919498 | + | NA | + | NA | NA | NA | + | NA | NA | + | + | + | 0,0929678574350453 | 8 | -523,421566364834 | 1063,96813272967 | 16,0065598143776 | 1.16304723020179e-05 |
| 1776 | 24,3437132352026 | + | + | + | + | NA | + | + | + | NA | + | + | NA | 0,138073650081718 | 11 | -519,929235688695 | 1063,97047137739 | 16,0088984620993 | 1.16168804614228e-05 |
| 718 | 25,099836279303 | + | NA | + | + | NA | NA | + | + | NA | + | NA | NA | 0,0929443751185769 | 8 | -523,423338898894 | 1063,97167779779 | 16,0101048824968 | 1.16098751536999e-05 |
| 1552 | 24,8436721006222 | + | + | + | + | NA | NA | NA | NA | NA | + | + | NA | 0,0929311634268903 | 8 | -523,424336147188 | 1063,97367229438 | 16,0120993790858 | 1.15983029966191e-05 |
| 3671 | 24,5364585174906 | NA | + | + | NA | + | NA | + | NA | NA | + | + | + | 0,107988962184139 | 9 | -522,278212740062 | 1063,97374831477 | 16,0121753994797 | 1.15978621512165e-05 |
| 113 | 25,5053844514151 | NA | NA | NA | NA | + | + | + | NA | NA | NA | NA | NA | 0,047671173008973 | 5 | -526,758177431693 | 1063,97437013056 | 16,0127972152702 | 1.15942568447938e-05 |
| 1185 | 26,1404578043449 | NA | NA | NA | NA | NA | + | NA | + | NA | NA | + | NA | 0,0476313164302018 | 5 | -526,761042901732 | 1063,98010107064 | 16,0185281553488 | 1.15610814034937e-05 |
| 2752 | 24,2605926558094 | + | + | + | + | + | + | NA | + | NA | + | NA | + | 0,137985477996001 | 11 | -519,936239102521 | 1063,98447820504 | 16,0229052897516 | 1.15358068675198e-05 |
| 3980 | 24,9043014176215 | + | + | NA | + | NA | NA | NA | + | + | + | + | + | 0,122951369875128 | 10 | -521,120032937476 | 1063,98609762098 | 16,0245247056941 | 1.1526470013317e-05 |
| 3768 | 24,516875395423 | + | + | + | NA | + | + | NA | + | NA | + | + | + | 0,137947864644133 | 11 | -519,939226472696 | 1063,99045294539 | 16,0288800301023 | 1.15013965658806e-05 |
| 3704 | 24,4716568223822 | + | + | + | NA | + | + | + | NA | NA | + | + | + | 0,137859844268257 | 11 | -519,946216817418 | 1064,00443363484 | 16,0428607195456 | 1.14212781928754e-05 |
| 2896 | 24,6126443060711 | + | + | + | + | NA | NA | + | NA | + | + | NA | + | 0,12282894292137 | 10 | -521,129589416097 | 1064,00521057823 | 16,0436376629361 | 1.14168422112563e-05 |
| 592 | 24,3317065368721 | + | + | + | + | NA | NA | + | NA | NA | + | NA | NA | 0,0926623844130974 | 8 | -523,444621042066 | 1064,01424208413 | 16,0526691688422 | 1.13654028035681e-05 |
| 2796 | 23,8752897738023 | + | + | NA | + | NA | + | + | + | NA | + | NA | + | 0,12271318029826 1 | 10 | -521,13862446011 | 1064,02328066625 | 16,061707750961 | 1.13141551290933e-05 |
| 162 | 25,9717070166779 | + | NA | NA | NA | NA | + | NA | + | NA | NA | NA | NA | 0,0473147027170755 | 5 | -526,783801438976 | 1064,02561814513 | 16,0640452298371 | 1.13009395540717e-05 |
| 177 | 25,9746443059252 | NA | NA | NA | NA | + | + | NA | + | NA | NA | NA | NA | 0,0473032129316688 | 5 | -526,784627195057 | 1064,02726965729 | 16,0656967419998 | 1.12916115863533e-05 |

| | | | | | | | | | | | | | | | | | | | |
|---|---|---|---|---|---|---|---|---|---|---|---|---|---|---|---|---|---|---|---|
| **3730** | 25,0311313455711 | + | NA | NA | NA | + | NA | NA | + | NA | + | + | + | 0,0925275220998479 | 8 | -523,454796909712 | 1064,03459381942 | 16,073020904134 | 1.12503364119911e-05 |
| **2710** | 25,0543434549422 | + | NA | + | NA | + | NA | NA | + | NA | + | NA | + | 0,0925222263060404 | 8 | -523,455196466385 | 1064,03539293277 | 16,0738200174801 | 1.12458421629168e-05 |
| **1121** | 25,456903790111 | NA | NA | NA | NA | NA | + | + | NA | NA | NA | + | NA | 0,047212314000253 | 5 | -526,791159633339 | 1064,04033453385 | 16,0787616185621 | 1.12180902288819e-05 |
| **726** | 25,2617344524222 | + | NA | + | NA | + | NA | + | + | NA | + | NA | NA | 0,0924852179664715 | 8 | -523,457988603897 | 1064,04097720779 | 16,079404292504 | 1.12144860208615e-05 |
| **3775** | 24,269656561719 | NA | + | + | + | + | + | NA | + | NA | + | + | + | 0,137587856681983 | 11 | -519,96781284313 | 1064,04762568626 | 16,0860527709688 | 1.11772682810685e-05 |
| **3718** | 25,1133206958746 | + | NA | + | NA | NA | NA | NA | + | NA | + | + | + | 0,0923968678431814 | 8 | -523,464653822268 | 1064,05430764454 | 16,0927347292461 | 1.11399875726264e-05 |
| **101** | 25,0239863449262 | NA | NA | + | NA | NA | + | + | NA | NA | NA | NA | NA | 0,0471090121441708 | 5 | -526,798582649787 | 1064,05518056675 | 16,0936076514588 | 1.11351264622458e-05 |
| **1626** | 24,9933912204161 | + | NA | NA | + | + | NA | + | NA | NA | + | + | NA | 0,0923764992787322 | 8 | -523,466190354393 | 1064,05738070879 | 16,0958077934961 | 1.11228837674872e-05 |
| **2688** | 24,1992120714033 | + | + | + | + | + | + | + | NA | NA | + | NA | + | 0,137498026377675 | 11 | -519,974943943669 | 1064,06188788734 | 16,100314972047 | 1.10978455794092e-05 |
| **2145** | 25,084652193027 | NA | NA | NA | NA | NA | + | + | NA | NA | NA | NA | + | 0,0470614945052259 | 5 | -526,801996879968 | 1064,06200902711 | 16,1004361118214 | 1.10971734045114e-05 |
| **1749** | 25,3562122597484 | NA | NA | + | NA | + | NA | + | + | NA | + | + | NA | 0,0922955880320687 | 8 | -523,472293670969 | 1064,06958734194 | 16,1080144266477 | 1.10552040321351e-05 |
| **1784** | 24,4575214121738 | + | + | + | NA | + | + | + | + | NA | + | + | NA | 0,137372074831069 | 11 | -519,984941247602 | 1064,0818824952 | 16,1203095799135 | 1.09874497937263e-05 |
| **3976** | 25,2562603392936 | + | + | + | NA | NA | NA | NA | + | + | + | + | + | 0,122306805491592 | 10 | -521,170331779777 | 1064,08669530559 | 16,125122390295 | 1.09610413249915e-05 |
| **1870** | 25,2973180744979 | + | NA | + | + | NA | NA | + | NA | + | + | + | NA | 0,107197793573702 | 9 | -522,338912199426 | 1064,0951472335 | 16,1335743182074 | 1.09148180970683e-05 |
| **2636** | 23,843703410879 | + | + | NA | + | NA | NA | + | NA | NA | + | NA | + | 0,0921229908471004 | 8 | -523,48531124626 | 1064,09562249252 | 16,1340495772299 | 1.09122247223216e-05 |
| **928** | 25,4619803723321 | + | + | + | + | + | NA | NA | + | + | + | NA | NA | 0,122147129519191 | 10 | -521,182786452881 | 1064,11160465179 | 16,1500317365035 | 1.08253717522343e-05 |
| **1824** | 25,4553841145667 | + | + | + | + | + | NA | NA | NA | + | + | + | NA | 0,122145031227889 | 10 | -521,182950103836 | 1064,1119319537 | 16,1503590384134 | 1.0823600314762e-05 |
| **2638** | 24,3923663481008 | + | NA | + | + | NA | NA | + | NA | NA | + | NA | + | 0,0919972455255472 | 8 | -523,49479361633 | 1064,11458723266 | 16,1530143173698 | 1.08092400104509e-05 |
| **193** | 26,5882951405241 | NA | NA | NA | NA | NA | NA | + | + | NA | NA | NA | NA | 0,0315255180565698 | 4 | -527,909260130016 | 1064,12155056306 | 16,1599776477719 | 1.07716712944524e-05 |
| **105** | 25,0862526365232 | NA | NA | NA | + | NA | + | + | NA | NA | NA | NA | NA | 0,0466436196808241 | 5 | -526,832014632092 | 1064,12204453136 | 16,1604716160682 | 1.0769011190909e-05 |
| **1951** | 25,5030870726818 | NA | + | + | + | + | NA | NA | + | + | + | + | NA | 0,122078417361769 | 10 | -521,18814528164 | 1064,12232230931 | 16,1607493940203 | 1.07675155978348e-05 |
| **2004** | 25,2797939633024 | + | + | NA | NA | + | NA | + | + | + | + | + | NA | 0,122031803469214 | 10 | -521,191780438487 | 1064,12959262301 | 16,1680197077151 | 1.07284450465952e-05 |
| **1612** | 24,3941837818188 | + | + | NA | + | NA | NA | + | NA | NA | + | + | NA | 0,0918537345100404 | 8 | -523,505614081755 | 1064,13622816351 | 16,1746552482189 | 1.06929095127358e-05 |
| **3658** | 24,4347757017879 | + | NA | NA | + | NA | NA | + | NA | NA | + | + | + | 0,0916871549896794 | 8 | -523,518171726762 | 1064,16134345352 | 16,1997705382325 | 1.05594713391256e-05 |
| **421** | 26,2245529298921 | NA | NA | + | NA | NA | + | NA | + | + | NA | NA | NA | 0,0614696114669478 | 6 | -525,758870582569 | 1064,16389501129 | 16,2023220960002 | 1.05460083782733e-05 |
| **3711** | 24,1872821111882 | NA | + | + | + | + | + | + | NA | NA | + | + | + | 0,136820924334137 | 11 | -520,028671226712 | 1064,16934245342 | 16,2077695381329 | 1.05173230762023e-05 |
| **1672** | 25,2489794433559 | + | + | + | NA | NA | NA | NA | + | NA | + | + | NA | 0,0916247902430316 | 8 | -523,522872518898 | 1064,1707450378 | 16,2091721225056 | 1.05099499453692e-05 |
| **1622** | 25,1852097458158 | + | NA | + | NA | + | NA | + | NA | NA | + | + | NA | 0,0916236094022129 | 8 | -523,522961522598 | 1064,1709230452 | 16,2093501299046 | 1.05090145625696e-05 |
| **98** | 25,3022281561898 | + | NA | NA | NA | NA | + | + | NA | NA | NA | NA | NA | 0,0462761994415183 | 5 | -526,858397150249 | 1064,17480956767 | 16,2132366523824 | 1.04886126314721e-05 |
| **53** | 25,4866391820448 | NA | NA | + | NA | + | + | NA | NA | NA | NA | NA | NA | 0,0461960593120255 | 5 | -526,864150242546 | 1064,18631575227 | 16,2247428369762 | 1.04284439189704e-05 |
| **1746** | 25,1564738449008 | + | NA | NA | NA | + | NA | + | + | NA | + | + | NA | 0,0915170035412838 | 8 | -523,530996266348 | 1064,1869925327 | 16,2254196174056 | 1.04249156325965e-05 |
| **768** | 24,1671173424473 | + | + | + | + | + | + | + | + | NA | + | NA | NA | 0,136676055807271 | 11 | -520,040160910307 | 1064,19232182061 | 16,2307489053237 | 1.03971739214385e-05 |
| **1693** | 25,3247059514638 | NA | NA | + | + | + | NA | NA | + | NA | + | + | NA | 0,0913443933500249 | 8 | -523,5440036718 | 1064,2130073436 | 16,251434428309 | 1.02901926260541e-05 |
| **2115** | 24,4858231093753 | NA | + | NA | NA | NA | NA | + | NA | NA | NA | NA | + | 0,0459387564005037 | 5 | -526,882618215054 | 1064,22325169728 | 16,2616787819934 | 1.02376191998226e-05 |
| **1791** | 24,1902978543379 | NA | + | + | + | + | + | + | + | NA | + | + | NA | 0,136457501335985 | 11 | -520,057491061863 | 1064,22698212373 | 16,2654092084351 | 1.02185416545166e-05 |
| **3668** | 24,51998662355 | + | + | NA | NA | + | NA | + | NA | NA | + | + | + | 0,106289332955752 | 9 | -522,408544175554 | 1064,23441118575 | 16,272838270462 | 1.01806549738167e-05 |
| **670** | 25,24064309528 | + | NA | + | + | + | NA | NA | + | NA | + | NA | NA | 0,0911942439868471 | 8 | -523,555316485699 | 1064,2356329714 | 16,2740600561081 | 1.01744375840321e-05 |
| **2775** | 24,5406383555709 | NA | + | + | NA | + | NA | + | + | NA | + | NA | + | 0,106268317978008 | 9 | -522,410154100867 | 1064,23763103638 | 16,2760581210898 | 1.01642780659985e-05 |
| **1676** | 24,8504034020201 | + | + | NA | + | NA | NA | NA | + | NA | + | + | NA | 0,091129934803382 | 8 | -523,560161208399 | 1064,2453224168 | 16,2837495015076 | 1.01252644665505e-05 |
| **1999** | 25,3151129858997 | NA | + | + | + | NA | NA | + | + | + | + | + | NA | 0,121278910852281 | 10 | -521,250467618917 | 1064,24696698387 | 16,2853940685761 | 1.01169420504614e-05 |
| **1065** | 25,4657919760521 | NA | NA | NA | + | NA | + | NA | NA | NA | NA | + | NA | 0,0457029108553792 | 5 | -526,899541707983 | 1064,25709868314 | 16,2955257678504 | 1.00658207390108e-05 |
| **2648** | 24,5212886491212 | + | + | + | NA | + | NA | + | NA | NA | + | NA | + | 0,106136690088768 | 9 | -522,420237051588 | 1064,25779693782 | 16,2962240225318 | 1.00623070991742e-05 |
| **1061** | 25,463752910743 | NA | NA | + | NA | NA | + | NA | NA | NA | NA | + | NA | 0,0455604199222636 | 5 | -526,90976435827 | 1064,27754398372 | 16,3159710684254 | 9.96344553821538e-06 |
| **57** | 25,4731837606838 | NA | NA | NA | + | + | + | NA | NA | NA | NA | NA | NA | 0,0455228585510506 | 5 | -526,912458849438 | 1064,28293296605 | 16,3213600507606 | 9.93663525846533e-06 |
| **1566** | 25,2153290876397 | + | NA | + | + | + | NA | NA | NA | NA | + | + | NA | 0,0908732758262622 | 8 | -523,579493162069 | 1064,28398632414 | 16,322413408848 | 9.93140321883383e-06 |
| **3795** | 24,4994336686152 | NA | + | NA | NA | + | NA | + | + | NA | + | + | + | 0,105903800517777 | 9 | -522,438073207627 | 1064,2934692499 | 16,3318963346092 | 9.88442529916176e-06 |
| **1** | 27,2678832116789 | NA | NA | NA | NA | NA | NA | NA | NA | NA | NA | NA | NA | 0 | 2 | -530,102559176374 | 1064,29467059155 | 16,3330976762625 | 9.87848979606653e-06 |
| **1734** | 25,2697964165883 | + | NA | + | NA | NA | NA | + | + | NA | + | + | NA | 0,0907876939310575 | 8 | -523,585938110903 | 1064,29687622181 | 16,3353033065164 | 9.8676016529959e-06 |
| **3916** | 24,5706443594505 | + | + | NA | + | NA | NA | + | NA | + | + | + | + | 0,120899630675747 | 10 | -521,280013072416 | 1064,30605789086 | 16,3444849755729 | 9.82240495128531e-06 |
| **115** | 24,8815751648382 | NA | + | NA | NA | + | + | + | NA | NA | NA | NA | NA | 0,0604712260980767 | 6 | -525,831666668856 | 1064,30948718387 | 16,3479142685751 | 9.80557742975128e-06 |
| **1948** | 25,3561118310245 | + | + | NA | + | + | NA | NA | + | + | + | + | NA | 0,120845826000722 | 10 | -521,284203357669 | 1064,31443846137 | 16,3528655460793 | 9.7813323856474e-06 |
| **3675** | 24,3423776490043 | NA | + | NA | + | + | NA | + | NA | NA | + | + | + | 0,10575844110907 | 9 | -522,449203396815 | 1064,31572962827 | 16,3541567129841 | 9.77501975720018e-06 |
| **976** | 25,2450097737675 | + | + | + | + | NA | NA | + | + | + | + | NA | NA | 0,120799949301838 | 10 | -521,28777601306 | 1064,32158377215 | 16,3600108568605 | 9.74644940530992e-06 |
| **83** | 25,3378229106595 | NA | + | NA | NA | + | NA | + | NA | NA | NA | NA | NA | 0,0452254602053206 | 5 | -526,933789186261 | 1064,3255936397 | 16,3640207244066 | 9.72692799583587e-06 |
| **3831** | 24,3858053757497 | NA | + | + | NA | + | + | + | + | NA | + | + | + | 0,135739109759764 | 11 | -520,114424638322 | 1064,34084927664 | 16,3792763613542 | 9.65301501056992e-06 |
| **1631** | 24,7100783289366 | NA | + | + | + | + | NA | + | NA | NA | + | + | NA | 0,105587069593482 | 9 | -522,462323014012 | 1064,34196886267 | 16,3803959473798 | 9.64761283240617e-06 |
| **45** | 25,0295409547717 | NA | NA | + | + | NA | + | NA | NA | NA | NA | NA | NA | 0,0450938585579718 | 5 | -526,943225947332 | 1064,34446716184 | 16,3828942465484 | 9.63556904465955e-06 |
| **3037** | 25,3655792851258 | NA | NA | + | + | + | NA | + | + | + | + | NA | + | 0,120578010768797 | 10 | -521,305056881774 | 1064,35614550958 | 16,3945725942888 | 9.57946922935538e-06 |
| **1123** | 24,8427984346789 | NA | + | NA | NA | NA | + | + | NA | NA | NA | + | NA | 0,060127079424075 | 6 | -525,856741797106 | 1064,35963744037 | 16,3980645250754 | 9.56275840010465e-06 |
| **1751** | 24,9538870845433 | NA | + | + | NA | + | NA | + | + | NA | + | + | NA | 0,105374937094608 | 9 | -522,478559673134 | 1064,37444218091 | 16,4128692656227 | 9.49223267240291e-06 |
| **2655** | 24,3083704650856 | NA | + | + | + | + | NA | + | NA | NA | + | NA | + | 0,105344948228498 | 9 | -522,480854715808 | 1064,37903226626 | 16,4174593509713 | 9.47047257307499e-06 |
| **323** | 25,4876233528144 | NA | + | NA | NA | NA | NA | + | NA | + | NA | NA | NA | 0,0448382135571818 | 5 | -526,961553772036 | 1064,38112281125 | 16,4195498959577 | 9.46057852048783e-06 |
| **1690** | 25,1627681897087 | + | NA | NA | + | + | NA | NA | + | NA | + | + | NA | 0,0902087135306181 | 8 | -523,629523691039 | 1064,38404738208 | 16,4224744667872 | 9.44675456425048e-06 |
| **2085** | 25,0995525126415 | NA | NA | + | NA | NA | + | NA | NA | NA | NA | NA | + | 0,0447447903300234 | 5 | -526,968250291823 | 1064,39451585082 | 16,4329429355303 | 9.39743721867995e-06 |
| **2089** | 25,108888032741 | NA | NA | NA | + | NA | + | NA | NA | NA | NA | NA | + | 0,0446829434711918 | 5 | -526,972683076724 | 1064,40338142062 | 16,4418085053339 | 9.35587259244544e-06 |
| **3708** | 24,1832229347797 | + | + | NA | + | + | + | + | NA | NA | + | + | + | 0,135300656110637 | 11 | -520,149149487779 | 1064,41029897556 | 16,4487260602673 | 9.32356860949052e-06 |
| **3105** | 25,6445361536433 | NA | NA | NA | NA | NA | + | NA | NA | NA | NA | + | + | 0,0445893849569385 | 5 | -526,979388203904 | 1064,41679167498 | 16,4552187596937 | 9.29335012163755e-06 |
| **1738** | 24,9511516626752 | + | NA | NA | + | NA | NA | + | + | NA | + | + | NA | 0,0899718701714249 | 8 | -523,647345243126 | 1064,41969048625 | 16,4581175709609 | 9.27989004452847e-06 |
| **3615** | 24,6178066116248 | NA | + | + | + | + | NA | NA | NA | NA | + | + | + | 0,105026598911352 | 9 | -522,505213193767 | 1064,42774922218 | 16,4661763068889 | 9.24257318504182e-06 |
| **2051** | 25,3683469859273 | NA | + | NA | NA | NA | NA | NA | NA | NA | NA | NA | + | 0,0293376363570788 | 4 | -528,063764568018 | 1064,43055943907 | 16,4689865237765 | 9.22959548708871e-06 |
| **735** | 24,6964919325021 | NA | + | + | + | + | NA | + | + | NA | + | NA | NA | 0,104979456511217 | 9 | -522,508819555348 | 1064,43496194534 | 16,4733890300504 | 9.20930115574499e-06 |
| **2767** | 24,3749794936056 | NA | + | + | + | NA | NA | + | + | NA | + | NA | + | 0,104970199594528 | 9 | -522,509527680884 | 1064,43637819641 | 16,4748052811231 | 9.20278212284405e-06 |
| **1624** | 24,9203680316203 | + | + | + | NA | + | NA | + | NA | NA | + | + | NA | 0,104923297220893 | 9 | -522,513115455104 | 1064,44355374485 | 16,4819808295622 | 9.16982377730029e-06 |
| **3772** | 24,1752205404896 | + | + | NA | + | + | + | NA | + | NA | + | + | + | 0,135087035370219 | 11 | -520,166061543916 | 1064,44412308783 | 16,482550172542 | 9.1672137614196e-06 |
| **42** | 25,3013490627618 | + | NA | NA | + | NA | + | NA | NA | NA | NA | NA | NA | 0,0443452457298926 | 5 | -526,996882019979 | 1064,45177930713 | 16,4902063918423 | 9.13218774644125e-06 |
| **301** | 25,5914766583413 | NA | NA | + | + | NA | + | NA | NA | + | NA | NA | NA | 0,0594427697604111 | 6 | -525,906574539943 | 1064,45930292604 | 16,4977300107489 | 9.09789873105445e-06 |

| | | | | | | | | | | | | | | | | | | | |
|---|---|---|---|---|---|---|---|---|---|---|---|---|---|---|---|---|---|---|---|
| **3997** | 25,3886228310462 | NA | NA | + | + | + | NA | NA | + | + | + | + | + | 0,119905438488168 | 10 | -521,357398977518 | 1064,46082970107 | 16,4992567857776 | 9.09095615903012e-06 |
| **2974** | 25,3901267034831 | + | NA | + | + | + | NA | NA | + | + | + | NA | + | 0,119888957266834 | 10 | -521,358681106021 | 1064,46339395807 | 16,5018210427827 | 9.07930785393071e-06 |
| **2808** | 24,3727097491015 | + | + | + | NA | + | + | + | + | NA | + | NA | + | 0,134886693461085 | 11 | -520,181918540211 | 1064,47583708042 | 16,5142641651316 | 9.02299574096662e-06 |
| **357** | 25,6401075666544 | NA | NA | + | NA | NA | + | + | NA | + | NA | NA | NA | 0,059328245715722 | 6 | -525,914910860533 | 1064,47597556722 | 16,5144026519295 | 9.0223709797037e-06 |
| **3022** | 25,2816186554478 | + | NA | + | + | NA | NA | + | + | + | + | NA | + | 0,119725295894492 | 10 | -521,371411561158 | 1064,48885486835 | 16,5272819530567 | 8.96445673721879e-06 |
| **4045** | 25,2933108388279 | NA | NA | + | + | NA | NA | + | + | + | + | + | + | 0,11969641829323 | 10 | -521,37365756959 | 1064,49334688521 | 16,5317739699208 | 8.94434508572218e-06 |
| **3823** | 24,2465878835538 | NA | + | + | + | NA | + | + | + | NA | + | + | + | 0,134751523202455 | 11 | -520,192615148353 | 1064,49723029671 | 16,5356573814145 | 8.92699464963944e-06 |
| **4057** | 25,2502905752203 | NA | NA | NA | + | + | NA | + | + | + | + | + | + | 0,119664760462926 | 10 | -521,376119730821 | 1064,49827120767 | 16,5366982923833 | 8.92234975514978e-06 |
| **2465** | 26,4572874374024 | NA | NA | NA | NA | NA | + | NA | + | + | NA | NA | + | 0,0591437104533515 | 6 | -525,928341234178 | 1064,50283631451 | 16,5412633992185 | 8.90200724049381e-06 |
| **608** | 24,6771391904651 | + | + | + | + | + | NA | + | NA | NA | + | NA | NA | 0,104527413256977 | 9 | -522,543390921477 | 1064,5041046776 | 16,542531762308 | 8.89636354154426e-06 |
| **3034** | 25,2613201313094 | + | NA | NA | + | + | NA | + | + | + | + | NA | + | 0,119584435654254 | 10 | -521,382366528408 | 1064,51076480285 | 16,5491918875568 | 8.8667873664108e-06 |
| **38** | 25,2475667818129 | + | NA | + | NA | NA | + | NA | NA | NA | NA | NA | NA | 0,0439304905742708 | 5 | -527,026591096146 | 1064,51119745947 | 16,5496245441766 | 8.86486943674453e-06 |
| **2341** | 25,6699811110977 | NA | NA | + | NA | NA | + | NA | NA | + | NA | NA | + | 0,0590571020145604 | 6 | -525,934643640305 | 1064,51544112676 | 16,5538682114725 | 8.84607959999251e-06 |
| **309** | 26,0125469520813 | NA | NA | + | NA | + | + | NA | NA | + | NA | NA | NA | 0,0590467261481857 | 6 | -525,935398642429 | 1064,51695113101 | 16,5553782157212 | 8.83940331172514e-06 |
| **2652** | 24,3139642048699 | + | + | NA | + | + | NA | + | NA | NA | + | NA | + | 0,104432971689504 | 9 | -522,550611421242 | 1064,51854567713 | 16,556972761839 | 8.83235870221998e-06 |
| **3933** | 25,2531967322478 | NA | NA | + | + | + | NA | + | NA | + | + | + | + | 0,119527603123821 | 10 | -521,386786005782 | 1064,5196037576 | 16,5580308423062 | 8.82768726490725e-06 |
| **2243** | 25,0051143657075 | NA | + | NA | NA | NA | NA | + | + | NA | NA | NA | + | 0,0589980242608826 | 6 | -525,938942334427 | 1064,52403851501 | 16,5624655997171 | 8.80813462496231e-06 |
| **2241** | 25,8886690219886 | NA | NA | NA | NA | NA | NA | + | + | NA | NA | NA | + | 0,0437837284503296 | 5 | -527,037100639615 | 1064,53221654641 | 16,570643631115 | 8.77219156016153e-06 |
| **2910** | 25,2740748183777 | + | NA | + | + | + | NA | + | NA | + | + | NA | + | 0,119403493057491 | 10 | -521,396436206086 | 1064,5389041582 | 16,5773312429126 | 8.74290804076477e-06 |
| **1788** | 24,1247898079938 | + | + | NA | + | + | + | + | + | NA | + | + | NA | 0,134480301878193 | 11 | -520,214073029795 | 1064,54014605959 | 16,5785731442998 | 8.73748081114827e-06 |
| **3727** | 24,6951500600175 | NA | + | + | + | NA | NA | NA | + | NA | + | + | + | 0,104287056464162 | 9 | -522,561765826844 | 1064,54085448833 | 16,5792815730438 | 8.73438641794143e-06 |
| **1678** | 25,2827983195939 | + | NA | + | + | NA | NA | NA | + | NA | + | + | NA | 0,0891342613197631 | 8 | -523,71033492022 | 1064,54566984044 | 16,584096925149 | 8.71338214082297e-06 |
| **427** | 25,7512926270346 | NA | + | NA | + | NA | + | NA | + | + | NA | NA | NA | 0,0739289887400286 | 7 | -524,843842577885 | 1064,55590221003 | 16,5943292947438 | 8.66891671124875e-06 |
| **2097** | 25,6103310040456 | NA | NA | NA | NA | + | + | NA | NA | NA | NA | NA | + | 0,0436074624012498 | 5 | -527,049720809774 | 1064,55745688672 | 16,5958839714328 | 8.66218064832297e-06 |
| **2760** | 24,547193131176 | + | + | + | NA | NA | NA | + | + | NA | + | NA | + | 0,104177132931719 | 9 | -522,570167668739 | 1064,55765817212 | 16,5960852568337 | 8.66130890693883e-06 |
| **1073** | 25,8668797285109 | NA | NA | NA | NA | + | + | NA | NA | NA | NA | + | NA | 0,0436002539440283 | 5 | -527,05023686624 | 1064,55848899966 | 16,5969160843656 | 8.65771162721924e-06 |
| **2815** | 24,1468014304249 | NA | + | + | + | + | + | + | + | NA | + | NA | + | 0,134345172397378 | 11 | -520,224761393267 | 1064,56152278653 | 16,5999498712442 | 8.64458875676674e-06 |
| **1743** | 24,8674403400965 | NA | + | + | + | NA | NA | + | + | NA | + | + | NA | 0,104150904342754 | 9 | -522,572172259699 | 1064,56166735404 | 16,600094438752 | 8.64396391602396e-06 |
| **3656** | 24,4293132568242 | + | + | + | NA | NA | NA | + | NA | NA | + | + | + | 0,104100847924781 | 9 | -522,575997794232 | 1064,56931842311 | 16,6077455078189 | 8.61095930394992e-06 |
| **3870** | 25,2880139647934 | + | NA | + | + | + | NA | NA | NA | + | + | + | + | 0,119201788977903 | 10 | -521,412116843118 | 1064,57026543227 | 16,608692516977 | 8.60688294045381e-06 |
| **3816** | 24,470100570003 | + | + | + | NA | NA | + | + | + | NA | + | + | + | 0,134271976154226 | 11 | -520,230550314815 | 1064,57310062963 | 16,6115277143385 | 8.59469047860282e-06 |
| **3994** | 25,2779834718862 | + | NA | NA | + | + | NA | NA | + | + | + | + | + | 0,119182664267944 | 10 | -521,41360342713 | 1064,57323860029 | 16,6116656850008 | 8.59409759148448e-06 |
| **171** | 25,2097045291272 | NA | + | NA | + | NA | + | NA | + | NA | NA | NA | NA | 0,0586373100438656 | 6 | -525,965183251903 | 1064,57652034996 | 16,6149474346685 | 8.58000731637768e-06 |
| **3** | 26,3088888888889 | NA | + | NA | NA | NA | NA | NA | NA | NA | NA | NA | NA | 0,0131085723822034 | 3 | -529,19908104804 | 1064,5786132239 | 16,6170403086085 | 8.5710335755649e-06 |
| **2772** | 24,4575275160357 | + | + | NA | NA | + | NA | + | + | NA | + | NA | + | 0,103962433389706 | 9 | -522,586574937802 | 1064,59047271025 | 16,6288997949587 | 8.52035993702926e-06 |
| **3930** | 25,1901778946738 | + | NA | NA | + | + | NA | + | NA | + | + | + | + | 0,119018754610784 | 10 | -521,426342975855 | 1064,59871769774 | 16,6371447824508 | 8.4853071086508e-06 |
| **2779** | 24,3073470375163 | NA | + | NA | + | + | NA | + | + | NA | + | NA | + | 0,103877589985614 | 9 | -522,593057559914 | 1064,60343795447 | 16,6418650391834 | 8.4653043085202e-06 |
| **1058** | 25,7647396823622 | + | NA | NA | NA | NA | + | NA | NA | NA | NA | + | NA | 0,0432692259605061 | 5 | -527,073931110309 | 1064,60587748779 | 16,6443045725032 | 8.45498490746877e-06 |
| **1187** | 25,6694851021394 | NA | + | NA | NA | NA | + | NA | + | NA | NA | + | NA | 0,0584081504278534 | 6 | -525,981848731969 | 1064,60985131009 | 16,6482783948011 | 8.43820228200598e-06 |
| **728** | 24,8716935496522 | + | + | + | NA | + | NA | + | + | NA | + | NA | NA | 0,103791945567031 | 9 | -522,599600762916 | 1064,61652436048 | 16,6549514451863 | 8.41009492404103e-06 |
| **418** | 26,4965097665301 | + | NA | NA | NA | NA | + | NA | + | + | NA | NA | NA | 0,0583343252546657 | 6 | -525,987216755008 | 1064,62058735617 | 16,659014440879 | 8.39302717673097e-06 |
| **3982** | 25,32233917814 | + | NA | + | + | NA | NA | NA | + | + | + | + | + | 0,118785333691502 | 10 | -521,44448105922 | 1064,63499386447 | 16,6734209491817 | 8.3327872909527e-06 |
| **2082** | 25,4512597766679 | + | NA | NA | NA | NA | + | NA | NA | NA | NA | NA | + | 0,043029892808649 | 5 | -527,091056939552 | 1064,64012914628 | 16,6785562309899 | 8.31141913008095e-06 |
| **50** | 25,7643777515301 | + | NA | NA | NA | + | + | NA | NA | NA | NA | NA | NA | 0,042884286031781 | 5 | -527,10147394825 | 1064,66096316368 | 16,6993902483853 | 8.22528839588139e-06 |
| **1992** | 25,3777259153738 | + | + | + | NA | NA | NA | + | + | + | + | + | NA | 0,118561928643346 | 10 | -521,461836357649 | 1064,66970446133 | 16,7081315460382 | 8.18941699655425e-06 |
| **2401** | 25,751707717234 | NA | NA | NA | NA | NA | + | + | NA | + | NA | NA | + | 0,0579963850598036 | 6 | -526,011783909057 | 1064,66972166427 | 16,7081487489779 | 8.18934655583375e-06 |
| **433** | 26,5160592612093 | NA | NA | NA | NA | + | + | NA | + | + | NA | NA | NA | 0,0579794279246442 | 6 | -526,013016405565 | 1064,67218665728 | 16,710613741993 | 8.17925943225372e-06 |
| **361** | 25,7167363897465 | NA | NA | NA | + | NA | + | + | NA | + | NA | NA | NA | 0,0579785795427229 | 6 | -526,013078067978 | 1064,67230998211 | 16,7107370668198 | 8.17875509492701e-06 |
| **3612** | 24,5969040411007 | + | + | NA | + | + | NA | NA | NA | NA | + | + | + | 0,103425054477648 | 9 | -522,627624039519 | 1064,67257091368 | 16,7109979983925 | 8.17768811681573e-06 |
| **2345** | 25,7106547749644 | NA | NA | NA | + | NA | + | NA | NA | + | NA | NA | + | 0,0579447761840034 | 6 | -526,015534931224 | 1064,6772237086 | 16,7156507933105 | 8.15868567614541e-06 |
| **3710** | 24,5173683943234 | + | NA | + | + | + | + | + | NA | NA | + | + | + | 0,133591874086014 | 11 | -520,284314636582 | 1064,68062927316 | 16,7190563578733 | 8.14480503194771e-06 |
| **294** | 25,7233385162581 | + | NA | + | NA | NA | + | NA | NA | + | NA | NA | NA | 0,0578772778896031 | 6 | -526,020440513779 | 1064,68703487371 | 16,7254619584212 | 8.11876057801592e-06 |
| **233** | 25,5604750191965 | NA | NA | NA | + | NA | + | + | + | NA | NA | NA | NA | 0,0578022899423774 | 6 | -526,025890010919 | 1064,69793386799 | 16,7363609527019 | 8.07463774827287e-06 |
| **1628** | 24,6876914647819 | + | + | NA | + | + | NA | + | NA | NA | + | + | NA | 0,103252394523187 | 9 | -522,640807904469 | 1064,69893864358 | 16,7373657282935 | 8.07058216763511e-06 |
| **4096** | 24,8468916370068 | + | + | + | + | + | + | + | + | + | + | + | + | 0,17864384209586 | 14 | -516,628355050123 | 1064,69933305107 | 16,737760135775 | 8.06899077556099e-06 |
| **1755** | 24,6760570800386 | NA | + | NA | + | + | NA | + | + | NA | + | + | NA | 0,103207459962156 | 9 | -522,644238574582 | 1064,70579998381 | 16,7442270685185 | 8.0429421016854e-06 |
| **371** | 25,4451338010909 | NA | + | NA | NA | + | + | + | NA | + | NA | NA | NA | 0,0729099512947791 | 7 | -524,919142298591 | 1064,70650165145 | 16,7449287361544 | 8.04012086052268e-06 |
| **3787** | 24,3159022337337 | NA | + | NA | + | NA | NA | + | + | NA | + | + | + | 0,103185075553395 | 9 | -522,645947518421 | 1064,70921787149 | 16,7476449561968 | 8.02920890330667e-06 |
| **3696** | 24,0820903822705 | + | + | + | + | NA | + | + | NA | NA | + | + | + | 0,133367795234516 | 11 | -520,302019573999 | 1064,716039148 | 16,7544662327066 | 8.00187082291679e-06 |
| **3828** | 24,2973361987993 | + | + | NA | NA | + | + | + | + | NA | + | + | + | 0,133359913742325 | 11 | -520,302642223834 | 1064,71728444767 | 16,755711532378 | 7.99689001017572e-06 |
| **1317** | 25,850295541544 | NA | NA | + | NA | NA | + | NA | NA | + | NA | + | NA | 0,0576361323919308 | 6 | -526,037963408336 | 1064,72208066283 | 16,7605077475346 | 7.97773558400638e-06 |
| **1944** | 25,5990759913282 | + | + | + | NA | + | NA | NA | + | + | + | + | NA | 0,118198334118369 | 10 | -521,490072925606 | 1064,72617759724 | 16,764604681953 | 7.96141018099406e-06 |
| **3663** | 24,2136153424159 | NA | + | + | + | NA | NA | + | NA | NA | + | + | + | 0,103036745716144 | 9 | -522,657270723428 | 1064,7318642815 | 16,7702913662117 | 7.93880531992332e-06 |
| **1441** | 26,5744508083023 | NA | NA | NA | NA | NA | + | NA | + | + | NA | + | NA | 0,0575310139593035 | 6 | -526,045600461425 | 1064,737354769 | 16,7757818537127 | 7.91704125173697e-06 |
| **3774** | 24,5812340204875 | + | NA | + | + | + | + | NA | + | NA | + | + | + | 0,133188783462392 | 11 | -520,31616038007 | 1064,74432076014 | 16,7827478448494 | 7.88951419827361e-06 |
| **1614** | 24,8913591394461 | + | NA | + | + | NA | NA | + | NA | NA | + | + | NA | 0,0878100824243807 | 8 | -523,809797514742 | 1064,74459502948 | 16,7830221141942 | 7.88843234651002e-06 |
| **1790** | 24,5237029596944 | + | NA | + | + | + | + | + | + | NA | + | + | NA | 0,133146515242326 | 11 | -520,319498878106 | 1064,75099775621 | 16,7894248409218 | 7.86321898829235e-06 |
| **313** | 25,9957276566708 | NA | NA | NA | + | + | + | NA | NA | + | NA | NA | NA | 0,057312955407335 | 6 | -526,06144011336 | 1064,76903407287 | 16,807461157583 | 7.79262602216707e-06 |
| **3780** | 24,4526339630547 | + | + | NA | NA | NA | NA | + | + | NA | + | + | + | 0,102782196721598 | 9 | -522,676698126671 | 1064,77071908799 | 16,8091461726967 | 7.78606344075606e-06 |
| **2592** | 24,5913658411655 | + | + | + | + | + | NA | NA | NA | NA | + | NA | + | 0,102771968832822 | 9 | -522,677478613012 | 1064,77228006067 | 16,810707145379 | 7.77998889544769e-06 |
| **2814** | 24,4769176415562 | + | NA | + | + | + | + | + | + | NA | + | NA | + | 0,132978899186422 | 11 | -520,332736203236 | 1064,77747240647 | 16,8158994911814 | 7.75981689543545e-06 |
| **1091** | 25,2553027487971 | NA | + | NA | NA | NA | NA | + | NA | NA | NA | + | NA | 0,0420066015888465 | 5 | -527,164231769434 | 1064,78647880604 | 16,8249058907531 | 7.72495145160234e-06 |
| **1686** | 25,4144605983575 | + | NA | + | NA | + | NA | NA | + | NA | + | + | NA | 0,0875244368197304 | 8 | -523,831234191013 | 1064,78746838203 | 16,8258954667363 | 7.72113018382539e-06 |
| **3739** | 24,563924988569 | NA | + | NA | + | + | NA | NA | + | NA | + | + | + | 0,102644856246136 | 9 | -522,687177784673 | 1064,79167840399 | 16,8301054887017 | 7.70489421443141e-06 |
| **4042** | 25,1077100587884 | + | NA | NA | + | NA | NA | + | + | + | + | + | + | 0,117670398415165 | 10 | -521,531051438119 | 1064,80813462227 | 16,8465617069789 | 7.64175760758425e-06 |
| **3608** | 24,9282277186423 | + | + | + | NA | + | NA | NA | NA | NA | + | + | + | 0,102513211797419 | 9 | -522,697221307118 | 1064,81176544888 | 16,8501925335906 | 7.62789724409418e-06 |

| | | | | | | | | | | | | | | | | | | | |
|---|---|---|---|---|---|---|---|---|---|---|---|---|---|---|---|---|---|---|---|
| **369** | 26,0557785543366 | NA | NA | NA | NA | + | + | + | NA | + | NA | NA | NA | 0,0569880253650534 | 6 | -526,085036057752 | 1064,81622596166 | 16,8546530463673 | 7.61090403418735e-06 |
| **1748** | 24,8144812663317 | + | + | NA | NA | + | NA | + | + | NA | + | + | NA | 0,102399246462119 | 9 | -522,705914851584 | 1064,82915253781 | 16,8675796225236 | 7.56187119624919e-06 |
| **2812** | 24,0836718738167 | + | + | NA | + | + | + | + | + | NA | + | NA | + | 0,132618932375228 | 11 | -520,361155611034 | 1064,83431122207 | 16,8727383067767 | 7.54239167629036e-06 |
| **3835** | 24,1050619562882 | NA | + | NA | + | + | + | + | + | NA | + | + | + | 0,132603809512527 | 11 | -520,362349304182 | 1064,83669860836 | 16,8751256930725 | 7.53339374647952e-06 |
| **3735** | 24,9007256406852 | NA | + | + | NA | + | NA | NA | + | NA | + | + | + | 0,102284479244915 | 9 | -522,714668450639 | 1064,84665973592 | 16,8850868206325 | 7.49596648025694e-06 |
| **3030** | 25,4588020852884 | + | NA | + | NA | + | NA | + | + | + | + | NA | + | 0,117417813074125 | 10 | -521,550648514131 | 1064,84732877429 | 16,8857558590037 | 7.49345935501902e-06 |
| **1321** | 25,8886211839499 | NA | NA | NA | + | NA | + | NA | NA | + | NA | + | NA | 0,0566907285552292 | 6 | -526,106618198987 | 1064,85939024413 | 16,8978173288378 | 7.4484042821908e-06 |
| **2800** | 24,165364878244 | + | + | + | + | NA | + | + | + | NA | + | NA | + | 0,132412244009091 | 11 | -520,377468346553 | 1064,86693669311 | 16,9053637778161 | 7.42035273641942e-06 |
| **2719** | 24,5871328604843 | NA | + | + | + | + | NA | NA | + | NA | + | NA | + | 0,102131825132498 | 9 | -522,726310050128 | 1064,8699429349 | 16,9083700196111 | 7.40920742762624e-06 |
| **298** | 25,8517194330754 | + | NA | NA | + | NA | + | NA | NA | + | NA | NA | NA | 0,0565869498237686 | 6 | -526,114150373044 | 1064,87445459224 | 16,9128816769512 | 7.39251236277133e-06 |
| **2305** | 26,6215970610982 | NA | NA | NA | NA | NA | NA | NA | NA | + | NA | NA | + | 0,0261753548736315 | 4 | -528,286465377644 | 1064,87596105832 | 16,9143881430282 | 7.38694617480372e-06 |
| **2371** | 25,0235839404694 | NA | + | NA | NA | NA | NA | + | NA | + | NA | NA | + | 0,0565694432340464 | 6 | -526,115420905135 | 1064,87699565642 | 16,9154227411325 | 7.38312590274597e-06 |
| **1736** | 25,0103553996367 | + | + | + | NA | NA | NA | + | + | NA | + | + | NA | 0,102074823117537 | 9 | -522,73065659003 | 1064,87863601471 | 16,9170630994149 | 7.3770728994964e-06 |
| **4050** | 25,3529350290805 | + | NA | NA | NA | + | NA | + | + | + | + | + | + | 0,117215590857269 | 10 | -521,566334078714 | 1064,87869990346 | 16,9171269881683 | 7.37683724726465e-06 |
| **732** | 24,6169328537459 | + | + | NA | + | + | NA | + | + | NA | + | NA | NA | 0,102028164805782 | 9 | -522,734214192576 | 1064,8857512198 | 16,9241783045079 | 7.35087483494545e-06 |
| **4053** | 25,4862657532214 | NA | NA | + | NA | + | NA | + | + | + | + | + | + | 0,117149888930564 | 10 | -521,571429540136 | 1064,8888908263 | 16,927317911013 | 7.33934446029812e-06 |
| **2273** | 25,6339323506757 | NA | NA | NA | NA | NA | + | + | + | NA | NA | NA | + | 0,0564798453907333 | 6 | -526,121923055388 | 1064,88999995693 | 16,928427041639 | 7.33527544281257e-06 |
| **3837** | 24,4731600452008 | NA | NA | + | + | + | + | + | + | NA | + | + | + | 0,132218434045787 | 11 | -520,392761134681 | 1064,89752226936 | 16,9359493540719 | 7.30773814448437e-06 |
| **354** | 25,8809507873076 | + | NA | NA | NA | NA | + | + | NA | + | NA | NA | NA | 0,0564243560692445 | 6 | -526,125949627517 | 1064,89805310119 | 16,9364801858976 | 7.30579881187014e-06 |
| **1377** | 25,9274622768002 | NA | NA | NA | NA | NA | + | + | NA | + | NA | + | NA | 0,056400718345749 | 6 | -526,12766482257 | 1064,90148349129 | 16,9399105760033 | 7.29327868219263e-06 |
| **3926** | 25,3565123324224 | + | NA | + | NA | + | NA | + | NA | + | + | + | + | 0,117051914221109 | 10 | -521,579027187966 | 1064,90408612196 | 16,9425132066731 | 7.28379399942184e-06 |
| **3600** | 24,5029643086266 | + | + | + | + | NA | NA | NA | NA | NA | + | + | + | 0,101873384275692 | 9 | -522,746014574146 | 1064,90935198294 | 16,9477790676469 | 7.26464150080331e-06 |
| **321** | 26,4715166441816 | NA | NA | NA | NA | NA | NA | + | NA | + | NA | NA | NA | 0,0259182839485599 | 4 | -528,304537583417 | 1064,91210546986 | 16,9505325545729 | 7.25464683473052e-06 |
| **3822** | 24,5372531737635 | + | NA | + | + | NA | + | + | + | NA | + | + | + | 0,131899645333014 | 11 | -520,417908084297 | 1064,94781616859 | 16,9862432533027 | 7.12626216839143e-06 |
| **1936** | 25,4655893131982 | + | + | + | + | NA | NA | NA | + | + | + | + | NA | 0,116675453700658 | 10 | -521,608212746306 | 1064,96245723864 | 17,0008843233531 | 7.07428460051599e-06 |
| **451** | 25,9269388639505 | NA | + | NA | NA | NA | NA | + | + | + | NA | NA | NA | 0,0559680236799914 | 6 | -526,159054327181 | 1064,96426250052 | 17,0026895852261 | 7.06790201337575e-06 |
| **229** | 25,6096763945374 | NA | NA | + | NA | NA | + | + | + | NA | NA | NA | NA | 0,0559220258859613 | 6 | -526,162390356412 | 1064,97093455898 | 17,009361643688 | 7.04436257162177e-06 |
| **2369** | 25,8700980158271 | NA | NA | NA | NA | NA | NA | + | NA | + | NA | NA | + | 0,0406107515797501 | 5 | -527,26392214254 | 1064,98585955225 | 17,0242866369642 | 6.99198969885522e-06 |
| **4038** | 25,3735943366608 | + | NA | + | NA | NA | NA | + | + | + | + | + | + | 0,11646071595907 | 10 | -521,62485498228 | 1064,99574171059 | 17,034168795301 | 6.95752693594398e-06 |
| **1075** | 25,4928575676193 | NA | + | NA | NA | + | + | NA | NA | NA | NA | + | NA | 0,0557493059263226 | 6 | -526,174915568538 | 1064,99598498323 | 17,034412067939 | 6.95668069944559e-06 |
| **720** | 24,7440550546642 | + | + | + | + | NA | NA | + | + | NA | + | NA | NA | 0,10124629320682 | 9 | -522,793802857504 | 1065,00492854965 | 17,0433556343623 | 6.92564138375428e-06 |
| **3039** | 24,981530791446 | NA | + | + | + | + | NA | + | + | + | + | NA | + | 0,131473703259104 | 11 | -520,451493190984 | 1065,01498638197 | 17,0534134666771 | 6.89090034200924e-06 |
| **1996** | 25,0923736551851 | + | + | NA | + | NA | NA | + | + | + | + | + | NA | 0,116310067042531 | 10 | -521,636527903313 | 1065,01908755266 | 17,0575146373671 | 6.87678444063032e-06 |
| **2211** | 25,359458714585 | NA | + | NA | NA | NA | + | NA | + | NA | NA | NA | + | 0,0555868799184464 | 6 | -526,186692200277 | 1065,01953824671 | 17,0579653314164 | 6.87523495231039e-06 |
| **1067** | 25,0562858557172 | NA | + | NA | + | NA | + | NA | NA | NA | NA | + | NA | 0,0555526263645672 | 6 | -526,189175481802 | 1065,02450480976 | 17,0629318944677 | 6.85818298954367e-06 |
| **100** | 24,6646995137937 | + | + | NA | NA | NA | + | + | NA | NA | NA | NA | NA | 0,0555409033305814 | 6 | -526,190025346478 | 1065,02620453911 | 17,0646316238192 | 6.8523569381095e-06 |
| **3107** | 25,1499411984052 | NA | + | NA | NA | NA | + | NA | NA | NA | NA | + | + | 0,0554897524326158 | 6 | -526,193733421974 | 1065,0336206901 | 17,0720477748102 | 6.82699493243684e-06 |
| **1379** | 25,3152184813884 | NA | + | NA | NA | NA | + | + | NA | + | NA | + | NA | 0,0706841344132091 | 7 | -525,083327268678 | 1065,03487159162 | 17,0732986763298 | 6.82272631831233e-06 |
| **1249** | 25,8521295222023 | NA | NA | NA | NA | NA | + | + | + | NA | NA | + | NA | 0,0553914097492519 | 6 | -526,200862001631 | 1065,04787784942 | 17,0863049341258 | 6.77850120628982e-06 |
| **241** | 25,7248307419627 | NA | NA | NA | NA | + | + | + | + | NA | NA | NA | NA | 0,0553883729170421 | 6 | -526,201082121099 | 1065,04831808835 | 17,0867451730617 | 6.77700929041675e-06 |
| **3990** | 25,4960000212562 | + | NA | + | NA | + | NA | NA | + | + | + | + | + | 0,116080184405346 | 10 | -521,654336358509 | 1065,05470446305 | 17,0931315477592 | 6.75540354404466e-06 |
| **69** | 25,753623574905 | NA | NA | + | NA | NA | NA | + | NA | NA | NA | NA | NA | 0,0248626916340904 | 4 | -528,378696269609 | 1065,06042284225 | 17,0988499269572 | 6.736116150791e-06 |
| **226** | 25,7684187633952 | + | NA | NA | NA | NA | + | + | + | NA | NA | NA | NA | 0,0552483274547869 | 6 | -526,211232302973 | 1065,0686184521 | 17,1070455368097 | 6.70856934012002e-06 |
| **1872** | 25,0300297881645 | + | + | + | + | NA | NA | + | NA | + | + | + | NA | 0,115869357266456 | 10 | -521,670664561026 | 1065,08736086808 | 17,1257879527923 | 6.64599559523117e-06 |
| **2467** | 25,8921050058853 | NA | + | NA | NA | NA | + | NA | + | + | NA | NA | + | 0,0703182418380308 | 7 | -525,110279315565 | 1065,08877568539 | 17,1272027701029 | 6.6412958229496e-06 |
| **179** | 25,5309003246553 | NA | + | NA | NA | + | + | NA | + | NA | NA | NA | NA | 0,0551049590124952 | 6 | -526,221621769256 | 1065,08939738467 | 17,1278244693749 | 6.63923169939325e-06 |
| **2353** | 26,1187339384188 | NA | NA | NA | NA | + | + | NA | NA | + | NA | NA | + | 0,0550758117824344 | 6 | -526,223733785871 | 1065,0936214179 | 17,132048502606 | 6.62522432883598e-06 |
| **3361** | 26,0895623240762 | NA | NA | NA | NA | NA | + | NA | NA | + | NA | + | + | 0,0550705997554494 | 6 | -526,224111443972 | 1065,0943767341 | 17,1328038188078 | 6.62272273160196e-06 |
| **2912** | 24,9397745372262 | + | + | + | + | + | NA | + | NA | + | + | NA | + | 0,130947276396735 | 11 | -520,492978683917 | 1065,09795736783 | 17,136384452544 | 6.61087656674138e-06 |
| **2338** | 26,0098032723478 | + | NA | NA | NA | NA | + | NA | NA | + | NA | NA | + | 0,0550392987099087 | 6 | -526,226379441657 | 1065,09891272947 | 17,1373398141775 | 6.60771943193499e-06 |
| **167** | 25,3793571050843 | NA | + | + | NA | NA | + | NA | + | NA | NA | NA | NA | 0,0550215396247893 | 6 | -526,227666188415 | 1065,10148622298 | 17,1399133076932 | 6.59922243829112e-06 |
| **3935** | 24,9302389218421 | NA | + | + | + | + | NA | + | NA | + | + | + | + | 0,130921078901105 | 11 | -520,495042542534 | 1065,10208508507 | 17,1405121697771 | 6.59724672205031e-06 |
| **2147** | 24,4060800281564 | NA | + | NA | NA | NA | + | + | NA | NA | NA | NA | + | 0,0549609377135851 | 6 | -526,232056959713 | 1065,11026776558 | 17,1486948502888 | 6.57031028160438e-06 |
| **3732** | 24,7870129278686 | + | + | NA | NA | + | NA | NA | + | NA | + | + | + | 0,100496551450116 | 9 | -522,85089417148 | 1065,11911117761 | 17,1575382623153 | 6.54132243621832e-06 |
| **3918** | 25,0275778134259 | + | NA | + | + | NA | NA | + | NA | + | + | + | + | 0,115643640956297 | 10 | -521,688141591419 | 1065,12231492887 | 17,1607420135788 | 6.53085243926356e-06 |
| **2307** | 25,9063135043194 | NA | + | NA | NA | NA | NA | NA | NA | + | NA | NA | + | 0,0396388011509917 | 5 | -527,333252398285 | 1065,12452006375 | 17,1629471484557 | 6.52365570219165e-06 |
| **1568** | 25,063116536015 | + | + | + | + | + | NA | NA | NA | NA | + | + | NA | 0,100414752860405 | 9 | -522,857120090247 | 1065,13156301514 | 17,169990099849 | 6.50072320899035e-06 |
| **423** | 25,8661265062483 | NA | + | + | NA | NA | + | NA | + | + | NA | NA | NA | 0,0700032205580336 | 7 | -525,133475634817 | 1065,1351683239 | 17,1735954086068 | 6.48901520774177e-06 |
| **3720** | 24,9195605994818 | + | + | + | NA | NA | NA | NA | + | NA | + | + | + | 0,100381406681142 | 9 | -522,859657998587 | 1065,13663883182 | 17,1750659165284 | 6.48424588715448e-06 |
| **2716** | 24,5142206121151 | + | + | NA | + | + | NA | NA | + | NA | + | NA | + | 0,100364708392017 | 9 | -522,860928835307 | 1065,13918050526 | 17,1776075899681 | 6.4760107032758e-06 |
| **1695** | 25,0577573916945 | NA | + | + | + | + | NA | NA | + | NA | + | + | NA | 0,100348975896998 | 9 | -522,862126147919 | 1065,14157513048 | 17,1800022151926 | 6.46826153399851e-06 |
| **164** | 25,4991351303791 | + | + | NA | NA | NA | + | NA | + | NA | NA | NA | NA | 0,0547189191560789 | 6 | -526,249589048713 | 1065,14533194358 | 17,1837590282901 | 6.45612291337566e-06 |
| **1443** | 26,103392997074 | NA | + | NA | NA | NA | + | NA | + | + | NA | + | NA | 0,0698774505596312 | 7 | -525,142734406164 | 1065,15368586659 | 17,1921129513012 | 6.42921217816989e-06 |
| **81** | 26,2861097612466 | NA | NA | NA | NA | + | NA | + | NA | NA | NA | NA | NA | 0,0241431679166662 | 4 | -528,429199103885 | 1065,1614285108 | 17,1998555955097 | 6.40437074257607e-06 |
| **2403** | 25,0319817184308 | NA | + | NA | NA | NA | + | + | NA | + | NA | NA | + | 0,0697420853498105 | 7 | -525,152698147129 | 1065,17361334852 | 17,2120404332302 | 6.36547124979884e-06 |
| **1251** | 25,2376349718741 | NA | + | NA | NA | NA | + | + | + | NA | NA | + | NA | 0,06971164820976 | 7 | -525,154938314791 | 1065,17809368385 | 17,2165207685548 | 6.35122748711465e-06 |
| **2057** | 25,612075984766 | NA | NA | NA | + | NA | NA | NA | NA | NA | NA | NA | + | 0,0240129755925142 | 4 | -528,438333229424 | 1065,17969676188 | 17,218123846588 | 6.34613877015543e-06 |
| **3830** | 24,697544144029 | + | NA | + | NA | + | + | + | + | NA | + | + | + | 0,13039359358122 | 11 | -520,536585011464 | 1065,18517002293 | 17,2235971076375 | 6.32879547500213e-06 |
| **73** | 25,8521684237106 | NA | NA | NA | + | NA | NA | + | NA | NA | NA | NA | NA | 0,0239527519233304 | 4 | -528,442558032397 | 1065,18814636782 | 17,2265734525338 | 6.31938414052751e-06 |
| **3834** | 24,3861651595029 | + | NA | NA | + | + | + | + | + | NA | + | + | + | 0,130266377999532 | 11 | -520,546600192431 | 1065,20520038486 | 17,2436274695706 | 6.2657277873324e-06 |
| **1063** | 25,1866775845893 | NA | + | + | NA | NA | + | NA | NA | NA | NA | + | NA | 0,0542641841232616 | 6 | -526,282518420436 | 1065,21119068703 | 17,2496177717348 | 6.24698906263425e-06 |
| **1060** | 25,3777523268778 | + | + | NA | NA | NA | + | NA | NA | NA | NA | + | NA | 0,0542466594651003 | 6 | -526,283787141591 | 1065,21372812934 | 17,2521552140449 | 6.23906840107307e-06 |
| **2704** | 24,5827992017759 | + | + | + | + | NA | NA | NA | + | NA | + | NA | + | 0,0998385744135873 | 9 | -522,900958614825 | 1065,2192400643 | 17,2576671490044 | 6.22189740367935e-06 |
| **2053** | 25,62797731569 | NA | NA | + | NA | NA | NA | NA | NA | NA | NA | NA | + | 0,0237102177312334 | 4 | -528,459569622141 | 1065,22216954731 | 17,2605966320223 | 6.21279060347024e-06 |
| **107** | 24,4994455861604 | NA | + | NA | + | NA | + | + | NA | NA | NA | NA | NA | 0,0541619482334655 | 6 | -526,289919593627 | 1065,22599303341 | 17,2644201181181 | 6.20092469012851e-06 |
| **103** | 24,5192476072669 | NA | + | + | NA | NA | + | + | NA | NA | NA | NA | NA | 0,0541230711779427 | 6 | -526,292733814422 | 1065,231621475 | 17,2700485597074 | 6.18349845106259e-06 |

| | | | | | | | | | | | | | | | | | | | |
|---|---|---|---|---|---|---|---|---|---|---|---|---|---|---|---|---|---|---|---|
| 59 | 25,0786693535644 | NA | + | NA | + | + | + | NA | NA | NA | NA | NA | NA | 0,0540445970030125 | 6 | -526,29841402758 | 1065,24298190131 | 17,2814089860224 | 6.1484744279305e-06 |
| 66 | 25,8812458137977 | + | NA | NA | NA | NA | NA | + | NA | NA | NA | NA | NA | 0,0235506242157607 | 4 | -528,470761366041 | 1065,24455303511 | 17,2829801198222 | 6.14364628659712e-06 |
| 306 | 26,2208554878551 | + | NA | NA | NA | + | + | NA | NA | + | NA | NA | NA | 0,0539150653119481 | 6 | -526,307788917678 | 1065,26173168151 | 17,3001587662193 | 6.09110250383112e-06 |
| 315 | 25,6009660174457 | NA | + | NA | + | + | + | NA | NA | + | NA | NA | NA | 0,0691355763677825 | 7 | -525,197323282709 | 1065,26286361968 | 17,3012907043906 | 6.0876561034865e-06 |
| 4047 | 24,9676710414836 | NA | + | + | + | NA | NA | + | + | + | + | + | + | 0,129860856595979 | 11 | -520,578515523194 | 1065,26903104639 | 17,3074581310977 | 6.06891243197643e-06 |
| 3724 | 24,5089043005769 | + | + | NA | + | NA | NA | NA | + | NA | + | + | + | 0,0995109438668826 | 9 | -522,925873871204 | 1065,26907057705 | 17,3074976617625 | 6.06879247909064e-06 |
| 3789 | 24,8800352848196 | NA | NA | + | + | NA | NA | + | + | NA | + | + | + | 0,0994431311436198 | 9 | -522,931029680382 | 1065,27938219541 | 17,3178092801197 | 6.0375834661156e-06 |
| 311 | 25,6913397428156 | NA | + | + | NA | + | + | NA | NA | + | NA | NA | NA | 0,0690082416003521 | 7 | -525,206688506663 | 1065,28159406759 | 17,3200211522983 | 6.03090997557554e-06 |
| 435 | 26,0660029823864 | NA | + | NA | NA | + | + | NA | + | + | NA | NA | NA | 0,0689447426107191 | 7 | -525,211358255169 | 1065,2909335646 | 17,3293606493103 | 6.00281279716432e-06 |
| 1314 | 26,1413311655639 | + | NA | NA | NA | NA | + | NA | NA | + | NA | + | NA | 0,0536926003709794 | 6 | -526,323886882652 | 1065,29392761146 | 17,3323546961667 | 5.9938331688058e-06 |
| 1329 | 26,295117998361 | NA | NA | NA | NA | + | + | NA | NA | + | NA | + | NA | 0,0536446432621093 | 6 | -526,327356650061 | 1065,30086714628 | 17,3392942309847 | 5.97307200086277e-06 |
| 420 | 26,0030542687168 | + | + | NA | NA | NA | + | NA | + | + | NA | NA | NA | 0,0688688249193042 | 7 | -525,216940863631 | 1065,30209878153 | 17,3405258662351 | 5.96933481020365e-06 |
| 2764 | 24,190164875973 | + | + | NA | + | NA | NA | + | + | NA | + | NA | + | 0,0992434840528217 | 9 | -522,946206619077 | 1065,3097360728 | 17,3481631575087 | 5.94664327443083e-06 |
| 2712 | 24,8120500978621 | + | + | + | NA | + | NA | NA | + | NA | + | NA | + | 0,0991728511163055 | 9 | -522,951575247254 | 1065,32047332915 | 17,3589004138637 | 5.91480350220953e-06 |
| 1089 | 26,0815383272358 | NA | NA | NA | NA | NA | NA | + | NA | NA | NA | + | NA | 0,0230063856526862 | 4 | -528,508913198984 | 1065,320856701 | 17,359283785707 | 5.91366982630708e-06 |
| 55 | 25,1659334496199 | NA | + | + | NA | + | + | NA | NA | NA | NA | NA | NA | 0,0535028971821334 | 6 | -526,337611160238 | 1065,32137616663 | 17,3598032513387 | 5.91213405164535e-06 |
| 359 | 25,1140104196341 | NA | + | + | NA | NA | + | + | NA | + | NA | NA | NA | 0,068707163133536 | 7 | -525,228827149498 | 1065,32587135326 | 17,3642984379687 | 5.89886090058381e-06 |
| 363 | 25,0996396796164 | NA | + | NA | + | NA | + | + | NA | + | NA | NA | NA | 0,0687054649035696 | 7 | -525,228952001992 | 1065,32612105825 | 17,3645481429578 | 5.89812445905969e-06 |
| 4055 | 25,1254792093475 | NA | + | + | NA | + | NA | + | + | + | + | + | + | 0,129492235624487 | 11 | -520,607513821305 | 1065,32702764261 | 17,3654547273188 | 5.8954514912262e-06 |
| 2781 | 24,8129176492329 | NA | NA | + | + | + | NA | + | + | NA | + | NA | + | 0,0991159427148352 | 9 | -522,955900402708 | 1065,32912364006 | 17,3675507247719 | 5.88927630193133e-06 |
| 3999 | 25,1346689037533 | NA | + | + | + | + | NA | NA | + | + | + | + | + | 0,129477026408267 | 11 | -520,608710020733 | 1065,32942004147 | 17,3678471261758 | 5.88840357172082e-06 |
| 9 | 26,5125 | NA | NA | NA | + | NA | NA | NA | NA | NA | NA | NA | NA | 0,0076278109643621 | 3 | -529,578280515032 | 1065,33701215788 | 17,3754392425938 | 5.86609322148972e-06 |
| 3660 | 24,1085791959298 | + | + | NA | + | NA | NA | + | NA | NA | + | + | + | 0,0990532116153502 | 9 | -522,960667778761 | 1065,33865839217 | 17,377085476877 | 5.86126672626267e-06 |
| 356 | 25,2273771855397 | + | + | NA | NA | NA | + | + | NA | + | NA | NA | NA | 0,068572674344939 | 7 | -525,238713952016 | 1065,3456449583 | 17,3840720430051 | 5.84082738346429e-06 |
| 3872 | 25,1081955850099 | + | + | + | + | + | NA | NA | NA | + | + | + | + | 0,129365583276577 | 11 | -520,617474346038 | 1065,34694869208 | 17,3853757767854 | 5.83702118218491e-06 |
| 4059 | 24,9030839897063 | NA | + | NA | + | + | NA | + | + | + | + | + | + | 0,129319709432103 | 11 | -520,621081719454 | 1065,35416343891 | 17,3925905236174 | 5.81600280051223e-06 |
| 2217 | 25,5589735188274 | NA | NA | NA | + | NA | + | NA | + | NA | NA | NA | + | 0,0531909887153952 | 6 | -526,360170532168 | 1065,36649491049 | 17,4049219952001 | 5.7802531884694e-06 |
| 68 | 25,0212306166842 | + | + | NA | NA | NA | NA | + | NA | NA | NA | NA | NA | 0,037940826140076 | 5 | -527,45425555811 | 1065,36652638339 | 17,4049534681042 | 5.78016222850801e-06 |
| 3932 | 24,8885598274707 | + | + | NA | + | + | NA | + | NA | + | + | + | + | 0,129205029702695 | 11 | -520,630098938123 | 1065,37219787625 | 17,4106249609561 | 5.76379437263998e-06 |
| 3036 | 24,899001802524 | + | + | NA | + | + | NA | + | + | + | + | NA | + | 0,129196898795556 | 11 | -520,630738222855 | 1065,37347644571 | 17,4119035304191 | 5.76011084443913e-06 |
| 75 | 24,941441372479 | NA | + | NA | + | NA | NA | + | NA | NA | NA | NA | NA | 0,0378516003019905 | 5 | -527,460552479308 | 1065,37912022579 | 17,4175473105013 | 5.74387935751822e-06 |
| 2976 | 25,1253567225693 | + | + | + | + | + | NA | NA | + | + | + | NA | + | 0,129137217812787 | 11 | -520,635430399958 | 1065,38286079992 | 17,4212878846261 | 5.73314669397737e-06 |
| 2177 | 26,4759714345726 | NA | NA | NA | NA | NA | NA | NA | + | NA | NA | NA | + | 0,0225616485407765 | 4 | -528,540074081303 | 1065,38317846564 | 17,4216055503462 | 5.7322361542046e-06 |
| 1193 | 25,808631161602 | NA | NA | NA | + | NA | + | NA | + | NA | NA | + | NA | 0,0530712149117667 | 6 | -526,368831425993 | 1065,38381669814 | 17,42224378285 | 5.73040719632842e-06 |
| 71 | 24,9646224741044 | NA | + | + | NA | NA | NA | + | NA | NA | NA | NA | NA | 0,0378061335800336 | 5 | -527,463787933253 | 1065,38559113368 | 17,4240182183914 | 5.72532533192407e-06 |
| 3928 | 25,083365820866 | + | + | + | NA | + | NA | + | NA | + | + | + | + | 0,129114144003315 | 11 | -520,637244399226 | 1065,38648879845 | 17,424915883161 | 5.72275619710022e-06 |
| 173 | 25,5145624416426 | NA | NA | + | + | NA | + | NA | + | NA | NA | NA | NA | 0,0530411326978182 | 6 | -526,37100651142 | 1065,38816686899 | 17,4265939537031 | 5.71795661659351e-06 |
| 3032 | 25,1066460845647 | + | + | + | NA | + | NA | + | + | + | + | NA | + | 0,129090968232444 | 11 | -520,639066366062 | 1065,39013273212 | 17,4285598168331 | 5.71233901785774e-06 |
| 2766 | 24,8000827020856 | + | NA | + | + | NA | NA | + | + | NA | + | NA | + | 0,0986010160323715 | 9 | -522,99502347871 | 1065,40736979207 | 17,4457968767756 | 5.6633185982363e-06 |
| 2010 | 25,5129222907504 | + | NA | NA | + | + | + | + | + | + | + | + | NA | 0,113774748774774 | 10 | -521,832677341739 | 1065,41138642951 | 17,4498135142185 | 5.65195626288442e-06 |
| 672 | 24,9658265470122 | + | + | + | + | + | NA | NA | + | NA | + | NA | NA | 0,0985563468119661 | 9 | -522,998416301672 | 1065,41415543799 | 17,4525825226992 | 5.64413651995476e-06 |
| 3139 | 24,771079811776 | NA | + | NA | NA | NA | NA | + | NA | NA | NA | + | + | 0,052832178977585 | 6 | -526,386112941566 | 1065,41837972929 | 17,4568068139954 | 5.63222786240376e-06 |
| 1886 | 25,5227858955724 | + | NA | + | + | + | NA | + | NA | + | + | + | NA | 0,113702064568171 | 10 | -521,838292411881 | 1065,42261656979 | 17,4610436545036 | 5.62030906579187e-06 |
| 2013 | 25,6805480540226 | NA | NA | + | + | + | NA | + | + | + | + | + | NA | 0,113658734015382 | 10 | -521,841639606391 | 1065,42931095881 | 17,4677380435242 | 5.60152824720169e-06 |
| 3024 | 24,9283022813452 | + | + | + | + | NA | NA | + | + | + | + | NA | + | 0,128838436635992 | 11 | -520,658916036562 | 1065,42983207312 | 17,4682591578344 | 5.60006891906474e-06 |
| 1323 | 25,4770677449986 | NA | + | NA | + | NA | + | NA | NA | + | NA | + | NA | 0,0679892679038577 | 7 | -525,281585951973 | 1065,43138895821 | 17,4698160429182 | 5.59571128348508e-06 |
| 990 | 25,6871315834024 | + | NA | + | + | + | NA | + | + | + | + | NA | NA | 0,113538866425416 | 10 | -521,85089827486 | 1065,44782829575 | 17,4862553804605 | 5.5499049047271e-06 |
| 2499 | 25,4628915396586 | NA | + | NA | NA | NA | NA | + | + | + | NA | NA | + | 0,0678601097614695 | 7 | -525,291073595068 | 1065,4503642444 | 17,4887913291084 | 5.54287222737708e-06 |
| 52 | 25,4173135827747 | + | + | NA | NA | + | + | NA | NA | NA | NA | NA | NA | 0,0524737440051567 | 6 | -526,412018449096 | 1065,47019074435 | 17,5086178290549 | 5.48819580769046e-06 |
| 170 | 25,7080659781583 | + | NA | NA | + | NA | + | NA | + | NA | NA | NA | NA | 0,0524520721108078 | 6 | -526,413584448239 | 1065,47332274263 | 17,5117498273403 | 5.47960802374675e-06 |
| 1219 | 25,6569505903379 | NA | + | NA | NA | NA | NA | + | + | NA | NA | + | NA | 0,0524246086595134 | 6 | -526,415568890666 | 1065,47729162749 | 17,5157187121947 | 5.46874483934687e-06 |
| 185 | 25,6385757612321 | NA | NA | NA | + | + | + | NA | + | NA | NA | NA | NA | 0,0523983246136589 | 6 | -526,417468058277 | 1065,48108996271 | 17,5195170474171 | 5.45836863247078e-06 |
| 3363 | 25,5924351506983 | NA | + | NA | NA | NA | + | NA | NA | + | NA | + | + | 0,0676417653213689 | 7 | -525,307109658847 | 1065,48243637196 | 17,5208634566663 | 5.45469527006562e-06 |
| 2099 | 25,2152240283098 | NA | + | NA | NA | + | + | NA | NA | NA | NA | NA | NA | 0,0523675139325334 | 6 | -526,419694233274 | 1065,4855423127 | 17,5239693974115 | 5.4462308641067e-06 |
| 489 | 26,1114340100036 | NA | NA | NA | + | NA | + | + | + | + | NA | NA | NA | 0,0675161544843468 | 7 | -525,316333305843 | 1065,50088366595 | 17,5393107506582 | 5.40461440558662e-06 |
| 235 | 24,9377386525886 | NA | + | NA | + | NA | + | + | + | NA | NA | NA | NA | 0,0674801887250744 | 7 | -525,318974055224 | 1065,50616516471 | 17,5445922494212 | 5.39036100155485e-06 |
| 2640 | 24,0447435709656 | + | + | + | + | + | NA | + | NA | NA | + | NA | + | 0,0979487933262765 | 9 | -523,044545971533 | 1065,50641477771 | 17,5448418624219 | 5.3896882914427e-06 |
| 1692 | 24,9299253472913 | + | + | NA | + | + | NA | NA | + | NA | + | + | NA | 0,0979093651735725 | 9 | -523,047538556904 | 1065,51239994845 | 17,5508270331629 | 5.37358329891169e-06 |
| 5 | 26,6011111111111 | NA | NA | + | NA | NA | NA | NA | NA | NA | NA | NA | NA | 0,0063369191070487 | 3 | -529,667289320104 | 1065,51502976803 | 17,5534568527376 | 5.36652216503343e-06 |
| 2275 | 24,9160436250878 | NA | + | NA | NA | NA | + | + | + | NA | NA | NA | + | 0,0674116873897559 | 7 | -525,324003414681 | 1065,51622388363 | 17,5546509683345 | 5.36331899745737e-06 |
| 1616 | 24,6191811239974 | + | + | + | + | NA | NA | + | NA | NA | + | + | NA | 0,0978831227836207 | 9 | -523,049530274237 | 1065,51638338312 | 17,5548104678294 | 5.36289129117683e-06 |
| 1950 | 25,6609657716942 | + | NA | + | + | + | NA | NA | + | + | + | + | NA | 0,113042137646809 | 10 | -521,889252670294 | 1065,52453708662 | 17,5629641713285 | 5.34107208548797e-06 |
| 1331 | 25,9297417599329 | NA | + | NA | NA | + | + | NA | NA | + | NA | + | NA | 0,0673068938575913 | 7 | -525,331696628213 | 1065,53161031069 | 17,5700373953996 | 5.32221614838841e-06 |
| 3801 | 24,7584216167527 | NA | NA | NA | + | + | NA | + | + | NA | + | + | + | 0,0976495434813949 | 9 | -523,067255679355 | 1065,55183419336 | 17,5902612780649 | 5.26866939800372e-06 |
| 2355 | 25,7247656943668 | NA | + | NA | NA | + | + | NA | NA | + | NA | NA | + | 0,0670799285536987 | 7 | -525,348355883402 | 1065,56492882107 | 17,6033559057762 | 5.23428644666543e-06 |
| 3820 | 24,0554275880851 | + | + | + | NA | + | NA | + | + | NA | + | + | + | 0,127976363713332 | 11 | -520,726634013249 | 1065,5652680265 | 17,6036951112083 | 5.23339877274569e-06 |
| 243 | 25,1118496590866 | NA | + | NA | NA | + | + | + | NA | + | NA | NA | NA | 0,0670495951855942 | 7 | -525,35058204598 | 1065,56938114622 | 17,6078082309336 | 5.22264703447037e-06 |
| 3677 | 24,7642282901151 | NA | NA | + | + | + | NA | + | NA | NA | + | + | + | 0,0974397524544177 | 9 | -523,083171977911 | 1065,58366679047 | 17,6220938751774 | 5.18547550853726e-06 |
| 1319 | 25,5733532992857 | NA | + | + | NA | NA | + | NA | NA | + | NA | + | NA | 0,0669215457197606 | 7 | -525,359978785072 | 1065,58817462441 | 17,626601709117 | 5.17380103887687e-06 |
| 3797 | 25,0387804002962 | NA | NA | + | NA | + | NA | + | + | NA | + | + | + | 0,0973865014760335 | 9 | -523,087211402457 | 1065,59174563956 | 17,6301727242694 | 5.16457142015338e-06 |
| 203 | 25,3807667105369 | NA | + | NA | + | NA | NA | + | + | NA | NA | NA | NA | 0,0515860107198982 | 6 | -526,476136282956 | 1065,59842641207 | 17,636853496776 | 5.1473485383309e-06 |
| 2347 | 25,2204085771855 | NA | + | NA | + | NA | + | NA | NA | + | NA | NA | + | 0,0668481991545141 | 7 | -525,365360644252 | 1065,59893834277 | 17,6373654274773 | 5.14603116406551e-06 |
| 2654 | 24,7533076001628 | + | NA | + | + | + | NA | + | NA | NA | + | NA | + | 0,0973172506783896 | 9 | -523,092464158514 | 1065,60225115167 | 17,6406782363822 | 5.13751431075716e-06 |
| 3741 | 24,9718710232109 | NA | NA | + | + | + | NA | NA | + | NA | + | + | + | 0,0972806039773114 | 9 | -523,095243691774 | 1065,60781021819 | 17,6462373029035 | 5.12325424619608e-06 |
| 228 | 25,1115314652301 | + | + | NA | NA | NA | + | + | + | NA | NA | NA | NA | 0,0667843749274725 | 7 | -525,370043450699 | 1065,60830395566 | 17,64673104037 | 5.12198963101357e-06 |
| 2778 | 24,7197940582788 | + | NA | NA | + | + | NA | + | + | NA | + | NA | + | 0,0972657288899431 | 9 | -523,096371886661 | 1065,61006660797 | 17,6484936926777 | 5.11747747622418e-06 |

| | | | | | | | | | | | | | | | | | | |
|---|---|---|---|---|---|---|---|---|---|---|---|---|---|---|---|---|---|---|
| 211 | 25,5671479502791 | NA | + | NA | NA | + | NA | + | + | NA | NA | NA | NA | 0,0514965158961782 | 6 | -526,482596848469 | 1065,61134754309 | 17,6497746278005 | 5.1142009472716e-06 |
| 1740 | 24,6531203418077 | + | + | NA | + | NA | NA | + | + | NA | + | + | NA | 0,0972423049404062 | 9 | -523,098148428818 | 1065,61361969228 | 17,6520467769913 | 5.10839413261167e-06 |
| 2131 | 24,8612780924361 | NA | + | NA | NA | + | NA | + | NA | NA | NA | NA | + | 0,0514280341464265 | 6 | -526,487540084135 | 1065,62123401442 | 17,6596610991335 | 5.08898262814281e-06 |
| 4052 | 25,0186147627024 | + | + | NA | NA | + | NA | + | + | + | + | + | + | 0,127570526946172 | 11 | -520,758490341595 | 1065,62898068319 | 17,6674077678988 | 5.06930942180568e-06 |
| 3996 | 25,034447445869 | + | + | NA | + | + | NA | NA | + | + | + | + | + | 0,127561550181221 | 11 | -520,759194809104 | 1065,63038961821 | 17,6688167029181 | 5.06573951561076e-06 |
| 3073 | 26,282162365794 | NA | NA | NA | NA | NA | NA | NA | NA | NA | NA | + | + | 0,0207099229336966 | 4 | -528,669664638971 | 1065,64235958097 | 17,6807866656807 | 5.0355117055974e-06 |
| 303 | 25,2413313861321 | NA | + | + | + | NA | + | NA | NA | + | NA | NA | NA | 0,066468562463049 | 7 | -525,393210014682 | 1065,65463708363 | 17,6930641683364 | 5.00469463736401e-06 |
| 231 | 25,0995494050176 | NA | + | + | NA | NA | + | + | + | NA | NA | NA | NA | 0,0664621538111327 | 7 | -525,393680043099 | 1065,65557714046 | 17,6940042251701 | 5.00234284141538e-06 |
| 339 | 25,7857023684415 | NA | + | NA | NA | + | NA | + | NA | + | NA | NA | NA | 0,0511503661832752 | 6 | -526,507579407113 | 1065,66131266038 | 17,6997397450898 | 4.98801787301228e-06 |
| 2343 | 25,2899759441984 | NA | + | + | NA | NA | + | NA | NA | + | NA | NA | + | 0,0664160797024657 | 7 | -525,397059151872 | 1065,66233535801 | 17,7007624427163 | 4.98546790800888e-06 |
| 2774 | 24,9763085511305 | + | NA | + | NA | + | NA | + | + | NA | + | NA | + | 0,0968982763693397 | 9 | -523,124235274853 | 1065,66579338435 | 17,7042204690606 | 4.97685541602813e-06 |
| 3782 | 25,023210188021 | + | NA | + | NA | NA | NA | + | + | NA | + | + | + | 0,0968512035844043 | 9 | -523,127803916586 | 1065,67293066782 | 17,7113577525276 | 4.95912645505895e-06 |
| 3726 | 24,9851075899275 | + | NA | + | + | NA | NA | NA | + | NA | + | + | + | 0,0968465509820794 | 9 | -523,128156625601 | 1065,67363608585 | 17,712063170557 | 4.9573776348837e-06 |
| 2117 | 25,1163427038158 | NA | NA | + | NA | NA | NA | + | NA | NA | NA | NA | + | 0,0356412943155139 | 5 | -527,617663298608 | 1065,69334186439 | 17,7317689491001 | 4.9087729833037e-06 |
| 429 | 26,0145236205562 | NA | NA | + | + | NA | + | NA | + | + | NA | NA | NA | 0,0661582171418694 | 7 | -525,415967903877 | 1065,70015286202 | 17,7385799467265 | 4.89208459500654e-06 |
| 3984 | 25,1136089441811 | + | + | + | + | NA | NA | NA | + | + | + | + | + | 0,127099969027267 | 11 | -520,795408446294 | 1065,70281689259 | 17,7412439772966 | 4.88557260155064e-06 |
| 308 | 25,881350005161 | + | + | NA | NA | + | + | NA | NA | + | NA | NA | NA | 0,0661366151827209 | 7 | -525,417551712688 | 1065,70332047964 | 17,741747564349 | 4.88434260085731e-06 |
| 4040 | 25,0912612266865 | + | + | + | NA | NA | NA | + | + | + | + | + | + | 0,12705209904743 | 11 | -520,799163019933 | 1065,71032603987 | 17,7487531245758 | 4.86726375191647e-06 |
| 3674 | 24,7331294221257 | + | NA | NA | + | + | NA | + | NA | NA | + | + | + | 0,0965939612249122 | 9 | -523,147302471069 | 1065,71192777678 | 17,7503548614923 | 4.86336727439099e-06 |
| 2718 | 24,9167502748121 | + | NA | + | + | + | NA | NA | + | NA | + | NA | + | 0,0965087126870477 | 9 | -523,153762948017 | 1065,72484873068 | 17,7632758153884 | 4.83204887704677e-06 |
| 2050 | 25,9840605950872 | + | NA | NA | NA | NA | NA | NA | NA | NA | NA | NA | + | 0,0201135285015897 | 4 | -528,711350369165 | 1065,72573104136 | 17,7641581260702 | 4.82991766300985e-06 |
| 2006 | 25,6709797191234 | + | NA | + | NA | + | NA | + | + | + | + | + | NA | 0,111706695517106 | 10 | -521,992261091856 | 1065,73055392974 | 17,7689810144523 | 4.818284617975e-06 |
| 3075 | 25,656831802775 | NA | + | NA | NA | NA | NA | NA | NA | NA | NA | + | + | 0,035350207071746 | 5 | -527,638327251782 | 1065,73466977074 | 17,7730968554486 | 4.80837916710479e-06 |
| 44 | 24,866995077975 | + | + | NA | + | NA | + | NA | NA | NA | NA | NA | NA | 0,050636765743284 | 6 | -526,544630555657 | 1065,73541495747 | 17,7738420421765 | 4.80658793065777e-06 |
| 300 | 25,3882821837867 | + | + | NA | + | NA | + | NA | NA | + | NA | NA | NA | 0,0659114826613948 | 7 | -525,434055761877 | 1065,73632857802 | 17,7747556627264 | 4.8043927333361e-06 |
| 1680 | 25,1138426068995 | + | + | + | + | NA | NA | NA | + | NA | + | + | NA | 0,0964049128202519 | 9 | -523,161628496717 | 1065,74057982808 | 17,7790069127893 | 4.79419124199496e-06 |
| 1998 | 25,5551838881192 | + | NA | + | + | NA | NA | + | + | + | + | + | NA | 0,11164093847378 | 10 | -521,997329217483 | 1065,740690181 | 17,7791172657076 | 4.7939267227956e-06 |
| 3137 | 25,5344882719054 | NA | NA | NA | NA | NA | NA | + | NA | NA | NA | + | + | 0,0352573827510572 | 5 | -527,644915434357 | 1065,74784613589 | 17,7862732205997 | 4.77680481027784e-06 |
| 199 | 25,5485532603486 | NA | + | + | NA | NA | NA | + | + | NA | NA | NA | NA | 0,0505330890379246 | 6 | -526,552107365334 | 1065,75036857682 | 17,7887956615302 | 4.77078400387276e-06 |
| 3614 | 24,9121742274107 | + | NA | + | + | + | NA | NA | NA | NA | + | + | + | 0,0963286429237207 | 9 | -523,167407356172 | 1065,75213754699 | 17,790564631699 | 4.76656618215569e-06 |
| 485 | 26,086112773445 | NA | NA | + | NA | NA | + | + | + | + | NA | NA | NA | 0,0657590535745355 | 7 | -525,445227799475 | 1065,75867265321 | 17,7970997379225 | 4.75101659238514e-06 |
| 1688 | 25,2380523508698 | + | + | + | NA | + | NA | NA | + | NA | + | + | NA | 0,0962609737012475 | 9 | -523,172534146901 | 1065,76239112845 | 17,8008182131564 | 4.74219153010393e-06 |
| 3786 | 24,6895725129595 | + | NA | NA | + | NA | NA | + | + | NA | + | + | + | 0,0962418015627734 | 9 | -523,173986606645 | 1065,76529604794 | 17,8037231326455 | 4.73530868754145e-06 |
| 2065 | 26,0963712071674 | NA | NA | NA | NA | + | NA | NA | NA | NA | NA | NA | + | 0,0198135525137325 | 4 | -528,732307978899 | 1065,76764626083 | 17,8060733455372 | 4.72974746393426e-06 |
| 196 | 25,4783785270653 | + | + | NA | NA | NA | NA | + | + | NA | NA | NA | NA | 0,0504076076079705 | 6 | -526,561155565764 | 1065,76846497768 | 17,8068920623919 | 4.72781169818892e-06 |
| 2 | 26,6909090909091 | + | NA | NA | NA | NA | NA | NA | NA | NA | NA | NA | NA | 0,0044501836397413 | 3 | -529,797174675407 | 1065,77480047863 | 17,8132275633436 | 4.71285886635604e-06 |
| 1316 | 25,7547281168203 | + | + | NA | NA | NA | + | NA | NA | + | NA | + | NA | 0,0656303422958725 | 7 | -525,454660060901 | 1065,77753717607 | 17,815964260775 | 4.70641444209441e-06 |
| 2129 | 25,5405755761707 | NA | NA | NA | NA | + | NA | + | NA | NA | NA | NA | + | 0,0350057289576172 | 5 | -527,662773311546 | 1065,78356189027 | 17,8219889749776 | 4.69225837339991e-06 |
| 2473 | 26,1157876856542 | NA | NA | NA | + | NA | + | NA | + | + | NA | NA | + | 0,0655387467984904 | 7 | -525,461371600757 | 1065,79096025578 | 17,8293873404873 | 4.67493291691527e-06 |
| 3670 | 24,9385989275948 | + | NA | + | NA | + | NA | + | NA | NA | + | + | + | 0,0959943431743251 | 9 | -523,192731013478 | 1065,8027848616 | 17,8412119463112 | 4.64737484353124e-06 |
| 2121 | 25,2134167900879 | NA | NA | NA | + | NA | NA | + | NA | NA | NA | NA | + | 0,0348518993931505 | 5 | -527,673687085507 | 1065,80538943819 | 17,8438165228995 | 4.64132656083271e-06 |
| 3794 | 24,9073830469067 | + | NA | NA | NA | + | NA | + | + | NA | + | + | + | 0,0959623334441134 | 9 | -523,195155302702 | 1065,80763344005 | 17,8460605247597 | 4.63612190847304e-06 |
| 201 | 26,3249554069675 | NA | NA | NA | + | NA | NA | + | + | NA | NA | NA | NA | 0,0348097988225681 | 5 | -527,676673699268 | 1065,81136266571 | 17,8497897504217 | 4.62748539045978e-06 |
| 2529 | 26,1801434511707 | NA | NA | NA | NA | NA | + | + | + | + | NA | NA | + | 0,0653888050986204 | 7 | -525,472356962897 | 1065,81293098006 | 17,8513580647666 | 4.62385813695469e-06 |
| 2091 | 24,6677422155924 | NA | + | NA | + | NA | + | NA | NA | NA | NA | NA | + | 0,0500753549485556 | 6 | -526,585107831677 | 1065,81636950951 | 17,8547965942175 | 4.61591533061413e-06 |
| 2213 | 25,5998620038657 | NA | NA | + | NA | NA | + | NA | + | NA | NA | NA | + | 0,0500454228456479 | 6 | -526,587265241253 | 1065,82068432866 | 17,8591114133687 | 4.60596764515402e-06 |
| 3776 | 24,3988705439448 | + | + | + | + | + | + | NA | + | NA | + | + | + | 0,141527146455079 | 12 | -519,654363409662 | 1065,82485585158 | 17,8632829362919 | 4.59637070728736e-06 |
| 117 | 25,3074502596601 | NA | NA | + | NA | + | + | + | NA | NA | NA | NA | NA | 0,0499697763967065 | 6 | -526,592717290823 | 1065,8315884278 | 17,8700155125098 | 4.58092401282966e-06 |
| 2084 | 24,968728498572 | + | + | NA | NA | NA | + | NA | NA | NA | NA | NA | + | 0,0498667340558114 | 6 | -526,600143140319 | 1065,84644012679 | 17,8848672115018 | 4.54703275199807e-06 |
| 3738 | 24,8664238805833 | + | NA | NA | + | + | NA | NA | + | NA | + | + | + | 0,0956686658376601 | 9 | -523,217392511871 | 1065,85210785839 | 17,8905349430968 | 4.53416531231213e-06 |
| 2340 | 25,4955164613453 | + | + | NA | NA | NA | + | NA | NA | + | NA | NA | + | 0,0650954179637945 | 7 | -525,493846648701 | 1065,85591035167 | 17,894337436376 | 4.52555293546931e-06 |
| 491 | 25,4766285245903 | NA | + | NA | + | NA | + | + | + | + | NA | NA | NA | 0,0803702590652791 | 8 | -524,365953045802 | 1065,8569060916 | 17,895333176313 | 4.52330035936335e-06 |
| 1189 | 25,8434837075746 | NA | NA | + | NA | NA | + | NA | + | NA | NA | + | NA | 0,049759251571793 | 6 | -526,607888115482 | 1065,86193007712 | 17,9003571618262 | 4.51195212093853e-06 |
| 2114 | 25,3843050210077 | + | NA | NA | NA | NA | NA | + | NA | NA | NA | NA | + | 0,0344198918942053 | 5 | -527,70432750441 | 1065,866670276 | 17,9050973607045 | 4.50127100839645e-06 |
| 296 | 25,4286371385731 | + | + | + | NA | NA | + | NA | NA | + | NA | NA | NA | 0,0650096214957262 | 7 | -525,500129696606 | 1065,86847644748 | 17,906903532185 | 4.49720780972044e-06 |
| 2015 | 25,3172817894925 | NA | + | + | + | + | NA | + | + | + | + | + | NA | 0,1259755122269 | 11 | -520,883548360456 | 1065,87909672091 | 17,9175238056221 | 4.47339031444344e-06 |
| 47 | 24,7096632382875 | NA | + | + | + | NA | + | NA | NA | NA | NA | NA | NA | 0,0496018658746271 | 6 | -526,619227437757 | 1065,88460872167 | 17,9230358063771 | 4.46107862235484e-06 |
| 992 | 25,3043220233644 | + | + | + | + | + | NA | + | + | + | + | NA | NA | 0,125917994727699 | 11 | -520,88805378981 | 1065,88810757962 | 17,9265346643288 | 4.45328110473983e-06 |
| 3712 | 24,3369460040751 | + | + | + | + | + | + | + | NA | NA | + | + | + | 0,141032431843539 | 12 | -519,693806713 | 1065,90374245826 | 17,9421695429667 | 4.41860357129198e-06 |
| 2087 | 24,7534443137079 | NA | + | + | NA | NA | + | NA | NA | NA | NA | NA | + | 0,0494656311625577 | 6 | -526,62904135877 | 1065,90423656369 | 17,9426636484031 | 4.41751207810261e-06 |
| 1792 | 24,3524412044411 | + | + | + | + | + | + | + | + | NA | + | + | NA | 0,140995160463816 | 12 | -519,696777418251 | 1065,90968386876 | 17,9481109534704 | 4.40549668045066e-06 |
| 2531 | 25,4509583280955 | NA | + | NA | NA | NA | + | + | + | + | NA | NA | + | 0,0800080537889213 | 8 | -524,392914362598 | 1065,9108287252 | 17,9492558099059 | 4.4029755714833e-06 |
| 166 | 25,6702543325177 | + | NA | + | NA | NA | + | NA | + | NA | NA | NA | NA | 0,0491978165691551 | 6 | -526,648329783558 | 1065,94281341327 | 17,9812404979798 | 4.33312172352565e-06 |
| 441 | 26,1716618692254 | NA | NA | NA | + | + | + | NA | + | + | NA | NA | NA | 0,0644963702118099 | 7 | -525,53770408937 | 1065,943625233 | 17,9820523177123 | 4.33136322358687e-06 |
| 259 | 26,8015274555298 | NA | + | NA | NA | NA | NA | NA | NA | + | NA | NA | NA | 0,018549924709782 | 4 | -528,820520113172 | 1065,94407052937 | 17,9824976140835 | 4.33039896077351e-06 |
| 40 | 24,9676069477774 | + | + | + | NA | NA | + | NA | NA | NA | NA | NA | NA | 0,0491555236872366 | 6 | -526,651375286375 | 1065,9489044189 | 17,9873315036136 | 4.31994526376766e-06 |
| 181 | 25,7358646137082 | NA | NA | + | NA | + | + | NA | + | NA | NA | NA | NA | 0,0491326472181347 | 6 | -526,653022560373 | 1065,9521989669 | 17,9906260516084 | 4.31283498815844e-06 |
| 1888 | 25,2593243334138 | + | + | + | + | + | NA | + | NA | + | + | + | NA | 0,125483373648489 | 11 | -520,922088714687 | 1065,95617742937 | 17,9946045140839 | 4.30426428945371e-06 |
| 1757 | 25,2126387424211 | NA | NA | + | + | + | NA | + | + | NA | + | + | NA | 0,0949508877026185 | 9 | -523,271714012561 | 1065,96075085977 | 17,9991779444761 | 4.29443290796389e-06 |
| 734 | 25,1725147564538 | + | NA | + | + | + | NA | + | + | NA | + | NA | NA | 0,0948670146601041 | 9 | -523,278058720412 | 1065,97344027547 | 18,0118673601792 | 4.26727224013934e-06 |
| 3233 | 26,0204523430242 | NA | NA | NA | NA | NA | + | NA | + | NA | NA | + | + | 0,0489405569518511 | 6 | -526,66685290948 | 1065,97985966511 | 18,0182867498236 | 4.25359755604647e-06 |
| 3992 | 25,3029075713615 | + | + | + | NA | + | NA | NA | + | + | + | + | + | 0,125319070355298 | 11 | -520,934950806859 | 1065,98190161372 | 18,0203286984281 | 4.24925695844593e-06 |
| 426 | 26,232158006334 | + | NA | NA | + | NA | + | NA | + | + | NA | NA | NA | 0,0641414787319215 | 7 | -525,56367313966 | 1065,99556333358 | 18,0339904182924 | 4.22032979054304e-06 |
| 1449 | 26,2340044524343 | NA | NA | NA | + | NA | + | NA | + | + | NA | + | NA | 0,0641188897224278 | 7 | -525,565325748697 | 1065,99886855166 | 18,0372956363665 | 4.21336099532499e-06 |
| 3791 | 24,5525422448782 | NA | + | + | + | NA | NA | + | + | NA | + | + | + | 0,109945033747412 | 10 | -522,127909159097 | 1066,00185006423 | 18,0402771489353 | 4.20708458040764e-06 |
| 3920 | 24,7329851529839 | + | + | + | + | NA | NA | + | NA | + | + | + | + | 0,125185318248694 | 11 | -520,94541948918 | 1066,00283897836 | 18,0412660630702 | 4.20500487190904e-06 |

| | | | | | | | | | | | | | | | | | | |
|---|---|---|---|---|---|---|---|---|---|---|---|---|---|---|---|---|---|---|
| 497 | 26,2481723079043 | NA | NA | NA | NA | + | + | + | + | + | NA | NA | NA | 0,0640861546850279 | 7 | -525,567720569082 | 1066,00365819243 | 18,0420852771369 | 4.20328282504352e-06 |
| 449 | 26,910313011532 | NA | NA | NA | NA | NA | NA | + | + | + | NA | NA | NA | 0,0334517555020564 | 5 | -527,772943488741 | 1066,00390224466 | 18,0423293293672 | 4.20276994606245e-06 |
| 1507 | 25,6719251558723 | NA | + | NA | NA | NA | + | + | + | + | NA | + | NA | 0,0793813011711571 | 8 | -524,439542621209 | 1066,00408524242 | 18,042512327128 | 4.20238541491037e-06 |
| 2179 | 25,7888888888889 | NA | + | NA | NA | NA | NA | NA | + | NA | NA | NA | + | 0,0333967872283786 | 5 | -527,776837265154 | 1066,01168979748 | 18,0501168821929 | 4.18643711840327e-06 |
| 1139 | 25,0620890719057 | NA | + | NA | NA | + | + | + | NA | NA | NA | + | NA | 0,0639948381855694 | 7 | -525,574400632597 | 1066,01701831946 | 18,0554454041676 | 4.17529820239142e-06 |
| 121 | 25,3612488306007 | NA | NA | NA | + | + | + | + | NA | NA | NA | NA | NA | 0,0486755847211871 | 6 | -526,685926121656 | 1066,01800608947 | 18,0564331741748 | 4.17323659436422e-06 |
| 482 | 26,2703680662797 | + | NA | NA | NA | NA | + | + | + | + | NA | NA | NA | 0,0639872842329944 | 7 | -525,574953196695 | 1066,01812344765 | 18,0565505323625 | 4.172991719807e-06 |
| 3662 | 24,6345145089238 | + | NA | + | + | NA | NA | + | NA | NA | + | + | + | 0,0945473086459043 | 9 | -523,302237995372 | 1066,02179882539 | 18,0602259100981 | 4.16533010136974e-06 |
| 3169 | 25,3309045602176 | NA | NA | NA | NA | NA | + | + | NA | NA | NA | + | + | 0,0486031501772737 | 6 | -526,691139175625 | 1066,0284321974 | 18,0668592821141 | 4.1515378941574e-06 |
| 194 | 26,3561814527121 | + | NA | NA | NA | NA | NA | + | + | NA | NA | NA | NA | 0,0332404184103546 | 5 | -527,787912721054 | 1066,03384070928 | 18,0722677939939 | 4.14032623957183e-06 |
| 2497 | 26,3123911149411 | NA | NA | NA | NA | NA | NA | + | + | + | NA | NA | + | 0,0485638306904434 | 6 | -526,693968800319 | 1066,03409144679 | 18,0725185315011 | 4.13980720456763e-06 |
| 11 | 25,7994499679406 | NA | + | NA | + | NA | NA | NA | NA | NA | NA | NA | NA | 0,017903162238349 | 4 | -528,865625807122 | 1066,03428191727 | 18,0727090019843 | 4.13941296780129e-06 |
| 1125 | 25,2589173075946 | NA | NA | + | NA | NA | + | + | NA | NA | NA | + | NA | 0,0485373519425328 | 6 | -526,695874276087 | 1066,03790239833 | 18,0763294830376 | 4.13192641296734e-06 |
| 4044 | 24,7909561051578 | + | + | NA | + | NA | NA | + | + | + | + | + | + | 0,124955438270302 | 11 | -520,963408294667 | 1066,03881658933 | 18,0772436740442 | 4.13003815957298e-06 |
| 2225 | 25,8046881529465 | NA | NA | NA | NA | + | + | NA | + | NA | NA | NA | + | 0,0485042823714325 | 6 | -526,698253969254 | 1066,04266178466 | 18,0810888693707 | 4.12210538606781e-06 |
| 1505 | 26,2818042398593 | NA | NA | NA | NA | NA | + | + | + | + | NA | + | NA | 0,0637582349362712 | 7 | -525,59170580804 | 1066,05162867034 | 18,0900557550528 | 4.10366553009323e-06 |
| 2161 | 25,3628422035588 | NA | NA | NA | NA | + | + | + | NA | NA | NA | NA | + | 0,0484068347191806 | 6 | -526,705265841742 | 1066,05668552964 | 18,095112614348 | 4.09330280673248e-06 |
| 3799 | 24,6785341368943 | NA | + | + | NA | + | NA | + | + | NA | + | + | + | 0,109571309484585 | 10 | -522,156651445947 | 1066,05933463793 | 18,0977617226358 | 4.08788459468722e-06 |
| 3672 | 24,6648581722027 | + | + | + | NA | + | NA | + | NA | NA | + | + | + | 0,109528155929542 | 10 | -522,159969511604 | 1066,06597076924 | 18,1043978539496 | 4.07434320324084e-06 |
| 197 | 26,3259515785003 | NA | NA | + | NA | NA | NA | + | + | NA | NA | NA | NA | 0,032945646276534 | 5 | -527,808786284842 | 1066,07558783686 | 18,1140149215685 | 4.05479861426451e-06 |
| 2210 | 25,8732798458574 | + | NA | NA | NA | NA | + | NA | + | NA | NA | NA | + | 0,0482595204138972 | 6 | -526,715864520406 | 1066,07788288697 | 18,1163099716757 | 4.05014829995363e-06 |
| 257 | 27,6566666666667 | NA | NA | NA | NA | NA | NA | NA | NA | + | NA | NA | NA | 0,0021544621755788 | 3 | -529,954883739105 | 1066,09021860603 | 18,1286456907385 | 4.02524443487938e-06 |
| 1129 | 25,3170632102996 | NA | NA | NA | + | NA | + | + | NA | NA | NA | + | NA | 0,0481061346301638 | 6 | -526,72689827543 | 1066,09995039701 | 18,1383774817236 | 4.00570559166342e-06 |
| 1137 | 25,6005418862301 | NA | NA | NA | NA | + | + | + | NA | NA | NA | + | NA | 0,0480939995701308 | 6 | -526,727771131051 | 1066,10169610826 | 18,1401231929651 | 4.00221071450617e-06 |
| 499 | 25,6436375272176 | NA | + | NA | NA | + | + | + | + | + | NA | NA | NA | 0,0786955423377699 | 8 | -524,490524404411 | 1066,10604880882 | 18,1444758935322 | 3.99350997344723e-06 |
| 2149 | 24,8961362495118 | NA | NA | + | NA | NA | + | + | NA | NA | NA | NA | + | 0,0480345363600114 | 6 | -526,732048064668 | 1066,11024997549 | 18,1486770601989 | 3.9851300773718e-06 |
| 61 | 25,2887798041517 | NA | NA | + | + | + | + | NA | NA | NA | NA | NA | NA | 0,047969838585139 | 6 | -526,736701194757 | 1066,11955623567 | 18,1579833203778 | 3.96662982412107e-06 |
| 1201 | 26,0830554192422 | NA | NA | NA | NA | + | + | NA | + | NA | NA | + | NA | 0,0479627306904041 | 6 | -526,737212382541 | 1066,12057861124 | 18,1590056959451 | 3.9646026495902e-06 |
| 2309 | 26,1276285752652 | NA | NA | + | NA | NA | NA | NA | NA | + | NA | NA | + | 0,0325962894419742 | 5 | -527,833516896318 | 1066,12504905981 | 18,1634761445212 | 3.95575077010896e-06 |
| 114 | 25,568467092944 | + | NA | NA | NA | + | + | + | NA | NA | NA | NA | NA | 0,0478259054248402 | 6 | -526,747051881099 | 1066,14025760835 | 18,178684693062 | 3.92578423757484e-06 |
| 1186 | 26,0838726399383 | + | NA | NA | NA | NA | + | NA | + | NA | NA | + | NA | 0,0478084963014283 | 6 | -526,748303719908 | 1066,14276128597 | 18,1811883706794 | 3.92087286327758e-06 |
| 3784 | 24,7359031010313 | + | + | + | NA | NA | NA | + | + | NA | + | + | + | 0,109017584907562 | 10 | -522,199214989763 | 1066,14446172556 | 18,1828888102677 | 3.91754067630512e-06 |
| 2469 | 26,082282903046 | NA | NA | + | NA | NA | + | NA | + | + | NA | NA | + | 0,063070585331724 | 7 | -525,641975737491 | 1066,15216852925 | 18,1905956139558 | 3.90247386572039e-06 |
| 487 | 25,575734338459 | NA | + | + | NA | NA | + | + | + | + | NA | NA | NA | 0,0783475245353304 | 8 | -524,516382795122 | 1066,15776559024 | 18,1961926749536 | 3.89156794098515e-06 |
| 3267 | 25,1734549921495 | NA | + | NA | NA | NA | NA | + | + | NA | NA | + | + | 0,0630139647984595 | 7 | -525,646113280962 | 1066,16044361619 | 18,1988707008977 | 3.88636056817852e-06 |
| 484 | 25,6117654718544 | + | + | NA | NA | NA | + | + | + | + | NA | NA | NA | 0,0783096785827015 | 8 | -524,519194233653 | 1066,16338846731 | 18,2018155520152 | 3.88064240236728e-06 |
| 3679 | 24,441105742613 | NA | + | + | + | + | NA | + | NA | NA | + | + | + | 0,108887080650059 | 10 | -522,20924270403 | 1066,16451715409 | 18,2029442388014 | 3.87845300531059e-06 |
| 2153 | 24,9642545969318 | NA | NA | NA | + | NA | + | + | NA | NA | NA | NA | + | 0,0475472069304282 | 6 | -526,767089521416 | 1066,18033288899 | 18,2187599736951 | 3.84790366232162e-06 |
| 2146 | 25,1843689644901 | + | NA | NA | NA | NA | + | + | NA | NA | NA | NA | + | 0,0475372942545956 | 6 | -526,76780210708 | 1066,18175806031 | 18,2201851450225 | 3.84516267804567e-06 |
| 1630 | 25,0918440666001 | + | NA | + | + | + | NA | + | NA | NA | + | + | NA | 0,0934262388508437 | 9 | -523,386956812506 | 1066,19123645966 | 18,2296635443672 | 3.82698279741052e-06 |
| 2116 | 24,5957364716982 | + | + | NA | NA | NA | NA | + | NA | NA | NA | NA | + | 0,0474531947752807 | 6 | -526,773847409906 | 1066,19384866597 | 18,2322757506745 | 3.82198762592462e-06 |
| 178 | 25,9145559096894 | + | NA | NA | NA | + | + | NA | + | NA | NA | NA | NA | 0,0474513835751589 | 6 | -526,773977598104 | 1066,19410904236 | 18,2325361270707 | 3.82149008063036e-06 |
| 109 | 24,9381248770432 | NA | NA | + | + | NA | + | + | NA | NA | NA | NA | NA | 0,04744666124199 | 6 | -526,774317035989 | 1066,19478791813 | 18,2332150028408 | 3.82019314224718e-06 |
| 1742 | 25,1915618984295 | + | NA | + | + | NA | NA | + | + | NA | + | + | NA | 0,0933619924497683 | 9 | -523,391808715642 | 1066,20094026593 | 18,2393673506381 | 3.80845962024498e-06 |
| 1195 | 25,3985355494604 | NA | + | NA | + | NA | + | NA | + | NA | NA | + | NA | 0,062691170221657 | 7 | -525,669696710741 | 1066,20761047575 | 18,2460375604544 | 3.79577916495095e-06 |
| 3734 | 25,1463470576617 | + | NA | + | NA | + | NA | NA | + | NA | + | + | + | 0,0933141422475609 | 9 | -523,395422150746 | 1066,20816713614 | 18,2465942208462 | 3.79472283200389e-06 |
| 2313 | 26,1246972446366 | NA | NA | NA | + | NA | NA | NA | NA | + | NA | NA | + | 0,0319879203950559 | 5 | -527,876561416009 | 1066,21113809919 | 18,2495651839033 | 3.78909002608132e-06 |
| 1122 | 25,5034131139048 | + | NA | NA | NA | NA | + | + | NA | NA | NA | + | NA | 0,0473158323853352 | 6 | -526,783720250635 | 1066,21359434742 | 18,2520214321341 | 3.78443940959648e-06 |
| 102 | 25,1005189320983 | + | NA | + | NA | NA | + | + | NA | NA | NA | NA | NA | 0,0472992298281256 | 6 | -526,784913453256 | 1066,21598075267 | 18,2544078373758 | 3.77992649951814e-06 |
| 1754 | 25,0570305434321 | + | NA | NA | + | + | NA | + | + | NA | + | + | NA | 0,0932398375821933 | 9 | -523,401032931821 | 1066,21938869829 | 18,2578157829976 | 3.77349109197262e-06 |
| 2059 | 25,0465115462905 | NA | + | NA | + | NA | NA | NA | NA | NA | NA | NA | + | 0,0319234360932521 | 5 | -527,881122350713 | 1066,2202599686 | 18,2586870533103 | 3.77184758460157e-06 |
| 1107 | 25,4927820394566 | NA | + | NA | NA | + | NA | + | NA | NA | NA | + | NA | 0,0472571516457853 | 6 | -526,78793746042 | 1066,22202876699 | 18,2604558517039 | 3.76851324029026e-06 |
| 129 | 27,5555555555556 | NA | NA | NA | NA | NA | NA | NA | + | NA | NA | NA | NA | 0,0011795586721072 | 3 | -530,021746955937 | 1066,22394503969 | 18,2623721244036 | 3.76490422001281e-06 |
| 2012 | 25,1943174306613 | + | + | NA | + | + | NA | + | + | + | + | + | NA | 0,123743960818258 | 11 | -521,058132108937 | 1066,22826421787 | 18,2666913025837 | 3.75678234704482e-06 |
| 106 | 25,1746980440557 | + | NA | NA | + | NA | + | + | NA | NA | NA | NA | NA | 0,047191175088188 | 6 | -526,792678688855 | 1066,23151122386 | 18,2699383085733 | 3.75068814796742e-06 |
| 209 | 26,5056523531054 | NA | NA | NA | NA | + | NA | + | + | NA | NA | NA | NA | 0,031818827221646 | 5 | -527,88852062552 | 1066,23505651822 | 18,2734836029249 | 3.74404539058657e-06 |
| 1069 | 25,2736946146985 | NA | NA | + | + | NA | + | NA | NA | NA | NA | + | NA | 0,0471384998084417 | 6 | -526,796463821037 | 1066,23908148823 | 18,2775085729374 | 3.73651813216307e-06 |
| 1217 | 26,4829581043005 | NA | NA | NA | NA | NA | NA | + | + | NA | NA | + | NA | 0,0317825487982708 | 5 | -527,891086165347 | 1066,24018759787 | 18,2786146825786 | 3.73445220413637e-06 |
| 422 | 26,1046133614505 | + | NA | + | NA | NA | + | NA | + | + | NA | NA | NA | 0,0624271276607901 | 7 | -525,688981672348 | 1066,24618039896 | 18,2846074836689 | 3.72327903751708e-06 |
| 2008 | 25,3967946314952 | + | + | + | NA | + | NA | + | + | + | + | + | NA | 0,12348382002122 | 11 | -521,078455100624 | 1066,26891020125 | 18,3073372859571 | 3.68120388218553e-06 |
| 3113 | 25,3575647758333 | NA | NA | NA | + | NA | + | NA | NA | NA | NA | + | + | 0,0469124091194995 | 6 | -526,812707836071 | 1066,2715695183 | 18,3099966030054 | 3.67631239079182e-06 |
| 1077 | 25,5991710489876 | NA | NA | + | NA | + | + | NA | NA | NA | NA | + | NA | 0,0468596965879446 | 6 | -526,816494537665 | 1066,27914292148 | 18,3175700061934 | 3.66241761708573e-06 |
| 116 | 25,040738351402 | + | + | NA | NA | + | + | + | NA | NA | NA | NA | NA | 0,0621946788954703 | 7 | -525,705954615152 | 1066,28012628457 | 18,3185533692761 | 3.66061731657023e-06 |
| 3109 | 25,3490290560196 | NA | NA | + | NA | NA | + | NA | NA | NA | NA | + | + | 0,0468327635759403 | 6 | -526,818429239289 | 1066,28301232473 | 18,3214394094407 | 3.65533878170061e-06 |
| 1347 | 25,6134344282631 | NA | + | NA | NA | NA | NA | + | NA | + | NA | + | NA | 0,0467907825665828 | 6 | -526,821444787354 | 1066,28904342086 | 18,3274705055719 | 3.64433253518149e-06 |
| 2101 | 25,3751644399803 | NA | NA | + | NA | + | + | NA | NA | NA | NA | NA | + | 0,046710795647001 | 6 | -526,827189979775 | 1066,3005338057 | 18,3389608904126 | 3.62345517317935e-06 |
| 3171 | 24,6918381859905 | NA | + | NA | NA | NA | + | + | NA | NA | NA | + | + | 0,0618851913465077 | 7 | -525,728546250198 | 1066,32530955466 | 18,3637366393691 | 3.57884514750009e-06 |
| 1750 | 25,2878209521946 | + | NA | + | NA | + | NA | + | + | NA | + | + | NA | 0,0925219006787076 | 9 | -523,455221034217 | 1066,32776490308 | 18,3661919877891 | 3.57445418749665e-06 |
| 1081 | 25,5926272050524 | NA | NA | NA | + | + | + | NA | NA | NA | NA | + | NA | 0,0464603492461116 | 6 | -526,845175589837 | 1066,33650502583 | 18,3749321105379 | 3.5588676851085e-06 |
| 2656 | 24,4232189121282 | + | + | + | + | + | NA | + | NA | NA | + | NA | + | 0,107723137013347 | 10 | -522,298613184925 | 1066,34325811588 | 18,3816852005918 | 3.54687127273558e-06 |
| 1027 | 26,5666650839925 | NA | + | NA | NA | NA | NA | NA | NA | NA | NA | + | NA | 0,0156276861005253 | 4 | -529,024083544652 | 1066,35119739233 | 18,389624477044 | 3.53281938585454e-06 |
| 2783 | 24,4465188964526 | NA | + | + | + | + | NA | + | + | NA | + | NA | + | 0,107633220660525 | 10 | -522,305512336396 | 1066,35705641882 | 18,3954835035336 | 3.52248508929233e-06 |
| 3824 | 24,3605167493111 | + | + | + | + | NA | + | + | + | NA | + | + | + | 0,138143940120777 | 12 | -519,923652113607 | 1066,36343325947 | 18,4018603441809 | 3.51127181206381e-06 |
| 131 | 26,6714345725688 | NA | + | NA | NA | NA | NA | NA | + | NA | NA | NA | NA | 0,015519935712071 | 4 | -529,031577895708 | 1066,36618609445 | 18,4046131791558 | 3.50644216070928e-06 |
| 2249 | 25,6821464529569 | NA | NA | NA | + | NA | NA | + | + | NA | NA | NA | + | 0,0462476325598091 | 6 | -526,860447962365 | 1066,36704977088 | 18,4054768555925 | 3.50492827187496e-06 |
| 2067 | 25,6282650351123 | NA | + | NA | NA | + | NA | NA | NA | NA | NA | NA | + | 0,0308738948598383 | 5 | -527,9553130888 | 1066,36864144477 | 18,4070685294841 | 3.5021400301038e-06 |

| | | | | | | | | | | | | | | | | | | |
|---|---|---|---|---|---|---|---|---|---|---|---|---|---|---|---|---|---|---|
| **19** | 26,6042307924437 | NA | + | NA | NA | + | NA | NA | NA | NA | NA | NA | NA | 0,0154953317997169 | 4 | -529,033289053968 | 1066,36960841097 | 18,4080354956754 | 3.50044721385764e-06 |
| **54** | 25,4937315786346 | + | NA | + | NA | + | + | NA | NA | NA | NA | NA | NA | 0,046197655055718 | 6 | -526,864035692158 | 1066,37422523047 | 18,4126523151788 | 3.49237606671548e-06 |
| **17** | 27,3068181818182 | NA | NA | NA | NA | + | NA | NA | NA | NA | NA | NA | NA | 2,02649229417243E- | 3 | -530,101171619814 | 1066,38279436745 | 18,4212214521563 | 3.47744475223078e-06 |
| **1445** | 26,2087201585995 | NA | NA | + | NA | NA | + | NA | + | + | NA | + | NA | 0,0614893027232168 | 7 | -525,757434039709 | 1066,38308513368 | 18,4215122183909 | 3.47693922722061e-06 |
| **1025** | 27,2351648351649 | NA | NA | NA | NA | NA | NA | NA | NA | NA | NA | + | NA | 1,57632486041663E- | 3 | -530,101479855724 | 1066,38341083927 | 18,4218379239767 | 3.47637304406033e-06 |
| **437** | 26,2295922407839 | NA | NA | + | NA | + | + | NA | + | + | NA | NA | NA | 0,0614782336692648 | 7 | -525,758241567843 | 1066,38470018995 | 18,4231272746583 | 3.47413263432937e-06 |
| **317** | 25,7948032847509 | NA | NA | + | + | + | + | NA | NA | + | NA | NA | NA | 0,0614775535897953 | 7 | -525,758291181827 | 1066,38479941792 | 18,4232265026269 | 3.47396027304324e-06 |
| **2816** | 24,2638512628515 | + | + | + | + | + | + | + | + | NA | + | NA | + | 0,138006731531041 | 12 | -519,934551022613 | 1066,38523107748 | 18,4236581621938 | 3.47321056985692e-06 |
| **3676** | 24,4288552129746 | + | + | NA | + | + | NA | + | NA | NA | + | + | + | 0,107438127396491 | 10 | -522,320479170865 | 1066,38699008776 | 18,4254171724715 | 3.47015720623367e-06 |
| **3832** | 24,5094996357808 | + | + | + | NA | + | + | + | + | NA | + | + | + | 0,137992536415686 | 12 | -519,935678486211 | 1066,38748600468 | 18,425913089389 | 3.46929685807083e-06 |
| **2105** | 25,3628838188647 | NA | NA | NA | + | + | + | NA | NA | NA | NA | NA | + | 0,0460459790240828 | 6 | -526,874922893003 | 1066,39599963216 | 18,4344267168697 | 3.4545600956157e-06 |
| **1066** | 25,5271358109843 | + | NA | NA | + | NA | + | NA | NA | NA | NA | + | NA | 0,0460199435560119 | 6 | -526,876791526937 | 1066,39973690003 | 18,4381639847372 | 3.44811081494373e-06 |
| **1124** | 24,9662403082292 | + | + | NA | NA | NA | + | + | NA | NA | NA | + | NA | 0,0613739748316315 | 7 | -525,765847165833 | 1066,39991138593 | 18,4383384706387 | 3.44781000470383e-06 |
| **2776** | 24,6296564326262 | + | + | + | NA | + | NA | + | + | NA | + | NA | + | 0,107339136239661 | 10 | -522,328072156276 | 1066,40217605858 | 18,4406031432939 | 3.44390813371688e-06 |
| **58** | 25,5561004428143 | + | NA | NA | + | + | + | NA | NA | NA | NA | NA | NA | 0,0459725699527026 | 6 | -526,880191524131 | 1066,40653689442 | 18,4449639791255 | 3.43640715532723e-06 |
| **2123** | 24,4767270492356 | NA | + | NA | + | NA | NA | + | NA | NA | NA | NA | + | 0,0459438333252186 | 6 | -526,882253865826 | 1066,4106615778 | 18,4490886625142 | 3.4293274125058e-06 |
| **2119** | 24,4861765019947 | NA | + | + | NA | NA | NA | + | NA | NA | NA | NA | + | 0,0459387655910594 | 6 | -526,882617555489 | 1066,41138895713 | 18,4498160418411 | 3.42808042834462e-06 |
| **7** | 26,0361096184847 | NA | + | + | NA | NA | NA | NA | NA | NA | NA | NA | NA | 0,0151667071869339 | 4 | -529,056140212787 | 1066,4153107286 | 18,4537378133127 | 3.42136494063534e-06 |
| **2055** | 25,1863054485808 | NA | + | + | NA | NA | NA | NA | NA | NA | NA | NA | + | 0,0304892577130839 | 5 | -527,982482492951 | 1066,42298025308 | 18,4614073377877 | 3.40826994370744e-06 |
| **2093** | 24,9242738828502 | NA | NA | + | + | NA | + | NA | NA | NA | NA | NA | + | 0,0458402727090547 | 6 | -526,889685586068 | 1066,42552501829 | 18,463952102999 | 3.40393607806809e-06 |
| **373** | 25,8464833265593 | NA | NA | + | NA | + | + | + | NA | + | NA | NA | NA | 0,061185970097446 | 7 | -525,779559825353 | 1066,42733670497 | 18,4657637896792 | 3.40085404137362e-06 |
| **1759** | 24,8497164450691 | NA | + | + | + | + | NA | + | + | NA | + | + | NA | 0,107122793528429 | 10 | -522,344663507918 | 1066,43535876187 | 18,4737858465767 | 3.38724043958732e-06 |
| **3839** | 24,2780266278674 | NA | + | + | + | + | + | + | + | NA | + | + | + | 0,137679876652666 | 12 | -519,960507148011 | 1066,43714332828 | 18,47557041299 | 3.38421940983363e-06 |
| **2185** | 26,0233859192401 | NA | NA | NA | + | NA | NA | NA | + | NA | NA | NA | + | 0,0303709859430581 | 5 | -527,99083462562 | 1066,43968451842 | 18,4781116031252 | 3.37992216794918e-06 |
| **87** | 25,2735711403245 | NA | + | + | NA | + | NA | + | NA | NA | NA | NA | NA | 0,0457143818887364 | 6 | -526,898718681458 | 1066,44359120907 | 18,4820182937801 | 3.37332645672322e-06 |
| **1062** | 25,462938202109 | + | NA | + | NA | NA | + | NA | NA | NA | NA | + | NA | 0,0455604433891677 | 6 | -526,909762674822 | 1066,4656791958 | 18,5041062805078 | 3.33627642843561e-06 |
| **119** | 24,8208849998685 | NA | + | + | NA | + | + | + | NA | NA | NA | NA | NA | 0,0609166165545257 | 7 | -525,799201108926 | 1066,46661927212 | 18,5050463568248 | 3.33470861970083e-06 |
| **91** | 25,2720191164584 | NA | + | NA | + | + | NA | + | NA | NA | NA | NA | NA | 0,0455181000349257 | 6 | -526,912800197348 | 1066,47175424085 | 18,5101813255583 | 3.32615778921112e-06 |
| **1952** | 25,473542659972 | + | + | + | + | + | NA | NA | + | + | + | + | NA | 0,122169255306358 | 11 | -521,181060783937 | 1066,47412156787 | 18,5125486525842 | 3.32222306675079e-06 |
| **84** | 25,4121330685457 | + | + | NA | NA | + | NA | + | NA | NA | NA | NA | NA | 0,0455003181008274 | 6 | -526,914075753475 | 1066,4743053531 | 18,5127324378122 | 3.32191779301523e-06 |
| **2405** | 25,4921010316652 | NA | NA | + | NA | NA | + | + | NA | + | NA | NA | + | 0,0608618954498177 | 7 | -525,803190688398 | 1066,47459843106 | 18,5130255157692 | 3.32143103824043e-06 |
| **2163** | 24,7922095445821 | NA | + | NA | NA | + | + | + | NA | NA | NA | NA | + | 0,0608115334590093 | 7 | -525,806862250409 | 1066,48194155508 | 18,5203686397915 | 3.30925855790004e-06 |
| **2090** | 25,1949936125942 | + | NA | NA | + | NA | + | NA | NA | NA | NA | NA | + | 0,0454341050222931 | 6 | -526,918825224666 | 1066,48380429549 | 18,522231380196 | 3.30617784795114e-06 |
| **46** | 25,1092967305183 | + | NA | + | + | NA | + | NA | NA | NA | NA | NA | NA | 0,0454317273961097 | 6 | -526,918995765922 | 1066,484145378 | 18,5225724627066 | 3.30561405630684e-06 |
| **2349** | 25,4630001886713 | NA | NA | + | + | NA | + | NA | NA | + | NA | NA | + | 0,060789910345168 | 7 | -525,808438589252 | 1066,48509423277 | 18,523521317476 | 3.30404615443252e-06 |
| **3803** | 24,4153594193247 | NA | + | NA | + | + | NA | + | + | NA | + | + | + | 0,106746396611873 | 10 | -522,37351986371 | 1066,49307147345 | 18,5314985581613 | 3.29089381600708e-06 |
| **123** | 24,8203909888918 | NA | + | NA | + | + | + | + | NA | NA | NA | NA | NA | 0,0607284608572671 | 7 | -525,812918098246 | 1066,49405325076 | 18,5324803354645 | 3.28927875002041e-06 |
| **3796** | 24,570948004907 | + | + | NA | NA | + | NA | + | + | NA | + | + | + | 0,106717644105131 | 10 | -522,375723666266 | 1066,49747907856 | 18,5359061632739 | 3.28364932145264e-06 |
| **1694** | 25,2976938890933 | + | NA | + | + | + | NA | NA | + | NA | + | + | NA | 0,091388067368074 | 9 | -523,540712757352 | 1066,49874834935 | 18,5371754340597 | 3.28156606250028e-06 |
| **4** | 26,0551618420628 | + | + | NA | NA | NA | NA | NA | NA | NA | NA | NA | NA | 0,0145498688634398 | 4 | -529,09901195579 | 1066,50105421461 | 18,539481299319 | 3.27778481808926e-06 |
| **1632** | 24,8290970105435 | + | + | + | + | + | NA | + | NA | NA | + | + | NA | 0,106667688582583 | 10 | -522,37955245454 | 1066,50513665511 | 18,5435637398211 | 3.27110096127269e-06 |
| **3616** | 24,7322713559881 | + | + | + | + | + | NA | NA | NA | NA | + | + | + | 0,106650378489277 | 10 | -522,380879118438 | 1066,50778998291 | 18,5462170676167 | 3.26676418707836e-06 |
| **2768** | 24,4567584585568 | + | + | + | + | NA | NA | + | + | NA | + | NA | + | 0,106643103312931 | 10 | -522,381436687995 | 1066,50890512202 | 18,5473322067317 | 3.26494324651384e-06 |
| **2387** | 25,3062614769115 | NA | + | NA | NA | + | NA | + | NA | + | NA | NA | + | 0,0605093115686229 | 7 | -525,828891130701 | 1066,52599931567 | 18,5644264003754 | 3.23715637815408e-06 |
| **331** | 25,4258585021284 | NA | + | NA | + | NA | NA | + | NA | + | NA | NA | NA | 0,0451317828110484 | 6 | -526,940506645749 | 1066,52716713765 | 18,5655942223616 | 3.23526671870865e-06 |
| **327** | 25,4447331173778 | NA | + | + | NA | NA | NA | + | NA | + | NA | NA | NA | 0,0450577086339538 | 6 | -526,945817922826 | 1066,53778969181 | 18,5762167765158 | 3.21812887295859e-06 |
| **365** | 25,5119145404315 | NA | NA | + | + | NA | + | + | NA | + | NA | NA | NA | 0,0603357299287661 | 7 | -525,841540252165 | 1066,55129755859 | 18,5897246433033 | 3.19646707839997e-06 |
| **1131** | 24,7881182536296 | NA | + | NA | + | NA | + | + | NA | NA | NA | + | NA | 0,0603189089671525 | 7 | -525,842765893539 | 1066,55374884134 | 18,5921759260507 | 3.19255175597969e-06 |
| **324** | 25,4698856222978 | + | + | NA | NA | NA | NA | + | NA | + | NA | NA | NA | 0,0448726588551344 | 6 | -526,959084588955 | 1066,56432302406 | 18,6027501087722 | 3.17571698587138e-06 |
| **2086** | 25,158466294637 | + | NA | + | NA | NA | + | NA | NA | NA | NA | NA | + | 0,0448699041399746 | 6 | -526,959282061738 | 1066,56471796963 | 18,6031450543387 | 3.17508993011456e-06 |
| **3395** | 25,1597014773535 | NA | + | NA | NA | NA | NA | + | NA | + | NA | + | + | 0,0601971028474332 | 7 | -525,851640511843 | 1066,57149807795 | 18,6099251626581 | 3.16434442749852e-06 |
| **2357** | 25,8623405502682 | NA | NA | + | NA | + | + | NA | NA | + | NA | NA | + | 0,0601599976693087 | 7 | -525,85434371318 | 1066,57690448062 | 18,6153315653339 | 3.15580211840096e-06 |
| **2052** | 25,3938933484689 | + | + | NA | NA | NA | NA | NA | NA | NA | NA | NA | + | 0,0293708283093736 | 5 | -528,06142321095 | 1066,58086168908 | 18,6192887737855 | 3.14956420820658e-06 |
| **1127** | 24,8418188185309 | NA | + | + | NA | NA | + | + | NA | NA | NA | + | NA | 0,0601271417685799 | 7 | -525,856737255408 | 1066,58169156508 | 18,6201186497883 | 3.14825760542698e-06 |
| **3121** | 25,7182504596035 | NA | NA | NA | NA | + | + | NA | NA | NA | NA | + | + | 0,0447240002461836 | 6 | -526,96974042327 | 1066,58563469269 | 18,6240617774033 | 3.14205672939541e-06 |
| **2257** | 25,7179737337372 | NA | NA | NA | NA | + | NA | + | + | NA | NA | NA | + | 0,0447090847994103 | 6 | -526,970809469517 | 1066,58777278519 | 18,6261998698978 | 3.1386995202658e-06 |
| **2245** | 25,6938122339089 | NA | NA | + | NA | NA | NA | + | + | NA | NA | NA | + | 0,0446749982978533 | 6 | -526,973252514836 | 1066,59265887583 | 18,6310859605353 | 3.13104089409052e-06 |
| **2481** | 26,3163341763414 | NA | NA | NA | NA | + | + | NA | + | + | NA | NA | + | 0,0600498189328542 | 7 | -525,862369870127 | 1066,59295679452 | 18,6313838792262 | 3.13057453102383e-06 |
| **3106** | 25,6790875850522 | + | NA | NA | NA | NA | + | NA | NA | NA | NA | + | + | 0,0446425515459773 | 6 | -526,97557795519 | 1066,59730975653 | 18,6357368412439 | 3.12376830450841e-06 |
| **3235** | 25,5294283844431 | NA | + | NA | NA | NA | + | NA | + | NA | NA | + | + | 0,0600184312889738 | 7 | -525,864656184182 | 1066,59752942263 | 18,6359565073365 | 3.12342523036044e-06 |
| **2000** | 25,3004469429858 | + | + | + | + | NA | NA | + | + | + | + | + | NA | 0,121311389765074 | 11 | -521,247936959592 | 1066,60787391918 | 18,6463010038942 | 3.10731180682444e-06 |
| **2475** | 25,6246438403526 | NA | + | NA | + | NA | + | NA | + | + | NA | NA | + | 0,0752902796317793 | 8 | -524,74312340986 | 1066,61124681972 | 18,6496739044298 | 3.10207589628797e-06 |
| **2251** | 24,9131498749394 | NA | + | NA | + | NA | NA | + | + | NA | NA | NA | + | 0,0599087462090779 | 7 | -525,872645179236 | 1066,61350741274 | 18,6519344974449 | 3.09857161154976e-06 |
| **325** | 26,1909199037564 | NA | NA | + | NA | NA | NA | + | NA | + | NA | NA | NA | 0,0291164884806355 | 5 | -528,079362275028 | 1066,61673981723 | 18,6551669019416 | 3.09356773786973e-06 |
| **3743** | 24,722625766599 | NA | + | + | + | + | NA | NA | + | NA | + | + | + | 0,10588853223273 | 10 | -522,439242387243 | 1066,62451652052 | 18,6629436052281 | 3.08156221464692e-06 |
| **1451** | 25,8211946584356 | NA | + | NA | + | NA | + | NA | + | + | NA | + | NA | 0,0751737273261322 | 8 | -524,751752679249 | 1066,6285053585 | 18,6669324432066 | 3.07542241315291e-06 |
| **736** | 24,793990312713 | + | + | + | + | + | NA | + | + | NA | + | NA | NA | 0,105854021954334 | 10 | -522,441884962446 | 1066,62980167092 | 18,6682287556328 | 3.07342970487093e-06 |
| **2466** | 26,3902592939796 | + | NA | NA | NA | NA | + | NA | + | + | NA | NA | + | 0,0596188055401149 | 7 | -525,893758739658 | 1066,65573453358 | 18,6941616182896 | 3.03383554189773e-06 |
| **1325** | 25,6446283283157 | NA | NA | + | + | NA | + | NA | NA | + | NA | + | NA | 0,0595980705840208 | 7 | -525,895268415635 | 1066,65875388553 | 18,6971809702432 | 3.02925888876355e-06 |
| **1752** | 25,0127183830684 | + | + | + | NA | + | NA | + | + | NA | + | + | NA | 0,105620935070031 | 10 | -522,459730588145 | 1066,66549292232 | 18,703920007031 | 3.0190689224984e-06 |
| **2244** | 25,0545892906419 | + | + | NA | NA | NA | NA | + | + | NA | NA | NA | + | 0,0595087632955663 | 7 | -525,901770343254 | 1066,67175774077 | 18,7101848254802 | 3.00962675923611e-06 |
| **302** | 25,5830909802042 | + | NA | + | + | NA | + | NA | NA | + | NA | NA | NA | 0,0594482482534992 | 7 | -525,906175729484 | 1066,68056851323 | 18,7189955979411 | 2.99639735267059e-06 |
| **3664** | 24,3330123126168 | + | + | + | + | NA | NA | + | NA | NA | + | + | + | 0,105522174287743 | 10 | -522,467290520838 | 1066,68061278771 | 18,7190398724167 | 2.99633102144403e-06 |
| **310** | 25,9168437721801 | + | NA | + | NA | + | + | NA | NA | + | NA | NA | NA | 0,0594360912147368 | 7 | -525,907060705868 | 1066,682338466 | 18,7207655507086 | 2.9937467847951e-06 |
| **2409** | 25,5784966274259 | NA | NA | NA | + | NA | + | + | NA | + | NA | NA | + | 0,0594320523287233 | 7 | -525,907354715622 | 1066,68292648551 | 18,7213535702172 | 2.99286672341821e-06 |
| **358** | 25,5961740623321 | + | NA | + | NA | NA | + | + | NA | + | NA | NA | NA | 0,0594248168033227 | 7 | -525,907881420831 | 1066,68397989593 | 18,7224069806348 | 2.99129077999094e-06 |

| | | | | | | | | | | | | | | | | | | |
|---|---|---|---|---|---|---|---|---|---|---|---|---|---|---|---|---|---|---|
| **2219** | 25,1068725357918 | NA | + | NA | + | NA | + | NA | + | NA | NA | NA | + | 0,0593868880145917 | 7 | -525,91064235526 | 1066,68950176478 | 18,7279288494929 | 2.98304341274827e-06 |
| **1381** | 25,6747788010083 | NA | NA | + | NA | NA | + | + | NA | + | NA | + | NA | 0,0593793424530654 | 7 | -525,911191602888 | 1066,69060026004 | 18,7290273447493 | 2.98140543309835e-06 |
| **3728** | 24,7750611215655 | + | + | + | + | NA | NA | NA | + | NA | + | + | + | 0,105445459564793 | 10 | -522,473162297787 | 1066,69235634161 | 18,7307834263145 | 2.97878878646798e-06 |
| **2259** | 25,0587163681731 | NA | + | NA | NA | + | NA | + | + | NA | NA | NA | + | 0,0592855656596332 | 7 | -525,91801732534 | 1066,70425170494 | 18,7426787896538 | 2.96112448190781e-06 |
| **172** | 25,2688219175694 | + | + | NA | + | NA | + | NA | + | NA | NA | NA | NA | 0,0592767931282961 | 7 | -525,918655815948 | 1066,70552868616 | 18,7439557708692 | 2.95923443519068e-06 |
| **4061** | 25,339760184939 | NA | NA | + | + | + | NA | + | + | + | + | + | + | 0,120682929812887 | 11 | -521,296888081079 | 1066,70577616216 | 18,7442032468675 | 2.95886828809624e-06 |
| **3265** | 25,9358046301525 | NA | NA | NA | NA | NA | NA | + | + | NA | NA | + | + | 0,0438630180754361 | 6 | -527,031422959633 | 1066,70899976542 | 18,7474268501298 | 2.95410302072761e-06 |
| **2242** | 25,8587933762788 | + | NA | NA | NA | NA | NA | + | + | NA | NA | NA | + | 0,043843964267963 | 6 | -527,032787385649 | 1066,71172861745 | 18,7501557021617 | 2.9500751142275e-06 |
| **3365** | 25,7271355877004 | NA | NA | + | NA | NA | + | NA | NA | + | NA | + | + | 0,0592327021697749 | 7 | -525,921864795343 | 1066,71194664495 | 18,7503737296597 | 2.94975353300814e-06 |
| **2247** | 25,0667746383447 | NA | + | + | NA | NA | NA | + | + | NA | NA | NA | + | 0,0592052301188923 | 7 | -525,923864159766 | 1066,71594537379 | 18,7543724585043 | 2.94386179257041e-06 |
| **3038** | 25,3532871543279 | + | NA | + | + | + | NA | + | + | + | + | NA | + | 0,120604195481865 | 11 | -521,30301828017 | 1066,71803656034 | 18,7564636450491 | 2.94078531913561e-06 |
| **2342** | 25,6255175340593 | + | NA | + | NA | NA | + | NA | NA | + | NA | NA | + | 0,0591546011967701 | 7 | -525,927548685048 | 1066,72331442436 | 18,761741509069 | 2.93303501737992e-06 |
| **3489** | 26,4479736957532 | NA | NA | NA | NA | NA | + | NA | + | + | NA | + | + | 0,0591509501275416 | 7 | -525,927814384345 | 1066,72384582295 | 18,7622729076625 | 2.93225581555978e-06 |
| **2780** | 24,3722438061422 | + | + | NA | + | + | NA | + | + | NA | + | NA | + | 0,105193716329459 | 10 | -522,492427295081 | 1066,73088633619 | 18,7693134209037 | 2.92195166987117e-06 |
| **187** | 25,2633580948491 | NA | + | NA | + | + | + | NA | + | NA | NA | NA | NA | 0,059086245538903 | 7 | -525,932522961042 | 1066,73326297635 | 18,7716900610567 | 2.91848151827307e-06 |
| **1333** | 25,9951615425671 | NA | NA | + | NA | + | + | NA | NA | + | NA | + | NA | 0,0590758014364777 | 7 | -525,933282951877 | 1066,73478295802 | 18,7732100427277 | 2.9162643416893e-06 |
| **2098** | 25,6481702208866 | + | NA | NA | NA | + | + | NA | NA | NA | NA | NA | + | 0,0436787220232405 | 6 | -527,044619094448 | 1066,73539203505 | 18,7738191197591 | 2.91537636209413e-06 |
| **1074** | 25,8641104490922 | + | NA | NA | NA | + | + | NA | NA | NA | NA | + | NA | 0,0436005465582077 | 6 | -527,050215917945 | 1066,74658568204 | 18,7850127667523 | 2.89910509135851e-06 |
| **2433** | 26,9224246982144 | NA | NA | NA | NA | NA | NA | NA | + | + | NA | NA | + | 0,0281653448070021 | 5 | -528,146406614128 | 1066,75082849543 | 18,7892555801404 | 2.89296142931783e-06 |
| **434** | 26,3888857696267 | + | NA | NA | NA | + | + | NA | + | + | NA | NA | NA | 0,0589621448413175 | 7 | -525,941552908834 | 1066,75132287193 | 18,7897499566402 | 2.89224641162076e-06 |
| **377** | 25,8848200935208 | NA | NA | NA | + | + | + | + | NA | + | NA | NA | NA | 0,0589594435417717 | 7 | -525,941749450446 | 1066,75171595516 | 18,7901430398647 | 2.89167802070608e-06 |
| **431** | 25,6718127664419 | NA | + | + | + | NA | + | NA | + | + | NA | NA | NA | 0,0743298134020953 | 8 | -524,81420174921 | 1066,75340349842 | 18,7918305831297 | 2.88923913389833e-06 |
| **2373** | 25,6088035481538 | NA | NA | + | NA | NA | NA | + | NA | + | NA | NA | + | 0,043511021396456 | 6 | -527,056624739711 | 1066,75940332558 | 18,7978304102851 | 2.88058465402766e-06 |
| **375** | 25,3639059416549 | NA | + | + | NA | + | + | + | NA | + | NA | NA | NA | 0,0742872728026438 | 8 | -524,81734821854 | 1066,75969643708 | 18,7981235217885 | 2.88016251871225e-06 |
| **2315** | 25,5462730604409 | NA | + | NA | + | NA | NA | NA | NA | + | NA | NA | + | 0,0435052325786989 | 6 | -527,057039122136 | 1066,76023209042 | 18,7986591751342 | 2.8793912376571e-06 |
| **2281** | 25,4320385375631 | NA | NA | NA | + | NA | + | + | + | NA | NA | NA | + | 0,0588550386786667 | 7 | -525,949345325188 | 1066,76690770464 | 18,8053347893494 | 2.86979640687352e-06 |
| **137** | 26,9100247450058 | NA | NA | NA | + | NA | NA | NA | + | NA | NA | NA | NA | 0,0125818584945908 | 4 | -529,235614305944 | 1066,77425891492 | 18,8126859996282 | 2.85926753030165e-06 |
| **443** | 25,7841202357173 | NA | + | NA | + | + | + | NA | + | + | NA | NA | NA | 0,0741341477692759 | 8 | -524,828672749403 | 1066,78234549881 | 18,8207725835161 | 2.84773001741535e-06 |
| **1191** | 25,5739166738939 | NA | + | + | NA | NA | + | NA | + | NA | NA | + | NA | 0,0587234259773814 | 7 | -525,958919479864 | 1066,78605601399 | 18,8244830987003 | 2.84245164256689e-06 |
| **1395** | 25,5105935390842 | NA | + | NA | NA | + | + | + | NA | + | NA | + | NA | 0,0740815650344666 | 8 | -524,832561131799 | 1066,7901222636 | 18,8285493483079 | 2.83667845442345e-06 |
| **175** | 25,1908208242968 | NA | + | + | + | NA | + | NA | + | NA | NA | NA | NA | 0,0586549788698351 | 7 | -525,963898129954 | 1066,79601331417 | 18,834440398882 | 2.82833523991165e-06 |
| **428** | 25,7688126569186 | + | + | NA | + | NA | + | NA | + | + | NA | NA | NA | 0,0740238580844392 | 8 | -524,836828185089 | 1066,79865637018 | 18,8370834548871 | 2.82459998435749e-06 |
| **3836** | 24,191260992299 | + | + | NA | + | + | + | + | + | NA | + | + | + | 0,13531298662909 | 12 | -520,148173170353 | 1066,81247537296 | 18,850902457674 | 2.80515067671644e-06 |
| **3998** | 25,3689140131738 | + | NA | + | + | + | NA | NA | + | + | + | + | + | 0,119965451060773 | 11 | -521,352730198999 | 1066,817460398 | 18,8558874827067 | 2.79816750997499e-06 |
| **1203** | 25,6887251324443 | NA | + | NA | NA | + | + | NA | + | NA | NA | + | NA | 0,0584980144604126 | 7 | -525,975313913433 | 1066,81884488113 | 18,8572719658398 | 2.79623117239849e-06 |
| **1744** | 24,9339510668529 | + | + | + | + | NA | NA | + | + | NA | + | + | NA | 0,104580594427868 | 10 | -522,539324637115 | 1066,82468102026 | 18,8631081049716 | 2.78808346887947e-06 |
| **1188** | 25,6907128646523 | + | + | NA | NA | NA | + | NA | + | NA | NA | + | NA | 0,0584456383537218 | 7 | -525,979122724964 | 1066,82646250419 | 18,8648895889007 | 2.78560111166706e-06 |
| **2361** | 25,8490625837256 | NA | NA | NA | + | + | + | NA | NA | + | NA | NA | + | 0,0584346571574623 | 7 | -525,979921255115 | 1066,82805956449 | 18,866486649202 | 2.78337761307524e-06 |
| **1442** | 26,4687373647596 | + | NA | NA | NA | NA | + | NA | + | + | NA | + | NA | 0,0583875686771054 | 7 | -525,983345327637 | 1066,83490770954 | 18,8733347942466 | 2.77386342420303e-06 |
| **2417** | 25,8808599366353 | NA | NA | NA | NA | + | + | + | NA | + | NA | NA | + | 0,0583776937313338 | 7 | -525,98406336967 | 1066,8363437936 | 18,8747708783137 | 2.77187238857829e-06 |
| **329** | 26,2786121074926 | NA | NA | NA | + | NA | NA | + | NA | + | NA | NA | NA | 0,0275504816425138 | 5 | -528,189712274271 | 1066,83743981572 | 18,8758669004264 | 2.77035378800547e-06 |
| **2419** | 25,3174012396615 | NA | + | NA | NA | + | + | + | NA | + | NA | NA | + | 0,0737579636208066 | 8 | -524,856485916473 | 1066,83797183295 | 18,8763989176562 | 2.76961694803889e-06 |
| **379** | 25,3511694649264 | NA | + | NA | + | + | + | + | NA | + | NA | NA | NA | 0,0737368968082904 | 8 | -524,858043156941 | 1066,84108631388 | 18,8795133985923 | 2.76530734486151e-06 |
| **85** | 26,0288143243964 | NA | NA | + | NA | + | NA | + | NA | NA | NA | NA | NA | 0,0274853917907474 | 5 | -528,194295040131 | 1066,84660534744 | 18,8850324321477 | 2.7576869519846e-06 |
| **322** | 26,2871725575604 | + | NA | NA | NA | NA | NA | + | NA | + | NA | NA | NA | 0,0274685826961634 | 5 | -528,195478464284 | 1066,84897219574 | 18,8873992804533 | 2.75442536893841e-06 |
| **4058** | 25,2330884629829 | + | NA | NA | + | + | NA | + | + | + | + | + | + | 0,119732528480664 | 11 | -521,370849021963 | 1066,85369804393 | 18,8921251286363 | 2.74792455436639e-06 |
| **4046** | 25,2805607710697 | + | NA | + | + | NA | NA | + | + | + | + | + | + | 0,119725409536978 | 11 | -521,371402722259 | 1066,85480544452 | 18,8932325292283 | 2.74640344888441e-06 |
| **3369** | 25,7737412078449 | NA | NA | NA | + | NA | + | NA | NA | + | NA | + | + | 0,05819697784399 | 7 | -525,99720252827 | 1066,8626221108 | 18,9010491955128 | 2.73569053771274e-06 |
| **2306** | 26,4580647146461 | + | NA | NA | NA | NA | NA | NA | NA | + | NA | NA | + | 0,0273352485747974 | 5 | -528,204864969023 | 1066,86774520522 | 18,9061722899314 | 2.72869190474147e-06 |
| **2435** | 26,2513989375956 | NA | + | NA | NA | NA | NA | NA | + | + | NA | NA | + | 0,042742938084083 | 6 | -527,111584756441 | 1066,86932335904 | 18,9077504437448 | 2.72653960625013e-06 |
| **1083** | 25,2488576569563 | NA | + | NA | + | + | + | NA | NA | NA | NA | + | NA | 0,0581401864034368 | 7 | -526,001331095554 | 1066,87087924537 | 18,9093063300802 | 2.72441933822296e-06 |
| **1257** | 25,6551514097879 | NA | NA | NA | + | NA | + | + | + | NA | NA | + | NA | 0,05812612038036 | 7 | -526,002353614769 | 1066,8729242838 | 18,9113513685118 | 2.72163499086472e-06 |
| **3425** | 25,7964518620614 | NA | NA | NA | NA | NA | + | + | NA | + | NA | + | + | 0,0580895495784506 | 7 | -526,005012030924 | 1066,87824111611 | 18,9166682008217 | 2.71440936105793e-06 |
| **3331** | 26,0453857017565 | NA | + | NA | NA | NA | NA | NA | NA | + | NA | + | + | 0,0426582860218697 | 6 | -527,117639317487 | 1066,88143248113 | 18,9198595658372 | 2.71008147940247e-06 |
| **2346** | 25,7370214289127 | + | NA | NA | + | NA | + | NA | NA | + | NA | NA | + | 0,05806409872255 | 7 | -526,006862051609 | 1066,88194115748 | 18,9203682421908 | 2.70939228986741e-06 |
| **1385** | 25,7579804617413 | NA | NA | NA | + | NA | + | + | NA | + | NA | + | NA | 0,0580600194107732 | 7 | -526,00715857181 | 1066,88253419788 | 18,9209612825939 | 2.70858901941832e-06 |
| **237** | 25,4681375089713 | NA | NA | + | + | NA | + | + | + | NA | NA | NA | NA | 0,0580556799350291 | 7 | -526,007474001605 | 1066,88316505747 | 18,9215921421826 | 2.70773478447363e-06 |
| **3934** | 25,2463606814672 | + | NA | + | + | + | NA | + | NA | + | + | + | + | 0,119534761892494 | 11 | -521,386229332977 | 1066,88445866595 | 18,9228857506639 | 2.70598397640853e-06 |
| **372** | 25,5126543474775 | + | + | NA | NA | + | + | + | NA | + | NA | NA | NA | 0,0734157873847296 | 8 | -524,881774905201 | 1066,8885498104 | 18,926976895111 | 2.70045434830401e-06 |
| **362** | 25,7336816186465 | + | NA | NA | + | NA | + | + | NA | + | NA | NA | NA | 0,058014083730845 | 7 | -526,010497491799 | 1066,88921203786 | 18,9276391225712 | 2.69956033880932e-06 |
| **2402** | 25,7584944770533 | + | NA | NA | NA | NA | + | + | NA | + | NA | NA | + | 0,0580003185791187 | 7 | -526,01149800566 | 1066,89121306558 | 18,9296401502932 | 2.69686074198907e-06 |
| **1457** | 26,5102016899464 | NA | NA | NA | NA | + | + | NA | + | + | NA | + | NA | 0,0579834306803012 | 7 | -526,012725475027 | 1066,89366800432 | 18,932095089026 | 2.69355245886623e-06 |
| **3788** | 24,3661404487024 | + | + | NA | + | NA | NA | + | + | NA | + | + | + | 0,104126451444777 | 10 | -522,574041085826 | 1066,89411391768 | 18,9325410023926 | 2.69295198028662e-06 |
| **1318** | 25,7635794352198 | + | NA | + | NA | NA | + | NA | NA | + | NA | + | NA | 0,057960248818616 | 7 | -526,014410374995 | 1066,89703780425 | 18,9354648889623 | 2.68901791361439e-06 |
| **234** | 25,600670267181 | + | NA | NA | + | NA | + | + | + | NA | NA | NA | NA | 0,0579491736423144 | 7 | -526,015215324369 | 1066,898647703 | 18,9370747877113 | 2.68685426125903e-06 |
| **249** | 25,5177986698939 | NA | NA | NA | + | + | + | + | + | NA | NA | NA | NA | 0,05789463956918 | 7 | -526,019178750234 | 1066,90657455473 | 18,9450016394403 | 2.67622618928621e-06 |
| **459** | 25,8112205671519 | NA | + | NA | + | NA | NA | + | + | + | NA | NA | NA | 0,0577783496916056 | 7 | -526,027629696923 | 1066,92347644811 | 18,9619035328187 | 2.65370484182226e-06 |
| **265** | 26,947352198197 | NA | NA | NA | + | NA | NA | NA | NA | + | NA | NA | NA | 0,0114493650437227 | 4 | -529,314098951896 | 1066,93122820682 | 18,9696552915311 | 2.64343930884318e-06 |
| **1099** | 25,2042197229636 | NA | + | NA | + | NA | NA | + | NA | NA | NA | + | NA | 0,0421756764905644 | 6 | -527,152146732534 | 1066,95044731122 | 18,9888743959318 | 2.61815870300844e-06 |
| **2311** | 25,6766988442496 | NA | + | + | NA | NA | NA | NA | NA | + | NA | NA | + | 0,0421040419982902 | 6 | -527,157267241165 | 1066,96068832848 | 18,9991154131928 | 2.60478666381215e-06 |
| **1092** | 25,2895376972336 | + | + | NA | NA | NA | NA | + | NA | NA | NA | + | NA | 0,0420789754507391 | 6 | -527,159058933871 | 1066,9642717139 | 19,002698798605 | 2.6001238649444e-06 |
| **1756** | 24,7364645464483 | + | + | NA | + | + | NA | + | + | NA | + | + | NA | 0,103650927225897 | 10 | -522,610373152201 | 1066,96677805043 | 19,0052051351438 | 2.59686751302498e-06 |
| **337** | 26,627307697413 | NA | NA | NA | NA | + | NA | + | NA | + | NA | NA | NA | 0,0265849107350772 | 5 | -528,257663557201 | 1066,97334238158 | 19,0117694662872 | 2.58835813611044e-06 |
| **1095** | 25,2549559596435 | NA | + | + | NA | NA | NA | + | NA | NA | NA | + | NA | 0,042006609416486 | 6 | -527,164231209984 | 1066,97461626612 | 19,0130433508311 | 2.58671002632967e-06 |
| **3427** | 25,1573998948362 | NA | + | NA | NA | NA | + | + | NA | + | NA | + | + | 0,0728248409862664 | 8 | -524,925427609739 | 1066,97585521948 | 19,014282304187 | 2.58510811602011e-06 |
| **2181** | 26,0159938246002 | NA | NA | + | NA | NA | NA | NA | + | NA | NA | NA | + | 0,0265440029467847 | 5 | -528,260540922336 | 1066,97909711185 | 19,0175241965578 | 2.58092118918882e-06 |

| | | | | | | | | | | | | | | | | | | |
|---|---|---|---|---|---|---|---|---|---|---|---|---|---|---|---|---|---|---|
| 3740 | 24,6260120261366 | + | + | NA | + | + | NA | NA | + | NA | + | + | + | 0,103548557075908 | 10 | -522,618192145524 | 1066,98241603708 | 19,0208431217889 | 2.57664179868809e-06 |
| 13 | 26,1410188143415 | NA | NA | + | + | NA | NA | NA | NA | NA | NA | NA | NA | 0,0110761702235385 | 4 | -529,339942605338 | 1066,98291551371 | 19,0213425984148 | 2.57599839285707e-06 |
| 314 | 26,0086460710676 | + | NA | NA | + | + | + | NA | NA | + | NA | NA | NA | 0,0573293786399487 | 7 | -526,06024726641 | 1066,98871158708 | 19,0271386717925 | 2.56854386197343e-06 |
| 1337 | 26,0016441694556 | NA | NA | NA | + | + | + | NA | NA | + | NA | + | NA | 0,0573176433338431 | 7 | -526,061099623508 | 1066,99041630128 | 19,0288433859882 | 2.56635547815622e-06 |
| 3838 | 24,5852014515586 | + | NA | + | + | + | + | + | + | NA | + | + | + | 0,134156585593263 | 12 | -520,239675293941 | 1066,99547962014 | 19,0339067048499 | 2.55986655744157e-06 |
| 2377 | 25,7117519309661 | NA | NA | NA | + | NA | NA | + | NA | + | NA | NA | + | 0,0418222946273242 | 6 | -527,177403126183 | 1067,00096009852 | 19,0393871832291 | 2.55286151291279e-06 |
| 2321 | 26,5163806450794 | NA | NA | NA | NA | + | NA | NA | NA | + | NA | NA | + | 0,0263567463714683 | 5 | -528,273710601152 | 1067,00543646948 | 19,0438635541889 | 2.5471541248216e-06 |
| 2061 | 25,3370371994873 | NA | NA | + | + | NA | NA | NA | NA | NA | NA | NA | + | 0,0263069189937527 | 5 | -528,277214513905 | 1067,01244429499 | 19,0508713796955 | 2.53824473697314e-06 |
| 2720 | 24,6680049811026 | + | + | + | + | + | NA | NA | + | NA | + | NA | + | 0,103341197409648 | 10 | -522,634027463998 | 1067,01408667403 | 19,0525137587374 | 2.53616121259485e-06 |
| 2277 | 25,4742502542147 | NA | NA | + | NA | NA | + | + | + | NA | NA | NA | + | 0,0570961768342619 | 7 | -526,077183158668 | 1067,0225833716 | 19,0610104563086 | 2.52540956972257e-06 |
| 3329 | 26,6527572905331 | NA | NA | NA | NA | NA | NA | NA | NA | + | NA | + | + | 0,0262195578184784 | 5 | -528,28335740957 | 1067,02473008631 | 19,0631571710242 | 2.52270035701633e-06 |
| 2372 | 25,0683314057223 | + | + | NA | NA | NA | NA | + | NA | + | NA | NA | + | 0,0570702692379036 | 7 | -526,079064396247 | 1067,02634584676 | 19,0647729314674 | 2.52066314031722e-06 |
| 3115 | 24,9288449288528 | NA | + | NA | + | NA | + | NA | NA | NA | NA | + | + | 0,0570605330908789 | 7 | -526,079771357224 | 1067,02775976871 | 19,0661868534219 | 2.51888175959773e-06 |
| 370 | 26,0285152173642 | + | NA | NA | NA | + | + | + | NA | + | NA | NA | NA | 0,0570339408306876 | 7 | -526,081702236887 | 1067,03162152804 | 19,0700486127466 | 2.51402279458418e-06 |
| 1393 | 26,0406108981478 | NA | NA | NA | NA | + | + | + | NA | + | NA | + | NA | 0,0570096839536665 | 7 | -526,083463495536 | 1067,03514404534 | 19,0735711300458 | 2.50959884719334e-06 |
| 2148 | 24,531281498871 | + | + | NA | NA | NA | + | + | NA | NA | NA | NA | + | 0,0569946513234634 | 7 | -526,084554971518 | 1067,0373269973 | 19,0757540820096 | 2.50686117465121e-06 |
| 261 | 27,0405259087394 | NA | NA | + | NA | NA | NA | NA | NA | + | NA | NA | NA | 0,0106660367468505 | 4 | -529,368333010904 | 1067,03969632484 | 19,0781234095475 | 2.50389314544919e-06 |
| 1345 | 26,3999492811111 | NA | NA | NA | NA | NA | NA | + | NA | + | NA | + | NA | 0,0260925870913637 | 5 | -528,292284511736 | 1067,04258429065 | 19,0810113753571 | 2.50028017571279e-06 |
| 1079 | 25,3572430570495 | NA | + | + | NA | + | + | NA | NA | NA | NA | + | NA | 0,0568764857588971 | 7 | -526,093134027216 | 1067,05448510869 | 19,0929121934043 | 2.48544666249702e-06 |
| 467 | 25,9834819440691 | NA | + | NA | NA | + | NA | + | + | + | NA | NA | NA | 0,0568592031939426 | 7 | -526,094388685845 | 1067,05699442595 | 19,0954215106633 | 2.48233023083157e-06 |
| 3123 | 25,3478642371021 | NA | + | NA | NA | + | + | NA | NA | NA | NA | + | + | 0,056828705313115 | 7 | -526,096602677846 | 1067,06142240996 | 19,0998494946657 | 2.47684045096299e-06 |
| 2375 | 24,9775768588443 | NA | + | + | NA | NA | NA | + | NA | + | NA | NA | + | 0,0568159796800315 | 7 | -526,09752647333 | 1067,06327000092 | 19,1016970856331 | 2.47455341348068e-06 |
| 2289 | 25,5367471320947 | NA | NA | NA | NA | + | + | + | + | NA | NA | NA | + | 0,0568045099773159 | 7 | -526,098359086031 | 1067,06493522632 | 19,1033623110341 | 2.4724939263771e-06 |
| 3297 | 25,7257545282676 | NA | NA | NA | NA | NA | + | + | + | NA | NA | + | + | 0,0568001812659582 | 7 | -526,09867331474 | 1067,06556368374 | 19,1039907684531 | 2.47171711985522e-06 |
| 1068 | 25,1538686435378 | + | + | NA | + | NA | + | NA | NA | NA | NA | + | NA | 0,0567732903562268 | 7 | -526,100625340502 | 1067,06946773527 | 19,1078948199763 | 2.46689697042031e-06 |
| 2379 | 24,9754312055521 | NA | + | NA | + | NA | NA | + | NA | + | NA | NA | + | 0,0567613073657325 | 7 | -526,101495174382 | 1067,07120740303 | 19,1096344877367 | 2.46475211282664e-06 |
| 1322 | 25,8975302914173 | + | NA | NA | + | NA | + | NA | NA | + | NA | + | NA | 0,0567001284292894 | 7 | -526,105935923034 | 1067,08008890033 | 19,1185159850415 | 2.45383103503098e-06 |
| 1475 | 25,9859197887902 | NA | + | NA | NA | NA | NA | + | + | + | NA | + | NA | 0,0566970784895991 | 7 | -526,10615729914 | 1067,08053165254 | 19,1189587372521 | 2.45328787559689e-06 |
| 2274 | 25,6507281725976 | + | NA | NA | NA | NA | + | + | + | NA | NA | NA | + | 0,0565013357097363 | 7 | -526,120363550863 | 1067,10894415599 | 19,1473712406998 | 2.4186822406657e-06 |
| 3081 | 25,8286658000841 | NA | NA | NA | + | NA | NA | NA | NA | NA | NA | + | + | 0,0255991981232835 | 5 | -528,326962835597 | 1067,11194093837 | 19,1503680230794 | 2.41506082233362e-06 |
| 89 | 26,1051958042474 | NA | NA | NA | + | + | NA | + | NA | NA | NA | NA | NA | 0,025590369887616 | 5 | -528,327583176886 | 1067,11318162095 | 19,1516087056577 | 2.41356312497947e-06 |
| 3491 | 25,9547212776322 | NA | + | NA | NA | NA | + | NA | + | + | NA | + | + | 0,0718854190394595 | 8 | -524,994764694049 | 1067,1145293881 | 19,1529564728064 | 2.41193721233337e-06 |
| 1378 | 25,9025432845328 | + | NA | NA | NA | NA | + | + | NA | + | NA | + | NA | 0,0564442970437445 | 7 | -526,124502641716 | 1067,1172223377 | 19,1556494224048 | 2.40869178509298e-06 |
| 6 | 26,1060175120773 | + | NA | + | NA | NA | NA | NA | NA | NA | NA | NA | NA | 0,0100908406351555 | 4 | -529,408129628369 | 1067,11928955977 | 19,1577166444779 | 2.40620342090222e-06 |
| 452 | 25,8772786418789 | + | + | NA | NA | NA | NA | + | + | + | NA | NA | NA | 0,0563634180033322 | 7 | -526,130371314079 | 1067,12895968242 | 19,1673867671316 | 2.39459736041476e-06 |
| 3736 | 24,9532522593486 | + | + | + | NA | + | NA | NA | + | NA | + | + | + | 0,102553875960513 | 10 | -522,694119081792 | 1067,13426990962 | 19,172696994325 | 2.38824786544987e-06 |
| 70 | 25,5951451733669 | + | NA | + | NA | NA | NA | + | NA | NA | NA | NA | NA | 0,0254396476345974 | 5 | -528,338173241498 | 1067,13436175017 | 19,1727888348814 | 2.38813819896136e-06 |
| 77 | 25,6397052327617 | NA | NA | + | + | NA | NA | + | NA | NA | NA | NA | NA | 0,0254167705228297 | 5 | -528,339780492706 | 1067,13757625259 | 19,176003337298 | 2.38430294389036e-06 |
| 3108 | 25,2600219212352 | + | + | NA | NA | NA | + | NA | NA | NA | NA | + | + | 0,0562930033660167 | 7 | -526,135480268424 | 1067,13917759111 | 19,1776046758218 | 2.38239466986217e-06 |
| 2385 | 25,9488003008237 | NA | NA | NA | NA | + | NA | + | NA | + | NA | NA | + | 0,0407506062162416 | 6 | -527,253940385491 | 1067,15403461714 | 19,1924617018444 | 2.36476259120415e-06 |
| 1253 | 25,6874668348262 | NA | NA | + | NA | NA | + | + | + | NA | NA | + | NA | 0,0561767324346679 | 7 | -526,143915505451 | 1067,15604806517 | 19,1944751498745 | 2.36238312584714e-06 |
| 1076 | 25,5841145838318 | + | + | NA | NA | + | + | NA | NA | NA | NA | + | NA | 0,0561469647653971 | 7 | -526,146074926777 | 1067,16036690782 | 19,1987939925264 | 2.35728724939752e-06 |
| 2471 | 25,7283343330307 | NA | + | + | NA | NA | + | NA | + | + | NA | NA | + | 0,0715359773685962 | 8 | -525,020538471723 | 1067,16607694345 | 19,2045040281553 | 2.3505667504438e-06 |
| 3393 | 25,904253610207 | NA | NA | NA | NA | NA | NA | + | NA | + | NA | + | + | 0,0406646897710066 | 6 | -527,260072618233 | 1067,16629908262 | 19,2047261673288 | 2.35030568846448e-06 |
| 2370 | 25,8456820249249 | + | NA | NA | NA | NA | NA | + | NA | + | NA | NA | + | 0,0406601113568595 | 6 | -527,260399384284 | 1067,16695261472 | 19,2053796994305 | 2.34953781382083e-06 |
| 3299 | 25,0857184691583 | NA | + | NA | NA | NA | + | + | + | NA | NA | + | + | 0,0715131882681461 | 8 | -525,022218991156 | 1067,16943798231 | 19,2078650670214 | 2.34661989465372e-06 |
| 4054 | 25,422079470706 | + | NA | + | NA | + | NA | + | + | + | + | + | + | 0,117613667828364 | 11 | -521,535453423347 | 1067,1829068467 | 19,2213339314044 | 2.33086983547223e-06 |
| 3077 | 25,8384426956946 | NA | NA | + | NA | NA | NA | NA | NA | NA | NA | + | + | 0,0250916128143193 | 5 | -528,362620651562 | 1067,1832565703 | 19,2216836550094 | 2.33046229100437e-06 |
| 455 | 25,9262334016581 | NA | + | + | NA | NA | NA | + | + | + | NA | NA | NA | 0,0559680641743052 | 7 | -526,159051390225 | 1067,18631983471 | 19,2247469194235 | 2.32689561201962e-06 |
| 230 | 25,5951426751121 | + | NA | + | NA | NA | + | + | + | NA | NA | NA | NA | 0,0559324790092838 | 7 | -526,161632248996 | 1067,19148155226 | 19,2299086369653 | 2.32089796593823e-06 |
| 245 | 25,6071246026595 | NA | NA | + | NA | + | + | + | + | NA | NA | NA | NA | 0,0559235588580113 | 7 | -526,162279178924 | 1067,19277541211 | 19,2312024968219 | 2.31939699314804e-06 |
| 2215 | 25,2662249741073 | NA | + | + | NA | NA | + | NA | + | NA | NA | NA | + | 0,0559015348877654 | 7 | -526,16387643122 | 1067,1959699167 | 19,2343970014135 | 2.31569528804128e-06 |
| 1093 | 25,8306377332263 | NA | NA | + | NA | NA | NA | + | NA | NA | NA | + | NA | 0,0249844971146942 | 5 | -528,370143147221 | 1067,19830156162 | 19,236728646327 | 2.31299717153948e-06 |
| 108 | 24,6024617856513 | + | + | NA | + | NA | + | + | NA | NA | NA | NA | NA | 0,0558743518975095 | 7 | -526,165847781624 | 1067,19991261751 | 19,2383397022218 | 2.31113473789751e-06 |
| 2323 | 26,0755834201793 | NA | + | NA | NA | + | NA | NA | NA | + | NA | NA | + | 0,0404163531289598 | 6 | -527,277794405339 | 1067,20174265683 | 19,2401697415419 | 2.30902097139174e-06 |
| 1387 | 25,229343765684 | NA | + | NA | + | NA | + | + | NA | + | NA | + | NA | 0,0712889764182062 | 8 | -525,03875067853 | 1067,20250135706 | 19,2409284417693 | 2.30814521014429e-06 |
| 183 | 25,4016357194305 | NA | + | + | NA | + | + | NA | + | NA | NA | NA | NA | 0,0558453541460554 | 7 | -526,167950678639 | 1067,20411841154 | 19,2425454962504 | 2.3062797660995e-06 |
| 3111 | 25,0535663138732 | NA | + | + | NA | NA | + | NA | NA | NA | NA | + | + | 0,0558446591898994 | 7 | -526,16800107559 | 1067,20421920544 | 19,2426462901522 | 2.30616353956016e-06 |
| 267 | 26,2807722996538 | NA | + | NA | + | NA | NA | NA | NA | + | NA | NA | NA | 0,0249339903800221 | 5 | -528,373689835496 | 1067,20539493817 | 19,2438220228778 | 2.30480822199992e-06 |
| 2227 | 25,4109069898574 | NA | + | NA | NA | + | + | NA | + | NA | NA | NA | + | 0,0558251907122882 | 7 | -526,169412878953 | 1067,20704281217 | 19,2454698968784 | 2.30290998734751e-06 |
| 319 | 25,4864378569737 | NA | + | + | + | + | + | NA | NA | + | NA | NA | NA | 0,0712299297011004 | 8 | -525,04310367359 | 1067,21120734718 | 19,2496344318899 | 2.29811970177094e-06 |
| 2212 | 25,3937178081139 | + | + | NA | NA | NA | + | NA | + | NA | NA | NA | + | 0,0557473217520827 | 7 | -526,175059442501 | 1067,21833593927 | 19,256763023976 | 2.28994310337663e-06 |
| 82 | 26,1254845066489 | + | NA | NA | NA | + | NA | + | NA | NA | NA | NA | NA | 0,0248196627324068 | 5 | -528,38171748318 | 1067,22145023353 | 19,259877318244 | 2.2863800998165e-06 |
| 1071 | 25,0085553101959 | NA | + | + | + | NA | + | NA | NA | NA | NA | + | NA | 0,0556874118394974 | 7 | -526,179403412932 | 1067,22702388013 | 19,2654509648369 | 2.28001723270099e-06 |
| 1265 | 25,8005689326183 | NA | NA | NA | NA | + | + | + | + | NA | NA | + | NA | 0,0556687767125801 | 7 | -526,180754559541 | 1067,22972617335 | 19,2681532580555 | 2.27693867540737e-06 |
| 60 | 25,2042958526921 | + | + | NA | + | + | + | NA | NA | NA | NA | NA | NA | 0,0556536486239382 | 7 | -526,181851407474 | 1067,23191986921 | 19,2703469539213 | 2.27444258909171e-06 |
| 3155 | 24,9978244787045 | NA | + | NA | NA | + | NA | + | NA | NA | NA | + | + | 0,0556414481890952 | 7 | -526,18273597581 | 1067,23368900588 | 19,2721160905933 | 2.2724315787645e-06 |
| 1259 | 25,1226121760702 | NA | + | NA | + | NA | + | + | + | NA | NA | + | NA | 0,0710719240961864 | 8 | -525,054750676653 | 1067,23450135331 | 19,2729284380148 | 2.2715087642219e-06 |
| 1380 | 25,3667176588862 | + | + | NA | NA | NA | + | + | NA | + | NA | + | NA | 0,0710289483589895 | 8 | -525,057918187202 | 1067,2408363744 | 19,2792634591135 | 2.26432511938985e-06 |
| 1447 | 25,9395190891838 | NA | + | + | NA | NA | + | NA | + | + | NA | + | NA | 0,0710240968137302 | 8 | -525,05827575932 | 1067,24155151864 | 19,2799786033502 | 2.26351560459891e-06 |
| 1383 | 25,2577284316618 | NA | + | + | NA | NA | + | + | NA | + | NA | + | NA | 0,0710098490225247 | 8 | -525,059325849604 | 1067,24365169921 | 19,2820787839171 | 2.26113995639691e-06 |
| 104 | 24,6730278083876 | + | + | + | NA | NA | + | + | NA | NA | NA | NA | NA | 0,0555447430259575 | 7 | -526,189746987833 | 1067,24771102993 | 19,2861381146395 | 2.25655525323111e-06 |
| 1250 | 25,8358392086654 | + | NA | NA | NA | NA | + | + | + | NA | NA | + | NA | 0,0554085825599563 | 7 | -526,199617247192 | 1067,26745154865 | 19,3058786333575 | 2.23439202595645e-06 |
| 242 | 25,7076382938366 | + | NA | NA | NA | + | + | + | + | NA | NA | NA | NA | 0,0554017873383536 | 7 | -526,200109794909 | 1067,26843664408 | 19,3068637287902 | 2.2332917522575e-06 |
| 439 | 25,8866505358588 | NA | + | + | NA | + | + | NA | + | + | NA | NA | NA | 0,0707679962437974 | 8 | -525,077148420508 | 1067,27929684102 | 19,3177239257252 | 2.22119762400694e-06 |
| 2069 | 25,7964052891951 | NA | NA | + | NA | + | NA | NA | NA | NA | NA | NA | + | 0,0243553070763493 | 5 | -528,414313084349 | 1067,28664143587 | 19,3250685205833 | 2.21305568464444e-06 |

| | | | | | | | | | | | | | | | | | | |
|---|---|---|---|---|---|---|---|---|---|---|---|---|---|---|---|---|---|---|
| **10** | 26,31370204285 | + | NA | NA | + | NA | NA | NA | NA | NA | NA | NA | NA | 0,0088498457217521 | 4 | -529,493912801356 | 1067,29085590574 | 19,3292829904508 | 2.2083971664154e-06 |
| **258** | 27,1101316681189 | + | NA | NA | NA | NA | NA | NA | NA | + | NA | NA | NA | 0,0088418383465268 | 4 | -529,494465958625 | 1067,29196222028 | 19,3303893049895 | 2.20717591327288e-06 |
| **2073** | 25,7379277524806 | NA | NA | NA | + | + | NA | NA | NA | NA | NA | NA | + | 0,0243112049240325 | 5 | -528,417408043377 | 1067,29283135393 | 19,3312584386399 | 2.2062169562241e-06 |
| **2308** | 25,8673328318615 | + | + | NA | NA | NA | NA | NA | NA | + | NA | NA | + | 0,0397543178742978 | 6 | -527,32501614116 | 1067,29618612847 | 19,334613213184 | 2.20251937798471e-06 |
| **180** | 25,5656691289656 | + | + | NA | NA | + | + | NA | + | NA | NA | NA | NA | 0,0551862787956269 | 7 | -526,215728970242 | 1067,29967499475 | 19,3381020794573 | 2.19868057943481e-06 |
| **2354** | 26,0855747984608 | + | NA | NA | NA | + | + | NA | NA | + | NA | NA | + | 0,0551530868909126 | 7 | -526,218134266899 | 1067,30448558806 | 19,3429126727719 | 2.1933984554707e-06 |
| **3362** | 26,0547246817122 | + | NA | NA | NA | NA | + | NA | NA | + | NA | + | + | 0,0551420019452945 | 7 | -526,218937533794 | 1067,30609212185 | 19,3445192065601 | 2.19163727854815e-06 |
| **1105** | 26,2443190989413 | NA | NA | NA | NA | + | NA | + | NA | NA | NA | + | NA | 0,0242014307134469 | 5 | -528,425111066682 | 1067,30823740054 | 19,3466644852494 | 2.18928770252306e-06 |
| **3377** | 26,1325815626696 | NA | NA | NA | NA | + | + | NA | NA | + | NA | + | + | 0,0551193216765481 | 7 | -526,220581022566 | 1067,3093790994 | 19,3478061841047 | 2.18803830553389e-06 |
| **74** | 25,7799735686385 | + | NA | NA | + | NA | NA | + | NA | NA | NA | NA | NA | 0,0241678889517195 | 5 | -528,427464570349 | 1067,31294440787 | 19,3513714925832 | 2.18414126434337e-06 |
| **63** | 24,9839578933421 | NA | + | + | + | + | + | NA | NA | NA | NA | NA | NA | 0,0550743492261852 | 7 | -526,223839761399 | 1067,31589657706 | 19,3543236617716 | 2.18091966533285e-06 |
| **168** | 25,3986366044253 | + | + | + | NA | NA | + | NA | + | NA | NA | NA | NA | 0,0550478364111609 | 7 | -526,225760828022 | 1067,31973871031 | 19,3581657950178 | 2.17673399511897e-06 |
| **1339** | 25,6638364161766 | NA | + | NA | + | + | + | NA | NA | + | NA | + | NA | 0,070469952982673 | 8 | -525,099105393782 | 1067,32321078756 | 19,3616378722741 | 2.17295837907623e-06 |
| **2404** | 25,0852716142653 | + | + | NA | NA | NA | + | + | NA | + | NA | NA | + | 0,0704495339403375 | 8 | -525,100609415702 | 1067,3262188314 | 19,3646459161143 | 2.16969265851581e-06 |
| **1097** | 25,9266265957376 | NA | NA | NA | + | NA | NA | + | NA | NA | NA | + | NA | 0,0240711645068536 | 5 | -528,434250920781 | 1067,32651710874 | 19,364944193446 | 2.16936909757574e-06 |
| **2058** | 25,6367098525475 | + | NA | NA | + | NA | NA | NA | NA | NA | NA | NA | + | 0,0240596852129556 | 5 | -528,435056282801 | 1067,32812783278 | 19,3665549174873 | 2.16762267344206e-06 |
| **2155** | 24,3860466940842 | NA | + | NA | + | NA | + | + | NA | NA | NA | NA | + | 0,0549852586668435 | 7 | -526,230294875111 | 1067,32880680449 | 19,3672338891959 | 2.16688692110282e-06 |
| **2151** | 24,4004198353077 | NA | + | + | NA | NA | + | + | NA | NA | NA | NA | + | 0,0549632764681124 | 7 | -526,231887515884 | 1067,33199208603 | 19,3704191707418 | 2.16343859534285e-06 |
| **4063** | 25,0152284230997 | NA | + | + | + | + | NA | + | + | + | + | + | + | 0,132010743634019 | 12 | -520,40914538605 | 1067,33441980436 | 19,3728468890681 | 2.16081407879895e-06 |
| **2483** | 25,912495718381 | NA | + | NA | NA | + | + | NA | + | + | NA | NA | + | 0,0703636666223197 | 8 | -525,106933852808 | 1067,33886770562 | 19,3772947903253 | 2.15601387465709e-06 |
| **139** | 26,2009147973208 | NA | + | NA | + | NA | NA | NA | + | NA | NA | NA | NA | 0,0239604449121912 | 5 | -528,442018369364 | 1067,3420520059 | 19,3804790906129 | 2.15258390809201e-06 |
| **2054** | 25,5364871287736 | + | NA | + | NA | NA | NA | NA | NA | NA | NA | NA | + | 0,0239550831862881 | 5 | -528,442394494791 | 1067,34280425676 | 19,3812313414662 | 2.15177441879543e-06 |
| **2468** | 25,8856198258273 | + | + | NA | NA | NA | + | NA | + | + | NA | NA | + | 0,0703272822308269 | 8 | -525,109613518108 | 1067,34422703622 | 19,382654120926 | 2.1502442129238e-06 |
| **3040** | 25,01348116156 | + | + | + | + | + | NA | + | + | + | + | NA | + | 0,13188747425882 | 12 | -520,418867989903 | 1067,35386501206 | 19,3922920967736 | 2.13990713914976e-06 |
| **424** | 25,8228037224304 | + | + | + | NA | NA | + | NA | + | + | NA | NA | NA | 0,070187167488684 | 8 | -525,119931816051 | 1067,3648636321 | 19,4032907168107 | 2.12817142512191e-06 |
| **2187** | 25,4540329367365 | NA | + | NA | + | NA | NA | NA | + | NA | NA | NA | + | 0,0392641333835413 | 6 | -527,359959120823 | 1067,3660720878 | 19,4044991725091 | 2.12688591308922e-06 |
| **2407** | 24,9708622054341 | NA | + | + | NA | NA | + | + | NA | + | NA | NA | + | 0,0701772881328161 | 8 | -525,12065929067 | 1067,36631858134 | 19,4047456660519 | 2.12662379742006e-06 |
| **25** | 26,6889242878561 | NA | NA | NA | + | + | NA | NA | NA | NA | NA | NA | NA | 0,0083030631878422 | 4 | -529,531674812715 | 1067,36637992846 | 19,4048070131698 | 2.12655856730003e-06 |
| **1252** | 25,2994226334721 | + | + | NA | NA | NA | + | + | + | NA | NA | + | NA | 0,0701557563232112 | 8 | -525,122244776687 | 1067,36948955337 | 19,4079166380834 | 2.12325473663361e-06 |
| **2193** | 26,279509455231 | NA | NA | NA | NA | + | NA | NA | + | NA | NA | NA | + | 0,0237419439752186 | 5 | -528,457344551957 | 1067,37270437109 | 19,4111314557986 | 2.11984453968769e-06 |
| **2411** | 24,9642900707539 | NA | + | NA | + | NA | + | + | NA | + | NA | NA | + | 0,0701211214542628 | 8 | -525,124795024144 | 1067,37459004829 | 19,4130171329975 | 2.11784681034693e-06 |
| **1064** | 25,2750288543005 | + | + | + | NA | NA | + | NA | NA | NA | NA | + | NA | 0,0545981847165397 | 7 | -526,258333506234 | 1067,38488406673 | 19,4233111514411 | 2.10697423790179e-06 |
| **1090** | 25,9462278808414 | + | NA | NA | NA | NA | NA | + | NA | NA | NA | + | NA | 0,0236436311734421 | 5 | -528,464239330546 | 1067,38649392827 | 19,4249210129767 | 2.10527895189621e-06 |
| **1267** | 25,2664903176955 | NA | + | NA | NA | + | + | + | + | NA | NA | + | NA | 0,0699756864769831 | 8 | -525,135502709806 | 1067,39600541961 | 19,4344325043207 | 2.09529055060618e-06 |
| **1444** | 26,0752987027884 | + | + | NA | NA | NA | + | NA | + | + | NA | + | NA | 0,0699709924743045 | 8 | -525,135848279016 | 1067,39669655803 | 19,435123642741 | 2.09456660779897e-06 |
| **1459** | 26,1194869277895 | NA | + | NA | NA | + | + | NA | + | + | NA | + | NA | 0,0699435989708692 | 8 | -525,1378649349 | 1067,4007298698 | 19,4391569545096 | 2.09034684406012e-06 |
| **2178** | 26,340644341739 | + | NA | NA | NA | NA | NA | NA | + | NA | NA | NA | + | 0,0234930655393119 | 5 | -528,474797309277 | 1067,40760988573 | 19,4460369704382 | 2.08316838831983e-06 |
| **3371** | 25,3411152415539 | NA | + | NA | + | NA | + | NA | NA | + | NA | + | + | 0,0698759479726391 | 8 | -525,142845014085 | 1067,41069002817 | 19,4491171128786 | 2.07996262981991e-06 |
| **2363** | 25,4873897072507 | NA | + | NA | + | + | + | NA | NA | + | NA | NA | + | 0,0698757103021954 | 8 | -525,142862509384 | 1067,41072501877 | 19,449152103477 | 2.07992624056975e-06 |
| **3936** | 24,975842439261 | + | + | + | + | + | NA | + | NA | + | + | + | + | 0,131518709297361 | 12 | -520,447945287789 | 1067,41201960784 | 19,4504466925459 | 2.07858035132249e-06 |
| **1255** | 25,2894791801054 | NA | + | + | NA | NA | + | + | + | NA | NA | + | NA | 0,0698477869548556 | 8 | -525,144917960269 | 1067,41483592054 | 19,4532630052474 | 2.07565544504622e-06 |
| **3140** | 24,8821732270812 | + | + | NA | NA | NA | NA | + | NA | NA | NA | + | + | 0,0543673426918206 | 7 | -526,275049641051 | 1067,41831633637 | 19,4567434210744 | 2.07204651407096e-06 |
| **2359** | 25,5685281098648 | NA | + | + | NA | + | + | NA | NA | + | NA | NA | + | 0,0698081079959025 | 8 | -525,147838641232 | 1067,42067728246 | 19,4591043671728 | 2.0696019621521e-06 |
| **1335** | 25,7561791783442 | NA | + | + | NA | + | + | NA | NA | + | NA | + | NA | 0,0697914164422473 | 8 | -525,149067232601 | 1067,4231344652 | 19,4615615499122 | 2.06706082836989e-06 |
| **2107** | 25,004552261507 | NA | + | NA | + | + | + | NA | NA | NA | NA | NA | + | 0,0543122268567593 | 7 | -526,279040181729 | 1067,42629741772 | 19,464724502431 | 2.06379440430995e-06 |
| **3523** | 25,5314209814782 | NA | + | NA | NA | NA | NA | + | + | + | NA | + | + | 0,069715167748715 | 8 | -525,154679281146 | 1067,43435856229 | 19,4727856470006 | 2.05549287298197e-06 |
| **316** | 25,6623978838135 | + | + | NA | + | + | + | NA | NA | + | NA | NA | NA | 0,0697013231591745 | 8 | -525,15569821989 | 1067,43639643978 | 19,4748235244899 | 2.0533995183355e-06 |
| **1696** | 25,1101760041644 | + | + | + | + | + | NA | NA | + | NA | + | + | NA | 0,10056663653832 | 10 | -522,845559349817 | 1067,43715044567 | 19,4755775303754 | 2.05262552658222e-06 |
| **3083** | 25,3063047354708 | NA | + | NA | + | NA | NA | NA | NA | NA | NA | + | + | 0,0387542050700872 | 6 | -527,396290639578 | 1067,43873512531 | 19,4771622100184 | 2.05099979379038e-06 |
| **1033** | 26,5930796010538 | NA | NA | NA | + | NA | NA | NA | NA | NA | NA | + | NA | 0,0077680940247264 | 4 | -529,5686008302 | 1067,44023196343 | 19,4786590481399 | 2.0494653607222e-06 |
| **3241** | 25,7021029622008 | NA | NA | NA | + | NA | + | NA | + | NA | NA | + | + | 0,0542048686896364 | 7 | -526,286812546056 | 1067,44184214638 | 19,4802692310855 | 2.04781601766162e-06 |
| **111** | 24,4946037387582 | NA | + | + | + | NA | + | + | NA | NA | NA | NA | NA | 0,0541641176097825 | 7 | -526,289762554051 | 1067,44774216236 | 19,486169247074 | 2.04178384589077e-06 |
| **133** | 26,8992413793104 | NA | NA | + | NA | NA | NA | NA | + | NA | NA | NA | NA | 0,0076777774568138 | 4 | -529,57483292915 | 1067,45269616133 | 19,4911232460406 | 2.03673260690035e-06 |
| **21** | 26,8153419035549 | NA | NA | + | NA | + | NA | NA | NA | NA | NA | NA | NA | 0,0075927243560507 | 4 | -529,580701315616 | 1067,46443293426 | 19,5028600189714 | 2.02481527479228e-06 |
| **2507** | 25,3607929155468 | NA | + | NA | + | NA | NA | + | + | + | NA | NA | + | 0,0694079602130585 | 8 | -525,177285681862 | 1067,47957136372 | 19,5179984484325 | 2.0095468709988e-06 |
| **1197** | 25,6478897320925 | NA | NA | + | + | NA | + | NA | + | NA | NA | + | NA | 0,0539212363961707 | 7 | -526,307342312945 | 1067,48290168015 | 19,5213287648639 | 2.00620344195843e-06 |
| **1330** | 26,2210243671706 | + | NA | NA | NA | + | + | NA | NA | + | NA | + | NA | 0,0539150691937691 | 7 | -526,307788636749 | 1067,48379432776 | 19,5222214124719 | 2.00530822539971e-06 |
| **56** | 25,2643808554699 | + | + | + | NA | + | + | NA | NA | NA | NA | NA | NA | 0,0539021692557154 | 7 | -526,308722202904 | 1067,48566146007 | 19,5240885447818 | 2.00343701109738e-06 |
| **3201** | 26,5719495091164 | NA | NA | NA | NA | NA | NA | NA | + | NA | NA | + | + | 0,0229309933938309 | 5 | -528,514196619355 | 1067,48640850589 | 19,5248355905944 | 2.00268882122408e-06 |
| **2221** | 25,4064384473338 | NA | NA | + | + | NA | + | NA | + | NA | NA | NA | + | 0,0538631194202323 | 7 | -526,311548154798 | 1067,49131336386 | 19,5297404485686 | 1.99778338668028e-06 |
| **364** | 25,1445636147321 | + | + | NA | + | NA | + | + | NA | + | NA | NA | NA | 0,0693198805228399 | 8 | -525,183765803112 | 1067,49253160622 | 19,5309586909343 | 1.99656686504291e-06 |
| **130** | 27,0309871881757 | + | NA | NA | NA | NA | NA | NA | + | NA | NA | NA | NA | 0,0073401725989868 | 4 | -529,598123600731 | 1067,49927750449 | 19,5377045892026 | 1.98984389106267e-06 |
| **2103** | 25,084170237971 | NA | + | + | NA | + | + | NA | NA | NA | NA | NA | + | 0,0538067885775275 | 7 | -526,315624490323 | 1067,49946603491 | 19,5378931196185 | 1.989656326855e-06 |
| **1227** | 25,5415084462908 | NA | + | NA | + | NA | NA | + | + | NA | NA | + | NA | 0,0537925986468211 | 7 | -526,316651294846 | 1067,50151964396 | 19,5399467286647 | 1.98761438725494e-06 |
| **2537** | 25,9699041300812 | NA | NA | NA | + | NA | + | + | + | + | NA | NA | + | 0,0690979053387615 | 8 | -525,200094046837 | 1067,52518809367 | 19,5636151783829 | 1.96423114583063e-06 |
| **367** | 25,0499481561536 | NA | + | + | + | NA | + | + | NA | + | NA | NA | NA | 0,06907270459838 | 8 | -525,201947538829 | 1067,52889507766 | 19,5673221623672 | 1.9605938310393e-06 |
| **436** | 26,0374440326298 | + | + | NA | NA | + | + | NA | + | + | NA | NA | NA | 0,0690242331736343 | 8 | -525,205512427918 | 1067,53602485584 | 19,5744519405459 | 1.95361697473054e-06 |
| **76** | 24,9863491381665 | + | + | NA | + | NA | NA | + | NA | NA | NA | NA | NA | 0,0380552108213549 | 6 | -527,446061491774 | 1067,5382768297 | 19,5767039144107 | 1.95141846552472e-06 |
| **312** | 25,6907664045303 | + | + | + | NA | + | + | NA | NA | + | NA | NA | NA | 0,0690082593636059 | 8 | -525,206687200299 | 1067,5383744006 | 19,5768014853079 | 1.95132326702165e-06 |
| **3805** | 24,9056760472863 | NA | NA | + | + | + | NA | + | + | NA | + | + | + | 0,0998618450008881 | 10 | -522,899188616263 | 1067,54440897856 | 19,5828360632665 | 1.94544443436156e-06 |
| **2218** | 25,6032139254546 | + | NA | NA | + | NA | + | NA | + | NA | NA | NA | + | 0,0534800102535699 | 7 | -526,339266753393 | 1067,54675056105 | 19,5851776457594 | 1.94316805788829e-06 |
| **360** | 25,1623837252233 | + | + | + | NA | NA | + | + | NA | + | NA | NA | NA | 0,0689351777352671 | 8 | -525,212061633464 | 1067,54912326693 | 19,5875503516384 | 1.94086414164999e-06 |
| **72** | 25,0189559995381 | + | + | + | NA | NA | NA | + | NA | NA | NA | NA | NA | 0,0379403624557468 | 6 | -527,454235642009 | 1067,55462513017 | 19,593052214881 | 1.93553229424627e-06 |
| **3790** | 24,9321727568616 | + | NA | + | + | NA | NA | + | + | NA | + | + | + | 0,0997849040234731 | 10 | -522,905040697254 | 1067,55611314054 | 19,5945402252491 | 1.93409278375347e-06 |
| **3367** | 25,4310328422022 | NA | + | + | NA | NA | + | NA | NA | + | NA | + | + | 0,0688831202859096 | 8 | -525,215889688424 | 1067,55677937685 | 19,5952064615574 | 1.93344860963446e-06 |
| **3203** | 25,9593159704301 | NA | + | NA | NA | NA | NA | NA | + | NA | NA | + | + | 0,037876965759829 | 6 | -527,458747383478 | 1067,56364861311 | 19,6020756978201 | 1.92681934300039e-06 |

| | | | | | | | | | | | | | | | | | | |
|---|---|---|---|---|---|---|---|---|---|---|---|---|---|---|---|---|---|---|
| 2233 | 25,4950752209246 | NA | NA | NA | + | + | + | NA | + | NA | NA | NA | + | 0,0533485486184144 | 7 | -526,348775641696 | 1067,56576833766 | 19,6041954223651 | 1.92477826169634e-06 |
| 1209 | 25,7584884897816 | NA | NA | NA | + | + | + | NA | + | NA | NA | + | NA | 0,0533420180167083 | 7 | -526,349247979029 | 1067,56671301232 | 19,6051400970312 | 1.92386933174325e-06 |
| 79 | 24,9383593679061 | NA | + | + | + | NA | NA | + | NA | NA | NA | NA | NA | 0,0378524870467482 | 6 | -527,460489376213 | 1067,56713259858 | 19,6055596832882 | 1.92346575951208e-06 |
| 493 | 25,9399380776453 | NA | NA | + | + | NA | + | + | + | + | NA | NA | NA | 0,0687870783905547 | 8 | -525,222951587059 | 1067,57090317412 | 19,6093302588276 | 1.91984288918984e-06 |
| 3379 | 25,7690358611475 | NA | + | NA | NA | + | + | NA | NA | + | NA | + | + | 0,0687549078319391 | 8 | -525,225316904723 | 1067,57563380945 | 19,614060894156 | 1.91530721715887e-06 |
| 1327 | 25,3792214983104 | NA | + | + | + | NA | + | NA | NA | + | NA | + | NA | 0,0687138503360041 | 8 | -525,228335509565 | 1067,58167101913 | 19,6200981038394 | 1.9095343788621e-06 |
| 1194 | 25,8361942711672 | + | NA | NA | + | NA | + | NA | + | NA | NA | + | NA | 0,0531528284998729 | 7 | -526,362930032177 | 1067,59407711862 | 19,6325042033268 | 1.89772610364502e-06 |
| 2782 | 24,8743151716262 | + | NA | + | + | + | NA | + | + | NA | + | NA | + | 0,0995294614532965 | 10 | -522,924465909516 | 1067,59496356506 | 19,6333906497725 | 1.89688517373883e-06 |
| 174 | 25,5477654572866 | + | NA | + | + | NA | + | NA | + | NA | NA | NA | NA | 0,0531185683028272 | 7 | -526,365407413369 | 1067,599031881 | 19,6374589657119 | 1.89303053144995e-06 |
| 2100 | 25,3165554304723 | + | + | NA | NA | + | + | NA | NA | NA | NA | NA | + | 0,0531041338144254 | 7 | -526,366451155499 | 1067,60111936526 | 19,6395464499708 | 1.89105572650474e-06 |
| 1235 | 25,6964772723636 | NA | + | NA | NA | + | NA | + | + | NA | NA | + | NA | 0,0530808097101184 | 7 | -526,368137662458 | 1067,60449237918 | 19,6429194638899 | 1.88786913572012e-06 |
| 189 | 25,522230745418 | NA | NA | + | + | + | + | NA | + | NA | NA | NA | NA | 0,0530542416172954 | 7 | -526,370058683433 | 1067,60833442113 | 19,6467615058389 | 1.88424598070597e-06 |
| 2441 | 26,485295475932 | NA | NA | NA | + | NA | NA | NA | + | + | NA | NA | + | 0,0375595412119789 | 6 | -527,481333012478 | 1067,60881987111 | 19,6472469558198 | 1.88378868261934e-06 |
| 2132 | 24,9780907136753 | + | + | NA | NA | + | NA | + | NA | NA | NA | NA | + | 0,0530403844284293 | 7 | -526,371060613933 | 1067,61033828213 | 19,6487653668382 | 1.88235904263725e-06 |
| 4048 | 24,9999626248149 | + | + | + | + | NA | NA | + | + | + | + | + | + | 0,130253885984016 | 12 | -520,547583560591 | 1067,61129615344 | 19,6497232381496 | 1.88145772962764e-06 |
| 457 | 26,6781042356887 | NA | NA | NA | + | NA | NA | + | + | + | NA | NA | NA | 0,037523234508002 | 6 | -527,483915859722 | 1067,6139855656 | 19,6524126503073 | 1.87892942227764e-06 |
| 236 | 25,0052567170698 | + | + | NA | + | NA | + | + | + | NA | NA | NA | NA | 0,0684567914765639 | 8 | -525,247231815415 | 1067,61946363083 | 19,6578907155388 | 1.87379001502249e-06 |
| 2283 | 24,8172009208305 | NA | + | NA | + | NA | + | + | + | NA | NA | NA | + | 0,0684486499434034 | 8 | -525,247830211484 | 1067,62066042297 | 19,6590875076772 | 1.8726690818574e-06 |
| 263 | 26,5111380828723 | NA | + | + | NA | NA | NA | NA | NA | + | NA | NA | NA | 0,0219431364124087 | 5 | -528,58338710536 | 1067,6247894779 | 19,6632165626045 | 1.86880689327662e-06 |
| 1324 | 25,5242436991812 | + | + | NA | + | NA | + | NA | NA | + | NA | + | NA | 0,0684046383774088 | 8 | -525,251064935303 | 1067,62712987061 | 19,6655569553147 | 1.86662130129695e-06 |
| 27 | 26,0983660991473 | NA | + | NA | + | + | NA | NA | NA | NA | NA | NA | NA | 0,0219251541335251 | 5 | -528,584645954618 | 1067,62730717641 | 19,6657342611204 | 1.86645582723509e-06 |
| 3147 | 24,7363190797516 | NA | + | NA | + | NA | NA | + | NA | NA | NA | + | + | 0,0529175362743087 | 7 | -526,379942388874 | 1067,62810183201 | 19,6665289167215 | 1.86571437975502e-06 |
| 1029 | 26,651438395054 | NA | NA | + | NA | NA | NA | NA | NA | NA | NA | + | NA | 0,0063863512798216 | 4 | -529,663883031221 | 1067,63079636547 | 19,6692234501807 | 1.86320245733901e-06 |
| 3143 | 24,7813519416273 | NA | + | + | NA | NA | NA | + | NA | NA | NA | + | + | 0,0528395012740067 | 7 | -526,38558362823 | 1067,63938431072 | 19,6778113954335 | 1.85521906948074e-06 |
| 2133 | 25,3614224296841 | NA | NA | + | NA | + | NA | + | NA | NA | NA | NA | + | 0,0372790041326998 | 6 | -527,501287799524 | 1067,6487294452 | 19,6871565299109 | 1.84657065452129e-06 |
| 2276 | 24,9777353747015 | + | + | NA | NA | NA | + | + | + | NA | NA | NA | + | 0,0682527352119129 | 8 | -525,262228208837 | 1067,64945641767 | 19,6878835023833 | 1.84589957347593e-06 |
| 251 | 25,0007663587892 | NA | + | NA | + | + | + | + | + | NA | NA | NA | NA | 0,068186387572809 | 8 | -525,267103486706 | 1067,65920697341 | 19,6976340581207 | 1.83692220151042e-06 |
| 219 | 25,4530894822651 | NA | + | NA | + | + | NA | + | + | NA | NA | NA | NA | 0,052680442178102 | 7 | -526,397080753695 | 1067,66237856165 | 19,7008056463635 | 1.83401152955498e-06 |
| 260 | 26,505062046424 | + | + | NA | NA | NA | NA | NA | NA | + | NA | NA | NA | 0,0216077200906992 | 5 | -528,606864113853 | 1067,67174349488 | 19,7101705795903 | 1.82544390625834e-06 |
| 1223 | 25,7115281918407 | NA | + | + | NA | NA | NA | + | + | NA | NA | + | NA | 0,0525765778816483 | 7 | -526,40458724184 | 1067,67739153794 | 19,7158186226525 | 1.82029608556275e-06 |
| 186 | 25,6757013754044 | + | NA | NA | + | + | + | NA | + | NA | NA | NA | NA | 0,0525004165092122 | 7 | -526,410091059505 | 1067,68839917327 | 19,7268262579835 | 1.81030502751224e-06 |
| 2014 | 25,5930538018002 | + | NA | + | + | + | NA | + | + | + | + | + | NA | 0,114344951264663 | 11 | -521,788611535182 | 1067,68922307036 | 19,7276501550732 | 1.80955942857514e-06 |
| 1220 | 25,6368688039239 | + | + | NA | NA | NA | NA | + | + | NA | NA | + | NA | 0,0524573066564795 | 7 | -526,413206206132 | 1067,69462946653 | 19,7330565512375 | 1.80467443652353e-06 |
| 2515 | 25,4879867228916 | NA | + | NA | NA | + | NA | + | + | + | NA | NA | + | 0,0679426345531625 | 8 | -525,285011676106 | 1067,69502335221 | 19,7334504369212 | 1.80431905380757e-06 |
| 505 | 26,0329195468922 | NA | NA | NA | + | + | + | + | + | + | NA | NA | NA | 0,0679294768455599 | 8 | -525,285978221094 | 1067,69695644219 | 19,7353835268973 | 1.80257594080355e-06 |
| 2500 | 25,4749041998146 | + | + | NA | NA | NA | NA | + | + | + | NA | NA | + | 0,0679278717478681 | 8 | -525,286096128171 | 1067,69719225634 | 19,7356193410521 | 1.80236341687181e-06 |
| 4000 | 25,1632180015458 | + | + | + | + | + | NA | NA | + | + | + | + | + | 0,129676218276812 | 12 | -520,5930419964 | 1067,70221302506 | 19,740640109768 | 1.79784446646718e-06 |
| 1035 | 26,0518515742932 | NA | + | NA | + | NA | NA | NA | NA | NA | NA | + | NA | 0,021379788169987 | 5 | -528,622813307609 | 1067,70364188239 | 19,7420689671028 | 1.79656049354899e-06 |
| 2291 | 24,9824446893855 | NA | + | NA | NA | + | + | + | + | NA | NA | NA | + | 0,067864956668175 | 8 | -525,290717576817 | 1067,70643515363 | 19,7448622383431 | 1.79405310453192e-06 |
| 2503 | 25,459372543043 | NA | + | + | NA | NA | NA | + | + | + | NA | NA | + | 0,0678610997293924 | 8 | -525,291000879428 | 1067,70700175886 | 19,7454288435663 | 1.79354491659096e-06 |
| 3141 | 25,3170637706385 | NA | NA | + | NA | NA | NA | + | NA | NA | NA | + | + | 0,0368534508874922 | 6 | -527,531546589176 | 1067,70924702451 | 19,747674109216 | 1.79153255397829e-06 |
| 3364 | 25,6354815325563 | + | + | NA | NA | NA | + | NA | NA | + | NA | + | + | 0,0678069651882502 | 8 | -525,294977084632 | 1067,71495416926 | 19,7533812539737 | 1.78642757334641e-06 |
| 239 | 25,0057232538333 | NA | + | + | + | NA | + | + | + | NA | NA | NA | NA | 0,0677769671546501 | 8 | -525,29718035365 | 1067,7193607073 | 19,7577877920098 | 1.78249592565027e-06 |
| 4056 | 25,1426131611055 | + | + | + | NA | + | NA | + | + | + | + | + | + | 0,129552409158526 | 12 | -520,602780989504 | 1067,72169101127 | 19,7601180959762 | 1.78042025645736e-06 |
| 2351 | 25,1168649064336 | NA | + | + | + | NA | + | NA | NA | + | NA | NA | + | 0,0677606179353126 | 8 | -525,298381126804 | 1067,72176225361 | 19,7601893383176 | 1.78035683693304e-06 |
| 343 | 25,7155304132734 | NA | + | + | NA | + | NA | + | NA | + | NA | NA | NA | 0,0522415866583398 | 7 | -526,428792148928 | 1067,72580135212 | 19,7642284368296 | 1.77676494682785e-06 |
| 4060 | 24,9218090762658 | + | + | NA | + | + | NA | + | + | + | + | + | + | 0,129516230507418 | 12 | -520,605626589849 | 1067,72738221196 | 19,7658092966662 | 1.77536109355298e-06 |
| 244 | 25,1965861773332 | + | + | NA | NA | + | + | + | + | NA | NA | NA | NA | 0,0676993608338856 | 8 | -525,302879985205 | 1067,73075997041 | 19,7691870551203 | 1.77236525359967e-06 |
| 3074 | 26,17101888864 | + | NA | NA | NA | NA | NA | NA | NA | NA | NA | + | + | 0,0211331668398668 | 5 | -528,640066082988 | 1067,73814743315 | 19,7765745178604 | 1.76583068835718e-06 |
| 1513 | 26,0770888491448 | NA | NA | NA | + | NA | + | + | + | + | NA | + | NA | 0,0675935820126838 | 8 | -525,310647922622 | 1067,74629584524 | 19,7847229299525 | 1.75865096605576e-06 |
| 2477 | 25,885066052745 | NA | NA | + | + | NA | + | NA | + | + | NA | NA | + | 0,0675688103595886 | 8 | -525,312466917992 | 1067,74993383598 | 19,788360920694 | 1.75545489578952e-06 |
| 2279 | 24,9697841174103 | NA | + | + | NA | NA | + | + | + | NA | NA | NA | + | 0,0675675377463432 | 8 | -525,31256036534 | 1067,75012073068 | 19,7885478153901 | 1.75529086084934e-06 |
| 2348 | 25,2627514021826 | + | + | NA | + | NA | + | NA | NA | + | NA | NA | + | 0,0675596157147981 | 8 | -525,31314207325 | 1067,7512841465 | 19,7897112312091 | 1.75427009119573e-06 |
| 490 | 26,1029032054379 | + | NA | NA | + | NA | + | + | + | + | NA | NA | NA | 0,0675281193491304 | 8 | -525,315454775219 | 1067,75590955044 | 19,7943366351465 | 1.7502176751238e-06 |
| 2533 | 25,9357563723703 | NA | NA | + | NA | NA | + | + | + | + | NA | NA | + | 0,0674943124974753 | 8 | -525,317937043706 | 1067,76087408741 | 19,799301172121 | 1.74587855260456e-06 |
| 2505 | 26,1112661826892 | NA | NA | NA | + | NA | NA | + | + | + | NA | NA | + | 0,0519954372439308 | 7 | -526,446572310578 | 1067,76136167542 | 19,7997887601289 | 1.74545296976119e-06 |
| 3079 | 25,4803891456484 | NA | + | + | NA | NA | NA | NA | NA | NA | NA | + | + | 0,0364034669218984 | 6 | -527,56352798249 | 1067,77320981113 | 19,8116368958438 | 1.73514335550451e-06 |
| 207 | 25,4515701623098 | NA | + | + | + | NA | NA | + | + | NA | NA | NA | NA | 0,051911682963505 | 7 | -526,452621098703 | 1067,77345925167 | 19,8118863363782 | 1.73492696145623e-06 |
| 3089 | 26,2010662213156 | NA | NA | NA | NA | + | NA | NA | NA | NA | NA | + | + | 0,0208480058472477 | 5 | -528,660009544365 | 1067,77803435591 | 19,816461440616 | 1.73096276149511e-06 |
| 450 | 26,6989078940137 | + | NA | NA | NA | NA | NA | + | + | + | NA | NA | NA | 0,0363408370686806 | 6 | -527,567978046425 | 1067,782109939 | 19,8205370237129 | 1.72743901176216e-06 |
| 1363 | 25,8394128079567 | NA | + | NA | NA | + | NA | + | NA | + | NA | + | NA | 0,0518498331181884 | 7 | -526,45708759157 | 1067,7823922374 | 19,8208193221126 | 1.727195202335e-06 |
| 1332 | 25,9462406404043 | + | + | NA | NA | + | + | NA | NA | + | NA | + | NA | 0,0673237381851879 | 8 | -525,330460092912 | 1067,78592018582 | 19,8243472705326 | 1.72415116013772e-06 |
| 347 | 25,7025786220941 | NA | + | NA | + | + | NA | + | NA | + | NA | NA | NA | 0,0518164174623706 | 7 | -526,45950058538 | 1067,78721822502 | 19,8256453097329 | 1.72303251529168e-06 |
| 2092 | 24,7540168216386 | + | + | NA | + | NA | + | NA | NA | NA | NA | NA | + | 0,0518052745673298 | 7 | -526,460305211374 | 1067,78882747701 | 19,8272545617208 | 1.72164667615729e-06 |
| 2135 | 24,8156638834079 | NA | + | + | NA | + | NA | + | NA | NA | NA | NA | + | 0,0517426528306628 | 7 | -526,464826937369 | 1067,797870929 | 19,8362980137108 | 1.71387943552851e-06 |
| 2356 | 25,7587479428771 | + | + | NA | NA | + | + | NA | NA | + | NA | NA | + | 0,0671934730602216 | 8 | -525,340022222124 | 1067,80504444425 | 19,8434715289573 | 1.70774317661306e-06 |
| 2317 | 25,8142098522794 | NA | NA | + | + | NA | NA | NA | NA | + | NA | NA | + | 0,0361512247370865 | 6 | -527,581448884782 | 1067,80905161572 | 19,8474787004268 | 1.70432499220199e-06 |
| 204 | 25,3933691516195 | + | + | NA | + | NA | NA | + | + | NA | NA | NA | NA | 0,0516056723293908 | 7 | -526,474716844184 | 1067,81765074263 | 19,8560778273409 | 1.6970128694789e-06 |
| 2139 | 24,8214639537032 | NA | + | NA | + | + | NA | + | NA | NA | NA | NA | + | 0,0515627441711956 | 7 | -526,477815936612 | 1067,82384892749 | 19,8622760121978 | 1.69176181070513e-06 |
| 247 | 25,1210731727195 | NA | + | + | NA | + | + | + | + | NA | NA | NA | NA | 0,0670553091678611 | 8 | -525,350162702975 | 1067,82532540595 | 19,8637524906596 | 1.69051334665633e-06 |
| 3145 | 25,4077671790643 | NA | NA | NA | + | NA | NA | + | NA | NA | NA | + | + | 0,0360365508035333 | 6 | -527,58959450639 | 1067,82534285893 | 19,8637699436431 | 1.69049859446996e-06 |
| 3678 | 24,8356143358805 | + | NA | + | + | + | NA | + | NA | NA | + | + | + | 0,0979746399535305 | 10 | -523,042584149066 | 1067,83120004416 | 19,8696271288725 | 1.68555505511073e-06 |
| 215 | 25,5700651669521 | NA | + | + | NA | + | NA | + | + | NA | NA | NA | NA | 0,0514970735894307 | 7 | -526,48255659089 | 1067,83333023604 | 19,8717573207521 | 1.68376073299746e-06 |
| 212 | 25,5691134426322 | + | + | NA | NA | + | NA | + | + | NA | NA | NA | NA | 0,0514967601864189 | 7 | -526,482579214165 | 1067,83337548259 | 19,8718025673033 | 1.68372264124517e-06 |
| 232 | 25,1756853268863 | + | + | + | NA | NA | + | + | + | NA | NA | NA | NA | 0,0669679081411922 | 8 | -525,356576689538 | 1067,83815337908 | 19,8765804637862 | 1.67970511574321e-06 |
| 2125 | 25,0430735353489 | NA | NA | + | + | NA | NA | + | NA | NA | NA | NA | + | 0,0359144682690062 | 6 | -527,598265317993 | 1067,84268448214 | 19,8811115668498 | 1.67590396472294e-06 |

| | | | | | | | | | | | | | | | | | |
|---|---|---|---|---|---|---|---|---|---|---|---|---|---|---|---|---|---|
| 1320 | 25,5755394920405 | + | + | + | NA | NA | + | NA | NA | + | NA | + | NA | 0,0669217987841863 | 8 | -525,359960215548 | 1067,8449204311 | 19,8833475158058 | 1.67403139380108e-06 |
| 3802 | 24,7971575947437 | + | NA | NA | + | + | NA | + | + | NA | + | + | + | 0,0978747793949123 | 10 | -523,050163499852 | 1067,84635874574 | 19,8847858304455 | 1.67282793466066e-06 |
| 275 | 27,0299466658128 | NA | + | NA | NA | + | NA | NA | NA | + | NA | NA | NA | 0,0203163705158179 | 5 | -528,697175320987 | 1067,85236590915 | 19,8907929938591 | 1.66781099743958e-06 |
| 453 | 26,6442642431212 | NA | NA | + | NA | NA | NA | + | + | + | NA | NA | NA | 0,0358412736090537 | 6 | -527,603463382276 | 1067,85308061071 | 19,8915076954158 | 1.66721511035819e-06 |
| 387 | 27,0839311195958 | NA | + | NA | NA | NA | NA | NA | + | + | NA | NA | NA | 0,0203036278065352 | 5 | -528,698065896023 | 1067,85414705922 | 19,8925741439316 | 1.66632634779418e-06 |
| 2137 | 25,4249047186558 | NA | NA | NA | + | + | NA | + | NA | NA | NA | NA | + | 0,035804112933256 | 6 | -527,606102270582 | 1067,85835838732 | 19,8967854720279 | 1.66282131581612e-06 |
| 2118 | 25,1591183645718 | + | NA | + | NA | NA | NA | + | NA | NA | NA | NA | + | 0,0357164328653293 | 6 | -527,61232828584 | 1067,87081041783 | 19,909237502543 | 1.65250072634534e-06 |
| 3091 | 25,7625269568816 | NA | + | NA | NA | + | NA | NA | NA | NA | NA | + | + | 0,0356798998290717 | 6 | -527,614922268755 | 1067,87599838366 | 19,9144254683736 | 1.64821972251955e-06 |
| 18 | 26,7833333333334 | + | NA | NA | NA | + | NA | NA | NA | NA | NA | NA | NA | 0,0045927157144868 | 4 | -529,78737117477 | 1067,87777265257 | 19,9161997372787 | 1.64675817840722e-06 |
| 2066 | 26,0093131057733 | + | NA | NA | NA | + | NA | NA | NA | NA | NA | NA | + | 0,0201231019538668 | 5 | -528,710681421116 | 1067,87937810941 | 19,9178051941162 | 1.64543680923983e-06 |
| 304 | 25,2820573117712 | + | + | + | + | NA | + | NA | NA | + | NA | NA | NA | 0,0666779855176254 | 8 | -525,377848568392 | 1067,88069713678 | 19,9191242214938 | 1.64435197890965e-06 |
| 340 | 25,7860926991465 | + | + | NA | NA | + | NA | + | NA | + | NA | NA | NA | 0,0511503800526995 | 7 | -526,507578406302 | 1067,88337386687 | 19,9218009515771 | 1.64215270774702e-06 |
| 1026 | 26,7459729017689 | + | NA | NA | NA | NA | NA | NA | NA | NA | NA | + | NA | 0,004515690613321 | 4 | -529,792669213571 | 1067,88836873017 | 19,9267958148814 | 1.63805666052498e-06 |
| 3742 | 25,0167074412148 | + | NA | + | + | + | NA | NA | + | NA | + | + | + | 0,0975137083172439 | 10 | -523,077561561359 | 1067,90115486875 | 19,9395819534591 | 1.62761785435475e-06 |
| 3237 | 25,7287596288589 | NA | NA | + | NA | NA | + | NA | + | NA | NA | + | + | 0,051022776637058 | 7 | -526,516785591211 | 1067,90178823668 | 19,9402153213944 | 1.62710249548208e-06 |
| 205 | 26,1645033104482 | NA | NA | + | + | NA | NA | + | + | NA | NA | NA | NA | 0,03549500176204 | 6 | -527,628049220173 | 1067,9022522865 | 19,9406793712085 | 1.62672501097138e-06 |
| 3153 | 25,6205506416773 | NA | NA | NA | NA | + | NA | + | NA | NA | NA | + | + | 0,0354872562987575 | 6 | -527,628599059186 | 1067,90335196452 | 19,9417790492344 | 1.62583081995012e-06 |
| 202 | 26,209297165523 | + | NA | NA | + | NA | NA | + | + | NA | NA | NA | NA | 0,0354809493442214 | 6 | -527,629046777325 | 1067,9042474008 | 19,9426744855123 | 1.62510306892697e-06 |
| 2344 | 25,3194821609625 | + | + | + | NA | NA | + | NA | NA | + | NA | NA | + | 0,0664808664791664 | 8 | -525,392307594761 | 1067,90961518952 | 19,9480422742317 | 1.62074731177697e-06 |
| 2539 | 25,3371197077215 | NA | + | NA | + | NA | + | + | + | + | NA | NA | + | 0,0819229351889931 | 9 | -524,250256769329 | 1067,9178363733 | 19,9562634580138 | 1.61409875513295e-06 |
| 3798 | 25,0522469847292 | + | NA | + | NA | + | NA | + | + | NA | + | + | + | 0,097401307454856 | 10 | -523,086088298939 | 1067,91820834391 | 19,9566354286189 | 1.61379858440217e-06 |
| 3076 | 25,6797203667152 | + | + | NA | NA | NA | NA | NA | NA | NA | NA | + | + | 0,0353771808068981 | 6 | -527,63641267896 | 1067,91897920407 | 19,9574062887834 | 1.61317669773567e-06 |
| 3138 | 25,5606627316302 | + | NA | NA | NA | NA | NA | + | NA | NA | NA | + | + | 0,0352975413717901 | 6 | -527,642065264221 | 1067,9302843746 | 19,968711459305 | 1.60408380230544e-06 |
| 2545 | 26,0378237895528 | NA | NA | NA | NA | + | + | + | + | + | NA | NA | + | 0,066308606284792 | 8 | -525,404940682953 | 1067,93488136591 | 19,9733084506151 | 1.60040105666428e-06 |
| 486 | 26,0032008423446 | + | NA | + | NA | NA | + | + | + | + | NA | NA | NA | 0,0662788253152203 | 8 | -525,407124501039 | 1067,93924900208 | 19,9776760867871 | 1.59690988532092e-06 |
| 430 | 25,9820706490171 | + | NA | + | + | NA | + | NA | + | + | NA | NA | NA | 0,0662687973855816 | 8 | -525,407859826578 | 1067,94071965316 | 19,9791467378648 | 1.59573606832076e-06 |
| 48 | 24,8237681452066 | + | + | + | + | NA | + | NA | NA | NA | NA | NA | NA | 0,0507333883657938 | 7 | -526,537661727555 | 1067,94354050937 | 19,9819675940821 | 1.59348698377485e-06 |
| 2489 | 25,9959365643539 | NA | NA | NA | + | + | + | NA | + | + | NA | NA | + | 0,0662480172763105 | 8 | -525,409383560154 | 1067,94376712031 | 19,9821942050164 | 1.59330644321606e-06 |
| 2122 | 25,2702641041898 | + | NA | NA | + | NA | NA | + | NA | NA | NA | NA | + | 0,0351619237406958 | 6 | -527,651689952558 | 1067,94953375127 | 19,987960835979 | 1.58871905470413e-06 |
| 2501 | 26,067837175093 | NA | NA | + | NA | NA | NA | + | + | + | NA | NA | + | 0,0506720317496055 | 7 | -526,542087105778 | 1067,95239126582 | 19,9908183505293 | 1.58645078158834e-06 |
| 445 | 26,0115037814592 | NA | NA | + | + | + | + | NA | + | + | NA | NA | NA | 0,0661606855360821 | 8 | -525,415786924233 | 1067,95657384847 | 19,995000933176 | 1.58313651758752e-06 |
| 1453 | 26,0094518510628 | NA | NA | + | + | NA | + | NA | + | + | NA | + | NA | 0,0661603928557793 | 8 | -525,41580838322 | 1067,95661676644 | 19,9950438511494 | 1.58310254544658e-06 |
| 125 | 25,2050098583552 | NA | NA | + | + | + | + | + | NA | NA | NA | NA | NA | 0,050617038736325 | 7 | -526,546053262851 | 1067,96032357997 | 19,9987506646755 | 1.58017112987407e-06 |
| 1283 | 26,9107074451718 | NA | + | NA | NA | NA | NA | NA | NA | + | NA | + | NA | 0,0195420529046419 | 5 | -528,751270580343 | 1067,96055642786 | 19,9989835125716 | 1.57998717082138e-06 |
| 2130 | 25,5706593951909 | + | NA | NA | NA | + | NA | + | NA | NA | NA | NA | + | 0,0350588604111634 | 6 | -527,659003380648 | 1067,96416060745 | 20,0025876921588 | 1.57714245605238e-06 |
| 200 | 25,5344258022002 | + | + | + | NA | NA | NA | + | + | NA | NA | NA | NA | 0,0505453475225099 | 7 | -526,551223367859 | 1067,97066378998 | 20,0090908746918 | 1.57202256182897e-06 |
| 217 | 26,2601873373292 | NA | NA | NA | + | + | NA | + | + | NA | NA | NA | NA | 0,035009011773721 | 6 | -527,662540386047 | 1067,97123461825 | 20,0096617029569 | 1.57157394839645e-06 |
| 3555 | 25,5136391076201 | NA | + | NA | NA | NA | + | + | + | + | NA | + | + | 0,081550845907865 | 9 | -524,278000478795 | 1067,97332379223 | 20,011750876944 | 1.56993315981297e-06 |
| 2165 | 25,1910295320002 | NA | NA | + | NA | + | + | + | NA | NA | NA | NA | + | 0,0505223442943714 | 7 | -526,552882192849 | 1067,97398143996 | 20,0124085246709 | 1.56941701319128e-06 |
| 1515 | 25,5459890839353 | NA | + | NA | + | NA | + | + | + | + | NA | + | NA | 0,0815415094580334 | 9 | -524,278696478386 | 1067,97471579142 | 20,0131428761265 | 1.56884086713718e-06 |
| 1225 | 26,2674338147533 | NA | NA | NA | + | NA | NA | + | + | NA | NA | + | NA | 0,0348993259534825 | 6 | -527,670322490148 | 1067,98679882645 | 20,0252259111585 | 1.55939126128069e-06 |
| 1141 | 25,4018419390535 | NA | NA | + | NA | + | + | + | NA | NA | NA | + | NA | 0,0503817103433143 | 7 | -526,56302281395 | 1067,99426268216 | 20,0326897668733 | 1.55358257113999e-06 |
| 1509 | 26,0484444628117 | NA | NA | + | NA | NA | + | + | + | + | NA | + | NA | 0,0658593970256063 | 8 | -525,43787349727 | 1068,00074699454 | 20,0391740792497 | 1.54855377027705e-06 |
| 23 | 26,3274873774874 | NA | + | + | NA | + | NA | NA | NA | NA | NA | NA | NA | 0,0192448959221165 | 5 | -528,772019178617 | 1068,00205362441 | 20,0404807091181 | 1.54754240737773e-06 |
| 2095 | 24,6117279731474 | NA | + | + | + | NA | + | NA | NA | NA | NA | NA | + | 0,0502795692041754 | 7 | -526,570386912372 | 1068,00899087901 | 20,0474179637179 | 1.54218385831314e-06 |
| 501 | 26,0788675948853 | NA | NA | + | NA | + | + | + | + | + | NA | NA | NA | 0,0657729889990801 | 8 | -525,444206501273 | 1068,01341300255 | 20,0518400872563 | 1.53877776147579e-06 |
| 2513 | 26,1169841522239 | NA | NA | NA | NA | + | NA | + | + | + | NA | NA | + | 0,0502481467363164 | 7 | -526,572652227643 | 1068,01352150955 | 20,0519485942584 | 1.53869427965951e-06 |
| 2088 | 24,8695849423694 | + | + | + | NA | NA | + | NA | NA | NA | NA | NA | + | 0,050215977940764 | 7 | -526,574971269726 | 1068,01815959372 | 20,0565866784259 | 1.53513011718967e-06 |
| 198 | 26,0989151242642 | + | NA | + | NA | NA | NA | + | + | NA | NA | NA | NA | 0,0346288269863879 | 6 | -527,689510355533 | 1068,02517455722 | 20,0636016419301 | 1.52975510820001e-06 |
| 118 | 25,3627920337442 | + | NA | + | NA | + | + | + | NA | NA | NA | NA | NA | 0,0500814371804629 | 7 | -526,584669435684 | 1068,03755592563 | 20,0759830103407 | 1.52031413062084e-06 |
| 2437 | 26,426401960777 | NA | NA | + | NA | NA | NA | NA | + | + | NA | NA | + | 0,0345317448478392 | 6 | -527,696395574162 | 1068,03894499448 | 20,0773720791872 | 1.51925858672119e-06 |
| 2214 | 25,5817625575201 | + | NA | + | NA | NA | + | NA | + | NA | NA | NA | + | 0,0500672710318633 | 7 | -526,585690501054 | 1068,03959805637 | 20,0780251410811 | 1.51876258276094e-06 |
| 2075 | 25,3229687704056 | NA | + | NA | + | + | NA | NA | NA | NA | NA | NA | + | 0,0345091091455219 | 6 | -527,698000834441 | 1068,04215551504 | 20,0805825997454 | 1.51682173767e-06 |
| 2229 | 25,6047149552568 | NA | NA | + | NA | + | + | NA | + | NA | NA | NA | + | 0,050048693140436 | 7 | -526,587029532243 | 1068,04227611875 | 20,0807032034597 | 1.51673027326e-06 |
| 2474 | 26,1185755855454 | + | NA | NA | + | NA | + | NA | + | + | NA | NA | + | 0,0655408564737144 | 8 | -525,461217024472 | 1068,04743404894 | 20,0858611336539 | 1.51282371842308e-06 |
| 3497 | 26,1204397407047 | NA | NA | NA | + | NA | + | NA | + | + | NA | + | + | 0,0655408011459614 | 8 | -525,461221078351 | 1068,0474421567 | 20,0858692414117 | 1.51281758563134e-06 |
| 2443 | 25,9126169690183 | NA | + | NA | + | NA | NA | NA | + | + | NA | NA | + | 0,0500124420553673 | 7 | -526,589642311717 | 1068,0475016777 | 20,0859287624066 | 1.51277256409737e-06 |
| 2530 | 26,1483252952539 | + | NA | NA | NA | NA | + | + | + | + | NA | NA | + | 0,0655322937682701 | 8 | -525,461844413282 | 1068,04868882656 | 20,0871159112739 | 1.51187488742471e-06 |
| 2183 | 25,6096713253366 | NA | + | + | NA | NA | NA | NA | + | NA | NA | NA | + | 0,0344520518783348 | 6 | -527,702047008275 | 1068,0502478627 | 20,0886749474137 | 1.51069681285546e-06 |
| 1473 | 26,7691586716303 | NA | NA | NA | NA | NA | NA | + | + | + | NA | + | NA | 0,0344402481308317 | 6 | -527,70288403245 | 1068,05192191105 | 20,0903489957632 | 1.50943285215899e-06 |
| 1140 | 25,2060920108472 | + | + | NA | NA | + | + | + | NA | NA | NA | + | NA | 0,0655081488842336 | 8 | -525,463613476248 | 1068,0522269525 | 20,0906540372052 | 1.50920264992779e-06 |
| 15 | 25,6623516298922 | NA | + | + | + | NA | NA | NA | NA | NA | NA | NA | NA | 0,0188408360975206 | 5 | -528,800222049183 | 1068,05845936554 | 20,0968864502506 | 1.50450698291996e-06 |
| 1190 | 25,7872954679468 | + | NA | + | NA | NA | + | NA | + | NA | NA | + | NA | 0,0499347031639564 | 7 | -526,59524496884 | 1068,05870699194 | 20,0971340766528 | 1.50432071662573e-06 |
| 3553 | 26,1507219906479 | NA | NA | NA | NA | NA | + | + | + | + | NA | + | + | 0,0654519621969783 | 8 | -525,467730021903 | 1068,06046004381 | 20,0988871285153 | 1.5030027182229e-06 |
| 3173 | 25,1384340497453 | NA | NA | + | NA | NA | + | + | NA | NA | NA | + | + | 0,0498822287503091 | 7 | -526,599026551065 | 1068,06627015639 | 20,1046972411029 | 1.49864274676427e-06 |
| 3792 | 24,6462418502744 | + | + | + | + | NA | NA | + | + | NA | + | + | + | 0,111900941131984 | 11 | -521,977287711773 | 1068,06657542355 | 20,1050025082552 | 1.49841402101854e-06 |
| 1205 | 25,8436803702176 | NA | NA | + | NA | + | + | NA | + | NA | NA | + | NA | 0,0497593507001012 | 7 | -526,607880972895 | 1068,08397900005 | 20,1224060847633 | 1.48543170608807e-06 |
| 507 | 25,517110837797 | NA | + | NA | + | + | + | + | + | + | NA | NA | NA | 0,0807498612240834 | 9 | -524,337685371555 | 1068,09269357775 | 20,1311206624641 | 1.47897333179641e-06 |
| 385 | 27,8688930896871 | NA | NA | NA | NA | NA | NA | NA | + | + | NA | NA | NA | 0,0029807589984341 | 4 | -529,898161483373 | 1068,09935326978 | 20,1377803544851 | 1.47405676858993e-06 |
| 3249 | 25,914712522086 | NA | NA | NA | NA | + | + | NA | + | NA | NA | + | + | 0,0496517642888819 | 7 | -526,615632560795 | 1068,09948217585 | 20,1379092605621 | 1.473961764214e-06 |
| 492 | 25,5013268234768 | + | + | NA | + | NA | + | + | + | + | NA | NA | NA | 0,080629246550999 | 9 | -524,346668398122 | 1068,11065963089 | 20,1490867155987 | 1.4657471694883e-06 |
| 465 | 26,8207592544584 | NA | NA | NA | NA | + | NA | + | + | + | NA | NA | NA | 0,0339552702103135 | 6 | -527,737265816502 | 1068,12068547916 | 20,1591125638681 | 1.45841787607134e-06 |
| 3187 | 24,9279313094216 | NA | + | NA | NA | + | + | + | NA | NA | NA | + | + | 0,0649593273802263 | 8 | -525,503812567917 | 1068,13262513583 | 20,1710522205426 | 1.44973730823759e-06 |
| 3177 | 25,2015075749409 | NA | NA | NA | + | NA | + | + | NA | NA | NA | + | + | 0,0494126282018249 | 7 | -526,632859145792 | 1068,13393534585 | 20,1723624305564 | 1.44878788908684e-06 |
| 3680 | 24,5639352315502 | + | + | + | + | + | NA | + | NA | NA | + | + | + | 0,111451524060974 | 11 | -522,01192595428 | 1068,13585190856 | 20,1742789932687 | 1.44740020766455e-06 |
| 2310 | 25,9557589894886 | + | NA | + | NA | NA | NA | NA | NA | + | NA | NA | + | 0,0338329944624144 | 6 | -527,745931649414 | 1068,13801714498 | 20,1764442296903 | 1.44583407375688e-06 |

| | | | | | | | | | | | | | | | | | | | |
|---|---|---|---|---|---|---|---|---|---|---|---|---|---|---|---|---|---|---|---|
| **495** | 25,4744940470567 | NA | + | + | + | NA | + | + | + | + | NA | NA | NA | 0,0803706932467332 | 9 | -524,365920720457 | 1068,14916427556 | 20,1875913602685 | 1.43779803863587e-06 |
| **2169** | 25,2417531133955 | NA | NA | NA | + | + | + | + | NA | NA | NA | NA | + | 0,0492748787148182 | 7 | -526,642780204159 | 1068,15377746258 | 20,1922045472907 | 1.43448544488201e-06 |
| **1281** | 27,5376074473639 | NA | NA | NA | NA | NA | NA | NA | NA | + | NA | + | NA | 0,0025782417488993 | 4 | -529,925798688193 | 1068,15462767942 | 20,1930547641257 | 1.43387576264429e-06 |
| **132** | 26,4121607642692 | + | + | NA | NA | NA | NA | NA | + | NA | NA | NA | NA | 0,0181462310495569 | 5 | -528,84867749159 | 1068,15537025036 | 20,1937973350655 | 1.43334348422772e-06 |
| **1145** | 25,4615263844324 | NA | NA | NA | + | + | + | + | NA | NA | NA | + | NA | 0,0492535894790886 | 7 | -526,644313379323 | 1068,15684381291 | 20,195270897618 | 1.43228781253342e-06 |
| **182** | 25,6855198020243 | + | NA | + | NA | + | + | NA | + | NA | NA | NA | NA | 0,0492456014852376 | 7 | -526,644888637423 | 1068,15799432911 | 20,1964214138184 | 1.43146411430972e-06 |
| **122** | 25,4520272524414 | + | NA | NA | + | + | + | + | NA | NA | NA | NA | NA | 0,0492388650662769 | 7 | -526,645373759176 | 1068,15896457261 | 20,1973916573243 | 1.43076984834468e-06 |
| **12** | 25,7430236417414 | + | + | NA | + | NA | NA | NA | NA | NA | NA | NA | NA | 0,0180786099181137 | 5 | -528,853392891347 | 1068,16480104987 | 20,2032281345789 | 1.42660060688597e-06 |
| **210** | 26,2598465936751 | + | NA | NA | NA | + | NA | + | + | NA | NA | NA | NA | 0,0336003528316204 | 6 | -527,762416221891 | 1068,17098628994 | 20,2094133746448 | 1.42219548845986e-06 |
| **3807** | 24,5810567531441 | NA | + | + | + | + | NA | + | + | NA | + | + | + | 0,11121221058233 | 11 | -522,030363588472 | 1068,17272717694 | 20,2111542616535 | 1.42095808625747e-06 |
| **3185** | 25,4596366541844 | NA | NA | NA | NA | + | + | + | NA | NA | NA | + | + | 0,0491375185230692 | 7 | -526,652671793916 | 1068,1735606421 | 20,2119877268044 | 1.42036605010353e-06 |
| **1133** | 25,1670267194237 | NA | NA | + | + | NA | + | + | NA | NA | NA | + | NA | 0,0491179340062628 | 7 | -526,654081998889 | 1068,17638105204 | 20,2148081367518 | 1.41836445449695e-06 |
| **2532** | 25,4698455703357 | + | + | NA | NA | NA | + | + | + | + | NA | NA | + | 0,0801763265506472 | 9 | -524,380390035953 | 1068,17810290655 | 20,2165299912608 | 1.41714387137328e-06 |
| **2547** | 25,4859990477189 | NA | + | NA | NA | + | + | + | + | + | NA | NA | + | 0,0801691921480621 | 9 | -524,380921086909 | 1068,17916500846 | 20,2175920931738 | 1.41639149555751e-06 |
| **498** | 26,1666234524366 | + | NA | NA | NA | + | + | + | + | + | NA | NA | NA | 0,0645787082648406 | 8 | -525,531677626057 | 1068,18835525211 | 20,2267823368234 | 1.40989793486114e-06 |
| **2498** | 26,2540221149897 | + | NA | NA | NA | NA | NA | + | + | + | NA | NA | + | 0,0490230882062857 | 7 | -526,660911065771 | 1068,19003918581 | 20,2284662705149 | 1.40871134714792e-06 |
| **2180** | 25,7636097655543 | + | + | NA | NA | NA | NA | NA | + | NA | NA | NA | + | 0,0334641550418653 | 6 | -527,772065114468 | 1068,19028407509 | 20,2287111597984 | 1.40853886855143e-06 |
| **442** | 26,1487050907031 | + | NA | NA | + | + | + | NA | + | + | NA | NA | NA | 0,0645578913429855 | 8 | -525,533201302347 | 1068,19140260469 | 20,2298296894041 | 1.40775134257709e-06 |
| **3234** | 25,9977993502603 | + | NA | NA | NA | NA | + | NA | + | NA | NA | + | + | 0,0489788470092987 | 7 | -526,66409627808 | 1068,19640961042 | 20,2348366951333 | 1.40423144093871e-06 |
| **2535** | 25,4273050709614 | NA | + | + | NA | NA | + | + | + | + | NA | NA | + | 0,0800525221501181 | 9 | -524,389604861899 | 1068,19653255844 | 20,2349596431527 | 1.40414511985473e-06 |
| **2195** | 25,8037749136552 | NA | + | NA | NA | + | NA | NA | + | NA | NA | NA | + | 0,0334144717294657 | 6 | -527,7755845759 | 1068,19732299795 | 20,2357500826629 | 1.40359028361269e-06 |
| **1465** | 26,1767982401621 | NA | NA | NA | + | + | + | NA | + | + | NA | + | NA | 0,0644999211715393 | 8 | -525,537444199476 | 1068,19988839895 | 20,2383154836623 | 1.40179105183897e-06 |
| **3170** | 25,4025668232634 | + | NA | NA | NA | NA | + | + | NA | NA | NA | + | + | 0,0489284279759759 | 7 | -526,66772609255 | 1068,20366923936 | 20,2420963240727 | 1.3991435809158e-06 |
| **1218** | 26,2936891442142 | + | NA | NA | NA | NA | NA | + | + | NA | NA | + | NA | 0,0333534287579165 | 6 | -527,779908484157 | 1068,20597081447 | 20,2443978991773 | 1.39753438999501e-06 |
| **273** | 27,6325986697259 | NA | NA | NA | NA | + | NA | NA | NA | + | NA | NA | NA | 0,0021651637361463 | 4 | -529,954149416236 | 1068,2113291355 | 20,2497562202122 | 1.39379518224107e-06 |
| **1147** | 24,9897596967133 | NA | + | NA | + | + | + | + | NA | NA | NA | + | NA | 0,0643931983383941 | 8 | -525,545254667407 | 1068,21550933481 | 20,2539364195236 | 1.39088505369933e-06 |
| **1031** | 26,2939099546526 | NA | + | + | NA | NA | NA | NA | NA | NA | NA | + | NA | 0,0176842230768733 | 5 | -528,880888060031 | 1068,21979138724 | 20,2582184719477 | 1.38791031797137e-06 |
| **1085** | 25,4056363764375 | NA | NA | + | + | + | + | NA | NA | NA | NA | + | NA | 0,0488139312714597 | 7 | -526,675968332708 | 1068,22015371968 | 20,2585808043896 | 1.3876588982791e-06 |
| **2162** | 25,438773460377 | + | NA | NA | NA | + | + | + | NA | NA | NA | NA | + | 0,0487684148594484 | 7 | -526,679244633629 | 1068,22670632152 | 20,2651334062305 | 1.38311994969426e-06 |
| **1143** | 25,0119677062757 | NA | + | + | NA | + | + | + | NA | NA | NA | + | NA | 0,0642463625489292 | 8 | -525,55599933068 | 1068,23699866136 | 20,2754257460697 | 1.37602046266524e-06 |
| **3521** | 26,2717254723274 | NA | NA | NA | NA | NA | NA | + | + | + | NA | + | + | 0,0486830399399475 | 7 | -526,685389553235 | 1068,23899616073 | 20,2774232454424 | 1.37464684872217e-06 |
| **135** | 26,3981718963165 | NA | + | + | NA | NA | NA | NA | + | NA | NA | NA | NA | 0,0175229092058946 | 5 | -528,89213107752 | 1068,24227742222 | 20,2807045069255 | 1.37239340987933e-06 |
| **2071** | 25,4576981538556 | NA | + | + | NA | + | NA | NA | NA | NA | NA | NA | + | 0,0330886193980476 | 6 | -527,798662789193 | 1068,24347942454 | 20,2819065092485 | 1.3715688476522e-06 |
| **1221** | 26,2658026161417 | NA | NA | + | NA | NA | NA | + | + | NA | NA | + | NA | 0,0330600689511377 | 6 | -527,80068447981 | 1068,24752280577 | 20,2859498904832 | 1.36879876085465e-06 |
| **1126** | 25,3012436779395 | + | NA | + | NA | NA | + | + | NA | NA | NA | + | NA | 0,0486161121261108 | 7 | -526,690206344077 | 1068,24862974242 | 20,2870568271262 | 1.36804138371343e-06 |
| **1450** | 26,2191157466548 | + | NA | NA | + | NA | + | NA | + | + | NA | + | NA | 0,0641543088674037 | 8 | -525,562734470839 | 1068,25046894168 | 20,2888960263881 | 1.36678391163589e-06 |
| **1506** | 26,2186270527234 | + | NA | NA | NA | NA | + | + | + | + | NA | + | NA | 0,0641516012231516 | 8 | -525,562932566563 | 1068,25086513313 | 20,2892922178357 | 1.36651318440346e-06 |
| **1521** | 26,2239658467217 | NA | NA | NA | NA | + | + | + | + | + | NA | + | NA | 0,0641440581919387 | 8 | -525,563484424159 | 1068,25196884832 | 20,2903959330281 | 1.36575927176771e-06 |
| **3800** | 24,7705699988686 | + | + | + | NA | + | NA | + | + | NA | + | + | + | 0,110690578008463 | 11 | -522,070534985987 | 1068,25306997197 | 20,2914970566831 | 1.36500754380249e-06 |
| **3243** | 25,2727548865607 | NA | + | NA | + | NA | + | NA | + | NA | NA | + | + | 0,0641120025696 | 8 | -525,565829603983 | 1068,25665920797 | 20,2950862926757 | 1.36256007349327e-06 |
| **3275** | 25,0767938888457 | NA | + | NA | + | NA | NA | + | + | NA | NA | + | + | 0,0640974768794757 | 8 | -525,566892272731 | 1068,25878454546 | 20,2972116301712 | 1.3611128925594e-06 |
| **2226** | 25,7837282329607 | + | NA | NA | NA | + | + | NA | + | NA | NA | NA | + | 0,0485314588102657 | 7 | -526,696298353156 | 1068,26081376058 | 20,2992408452847 | 1.35973259748037e-06 |
| **1028** | 26,3022856520787 | + | + | NA | NA | NA | NA | NA | NA | NA | NA | + | NA | 0,0173813866163846 | 5 | -528,901993191919 | 1068,26200165101 | 20,300428735723 | 1.35892523059432e-06 |
| **1130** | 25,3840424901333 | + | NA | NA | + | NA | + | + | NA | NA | NA | + | NA | 0,0485193239330738 | 7 | -526,697171585621 | 1068,26256022551 | 20,3009873102149 | 1.35854575310334e-06 |
| **213** | 26,3234516433953 | NA | NA | + | NA | + | NA | + | + | NA | NA | NA | NA | 0,0329471186768027 | 6 | -527,808682036242 | 1068,26351791864 | 20,301945003348 | 1.35789537386228e-06 |
| **3840** | 24,4026593005916 | + | + | + | + | + | + | + | + | NA | + | + | + | 0,141554112289468 | 13 | -519,652212786767 | 1068,26377516703 | 20,3022022517384 | 1.35772072689479e-06 |
| **1523** | 25,6964667562701 | NA | + | NA | NA | + | + | + | + | + | NA | + | NA | 0,0795906780523615 | 9 | -524,42396922514 | 1068,26526128493 | 20,303688369635 | 1.35671223509024e-06 |
| **2150** | 24,9954686970178 | + | NA | + | NA | NA | + | + | NA | NA | NA | NA | + | 0,0484723156303957 | 7 | -526,700554223963 | 1068,26932550219 | 20,3077525868982 | 1.35395804779835e-06 |
| **2154** | 25,055236584745 | + | NA | NA | + | NA | + | + | NA | NA | NA | NA | + | 0,0484660355648152 | 7 | -526,701006114264 | 1068,27022928279 | 20,3086563675004 | 1.35334634551007e-06 |
| **3172** | 24,8183276002894 | + | + | NA | NA | NA | + | + | NA | NA | NA | + | + | 0,0639439650674846 | 8 | -525,578121864381 | 1068,28124372876 | 20,319670813471 | 1.34591365094796e-06 |
| **2325** | 26,2176063353021 | NA | NA | + | NA | + | NA | NA | NA | + | NA | NA | + | 0,03281599458764 | 6 | -527,817965236666 | 1068,28208431949 | 20,3205114041957 | 1.3453480885423e-06 |
| **2016** | 25,322631766363 | + | + | + | + | + | NA | + | + | + | + | + | NA | 0,125979767082921 | 12 | -520,883215059625 | 1068,28255915151 | 20,3209862362175 | 1.34502871927901e-06 |
| **2109** | 25,1977438455385 | NA | NA | + | + | + | + | NA | NA | NA | NA | NA | + | 0,0483619564800058 | 7 | -526,708494826348 | 1068,28520670696 | 20,3236337916692 | 1.34324937820573e-06 |
| **1155** | 26,8243956933254 | NA | + | NA | NA | NA | NA | NA | + | NA | NA | + | NA | 0,017210284745269 | 5 | -528,913914666401 | 1068,28584459998 | 20,3242716846862 | 1.3428210218215e-06 |
| **1758** | 25,2110158877168 | + | NA | + | + | + | NA | + | + | NA | + | + | NA | 0,0949510594659356 | 10 | -523,271701018652 | 1068,28943378334 | 20,3278608680453 | 1.3404133674131e-06 |
| **3117** | 25,1675312713123 | NA | NA | + | + | NA | + | NA | NA | NA | NA | + | + | 0,0483295101033245 | 7 | -526,71082924495 | 1068,28987554416 | 20,3283026288739 | 1.34011732904891e-06 |
| **1511** | 25,6499865342141 | NA | + | + | NA | NA | + | + | + | + | NA | + | NA | 0,0794161252466704 | 9 | -524,436952661301 | 1068,29122815725 | 20,3296552419577 | 1.33921130534178e-06 |
| **262** | 26,5083229391462 | + | NA | + | NA | NA | NA | NA | NA | + | NA | NA | NA | 0,0171674387164866 | 5 | -528,916899626543 | 1068,29181452026 | 20,330241604971 | 1.33881873090414e-06 |
| **1508** | 25,6839874748674 | + | + | NA | NA | NA | + | + | + | + | NA | + | NA | 0,0794099878107159 | 9 | -524,437409125985 | 1068,29214108662 | 20,3305681713246 | 1.33860014217503e-06 |
| **2157** | 24,822399839634 | NA | NA | + | + | NA | + | + | NA | NA | NA | NA | + | 0,0483109862952215 | 7 | -526,712161941208 | 1068,29254093668 | 20,3309680213881 | 1.33833254924922e-06 |
| **3333** | 26,1760903576371 | NA | NA | + | NA | NA | NA | NA | NA | + | NA | + | + | 0,0327204505837625 | 6 | -527,824728679307 | 1068,29561120477 | 20,3340382894767 | 1.33627960556534e-06 |
| **62** | 25,3615050055443 | + | NA | + | + | + | + | NA | NA | NA | NA | NA | NA | 0,0482651698712692 | 7 | -526,715458094643 | 1068,29913324355 | 20,3375603282593 | 1.33392846207996e-06 |
| **145** | 27,4719497058982 | NA | NA | NA | NA | + | NA | NA | + | NA | NA | NA | NA | 0,0014790349514701 | 4 | -530,001214485303 | 1068,30545927364 | 20,3438863583447 | 1.32971589201191e-06 |
| **1138** | 25,646394110012 | + | NA | NA | NA | + | + | + | NA | NA | NA | + | NA | 0,0481953758950616 | 7 | -526,720478951118 | 1068,3091749565 | 20,347602041209 | 1.32724778412115e-06 |
| **1202** | 26,0156915633557 | + | NA | NA | NA | + | + | NA | + | NA | NA | + | NA | 0,048183993824249 | 7 | -526,721297722467 | 1068,3108124992 | 20,3492395839078 | 1.32616151642511e-06 |
| **1043** | 26,7480181968397 | NA | + | NA | NA | + | NA | NA | NA | NA | NA | + | NA | 0,0170069057578786 | 5 | -528,928082341591 | 1068,31417995036 | 20,3526070350672 | 1.32393050309125e-06 |
| **20** | 26,3487835632696 | + | + | NA | NA | + | NA | NA | NA | NA | NA | NA | NA | 0,0170041410539769 | 5 | -528,928274914688 | 1068,31456509655 | 20,352992181261 | 1.32367557424118e-06 |
| **2060** | 25,1060406803221 | + | + | NA | + | NA | NA | NA | NA | NA | NA | NA | + | 0,0325370704530787 | 6 | -527,837708062856 | 1068,32156997187 | 20,3599970565745 | 1.31904759240459e-06 |
| **1153** | 27,4756074590319 | NA | NA | NA | NA | NA | NA | NA | + | NA | NA | + | NA | 0,0013330570896470 | 4 | -530,011223680065 | 1068,32547766316 | 20,3639047478694 | 1.31647289310898e-06 |
| **2470** | 26,006571014351 | + | NA | + | NA | NA | + | NA | + | + | NA | NA | + | 0,0636195385543065 | 8 | -525,601848037359 | 1068,32869607472 | 20,3671231594269 | 1.31435612093957e-06 |
| **3268** | 25,2291484195004 | + | + | NA | NA | NA | NA | + | + | NA | NA | + | + | 0,0636155872007787 | 8 | -525,602136959693 | 1068,32927391939 | 20,3677010040951 | 1.31397642895465e-06 |
| **2063** | 24,9496150213737 | NA | + | + | + | NA | NA | NA | NA | NA | NA | NA | + | 0,0324805235217525 | 6 | -527,841709877782 | 1068,32957360172 | 20,3680006864272 | 1.31377955594461e-06 |
| **110** | 25,0274409102818 | + | NA | + | + | NA | + | + | NA | NA | NA | NA | NA | 0,0478898296967918 | 7 | -526,742455078673 | 1068,35312721161 | 20,3915542963184 | 1.29839817987943e-06 |
| **8** | 25,7742458703673 | + | + | + | NA | NA | NA | NA | NA | NA | NA | NA | NA | 0,0166607602123591 | 5 | -528,952188606619 | 1068,36239248041 | 20,4008195651238 | 1.29239708697026e-06 |
| **503** | 25,5966319009869 | NA | + | + | NA | + | + | + | + | + | NA | NA | NA | 0,0789242713010792 | 9 | -524,473524085853 | 1068,36437100635 | 20,4027980910616 | 1.29111919857903e-06 |
| **1196** | 25,4604832964086 | + | + | NA | + | NA | + | NA | + | NA | NA | + | NA | 0,0633715454227363 | 8 | -525,619978893788 | 1068,36495778758 | 20,4033848722852 | 1.29074045189057e-06 |

| | | | | | | | | | | | | | | | | | | |
|---|---|---|---|---|---|---|---|---|---|---|---|---|---|---|---|---|---|---|
| 3337 | 26,1855496395807 | NA | NA | NA | + | NA | NA | NA | NA | + | NA | + | + | 0,0322214548593015 | 6 | -527,860041126213 | 1068,36623609858 | 20,4046631832894 | 1.2899157316193e-06 |
| 269 | 26,5824868149411 | NA | NA | + | + | NA | NA | NA | NA | + | NA | NA | NA | 0,0165801235509161 | 5 | -528,957803086176 | 1068,37362143953 | 20,412048524237 | 1.28516128159463e-06 |
| 3125 | 25,4815083686775 | NA | NA | + | NA | + | + | NA | NA | NA | NA | + | + | 0,0476979288245512 | 7 | -526,756253767634 | 1068,38072458953 | 20,4191516742419 | 1.28060503063555e-06 |
| 2189 | 25,7661658113187 | NA | NA | + | + | NA | NA | NA | + | NA | NA | NA | + | 0,0321066063759141 | 6 | -527,868166036213 | 1068,38248591858 | 20,4209130032896 | 1.2794777436697e-06 |
| 3271 | 25,2379682849182 | NA | + | + | NA | NA | NA | + | + | NA | NA | + | + | 0,063238218073898 | 8 | -525,629724515308 | 1068,38444903062 | 20,4228761153245 | 1.27822248074743e-06 |
| 1115 | 25,4197854134802 | NA | + | NA | + | + | NA | + | NA | NA | NA | + | NA | 0,0476619693280547 | 7 | -526,75883913666 | 1068,38589532758 | 20,4243224122931 | 1.27729847023651e-06 |
| 2124 | 24,5495321733489 | + | + | NA | + | NA | NA | + | NA | NA | NA | NA | + | 0,0476397198478482 | 7 | -526,760438751933 | 1068,38909455813 | 20,4275216428396 | 1.27525691737995e-06 |
| 500 | 25,662556515132 | + | + | NA | NA | + | + | + | + | + | NA | NA | NA | 0,0787513493910924 | 9 | -524,486376924923 | 1068,39007668449 | 20,4285037692018 | 1.27463083939584e-06 |
| 2314 | 26,1032342685364 | + | NA | NA | + | NA | NA | NA | NA | + | NA | NA | + | 0,0320512404251082 | 6 | -527,872082534 | 1068,39031891415 | 20,4287459988632 | 1.27447647204583e-06 |
| 1111 | 25,4341076452015 | NA | + | + | NA | + | NA | + | NA | NA | NA | + | NA | 0,0475890687150685 | 7 | -526,764080150534 | 1068,39637735533 | 20,4348044400415 | 1.2706216431875e-06 |
| 2329 | 26,1503867556845 | NA | NA | NA | + | + | NA | NA | NA | + | NA | NA | + | 0,0320035316269369 | 6 | -527,875457197593 | 1068,39706824134 | 20,4354953260486 | 1.27018279163418e-06 |
| 3114 | 25,4302077498954 | + | NA | NA | + | NA | + | NA | NA | NA | NA | + | + | 0,0475445703522468 | 7 | -526,76727905601 | 1068,40277516628 | 20,4412022509923 | 1.26656353886255e-06 |
| 1233 | 26,4442650574812 | NA | NA | NA | NA | + | NA | + | + | NA | NA | + | NA | 0,0319494208000084 | 6 | -527,879284505038 | 1068,40472285623 | 20,4431499409382 | 1.26533070272042e-06 |
| 2485 | 26,0705150672911 | NA | NA | + | NA | + | + | NA | + | + | NA | NA | + | 0,0630937968406462 | 8 | -525,640279485066 | 1068,40555897013 | 20,4439860548423 | 1.26480183298002e-06 |
| 3493 | 26,0854486281413 | NA | NA | + | NA | NA | + | NA | + | + | NA | + | + | 0,0630716194541082 | 8 | -525,641900166711 | 1068,40880033342 | 20,4472274181308 | 1.26275365203866e-06 |
| 124 | 24,9575832112326 | + | + | NA | + | + | + | + | NA | NA | NA | NA | NA | 0,0630713608814972 | 8 | -525,641919062484 | 1068,40883812497 | 20,4472652096774 | 1.26272979155731e-06 |
| 2120 | 24,6156720493799 | + | + | + | NA | NA | NA | + | NA | NA | NA | NA | + | 0,0474772652082319 | 7 | -526,772117215208 | 1068,41245148468 | 20,4508785693884 | 1.26045050267265e-06 |
| 3283 | 25,1815560825352 | NA | + | NA | NA | + | NA | + | + | NA | NA | + | + | 0,063023376878919 | 8 | -525,645425510834 | 1068,41585102167 | 20,4542781063765 | 1.25830984844359e-06 |
| 1108 | 25,5510281144454 | + | + | NA | NA | + | NA | + | NA | NA | NA | + | NA | 0,0474435676630188 | 7 | -526,774539399291 | 1068,41729585285 | 20,4557229375548 | 1.25740115406008e-06 |
| 2784 | 24,5402157051597 | + | + | + | + | + | NA | + | + | NA | + | NA | + | 0,109622192198562 | 11 | -522,152738880771 | 1068,41747776154 | 20,455904846252 | 1.25728679315808e-06 |
| 2164 | 24,9256302504584 | + | + | NA | NA | + | + | + | NA | NA | NA | NA | + | 0,062974069900019 | 8 | -525,649028449183 | 1068,42305689837 | 20,4614839830756 | 1.25378439299896e-06 |
| 147 | 26,7344618567817 | NA | + | NA | NA | + | NA | NA | + | NA | NA | NA | NA | 0,0162105098571659 | 5 | -528,983532249059 | 1068,42507976529 | 20,4635068500029 | 1.25251691460077e-06 |
| 3129 | 25,4699238866908 | NA | NA | NA | + | + | + | NA | NA | NA | NA | + | + | 0,0473581064165604 | 7 | -526,780681981157 | 1068,42958101658 | 20,4680081012864 | 1.24970113773442e-06 |
| 1070 | 25,330578033884 | + | NA | + | + | NA | + | NA | NA | NA | NA | + | NA | 0,0473517295214496 | 7 | -526,781140302497 | 1068,43049765926 | 20,4689247439669 | 1.24912850426914e-06 |
| 488 | 25,5844318205604 | + | + | + | NA | NA | + | + | + | + | NA | NA | NA | 0,0783582526498316 | 9 | -524,515585821422 | 1068,44849447749 | 20,4869215621991 | 1.23793875542217e-06 |
| 1211 | 25,4179806071314 | NA | + | NA | + | + | + | NA | + | NA | NA | + | NA | 0,0627837901778819 | 8 | -525,662930710678 | 1068,45086142136 | 20,4892885060656 | 1.23647455624027e-06 |
| 1355 | 25,5449409526514 | NA | + | NA | + | NA | NA | + | NA | + | NA | + | NA | 0,0471889756795786 | 7 | -526,792836737806 | 1068,45389052988 | 20,4923176145855 | 1.23460326587686e-06 |
| 1199 | 25,3855018545819 | NA | + | + | + | NA | + | NA | + | NA | NA | + | NA | 0,062699354125377 | 8 | -525,669098893571 | 1068,46319778714 | 20,5016248718507 | 1.22887122850006e-06 |
| 3339 | 25,6729400576458 | NA | + | NA | + | NA | NA | NA | NA | + | NA | + | + | 0,0470395664501665 | 7 | -526,803572393869 | 1068,475361842 | 20,5137889267103 | 1.22141988259516e-06 |
| 393 | 27,2885033281891 | NA | NA | NA | + | NA | NA | NA | + | + | NA | NA | NA | 0,0158233416942268 | 5 | -529,010473035888 | 1068,47896133895 | 20,517388423661 | 1.21922361098473e-06 |
| 2265 | 25,5351375084569 | NA | NA | NA | + | + | NA | + | + | NA | NA | NA | + | 0,0469986251467157 | 7 | -526,806513898065 | 1068,48124485039 | 20,5196719351031 | 1.21783234984297e-06 |
| 3411 | 25,3571641443707 | NA | + | NA | NA | + | NA | + | NA | + | NA | + | + | 0,0625700825204159 | 8 | -525,678541306215 | 1068,48208261243 | 20,5205096971401 | 1.21732232981422e-06 |
| 381 | 25,7167495746032 | NA | NA | + | + | + | + | + | NA | + | NA | NA | NA | 0,0625619883804701 | 8 | -525,67913248486 | 1068,48326496972 | 20,5216920544292 | 1.21660288752973e-06 |
| 1351 | 25,5770528554355 | NA | + | + | NA | NA | NA | + | NA | + | NA | + | NA | 0,0469281578543981 | 7 | -526,811576456138 | 1068,49136996654 | 20,5297970512493 | 1.21168258275975e-06 |
| 120 | 24,9852405098927 | + | + | + | NA | + | + | + | NA | NA | NA | NA | NA | 0,0624779891035096 | 8 | -525,68526731093 | 1068,49553462186 | 20,5339617065692 | 1.20916208776152e-06 |
| 3110 | 25,3823629602727 | + | NA | + | NA | NA | + | NA | NA | NA | NA | + | + | 0,0468813805946088 | 7 | -526,814936852533 | 1068,49809075933 | 20,5365178440397 | 1.20761768263949e-06 |
| 1041 | 27,2739082161362 | NA | NA | NA | NA | + | NA | NA | NA | NA | NA | + | NA | 5,72808969298304E- | 4 | -530,098637031939 | 1068,50030436691 | 20,538731451617 | 1.20628182621499e-06 |
| 1078 | 25,5918744457427 | + | NA | + | NA | + | + | NA | NA | NA | NA | + | NA | 0,0468616723702473 | 7 | -526,816352607488 | 1068,50092226924 | 20,5393493539484 | 1.20590920160296e-06 |
| 1446 | 26,0930416719352 | + | NA | + | NA | NA | + | NA | + | + | NA | + | NA | 0,0624384709074855 | 8 | -525,688153303586 | 1068,50130660717 | 20,5397336918813 | 1.20567748554307e-06 |
| 438 | 26,1059499600177 | + | NA | + | NA | + | + | NA | + | + | NA | NA | NA | 0,0624275715399807 | 8 | -525,688949257156 | 1068,50289851431 | 20,5413255990213 | 1.20471820406659e-06 |
| 1348 | 25,5963794956452 | + | + | NA | NA | NA | NA | + | NA | + | NA | + | NA | 0,0468222800547807 | 7 | -526,819182296076 | 1068,50658164642 | 20,5450087311256 | 1.20250167748609e-06 |
| 2365 | 25,6668996119784 | NA | NA | + | + | + | + | NA | NA | + | NA | NA | + | 0,0623932398703387 | 8 | -525,691456352744 | 1068,50791270549 | 20,5463397901979 | 1.20170164335515e-06 |
| 1082 | 25,659806919598 | + | NA | NA | + | + | + | NA | NA | NA | NA | + | NA | 0,0468097507990283 | 7 | -526,820082292089 | 1068,50838163844 | 20,5468087231507 | 1.20141991763406e-06 |
| 2201 | 25,8697845065743 | NA | NA | NA | + | + | NA | NA | + | NA | NA | NA | + | 0,0311836952991144 | 6 | -527,93342204635 | 1068,51299793885 | 20,5514250235635 | 1.19865005785908e-06 |
| 2102 | 25,4053053543326 | + | NA | + | NA | + | + | NA | NA | NA | NA | NA | + | 0,0467531340947745 | 7 | -526,82414901096 | 1068,51651507618 | 20,5549421608937 | 1.1965440017918e-06 |
| 2106 | 25,4417941314209 | + | NA | NA | + | + | + | NA | NA | NA | NA | NA | + | 0,0467492416818713 | 7 | -526,824428590054 | 1068,51707423437 | 20,5555013190801 | 1.19620951986404e-06 |
| 341 | 26,4053879762017 | NA | NA | + | NA | + | NA | + | NA | + | NA | NA | NA | 0,0311547078835316 | 6 | -527,935470644678 | 1068,51709513551 | 20,5555222202186 | 1.19619701885897e-06 |
| 2421 | 25,6961124455426 | NA | NA | + | NA | + | + | + | NA | + | NA | NA | + | 0,0623007893585598 | 8 | -525,698207163589 | 1068,52141432718 | 20,5598414118883 | 1.19361650419663e-06 |
| 141 | 26,5470454607138 | NA | NA | + | + | NA | NA | NA | + | NA | NA | NA | NA | 0,015467040791599 | 5 | -529,035256590335 | 1068,52852844785 | 20,5669555325551 | 1.18937828055363e-06 |
| 3209 | 26,1443000625412 | NA | NA | NA | + | NA | NA | NA | + | NA | NA | + | + | 0,0310603490606755 | 6 | -527,942138746131 | 1068,53043133842 | 20,5688584231257 | 1.1882471903667e-06 |
| 2253 | 25,5662897314849 | NA | NA | + | + | NA | NA | + | + | NA | NA | NA | + | 0,0466479152583466 | 7 | -526,831706129125 | 1068,53162931251 | 20,5700563972221 | 1.18753565880999e-06 |
| 326 | 25,976601187613 | + | NA | + | NA | NA | NA | + | NA | + | NA | NA | NA | 0,030957827740383 | 6 | -527,949382935133 | 1068,54491971642 | 20,5833468011294 | 1.17967040656803e-06 |
| 2068 | 25,6444704708089 | + | + | NA | NA | + | NA | NA | NA | NA | NA | NA | + | 0,0308893098548312 | 6 | -527,954224003588 | 1068,55460185333 | 20,5930289380394 | 1.17397334244056e-06 |
| 3273 | 25,751048178085 | NA | NA | NA | + | NA | NA | + | + | NA | NA | + | + | 0,0464460601652488 | 7 | -526,846201606482 | 1068,56062026723 | 20,5990473519371 | 1.17044592372902e-06 |
| 2094 | 25,0132490988936 | + | NA | + | + | NA | + | NA | NA | NA | NA | NA | + | 0,0464454690258167 | 7 | -526,846244052471 | 1068,56070515921 | 20,5991322439147 | 1.17039624404879e-06 |
| 3179 | 24,6465764189115 | NA | + | NA | + | NA | + | + | NA | NA | NA | + | + | 0,0620279897273844 | 8 | -525,718123335304 | 1068,56124667061 | 20,5996737553173 | 1.17007939548914e-06 |
| 1132 | 24,8893593007994 | + | + | NA | + | NA | + | + | NA | NA | NA | + | NA | 0,0619981605217095 | 8 | -525,720300712831 | 1068,56560142566 | 20,6040285103704 | 1.16753446255447e-06 |
| 3175 | 24,6959670274917 | NA | + | + | NA | NA | + | + | NA | NA | NA | + | + | 0,0618863651963239 | 8 | -525,728460576863 | 1068,58192115373 | 20,6203482384344 | 1.15804630377621e-06 |
| 2250 | 25,7003652224606 | + | NA | NA | + | NA | NA | + | + | NA | NA | NA | + | 0,0462903685213564 | 7 | -526,857379931371 | 1068,58297691701 | 20,6214040017151 | 1.15743515371544e-06 |
| 2413 | 25,3770652161166 | NA | NA | + | + | NA | + | + | NA | + | NA | NA | + | 0,0617519501540276 | 8 | -525,738270151751 | 1068,6015403035 | 20,639967388212 | 1.14674189819088e-06 |
| 2056 | 25,2055830594037 | + | + | + | NA | NA | NA | NA | NA | NA | NA | NA | + | 0,0305060562576118 | 6 | -527,981296128349 | 1068,60874610285 | 20,6471731875602 | 1.142617736112e-06 |
| 92 | 25,3512207382566 | + | + | NA | + | + | NA | + | NA | NA | NA | NA | NA | 0,0460259084129346 | 7 | -526,876363417989 | 1068,62094389024 | 20,6593709749502 | 1.13567023956206e-06 |
| 2186 | 26,0140961011649 | + | NA | NA | + | NA | NA | NA | + | NA | NA | NA | + | 0,0303806668203081 | 6 | -527,99015101832 | 1068,62645588279 | 20,6648829675023 | 1.13254464565196e-06 |
| 2127 | 24,4779634290048 | NA | + | + | + | NA | NA | + | NA | NA | NA | NA | + | 0,0459439828887358 | 7 | -526,882243132261 | 1068,63270331878 | 20,6711304034941 | 1.12901241527916e-06 |
| 95 | 25,2206868994256 | NA | + | + | + | + | NA | + | NA | NA | NA | NA | NA | 0,045935076625859 | 7 | -526,882882295571 | 1068,63398164541 | 20,6724087301154 | 1.12829102253447e-06 |
| 88 | 25,3427605655972 | + | + | + | NA | + | NA | + | NA | NA | NA | NA | NA | 0,0459219780129113 | 7 | -526,88382231457 | 1068,6358616834 | 20,674288768113 | 1.12723090588009e-06 |
| 1461 | 26,2119829394519 | NA | NA | + | NA | + | + | NA | + | + | NA | + | NA | 0,0615081981789495 | 8 | -525,756055524829 | 1068,63711104966 | 20,6755381343676 | 1.12652696364594e-06 |
| 318 | 25,7849104671746 | + | NA | + | + | + | + | NA | NA | + | NA | NA | NA | 0,0614853752191987 | 8 | -525,757720566639 | 1068,64044113328 | 20,6788682179881 | 1.12465280985627e-06 |
| 1341 | 25,7878150932689 | NA | NA | + | + | + | + | NA | NA | + | NA | + | NA | 0,0614831347702221 | 8 | -525,757884015704 | 1068,64076803141 | 20,6791951161181 | 1.12446900142813e-06 |
| 395 | 26,6336772348896 | NA | + | NA | + | NA | NA | NA | + | + | NA | NA | NA | 0,0302704082719188 | 6 | -527,997936432753 | 1068,64202671166 | 20,680453796369 | 1.12376155060266e-06 |
| 2449 | 26,6981282152606 | NA | NA | NA | NA | + | NA | NA | + | + | NA | NA | + | 0,030260034924467 | 6 | -527,998668854766 | 1068,64349155569 | 20,6819186403948 | 1.12293878424862e-06 |
| 138 | 26,6861473644008 | + | NA | NA | + | NA | NA | NA | + | NA | NA | NA | NA | 0,014634017145858 | 5 | -529,093165028603 | 1068,64434532438 | 20,682772409092 | 1.12245952155934e-06 |
| 3804 | 24,4832787322771 | + | + | NA | + | + | NA | + | + | NA | + | + | + | 0,108132103307284 | 11 | -522,267225023204 | 1068,64645004641 | 20,6848771311177 | 1.12127891024397e-06 |
| 2391 | 25,2461071915346 | NA | + | + | NA | + | NA | + | NA | + | NA | NA | + | 0,0614278672749015 | 8 | -525,761915861064 | 1068,64883172213 | 20,6872588068377 | 1.11994444359042e-06 |
| 2434 | 26,769483908456 | + | NA | NA | NA | NA | NA | NA | + | + | NA | NA | + | 0,0301898570463921 | 6 | -528,003623638095 | 1068,65340112234 | 20,691828207053 | 1.11738862714836e-06 |
| 1128 | 24,9824757275791 | + | + | + | NA | NA | + | + | NA | NA | NA | + | NA | 0,0613879480744198 | 8 | -525,764827878062 | 1068,65465575612 | 20,6930828408324 | 1.11668789020525e-06 |

| | | | | | | | | | | | | | | | | | | |
|---|---|---|---|---|---|---|---|---|---|---|---|---|---|---|---|---|---|---|
| 3499 | 25,6864735560969 | NA | + | NA | + | NA | + | NA | + | + | NA | + | + | 0,0769632094725861 | 9 | -524,619143532593 | 1068,65560989983 | 20,6940369845413 | 1.11615527690015e-06 |
| 333 | 26,06219577122 | NA | NA | + | + | NA | NA | + | NA | + | NA | NA | NA | 0,0301312847440271 | 6 | -528,007758756175 | 1068,6616713585 | 20,7000984432127 | 1.1127776333064e-06 |
| 374 | 25,7967725312467 | + | NA | + | NA | + | + | + | NA | + | NA | NA | NA | 0,0613211527242949 | 8 | -525,76970017362 | 1068,66440034724 | 20,7028274319503 | 1.11126028993072e-06 |
| 1397 | 25,8191887044002 | NA | NA | + | NA | + | + | + | NA | + | NA | + | NA | 0,061249367103968 | 8 | -525,774936091288 | 1068,67487218258 | 20,7132992672846 | 1.10545702850332e-06 |
| 1760 | 24,9395160644254 | + | + | + | + | + | NA | + | + | NA | + | + | NA | 0,107913485203026 | 11 | -522,284005751222 | 1068,68001150244 | 20,7184385871526 | 1.1026200264947e-06 |
| 2167 | 24,740344015512 | NA | + | + | NA | + | + | + | NA | NA | NA | NA | + | 0,0612127959567973 | 8 | -525,777603372964 | 1068,68020674593 | 20,7186338306369 | 1.10251239206059e-06 |
| 2388 | 25,3595725597363 | + | + | NA | NA | + | NA | + | NA | + | NA | NA | + | 0,0611169405588621 | 8 | -525,784594001996 | 1068,69418800399 | 20,7326150887009 | 1.09483201354068e-06 |
| 127 | 24,7718378265214 | NA | + | + | + | + | + | + | NA | NA | NA | NA | NA | 0,0611095426829238 | 8 | -525,78513349132 | 1068,69526698264 | 20,7336940673499 | 1.09424152265331e-06 |
| 3373 | 25,5302351116272 | NA | NA | + | + | NA | + | NA | NA | + | NA | + | + | 0,0610982425272009 | 8 | -525,785957545786 | 1068,69691509157 | 20,7353421762809 | 1.09334017946917e-06 |
| 2171 | 24,7415272128013 | NA | + | NA | + | + | + | + | NA | NA | NA | NA | + | 0,061026774973205 | 8 | -525,791169028172 | 1068,70733805634 | 20,7457651410539 | 1.08765707793575e-06 |
| 3429 | 25,5474819533282 | NA | NA | + | NA | NA | + | + | NA | + | NA | + | + | 0,0610224359177057 | 8 | -525,791485423482 | 1068,70797084696 | 20,7463979316742 | 1.08731300277198e-06 |
| 2395 | 25,2432247836996 | NA | + | NA | + | + | NA | + | NA | + | NA | NA | + | 0,0609557486098361 | 8 | -525,79634794572 | 1068,71769589144 | 20,7561229761498 | 1.08203875258806e-06 |
| 335 | 25,3933612795019 | NA | + | + | + | NA | NA | + | NA | + | NA | NA | NA | 0,045300020099792 | 7 | -526,928442141943 | 1068,72510133815 | 20,7635284228591 | 1.07803967074409e-06 |
| 2252 | 24,9589151318297 | + | + | NA | + | NA | NA | + | + | NA | NA | NA | + | 0,0608894816224023 | 8 | -525,801179478296 | 1068,72735895659 | 20,7657860413015 | 1.07682345618918e-06 |
| 2406 | 25,4854207400693 | + | NA | + | NA | NA | + | + | NA | + | NA | NA | + | 0,0608650336585081 | 8 | -525,802961895698 | 1068,7309237914 | 20,7693508761058 | 1.07490581684636e-06 |
| 3396 | 25,2123139802024 | + | + | NA | NA | NA | NA | + | NA | + | NA | + | + | 0,060824377969289 | 8 | -525,805925860212 | 1068,73685172042 | 20,7752788051343 | 1.07172455105747e-06 |
| 2350 | 25,4778326805444 | + | NA | + | + | NA | + | NA | NA | + | NA | NA | + | 0,0608152337560185 | 8 | -525,806592492753 | 1068,73818498551 | 20,7766120702161 | 1.07101034268063e-06 |
| 2331 | 25,7522116941915 | NA | + | NA | + | + | NA | NA | NA | + | NA | NA | + | 0,0451847038107046 | 7 | -526,936711844556 | 1068,74164074338 | 20,7800678280844 | 1.06916136433663e-06 |
| 266 | 26,7361573964562 | + | NA | NA | + | NA | NA | NA | NA | + | NA | NA | NA | 0,013907187612268 | 5 | -529,143651320919 | 1068,74531790901 | 20,7837449937231 | 1.06719742860354e-06 |
| 332 | 25,4229029178046 | + | + | NA | + | NA | NA | + | NA | + | NA | NA | NA | 0,0451336835623905 | 7 | -526,940370352389 | 1068,74895775904 | 20,7873848437519 | 1.06525697558301e-06 |
| 328 | 25,4162700672886 | + | + | + | NA | NA | NA | + | NA | + | NA | NA | NA | 0,0451242215891726 | 7 | -526,941048820418 | 1068,7503146951 | 20,7887417798097 | 1.06453447790599e-06 |
| 3281 | 25,7914825191155 | NA | NA | NA | NA | + | NA | + | + | NA | NA | + | + | 0,0451224545701934 | 7 | -526,941175523254 | 1068,75056810077 | 20,7889951854811 | 1.06439960691339e-06 |
| 2261 | 25,6347295917747 | NA | NA | + | NA | + | NA | + | + | NA | NA | NA | + | 0,0450472586097864 | 7 | -526,946567178785 | 1068,76135141183 | 20,799778496543 | 1.0586761741396e-06 |
| 2439 | 26,020159379812 | NA | + | + | NA | NA | NA | NA | + | + | NA | NA | + | 0,0450188116952609 | 7 | -526,948606751708 | 1068,76543055768 | 20,8038576423899 | 1.05651912735501e-06 |
| 2482 | 26,2403228113455 | + | NA | NA | NA | + | + | NA | + | + | NA | NA | + | 0,0605854391928257 | 8 | -525,82334287409 | 1068,77168574818 | 20,8101128328897 | 1.05321992512483e-06 |
| 2319 | 25,4123750571848 | NA | + | + | + | NA | NA | NA | NA | + | NA | NA | + | 0,0449427079022879 | 7 | -526,954062905592 | 1068,77634286545 | 20,8147699501562 | 1.05077029393722e-06 |
| 3744 | 24,8109588849615 | + | + | + | + | + | NA | NA | + | NA | + | + | + | 0,107243691407793 | 11 | -522,335392328652 | 1068,78278465731 | 20,8212117420144 | 1.04739131676317e-06 |
| 3403 | 25,1012704412396 | NA | + | NA | + | NA | NA | + | NA | + | NA | + | + | 0,0605016323619889 | 8 | -525,829450773931 | 1068,78390154786 | 20,822328632572 | 1.04680656931721e-06 |
| 3459 | 26,3203397350835 | NA | + | NA | NA | NA | NA | NA | + | + | NA | + | + | 0,0448692704135641 | 7 | -526,959327490571 | 1068,78687203541 | 20,8252991201155 | 1.04525296040947e-06 |
| 3269 | 25,7517693150406 | NA | NA | + | NA | NA | NA | + | + | NA | NA | + | + | 0,0448431609577882 | 7 | -526,961199127134 | 1068,79061530853 | 20,829042393241 | 1.04329845638257e-06 |
| 1389 | 25,5601091353657 | NA | NA | + | + | NA | + | + | NA | + | NA | + | NA | 0,0604552354531412 | 8 | -525,832831978562 | 1068,79066395712 | 20,829091041833 | 1.04327307919078e-06 |
| 1349 | 26,1401521539372 | NA | NA | + | NA | NA | NA | + | NA | + | NA | + | NA | 0,0292141582216593 | 6 | -528,072474001053 | 1068,79110184826 | 20,8295289329685 | 1.04304468417806e-06 |
| 3335 | 25,8228237687138 | NA | + | + | NA | NA | NA | NA | NA | + | NA | + | + | 0,0448292817090551 | 7 | -526,962194029751 | 1068,79260511376 | 20,8310321984741 | 1.04226099219199e-06 |
| 2258 | 25,6867556573987 | + | NA | NA | NA | + | NA | + | + | NA | NA | NA | + | 0,0447732934673013 | 7 | -526,966207273771 | 1068,8006316018 | 20,8390586865144 | 1.03808652667491e-06 |
| 2220 | 25,1574490337215 | + | + | NA | + | NA | + | NA | + | NA | NA | NA | + | 0,0603834011989141 | 8 | -525,838066616914 | 1068,80113323383 | 20,8395603185384 | 1.03782619060177e-06 |
| 3122 | 25,7461449641462 | + | NA | NA | NA | + | + | NA | NA | NA | NA | + | + | 0,0447660834332838 | 7 | -526,966724072803 | 1068,80166519987 | 20,8400922845783 | 1.03755018316566e-06 |
| 2246 | 25,6618648811339 | + | NA | + | NA | NA | NA | + | + | NA | NA | NA | + | 0,0447410811336584 | 7 | -526,968516151238 | 1068,80524935674 | 20,8436764414496 | 1.03569247693133e-06 |
| 366 | 25,5124997364142 | + | NA | + | + | NA | + | + | NA | + | NA | NA | NA | 0,0603357600707931 | 8 | -525,841538055879 | 1068,80807611176 | 20,8465031964672 | 1.03422968645948e-06 |
| 3399 | 25,1240956358904 | NA | + | + | NA | NA | NA | + | NA | + | NA | + | + | 0,060329321465966 | 8 | -525,842007200538 | 1068,80901440108 | 20,8474414857844 | 1.03374459692378e-06 |
| 2358 | 25,8105914999907 | + | NA | + | NA | + | + | NA | NA | + | NA | NA | + | 0,060328316754298 | 8 | -525,842080407895 | 1068,80916081579 | 20,8475879004998 | 1.03366892198335e-06 |
| 1135 | 24,7935700767435 | NA | + | + | + | NA | + | + | NA | NA | NA | + | NA | 0,060321430337078 | 8 | -525,842582178014 | 1068,81016435603 | 20,8485914407368 | 1.03315038790921e-06 |
| 3239 | 25,4407987114885 | NA | + | + | NA | NA | + | NA | + | NA | NA | + | + | 0,0602985997248228 | 8 | -525,844245675533 | 1068,81349135107 | 20,8519184357751 | 1.03143317348894e-06 |
| 2255 | 24,9882798066747 | NA | + | + | + | NA | NA | + | + | NA | NA | NA | + | 0,0602918814622636 | 8 | -525,844735177844 | 1068,81447035569 | 20,852897440398 | 1.0309284081184e-06 |
| 3236 | 25,5711386095652 | + | + | NA | NA | NA | + | NA | + | NA | NA | + | + | 0,0602414607534816 | 8 | -525,84840879186 | 1068,82181758372 | 20,8602446684301 | 1.02714812297441e-06 |
| 2267 | 24,966827679983 | NA | + | NA | + | + | NA | + | + | NA | NA | NA | + | 0,0601980684347764 | 8 | -525,851570165062 | 1068,82814033012 | 20,8665674148324 | 1.02390605182346e-06 |
| 29 | 26,3781250226405 | NA | NA | + | + | + | NA | NA | NA | NA | NA | NA | NA | 0,0132832556321542 | 5 | -529,186960587132 | 1068,83193644144 | 20,8703635261486 | 1.02196446435187e-06 |
| 3381 | 25,865407787184 | NA | NA | + | NA | + | + | NA | NA | + | NA | + | + | 0,0601616675630783 | 8 | -525,854222059701 | 1068,8334441194 | 20,871871204112 | 1.02119435800527e-06 |
| 2425 | 25,7342007840667 | NA | NA | NA | + | + | + | + | NA | + | NA | NA | + | 0,0601210996521282 | 8 | -525,857177412622 | 1068,83935482524 | 20,8777819099535 | 1.01818082349911e-06 |
| 3505 | 26,3310948937915 | NA | NA | NA | NA | + | + | NA | + | + | NA | + | + | 0,0600985437797541 | 8 | -525,858820541989 | 1068,84264108398 | 20,8810681686875 | 1.01650919441458e-06 |
| 2479 | 25,5460855473484 | NA | + | + | + | NA | + | NA | + | + | NA | NA | + | 0,0756847840143037 | 9 | -524,713907127133 | 1068,84513708891 | 20,8835641736205 | 1.01524137971525e-06 |
| 2389 | 25,7709983851792 | NA | NA | + | NA | + | NA | + | NA | + | NA | NA | + | 0,0444059137453238 | 7 | -526,992535265563 | 1068,85328758539 | 20,8917146700994 | 1.01111243801546e-06 |
| 3251 | 25,5320002470199 | NA | + | NA | NA | + | + | NA | + | NA | NA | + | + | 0,060019249485894 | 8 | -525,864596586694 | 1068,85419317339 | 20,8926202580978 | 1.0106547160057e-06 |
| 2476 | 25,6445031418265 | + | + | NA | + | NA | + | NA | + | + | NA | NA | + | 0,0755918710809867 | 9 | -524,720789213221 | 1068,85890126109 | 20,8973283457967 | 1.00827838857444e-06 |
| 134 | 26,4466512243548 | + | NA | + | NA | NA | NA | NA | + | NA | NA | NA | NA | 0,0130649242878586 | 5 | -529,202109252104 | 1068,86223377138 | 20,9006608560931 | 1.0065997384368e-06 |
| 188 | 25,3418785441478 | + | + | NA | + | + | + | NA | + | NA | NA | NA | NA | 0,0599093424605988 | 8 | -525,872601753328 | 1068,87020350666 | 20,9086305913652 | 1.00259655309575e-06 |
| 1455 | 25,7470376115132 | NA | + | + | + | NA | + | NA | + | + | NA | + | NA | 0,0755039309583121 | 9 | -524,727302325029 | 1068,8719274847 | 20,9103545694129 | 1.00173269824182e-06 |
| 2260 | 25,1214403263485 | + | + | NA | NA | + | NA | + | + | NA | NA | NA | + | 0,0598906958048921 | 8 | -525,873959804684 | 1068,87291960937 | 20,9113466940773 | 1.001235899615e-06 |
| 345 | 26,4531243852994 | NA | NA | NA | + | + | NA | + | NA | + | NA | NA | NA | 0,0286177772575968 | 6 | -528,114523679549 | 1068,87520120525 | 20,9136282899619 | 1.00009434302747e-06 |
| 2381 | 25,5065813583552 | NA | NA | + | + | NA | NA | + | NA | + | NA | NA | + | 0,0442220599760675 | 7 | -527,005707209591 | 1068,87963147345 | 20,918058558156 | 9.97881451778281e-07 |
| 2248 | 25,1313648756309 | + | + | + | NA | NA | NA | + | + | NA | NA | NA | + | 0,0597943849403335 | 8 | -525,880973775962 | 1068,88694755192 | 20,9253746366342 | 9.94237830589364e-07 |
| 3443 | 25,3624961117218 | NA | + | NA | NA | + | + | + | NA | + | NA | + | + | 0,0753845569212189 | 9 | -524,736142536971 | 1068,88960790859 | 20,9280349932978 | 9.92916196170024e-07 |
| 2491 | 25,6473843969871 | NA | + | NA | + | + | + | NA | + | + | NA | NA | + | 0,0753486147234401 | 9 | -524,738804003232 | 1068,89493084111 | 20,9333579258196 | 9.90277096708959e-07 |
| 3211 | 25,6209378429247 | NA | + | NA | + | NA | NA | NA | + | NA | NA | + | + | 0,0441063965665911 | 7 | -527,013992453711 | 1068,89620196169 | 20,9346290463957 | 9.8964791587447e-07 |
| 1452 | 25,8425438893096 | + | + | NA | + | NA | + | NA | + | + | NA | + | NA | 0,0753015931610698 | 9 | -524,742285723828 | 1068,9018942823 | 20,94032136701 | 9.86835223843155e-07 |
| 93 | 25,9017367835905 | NA | NA | + | + | + | NA | + | NA | NA | NA | NA | NA | 0,0284082312658623 | 6 | -528,129292239664 | 1068,90473832548 | 20,943165410192 | 9.85432920137472e-07 |
| 2235 | 25,1607888132728 | NA | + | NA | + | + | + | NA | + | NA | NA | NA | + | 0,059660366996084 | 8 | -525,890732622375 | 1068,90646524475 | 20,9448923294594 | 9.84582405834435e-07 |
| 330 | 26,1801675301267 | + | NA | NA | + | NA | NA | + | NA | + | NA | NA | NA | 0,0283762064976968 | 6 | -528,131549027964 | 1068,90925190208 | 20,9476789867917 | 9.83211514217273e-07 |
| 153 | 26,8561291223883 | NA | NA | NA | + | + | NA | NA | + | NA | NA | NA | NA | 0,0127234076998117 | 5 | -529,2257982646 | 1068,90961179637 | 20,9480388810844 | 9.83034604028792e-07 |
| 1084 | 25,3677948915002 | + | + | NA | + | + | + | NA | NA | NA | NA | + | NA | 0,0596377803742883 | 8 | -525,892377185932 | 1068,90975437186 | 20,9481814565731 | 9.82964528207017e-07 |
| 3433 | 25,6372139731101 | NA | NA | NA | + | NA | + | + | NA | + | NA | + | + | 0,0596348538175723 | 8 | -525,892590269735 | 1068,91018053947 | 20,9486076241785 | 9.82755096701459e-07 |
| 1467 | 25,8374416463335 | NA | + | NA | + | + | + | NA | + | + | NA | + | NA | 0,0752415019295544 | 9 | -524,746734933006 | 1068,91079270066 | 20,9492197853676 | 9.8245434046733e-07 |
| 3490 | 26,381457183957 | + | NA | NA | NA | NA | + | NA | + | + | NA | + | + | 0,0596253426036415 | 8 | -525,89328278057 | 1068,91156556114 | 20,9499926458491 | 9.82074763744728e-07 |
| 2410 | 25,6084231342592 | + | NA | NA | + | NA | + | + | NA | + | NA | NA | + | 0,0596229469615509 | 8 | -525,893457206003 | 1068,91191441201 | 20,9503414967148 | 9.81903479867514e-07 |
| 1326 | 25,6349603040933 | + | NA | + | + | NA | + | NA | NA | + | NA | + | NA | 0,0596060648383088 | 8 | -525,894686371869 | 1068,91437274374 | 20,9527998284475 | 9.80697299076018e-07 |
| 2327 | 25,8919685376041 | NA | + | + | NA | + | NA | NA | NA | + | NA | NA | + | 0,0439592465896717 | 7 | -527,024531707023 | 1068,91728046831 | 20,9557075530199 | 9.79272536214159e-07 |
| 3266 | 25,906484563793 | + | NA | NA | NA | NA | NA | + | + | NA | NA | + | + | 0,0439335226351184 | 7 | -527,026373955173 | 1068,92096496461 | 20,9593920493187 | 9.77470133942459e-07 |

| | | | | | | | | | | | | | | | | | | | |
|---|---|---|---|---|---|---|---|---|---|---|---|---|---|---|---|---|---|---|---|
| 1399 | 25,4296161297336 | NA | + | + | NA | + | + | + | NA | + | NA | + | NA | 0,0751728032791795 | 9 | -524,751821089254 | 1068,92096501315 | 20,9593920978632 | 9.77470110217056e-07 |
| 283 | 26,5294027346715 | NA | + | NA | + | + | NA | NA | NA | + | NA | NA | NA | 0,0282522259197962 | 6 | -528,14028524943 | 1068,92672434501 | 20,9651514297236 | 9.74659371779057e-07 |
| 1161 | 26,902583305503 | NA | NA | NA | + | NA | NA | NA | + | NA | NA | + | NA | 0,0125834738593221 | 5 | -529,235502292854 | 1068,92901985288 | 20,9674469375932 | 9.73541344382926e-07 |
| 86 | 25,8576860925892 | + | NA | + | NA | + | NA | + | NA | NA | NA | NA | NA | 0,02822934094086 | 6 | -528,141897704753 | 1068,92994925566 | 20,9683763403698 | 9.73089043469199e-07 |
| 3085 | 25,5557860081733 | NA | NA | + | + | NA | NA | NA | NA | NA | NA | + | + | 0,0282211349652423 | 6 | -528,142475881274 | 1068,9311056087 | 20,9695326934107 | 9.7252658884667e-07 |
| 1382 | 25,6320175219132 | + | NA | + | NA | NA | + | + | NA | + | NA | + | NA | 0,0594877998736329 | 8 | -525,903296474743 | 1068,93159294949 | 20,9700200341952 | 9.72289641780955e-07 |
| 1334 | 25,9041926433935 | + | NA | + | NA | + | + | NA | NA | + | NA | + | NA | 0,0594544276689638 | 8 | -525,90572589214 | 1068,93645178428 | 20,9748788689885 | 9.69930411348686e-07 |
| 338 | 26,4450586602761 | + | NA | NA | NA | + | NA | + | NA | + | NA | NA | NA | 0,0281725935569427 | 6 | -528,145895911359 | 1068,93794566887 | 20,9763727535817 | 9.69206199805492e-07 |
| 3457 | 26,915549274998 | NA | NA | NA | NA | NA | NA | NA | + | + | NA | + | + | 0,0281693140846335 | 6 | -528,146126963474 | 1068,9384077731 | 20,976834857812 | 9.68982288531575e-07 |
| 2423 | 25,2464929997732 | NA | + | + | NA | + | + | + | NA | + | NA | NA | + | 0,0750373402073724 | 9 | -524,761849093275 | 1068,94102102119 | 20,9794481059043 | 9.67717019768393e-07 |
| 2223 | 25,0898931070763 | NA | + | + | + | NA | + | NA | + | NA | NA | NA | + | 0,0594015077579714 | 8 | -525,909578159585 | 1068,94415631917 | 20,9825834038786 | 9.66201167634988e-07 |
| 14 | 25,9201256982475 | + | NA | + | + | NA | NA | NA | NA | NA | NA | NA | NA | 0,012462290521022 | 5 | -529,243904914573 | 1068,94582509632 | 20,9842521810315 | 9.65395316661351e-07 |
| 3305 | 25,5409245478254 | NA | NA | NA | + | NA | + | + | + | NA | NA | + | + | 0,0593848991073934 | 8 | -525,910787129878 | 1068,94657425976 | 20,985001344465 | 9.65033764945847e-07 |
| 1403 | 25,4142813708507 | NA | + | NA | + | + | + | + | NA | + | NA | + | NA | 0,0749914658396027 | 9 | -524,765244729192 | 1068,94781229303 | 20,9862393777398 | 9.64436577843123e-07 |
| 2263 | 25,0891438968009 | NA | + | + | NA | + | NA | + | + | NA | NA | NA | + | 0,0593663358911322 | 8 | -525,912138350888 | 1068,94927670178 | 20,987703786486 | 9.63730671628601e-07 |
| 3366 | 25,6818915771735 | + | NA | + | NA | NA | + | NA | NA | + | NA | + | + | 0,0593432021693854 | 8 | -525,913822222796 | 1068,95264444559 | 20,9910715303013 | 9.62109238150387e-07 |
| 383 | 25,2854125281642 | NA | + | + | + | + | + | + | NA | + | NA | NA | NA | 0,0749364143855669 | 9 | -524,769319433888 | 1068,95596170242 | 20,99438878713 | 9.60514779103348e-07 |
| 176 | 25,2641273924607 | + | + | + | + | NA | + | NA | + | NA | NA | NA | NA | 0,0592777371221449 | 8 | -525,918587109588 | 1068,96217421918 | 21,0006013038849 | 9.57535801157008e-07 |
| 2374 | 25,567488613286 | + | NA | + | NA | NA | NA | + | NA | + | NA | NA | + | 0,0436278178236642 | 7 | -527,048263535582 | 1068,96474412543 | 21,0031712101377 | 9.56306202693049e-07 |
| 3397 | 25,6543215229086 | NA | NA | + | NA | NA | NA | + | NA | + | NA | + | + | 0,0436187437587756 | 7 | -527,048913164868 | 1068,966043384 | 21,0044704687098 | 9.5568515992298e-07 |
| 2316 | 25,5652957317049 | + | + | NA | + | NA | NA | NA | NA | + | NA | NA | + | 0,0436167032157289 | 7 | -527,049059250315 | 1068,96633555489 | 21,0047626396033 | 9.55545558426541e-07 |
| 191 | 25,21541054012 | NA | + | + | + | + | + | NA | + | NA | NA | NA | NA | 0,059241574810142 | 8 | -525,921219048724 | 1068,96743809745 | 21,0058651821569 | 9.55018938775009e-07 |
| 447 | 25,6938402326239 | NA | + | + | + | + | + | NA | + | + | NA | NA | NA | 0,0748461630464328 | 9 | -524,775998979329 | 1068,9693207933 | 21,0077478780124 | 9.54120356681555e-07 |
| 2282 | 25,4811733217951 | + | NA | NA | + | NA | + | + | + | NA | NA | NA | + | 0,0592253464875829 | 8 | -525,922400133841 | 1068,96980026768 | 21,0082273523908 | 9.53891645965423e-07 |
| 380 | 25,4203399761308 | + | + | NA | + | + | + | + | NA | + | NA | NA | NA | 0,0747532376477289 | 9 | -524,782875752718 | 1068,98307434008 | 21,0215014247917 | 9.47581595758915e-07 |
| 2285 | 25,3461971011898 | NA | NA | + | + | NA | + | + | + | NA | NA | NA | + | 0,0590819973717331 | 8 | -525,932832090406 | 1068,99066418081 | 21,0290912655214 | 9.43992413701307e-07 |
| 2297 | 25,3544909540967 | NA | NA | NA | + | + | + | + | + | NA | NA | NA | + | 0,059081827308651 | 8 | -525,932844465477 | 1068,99068893095 | 21,0291160156628 | 9.43980731800733e-07 |
| 3131 | 25,1278864857485 | NA | + | NA | + | + | + | NA | NA | NA | NA | + | + | 0,0590171607890722 | 8 | -525,937549926438 | 1069,00009985288 | 21,0385269375854 | 9.39549301456754e-07 |
| 378 | 25,9055321607005 | + | NA | NA | + | + | + | + | NA | + | NA | NA | NA | 0,059006957044295 | 8 | -525,938292372665 | 1069,00158474533 | 21,0400118300392 | 9.38851995510994e-07 |
| 1207 | 25,576229581042 | NA | + | + | NA | + | + | NA | + | NA | NA | + | NA | 0,0590068931710891 | 8 | -525,93829702019 | 1069,00159404038 | 21,0400211250899 | 9.38847632182717e-07 |
| 3116 | 25,023755054018 | + | + | NA | + | NA | + | NA | NA | NA | NA | + | + | 0,0589889707537826 | 8 | -525,93960107392 | 1069,00420214784 | 21,0426292325494 | 9.37624122360945e-07 |
| 376 | 25,4227715374401 | + | + | + | NA | + | + | + | NA | + | NA | NA | NA | 0,074601213539619 | 9 | -524,794124529153 | 1069,00557189295 | 21,043998977661 | 9.36982189178204e-07 |
| 2420 | 25,3800647006076 | + | + | NA | NA | + | + | + | NA | + | NA | NA | + | 0,0745935924543779 | 9 | -524,794688390331 | 1069,00669961531 | 21,0451267000174 | 9.36454010220867e-07 |
| 1458 | 26,3882996276866 | + | NA | NA | NA | + | + | NA | + | + | NA | + | NA | 0,0589621919245368 | 8 | -525,94154948314 | 1069,00809896628 | 21,0465260509889 | 9.35799025471491e-07 |
| 1401 | 25,8809378926148 | NA | NA | NA | + | + | + | + | NA | + | NA | + | NA | 0,0589611443121229 | 8 | -525,941625705573 | 1069,00825141115 | 21,0466784958551 | 9.35727699311231e-07 |
| 1396 | 25,5774962444107 | + | + | NA | NA | + | + | + | NA | + | NA | + | NA | 0,074580880303813 | 9 | -524,795628913795 | 1069,00858066223 | 21,047007746944 | 9.35573667308404e-07 |
| 1353 | 26,2384381804239 | NA | NA | NA | + | NA | NA | + | NA | + | NA | + | NA | 0,0276186386178301 | 6 | -528,184913233536 | 1069,01598031323 | 21,0544073979356 | 9.32118603508744e-07 |
| 2436 | 26,1968247624297 | + | + | NA | NA | NA | NA | NA | + | + | NA | NA | + | 0,0432604151757038 | 7 | -527,074561654654 | 1069,01734036357 | 21,0557674482809 | 9.31484954867402e-07 |
| 2427 | 25,2372422324049 | NA | + | NA | + | + | + | + | NA | + | NA | NA | + | 0,0744804107322776 | 9 | -524,80306181931 | 1069,02344647327 | 21,0618735579746 | 9.28645417049148e-07 |
| 1346 | 26,2386559342402 | + | NA | NA | NA | NA | NA | + | NA | + | NA | + | NA | 0,0275621078339808 | 6 | -528,188893679473 | 1069,0239412051 | 21,0623682898099 | 9.28415730232923e-07 |
| 1109 | 25,9892337353139 | NA | NA | + | NA | + | NA | + | NA | NA | NA | + | NA | 0,0275381958951294 | 6 | -528,190577297467 | 1069,02730844109 | 21,0657355257963 | 9.26853947895676e-07 |
| 2077 | 25,5358999379587 | NA | NA | + | + | + | NA | NA | NA | NA | NA | NA | + | 0,0275090960050939 | 6 | -528,192626138592 | 1069,03140612334 | 21,0698332080467 | 9.2495691543284e-07 |
| 1087 | 25,1656756119637 | NA | + | + | + | + | + | NA | NA | NA | NA | + | NA | 0,0587988454547357 | 8 | -525,95343326015 | 1069,0318665203 | 21,0702936050086 | 9.24744016261437e-07 |
| 432 | 25,6883719857794 | + | + | + | + | NA | + | NA | + | + | NA | NA | NA | 0,0743771766722258 | 9 | -524,810698406227 | 1069,0387196471 | 21,0771467318098 | 9.2158074493038e-07 |
| 1192 | 25,5933849489479 | + | + | + | NA | NA | + | NA | + | NA | NA | + | NA | 0,0587502051734978 | 8 | -525,956971537972 | 1069,03894307594 | 21,0773701606543 | 9.21477796820408e-07 |
| 22 | 26,3256308008022 | + | NA | + | NA | + | NA | NA | NA | NA | NA | NA | NA | 0,0117895393485112 | 5 | -529,290533463971 | 1069,03908219512 | 21,0775092798278 | 9.21413701434874e-07 |
| 2322 | 26,3824659163009 | + | NA | NA | NA | + | NA | NA | NA | + | NA | NA | + | 0,0274400591074651 | 6 | -528,19748658657 | 1069,04112701929 | 21,0795541040025 | 9.20472118354098e-07 |
| 281 | 27,0534275039079 | NA | NA | NA | + | + | NA | NA | NA | + | NA | NA | NA | 0,0117716033581213 | 5 | -529,291776177594 | 1069,04156762236 | 21,0799947070725 | 9.20269359268368e-07 |
| 3330 | 26,4950428295612 | + | NA | NA | NA | NA | NA | NA | NA | + | NA | + | + | 0,0274039213212866 | 6 | -528,200030680276 | 1069,04621520671 | 21,0846422914155 | 9.18133327343955e-07 |
| 2182 | 25,8928051316997 | + | NA | + | NA | NA | NA | NA | + | NA | NA | NA | + | 0,0273825484186937 | 6 | -528,201535284344 | 1069,04922441484 | 21,0876514995516 | 9.1675293893406e-07 |
| 444 | 25,8104857134856 | + | + | NA | + | + | + | NA | + | + | NA | NA | NA | 0,0742805349063941 | 9 | -524,817846566474 | 1069,05301596759 | 21,0914430523032 | 9.15016626719827e-07 |
| 475 | 25,8677624430491 | NA | + | NA | + | + | NA | + | + | + | NA | NA | NA | 0,0586397222002202 | 8 | -525,965007807877 | 1069,05501561575 | 21,0934427004624 | 9.14102228258171e-07 |
| 1483 | 25,8706652601039 | NA | + | NA | + | NA | NA | + | + | + | NA | + | NA | 0,0585681165814768 | 8 | -525,970215727568 | 1069,06543145514 | 21,1038585398451 | 9.09354032107744e-07 |
| 1204 | 25,7171945036172 | + | + | NA | NA | + | + | NA | + | NA | NA | + | NA | 0,0585545305847752 | 8 | -525,971203800587 | 1069,06740760117 | 21,1058346858824 | 9.08455967673877e-07 |
| 2362 | 25,874748403591 | + | NA | NA | + | + | + | NA | NA | + | NA | NA | + | 0,0585512738683548 | 8 | -525,971440650699 | 1069,0678813014 | 21,1063083861084 | 9.08240825254565e-07 |
| 3205 | 26,1271751091273 | NA | NA | + | NA | NA | NA | NA | + | NA | NA | + | + | 0,0272426656354397 | 6 | -528,211381900446 | 1069,06891764704 | 21,1073447317542 | 9.07770321453919e-07 |
| 3385 | 25,8659493444497 | NA | NA | NA | + | + | + | NA | NA | + | NA | + | + | 0,058511429512154 | 8 | -525,974338331409 | 1069,07367666282 | 21,1121037475282 | 9.05612842701283e-07 |
| 277 | 27,1998320631329 | NA | NA | + | NA | + | NA | NA | NA | + | NA | NA | NA | 0,0115252484316439 | 5 | -529,308842852088 | 1069,07570097135 | 21,1141280560616 | 9.04696686522442e-07 |
| 3347 | 26,1233891517182 | NA | + | NA | NA | + | NA | NA | NA | + | NA | + | + | 0,0428521710635987 | 7 | -527,103771306 | 1069,07575966626 | 21,114186750973 | 9.04670136366135e-07 |
| 1289 | 26,9092790749229 | NA | NA | NA | + | NA | NA | NA | NA | + | NA | + | NA | 0,0115110057108816 | 5 | -529,309829411821 | 1069,07767409082 | 21,1161011755275 | 9.03804589326927e-07 |
| 389 | 27,2529613723666 | NA | NA | + | NA | NA | NA | NA | + | + | NA | NA | NA | 0,0114946026910015 | 5 | -529,310965592783 | 1069,07994645274 | 21,1183735374511 | 9.02778286902394e-07 |
| 1261 | 25,5604682128913 | NA | NA | + | + | NA | + | + | + | NA | NA | + | NA | 0,0584530319052877 | 8 | -525,97858507576 | 1069,08217015152 | 21,1205972362291 | 9.01775091220333e-07 |
| 1361 | 26,543591134622 | NA | NA | NA | NA | + | NA | + | NA | + | NA | + | NA | 0,0271282274476278 | 6 | -528,219436370406 | 1069,08502658697 | 21,1234536716752 | 9.00488079338199e-07 |
| 2451 | 26,2394793765148 | NA | + | NA | NA | + | NA | NA | + | + | NA | NA | + | 0,0427572003394084 | 7 | -527,11056462589 | 1069,08934630604 | 21,1277733907525 | 8.98545250444619e-07 |
| 3332 | 26,0120976871918 | + | + | NA | NA | NA | NA | NA | NA | + | NA | + | + | 0,0427387545239412 | 7 | -527,111883989368 | 1069,091985033 | 21,1304121177095 | 8.97360524371415e-07 |
| 3441 | 25,8873218040036 | NA | NA | NA | NA | + | + | + | NA | + | NA | + | + | 0,05838503260459 | 8 | -525,983529735106 | 1069,09205947021 | 21,1304865549216 | 8.97327126485086e-07 |
| 2418 | 25,8841340691351 | + | NA | NA | NA | + | + | + | NA | + | NA | NA | + | 0,0583788021370186 | 8 | -525,98398277397 | 1069,09296554794 | 21,1313926326497 | 8.96920694494703e-07 |
| 1273 | 25,6079457981077 | NA | NA | NA | + | + | + | + | + | NA | NA | + | NA | 0,0583701436706524 | 8 | -525,984612356056 | 1069,09422471211 | 21,1326517968214 | 8.96356187013591e-07 |
| 1037 | 26,2501178157186 | NA | NA | + | + | NA | NA | NA | NA | NA | NA | + | NA | 0,0113828886968002 | 5 | -529,318703137041 | 1069,09542154126 | 21,133848625966 | 8.95819954869857e-07 |
| 3370 | 25,7993767961081 | + | NA | NA | + | NA | + | NA | NA | + | NA | + | + | 0,0583131961388048 | 8 | -525,988753031321 | 1069,10250606264 | 21,1409331473512 | 8.92652340625955e-07 |
| 386 | 27,4067558111233 | + | NA | NA | NA | NA | NA | NA | + | + | NA | NA | NA | 0,0113246433142352 | 5 | -529,322736987116 | 1069,10348924141 | 21,1419163261173 | 8.92213630054117e-07 |
| 4062 | 25,3277347667791 | + | NA | + | + | + | NA | + | + | + | + | + | + | 0,120708370285366 | 12 | -521,294907186685 | 1069,10594340563 | 21,1443704903381 | 8.91119482111993e-07 |
| 1258 | 25,6881018116085 | + | NA | NA | + | NA | + | + | + | NA | NA | + | NA | 0,0582473815847117 | 8 | -525,993538119156 | 1069,11207623831 | 21,1505033230214 | 8.8839112404843e-07 |
| 238 | 25,5093463550733 | + | NA | + | + | NA | + | + | + | NA | NA | NA | NA | 0,0581761267946846 | 8 | -525,998718365884 | 1069,12243673177 | 21,1608638164773 | 8.83800938247212e-07 |
| 3426 | 25,8013758687265 | + | NA | NA | NA | NA | + | + | NA | + | NA | + | + | 0,0580919040142924 | 8 | -526,004840884646 | 1069,13468176929 | 21,1731088540014 | 8.78406381409964e-07 |
| 1386 | 25,7729751463805 | + | NA | NA | + | NA | + | + | NA | + | NA | + | NA | 0,0580915558933706 | 8 | -526,004866189928 | 1069,13473237986 | 21,173159464566 | 8.78384153369759e-07 |

| | | | | | | | | | | | | | | | | | | | |
|---|---|---|---|---|---|---|---|---|---|---|---|---|---|---|---|---|---|---|---|
| **271** | 26,1222093836701 | NA | + | + | + | NA | NA | NA | NA | + | NA | NA | NA | 0,0267530684537591 | 6 | -528,245834438122 | 1069,1378227224 | 21,1762498071075 | 8.770279474663e-07 |
| **253** | 25,4611986521143 | NA | NA | + | + | + | + | + | + | NA | NA | NA | NA | 0,058065712407771 | 8 | -526,006744754442 | 1069,13848950888 | 21,1769165935943 | 8.76735601010209e-07 |
| **3428** | 25,2182919820575 | + | + | NA | NA | NA | + | + | NA | + | NA | + | + | 0,0736646282828486 | 9 | -524,863384914514 | 1069,14409266367 | 21,182519748382 | 8.74282795836126e-07 |
| **250** | 25,5595670568688 | + | NA | NA | + | + | + | + | + | NA | NA | NA | NA | 0,0580259347507822 | 8 | -526,009636094039 | 1069,14427218808 | 21,1826992727881 | 8.74204321808327e-07 |
| **1100** | 25,2427536171817 | + | + | NA | + | NA | NA | + | NA | NA | NA | + | NA | 0,0423354204754256 | 7 | -527,14072668628 | 1069,14967042682 | 21,1880975115328 | 8.71847921521376e-07 |
| **2312** | 25,6188567646431 | + | + | + | NA | NA | NA | NA | NA | + | NA | NA | + | 0,0423147162631042 | 7 | -527,142206931174 | 1069,15263091661 | 21,1910580013214 | 8.70558327779677e-07 |
| **2197** | 25,9935747096428 | NA | NA | + | NA | + | NA | NA | + | NA | NA | NA | + | 0,026605904694699 | 6 | -528,256186840667 | 1069,15852752749 | 21,1969546121973 | 8.67995435878401e-07 |
| **460** | 25,7926237142198 | + | + | NA | + | NA | NA | + | + | + | NA | NA | NA | 0,0578749103264578 | 8 | -526,020612575105 | 1069,16622515021 | 21,2046522349187 | 8.64661105903314e-07 |
| **1285** | 26,9643689542507 | NA | NA | + | NA | NA | NA | NA | NA | + | NA | + | NA | 0,0108696903521151 | 5 | -529,354237099281 | 1069,16648946574 | 21,2049165504475 | 8.64546841775233e-07 |
| **1103** | 25,2098418008948 | NA | + | + | + | NA | NA | + | NA | NA | NA | + | NA | 0,0421783595962434 | 7 | -527,15195493389 | 1069,17212692204 | 21,2105540067521 | 8.62113350539681e-07 |
| **3345** | 26,5412107231189 | NA | NA | NA | NA | + | NA | NA | NA | + | NA | + | + | 0,0265079692405909 | 6 | -528,263075354987 | 1069,17230455613 | 21,2107316408371 | 8.62036783581841e-07 |
| **463** | 25,8307797503745 | NA | + | + | + | NA | NA | + | + | + | NA | NA | NA | 0,05781730593986 | 8 | -526,024798808463 | 1069,17459761693 | 21,2130247016355 | 8.61048998575226e-07 |
| **2393** | 25,8185454229531 | NA | NA | NA | + | + | NA | + | NA | + | NA | NA | + | 0,042140890771658 | 7 | -527,154633300008 | 1069,17748365428 | 21,2159107389884 | 8.59807384840283e-07 |
| **3127** | 25,2336443228025 | NA | + | + | NA | + | + | NA | NA | NA | NA | + | + | 0,0577878657479964 | 8 | -526,026938191073 | 1069,17887638215 | 21,2173034668558 | 8.59208854409255e-07 |
| **1291** | 26,3930957000536 | NA | + | NA | + | NA | NA | NA | NA | + | NA | + | NA | 0,0264319016568252 | 6 | -528,268425264659 | 1069,18300437547 | 21,2214314601808 | 8.57437279093043e-07 |
| **1096** | 25,2932943270848 | + | + | + | NA | NA | NA | + | NA | NA | NA | + | NA | 0,042079707514801 | 7 | -527,159006608469 | 1069,1862302712 | 21,22465735591 | 8.56055392220034e-07 |
| **3435** | 25,0813217042005 | NA | + | NA | + | NA | + | + | NA | + | NA | + | + | 0,073339689737059 | 9 | -524,887397734006 | 1069,19211830266 | 21,2305453873669 | 8.5353885785693e-07 |
| **2062** | 25,3510409672097 | + | NA | + | + | NA | NA | NA | NA | NA | NA | NA | + | 0,0263190738861753 | 6 | -528,276359785822 | 1069,1988734178 | 21,2373005025081 | 8.50660844290022e-07 |
| **3401** | 25,7595268574143 | NA | NA | NA | + | NA | NA | + | NA | + | NA | + | + | 0,0419552835421814 | 7 | -527,167899429008 | 1069,20401591228 | 21,2424429969899 | 8.48476394524927e-07 |
| **3124** | 25,4502414097323 | + | + | NA | NA | + | + | NA | NA | NA | NA | + | + | 0,0575778557258792 | 8 | -526,042197423597 | 1069,20939484719 | 21,2478219319034 | 8.4619751073954e-07 |
| **3301** | 25,5665290339999 | NA | NA | + | NA | NA | + | + | + | NA | NA | + | + | 0,0575492579874722 | 8 | -526,04427505931 | 1069,21355011862 | 21,2519772033299 | 8.44441245642063e-07 |
| **2378** | 25,719230070465 | + | NA | NA | + | NA | NA | + | NA | + | NA | NA | + | 0,041833457399164 | 7 | -527,176605459235 | 1069,22142797273 | 21,2598550574423 | 8.41121595394394e-07 |
| **2380** | 25,0055722755138 | + | + | NA | + | NA | NA | + | NA | + | NA | NA | + | 0,0574924454778064 | 8 | -526,048402321106 | 1069,22180464221 | 21,2602317269211 | 8.40963197894273e-07 |
| **3431** | 25,105008600965 | NA | + | + | NA | NA | + | + | NA | + | NA | + | + | 0,0731102700217257 | 9 | -524,904346684502 | 1069,22601620365 | 21,2644432883592 | 8.39194177045427e-07 |
| **3313** | 25,6263882389607 | NA | NA | NA | NA | + | + | + | + | NA | NA | + | + | 0,0574438384629269 | 8 | -526,051933280446 | 1069,22886656089 | 21,2672936456011 | 8.37999027298716e-07 |
| **1163** | 26,3601619310009 | NA | + | NA | + | NA | NA | NA | + | NA | NA | + | NA | 0,026099130920003 | 6 | -528,291824454383 | 1069,22980275492 | 21,2682298396298 | 8.37606854250652e-07 |
| **1030** | 26,2102661265554 | + | NA | + | NA | NA | NA | NA | NA | NA | NA | + | NA | 0,0104072466718377 | 5 | -529,386241016769 | 1069,23049730071 | 21,2689243854222 | 8.3731602659371e-07 |
| **1491** | 26,0251978912413 | NA | + | NA | NA | + | NA | + | + | + | NA | + | NA | 0,0573496562854158 | 8 | -526,058774438327 | 1069,24254887665 | 21,2809759613642 | 8.32285708787944e-07 |
| **1338** | 26,0137940191249 | + | NA | NA | + | + | + | NA | NA | + | NA | + | NA | 0,0573333217299381 | 8 | -526,059960870073 | 1069,24492174015 | 21,2833488248559 | 8.31298844140955e-07 |
| **2156** | 24,4684637633478 | + | + | NA | + | NA | + | + | NA | NA | NA | NA | + | 0,0573322696499735 | 8 | -526,060037285349 | 1069,2450745707 | 21,2835016554077 | 8.31235322637475e-07 |
| **3495** | 25,7971058228945 | NA | + | + | NA | NA | + | NA | + | + | NA | + | + | 0,0729744747047463 | 9 | -524,914376924938 | 1069,24607668452 | 21,2845037692305 | 8.30818930760802e-07 |
| **3156** | 25,1142234910274 | + | + | NA | NA | + | NA | + | NA | NA | NA | + | + | 0,0572459116832693 | 8 | -526,066309396101 | 1069,2576187922 | 21,2960458769105 | 8.26038038641968e-07 |
| **2376** | 25,0275812317417 | + | + | + | NA | NA | NA | + | NA | + | NA | NA | + | 0,0572255780914745 | 8 | -526,067786125247 | 1069,26057225049 | 21,2989993352041 | 8.24819104433277e-07 |
| **1080** | 25,4441047491852 | + | + | + | NA | + | + | NA | NA | NA | NA | + | NA | 0,0572014860502414 | 8 | -526,069535770957 | 1069,26407154191 | 21,3024986266239 | 8.23377224982707e-07 |
| **3119** | 24,8847474223881 | NA | + | + | + | NA | + | NA | NA | NA | NA | + | + | 0,057179047885272 | 8 | -526,071165266374 | 1069,26733053275 | 21,3057576174565 | 8.22036628113578e-07 |
| **2293** | 25,4512598793643 | NA | NA | + | NA | + | + | + | + | NA | NA | NA | + | 0,0571600500441432 | 8 | -526,072544889669 | 1069,27008977934 | 21,3085168640473 | 8.2090330918811e-07 |
| **90** | 26,03295821024 | + | NA | NA | + | + | NA | + | NA | NA | NA | NA | NA | 0,0258059796050489 | 6 | -528,312431134879 | 1069,27101611591 | 21,3094432006203 | 8.20523180847502e-07 |
| **268** | 26,1860238950438 | + | + | NA | + | NA | NA | NA | NA | + | NA | NA | NA | 0,0257727500718577 | 6 | -528,314766569454 | 1069,27568698506 | 21,3141140697717 | 8.18609138571029e-07 |
| **2278** | 25,4899595039941 | + | NA | + | NA | NA | + | + | + | NA | NA | NA | + | 0,0571138645173139 | 8 | -526,075898766921 | 1069,27679753384 | 21,3152246185523 | 8.18154712058087e-07 |
| **468** | 25,9403458589526 | + | + | NA | NA | + | NA | + | + | + | NA | NA | NA | 0,0570877735297208 | 8 | -526,077793356493 | 1069,28058671299 | 21,319013797696 | 8.16606112116323e-07 |
| **3307** | 24,9822231388157 | NA | + | NA | + | NA | + | + | + | NA | NA | + | + | 0,0727369522854213 | 9 | -524,931917497173 | 1069,28115782899 | 21,3195849137019 | 8.16372956997007e-07 |
| **78** | 25,5482857572409 | + | NA | + | + | NA | NA | + | NA | NA | NA | NA | NA | 0,0257140787676586 | 6 | -528,318889905522 | 1069,2839336572 | 21,3223607419066 | 8.15240687385463e-07 |
| **1476** | 25,9369465847946 | + | + | NA | NA | NA | NA | + | + | + | NA | + | NA | 0,0570629662405224 | 8 | -526,079594682079 | 1069,28418936416 | 21,3226164488674 | 8.15136462689086e-07 |
| **1394** | 26,0160528307284 | + | NA | NA | NA | + | + | + | NA | + | NA | + | NA | 0,0570514286074862 | 8 | -526,080432445249 | 1069,2858648905 | 21,3242919752074 | 8.14453857352841e-07 |
| **1094** | 25,6863248862461 | + | NA | + | NA | NA | NA | + | NA | NA | NA | + | NA | 0,0256750070146361 | 6 | -528,321635675249 | 1069,28942519665 | 21,3278522813603 | 8.13005294527388e-07 |
| **2152** | 24,547537893309 | + | + | + | NA | NA | + | + | NA | NA | NA | NA | + | 0,057010607217185 | 8 | -526,083396459465 | 1069,29179291893 | 21,3302200036387 | 8.1204337865297e-07 |
| **1101** | 25,732708432534 | NA | NA | + | + | NA | NA | + | NA | NA | NA | + | NA | 0,0256326397994085 | 6 | -528,324612909402 | 1069,29537966496 | 21,3338067496663 | 8.10588387032964e-07 |
| **471** | 25,949977785517 | NA | + | + | NA | + | NA | + | + | + | NA | NA | NA | 0,0569804339287357 | 8 | -526,085587239598 | 1069,2961744792 | 21,3346015639058 | 8.10266317437385e-07 |
| **3082** | 25,8440251391714 | + | NA | NA | + | NA | NA | NA | NA | NA | NA | + | + | 0,0256207428698825 | 6 | -528,325448908625 | 1069,2970516634 | 21,3354787481139 | 8.09911018949633e-07 |
| **2383** | 24,9390821077227 | NA | + | + | + | NA | NA | + | NA | + | NA | NA | + | 0,0569638053694379 | 8 | -526,086794552943 | 1069,29858910589 | 21,3370161905946 | 8.092886623866e-07 |
| **3097** | 25,8444366538907 | NA | NA | NA | + | + | NA | NA | NA | NA | NA | + | + | 0,0256069007776262 | 6 | -528,326421581918 | 1069,29899700999 | 21,3374240946985 | 8.09123623133907e-07 |
| **1113** | 26,0881631534469 | NA | NA | NA | + | + | NA | + | NA | NA | NA | + | NA | 0,0256020825058041 | 6 | -528,326760154972 | 1069,2996741561 | 21,3381012408081 | 8.08849722047477e-07 |
| **279** | 26,7654151906938 | NA | + | + | NA | + | NA | NA | NA | + | NA | NA | NA | 0,0254971831475852 | 6 | -528,334130867739 | 1069,31441558163 | 21,3528426663415 | 8.02909840546539e-07 |
| **3078** | 25,7349576596156 | + | NA | + | NA | NA | NA | NA | NA | NA | NA | + | + | 0,0254731961745219 | 6 | -528,335816191702 | 1069,31778622956 | 21,3562133142668 | 8.015578169713e-07 |
| **2290** | 25,5526737896418 | + | NA | NA | NA | + | + | + | + | NA | NA | NA | + | 0,0568226870801477 | 8 | -526,097039562688 | 1069,31907912538 | 21,3575062100854 | 8.01039819043733e-07 |
| **1072** | 25,1227581745774 | + | + | + | + | NA | + | NA | NA | NA | NA | + | NA | 0,0568202409104852 | 8 | -526,097217138009 | 1069,31943427602 | 21,3578613607278 | 8.00897586769392e-07 |
| **3300** | 25,1542060382773 | + | + | NA | NA | NA | + | + | + | NA | NA | + | + | 0,0724735776032271 | 9 | -524,951361959105 | 1069,32004675286 | 21,358473837565 | 8.00652358710001e-07 |
| **3298** | 25,7375318297211 | + | NA | NA | NA | NA | + | + | + | NA | NA | + | + | 0,0568127919390567 | 8 | -526,097757879961 | 1069,32051575992 | 21,3589428446323 | 8.00464624915639e-07 |
| **264** | 26,1820113816598 | + | + | + | NA | NA | NA | NA | NA | + | NA | NA | NA | 0,0253588561760809 | 6 | -528,343849146494 | 1069,33385213914 | 21,3722792238507 | 7.95144731761689e-07 |
| **1479** | 25,9899599306898 | NA | + | + | NA | NA | NA | + | + | + | NA | + | NA | 0,0566983264469043 | 8 | -526,106066717786 | 1069,33713343557 | 21,3755605202823 | 7.93841248547302e-07 |
| **26** | 26,4917532274818 | + | NA | NA | + | + | NA | NA | NA | NA | NA | NA | NA | 0,0095588825927923 | 5 | -529,444914126434 | 1069,34784352004 | 21,3862706047537 | 7.89601557127819e-07 |
| **3093** | 25,8981477710261 | NA | NA | + | NA | + | NA | NA | NA | NA | NA | + | + | 0,0252428340514586 | 6 | -528,351999316238 | 1069,35015247863 | 21,3885795633398 | 7.88690504477243e-07 |
| **2445** | 26,179620794653 | NA | NA | + | + | NA | NA | NA | + | + | NA | NA | + | 0,0409149328808899 | 7 | -527,242210142883 | 1069,35263734003 | 21,391064424738 | 7.8771121965512e-07 |
| **3112** | 25,1684425853154 | + | + | + | NA | NA | + | NA | NA | NA | NA | + | + | 0,0565825018545868 | 8 | -526,114473184441 | 1069,35394636888 | 21,3923734535906 | 7.87195819984766e-07 |
| **1343** | 25,5500917858866 | NA | + | + | + | + | + | NA | NA | + | NA | + | NA | 0,0721755527624541 | 9 | -524,973357923193 | 1069,36403868103 | 21,4024657657417 | 7.8323351263368e-07 |
| **2386** | 25,9248207149011 | + | NA | NA | NA | + | NA | + | NA | + | NA | NA | + | 0,0408090483533598 | 7 | -527,249768800635 | 1069,36775465553 | 21,4061817402433 | 7.81779625822646e-07 |
| **2231** | 25,3026962057417 | NA | + | + | NA | + | + | NA | + | NA | NA | NA | + | 0,0564421686269629 | 8 | -526,124657088427 | 1069,37431417685 | 21,4127412615626 | 7.7921977590353e-07 |
| **3409** | 25,9565376887187 | NA | NA | NA | NA | + | NA | + | NA | + | NA | + | + | 0,0407611467850962 | 7 | -527,253188021149 | 1069,37459309656 | 21,4130201812713 | 7.79111113604242e-07 |
| **1260** | 25,192407707926 | + | + | NA | + | NA | + | + | + | NA | NA | + | NA | 0,0720863314992379 | 9 | -524,979941597048 | 1069,37720602874 | 21,4156331134513 | 7.78093895978829e-07 |
| **3394** | 25,8802021687839 | + | NA | NA | NA | NA | NA | + | NA | + | NA | + | + | 0,0407192254907381 | 7 | -527,256180228857 | 1069,38057751198 | 21,4190045966864 | 7.7678333565944e-07 |
| **64** | 25,1123618629502 | + | + | + | + | + | + | NA | NA | NA | NA | NA | NA | 0,0563972668918755 | 8 | -526,127915262579 | 1069,38083052516 | 21,4192576098665 | 7.76685073663965e-07 |
| **456** | 25,8662051469862 | + | + | + | NA | NA | NA | + | + | + | NA | NA | NA | 0,0563716024373325 | 8 | -526,129777465388 | 1069,38455493078 | 21,4229820154844 | 7.75240074396523e-07 |
| **1388** | 25,2790518553253 | + | + | NA | + | NA | + | + | NA | + | NA | + | NA | 0,0719813050976102 | 9 | -524,987690727843 | 1069,39270429033 | 21,4311313750406 | 7.72087646271413e-07 |
| **2487** | 25,7582159234998 | NA | + | + | NA | + | + | NA | + | + | NA | NA | + | 0,0719442796065883 | 9 | -524,990422359315 | 1069,39816755328 | 21,4365946379844 | 7.69981465322555e-07 |
| **2108** | 25,1054460779923 | + | + | NA | + | + | + | NA | NA | NA | NA | NA | + | 0,0562672487273146 | 8 | -526,137348808505 | 1069,39969761701 | 21,4381247017195 | 7.69392630231677e-07 |
| **2324** | 26,0356639431365 | + | + | NA | NA | + | NA | NA | NA | + | NA | NA | + | 0,0405501754625058 | 7 | -527,268245152269 | 1069,4047073588 | 21,4431344435111 | 7.67467812740849e-07 |

| | | | | | | | | | | | | | | | | | | |
|---|---|---|---|---|---|---|---|---|---|---|---|---|---|---|---|---|---|---|
| **3507** | 25,9504870889569 | NA | + | NA | NA | + | + | NA | + | + | NA | + | + | 0,0718877785413665 | 9 | -524,994590631244 | 1069,40650409713 | 21,4449311818439 | 7.66778652927986e-07 |
| **3492** | 25,9538319114722 | + | + | NA | NA | NA | + | NA | + | + | NA | + | + | 0,0718855730373328 | 9 | -524,994753333486 | 1069,40682950162 | 21,4452565863262 | 7.66653906471235e-07 |
| **1106** | 26,109250244845 | + | NA | NA | NA | + | NA | + | NA | NA | NA | + | NA | 0,0248323625235476 | 6 | -528,380825799138 | 1069,40780544443 | 21,4462325291397 | 7.66279892547471e-07 |
| **1269** | 25,6830082374671 | NA | NA | + | NA | + | + | + | + | NA | NA | + | NA | 0,0562099048904934 | 8 | -526,141509012174 | 1069,40801802435 | 21,4464451090573 | 7.66198449017648e-07 |
| **2367** | 25,3831101043539 | NA | + | + | + | + | + | NA | NA | + | NA | NA | + | 0,071872665054002 | 9 | -524,995705560592 | 1069,40873395583 | 21,4471610405399 | 7.65924225306159e-07 |
| **1254** | 25,6682145380803 | + | NA | + | NA | NA | + | + | + | NA | NA | + | NA | 0,0561992440768986 | 8 | -526,142282409302 | 1069,4095648186 | 21,4479919033133 | 7.65606102426802e-07 |
| **1034** | 26,4132427166767 | + | NA | NA | + | NA | NA | NA | NA | NA | NA | + | NA | 0,0091103111954747 | 5 | -529,475917156814 | 1069,4098495808 | 21,4482766655124 | 7.65497102348097e-07 |
| **3217** | 26,3860554141857 | NA | NA | NA | NA | + | NA | NA | + | NA | NA | + | + | 0,0248014800531072 | 6 | -528,382994114281 | 1069,41214207472 | 21,450569159425 | 7.64620156318853e-07 |
| **2070** | 25,7003971082016 | + | NA | + | NA | + | NA | NA | NA | NA | NA | NA | + | 0,0247175894640962 | 6 | -528,388883880645 | 1069,42392160744 | 21,4623486921541 | 7.60129958329186e-07 |
| **388** | 26,8218118686869 | + | + | NA | NA | NA | NA | NA | + | + | NA | NA | NA | 0,0247122671289548 | 6 | -528,389257532539 | 1069,42466891123 | 21,4630959959402 | 7.59845987387753e-07 |
| **1282** | 27,0462845206758 | + | NA | NA | NA | NA | NA | NA | NA | + | NA | + | NA | 0,0089962766592879 | 5 | -529,483796421935 | 1069,42560811105 | 21,4640351957555 | 7.59489247551293e-07 |
| **2194** | 26,1428977349816 | + | NA | NA | NA | + | NA | NA | + | NA | NA | NA | + | 0,0246816022905879 | 6 | -528,391410302574 | 1069,4289744513 | 21,4674015360106 | 7.58211973174218e-07 |
| **155** | 26,2643819794567 | NA | + | NA | + | + | NA | NA | + | NA | NA | NA | NA | 0,0246766739797681 | 6 | -528,391756279494 | 1069,42966640514 | 21,468093489852 | 7.57949694704267e-07 |
| **2216** | 25,3013912008607 | + | + | + | NA | NA | + | NA | + | NA | NA | NA | + | 0,0560366241925884 | 8 | -526,15407871466 | 1069,43315742932 | 21,4715845140286 | 7.56627838343593e-07 |
| **2228** | 25,4548905376059 | + | + | NA | NA | + | + | NA | + | NA | NA | NA | + | 0,0560309193342809 | 8 | -526,154492503253 | 1069,43398500651 | 21,4724120912156 | 7.56314819140912e-07 |
| **3303** | 25,1422559531105 | NA | + | + | NA | NA | + | + | + | NA | NA | + | + | 0,0716837747869594 | 9 | -525,009638557399 | 1069,43659994944 | 21,4750270341538 | 7.55326605264737e-07 |
| **3387** | 25,5284440099018 | NA | + | NA | + | + | + | NA | NA | + | NA | + | + | 0,0716594010676744 | 9 | -525,011436214672 | 1069,44019526399 | 21,4786223486988 | 7.5397000661471e-07 |
| **274** | 27,1271464815062 | + | NA | NA | NA | + | NA | NA | NA | + | NA | NA | NA | 0,0088479084869023 | 5 | -529,494046627833 | 1069,44610852284 | 21,4845356075509 | 7.51744088935144e-07 |
| **246** | 25,5918601870968 | + | NA | + | NA | + | + | + | + | NA | NA | NA | NA | 0,0559344928018325 | 8 | -526,161486198759 | 1069,44797239752 | 21,4863994822272 | 7.51043836896144e-07 |
| **184** | 25,4359652536907 | + | + | + | NA | + | + | NA | + | NA | NA | NA | NA | 0,0559232779127483 | 8 | -526,162299554253 | 1069,44959910851 | 21,4880261932155 | 7.50433219623148e-07 |
| **112** | 24,6215600153491 | + | + | + | + | NA | + | + | NA | NA | NA | NA | NA | 0,0559006091760832 | 8 | -526,163943566125 | 1069,45288713225 | 21,4913142169596 | 7.4920051207149e-07 |
| **140** | 26,1159043009077 | + | + | NA | + | NA | NA | NA | + | NA | NA | NA | NA | 0,0245096172691787 | 6 | -528,40348294921 | 1069,45311974457 | 21,4915468292829 | 7.49113380502687e-07 |
| **2472** | 25,7136377780242 | + | + | + | NA | NA | + | NA | + | + | NA | NA | + | 0,0715695321744464 | 9 | -525,018063989957 | 1069,45345081456 | 21,491877899268 | 7.48989386287791e-07 |
| **3315** | 25,1048597458813 | NA | + | NA | NA | + | + | + | + | NA | NA | + | + | 0,0715674239927232 | 9 | -525,018219459357 | 1069,45376175336 | 21,4921888380684 | 7.48872950408514e-07 |
| **143** | 26,0853371700126 | NA | + | + | + | NA | NA | NA | + | NA | NA | NA | NA | 0,0244927987549674 | 6 | -528,40466342607 | 1069,45548069829 | 21,493907783004 | 7.4822959124009e-07 |
| **3163** | 24,9480789788871 | NA | + | NA | + | + | NA | + | NA | NA | NA | + | + | 0,0558650237518374 | 8 | -526,166524259384 | 1069,45804851877 | 21,4964756034765 | 7.47269548045373e-07 |
| **320** | 25,5382444221074 | + | + | + | + | + | + | NA | NA | + | NA | NA | NA | 0,071534337269682 | 9 | -525,020659417676 | 1069,45864167 | 21,497068754707 | 7.47047958979973e-07 |
| **1391** | 25,186561706976 | NA | + | + | + | NA | + | + | NA | + | NA | + | NA | 0,0715227436695004 | 9 | -525,021514359426 | 1069,4603515535 | 21,4987786382064 | 7.46409549431181e-07 |
| **1463** | 25,9455094243204 | NA | + | + | NA | + | + | NA | + | + | NA | + | NA | 0,0714893408094606 | 9 | -525,023977512158 | 1069,46527785896 | 21,5037049436705 | 7.44573291130997e-07 |
| **3159** | 24,9667876538351 | NA | + | + | NA | + | NA | + | NA | NA | NA | + | + | 0,0557638853692084 | 8 | -526,173858392922 | 1069,47271678584 | 21,5111438705537 | 7.41809021980903e-07 |
| **2074** | 25,7580354463422 | + | NA | NA | + | + | NA | NA | NA | NA | NA | NA | + | 0,0243484307168041 | 6 | -528,414795656191 | 1069,47574515854 | 21,5141722432454 | 7.40686634854674e-07 |
| **1098** | 25,8602788376736 | + | NA | NA | + | NA | NA | + | NA | NA | NA | + | NA | 0,0243310161698002 | 6 | -528,416017765856 | 1069,47818937787 | 21,5166164625748 | 7.39781987461463e-07 |
| **1263** | 25,1942364750782 | NA | + | + | + | NA | + | + | + | NA | NA | + | NA | 0,0713949719410166 | 9 | -525,030935866953 | 1069,47919456855 | 21,5176216532604 | 7.39410269899443e-07 |
| **1266** | 25,7770437819991 | + | NA | NA | NA | + | + | + | + | NA | NA | + | NA | 0,0556999890237792 | 8 | -526,178491484578 | 1069,48198296916 | 21,5204100538656 | 7.38380102174372e-07 |
| **3531** | 25,4282401654008 | NA | + | NA | + | NA | NA | + | + | + | NA | + | + | 0,0713381137710812 | 9 | -525,035128002835 | 1069,48757884032 | 21,5260059250247 | 7.36317049705083e-07 |
| **1275** | 25,1512009142568 | NA | + | NA | + | + | + | + | + | NA | NA | + | NA | 0,0713167150863 | 9 | -525,036705655026 | 1069,4907341447 | 21,5291612294075 | 7.35156313357946e-07 |
| **1384** | 25,3104432980858 | + | + | + | NA | NA | + | + | NA | + | NA | + | NA | 0,0712700058579845 | 9 | -525,040149241225 | 1069,4976213171 | 21,5360484018045 | 7.32629093071351e-07 |
| **31** | 25,9353338887537 | NA | + | + | + | + | NA | NA | NA | NA | NA | NA | NA | 0,0241803381775528 | 6 | -528,426591062798 | 1069,49933597175 | 21,5377630564599 | 7.32001259296768e-07 |
| **2412** | 25,0009984280169 | + | + | NA | + | NA | + | + | NA | + | NA | NA | + | 0,0712123981734022 | 9 | -525,044396065658 | 1069,50611496596 | 21,5445420506717 | 7.29524343268099e-07 |
| **1448** | 25,899036321435 | + | + | + | NA | NA | + | NA | + | + | NA | + | NA | 0,0711757668476326 | 9 | -525,047096381111 | 1069,51151559687 | 21,5499426815775 | 7.27557054755898e-07 |
| **1049** | 26,7034797151489 | NA | NA | NA | + | + | NA | NA | NA | NA | NA | + | NA | 0,0083124453812112 | 5 | -529,53102703319 | 1069,52006933355 | 21,5584964182642 | 7.24452033636667e-07 |
| **1340** | 25,7265358271744 | + | + | NA | + | + | + | NA | NA | + | NA | + | NA | 0,0710520077018378 | 9 | -525,056218625219 | 1069,52976008508 | 21,5681871697939 | 7.2095028181289e-07 |
| **1051** | 26,2439746946051 | NA | + | NA | + | + | NA | NA | NA | NA | NA | + | NA | 0,0239476268016638 | 6 | -528,442917557232 | 1069,53198896062 | 21,5704160453272 | 7.20147275124553e-07 |
| **3202** | 26,4403342089635 | + | NA | NA | NA | NA | NA | NA | + | NA | NA | + | + | 0,0239336756017459 | 6 | -528,443896217678 | 1069,53394628151 | 21,5723733662187 | 7.19442840228781e-07 |
| **3099** | 25,4482174791242 | NA | + | NA | + | + | NA | NA | NA | NA | NA | + | + | 0,0396400208750302 | 7 | -527,333165438034 | 1069,53454793033 | 21,5729750150417 | 7.19226446809596e-07 |
| **2111** | 24,9191427476494 | NA | + | + | + | + | + | NA | NA | NA | NA | NA | + | 0,0552925116645369 | 8 | -526,208030092676 | 1069,54106018535 | 21,5794872700621 | 7.16888362406309e-07 |
| **3383** | 25,6198509562164 | NA | + | + | NA | + | + | NA | NA | + | NA | + | + | 0,0709712088346929 | 9 | -525,062173626553 | 1069,54167008775 | 21,58009717246 | 7.16669779770909e-07 |
| **3378** | 26,0992908457328 | + | NA | NA | NA | + | + | NA | NA | + | NA | + | + | 0,0551988857432849 | 8 | -526,214815368449 | 1069,5546307369 | 21,5930578216069 | 7.12040542665746e-07 |
| **2188** | 25,482282556779 | + | + | NA | + | NA | NA | NA | + | NA | NA | NA | + | 0,0394917866292819 | 7 | -527,343732993239 | 1069,55568304074 | 21,59411012545 | 7.11665999707871e-07 |
| **2191** | 25,3853004907961 | NA | + | + | + | NA | NA | NA | + | NA | NA | NA | + | 0,0394856381753441 | 7 | -527,344171278647 | 1069,55655961156 | 21,5949866962678 | 7.11354155227621e-07 |
| **440** | 25,8532259469235 | + | + | + | NA | + | + | NA | + | + | NA | NA | NA | 0,0708694777801877 | 9 | -525,069670626265 | 1069,55666408718 | 21,5950911718858 | 7.11316996615699e-07 |
| **3084** | 25,3733982333969 | + | + | NA | + | NA | NA | NA | NA | NA | NA | + | + | 0,0394817800239819 | 7 | -527,344446301061 | 1069,55710965639 | 21,5955367410954 | 7.11158543790779e-07 |
| **149** | 26,9275305178558 | NA | NA | + | NA | + | NA | NA | + | NA | NA | NA | NA | 0,0079814768872277 | 5 | -529,553874555774 | 1069,56576437872 | 21,6041914634327 | 7.0808775292619e-07 |
| **3372** | 25,3922576267841 | + | + | NA | + | NA | + | NA | NA | + | NA | + | + | 0,0707659131593485 | 9 | -525,077301906647 | 1069,57192664794 | 21,6103537326499 | 7.05909396872184e-07 |
| **2364** | 25,5412429851685 | + | + | NA | + | + | + | NA | NA | + | NA | NA | + | 0,0707418821228929 | 9 | -525,079072540197 | 1069,57546791504 | 21,6138949997494 | 7.0466059592168e-07 |
| **2408** | 25,0295069092168 | + | + | + | NA | NA | + | + | NA | + | NA | NA | + | 0,0707409523773525 | 9 | -525,079141043965 | 1069,57560492258 | 21,6140320072855 | 7.0461232566904e-07 |
| **3245** | 25,5429521799859 | NA | NA | + | + | NA | + | NA | + | NA | NA | + | + | 0,055043654855414 | 8 | -526,226063810439 | 1069,57712762088 | 21,6155547055878 | 7.04076073836336e-07 |
| **2203** | 25,4751272424139 | NA | + | NA | + | + | NA | NA | + | NA | NA | NA | + | 0,0393019536592239 | 7 | -527,357263723217 | 1069,5827445007 | 21,6211715854079 | 7.02101492535184e-07 |
| **2159** | 24,3831284662811 | NA | + | + | + | NA | + | + | NA | NA | NA | NA | + | 0,0549860850019039 | 8 | -526,230235005269 | 1069,58547001054 | 21,6238970952465 | 7.01145351912993e-07 |
| **391** | 26,7880762069141 | NA | + | + | NA | NA | NA | NA | + | + | NA | NA | NA | 0,0235566759607597 | 6 | -528,470337011348 | 1069,58682786885 | 21,6252549535591 | 7.00669485448887e-07 |
| **2457** | 26,2948307984417 | NA | NA | NA | + | + | NA | NA | + | + | NA | NA | + | 0,039221590517933 | 7 | -527,362990963991 | 1069,59419898225 | 21,6326260669555 | 6.98091881204563e-07 |
| **146** | 26,9250294464076 | + | NA | NA | NA | + | NA | NA | + | NA | NA | NA | NA | 0,0077680896205334 | 5 | -529,568601134116 | 1069,59521753541 | 21,6336446201165 | 6.97736449872546e-07 |
| **3375** | 25,2464428400252 | NA | + | + | + | NA | + | NA | NA | + | NA | + | + | 0,0705699419235551 | 9 | -525,091739950649 | 1069,60080273594 | 21,6392298206531 | 6.95790669033993e-07 |
| **3087** | 25,2203630731149 | NA | + | + | + | NA | NA | NA | NA | NA | NA | + | + | 0,0391659604804643 | 7 | -527,366955269965 | 1069,60212759419 | 21,6405546789024 | 6.95329909657232e-07 |
| **1268** | 25,3434116796187 | + | + | NA | NA | + | + | + | + | NA | NA | + | NA | 0,0705411404426708 | 9 | -525,093861623182 | 1069,60504608101 | 21,6434731657196 | 6.9431599402454e-07 |
| **1157** | 26,8908045482932 | NA | NA | + | NA | NA | NA | NA | + | NA | NA | + | NA | 0,0076802122507767 | 5 | -529,574664928908 | 1069,60734512499 | 21,6457722097018 | 6.93518321079753e-07 |
| **3148** | 24,8170399853226 | + | + | NA | + | NA | NA | + | NA | NA | NA | + | + | 0,0548112249071726 | 8 | -526,242902848482 | 1069,61080569696 | 21,6492327816734 | 6.92319373608033e-07 |
| **3257** | 25,608885681352 | NA | NA | NA | + | + | + | NA | + | NA | NA | + | + | 0,0547982950717791 | 8 | -526,24383946506 | 1069,61267893012 | 21,6511060148287 | 6.91671239379708e-07 |
| **276** | 26,7332653234427 | + | + | NA | NA | + | NA | NA | NA | + | NA | NA | NA | 0,0233608926670381 | 6 | -528,484064207806 | 1069,61428226177 | 21,6527093464751 | 6.91116972384313e-07 |
| **2415** | 24,9161081966565 | NA | + | + | + | NA | + | + | NA | + | NA | NA | + | 0,0704753011840671 | 9 | -525,098711451528 | 1069,6147457377 | 21,6531728224111 | 6.90956832897422e-07 |
| **1045** | 26,7845795742768 | NA | NA | + | NA | + | NA | NA | NA | NA | NA | + | NA | 0,0076253565999604 | 5 | -529,57844985669 | 1069,61491498056 | 21,6533420652654 | 6.90898365617941e-07 |
| **454** | 26,4086469024536 | + | NA | + | NA | NA | NA | + | + | + | NA | NA | NA | 0,0389930333968198 | 7 | -527,379276929491 | 1069,62677091325 | 21,6651979979558 | 6.86814858762331e-07 |
| **458** | 26,5611802100914 | + | NA | NA | + | NA | NA | + | + | + | NA | NA | NA | 0,0389723686083411 | 7 | -527,380749219702 | 1069,62971549367 | 21,668142578377 | 6.85804411987262e-07 |
| **2484** | 25,906823013665 | + | + | NA | NA | + | + | NA | + | + | NA | NA | + | 0,0703687396510817 | 9 | -525,106560222206 | 1069,63044327906 | 21,668870363766 | 6.85554898172758e-07 |
| **1256** | 25,3647349812935 | + | + | + | NA | NA | + | + | + | NA | NA | + | NA | 0,0703450368443906 | 9 | -525,108305925939 | 1069,63393468652 | 21,6723617712325 | 6.84359166428011e-07 |
| **461** | 26,4958118935486 | NA | NA | + | + | NA | NA | + | + | + | NA | NA | NA | 0,0389375445889397 | 7 | -527,383230231469 | 1069,6346775172 | 21,673104601912 | 6.84105032133334e-07 |

| | | | | | | | | | | | | | | | | | |
|---|---|---|---|---|---|---|---|---|---|---|---|---|---|---|---|---|---|
| 2541 | 25,8023565817306 | NA | NA | + | + | NA | + | + | + | + | NA | NA | + | 0,0703336509038499 | 9 | -525,109144480809 | 1069,63561179626 | 21,6740388809721 | 6.83785534260946e-07 |
| 3207 | 25,7844597794903 | NA | + | + | NA | NA | NA | NA | + | NA | NA | + | + | 0,0388672967340379 | 7 | -527,388234715017 | 1069,6446864843 | 21,6831135690065 | 6.80689992138573e-07 |
| 1154 | 27,016228888051 | + | NA | NA | NA | NA | NA | NA | + | NA | NA | + | NA | 0,0073469655084105 | 5 | -529,597655049802 | 1069,65332536678 | 21,6917524514897 | 6.77756132592309e-07 |
| 3242 | 25,7417849316994 | + | NA | NA | + | NA | + | NA | + | NA | NA | + | + | 0,0544629314547949 | 8 | -526,268128195878 | 1069,66125639176 | 21,6996834764657 | 6.750738041033e-07 |
| 3144 | 24,913807905291 | + | + | + | NA | NA | NA | + | NA | NA | NA | + | + | 0,0544247302729699 | 8 | -526,270894372541 | 1069,66678874508 | 21,7052158297906 | 6.73209011063509e-07 |
| 1243 | 25,5813907843709 | NA | + | NA | + | + | NA | + | + | NA | NA | + | NA | 0,0544181974702609 | 8 | -526,271367406574 | 1069,66773481315 | 21,7061618978582 | 6.72890635596566e-07 |
| 2104 | 25,1816594353831 | + | + | + | NA | + | + | NA | NA | NA | NA | NA | + | 0,0543983345424848 | 8 | -526,272805642227 | 1069,67061128445 | 21,7090383691641 | 6.71923555904878e-07 |
| 1460 | 26,0917968838302 | + | + | NA | NA | + | + | NA | + | + | NA | + | NA | 0,0700175309309956 | 9 | -525,132422073183 | 1069,68216698101 | 21,7205940657218 | 6.68052477564717e-07 |
| 1271 | 25,2925463848809 | NA | + | + | NA | + | + | + | + | NA | NA | + | NA | 0,0700164732052123 | 9 | -525,132499945874 | 1069,68232272639 | 21,7207498111022 | 6.6800045654662e-07 |
| 2284 | 24,8706274716737 | + | + | NA | + | NA | + | + | + | NA | NA | NA | + | 0,0699359258577627 | 9 | -525,138429803762 | 1069,69418244217 | 21,7326095268797 | 6.64051030111295e-07 |
| 1287 | 26,6178130461668 | NA | + | + | NA | NA | NA | NA | NA | + | NA | + | NA | 0,0227726853317734 | 6 | -528,525289377134 | 1069,69673260042 | 21,7351596851313 | 6.63204852090961e-07 |
| 4064 | 25,052298506597 | + | + | + | + | + | NA | + | + | + | + | + | + | 0,132497738447317 | 13 | -520,370721228589 | 1069,70079205067 | 21,7392191353836 | 6.61860093745474e-07 |
| 2553 | 25,8449062933807 | NA | NA | NA | + | + | + | + | + | + | NA | NA | + | 0,0698566035650457 | 9 | -525,14426897206 | 1069,70586077877 | 21,7442878634752 | 6.60184823090082e-07 |
| 1231 | 25,6155549129279 | NA | + | + | + | NA | NA | + | + | NA | NA | + | NA | 0,054140775780835 | 8 | -526,291452233203 | 1069,70790446641 | 21,7463315511145 | 6.59510561862271e-07 |
| 2360 | 25,5887455951288 | + | + | + | NA | + | + | NA | NA | + | NA | NA | + | 0,0698417888616858 | 9 | -525,145359474714 | 1069,70804178407 | 21,7464688687837 | 6.59465282190125e-07 |
| 3524 | 25,5489789839822 | + | + | NA | NA | NA | NA | + | + | + | NA | + | + | 0,0698401097603912 | 9 | -525,145483071397 | 1069,70828897744 | 21,7467160621493 | 6.59383779505669e-07 |
| 1284 | 26,614595612394 | + | + | NA | NA | NA | NA | NA | NA | + | NA | + | NA | 0,0226647181082505 | 6 | -528,532853686587 | 1069,71186121933 | 21,7502883040377 | 6.58207091493213e-07 |
| 2222 | 25,4525350361759 | + | NA | + | + | NA | + | NA | + | NA | NA | NA | + | 0,0540905038603548 | 8 | -526,295091198783 | 1069,71518239757 | 21,753609482276 | 6.57114986978309e-07 |
| 1336 | 25,7564081172775 | + | + | + | NA | + | + | NA | NA | + | NA | + | NA | 0,0697914192542256 | 9 | -525,149067025626 | 1069,7154568859 | 21,7538839706069 | 6.57024807968734e-07 |
| 2508 | 25,3764560596273 | + | + | NA | + | NA | NA | + | + | + | NA | NA | + | 0,0697460120998644 | 9 | -525,152409133743 | 1069,72214110213 | 21,7605681868415 | 6.54832625320967e-07 |
| 3527 | 25,5353415122564 | NA | + | + | NA | NA | NA | + | + | + | NA | + | + | 0,0697163421485512 | 9 | -525,154592846575 | 1069,7265085278 | 21,7649356125053 | 6.53404219096447e-07 |
| 3539 | 25,53180578632 | NA | + | NA | NA | + | NA | + | + | + | NA | + | + | 0,0697151916978191 | 9 | -525,154677518518 | 1069,72667787168 | 21,7651049563915 | 6.53348896433749e-07 |
| 1198 | 25,671678805444 | + | NA | + | + | NA | + | NA | + | NA | NA | + | NA | 0,0539679429784697 | 8 | -526,303962037677 | 1069,73292407535 | 21,7713511600632 | 6.51311604287204e-07 |
| 1481 | 26,5661148632393 | NA | NA | NA | + | NA | NA | + | + | + | NA | + | NA | 0,0382279234805629 | 7 | -527,433767105532 | 1069,73575126533 | 21,7741783500376 | 6.50391563903731e-07 |
| 1213 | 25,6434906833223 | NA | NA | + | + | + | + | NA | + | NA | NA | + | NA | 0,0539463816935054 | 8 | -526,305522503951 | 1069,7360450079 | 21,7744720926116 | 6.50296047072179e-07 |
| 2237 | 25,4014851503301 | NA | NA | + | + | + | + | NA | + | NA | NA | NA | + | 0,053866103796398 | 8 | -526,311332186082 | 1069,74766437216 | 21,7860914568741 | 6.46528987053721e-07 |
| 368 | 25,1002635863827 | + | + | + | + | NA | + | + | NA | + | NA | NA | NA | 0,0695420572729504 | 9 | -525,167418834046 | 1069,75216050274 | 21,7905875874478 | 6.45077180170749e-07 |
| 1228 | 25,552480991687 | + | + | NA | + | NA | NA | + | + | NA | NA | + | NA | 0,0538079081945909 | 8 | -526,315543472509 | 1069,75608694502 | 21,7945140297277 | 6.43811993340877e-07 |
| 80 | 24,9912646672634 | + | + | + | + | NA | NA | + | NA | NA | NA | NA | NA | 0,0380568770266961 | 7 | -527,445942895073 | 1069,76010284441 | 21,7985299291192 | 6.42520548257e-07 |
| 2523 | 25,3853463331775 | NA | + | NA | + | + | NA | + | + | + | NA | NA | + | 0,0694860260776899 | 9 | -525,171541776465 | 1069,76040638758 | 21,798833472285 | 6.42423039296101e-07 |
| 252 | 25,0914979016233 | + | + | NA | + | + | + | + | + | NA | NA | NA | NA | 0,0694569096376709 | 9 | -525,173684152962 | 1069,76469114057 | 21,8031182252789 | 6.41048200512595e-07 |
| 2511 | 25,3765977359669 | NA | + | + | + | NA | NA | + | + | + | NA | NA | + | 0,0694326087332112 | 9 | -525,175472152902 | 1069,76826714045 | 21,8066942251585 | 6.39903030455766e-07 |
| 3219 | 25,9268008608122 | NA | + | NA | NA | + | NA | NA | + | NA | NA | + | + | 0,0379899756379695 | 7 | -527,450704622139 | 1069,76962629854 | 21,8080533832504 | 6.39468313493837e-07 |
| 1039 | 25,918607885758 | NA | + | + | + | NA | NA | NA | NA | NA | NA | + | NA | 0,0222187753630791 | 6 | -528,564088107631 | 1069,77433006142 | 21,8127571461246 | 6.37966127012389e-07 |
| 473 | 26,6007778281221 | NA | NA | NA | + | + | NA | + | + | + | NA | NA | NA | 0,0379252450553672 | 7 | -527,455311526356 | 1069,77884010698 | 21,8172671916848 | 6.36529119714084e-07 |
| 3204 | 25,9404997446814 | + | + | NA | NA | NA | NA | NA | + | NA | NA | + | + | 0,0379122397066761 | 7 | -527,456237085402 | 1069,78069122507 | 21,8191183097765 | 6.35940246989651e-07 |
| 2234 | 25,541481756124 | + | NA | NA | + | + | + | NA | + | NA | NA | NA | + | 0,0536208949519151 | 8 | -526,329074810191 | 1069,78314962038 | 21,8215767050915 | 6.35159030960565e-07 |
| 2140 | 24,9129565083967 | + | + | NA | + | + | NA | + | NA | NA | NA | NA | + | 0,0536149943889168 | 8 | -526,329501701758 | 1069,78400340352 | 21,8224304882258 | 6.34887944792769e-07 |
| 2442 | 26,449252186696 | + | NA | NA | + | NA | NA | NA | + | + | NA | NA | + | 0,0378409480940653 | 7 | -527,461310513974 | 1069,79083808221 | 21,8292651669217 | 6.3272202019501e-07 |
| 2141 | 25,2803713591297 | NA | NA | + | + | + | NA | + | NA | NA | NA | NA | + | 0,0377752636339251 | 7 | -527,465984580042 | 1069,80018621435 | 21,8386132990574 | 6.2977153641964e-07 |
| 28 | 26,0550684703761 | + | + | NA | + | + | NA | NA | NA | NA | NA | NA | NA | 0,0220158820423474 | 6 | -528,578294312138 | 1069,80274247043 | 21,8411695551385 | 6.28967121941658e-07 |
| 2521 | 25,9338055267512 | NA | NA | NA | + | + | NA | + | + | + | NA | NA | + | 0,0534753539515958 | 8 | -526,339603575765 | 1069,80420715153 | 21,8426342362397 | 6.28506672436768e-07 |
| 3157 | 25,4393354985354 | NA | NA | + | NA | + | NA | + | NA | NA | NA | + | + | 0,0377157240755533 | 7 | -527,470221103098 | 1069,80865926046 | 21,847086345168 | 6.27109138423371e-07 |
| 2538 | 25,9780126027092 | + | NA | NA | + | NA | + | + | + | + | NA | NA | + | 0,0691182755686789 | 9 | -525,198595797929 | 1069,8145144305 | 21,8529415152127 | 6.25275907880831e-07 |
| 1210 | 25,7836866539224 | + | NA | NA | + | + | + | NA | + | NA | NA | + | NA | 0,0533984663843219 | 8 | -526,345165143284 | 1069,81533028657 | 21,8537573712767 | 6.25020892327794e-07 |
| 3561 | 25,9575977828045 | NA | NA | NA | + | NA | + | + | + | + | NA | + | + | 0,0691105432868045 | 9 | -525,199164518141 | 1069,81565187093 | 21,8540789556366 | 6.24920401935259e-07 |
| 3465 | 26,4947957053407 | NA | NA | NA | + | NA | NA | NA | + | + | NA | + | + | 0,0375682157917534 | 7 | -527,480715891249 | 1069,82964883676 | 21,8680759214708 | 6.20562175452935e-07 |
| 2509 | 25,9413304545607 | NA | NA | + | + | NA | NA | + | + | + | NA | NA | + | 0,05326459834952 | 8 | -526,354847245399 | 1069,8346944908 | 21,8731215755074 | 6.18998577602925e-07 |
| 1328 | 25,427005067715 | + | + | + | + | NA | + | NA | NA | + | NA | + | NA | 0,0689800201821014 | 9 | -525,208763963644 | 1069,83485076193 | 21,8732778466426 | 6.18950213687212e-07 |
| 3368 | 25,4679068767238 | + | + | + | NA | NA | + | NA | NA | + | NA | + | + | 0,0689789285147931 | 9 | -525,208844245679 | 1069,835011326 | 21,8734384107124 | 6.18900525099133e-07 |
| 240 | 25,1001635277648 | + | + | + | + | NA | + | + | + | NA | NA | NA | NA | 0,0689420802372309 | 9 | -525,211554040537 | 1069,84043091572 | 21,8788580004293 | 6.17225701876858e-07 |
| 2136 | 24,9399624354645 | + | + | + | NA | + | NA | + | NA | NA | NA | NA | + | 0,0532075346566485 | 8 | -526,358974002147 | 1069,84294800429 | 21,8813750890038 | 6.1644938461313e-07 |
| 2299 | 24,8836990172002 | NA | + | NA | + | + | + | + | + | NA | NA | NA | + | 0,0689057326549918 | 9 | -525,214226909617 | 1069,84577665388 | 21,8842037385891 | 6.15578141220837e-07 |
| 3380 | 25,8080818987197 | + | + | NA | NA | + | + | NA | NA | + | NA | + | + | 0,0689011438045974 | 9 | -525,214564349581 | 1069,84645153381 | 21,8848786185163 | 6.15370455597972e-07 |
| 2292 | 25,0613112987652 | + | + | NA | NA | + | + | + | + | NA | NA | NA | + | 0,0688639152758315 | 9 | -525,217301878668 | 1069,85192659198 | 21,89035367669 | 6.13688164787193e-07 |
| 190 | 25,5570705002594 | + | NA | + | + | + | + | NA | + | NA | NA | NA | NA | 0,0531349288833318 | 8 | -526,36422437839 | 1069,85344875678 | 21,8918758414891 | 6.13221275219415e-07 |
| 494 | 25,9179122336173 | + | NA | + | + | NA | + | + | + | + | NA | NA | NA | 0,0688409318393385 | 9 | -525,218991867358 | 1069,85530656936 | 21,8937336540719 | 6.12651914601119e-07 |
| 3806 | 24,959751284773 | + | NA | + | + | + | NA | + | + | NA | + | + | + | 0,10022176391507 | 11 | -522,871806774978 | 1069,85561354996 | 21,8940406346646 | 6.12557885693613e-07 |
| 1517 | 25,9166138103141 | NA | NA | + | + | NA | + | + | + | + | NA | + | NA | 0,068829132693936 | 9 | -525,219859450848 | 1069,85704173634 | 21,8954688210515 | 6.12120618420226e-07 |
| 509 | 25,9272399934723 | NA | NA | + | + | + | + | + | + | + | NA | NA | NA | 0,0688250777103087 | 9 | -525,220157608621 | 1069,85763805189 | 21,8960651365967 | 6.11938137105544e-07 |
| 1036 | 25,9891165640633 | + | + | NA | + | NA | NA | NA | NA | NA | NA | + | NA | 0,0216117891973382 | 6 | -528,606579350486 | 1069,85931254713 | 21,8977396318355 | 6.1142600777647e-07 |
| 1239 | 25,7114427368739 | NA | + | + | NA | + | NA | + | + | NA | NA | + | NA | 0,0530938159095307 | 8 | -526,367197222213 | 1069,85939444443 | 21,8978215291359 | 6.11400971219385e-07 |
| 1236 | 25,6903121836185 | + | + | NA | NA | + | NA | + | + | NA | NA | + | NA | 0,0530834284035172 | 8 | -526,367948313435 | 1069,86089662687 | 21,899323711579 | 6.10941925731148e-07 |
| 3149 | 25,2379845989673 | NA | NA | + | + | NA | NA | + | NA | NA | NA | + | + | 0,0373158950125607 | 7 | -527,49866405889 | 1069,86554517204 | 21,9039722567536 | 6.09523579111334e-07 |
| 2287 | 24,8864655350361 | NA | + | + | + | NA | + | + | + | NA | NA | NA | + | 0,0687711351313306 | 9 | -525,224123814173 | 1069,86557046299 | 21,9039975477012 | 6.09515871445607e-07 |
| 2134 | 25,3814522476912 | + | NA | + | NA | + | NA | + | NA | NA | NA | NA | + | 0,0373002457977391 | 7 | -527,499777069308 | 1069,86777119288 | 21,9061982775881 | 6.08845550414707e-07 |
| 3095 | 25,6090030392276 | NA | + | + | NA | + | NA | NA | NA | NA | NA | + | + | 0,0372653057056933 | 7 | -527,502262028813 | 1069,87274111189 | 21,911168196599 | 6.07334472138619e-07 |
| 3151 | 24,7494781243031 | NA | + | + | + | NA | NA | + | NA | NA | NA | + | + | 0,0529331288773976 | 8 | -526,378815126085 | 1069,88263025217 | 21,9210573368796 | 6.04338876312556e-07 |
| 1474 | 26,5786935975741 | + | NA | NA | NA | NA | NA | + | + | + | NA | + | NA | 0,0371376823837302 | 7 | -527,511337907192 | 1069,89089286865 | 21,9293199533574 | 6.01847316378552e-07 |
| 466 | 26,5846027591034 | + | NA | NA | NA | + | NA | + | + | + | NA | NA | NA | 0,0370472680517218 | 7 | -527,517766956398 | 1069,90375096706 | 21,9421780517685 | 5.97990421734712e-07 |
| 220 | 25,4845942176059 | + | + | NA | + | + | NA | + | + | NA | NA | NA | NA | 0,0527757819538549 | 8 | -526,390189626178 | 1069,90537925236 | 21,943806337066 | 5.97503770357519e-07 |
| 2280 | 25,0489176094507 | + | + | + | NA | NA | + | + | + | NA | NA | NA | + | 0,0684970060311388 | 9 | -525,244276001661 | 1069,90587483797 | 21,9443019226771 | 5.97355731564156e-07 |
| 351 | 25,6460152249635 | NA | + | + | + | + | NA | + | NA | + | NA | NA | NA | 0,052764866607656 | 8 | -526,39097861894 | 1069,90695723788 | 21,9453843225899 | 5.97032530134475e-07 |
| 1367 | 25,7675447174024 | NA | + | + | NA | + | NA | + | NA | + | NA | + | NA | 0,0527432517444086 | 8 | -526,392540976894 | 1069,91008195379 | 21,9485090384981 | 5.9610047989982e-07 |
| 223 | 25,4765218781423 | NA | + | + | + | + | NA | + | + | NA | NA | NA | NA | 0,0527287087972895 | 8 | -526,393592145037 | 1069,91218429007 | 21,9506113747841 | 5.95474207281889e-07 |
| 3090 | 26,1080028131073 | + | NA | NA | NA | + | NA | NA | NA | NA | NA | + | + | 0,0212297118977968 | 6 | -528,633312642183 | 1069,91277913052 | 21,9512062152282 | 5.95297127545775e-07 |

| | | | | | | | | | | | | | | | | | | |
|---|---|---|---|---|---|---|---|---|---|---|---|---|---|---|---|---|---|---|
| **3142** | 25,338248218562 | + | NA | + | NA | NA | NA | + | NA | NA | NA | + | + | 0,0368773815216136 | 7 | -527,529845365856 | 1069,92790778597 | 21,966334870684 | 5.90811093284754e-07 |
| **1224** | 25,6934891183992 | + | + | + | NA | NA | NA | + | + | NA | NA | + | NA | 0,0525974198075177 | 8 | -526,403081018596 | 1069,93116203719 | 21,9695891219005 | 5.89850551098548e-07 |
| **1371** | 25,754953560772 | NA | + | NA | + | + | NA | + | NA | + | NA | + | NA | 0,0525717320827002 | 8 | -526,404937437698 | 1069,9348748754 | 21,9733019601047 | 5.88756557038044e-07 |
| **255** | 25,029024929675 | NA | + | + | + | + | + | + | + | NA | NA | NA | NA | 0,0682584843784453 | 9 | -525,261805739121 | 1069,94093431289 | 21,9793613975969 | 5.86975489689249e-07 |
| **2352** | 25,1647432192553 | + | + | + | + | NA | + | NA | NA | + | NA | NA | + | 0,0682499126929236 | 9 | -525,262435616869 | 1069,94219406838 | 21,980621153092 | 5.86605883305579e-07 |
| **2333** | 25,9457507234864 | NA | NA | + | + | + | NA | NA | NA | + | NA | NA | + | 0,0367713627240662 | 7 | -527,537381896972 | 1069,94298084821 | 21,9814079329162 | 5.86375163853025e-07 |
| **1411** | 27,1390723367136 | NA | + | NA | NA | NA | NA | NA | + | + | NA | + | NA | 0,0208921208875799 | 6 | -528,656924626123 | 1069,9600030984 | 21,9984301831098 | 5.81405629642566e-07 |
| **2438** | 26,2665204066024 | + | NA | + | NA | NA | NA | NA | + | + | NA | NA | + | 0,0366467054492878 | 7 | -527,546242316215 | 1069,96070168669 | 21,9991287714033 | 5.81202583522714e-07 |
| **24** | 26,0597617453652 | + | + | + | NA | + | NA | NA | NA | NA | NA | NA | NA | 0,0208550978929049 | 6 | -528,659513614465 | 1069,96518107508 | 22,0036081597925 | 5.79902324105779e-07 |
| **1477** | 26,5317167115113 | NA | NA | + | NA | NA | NA | + | + | + | NA | + | NA | 0,0366057363310196 | 7 | -527,549154078676 | 1069,96652521162 | 22,004952296325 | 5.79512721090813e-07 |
| **403** | 27,132549228717 | NA | + | NA | NA | + | NA | NA | + | + | NA | NA | NA | 0,0208321106842797 | 6 | -528,661121042301 | 1069,96839593076 | 22,0068230154645 | 5.78970921749432e-07 |
| **1299** | 27,0784729428837 | NA | + | NA | NA | + | NA | NA | NA | + | NA | + | NA | 0,0207977795536417 | 6 | -528,66352164698 | 1069,97319714011 | 22,0116242248223 | 5.77582708388167e-07 |
| **2516** | 25,5051151086269 | + | + | NA | NA | + | NA | + | + | + | NA | NA | + | 0,0680315906046801 | 9 | -525,278476743403 | 1069,97427632145 | 22,0127034061604 | 5.77271134216925e-07 |
| **344** | 25,6995501233532 | + | + | + | NA | + | NA | + | NA | + | NA | NA | NA | 0,0522609081134016 | 8 | -526,427396302894 | 1069,97979260579 | 22,0182196904968 | 5.75681132093082e-07 |
| **3467** | 25,9806689883978 | NA | + | NA | + | NA | NA | NA | + | + | NA | + | + | 0,0522577869202495 | 8 | -526,427621790171 | 1069,98024358034 | 22,0186706650522 | 5.75551337955795e-07 |
| **2519** | 25,4758739000371 | NA | + | + | NA | + | NA | + | + | + | NA | NA | + | 0,067962000513332 | 9 | -525,283589057612 | 1069,98450094987 | 22,0229280345793 | 5.74327473664832e-07 |
| **506** | 26,0178767393693 | + | NA | NA | + | + | + | + | + | + | NA | NA | NA | 0,0679577849807747 | 9 | -525,283898732074 | 1069,98512029879 | 22,0235473835032 | 5.741496466491e-07 |
| **1529** | 26,0232828221412 | NA | NA | NA | + | + | + | + | + | + | NA | + | NA | 0,0679401844200508 | 9 | -525,285191660154 | 1069,98770615495 | 22,0261332396626 | 5.73407792134613e-07 |
| **3341** | 25,8805955285001 | NA | NA | + | + | NA | NA | NA | NA | + | NA | + | + | 0,0364512878788593 | 7 | -527,560129946314 | 1069,98847694689 | 22,0269040316011 | 5.73186845661456e-07 |
| **2504** | 25,475349976016 | + | + | + | NA | NA | NA | + | + | + | NA | NA | + | 0,0679278857404171 | 9 | -525,286095100309 | 1069,98951303526 | 22,0279401199734 | 5.72889986448217e-07 |
| **3080** | 25,4972685927737 | + | + | + | NA | NA | NA | NA | NA | NA | NA | + | + | 0,0364164802091812 | 7 | -527,562603307924 | 1069,99342367011 | 22,0318507548209 | 5.71770899117864e-07 |
| **2295** | 25,0005108183834 | NA | + | + | NA | + | + | + | + | NA | NA | NA | + | 0,0678928388839477 | 9 | -525,288669517614 | 1069,99466186987 | 22,0330889545837 | 5.7141702537492e-07 |
| **3161** | 25,5057732384315 | NA | NA | NA | + | + | NA | + | NA | NA | NA | + | + | 0,0364068835658629 | 7 | -527,563285210008 | 1069,99478747428 | 22,0332145589898 | 5.71381140253713e-07 |
| **2318** | 25,7674292384953 | + | NA | + | + | NA | NA | NA | NA | + | NA | NA | + | 0,0363644382740652 | 7 | -527,566301134512 | 1070,00081932329 | 22,0392464079964 | 5.69660493846149e-07 |
| **3563** | 25,3991214481418 | NA | + | NA | + | NA | + | + | + | + | NA | + | + | 0,0835370658602374 | 10 | -524,129773737097 | 1070,00557922023 | 22,0440063049352 | 5.68306343269209e-07 |
| **3529** | 26,0878068416838 | NA | NA | NA | + | NA | NA | + | + | + | NA | + | + | 0,0520407005429841 | 8 | -526,443303143598 | 1070,0116062872 | 22,0500333719056 | 5.66596310987831e-07 |
| **2506** | 26,0998750757566 | + | NA | NA | + | NA | NA | + | + | + | NA | NA | + | 0,0520322279747947 | 8 | -526,443915091423 | 1070,01283018285 | 22,051257267555 | 5.66249689675721e-07 |
| **3146** | 25,4526582301984 | + | NA | NA | + | NA | NA | + | NA | NA | NA | + | + | 0,0362779281283238 | 7 | -527,572447649437 | 1070,01311235314 | 22,0515394378469 | 5.66169805890939e-07 |
| **206** | 26,0329662001956 | + | NA | + | + | NA | NA | + | + | NA | NA | NA | NA | 0,0362442715891469 | 7 | -527,574838785496 | 1070,01789462526 | 22,0563217099652 | 5.64817634110082e-07 |
| **248** | 25,2112996539388 | + | + | + | NA | + | + | + | + | NA | NA | NA | NA | 0,0677127065278122 | 9 | -525,301899872798 | 1070,02112258024 | 22,0595496649501 | 5.63906766419016e-07 |
| **2534** | 25,8925528645573 | + | NA | + | NA | NA | + | + | + | + | NA | NA | + | 0,0677071645362057 | 9 | -525,302306880293 | 1070,02193659523 | 22,060363679941 | 5.6367729883901e-07 |
| **208** | 25,4760654473735 | + | + | + | + | NA | NA | + | + | NA | NA | NA | NA | 0,0519622643231462 | 8 | -526,448968144184 | 1070,02293628837 | 22,0613633730775 | 5.6339561708029e-07 |
| **1047** | 26,4665306472432 | NA | + | + | NA | + | NA | NA | NA | NA | NA | + | NA | 0,0204179205661955 | 6 | -528,690077677058 | 1070,02630920027 | 22,0647362849793 | 5.62446275925177e-07 |
| **1042** | 26,8008035714286 | + | NA | NA | NA | + | NA | NA | NA | NA | NA | + | NA | 0,0046092140128146 | 5 | -529,786236314473 | 1070,03048789612 | 22,0689149808316 | 5.61272356754855e-07 |
| **2493** | 25,8618268567409 | NA | NA | + | + | + | + | NA | + | + | NA | NA | + | 0,067647264506736 | 9 | -525,306705823394 | 1070,03073448143 | 22,0691615661433 | 5.61203160261121e-07 |
| **2549** | 25,9031333744191 | NA | NA | + | NA | + | + | + | + | + | NA | NA | + | 0,0676406120696443 | 9 | -525,307194348167 | 1070,03171153098 | 22,0701386156891 | 5.60929065571183e-07 |
| **2126** | 25,0978965307991 | + | NA | + | + | NA | NA | + | NA | NA | NA | NA | + | 0,03613436222692 | 7 | -527,582646738461 | 1070,03351053119 | 22,0719376158954 | 5.6042473667478e-07 |
| **2096** | 24,7182230254572 | + | + | + | + | NA | + | NA | NA | NA | NA | NA | + | 0,0518757996411171 | 8 | -526,455212451945 | 1070,03542490389 | 22,0738519885995 | 5.59888562415776e-07 |
| **348** | 25,7215565573873 | + | + | NA | + | + | NA | + | NA | + | NA | NA | NA | 0,0518729079096782 | 8 | -526,455421277215 | 1070,03584255443 | 22,0742696391392 | 5.59771655742602e-07 |
| **1514** | 26,0692945187471 | + | NA | NA | + | NA | + | + | + | + | NA | + | NA | 0,0676042803587917 | 9 | -525,309862322567 | 1070,03704747978 | 22,0754745644892 | 5.59434515780794e-07 |
| **1364** | 25,8387505691073 | + | + | NA | NA | + | NA | + | NA | + | NA | + | NA | 0,0518498736319947 | 8 | -526,457084665956 | 1070,03916933191 | 22,0775964166223 | 5.58841311848876e-07 |
| **3501** | 25,8959254117507 | NA | NA | + | + | NA | + | NA | + | + | NA | + | + | 0,0675819011162854 | 9 | -525,311505662938 | 1070,04033416052 | 22,0787612452305 | 5.58515929438037e-07 |
| **2143** | 24,7843024828025 | NA | + | + | + | + | NA | + | NA | NA | NA | NA | + | 0,0518396351200039 | 8 | -526,457824013078 | 1070,04064802616 | 22,0790751108646 | 5.58428286837044e-07 |
| **2478** | 25,8773740792325 | + | NA | + | + | NA | + | NA | + | + | NA | NA | + | 0,067578652331476 | 9 | -525,311744222661 | 1070,04081127997 | 22,0792383646767 | 5.58382705924061e-07 |
| **2138** | 25,4753699847917 | + | NA | NA | + | + | NA | + | NA | NA | NA | NA | + | 0,0360826017438014 | 7 | -527,586323491121 | 1070,04086403651 | 22,079291121215 | 5.58367976949032e-07 |
| **3557** | 25,9210330804397 | NA | NA | + | NA | NA | + | + | + | + | NA | + | + | 0,0675141588716965 | 9 | -525,316479834344 | 1070,05028250333 | 22,0887095880435 | 5.55744673536636e-07 |
| **136** | 26,1321071043868 | + | + | + | NA | NA | NA | NA | + | NA | NA | NA | NA | 0,0202069785561584 | 6 | -528,704820237251 | 1070,05579432066 | 22,0942214053653 | 5.54215200489628e-07 |
| **397** | 26,9205442398129 | NA | NA | + | + | NA | NA | NA | + | + | NA | NA | NA | 0,0201843692801788 | 6 | -528,706400192342 | 1070,05895423084 | 22,0973813155472 | 5.53340256730058e-07 |
| **2079** | 25,2166045257375 | NA | + | + | + | + | NA | NA | NA | NA | NA | NA | + | 0,0359245523450406 | 7 | -527,597549146371 | 1070,06331534701 | 22,1017424317147 | 5.52134980724922e-07 |
| **469** | 26,639444546815 | NA | NA | + | NA | + | NA | + | + | + | NA | NA | NA | 0,0358475879820751 | 7 | -527,60301497007 | 1070,0742469944 | 22,1126740791126 | 5.49125340835573e-07 |
| **216** | 25,5722658885867 | + | + | + | NA | + | NA | + | + | NA | NA | NA | NA | 0,0514973534546068 | 8 | -526,482536388566 | 1070,09007277713 | 22,1284998618416 | 5.44797317838756e-07 |
| **1156** | 26,5645415918551 | + | + | NA | NA | NA | NA | NA | + | NA | NA | + | NA | 0,019960128539065 | 6 | -528,722068349882 | 1070,09029054592 | 22,1287176306262 | 5.44738001143287e-07 |
| **218** | 26,1324665836115 | + | NA | NA | + | + | NA | + | + | NA | NA | NA | NA | 0,0357292232134561 | 7 | -527,611420099823 | 1070,09105725391 | 22,1294843386197 | 5.4452921367567e-07 |
| **3092** | 25,7805310950646 | + | + | NA | NA | + | NA | NA | NA | NA | NA | + | + | 0,0356988174400098 | 7 | -527,613579059524 | 1070,09537517331 | 22,1338022580201 | 5.43354865189447e-07 |
| **3188** | 25,0608275139905 | + | + | NA | NA | + | + | + | NA | NA | NA | + | + | 0,0671107557073505 | 9 | -525,346093407829 | 1070,1095096503 | 22,1479367350137 | 5.39528384022655e-07 |
| **1229** | 26,1308753195237 | NA | NA | + | + | NA | NA | + | + | NA | NA | + | NA | 0,0355343323191906 | 7 | -527,62525713383 | 1070,11873132192 | 22,1571584066326 | 5.37046433559575e-07 |
| **1226** | 26,170931184314 | + | NA | NA | + | NA | NA | + | + | NA | NA | + | NA | 0,0355271497308768 | 7 | -527,625767036111 | 1070,11975112649 | 22,158178211195 | 5.36772662162234e-07 |
| **3154** | 25,6421194731623 | + | NA | NA | NA | + | NA | + | NA | NA | NA | + | + | 0,0355185638694253 | 7 | -527,626376553791 | 1070,12097016185 | 22,159397246556 | 5.36445589422694e-07 |
| **221** | 26,1572109993703 | NA | NA | + | + | + | NA | + | + | NA | NA | NA | NA | 0,035506034383239 | 7 | -527,627266023048 | 1070,12274910036 | 22,1611761850681 | 5.35968649706387e-07 |
| **2502** | 25,9950161883392 | + | NA | + | NA | NA | NA | + | + | + | NA | NA | + | 0,051250427537113 | 8 | -526,500358645437 | 1070,12571729087 | 22,1641443755843 | 5.35173811128842e-07 |
| **2517** | 26,0012656519198 | NA | NA | + | NA | + | NA | + | + | + | NA | NA | + | 0,0512457837885581 | 8 | -526,500693770695 | 1070,12638754139 | 22,1648146260989 | 5.3499449091675e-07 |
| **270** | 26,3210007982224 | + | NA | + | + | NA | NA | NA | NA | + | NA | NA | NA | 0,0196801607347328 | 6 | -528,741625241806 | 1070,12940432977 | 22,1678314144754 | 5.34188116653923e-07 |
| **3189** | 25,2863376873664 | NA | NA | + | NA | + | + | + | NA | NA | NA | + | + | 0,0511980100840947 | 8 | -526,504141358943 | 1070,13328271789 | 22,1717098025954 | 5.33153225982406e-07 |
| **390** | 26,8043774489878 | + | NA | + | NA | NA | NA | NA | + | + | NA | NA | NA | 0,0196262663036392 | 6 | -528,745389346631 | 1070,13693253942 | 22,1753596241258 | 5.32181156160919e-07 |
| **1149** | 25,3032796116596 | NA | NA | + | + | + | + | + | NA | NA | NA | + | NA | 0,0511533818612908 | 8 | -526,507361796834 | 1070,13972359367 | 22,1781506783773 | 5.31439000891041e-07 |
| **3253** | 25,716354178009 | NA | NA | + | NA | + | + | NA | + | NA | NA | + | + | 0,0510959662893837 | 8 | -526,511504765876 | 1070,14800953175 | 22,1864366164609 | 5.29241820131401e-07 |
| **2173** | 25,1043361348289 | NA | NA | + | + | + | + | + | NA | NA | NA | NA | + | 0,0510843270181369 | 8 | -526,512344597041 | 1070,14968919408 | 22,1881162787909 | 5.28797532946276e-07 |
| **3238** | 25,7058167637215 | + | NA | + | NA | NA | + | NA | + | NA | NA | + | + | 0,051061861489374 | 8 | -526,51396556728 | 1070,15293113456 | 22,1913582192699 | 5.2794106222695e-07 |
| **1409** | 27,7509088201308 | NA | NA | NA | NA | NA | NA | NA | + | + | NA | + | NA | 0,0037156426491310 | 5 | -529,847674914763 | 1070,1533650967 | 22,1917921814118 | 5.27826521436919e-07 |
| **1032** | 26,0212810751393 | + | + | + | NA | NA | NA | NA | NA | NA | NA | + | NA | 0,0194961490875309 | 6 | -528,754476164896 | 1070,15510617595 | 22,1935332606549 | 5.27367227482482e-07 |
| **2540** | 25,3577357985594 | + | + | NA | + | NA | + | + | + | + | NA | NA | + | 0,082522930717869 | 10 | -524,205496239503 | 1070,15702422504 | 22,1954513097471 | 5.26861711806238e-07 |
| **126** | 25,2878463982146 | + | NA | + | + | + | + | + | NA | NA | NA | NA | NA | 0,0510152789639194 | 8 | -526,517326545 | 1070,15965309 | 22,1980801747097 | 5.26169642597848e-07 |
| **151** | 26,4138270820089 | NA | + | + | NA | + | NA | NA | + | NA | NA | NA | NA | 0,0194582236904951 | 6 | -528,75712448218 | 1070,16040281051 | 22,198829895224 | 5.25972439474448e-07 |
| **2076** | 25,3895788095087 | + | + | NA | + | + | NA | NA | NA | NA | NA | NA | + | 0,0351921201615851 | 7 | -527,649547050736 | 1070,16731115574 | 22,2057382404446 | 5.24158774040257e-07 |
| **2447** | 25,7940995403075 | NA | + | + | + | NA | NA | NA | + | + | NA | NA | + | 0,0508665505875231 | 8 | -526,528056348581 | 1070,18111269716 | 22,2195397818723 | 5.20554126251521e-07 |
| **401** | 27,7792962293138 | NA | NA | NA | NA | + | NA | NA | + | + | NA | NA | NA | 0,0034846856155258 | 5 | -529,863545692283 | 1070,18510665174 | 22,2235337364509 | 5.19515628756617e-07 |

| | | | | | | | | | | | | | | | | | | |
|---|---|---|---|---|---|---|---|---|---|---|---|---|---|---|---|---|---|---|
| 1148 | 25,1189854694441 | + | + | NA | + | + | + | + | NA | NA | NA | + | NA | 0,0665841373492448 | 9 | -525,384732876381 | 1070,18678858741 | 22,2252156721163 | 5.1907891648048e-07 |
| 1241 | 26,232825455937 | NA | NA | NA | + | + | NA | + | + | NA | NA | + | NA | 0,0350396498265244 | 7 | -527,66036648751 | 1070,18895002928 | 22,227377113994 | 5.18518240048485e-07 |
| 394 | 27,0760918122533 | + | NA | NA | + | NA | NA | NA | + | + | NA | NA | NA | 0,0192519642742552 | 6 | -528,77152571312 | 1070,18920527239 | 22,2276323571025 | 5.18452070167214e-07 |
| 2166 | 25,2600655239915 | + | NA | + | NA | + | + | + | NA | NA | NA | NA | + | 0,0507902902947425 | 8 | -526,533557390461 | 1070,19211478092 | 22,230541865631 | 5.17698398144029e-07 |
| 1159 | 26,5509440017064 | NA | + | + | NA | NA | NA | NA | + | NA | NA | + | NA | 0,0192214797788331 | 6 | -528,773653913142 | 1070,19346167244 | 22,231888757148 | 5.17349873722946e-07 |
| 2514 | 26,050200982669 | + | NA | NA | NA | + | NA | + | + | + | NA | NA | + | 0,0507767044828533 | 8 | -526,53453735765 | 1070,1940747153 | 22,2325018000092 | 5.17191319200944e-07 |
| 2546 | 25,9996405334989 | + | NA | NA | NA | + | + | + | + | + | NA | NA | + | 0,0664861769879487 | 9 | -525,391918099652 | 1070,20115903395 | 22,2395861186583 | 5.15362585891521e-07 |
| 3525 | 26,0431033874333 | NA | NA | + | NA | NA | NA | + | + | + | NA | + | + | 0,0507272844791693 | 8 | -526,538101986395 | 1070,20120397279 | 22,2396310574995 | 5.15351006122899e-07 |
| 16 | 25,5827958046574 | + | + | + | + | NA | NA | NA | NA | NA | NA | NA | NA | 0,0191185668569835 | 6 | -528,780838037156 | 1070,20782992047 | 22,246257005175 | 5.13646486795362e-07 |
| 1510 | 25,9709004566586 | + | NA | + | NA | NA | + | + | + | + | NA | + | NA | 0,0663594562390396 | 9 | -525,401211729728 | 1070,2197462941 | 22,2581733788111 | 5.10595184210796e-07 |
| 3513 | 26,0157403091252 | NA | NA | NA | + | + | + | NA | + | + | NA | + | + | 0,0663508022550151 | 9 | -525,401846362168 | 1070,22101555898 | 22,2594426436901 | 5.10271246744913e-07 |
| 2555 | 25,3717163952409 | NA | + | NA | + | + | + | + | + | + | NA | NA | + | 0,0820785311208842 | 10 | -524,23865189758 | 1070,22333554119 | 22,2617626259012 | 5.09679679809817e-07 |
| 3569 | 26,0414656443591 | NA | NA | NA | NA | + | + | + | + | + | NA | + | + | 0,0663109827940206 | 9 | -525,40476641216 | 1070,22685565897 | 22,2652827436759 | 5.08783402541408e-07 |
| 502 | 25,9920563500823 | + | NA | + | NA | + | + | + | + | + | NA | NA | NA | 0,0663052516345636 | 9 | -525,405186680634 | 1070,22769619591 | 22,2661232806236 | 5.08569621843109e-07 |
| 2199 | 25,6380572137776 | NA | + | + | NA | + | NA | NA | + | NA | NA | NA | + | 0,0347422327151203 | 7 | -527,681466565613 | 1070,23115018549 | 22,2695772701982 | 5.07692082728049e-07 |
| 446 | 25,9775280192741 | + | NA | + | + | + | + | NA | + | + | NA | NA | NA | 0,0662734829327192 | 9 | -525,407516246921 | 1070,23235532849 | 22,2707824131971 | 5.07386254099544e-07 |
| 1454 | 25,9778960515005 | + | NA | + | + | NA | + | NA | + | + | NA | + | NA | 0,0662703320294968 | 9 | -525,407747295101 | 1070,23281742485 | 22,2712445095563 | 5.07269036971126e-07 |
| 2453 | 26,3910526476439 | NA | NA | + | NA | + | NA | NA | + | + | NA | NA | + | 0,0347241699831215 | 7 | -527,682747805947 | 1070,23371266616 | 22,272139750866 | 5.07042023684304e-07 |
| 2490 | 25,9956601596142 | + | NA | NA | + | + | + | NA | + | + | NA | NA | + | 0,0662480332059396 | 9 | -525,409382392102 | 1070,23608761885 | 22,2745147035587 | 5.06440280623119e-07 |
| 1142 | 25,4406187166931 | + | NA | + | NA | + | + | + | NA | NA | NA | + | NA | 0,0504492356246616 | 8 | -526,558153990032 | 1070,24130798006 | 22,2797350647738 | 5.05120103723347e-07 |
| 1222 | 26,0727587369978 | + | NA | + | NA | NA | NA | + | + | NA | NA | + | NA | 0,0346569450225182 | 7 | -527,687516050737 | 1070,24324915574 | 22,2816762404468 | 5.04630078139611e-07 |
| 3808 | 24,681892191511 | + | + | + | + | + | NA | + | + | NA | + | + | + | 0,113369914024015 | 12 | -521,863946168738 | 1070,24402136973 | 22,2824484544442 | 5.04435274544627e-07 |
| 1489 | 26,7285026324128 | NA | NA | NA | NA | + | NA | + | + | + | NA | + | NA | 0,034643234102632 | 7 | -527,688488521116 | 1070,24519409649 | 22,2836211812041 | 5.041395788731e-07 |
| 214 | 26,090851617689 | + | NA | + | NA | + | NA | + | + | NA | NA | NA | NA | 0,0346404998277137 | 7 | -527,688682452577 | 1070,24558195942 | 22,2840090441264 | 5.04041819827541e-07 |
| 2543 | 25,3365444954633 | NA | + | + | + | NA | + | + | + | + | NA | NA | + | 0,0819229675705333 | 10 | -524,250254354409 | 1070,24654045485 | 22,2849675395591 | 5.03800316810933e-07 |
| 3181 | 25,0543628561534 | NA | NA | + | + | NA | + | + | NA | NA | NA | + | + | 0,0504000055991581 | 8 | -526,561703692141 | 1070,24840738428 | 22,2868344689914 | 5.03330256417745e-07 |
| 1469 | 26,0082601216864 | NA | NA | + | + | + | + | NA | + | + | NA | + | NA | 0,0661618405510144 | 9 | -525,415702239787 | 1070,24872731422 | 22,2871543989299 | 5.03249747648207e-07 |
| 1516 | 25,5745502033124 | + | + | NA | + | NA | + | + | + | + | NA | + | NA | 0,0818508731358187 | 10 | -524,255630735348 | 1070,25729321673 | 22,2957203014375 | 5.01098962651965e-07 |
| 1044 | 26,4836907437132 | + | + | NA | NA | + | NA | NA | NA | NA | NA | + | NA | 0,0187551935604004 | 6 | -528,80619830008 | 1070,25855044631 | 22,2969775310223 | 5.00784063417119e-07 |
| 2184 | 25,5773794917197 | + | + | + | NA | NA | NA | NA | + | NA | NA | NA | + | 0,034545915531286 | 7 | -527,695390610041 | 1070,25899827435 | 22,2974253590558 | 5.00671943399062e-07 |
| 3461 | 26,4346791617593 | NA | NA | + | NA | NA | NA | NA | + | + | NA | + | + | 0,0345389068634112 | 7 | -527,695887656347 | 1070,25999236696 | 22,2984194516678 | 5.00423148095589e-07 |
| 3556 | 25,5380839839588 | + | + | NA | NA | NA | + | + | + | + | NA | + | + | 0,0817946823531442 | 10 | -524,259820822413 | 1070,26567339086 | 22,304100475568 | 4.99003707087275e-07 |
| 3537 | 26,1215780916419 | NA | NA | NA | NA | + | NA | + | + | + | NA | + | + | 0,0502519346745171 | 8 | -526,572379150763 | 1070,26975830153 | 22,3081853862359 | 4.97985554423696e-07 |
| 1531 | 25,5697742417775 | NA | + | NA | + | + | + | + | + | + | NA | + | NA | 0,081727139961889 | 10 | -524,264857047712 | 1070,27574584146 | 22,3141729261658 | 4.96496929639791e-07 |
| 3178 | 25,2776802285679 | + | NA | NA | + | NA | + | + | NA | NA | NA | + | + | 0,0502001873707064 | 8 | -526,57610958006 | 1070,27721916012 | 22,3156462448285 | 4.96131315226767e-07 |
| 3193 | 25,337847626929 | NA | NA | NA | + | + | + | + | NA | NA | NA | + | + | 0,0501619819816271 | 8 | -526,578863651218 | 1070,28272730244 | 22,3211543871462 | 4.94766814109554e-07 |
| 3174 | 25,2080899723064 | + | NA | + | NA | NA | + | + | NA | NA | NA | + | + | 0,0501582223374446 | 8 | -526,579134662697 | 1070,28326932539 | 22,3216964101032 | 4.94632744791731e-07 |
| 2170 | 25,3263766682281 | + | NA | NA | + | + | + | + | NA | NA | NA | NA | + | 0,0501446525799425 | 8 | -526,580112820824 | 1070,28522564165 | 22,3236527263571 | 4.9414915230608e-07 |
| 148 | 26,4727218229291 | + | + | NA | NA | + | NA | NA | + | NA | NA | NA | NA | 0,0185194564899545 | 6 | -528,822645656164 | 1070,29144515848 | 22,3298722431909 | 4.92614854704377e-07 |
| 1525 | 26,0474550860147 | NA | NA | + | NA | + | + | + | + | + | NA | + | NA | 0,0658604788938255 | 9 | -525,437794201437 | 1070,29291123752 | 22,3313383222289 | 4.92253880868444e-07 |
| 2230 | 25,5864581669368 | + | NA | + | NA | + | + | NA | + | NA | NA | NA | + | 0,0500701079434453 | 8 | -526,585486023818 | 1070,29597204764 | 22,3343991323457 | 4.91501109208868e-07 |
| 3559 | 25,496548264027 | NA | + | + | NA | NA | + | + | + | + | NA | + | + | 0,0815729942215973 | 10 | -524,276349371395 | 1070,29873048882 | 22,3371575735312 | 4.90823688021697e-07 |
| 3571 | 25,525043838283 | NA | + | NA | NA | + | + | + | + | + | NA | + | + | 0,0815719607055758 | 10 | -524,276426418595 | 1070,29888458322 | 22,3373116679311 | 4.90785872887652e-07 |
| 1519 | 25,5496861628777 | NA | + | + | + | NA | + | + | + | + | NA | + | NA | 0,081542728785792 | 10 | -524,278605582179 | 1070,30324291039 | 22,3416699950994 | 4.89717534650492e-07 |
| 2459 | 25,9058014861362 | NA | + | NA | + | + | NA | NA | + | + | NA | NA | + | 0,0500173268279087 | 8 | -526,589290249901 | 1070,3035804998 | 22,342007584512 | 4.89634879899115e-07 |
| 2444 | 25,9149559292814 | + | + | NA | + | NA | NA | NA | + | + | NA | NA | + | 0,050015607043915 | 8 | -526,589414200664 | 1070,30382840133 | 22,3422554860379 | 4.89574193043357e-07 |
| 1297 | 27,5604352449878 | NA | NA | NA | NA | + | NA | NA | NA | + | NA | + | NA | 0,0025941194147640 | 5 | -529,924708724263 | 1070,3074327157 | 22,3458598004104 | 4.88692697928109e-07 |
| 278 | 26,68299330039 | + | NA | + | NA | + | NA | NA | NA | + | NA | NA | NA | 0,0183949444204174 | 6 | -528,831331259526 | 1070,30881636521 | 22,3472434499151 | 4.88354725145823e-07 |
| 2326 | 26,0601120680725 | + | NA | + | NA | + | NA | NA | NA | + | NA | NA | + | 0,034183045601689 | 7 | -527,721120170094 | 1070,31045739445 | 22,3488844791605 | 4.87954187298848e-07 |
| 1206 | 25,7863824847414 | + | NA | + | NA | + | + | NA | + | NA | NA | + | NA | 0,0499355103080916 | 8 | -526,595186800158 | 1070,31537360032 | 22,3538006850258 | 4.86756218648366e-07 |
| 1144 | 25,1635505283841 | + | + | + | NA | + | + | + | NA | NA | NA | + | NA | 0,0656513388051681 | 9 | -525,453121476588 | 1070,32356578782 | 22,3619928725311 | 4.84766497366511e-07 |
| 285 | 26,7722169090538 | NA | NA | + | + | + | NA | NA | NA | + | NA | NA | NA | 0,0182851931757757 | 6 | -528,838986277154 | 1070,32412640046 | 22,3625534851706 | 4.84630633296378e-07 |
| 3554 | 26,1198471589716 | + | NA | NA | NA | NA | + | + | + | + | NA | + | + | 0,0655927131067332 | 9 | -525,457417369933 | 1070,33215757451 | 22,3705846592204 | 4.82688458908355e-07 |
| 3334 | 26,0087334586692 | + | NA | + | NA | NA | NA | NA | NA | + | NA | + | + | 0,0339986298273894 | 7 | -527,734192603693 | 1070,33660226165 | 22,3750293463593 | 4.81616950386697e-07 |
| 3498 | 26,1232970154627 | + | NA | NA | + | NA | + | NA | + | + | NA | + | + | 0,0655429495147409 | 9 | -525,461063666633 | 1070,33945016791 | 22,3778772526216 | 4.80931638463998e-07 |
| 1146 | 25,5364766962273 | + | NA | NA | + | + | + | + | NA | NA | NA | + | NA | 0,0497222701862244 | 8 | -526,610552718495 | 1070,34610543699 | 22,3845325216992 | 4.79333933496472e-07 |
| 3250 | 25,886754676378 | + | NA | NA | NA | + | + | NA | + | NA | NA | + | + | 0,0497050819528861 | 8 | -526,611791139199 | 1070,3485822784 | 22,3870093631081 | 4.78740683851027e-07 |
| 508 | 25,5539562245942 | + | + | NA | + | + | + | + | + | + | NA | NA | NA | 0,0811289338244642 | 10 | -524,309445486179 | 1070,36492271839 | 22,4033498030988 | 4.7484520228334e-07 |
| 3244 | 25,3323193591741 | + | + | NA | + | NA | + | NA | + | NA | NA | + | + | 0,0653193021704679 | 9 | -525,477448443414 | 1070,37221972147 | 22,4106468061823 | 4.73115885456591e-07 |
| 3133 | 25,3038294461367 | NA | NA | + | + | + | + | NA | NA | NA | NA | + | + | 0,0495202797010025 | 8 | -526,625104818484 | 1070,37520963697 | 22,4136367216781 | 4.72409125617639e-07 |
| 3195 | 24,8676558893772 | NA | + | NA | + | + | + | + | NA | NA | NA | + | + | 0,0652828221904069 | 9 | -525,480120655757 | 1070,37756414616 | 22,415991230868 | 4.71853307037991e-07 |
| 3276 | 25,1303515625833 | + | + | NA | + | NA | NA | + | + | NA | NA | + | + | 0,0652605142484012 | 9 | -525,481754694047 | 1070,38083222274 | 22,4192593074495 | 4.71082910267019e-07 |
| 3186 | 25,5234182768508 | + | NA | NA | NA | + | + | + | NA | NA | NA | + | + | 0,0494286058639302 | 8 | -526,631708302089 | 1070,38841660418 | 22,4268436888874 | 4.69299857002658e-07 |
| 1134 | 25,2324318072975 | + | NA | + | + | NA | + | + | NA | NA | NA | + | NA | 0,0494217544661987 | 8 | -526,632201798929 | 1070,38940359786 | 22,4278306825684 | 4.69068316143031e-07 |
| 1234 | 26,2370631162933 | + | NA | NA | NA | + | NA | + | + | NA | NA | + | NA | 0,0336256073641712 | 7 | -527,760626922535 | 1070,38947089933 | 22,4278979840424 | 4.69052531914062e-07 |
| 3191 | 24,8900567861673 | NA | + | + | NA | + | + | + | NA | NA | NA | + | + | 0,0651392828754687 | 9 | -525,490634110345 | 1070,39859105533 | 22,4370181400443 | 4.66918485189746e-07 |
| 2196 | 25,7769158079125 | + | + | NA | NA | + | NA | NA | + | NA | NA | NA | + | 0,0334761686923509 | 7 | -527,771214065567 | 1070,4106451854 | 22,4490722701062 | 4.64112800630166e-07 |
| 142 | 26,3065032167338 | + | NA | + | + | NA | NA | NA | + | NA | NA | NA | NA | 0,0176610398791258 | 6 | -528,882503962081 | 1070,41116177032 | 22,4495888550246 | 4.63992939273857e-07 |
| 511 | 25,4960387053293 | NA | + | + | + | + | + | + | + | + | NA | NA | NA | 0,0808112799660962 | 10 | -524,333110631351 | 1070,41225300873 | 22,4506800934441 | 4.63739844865847e-07 |
| 1171 | 26,8580441027492 | NA | + | NA | NA | + | NA | NA | + | NA | NA | + | NA | 0,0175777297969377 | 6 | -528,888310479397 | 1070,42277480495 | 22,4612018896578 | 4.61306563030004e-07 |
| 3522 | 26,2151101184746 | + | NA | NA | NA | NA | NA | + | + | + | NA | + | + | 0,0491348478158007 | 8 | -526,652864102881 | 1070,43072820576 | 22,4691552904706 | 4.59475727787718e-07 |
| 2158 | 24,9197228179891 | + | NA | + | + | NA | + | + | NA | NA | NA | NA | + | 0,0490879527250359 | 8 | -526,656240778241 | 1070,43748155648 | 22,4759086411909 | 4.57926843929763e-07 |
| 496 | 25,5088766971222 | + | + | + | + | NA | + | + | + | + | NA | NA | NA | 0,0806337590359069 | 10 | -524,346332342712 | 1070,43869643146 | 22,4771235161645 | 4.5764876646444e-07 |
| 1086 | 25,4637935059049 | + | NA | + | + | + | + | NA | NA | NA | NA | + | NA | 0,0490345635953676 | 8 | -526,660084853478 | 1070,44516970696 | 22,4835967916647 | 4.56169917730451e-07 |
| 1169 | 27,4323075236086 | NA | NA | NA | NA | + | NA | NA | + | NA | NA | + | NA | 0,0015361120034329 | 5 | -529,997300512417 | 1070,45261629201 | 22,4910433767186 | 4.54474621684769e-07 |
| 342 | 26,1902352052762 | + | NA | + | NA | + | NA | + | NA | + | NA | NA | NA | 0,0331511889874105 | 7 | -527,794231954534 | 1070,45668096333 | 22,4951080480414 | 4.53551914646723e-07 |

| | | | | | | | | | | | | | | | | | | | |
|---|---|---|---|---|---|---|---|---|---|---|---|---|---|---|---|---|---|---|---|
| 2072 | 25,4614382139448 | + | + | + | NA | + | NA | NA | NA | NA | NA | NA | + | 0,0330893507011874 | 7 | -527,798611003976 | 1070,46543906222 | 22,5038661469255 | 4.51570130719756e-07 |
| 1237 | 26,266794676253 | NA | NA | + | NA | + | NA | + | + | NA | NA | + | NA | 0,0330629253133398 | 7 | -527,80048222014 | 1070,46918149454 | 22,5076085792527 | 4.50725935474358e-07 |
| 2110 | 25,2706668358832 | + | NA | + | + | + | + | NA | NA | NA | NA | NA | + | 0,0488423312457185 | 8 | -526,673924004709 | 1070,47284800942 | 22,5112750941269 | 4.49900395745708e-07 |
| 3213 | 25,8886900368576 | NA | NA | + | + | NA | NA | NA | + | NA | NA | + | + | 0,0330273448597918 | 7 | -527,803001637696 | 1070,47422032966 | 22,5126474143656 | 4.49591797922331e-07 |
| 1286 | 26,4789674741999 | + | NA | + | NA | NA | NA | NA | NA | + | NA | + | NA | 0,017206439723285 | 6 | -528,914182543385 | 1070,47451893292 | 22,5129460176338 | 4.4952467814289e-07 |
| 1522 | 26,1484940533148 | + | NA | NA | NA | + | + | + | + | + | NA | + | NA | 0,0646166105933542 | 9 | -525,528903311415 | 1070,47512945748 | 22,5135565421849 | 4.4938747615908e-07 |
| 2548 | 25,5126821996808 | + | + | NA | NA | + | + | + | + | + | NA | NA | + | 0,0803847924786119 | 10 | -524,364871006954 | 1070,47577375994 | 22,5142008446496 | 4.49242728746361e-07 |
| 3180 | 24,7375305650447 | + | + | NA | + | NA | + | + | NA | NA | NA | + | + | 0,064611687948644 | 9 | -525,529263637719 | 1070,47585011008 | 22,5142771947926 | 4.49225579200418e-07 |
| 3118 | 25,2408659660604 | + | NA | + | + | NA | + | NA | NA | NA | NA | + | + | 0,0488080578678812 | 8 | -526,676391112921 | 1070,47778222584 | 22,5162093105519 | 4.48791810846306e-07 |
| 1151 | 24,9517576189846 | NA | + | + | + | + | + | + | NA | NA | NA | + | NA | 0,0645811640995808 | 9 | -525,531497871158 | 1070,48031857696 | 22,5187456616702 | 4.48223024777159e-07 |
| 1466 | 26,1545169751104 | + | NA | NA | + | + | + | NA | + | + | NA | + | NA | 0,064562089197508 | 9 | -525,532834833129 | 1070,4829925009 | 22,5214195856124 | 4.47624168051994e-07 |
| 2064 | 25,0145919529282 | + | + | + | + | NA | NA | NA | NA | NA | NA | NA | + | 0,0329561487263785 | 7 | -527,808042688947 | 1070,48430243216 | 22,5227295168672 | 4.47331085597869e-07 |
| 2551 | 25,4533906322186 | NA | + | + | NA | + | + | + | + | + | NA | NA | + | 0,0803087921816179 | 10 | -524,370529177204 | 1070,48709010044 | 22,5255171851484 | 4.4670801458832e-07 |
| 3279 | 25,1568589073081 | NA | + | + | + | NA | NA | + | + | NA | NA | + | + | 0,0645232635227121 | 9 | -525,535735780127 | 1070,4887943949 | 22,5272214796087 | 4.4632751573455e-07 |
| 3349 | 26,2319175552752 | NA | NA | + | NA | + | NA | NA | NA | + | NA | + | + | 0,0328544174445075 | 7 | -527,815245141551 | 1070,49870733737 | 22,5371344220753 | 4.44120779572003e-07 |
| 2536 | 25,4510291659084 | + | + | + | NA | NA | + | + | + | + | NA | NA | + | 0,0802011766844411 | 10 | -524,378540278177 | 1070,50311230239 | 22,5415393870958 | 4.43143687731383e-07 |
| 3225 | 25,9829244526125 | NA | NA | NA | + | + | NA | NA | + | NA | NA | + | + | 0,0326118053013593 | 7 | -527,832418734012 | 1070,53305452229 | 22,5714816069976 | 4.36558749765442e-07 |
| 349 | 26,2741511172874 | NA | NA | + | + | + | NA | + | NA | + | NA | NA | NA | 0,032557445193166 | 7 | -527,836266089312 | 1070,54074923289 | 22,5791763175964 | 4.34882380008887e-07 |
| 3247 | 25,2628288385103 | NA | + | + | + | NA | + | NA | + | NA | NA | + | + | 0,0641168707565975 | 9 | -525,565473454037 | 1070,54826974272 | 22,5866968274288 | 4.33250182064874e-07 |
| 3259 | 25,2771728131573 | NA | + | NA | + | + | + | NA | + | NA | NA | + | + | 0,0641144898749776 | 9 | -525,565647636333 | 1070,54861810731 | 22,5870451920202 | 4.33174724125449e-07 |
| 3291 | 25,0842753062292 | NA | + | NA | + | + | NA | + | + | NA | NA | + | + | 0,0641053647290333 | 9 | -525,566315216396 | 1070,54995326744 | 22,5883803521476 | 4.32885641819056e-07 |
| 1293 | 26,5589550969434 | NA | NA | + | + | NA | NA | NA | NA | + | NA | + | NA | 0,0166056902117547 | 6 | -528,956023009042 | 1070,55819986424 | 22,5966269489481 | 4.31104399964156e-07 |
| 2450 | 26,5329504986931 | + | NA | NA | NA | + | NA | NA | + | + | NA | NA | + | 0,032418548025028 | 7 | -527,846095602672 | 1070,56040825961 | 22,5988353443163 | 4.30628638199981e-07 |
| 3176 | 24,8462164870904 | + | + | + | NA | NA | + | + | NA | NA | NA | + | + | 0,063988434233235 | 9 | -525,574869075612 | 1070,56706098587 | 22,6054880705783 | 4.29198590721905e-07 |
| 3343 | 25,5484539273241 | NA | + | + | + | NA | NA | NA | NA | + | NA | + | + | 0,0481737077791735 | 8 | -526,722037642586 | 1070,56907528517 | 22,6075023698818 | 4.28766541116557e-07 |
| 1527 | 25,6572850742509 | NA | + | + | NA | + | + | + | + | + | NA | + | NA | 0,0797288009731467 | 10 | -524,413693740881 | 1070,57341922779 | 22,611846312504 | 4.27836283102656e-07 |
| 3272 | 25,310161785318 | + | + | + | NA | NA | NA | + | + | NA | NA | + | + | 0,063928865363213 | 9 | -525,579226328239 | 1070,57577549112 | 22,6142025758329 | 4.27332532430903e-07 |
| 3338 | 26,1640004789592 | + | NA | NA | + | NA | NA | NA | NA | + | NA | + | + | 0,0322870889722554 | 7 | -527,855397434375 | 1070,57901192301 | 22,6174390077238 | 4.26641575322609e-07 |
| 1116 | 25,4880976814339 | + | + | NA | + | + | NA | + | NA | NA | NA | + | NA | 0,0480744351405508 | 8 | -526,729178346019 | 1070,58335669204 | 22,6217837767479 | 4.25715751758666e-07 |
| 2205 | 25,7306869013357 | NA | NA | + | + | + | NA | NA | + | NA | NA | NA | + | 0,0322440724918285 | 7 | -527,858440937262 | 1070,58509892879 | 22,6235260134979 | 4.25345064424491e-07 |
| 1524 | 25,7160421353653 | + | + | NA | NA | + | + | + | + | + | NA | + | NA | 0,0796498151092447 | 10 | -524,419569985458 | 1070,58517171695 | 22,6235988016572 | 4.25329584664028e-07 |
| 3353 | 26,1779182153372 | NA | NA | NA | + | + | NA | NA | NA | + | NA | + | + | 0,0322234049174956 | 7 | -527,859903161806 | 1070,58802337787 | 22,6264504625842 | 4.24723568925698e-07 |
| 2172 | 24,8453698640868 | + | + | NA | + | + | + | + | NA | NA | NA | NA | + | 0,0638268863020741 | 9 | -525,586685093348 | 1070,59069302134 | 22,6291201060501 | 4.24157016882515e-07 |
| 3130 | 25,5412141073511 | + | NA | NA | + | + | + | NA | NA | NA | NA | + | + | 0,0479828992917909 | 8 | -526,73576188085 | 1070,5965237617 | 22,6349508464098 | 4.22922242950047e-07 |
| 2190 | 25,7471218833536 | + | NA | + | + | NA | NA | NA | + | NA | NA | NA | + | 0,0321385893319198 | 7 | -527,865903511606 | 1070,60002407747 | 22,6384511621841 | 4.22182709589734e-07 |
| 1119 | 25,3742872043229 | NA | + | + | + | + | NA | + | NA | NA | NA | + | NA | 0,0479200804704644 | 8 | -526,740279634202 | 1070,6055592684 | 22,6439863531141 | 4.21015894018789e-07 |
| 2330 | 26,1296910251021 | + | NA | NA | + | + | NA | NA | NA | + | NA | NA | + | 0,0320680512259911 | 7 | -527,870893388816 | 1070,6100038319 | 22,6484309166049 | 4.20081316914432e-07 |
| 1417 | 27,2098541735955 | NA | NA | NA | + | NA | NA | NA | + | + | NA | + | NA | 0,016206369880133 | 6 | -528,983820382099 | 1070,61379461035 | 22,6522216950621 | 4.19285853405844e-07 |
| 2486 | 25,9920187246495 | + | NA | + | NA | + | + | NA | + | + | NA | NA | + | 0,0636517886650206 | 9 | -525,599489868889 | 1070,61630257242 | 22,6547296571339 | 4.18760406416179e-07 |
| 1512 | 25,6627476141342 | + | + | + | NA | NA | + | + | + | + | NA | + | NA | 0,0794376255014616 | 10 | -524,435353580786 | 1070,6167389076 | 22,6551659923132 | 4.18669056432835e-07 |
| 3284 | 25,2455960459021 | + | + | NA | NA | + | NA | + | + | NA | NA | + | + | 0,0636478526177106 | 9 | -525,599777681946 | 1070,61687819854 | 22,6553052832464 | 4.18639899046391e-07 |
| 409 | 27,2184524351454 | NA | NA | NA | + | + | NA | NA | + | + | NA | NA | NA | 0,0161614483345149 | 6 | -528,986946742181 | 1070,62004733052 | 22,6584744152249 | 4.17977061794005e-07 |
| 3494 | 26,0102684427079 | + | NA | + | NA | NA | + | NA | + | + | NA | + | + | 0,0636209719467613 | 9 | -525,601743227632 | 1070,62080928991 | 22,6592363746188 | 4.17817851349567e-07 |
| 3355 | 25,796779870526 | NA | + | NA | + | + | NA | NA | NA | + | NA | + | + | 0,047753344052846 | 8 | -526,752269404692 | 1070,62953880938 | 22,6679658940927 | 4.15998150981004e-07 |
| 1212 | 25,4929990379508 | + | + | NA | + | + | + | NA | + | NA | NA | + | NA | 0,0635560897401084 | 9 | -525,606487257126 | 1070,6302973489 | 22,6687244336065 | 4.15840405379379e-07 |
| 1112 | 25,4890674677263 | + | + | + | NA | + | NA | + | NA | NA | NA | + | NA | 0,0477319589005702 | 8 | -526,7538070277 | 1070,6326140554 | 22,6710411401084 | 4.15358994169877e-07 |
| 3126 | 25,502414085593 | + | NA | + | NA | + | + | NA | NA | NA | NA | + | + | 0,0477200537082127 | 8 | -526,754663013087 | 1070,63432602617 | 22,6727531108831 | 4.15003605065993e-07 |
| 2429 | 25,5843535128864 | NA | NA | + | + | + | + | + | NA | + | NA | NA | + | 0,0635201478657201 | 9 | -525,609115098351 | 1070,63555303135 | 22,6739801160577 | 4.14749077365243e-07 |
| 2128 | 24,5754920996466 | + | + | + | + | NA | NA | + | NA | NA | NA | NA | + | 0,047693495875223 | 8 | -526,756572487323 | 1070,63814497465 | 22,6765720593544 | 4.14211922463742e-07 |
| 1365 | 26,3052971542744 | NA | NA | + | NA | + | NA | NA | + | NA | NA | + | NA | 0,0318646484071305 | 7 | -527,885280102711 | 1070,63877725969 | 22,6772043443959 | 4.14080993159691e-07 |
| 2335 | 25,6346453413083 | NA | + | + | + | + | NA | NA | NA | + | NA | NA | + | 0,0476194041767979 | 8 | -526,761899304338 | 1070,64879860868 | 22,687225693385 | 4.12011357553931e-07 |
| 3289 | 25,6161942197181 | NA | NA | NA | + | + | NA | + | + | NA | NA | + | + | 0,0476040117105994 | 8 | -526,763005892557 | 1070,65101178511 | 22,6894388698242 | 4.11555682807967e-07 |
| 1200 | 25,4628999086732 | + | + | + | + | NA | + | NA | + | NA | NA | + | NA | 0,0633717884955014 | 9 | -525,61996112501 | 1070,65724508467 | 22,6956721693757 | 4.10275004630774e-07 |
| 396 | 26,5306376765663 | + | + | NA | + | NA | NA | NA | + | + | NA | NA | NA | 0,0317145391313665 | 7 | -527,895895418357 | 1070,66000789098 | 22,6984349756876 | 4.09708640723075e-07 |
| 3412 | 25,4151109505959 | + | + | NA | NA | + | NA | + | NA | + | NA | + | + | 0,0632612124542595 | 9 | -525,628043829847 | 1070,67341049434 | 22,7118375790499 | 4.06972238489069e-07 |
| 287 | 26,3686432923582 | NA | + | + | + | + | NA | NA | NA | + | NA | NA | NA | 0,031606287255603 | 7 | -527,90354967286 | 1070,67531639998 | 22,7137434846923 | 4.06584597877196e-07 |
| 3287 | 25,2350223448311 | NA | + | + | NA | + | NA | + | + | NA | NA | + | + | 0,0632447632838298 | 9 | -525,629246122654 | 1070,67581507995 | 22,7142421646638 | 4.06483232717065e-07 |
| 128 | 24,9204208769194 | + | + | + | + | + | + | + | NA | NA | NA | NA | NA | 0,0632328893220383 | 9 | -525,630113993967 | 1070,67755082258 | 22,7159779072892 | 4.06130610617723e-07 |
| 2168 | 24,8826988110737 | + | + | + | NA | + | + | + | NA | NA | NA | NA | + | 0,0631847355796855 | 9 | -525,633633452258 | 1070,68458973916 | 22,7230168238705 | 4.04703763210895e-07 |
| 504 | 25,6141866668902 | + | + | + | NA | + | + | + | + | + | NA | NA | NA | 0,0789640065007644 | 10 | -524,470570330282 | 1070,6871724066 | 22,7255994913046 | 4.04181492881257e-07 |
| 3415 | 25,304608678576 | NA | + | + | NA | + | NA | + | NA | + | NA | + | + | 0,0631635931044435 | 9 | -525,635178655297 | 1070,68768014524 | 22,7261072299482 | 4.04078896623349e-07 |
| 399 | 26,4845423057096 | NA | + | + | + | NA | NA | NA | + | + | NA | NA | NA | 0,0314942070007588 | 7 | -527,911473722616 | 1070,6911644995 | 22,7295915842055 | 4.03375532481835e-07 |
| 3509 | 26,077096308891 | NA | NA | + | NA | + | + | NA | + | + | NA | + | + | 0,0631008873110291 | 9 | -525,639761318754 | 1070,69684547215 | 22,7352725568628 | 4.02231375546869e-07 |
| 1359 | 25,5192598935074 | NA | + | + | + | NA | NA | + | NA | + | NA | + | NA | 0,0472781360126416 | 8 | -526,786429406365 | 1070,69785881273 | 22,7362858974384 | 4.02027628480713e-07 |
| 157 | 26,5686588375186 | NA | NA | + | + | + | NA | NA | + | NA | NA | NA | NA | 0,0155959394190378 | 6 | -529,026291703341 | 1070,69873725284 | 22,7371643375448 | 4.01851088657105e-07 |
| 3419 | 25,2906749278071 | NA | + | NA | + | NA | NA | + | NA | + | NA | + | + | 0,0630779689684746 | 9 | -525,641436160076 | 1070,7001951548 | 22,7386222395062 | 4.01558265651701e-07 |
| 3340 | 25,7001850671019 | + | + | NA | + | NA | NA | NA | NA | + | NA | + | + | 0,0472385534126001 | 8 | -526,789274005803 | 1070,70354801161 | 22,7419750963161 | 4.00885645925856e-07 |
| 1356 | 25,5442377151157 | + | + | NA | + | NA | NA | + | NA | + | NA | + | NA | 0,047189077326738 | 8 | -526,792829433475 | 1070,71065886695 | 22,7490859516593 | 3.9946285681787e-07 |
| 334 | 25,9252723690956 | + | NA | + | + | NA | NA | + | NA | + | NA | NA | NA | 0,0313400070598737 | 7 | -527,922374127183 | 1070,71296530863 | 22,7513923933386 | 3.99002453451112e-07 |
| 1165 | 26,5713019388485 | NA | NA | + | + | NA | NA | NA | + | NA | NA | + | NA | 0,0154893430446055 | 6 | -529,033705554745 | 1070,71356495564 | 22,751992040354 | 3.98882841068134e-07 |
| 2269 | 25,5007088750389 | NA | NA | + | + | + | NA | + | + | NA | NA | NA | + | 0,0470790930337031 | 8 | -526,800732413013 | 1070,72646482603 | 22,7648919107351 | 3.96318351879745e-07 |
| 2202 | 25,857718565913 | + | NA | NA | + | + | NA | NA | + | NA | NA | NA | + | 0,0311985784004315 | 7 | -527,932370204167 | 1070,7329574626 | 22,7713845473077 | 3.95033862425701e-07 |
| 2266 | 25,5521517388354 | + | NA | NA | + | NA | NA | + | + | NA | NA | NA | + | 0,0470322946488685 | 8 | -526,804094859201 | 1070,7331897184 | 22,771616803112 | 3.94987990635518e-07 |
| 1215 | 25,3900394354857 | NA | + | + | + | + | + | NA | + | NA | NA | + | NA | 0,0628301712688571 | 9 | -525,659542264622 | 1070,73640736389 | 22,7748344485999 | 3.94353035874384e-07 |
| 1352 | 25,5506996428634 | + | + | + | NA | NA | NA | + | NA | + | NA | + | NA | 0,0469835445446885 | 8 | -526,807597360172 | 1070,74019472034 | 22,7786218050542 | 3.93606964748819e-07 |
| 3463 | 26,0965619624103 | NA | + | + | NA | NA | NA | NA | + | + | NA | + | + | 0,0469286523828711 | 8 | -526,811540929198 | 1070,7480818584 | 22,7865089431061 | 3.92057805130846e-07 |

| | | | | | | | | | | | | | | | | | |
|---|---|---|---|---|---|---|---|---|---|---|---|---|---|---|---|---|---|
| **3277** | 25,6364549606172 | NA | NA | + | + | NA | NA | + | + | NA | NA | + | + | 0,046919359310553 | 8 | -526,812208540702 | 1070,7494170814 | 22,7878441661128 | 3.91796150181815e-07 |
| **3210** | 26,132946608086 | + | NA | NA | + | NA | NA | NA | + | NA | NA | + | + | 0,0310764458557169 | 7 | -527,9410012719 | 1070,75021959806 | 22,7886466827733 | 3.91638970249774e-07 |
| **1419** | 26,6918906067113 | NA | + | NA | + | NA | NA | NA | + | + | NA | + | NA | 0,0310080179226364 | 7 | -527,945836576661 | 1070,75989020759 | 22,798317292295 | 3.8974984739543e-07 |
| **1350** | 25,9492045872704 | + | NA | + | NA | NA | NA | + | NA | + | NA | + | NA | 0,0309920197440454 | 7 | -527,946967002457 | 1070,76215105918 | 22,800578143886 | 3.89309513043684e-07 |
| **1405** | 25,7012061976869 | NA | NA | + | + | + | + | + | NA | + | NA | + | NA | 0,0625863148952355 | 9 | -525,677355712951 | 1070,77203426055 | 22,8104613452574 | 3.8739044642275e-07 |
| **382** | 25,716606643089 | + | NA | + | + | + | + | + | NA | + | NA | NA | NA | 0,0625619902000032 | 9 | -525,679132351966 | 1070,77558753858 | 22,8140146232863 | 3.86702804468447e-07 |
| **2254** | 25,5825892793496 | + | NA | + | + | NA | NA | + | + | NA | NA | NA | + | 0,046673327444508 | 8 | -526,8298810295 | 1070,784762059 | 22,8231891437085 | 3.84932960550837e-07 |
| **411** | 26,6823101882255 | NA | + | NA | + | + | NA | NA | + | + | NA | NA | NA | 0,0307995257729235 | 7 | -527,960567097823 | 1070,78935124991 | 22,8277783346184 | 3.84050707722865e-07 |
| **1462** | 26,0948622371737 | + | NA | + | NA | + | + | NA | + | + | NA | + | NA | 0,0624413141235474 | 9 | -525,687945669102 | 1070,79321417285 | 22,831641257558 | 3.83309644476391e-07 |
| **3389** | 25,6748235332445 | NA | NA | + | + | + | + | NA | NA | + | NA | + | + | 0,0624065711312696 | 9 | -525,690482838444 | 1070,79828851153 | 22,8367155962437 | 3.8233835568071e-07 |
| **2366** | 25,6761753830511 | + | NA | + | + | + | + | NA | NA | + | NA | NA | + | 0,0624047001541826 | 9 | -525,690619467236 | 1070,79856176912 | 22,8369888538275 | 3.82286120821553e-07 |
| **154** | 26,6142561279415 | + | NA | NA | + | + | NA | NA | + | NA | NA | NA | NA | 0,0148533475690143 | 6 | -529,07792281096 | 1070,80199946807 | 22,8404265527829 | 3.81629592920407e-07 |
| **2422** | 25,6806772896151 | + | NA | + | NA | + | + | + | NA | + | NA | NA | + | 0,0623215600017919 | 9 | -525,696690532433 | 1070,81070389951 | 22,8491309842209 | 3.79972267741669e-07 |
| **30** | 26,1543380704103 | + | NA | + | + | + | NA | NA | NA | NA | NA | NA | NA | 0,0147851507131287 | 6 | -529,082662468126 | 1070,81147878241 | 22,8499058671159 | 3.79825079251542e-07 |
| **3274** | 25,7664266219111 | + | NA | NA | + | NA | NA | + | + | NA | NA | + | + | 0,0464814805657241 | 8 | -526,843658243245 | 1070,81231648649 | 22,8507435711997 | 3.79666022054578e-07 |
| **3445** | 25,6927158805916 | NA | NA | + | NA | + | + | + | NA | + | NA | + | + | 0,0623025224308379 | 9 | -525,698080619374 | 1070,81348407339 | 22,8519111581038 | 3.79444440202246e-07 |
| **1162** | 26,7028650113974 | + | NA | NA | + | NA | NA | NA | + | NA | NA | + | NA | 0,0146435823411898 | 6 | -529,092500372626 | 1070,83115459141 | 22,8695816761151 | 3.761067168324e-07 |
| **96** | 25,2982940811151 | + | + | + | + | + | NA | + | NA | NA | NA | NA | NA | 0,0463306427736842 | 8 | -526,854488501648 | 1070,8339770033 | 22,8724040880061 | 3.75576327130042e-07 |
| **3473** | 26,7241051347872 | NA | NA | NA | NA | + | NA | NA | + | + | NA | + | + | 0,0304222440047182 | 7 | -527,987215001571 | 1070,84264705741 | 22,8810741421155 | 3.73951717498378e-07 |
| **1136** | 24,920937574731 | + | + | + | + | NA | + | + | NA | NA | NA | + | NA | 0,062064682724777 | 9 | -525,715444841594 | 1070,84821251783 | 22,8866396025439 | 3.72912557277277e-07 |
| **3183** | 24,6554912589434 | NA | + | + | + | NA | + | + | NA | NA | NA | + | + | 0,0620350908578004 | 9 | -525,717604979361 | 1070,85253279337 | 22,8909598780776 | 3.72107884193825e-07 |
| **3437** | 25,4408802857392 | NA | NA | + | + | NA | + | + | NA | + | NA | + | + | 0,0620107636976395 | 9 | -525,719380754704 | 1070,85608434405 | 22,8945114287631 | 3.7144769053954e-07 |
| **2396** | 25,2875751884076 | + | + | NA | + | + | NA | + | NA | + | NA | NA | + | 0,0619537817834083 | 9 | -525,723540002809 | 1070,86440284026 | 22,9028299249726 | 3.69905955887689e-07 |
| **3458** | 26,7635050893508 | + | NA | NA | NA | NA | NA | NA | + | + | NA | + | + | 0,0301928916546868 | 7 | -528,003409392428 | 1070,87503583912 | 22,9134629238281 | 3.67944569559444e-07 |
| **1357** | 26,0329761649919 | NA | NA | + | + | NA | NA | + | NA | + | NA | + | NA | 0,0301699557554473 | 7 | -528,005028667855 | 1070,87827438997 | 22,9167014746836 | 3.67349248083984e-07 |
| **2392** | 25,2967713163637 | + | + | + | NA | + | NA | + | NA | + | NA | NA | + | 0,0618447942522034 | 9 | -525,731494563621 | 1070,88031196189 | 22,9187390465979 | 3.66975188404386e-07 |
| **2414** | 25,3984816983524 | + | NA | + | + | NA | + | + | NA | + | NA | NA | + | 0,0618146816372528 | 9 | -525,733692199211 | 1070,88470723307 | 22,9231343177771 | 3.66169596193131e-07 |
| **282** | 26,8398572517497 | + | NA | NA | + | + | NA | NA | NA | + | NA | NA | NA | 0,0142103295298712 | 6 | -529,122599309076 | 1070,89135246431 | 22,9297795490147 | 3.6495497435187e-07 |
| **2399** | 25,1941261625737 | NA | + | + | + | + | NA | + | NA | + | NA | NA | + | 0,0617687516892956 | 9 | -525,73704405689 | 1070,89141094842 | 22,9298380331343 | 3.64944302472727e-07 |
| **3500** | 25,7122331144826 | + | + | NA | + | NA | + | NA | + | + | NA | + | + | 0,0773817168294888 | 10 | -524,588093069782 | 1070,9222178856 | 22,9606449703047 | 3.59365967523542e-07 |
| **2440** | 25,9485706516623 | + | + | + | NA | NA | NA | NA | + | + | NA | NA | + | 0,045700785458233 | 8 | -526,899694200414 | 1070,92438840083 | 22,9628154855382 | 3.58976174421311e-07 |
| **3351** | 25,9504915113726 | NA | + | + | NA | + | NA | NA | NA | + | NA | + | + | 0,0456811691955525 | 8 | -526,90110160681 | 1070,92720321362 | 22,9656302983292 | 3.58471304419564e-07 |
| **1290** | 26,7126210573624 | + | NA | NA | + | NA | NA | NA | NA | + | NA | + | NA | 0,0139331513721764 | 6 | -529,141848493404 | 1070,92985083296 | 22,9682779176708 | 3.57997070606982e-07 |
| **2256** | 25,06069989422 | + | + | + | + | NA | NA | + | + | NA | NA | NA | + | 0,0615013008933241 | 9 | -525,756558718388 | 1070,93044027142 | 22,9688673561307 | 3.57891577532194e-07 |
| **1342** | 25,7785253564714 | + | NA | + | + | + | + | NA | NA | + | NA | + | NA | 0,0614904771588584 | 9 | -525,757348359758 | 1070,93201955416 | 22,97044663887 | 3.57609083086039e-07 |
| **3404** | 25,1397153392479 | + | + | NA | + | NA | NA | + | NA | + | NA | + | + | 0,0614557686992389 | 9 | -525,759880440637 | 1070,93708371592 | 22,9755108006282 | 3.56704733391044e-07 |
| **3503** | 25,6150395913304 | NA | + | + | + | NA | + | NA | + | + | NA | + | + | 0,0772753546444709 | 10 | -524,595985770694 | 1070,93800328742 | 22,9764303721283 | 3.56540763336081e-07 |
| **2175** | 24,6995754462948 | NA | + | + | + | + | + | + | NA | NA | NA | NA | + | 0,0613742367041558 | 9 | -525,765828063502 | 1070,94897896165 | 22,987406046358 | 3.54589484748727e-07 |
| **1398** | 25,7741506517569 | + | NA | + | NA | + | + | + | NA | + | NA | + | NA | 0,0613728685115731 | 9 | -525,76592786646 | 1070,94917856757 | 22,9876056522744 | 3.5455409743512e-07 |
| **2268** | 25,0281022481811 | + | + | NA | + | + | NA | + | + | NA | NA | NA | + | 0,0613177833625412 | 9 | -525,769945937929 | 1070,9572147105 | 22,9956417952128 | 3.53132332022301e-07 |
| **3285** | 25,7086922639136 | NA | NA | + | NA | + | NA | + | + | NA | NA | + | + | 0,0454474924408887 | 8 | -526,917864970261 | 1070,96072994052 | 22,9991570252309 | 3.52512206465985e-07 |
| **2397** | 25,6735218075208 | NA | NA | + | + | + | NA | + | NA | + | NA | NA | + | 0,0453449027049834 | 8 | -526,925223195796 | 1070,97544639159 | 23,013873476302 | 3.49927861902299e-07 |
| **2332** | 25,7766137461959 | + | + | NA | + | + | NA | NA | NA | + | NA | NA | + | 0,0453332619713602 | 8 | -526,926058074845 | 1070,97711614969 | 23,015543234399 | 3.49635836381863e-07 |
| **336** | 25,3831207662488 | + | + | + | + | NA | NA | + | NA | + | NA | NA | NA | 0,0453148900944219 | 8 | -526,927375694047 | 1070,97975138809 | 23,0181784728034 | 3.49175452861775e-07 |
| **3515** | 25,6840505979724 | NA | + | NA | + | + | + | NA | + | + | NA | + | + | 0,0769640015337511 | 10 | -524,619084780214 | 1070,98420130646 | 23,0226283911686 | 3.48399415377589e-07 |
| **3374** | 25,5441859949648 | + | NA | + | + | NA | + | NA | NA | + | NA | + | + | 0,0611213928242852 | 9 | -525,78426931896 | 1070,98586147257 | 23,0242885572752 | 3.48110334924096e-07 |
| **346** | 26,3534745652003 | + | NA | NA | + | + | NA | + | NA | + | NA | NA | NA | 0,029387619727136 | 7 | -528,0602387155 | 1070,98869448526 | 23,0271215699725 | 3.47617583499808e-07 |
| **3460** | 26,2712447877373 | + | + | NA | NA | NA | NA | NA | + | + | NA | + | + | 0,0452517708438675 | 8 | -526,931902373006 | 1070,98880474601 | 23,0272318307223 | 3.47598419740364e-07 |
| **3132** | 25,2329902104938 | + | + | NA | + | + | + | NA | NA | NA | NA | + | + | 0,0610806529974983 | 9 | -525,787240227897 | 1070,99180329044 | 23,0302303751498 | 3.47077665561776e-07 |
| **3282** | 25,7566537734036 | + | NA | NA | NA | + | NA | + | + | NA | NA | + | + | 0,0452144907840799 | 8 | -526,934575820362 | 1070,99415164072 | 23,0325787254333 | 3.46670374756788e-07 |
| **2455** | 26,0417157832314 | NA | + | + | NA | + | NA | NA | + | + | NA | NA | + | 0,0452130934509754 | 8 | -526,934676024624 | 1070,99435204925 | 23,0327791339564 | 3.46635638648259e-07 |
| **3430** | 25,5388882007744 | + | NA | + | NA | NA | + | + | NA | + | NA | + | + | 0,0610282845575114 | 9 | -525,79105895194 | 1070,99944073853 | 23,0378678232346 | 3.45754799174134e-07 |
| **2262** | 25,6024584239601 | + | NA | + | NA | + | NA | + | + | NA | NA | NA | + | 0,0451143515592723 | 8 | -526,941756540872 | 1071,00851308174 | 23,0469401664529 | 3.44189947970494e-07 |
| **3475** | 26,280968565009 | NA | + | NA | NA | + | NA | NA | + | + | NA | + | + | 0,0450521391376368 | 8 | -526,94621725099 | 1071,01743450198 | 23,0558615866894 | 3.42658035635574e-07 |
| **1053** | 26,387535777307 | NA | NA | + | + | + | NA | NA | NA | NA | NA | + | NA | 0,0132870645961993 | 6 | -529,186696276882 | 1071,01954639992 | 23,0579734846265 | 3.42296397205749e-07 |
| **3400** | 25,1855916951326 | + | + | + | NA | NA | NA | + | NA | + | NA | + | + | 0,0608839111802713 | 9 | -525,801585604253 | 1071,02049404315 | 23,0589211278614 | 3.42134248191066e-07 |
| **3336** | 25,771283482373 | + | + | + | NA | NA | NA | NA | NA | + | NA | + | + | 0,044987995840035 | 8 | -526,950816103258 | 1071,02663220652 | 23,0650592912257 | 3.41085819916063e-07 |
| **2320** | 25,4237286774875 | + | + | + | + | NA | NA | NA | NA | + | NA | NA | + | 0,0449655980987368 | 8 | -526,952421871006 | 1071,02984374201 | 23,0682708267213 | 3.40538554815116e-07 |
| **150** | 26,471537888091 | + | NA | + | NA | + | NA | NA | + | NA | NA | NA | NA | 0,0132051475898973 | 6 | -529,192380407168 | 1071,03091466049 | 23,0693417451989 | 3.40356259110091e-07 |
| **2236** | 25,2275969705327 | + | + | NA | + | + | + | NA | + | NA | NA | NA | + | 0,0607933963468926 | 9 | -525,808184459947 | 1071,03369175454 | 23,0721188392483 | 3.39883986401478e-07 |
| **3270** | 25,7189054663689 | + | NA | + | NA | NA | NA | + | + | NA | NA | + | + | 0,0449255689853143 | 8 | -526,955291596068 | 1071,03558319214 | 23,0740102768452 | 3.39562703671751e-07 |
| **1369** | 26,3870662064089 | NA | NA | NA | + | + | NA | + | NA | + | NA | + | NA | 0,0290162353646092 | 7 | -528,086432024254 | 1071,04108110277 | 23,0795081874808 | 3.38630542793088e-07 |
| **1158** | 26,4801768017267 | + | NA | + | NA | NA | NA | NA | + | NA | NA | + | NA | 0,0131154769110269 | 6 | -529,198602015499 | 1071,04335787715 | 23,081784961862 | 3.38245269457541e-07 |
| **280** | 26,43528216693 | + | + | + | NA | + | NA | NA | NA | + | NA | NA | NA | 0,0289941473907235 | 7 | -528,087989547942 | 1071,04419615015 | 23,0826232348575 | 3.38103528226465e-07 |
| **1307** | 26,5803693188753 | NA | + | NA | + | + | NA | NA | NA | + | NA | + | NA | 0,0289316235877522 | 7 | -528,0923981942 | 1071,05301344266 | 23,0914405273736 | 3.36616230266149e-07 |
| **3506** | 26,2555988634333 | + | NA | NA | NA | + | + | NA | + | + | NA | + | + | 0,0606411480641065 | 9 | -525,819282471633 | 1071,05588777791 | 23,0943148626209 | 3.36132803784072e-07 |
| **284** | 26,4401567882863 | + | + | NA | + | + | NA | NA | NA | + | NA | NA | NA | 0,02886770567299 | 7 | -528,096904848072 | 1071,06202675041 | 23,100453835116 | 3.35102630625507e-07 |
| **3407** | 25,0753685178197 | NA | + | + | + | NA | NA | + | NA | + | NA | + | + | 0,0605921714216831 | 9 | -525,822852200958 | 1071,06302723656 | 23,1014543212716 | 3.34935039775762e-07 |
| **1038** | 26,0435842175444 | + | NA | + | + | NA | NA | NA | NA | NA | NA | + | NA | 0,0129593868173351 | 6 | -529,209430645158 | 1071,06501513647 | 23,1034422211799 | 3.34602296500814e-07 |
| **3240** | 25,4843016635212 | + | + | + | NA | NA | + | NA | + | NA | NA | + | + | 0,0604930863092526 | 9 | -525,830073585571 | 1071,07747000579 | 23,1158970904976 | 3.32525057211728e-07 |
| **94** | 25,80590878605 | + | NA | + | + | + | NA | + | NA | NA | NA | NA | NA | 0,0287479021161018 | 7 | -528,105351026594 | 1071,07891910745 | 23,1173461921605 | 3.32284213167451e-07 |
| **3447** | 25,2967199525553 | NA | + | + | NA | + | + | + | NA | + | NA | + | + | 0,0763137269215878 | 10 | -524,66730296564 | 1071,08063767731 | 23,119064762021 | 3.31998808989834e-07 |
| **3444** | 25,4293813430435 | + | + | NA | NA | + | + | + | NA | + | NA | + | + | 0,0763028161822641 | 10 | -524,668111712925 | 1071,08225517188 | 23,1206822565914 | 3.31730414400704e-07 |
| **1390** | 25,5591491934379 | + | NA | + | + | NA | + | + | NA | + | NA | + | NA | 0,0604553264570829 | 9 | -525,832825346754 | 1071,08297352815 | 23,1214006128635 | 3.31611285484329e-07 |
| **2390** | 25,7260864229549 | + | NA | + | NA | + | NA | + | NA | + | NA | NA | + | 0,044578795914008 | 8 | -526,980147055325 | 1071,08529411065 | 23,1237211953594 | 3.31226742945866e-07 |
| **2271** | 25,0087527873227 | NA | + | + | + | + | NA | + | + | NA | NA | NA | + | 0,0604068036993483 | 9 | -525,836361295799 | 1071,09004542624 | 23,1284725109538 | 3.30440795492088e-07 |

| | | | | | | | | | | | | | | | | | | | |
|---|---|---|---|---|---|---|---|---|---|---|---|---|---|---|---|---|---|---|---|
| **2224** | 25,159348092244 | + | + | + | + | NA | + | NA | + | NA | NA | NA | + | 0,0603835646430168 | 9 | -525,838054707023 | 1071,09343224869 | 23,1318593334015 | 3.29881696865731e-07 |
| **3101** | 25,6404171178522 | NA | NA | + | + | + | NA | NA | NA | NA | NA | + | + | 0,0286263583411009 | 7 | -528,113918826798 | 1071,09605470786 | 23,1344817925692 | 3.2944942968832e-07 |
| **1177** | 26,8672693002196 | NA | NA | NA | + | + | NA | NA | + | NA | NA | + | NA | 0,0127289652445405 | 6 | -529,225412835723 | 1071,0969795176 | 23,1354066023082 | 3.29297125883504e-07 |
| **3255** | 25,4481224835657 | NA | + | + | NA | + | + | NA | + | NA | NA | + | + | 0,0603560990821105 | 9 | -525,84005604624 | 1071,09743492712 | 23,135862011834 | 3.29222151895824e-07 |
| **3382** | 25,814034016982 | + | NA | + | NA | + | + | NA | NA | + | NA | + | + | 0,060330530456683 | 9 | -525,841919108478 | 1071,1011610516 | 23,139588136311 | 3.28609361547079e-07 |
| **3451** | 25,2797233929327 | NA | + | NA | + | + | + | + | NA | + | NA | + | + | 0,0761710455051714 | 10 | -524,677878324344 | 1071,10178839472 | 23,1402154794287 | 3.28506302300614e-07 |
| **2426** | 25,7644076356324 | + | NA | NA | + | + | + | + | NA | + | NA | NA | + | 0,0603134588553534 | 9 | -525,843163005401 | 1071,10364884545 | 23,142075930157 | 3.28200859493268e-07 |
| **1362** | 26,3818124667678 | + | NA | NA | NA | + | NA | + | NA | + | NA | + | NA | 0,0285648868854646 | 7 | -528,118251632476 | 1071,10472031921 | 23,1431474039241 | 3.28025077278342e-07 |
| **3212** | 25,6572933149896 | + | + | NA | + | NA | NA | NA | + | NA | NA | + | + | 0,0444418479927582 | 8 | -526,989960511153 | 1071,10492102231 | 23,1433481070146 | 3.27992161106576e-07 |
| **3413** | 25,7693665380277 | NA | NA | + | NA | + | NA | + | NA | + | NA | + | + | 0,0444063146768779 | 8 | -526,99250653862 | 1071,11001307724 | 23,1484401619484 | 3.271581462184e-07 |
| **3405** | 25,5600450916223 | NA | NA | + | + | NA | NA | + | NA | + | NA | + | + | 0,0444019990895887 | 8 | -526,992815751945 | 1071,11063150389 | 23,1490585886002 | 3.27057000198579e-07 |
| **3252** | 25,5786264695327 | + | + | NA | NA | + | + | NA | + | NA | NA | + | + | 0,0602471690649531 | 9 | -525,847992898572 | 1071,11330863179 | 23,1517357164985 | 3.26619506360856e-07 |
| **1404** | 25,4844527452089 | + | + | NA | + | + | + | + | NA | + | NA | + | NA | 0,076024720263852 | 10 | -524,688722063067 | 1071,12347587217 | 23,1619029568753 | 3.2496331014742e-07 |
| **1117** | 25,8821606774854 | NA | NA | + | + | + | NA | + | NA | NA | NA | + | NA | 0,0284234871229257 | 7 | -528,128217131895 | 1071,12465131805 | 23,1630784027629 | 3.24772377867288e-07 |
| **1354** | 26,149168580322 | + | NA | NA | + | NA | NA | + | NA | + | NA | + | NA | 0,0284201127557119 | 7 | -528,128454931093 | 1071,12512691645 | 23,1635540011598 | 3.2469515643812e-07 |
| **3449** | 25,7457562601816 | NA | NA | NA | + | + | + | + | NA | + | NA | + | + | 0,0601494454635961 | 9 | -525,855112447227 | 1071,1275477291 | 23,1659748138086 | 3.24302381124084e-07 |
| **3215** | 25,5591013820894 | NA | + | + | + | NA | NA | NA | + | NA | NA | + | + | 0,0442812086843781 | 8 | -527,001469860142 | 1071,12793972028 | 23,1663668049937 | 3.24238825515244e-07 |
| **2328** | 25,8294001061734 | + | + | + | NA | + | NA | NA | NA | + | NA | NA | + | 0,0442404660440961 | 8 | -527,00438864689 | 1071,13377729378 | 23,1722043784896 | 3.232938213319e-07 |
| **2382** | 25,4991060406156 | + | NA | + | + | NA | NA | + | NA | + | NA | NA | + | 0,0442293240562626 | 8 | -527,005186832866 | 1071,13537366573 | 23,1738007504423 | 3.23035875695462e-07 |
| **392** | 26,4977652115845 | + | + | + | NA | NA | NA | NA | + | + | NA | NA | NA | 0,0283407388336815 | 7 | -528,134048350677 | 1071,13631375562 | 23,1747408403264 | 3.22884069996532e-07 |
| **2495** | 25,5725246215274 | NA | + | + | + | + | + | NA | + | + | NA | NA | + | 0,0759319642165375 | 10 | -524,695595055518 | 1071,13722185707 | 23,1756489417774 | 3.22737497528495e-07 |
| **1088** | 25,2910309484633 | + | + | + | + | + | + | NA | NA | NA | NA | + | NA | 0,0600800301548771 | 9 | -525,860169175991 | 1071,13766118663 | 23,1760882713368 | 3.22666611253091e-07 |
| **3227** | 25,5935751703735 | NA | + | NA | + | + | NA | NA | + | NA | NA | + | + | 0,0441906897728065 | 8 | -527,007954430633 | 1071,14090886127 | 23,1793359459762 | 3.22143078349091e-07 |
| **1407** | 25,3492178326668 | NA | + | + | + | + | + | + | NA | + | NA | + | NA | 0,0759062421888128 | 10 | -524,697500871684 | 1071,1410334894 | 23,1794605741095 | 3.22123004929265e-07 |
| **2428** | 25,2901858582972 | + | + | NA | + | + | + | + | NA | + | NA | NA | + | 0,0759042236098312 | 10 | -524,697650431539 | 1071,14133260911 | 23,1797596938197 | 3.22074831861787e-07 |
| **2480** | 25,5716567601326 | + | + | + | + | NA | + | NA | + | + | NA | NA | + | 0,0758963972095533 | 10 | -524,698230299392 | 1071,14249234482 | 23,1809194295251 | 3.21888125158463e-07 |
| **192** | 25,3005027488393 | + | + | + | + | + | + | NA | + | NA | NA | NA | NA | 0,0600117877663288 | 9 | -525,865140096645 | 1071,14760302794 | 23,1860301126451 | 3.21066641087787e-07 |
| **2264** | 25,1601811115822 | + | + | + | NA | + | NA | + | + | NA | NA | NA | + | 0,0600076820328962 | 9 | -525,865439155492 | 1071,14820114563 | 23,1866282303397 | 3.20970637624254e-07 |
| **1110** | 25,8442230658284 | + | NA | + | NA | + | NA | + | NA | NA | NA | + | NA | 0,0282382511099569 | 7 | -528,141269906652 | 1071,15075686757 | 23,1891839522766 | 3.20560743723898e-07 |
| **3086** | 25,5588119677605 | + | NA | + | + | NA | NA | NA | NA | NA | NA | + | + | 0,0282217973206432 | 7 | -528,142429213227 | 1071,15307548072 | 23,1915025654266 | 3.20189330878286e-07 |
| **3321** | 25,4498130998659 | NA | NA | NA | + | + | + | + | + | NA | NA | + | + | 0,0599560729199894 | 9 | -525,869198217033 | 1071,15571926871 | 23,1941463534201 | 3.19766354150762e-07 |
| **3206** | 26,0044624999935 | + | NA | + | NA | NA | NA | NA | + | NA | NA | + | + | 0,02816953087097 | 7 | -528,146111690023 | 1071,16044043431 | 23,198867519018 | 3.19012409420063e-07 |
| **1471** | 25,7553895854427 | NA | + | + | + | + | + | NA | + | + | NA | + | NA | 0,0757580412340704 | 10 | -524,708480457523 | 1071,16299266108 | 23,2014197457875 | 3.18605573054381e-07 |
| **1301** | 27,1042990058638 | NA | NA | + | NA | + | NA | NA | NA | + | NA | + | NA | 0,0121854709381182 | 6 | -529,263095191357 | 1071,17234422887 | 23,2107713135763 | 3.17119319654901e-07 |
| **402** | 27,2787229472786 | + | NA | NA | NA | + | NA | NA | + | + | NA | NA | NA | 0,012180699348267 | 6 | -529,263425930543 | 1071,17300570724 | 23,2114327919496 | 3.17014453211719e-07 |
| **2492** | 25,6754128933999 | + | + | NA | + | + | + | NA | + | + | NA | NA | + | 0,0756886984604103 | 10 | -524,713617167872 | 1071,17326608178 | 23,2116931664855 | 3.16973184652541e-07 |
| **3434** | 25,6660777222417 | + | NA | NA | + | NA | + | + | NA | + | NA | + | + | 0,059820985402139 | 9 | -525,879036632503 | 1071,17539609965 | 23,2138231843605 | 3.16635785076601e-07 |
| **1295** | 26,2356743345413 | NA | + | + | + | NA | NA | NA | NA | + | NA | + | NA | 0,0280443779102517 | 7 | -528,154928642958 | 1071,17807434018 | 23,2165014248894 | 3.16212055456138e-07 |
| **384** | 25,3521861529246 | + | + | + | + | + | + | + | NA | + | NA | NA | NA | 0,0756387676593318 | 10 | -524,717315657407 | 1071,18066306084 | 23,2190901455542 | 3.15803027886567e-07 |
| **2239** | 25,1240607940314 | NA | + | + | + | + | + | NA | + | NA | NA | NA | + | 0,0597704426422144 | 9 | -525,882717295324 | 1071,18275742529 | 23,2211845100028 | 3.15472497662281e-07 |
| **2424** | 25,3059722021967 | + | + | + | NA | + | + | + | NA | + | NA | NA | + | 0,0756113990098242 | 10 | -524,719342831641 | 1071,18471740931 | 23,2231444940232 | 3.15163488573075e-07 |
| **2431** | 25,1793861199851 | NA | + | + | + | + | + | + | NA | + | NA | NA | + | 0,0756043525874791 | 10 | -524,719864744863 | 1071,18576123576 | 23,2241883204672 | 3.14999043498068e-07 |
| **272** | 25,9880276024254 | + | + | + | + | NA | NA | NA | NA | + | NA | NA | NA | 0,0279883323430886 | 7 | -528,158876652538 | 1071,18597035934 | 23,2243974440491 | 3.14966108355832e-07 |
| **3306** | 25,5852232572602 | + | NA | NA | + | NA | + | + | + | NA | NA | + | + | 0,0597265683824105 | 9 | -525,885912179277 | 1071,1891471932 | 23,2275742779095 | 3.14466207987095e-07 |
| **1456** | 25,7689912091821 | + | + | + | + | NA | + | NA | + | + | NA | + | NA | 0,0755795268011357 | 10 | -524,721703505335 | 1071,1894387567 | 23,2278658414111 | 3.14420367894133e-07 |
| **3309** | 25,4486382982624 | NA | NA | + | + | NA | + | + | + | NA | NA | + | + | 0,0596992553325265 | 9 | -525,887901015636 | 1071,19312486592 | 23,2315519506262 | 3.13841407677912e-07 |
| **1305** | 27,0086139034646 | NA | NA | NA | + | + | NA | NA | NA | + | NA | + | NA | 0,0119793412503134 | 6 | -529,277381461336 | 1071,20091676883 | 23,2393438535357 | 3.12621075510111e-07 |
| **1400** | 25,4902722340327 | + | + | + | NA | + | + | + | NA | + | NA | + | NA | 0,0755004678870174 | 10 | -524,727558797903 | 1071,20114934184 | 23,2395764265482 | 3.12584724011103e-07 |
| **3398** | 25,6134194153121 | + | NA | + | NA | NA | NA | + | NA | + | NA | + | + | 0,0437478146643666 | 8 | -527,039672157716 | 1071,20434431543 | 23,242771400141 | 3.12085772681924e-07 |
| **1413** | 27,1724613053321 | NA | NA | + | NA | NA | NA | NA | + | + | NA | + | NA | 0,0119221948580802 | 6 | -529,281341589891 | 1071,20883702594 | 23,2472641106456 | 3.11385503987896e-07 |
| **3135** | 25,0582084969503 | NA | + | + | + | + | + | NA | NA | NA | NA | + | + | 0,0595735555802779 | 9 | -525,897053267364 | 1071,21142936937 | 23,2498564540831 | 3.10982156364356e-07 |
| **1468** | 25,8654264937284 | + | + | NA | + | + | + | NA | + | + | NA | + | NA | 0,0754022452430945 | 10 | -524,73483270509 | 1071,21569715621 | 23,2541242409206 | 3.10319261112982e-07 |
| **1410** | 27,3158671230833 | + | NA | NA | NA | NA | NA | NA | + | + | NA | + | NA | 0,011819263067651 | 6 | -529,288473975745 | 1071,22310179764 | 23,2615288823536 | 3.09172483862357e-07 |
| **1046** | 26,3428720060613 | + | NA | + | NA | + | NA | NA | NA | NA | NA | + | NA | 0,0118055810295841 | 6 | -529,289421980428 | 1071,22499780701 | 23,2634248917182 | 3.08879525784773e-07 |
| **2298** | 25,4059977753657 | + | NA | NA | + | + | + | + | + | NA | NA | NA | + | 0,059429979852614 | 9 | -525,907505580542 | 1071,23233399573 | 23,270761080438 | 3.07748601974016e-07 |
| **405** | 27,275611366234 | NA | NA | + | NA | + | NA | NA | + | + | NA | NA | NA | 0,0117376161980454 | 6 | -529,294130951254 | 1071,23441574866 | 23,272842833371 | 3.07428440349878e-07 |
| **2286** | 25,4008960671542 | + | NA | + | + | NA | + | + | + | NA | NA | NA | + | 0,0594089548528964 | 9 | -525,909036066681 | 1071,23539496801 | 23,2738220527176 | 3.07277957253628e-07 |
| **3346** | 26,4075088874791 | + | NA | NA | NA | + | NA | NA | NA | + | NA | + | + | 0,0276031308915681 | 7 | -528,186005186686 | 1071,24022742764 | 23,2786545123447 | 3.06536399340213e-07 |
| **3221** | 26,0998106068261 | NA | NA | + | NA | + | NA | NA | + | NA | NA | + | + | 0,0275379109484374 | 7 | -528,190597360056 | 1071,24941177438 | 23,287838859085 | 3.05131958236774e-07 |
| **2078** | 25,5398257154299 | + | NA | + | + | + | NA | NA | NA | NA | NA | NA | + | 0,0275102673284857 | 7 | -528,192543670201 | 1071,25330439467 | 23,2917314793758 | 3.04538654374728e-07 |
| **2198** | 25,8684605356957 | + | NA | + | NA | + | NA | NA | + | NA | NA | NA | + | 0,0274521016150523 | 7 | -528,196638775191 | 1071,26149460465 | 23,2999216893559 | 3.03294086668524e-07 |
| **1499** | 25,9099482864759 | NA | + | NA | + | + | NA | + | + | + | NA | + | NA | 0,0591846801130144 | 9 | -525,925359712561 | 1071,26804225977 | 23,3064693444778 | 3.02302777701217e-07 |
| **2452** | 26,1755255275384 | + | + | NA | NA | + | NA | NA | + | + | NA | NA | + | 0,0432990073702516 | 8 | -527,07179975896 | 1071,26859951792 | 23,3070266026302 | 3.02218559090953e-07 |
| **2301** | 25,3180648754293 | NA | NA | + | + | + | + | + | + | NA | NA | NA | + | 0,0591715501342182 | 9 | -525,92631524636 | 1071,26995332737 | 23,3083804120752 | 3.02014055143739e-07 |
| **448** | 25,7178214381632 | + | + | + | + | + | + | NA | + | + | NA | NA | NA | 0,0749359978272908 | 10 | -524,769350265064 | 1071,28473227616 | 23,3231593608693 | 2.99790555380697e-07 |
| **1208** | 25,6044607271982 | + | + | + | NA | + | + | NA | + | NA | NA | + | NA | 0,0590617519364393 | 9 | -525,93430528505 | 1071,28593340474 | 23,324360489454 | 2.99610565930991e-07 |
| **1292** | 26,2989412543765 | + | + | NA | + | NA | NA | NA | NA | + | NA | + | NA | 0,0272400082967334 | 7 | -528,21156894189 | 1071,29135493804 | 23,3297820227531 | 2.9879949141455e-07 |
| **3120** | 25,0036234629021 | + | + | + | + | NA | + | NA | NA | NA | NA | + | + | 0,0590098015557223 | 9 | -525,938085400757 | 1071,29349363616 | 23,3319207208694 | 2.98480141238657e-07 |
| **1402** | 25,9008446923089 | + | NA | NA | + | + | + | + | NA | + | NA | + | NA | 0,0590097059492797 | 9 | -525,938092357269 | 1071,29350754918 | 23,3319346338926 | 2.98478064865319e-07 |
| **3348** | 26,09011418051 | + | + | NA | NA | + | NA | NA | NA | + | NA | + | + | 0,0429428606396013 | 8 | -527,097283590624 | 1071,31956718125 | 23,3579942659567 | 2.94614178123384e-07 |
| **3436** | 25,1265014443144 | + | + | NA | + | NA | + | + | NA | + | NA | + | + | 0,0746727079827279 | 10 | -524,788834644291 | 1071,32370103461 | 23,3621281193239 | 2.94005861107162e-07 |
| **476** | 25,8566473496379 | + | + | NA | + | + | NA | + | + | + | NA | NA | NA | 0,0586641191925686 | 9 | -525,963233309521 | 1071,34378945369 | 23,3822165383961 | 2.91067585663046e-07 |
| **479** | 25,8568107406593 | NA | + | + | + | + | NA | + | + | + | NA | NA | NA | 0,0586570653087421 | 9 | -525,963746373603 | 1071,34481558185 | 23,383242666562 | 2.90918287642132e-07 |
| **1484** | 25,8532760144792 | + | + | NA | + | NA | NA | + | + | + | NA | + | NA | 0,0586461025209769 | 9 | -525,9645437441 | 1071,34641032285 | 23,3848374075556 | 2.90686410440892e-07 |
| **1487** | 25,8958559337234 | NA | + | + | + | NA | NA | + | + | + | NA | + | NA | 0,0586281586743254 | 9 | -525,965848856874 | 1071,34902054839 | 23,3874476331021 | 2.90307279351844e-07 |
| **3386** | 25,891323084141 | + | NA | NA | + | + | + | NA | NA | + | NA | + | + | 0,058626628375662 | 9 | -525,965960159189 | 1071,34924315302 | 23,3876702377333 | 2.90274969277529e-07 |

| | | | | | | | | | | | | | | | | | | |
|---|---|---|---|---|---|---|---|---|---|---|---|---|---|---|---|---|---|---|
| **3308** | 25,044221223802 | + | + | NA | + | NA | + | + | + | NA | NA | + | + | 0,074454450877651 | 10 | -524,80498224129 | 1071,35599622861 | 23,3944233133211 | 2.89296497725863e-07 |
| **1262** | 25,5932974539083 | + | NA | + | + | NA | + | + | + | NA | NA | + | NA | 0,0585445810713423 | 9 | -525,971927392912 | 1071,36117762047 | 23,3996047051789 | 2.88547988465549e-07 |
| **1277** | 25,5518823096186 | NA | NA | + | + | + | + | + | + | NA | NA | + | NA | 0,0585338153050309 | 9 | -525,972710339764 | 1071,36274351417 | 23,4011705988821 | 2.88322159144163e-07 |
| **1164** | 26,2741444256641 | + | + | NA | + | NA | NA | NA | + | NA | NA | + | NA | 0,0266739918191155 | 7 | -528,251397368691 | 1071,37101179165 | 23,4094388763551 | 2.87132655817502e-07 |
| **1274** | 25,6394782970028 | + | NA | NA | + | + | + | + | + | NA | NA | + | NA | 0,0584617992280923 | 9 | -525,977947522275 | 1071,3732178792 | 23,4116449639059 | 2.86816110542531e-07 |
| **3128** | 25,3347972701664 | + | + | + | NA | + | + | NA | NA | NA | NA | + | + | 0,0584159149384129 | 9 | -525,98128412948 | 1071,37989109361 | 23,4183181783151 | 2.85860712621866e-07 |
| **1167** | 26,2463491762417 | NA | + | + | + | NA | NA | NA | + | NA | NA | + | NA | 0,02660717852947 | 7 | -528,256097238019 | 1071,3804115303 | 23,4188386150111 | 2.8578633609696e-07 |
| **3442** | 25,8902611499684 | + | NA | NA | NA | + | + | + | NA | + | NA | + | + | 0,0583859697267059 | 9 | -525,983461593454 | 1071,38424602155 | 23,4226731062629 | 2.8523893840972e-07 |
| **1179** | 26,3934609347914 | NA | + | NA | + | + | NA | NA | + | NA | NA | + | NA | 0,0264495261438233 | 7 | -528,267185753951 | 1071,40258856217 | 23,4410156468759 | 2.82634894451763e-07 |
| **1104** | 25,2571152630533 | + | + | + | + | NA | NA | + | NA | NA | NA | + | NA | 0,0423486339200604 | 8 | -527,139781976099 | 1071,4045639522 | 23,4429910369076 | 2.82355875190857e-07 |
| **254** | 25,5030245488015 | + | NA | + | + | + | + | + | + | NA | NA | NA | NA | 0,0581832903417848 | 9 | -525,99819759136 | 1071,41371801737 | 23,4521451020751 | 2.81066476211469e-07 |
| **3417** | 25,8306651746588 | NA | NA | NA | + | + | NA | + | NA | + | NA | + | + | 0,0421721081581181 | 8 | -527,152401809738 | 1071,42980361948 | 23,4682307041862 | 2.78814980761692e-07 |
| **1288** | 26,2894885960705 | + | + | + | NA | NA | NA | NA | NA | + | NA | + | NA | 0,0262426848903434 | 7 | -528,281731258074 | 1071,43167957041 | 23,4701066551215 | 2.78553581761813e-07 |
| **2394** | 25,8258482070435 | + | NA | NA | + | + | NA | + | NA | + | NA | NA | + | 0,0421517055045845 | 8 | -527,153860246363 | 1071,43272049273 | 23,4711475774352 | 2.78408643163135e-07 |
| **3164** | 25,0425996583158 | + | + | NA | + | + | NA | + | NA | NA | NA | + | + | 0,0579991876587986 | 9 | -526,011580205445 | 1071,44048324554 | 23,4789103302453 | 2.77330128838618e-07 |
| **3432** | 25,1746433602521 | + | + | + | NA | NA | + | + | NA | + | NA | + | + | 0,0738233800678213 | 10 | -524,851650163926 | 1071,44933207388 | 23,4877591585923 | 2.76105815916769e-07 |
| **464** | 25,8087630327512 | + | + | + | + | NA | NA | + | + | + | NA | NA | NA | 0,0578959068929573 | 9 | -526,019086646314 | 1071,45549612727 | 23,4939232119825 | 2.75256160424462e-07 |
| **3402** | 25,7664676439804 | + | NA | NA | + | NA | NA | + | NA | + | NA | + | + | 0,0419654090763169 | 8 | -527,167175780768 | 1071,45935156154 | 23,4977786462448 | 2.74726055518874e-07 |
| **1102** | 25,6454390201516 | + | NA | + | + | NA | NA | + | NA | NA | NA | + | NA | 0,0260055870751792 | 7 | -528,2984006676 | 1071,46501838946 | 23,5034454741724 | 2.73948744618763e-07 |
| **3317** | 25,5417488970832 | NA | NA | + | NA | + | + | + | + | NA | NA | + | + | 0,0577827977289485 | 9 | -526,027306471056 | 1071,47193577676 | 23,5103628614675 | 2.73002876512922e-07 |
| **3218** | 26,2465533182739 | + | NA | NA | NA | + | NA | NA | + | NA | NA | + | + | 0,0258726222272961 | 7 | -528,307747125086 | 1071,48371130444 | 23,5221383891453 | 2.71400222702009e-07 |
| **159** | 26,1103485823866 | NA | + | + | + | + | NA | NA | + | NA | NA | NA | NA | 0,0258675302548896 | 7 | -528,308105028196 | 1071,48442711066 | 23,5228541953659 | 2.71303105098555e-07 |
| **1055** | 26,0808511379933 | NA | + | + | + | + | NA | NA | NA | NA | NA | + | NA | 0,0258410436639534 | 7 | -528,309966680016 | 1071,4881504143 | 23,5265774990055 | 2.70798503021651e-07 |
| **3439** | 25,0420581046585 | NA | + | + | + | NA | + | + | NA | + | NA | + | + | 0,0735462089980495 | 10 | -524,872137023888 | 1071,49030579381 | 23,528732878518 | 2.70506823447213e-07 |
| **1114** | 26,0263061606563 | + | NA | NA | + | + | NA | + | NA | NA | NA | + | NA | 0,0258082936912221 | 7 | -528,312268493551 | 1071,49275404136 | 23,5311811260742 | 2.70175892204215e-07 |
| **1492** | 25,9820365326767 | + | + | NA | NA | + | NA | + | + | + | NA | + | NA | 0,0575787145847657 | 9 | -526,042135026223 | 1071,50159288709 | 23,5400199718017 | 2.68984505249459e-07 |
| **2384** | 24,9794289160325 | + | + | + | + | NA | NA | + | NA | + | NA | NA | + | 0,0575786351647394 | 9 | -526,042140796209 | 1071,50160442706 | 23,5400315117722 | 2.68982953217308e-07 |
| **3302** | 25,5764220517484 | + | NA | + | NA | NA | + | + | + | NA | NA | + | + | 0,0575574054870594 | 9 | -526,04368314703 | 1071,50468912871 | 23,5431162134155 | 2.68568406908506e-07 |
| **1303** | 26,8013567027965 | NA | + | + | NA | + | NA | NA | NA | + | NA | + | NA | 0,0256786646270404 | 7 | -528,321378640992 | 1071,51097433625 | 23,5494014209578 | 2.67725727617224e-07 |
| **3094** | 25,797141680482 | + | NA | + | NA | + | NA | NA | NA | NA | NA | + | + | 0,0256761660658133 | 7 | -528,321554224443 | 1071,51132550315 | 23,5497525878598 | 2.67678723536742e-07 |
| **3098** | 25,858955207169 | + | NA | NA | + | + | NA | NA | NA | NA | NA | + | + | 0,0256278402412375 | 7 | -528,324950176383 | 1071,51811740703 | 23,5565444917395 | 2.66771241211596e-07 |
| **3314** | 25,6349456283613 | + | NA | NA | NA | + | + | + | + | NA | NA | + | + | 0,0574498513762658 | 9 | -526,051496494252 | 1071,52031582315 | 23,5587429078596 | 2.66478165218321e-07 |
| **1495** | 25,99731868637 | NA | + | + | NA | + | NA | + | + | + | NA | + | NA | 0,057421688470772 | 9 | -526,05354226194 | 1071,52440735853 | 23,5628344432357 | 2.65933570045883e-07 |
| **1050** | 26,5221460668174 | + | NA | NA | + | + | NA | NA | NA | NA | NA | + | NA | 0,0096135527002876 | 6 | -529,441134640258 | 1071,52842312667 | 23,5668502113788 | 2.65400141976578e-07 |
| **407** | 26,8059545264228 | NA | + | + | NA | + | NA | NA | + | + | NA | NA | NA | 0,0255528935843592 | 7 | -528,330216494236 | 1071,52865004274 | 23,5670771274458 | 2.65370031906519e-07 |
| **2160** | 24,4940174146795 | + | + | + | + | NA | + | + | NA | NA | NA | NA | + | 0,0573843468384824 | 9 | -526,056254682774 | 1071,52983220019 | 23,568259284903 | 2.65213223673029e-07 |
| **2446** | 26,1208629130618 | + | NA | + | + | NA | NA | NA | + | + | NA | NA | + | 0,0414080466192999 | 8 | -527,20699780491 | 1071,53899560982 | 23,5774226945296 | 2.64000874403624e-07 |
| **3391** | 25,4308582827082 | NA | + | + | + | + | + | NA | NA | + | NA | + | + | 0,0731865775500655 | 10 | -524,898709742027 | 1071,54345123009 | 23,5818783147954 | 2.63413385230676e-07 |
| **3160** | 25,0944797327914 | + | + | + | NA | + | NA | + | NA | NA | NA | + | + | 0,0572843277567153 | 9 | -526,063519338014 | 1071,54436151067 | 23,5827885953829 | 2.632935224644e-07 |
| **472** | 25,8999568255857 | + | + | + | NA | + | NA | + | + | + | NA | NA | NA | 0,0572387470243218 | 9 | -526,066829733809 | 1071,55098230226 | 23,5894093869726 | 2.62423357784224e-07 |
| **3511** | 25,8093614606663 | NA | + | + | NA | + | + | NA | + | + | NA | + | + | 0,0731047123807191 | 10 | -524,904757216995 | 1071,55554618002 | 23,5939732647303 | 2.61825206453337e-07 |
| **3311** | 25,0563771877707 | NA | + | + | + | NA | + | + | + | NA | NA | + | + | 0,0730987737782038 | 10 | -524,905195887678 | 1071,55642352139 | 23,5948506060977 | 2.61710376599097e-07 |
| **2294** | 25,4667081013432 | + | NA | + | NA | + | + | + | + | NA | NA | NA | + | 0,0571767726802372 | 9 | -526,071330493236 | 1071,55998382112 | 23,5984109058279 | 2.61244907332942e-07 |
| **2461** | 26,1296539559777 | NA | NA | + | + | + | NA | NA | + | + | NA | NA | + | 0,0412513495637032 | 8 | -527,218189213367 | 1071,56137842673 | 23,5998055114439 | 2.6106280402343e-07 |
| **3496** | 25,7887284774801 | + | + | + | NA | NA | + | NA | + | + | NA | + | + | 0,0729843112867952 | 10 | -524,913650415473 | 1071,57333257698 | 23,6117596616871 | 2.59507066045333e-07 |
| **1412** | 26,8755239154185 | + | + | NA | NA | NA | NA | NA | + | + | NA | + | NA | 0,0252269519152575 | 7 | -528,353114908218 | 1071,5744468707 | 23,6128739554094 | 2.59362522767686e-07 |
| **1480** | 25,9311095810097 | + | + | + | NA | NA | NA | + | + | + | NA | + | NA | 0,0570651000001542 | 9 | -526,079439745778 | 1071,5762023262 | 23,6146294109108 | 2.5913497296197e-07 |
| **144** | 25,9787076017147 | + | + | + | + | NA | NA | NA | + | NA | NA | NA | NA | 0,0251582594205293 | 7 | -528,357939792755 | 1071,58409663977 | 23,6225237244832 | 2.58114142609e-07 |
| **156** | 26,1851987851086 | + | + | NA | + | + | NA | NA | + | NA | NA | NA | NA | 0,0250636591643966 | 7 | -528,364583853447 | 1071,59738476116 | 23,6358118458675 | 2.56404901020101e-07 |
| **2112** | 25,0272739883013 | + | + | + | + | + | + | NA | NA | NA | NA | NA | + | 0,0569143023027731 | 9 | -526,090388587601 | 1071,59810000985 | 23,6365270945566 | 2.56313220779945e-07 |
| **3323** | 25,0008865896208 | NA | + | NA | + | + | + | + | + | NA | NA | + | + | 0,0727877164459343 | 10 | -524,928169039273 | 1071,60236982458 | 23,6407969092863 | 2.55766599497712e-07 |
| **3469** | 26,1966131036555 | NA | NA | + | + | NA | NA | NA | + | + | NA | + | + | 0,0409474892745079 | 8 | -527,239885909159 | 1071,60477181832 | 23,643198903028 | 2.55459608996274e-07 |
| **1298** | 27,0850745681843 | + | NA | NA | NA | + | NA | NA | NA | + | NA | + | NA | 0,0090460489795901 | 6 | -529,480357493605 | 1071,60686883336 | 23,645295918074 | 2.55191898047411e-07 |
| **3304** | 25,2295966634565 | + | + | + | NA | NA | + | + | + | NA | NA | + | + | 0,072743560175831 | 10 | -524,931429577955 | 1071,60889090194 | 23,6473179866512 | 2.54934020671718e-07 |
| **3410** | 25,9327927335162 | + | NA | NA | NA | + | NA | + | NA | + | NA | + | + | 0,0408212026584802 | 8 | -527,248901197521 | 1071,62280239504 | 23,6612294797508 | 2.53166917123927e-07 |
| **404** | 26,8576149987652 | + | + | NA | NA | + | NA | NA | + | + | NA | NA | NA | 0,0248819921419605 | 7 | -528,377341068412 | 1071,62289919109 | 23,6613262757971 | 2.53154664642103e-07 |
| **3388** | 25,5873061822955 | + | + | NA | + | + | + | NA | NA | + | NA | + | + | 0,0726440354125738 | 10 | -524,93877800688 | 1071,62358775979 | 23,6620148445004 | 2.53067522454232e-07 |
| **1264** | 25,2919749223105 | + | + | + | + | NA | + | + | + | NA | NA | + | NA | 0,0726096973967825 | 10 | -524,941313177596 | 1071,62865810122 | 23,6670851859324 | 2.52426765639279e-07 |
| **3316** | 25,1850601599171 | + | + | NA | NA | + | + | + | + | NA | NA | + | + | 0,0725902399762414 | 10 | -524,942749674482 | 1071,631531095 | 23,6699581797045 | 2.52064415696069e-07 |
| **462** | 26,344380143869 | + | NA | + | + | NA | NA | + | + | + | NA | NA | NA | 0,0407160735027453 | 8 | -527,25640520238 | 1071,63781040476 | 23,6762374894693 | 2.51274261477506e-07 |
| **2232** | 25,3462975484666 | + | + | + | NA | + | + | NA | + | NA | NA | NA | + | 0,0566305968329386 | 9 | -526,110982610319 | 1071,63928805528 | 23,6777151399922 | 2.51088682269315e-07 |
| **1276** | 25,2375749786446 | + | + | NA | + | + | + | + | + | NA | NA | + | NA | 0,0725144165786401 | 10 | -524,948347255213 | 1071,64272625646 | 23,6811533411678 | 2.50657406377563e-07 |
| **1344** | 25,6056463800141 | + | + | + | + | + | + | NA | NA | + | NA | + | NA | 0,0725117504555157 | 10 | -524,948544070559 | 1071,64311988715 | 23,6815469718601 | 2.50608078007809e-07 |
| **2368** | 25,4337051670595 | + | + | + | + | + | + | NA | NA | + | NA | NA | + | 0,0723954754840949 | 10 | -524,957127032083 | 1071,6602858102 | 23,6987128949065 | 2.48466322965509e-07 |
| **3103** | 25,3581403827198 | NA | + | + | + | + | NA | NA | NA | NA | NA | + | + | 0,0405123035081695 | 8 | -527,270947742644 | 1071,66689548529 | 23,7053225699972 | 2.4764653750764e-07 |
| **3100** | 25,5195020107208 | + | + | NA | + | + | NA | NA | NA | NA | NA | + | + | 0,0404007850468128 | 8 | -527,278905221339 | 1071,68281044268 | 23,7212375273871 | 2.45683715376578e-07 |
| **32** | 25,8656755989263 | + | + | + | + | + | NA | NA | NA | NA | NA | NA | NA | 0,0243822216115296 | 7 | -528,412424232907 | 1071,69306552008 | 23,7314926047877 | 2.44427186820362e-07 |
| **1270** | 25,6613003458785 | + | NA | + | NA | + | + | + | + | NA | NA | + | NA | 0,0562374343769766 | 9 | -526,139511823482 | 1071,69634648161 | 23,7347735663182 | 2.44026537440539e-07 |
| **1392** | 25,2450711980623 | + | + | + | + | NA | + | + | NA | + | NA | + | NA | 0,0720939104102363 | 10 | -524,979382370668 | 1071,70479648737 | 23,7432235720769 | 2.42997699569147e-07 |
| **398** | 26,6641256320107 | + | NA | + | + | NA | NA | NA | + | + | NA | NA | NA | 0,0242137367284478 | 7 | -528,424247578003 | 1071,71671221027 | 23,7551392949797 | 2.4155425712686e-07 |
| **2488** | 25,7476344267948 | + | + | + | NA | + | + | NA | + | + | NA | NA | + | 0,0719598060382113 | 10 | -524,989276878267 | 1071,72458550257 | 23,7630125872761 | 2.40605212742548e-07 |
| **1052** | 26,1923447737011 | + | + | NA | + | + | NA | NA | NA | NA | NA | + | NA | 0,0240858620655664 | 7 | -528,433219758553 | 1071,73465657137 | 23,77308365608 | 2.3939668226857e-07 |
| **3508** | 25,9490687287234 | + | + | NA | NA | + | + | NA | + | + | NA | + | + | 0,0718880847006547 | 10 | -524,994568045537 | 1071,73516783711 | 23,7735949218145 | 2.39335492429669e-07 |
| **3167** | 24,9246994318189 | NA | + | + | + | + | NA | + | NA | NA | NA | + | + | 0,0559553647532356 | 9 | -526,159972442842 | 1071,73726772033 | 23,775694805038 | 2.39084336009854e-07 |
| **1415** | 26,842628372858 | NA | + | + | NA | NA | NA | NA | + | + | NA | + | NA | 0,0240351748330987 | 7 | -528,436775845469 | 1071,7417687452 | 23,7801958299119 | 2.38546878742161e-07 |
| **3532** | 25,4493559627154 | + | + | NA | + | NA | NA | + | + | + | NA | + | + | 0,0718116145027788 | 10 | -525,00020910489 | 1071,74644995581 | 23,7848770405208 | 2.37989187574482e-07 |

| | | | | | | | | | | | | | | | | | | |
|---|---|---|---|---|---|---|---|---|---|---|---|---|---|---|---|---|---|---|
| 1173 | 26,8939293709446 | NA | NA | + | NA | + | NA | NA | + | NA | NA | + | NA | 0,0080238774528760 | 6 | -529,550947971977 | 1071,74804979011 | 23,7864768748163 | 2.37798892062821e-07 |
| 1300 | 26,7817763285853 | + | + | NA | NA | + | NA | NA | NA | + | NA | + | NA | 0,0238909031683693 | 7 | -528,446896566982 | 1071,76201018823 | 23,8004372729363 | 2.3614478816332e-07 |
| 3319 | 25,1452986526343 | NA | + | + | NA | + | + | + | + | NA | NA | + | + | 0,0716915265305589 | 10 | -525,009066826056 | 1071,76416539814 | 23,8025924828539 | 2.35890454428853e-07 |
| 3088 | 25,295879331064 | + | + | + | + | NA | NA | NA | NA | NA | NA | + | + | 0,0397641035368368 | 8 | -527,324318385142 | 1071,77363677028 | 23,8120638549933 | 2.34775992244173e-07 |
| 1464 | 25,9129789325394 | + | + | + | NA | + | + | NA | + | + | NA | + | NA | 0,0715837862261708 | 10 | -525,017012805248 | 1071,78005735653 | 23,8184844412374 | 2.34023500996054e-07 |
| 1170 | 26,941972992861 | + | NA | NA | NA | + | NA | NA | + | NA | NA | + | NA | 0,0077836460229835 | 6 | -529,567527641054 | 1071,78120912826 | 23,819636212972 | 2.33888768968072e-07 |
| 2192 | 25,4181696382198 | + | + | + | + | NA | NA | NA | + | NA | NA | NA | + | 0,0396638967882618 | 8 | -527,331463182067 | 1071,78792636413 | 23,8263534488428 | 2.33104543647529e-07 |
| 2207 | 25,4086200836329 | NA | + | + | + | + | NA | NA | + | NA | NA | NA | + | 0,0396399709558825 | 8 | -527,333168997023 | 1071,79133799405 | 23,8297650787558 | 2.32707249382135e-07 |
| 1279 | 25,1984940612676 | NA | + | + | + | + | + | + | + | NA | NA | + | NA | 0,0714924515294723 | 10 | -525,023748128998 | 1071,79352800403 | 23,8319550887381 | 2.32452573243852e-07 |
| 1482 | 26,4588616466057 | + | NA | NA | + | NA | NA | + | + | + | NA | + | NA | 0,0395967542454829 | 8 | -527,336250065804 | 1071,79750013161 | 23,8359272163166 | 2.31991365750773e-07 |
| 2458 | 26,2500079373244 | + | NA | NA | + | + | NA | NA | + | + | NA | NA | + | 0,0395755408658979 | 8 | -527,337762390309 | 1071,80052478062 | 23,838951865328 | 2.31640784686221e-07 |
| 1478 | 26,3170467156928 | + | NA | + | NA | NA | NA | + | + | + | NA | + | NA | 0,0395720498854361 | 8 | -527,338011262839 | 1071,80102252568 | 23,8394496103865 | 2.31583142831292e-07 |
| 2204 | 25,5105689833696 | + | + | NA | + | + | NA | NA | + | NA | NA | NA | + | 0,039552023489882 | 8 | -527,339438930159 | 1071,80387786032 | 23,8423049450275 | 2.31252755044331e-07 |
| 474 | 26,4636122704474 | + | NA | NA | + | + | NA | + | + | + | NA | NA | NA | 0,0395260676702737 | 8 | -527,341289257578 | 1071,80757851516 | 23,8460055998655 | 2.30825257358269e-07 |
| 3535 | 25,4519742416244 | NA | + | + | + | NA | NA | + | + | + | NA | + | + | 0,071390864415431 | 10 | -525,031238722185 | 1071,8085091904 | 23,8469362751105 | 2.30717870669274e-07 |
| 1485 | 26,4016119418441 | NA | NA | + | + | NA | NA | + | + | + | NA | + | NA | 0,0395189401751065 | 8 | -527,341797350673 | 1071,80859470135 | 23,8470217860547 | 2.30708006428663e-07 |
| 2416 | 24,9647689320041 | + | + | + | + | NA | + | + | NA | + | NA | NA | + | 0,0713776279488492 | 10 | -525,032214661507 | 1071,81046106905 | 23,8488881537558 | 2.30492813865607e-07 |
| 3481 | 26,3253959291793 | NA | NA | NA | + | + | NA | NA | + | + | NA | + | + | 0,0394845939491741 | 8 | -527,34424571482 | 1071,81349142964 | 23,8519185143502 | 2.301438401412e-07 |
| 3547 | 25,4275079159878 | NA | + | NA | + | + | NA | + | + | + | NA | + | + | 0,071338198538079 | 10 | -525,03512175318 | 1071,81627525239 | 23,854702337102 | 2.2982372315094e-07 |
| 3376 | 25,3068853935792 | + | + | + | + | NA | + | NA | NA | + | NA | + | + | 0,0712377639658732 | 10 | -525,042526138077 | 1071,83108402219 | 23,8695111068951 | 2.28128304358954e-07 |
| 3246 | 25,5832145499995 | + | NA | + | + | NA | + | NA | + | NA | NA | + | + | 0,0552347419060203 | 9 | -526,212216873138 | 1071,84175658092 | 23,8801836656312 | 2.26914190302106e-07 |
| 3261 | 25,5203101826443 | NA | NA | + | + | + | + | NA | + | NA | NA | + | + | 0,0552251900229789 | 9 | -526,21290910752 | 1071,84314104969 | 23,8815681343954 | 2.26757166852581e-07 |
| 3384 | 25,6464194400349 | + | + | + | NA | + | + | NA | NA | + | NA | + | + | 0,0710268869213522 | 10 | -525,058070121007 | 1071,86217198805 | 23,9005990727544 | 2.24609699304158e-07 |
| 3258 | 25,648483750534 | + | NA | NA | + | + | + | NA | + | NA | NA | + | + | 0,0550168373010628 | 9 | -526,228006894476 | 1071,8733366236 | 23,9117637083064 | 2.233593497583e-07 |
| 470 | 26,3956138904689 | + | NA | + | NA | + | NA | + | + | + | NA | NA | NA | 0,0390287646405662 | 8 | -527,376731135066 | 1071,87846227013 | 23,9168893548417 | 2.22787652112611e-07 |
| 3152 | 24,8591835770655 | + | + | + | + | NA | NA | + | NA | NA | NA | + | + | 0,0549412672992024 | 9 | -526,233482073588 | 1071,88428698182 | 23,9227140665303 | 2.22139759097396e-07 |
| 477 | 26,485681604189 | NA | NA | + | + | + | NA | + | + | + | NA | NA | NA | 0,0389622607099209 | 8 | -527,381469358781 | 1071,88793871756 | 23,9263658022712 | 2.21734531307284e-07 |
| 3208 | 25,7591431298398 | + | + | + | NA | NA | NA | NA | + | NA | NA | + | + | 0,0389218917846063 | 8 | -527,384345374783 | 1071,89369074957 | 23,9321178342748 | 2.21097735403638e-07 |
| 3223 | 25,7857015164529 | NA | + | + | NA | + | NA | NA | + | NA | NA | + | + | 0,0388686492113545 | 8 | -527,388138367477 | 1071,90127673495 | 23,9397038196632 | 2.2026070174334e-07 |
| 2300 | 24,9566334266729 | + | + | NA | + | + | + | + | + | NA | NA | NA | + | 0,0706167996555974 | 10 | -525,088288016767 | 1071,92260777957 | 23,9610348642752 | 2.17923989594233e-07 |
| 1272 | 25,375260922047 | + | + | + | NA | + | + | + | + | NA | NA | + | NA | 0,0705976390646028 | 10 | -525,089699567763 | 1071,92543088156 | 23,9638579662669 | 2.17616595771772e-07 |
| 2557 | 25,7631683779172 | NA | NA | + | + | + | + | + | + | + | NA | NA | + | 0,0705309183418254 | 10 | -525,094614622502 | 1071,93526099104 | 23,9736880757459 | 2.16549622555182e-07 |
| 2288 | 24,9703425547914 | + | + | + | + | NA | + | + | + | NA | NA | NA | + | 0,070521612700226 | 10 | -525,095300104707 | 1071,93663195545 | 23,9750590401552 | 2.1640123250768e-07 |
| 1247 | 25,6193017613502 | NA | + | + | + | + | NA | + | + | NA | NA | + | NA | 0,0545267403993819 | 9 | -526,263507492481 | 1071,94433781961 | 23,9827649043161 | 2.15569057443177e-07 |
| 1040 | 25,8329585286081 | + | + | + | + | NA | NA | NA | NA | NA | NA | + | NA | 0,0225593967055169 | 7 | -528,540231821985 | 1071,94868069823 | 23,9871077829423 | 2.15101470169582e-07 |
| 1244 | 25,6064524686568 | + | + | NA | + | + | NA | + | + | NA | NA | + | NA | 0,0544833925488451 | 9 | -526,266646546539 | 1071,95061592772 | 23,9890430124337 | 2.14893435480168e-07 |
| 3565 | 25,7982445766105 | NA | NA | + | + | NA | + | + | + | + | NA | + | + | 0,0703353418636301 | 10 | -525,109019945188 | 1071,96407163641 | 24,0024987211177 | 2.13452516329436e-07 |
| 2542 | 25,8036881638291 | + | NA | + | + | NA | + | + | + | + | NA | NA | + | 0,0703339728189799 | 10 | -525,109120772456 | 1071,96427329094 | 24,0027003756529 | 2.13430995580409e-07 |
| 1497 | 26,5291004815699 | NA | NA | NA | + | + | NA | + | + | + | NA | + | NA | 0,0384027980356935 | 8 | -527,421316578179 | 1071,96763315636 | 24,0060602410676 | 2.13072746871284e-07 |
| 3165 | 25,3589123862249 | NA | NA | + | + | + | NA | + | NA | NA | NA | + | + | 0,0383069041765293 | 8 | -527,428144204533 | 1071,98128840907 | 24,019715493775 | 2.11622920836857e-07 |
| 1048 | 26,1933791799809 | + | + | + | NA | + | NA | NA | NA | NA | NA | + | NA | 0,0222418325866149 | 7 | -528,562473498235 | 1071,99316405073 | 24,0315911354435 | 2.10370065145892e-07 |
| 1232 | 25,6383197372932 | + | + | + | + | NA | NA | + | + | NA | NA | + | NA | 0,0541852924828711 | 9 | -526,28822970145 | 1071,99378223755 | 24,0322093222546 | 2.10305051194234e-07 |
| 2238 | 25,4479479910305 | + | NA | + | + | + | + | NA | + | NA | NA | NA | + | 0,0540930116570644 | 9 | -526,294909674873 | 1072,00714218439 | 24,0455692691 | 2.08904900731995e-07 |
| 3220 | 25,9030741474951 | + | + | NA | NA | + | NA | NA | + | NA | NA | + | + | 0,0380373852961415 | 8 | -527,447330259204 | 1072,01966051841 | 24,0580876031177 | 2.07601413696455e-07 |
| 1214 | 25,6665720386305 | + | NA | + | + | + | + | NA | + | NA | NA | + | NA | 0,0539886400985757 | 9 | -526,302464080622 | 1072,02225099589 | 24,0606780805997 | 2.07332694368154e-07 |
| 1160 | 26,284227799617 | + | + | + | NA | NA | NA | NA | + | NA | NA | + | NA | 0,0220308950389158 | 7 | -528,577243231597 | 1072,02270351746 | 24,061130602166 | 2.07285788417039e-07 |
| 2525 | 25,866845702515 | NA | NA | + | + | + | NA | + | + | + | NA | NA | + | 0,0539355567576738 | 9 | -526,306305929351 | 1072,02993469335 | 24,0683617780564 | 2.06537681657085e-07 |
| 3577 | 25,8561523444781 | NA | NA | NA | + | + | + | + | + | + | NA | + | + | 0,0698834906885353 | 10 | -525,142289780456 | 1072,03061130694 | 24,0690383916531 | 2.06467820373213e-07 |
| 2524 | 25,4091880478414 | + | + | NA | + | + | NA | + | + | + | NA | NA | + | 0,0698725408795737 | 10 | -525,143095815194 | 1072,03222337642 | 24,0706504611296 | 2.06301467189885e-07 |
| 2554 | 25,8509077152131 | + | NA | NA | + | + | + | + | + | + | NA | NA | + | 0,0698650424761286 | 10 | -525,143647780417 | 1072,03332730687 | 24,0717543915762 | 2.06187627375184e-07 |
| 2142 | 25,3216714504633 | + | NA | + | + | + | NA | + | NA | NA | NA | NA | + | 0,0379253231429309 | 8 | -527,455305969016 | 1072,03561193803 | 24,0740390227415 | 2.05952230509828e-07 |
| 3528 | 25,5592884234523 | + | + | + | NA | NA | NA | + | + | + | NA | + | + | 0,0698471939440301 | 10 | -525,144961611409 | 1072,03595496885 | 24,0743820535597 | 2.05916909557891e-07 |
| 3540 | 25,5527951814658 | + | + | NA | NA | + | NA | + | + | + | NA | + | + | 0,0698421364110577 | 10 | -525,145333891984 | 1072,03669953 | 24,0751266147086 | 2.0584026496005e-07 |
| 2512 | 25,4049163187957 | + | + | + | + | NA | NA | + | + | + | NA | NA | + | 0,0698164601369904 | 10 | -525,14722386883 | 1072,04047948369 | 24,0789065684012 | 2.05451599025738e-07 |
| 3466 | 26,4582652742714 | + | NA | NA | + | NA | NA | NA | + | + | NA | + | + | 0,0378486506963981 | 8 | -527,460762380603 | 1072,04652476121 | 24,0849518459154 | 2.04831530653258e-07 |
| 3543 | 25,5350187028071 | NA | + | + | NA | + | NA | + | + | + | NA | + | + | 0,0697164080292149 | 10 | -525,154587997826 | 1072,05520774168 | 24,0936348263924 | 2.03944184159222e-07 |
| 2144 | 24,8881692255941 | + | + | + | + | + | NA | + | NA | NA | NA | NA | + | 0,0537068867455817 | 9 | -526,322853208445 | 1072,06302925153 | 24,1014563362442 | 2.03148165966978e-07 |
| 3158 | 25,4515965150861 | + | NA | + | NA | + | NA | + | NA | NA | NA | + | + | 0,0377247931755666 | 8 | -527,469575810368 | 1072,06415162074 | 24,1025787054446 | 2.03034194327234e-07 |
| 256 | 25,13531949913 | + | + | + | + | + | + | + | + | NA | NA | NA | NA | 0,069604751057402 | 10 | -525,162805344258 | 1072,07164243455 | 24,1100695192576 | 2.02275170966507e-07 |
| 152 | 26,1729357562529 | + | + | + | NA | + | NA | NA | + | NA | NA | NA | NA | 0,0215706535107153 | 7 | -528,609458044966 | 1072,08713314419 | 24,1255602289041 | 2.00714529657755e-07 |
| 2522 | 25,9167531809461 | + | NA | NA | + | + | NA | + | + | + | NA | NA | + | 0,0535401907518788 | 9 | -526,334913334679 | 1072,087149504 | 24,1255765887136 | 2.00712887838738e-07 |
| 2527 | 25,3900621502001 | NA | + | + | + | + | NA | + | + | + | NA | NA | + | 0,0694900191458457 | 10 | -525,171247962816 | 1072,08852767166 | 24,1269547563729 | 2.0057462747528e-07 |
| 3545 | 25,9455651316227 | NA | NA | NA | + | + | NA | + | + | + | NA | + | + | 0,0535047882125252 | 9 | -526,337474365214 | 1072,09227156507 | 24,1306986497832 | 2.0019951366831e-07 |
| 1490 | 26,5163394077986 | + | NA | NA | NA | + | NA | + | + | + | NA | + | NA | 0,0375120858366284 | 8 | -527,484708953074 | 1072,09441790615 | 24,1328449908581 | 1.99984780691793e-07 |
| 3150 | 25,2790158505045 | + | NA | + | + | NA | NA | + | NA | NA | NA | + | + | 0,0374663839431179 | 8 | -527,487959994995 | 1072,10091998999 | 24,1393470746989 | 1.99335677488894e-07 |
| 286 | 26,5116141910056 | + | NA | + | + | + | NA | NA | NA | + | NA | NA | NA | 0,0214529770684652 | 7 | -528,617692429135 | 1072,10360191253 | 24,1420289972425 | 1.99068555205638e-07 |
| 2510 | 25,9168090299239 | + | NA | + | + | NA | NA | + | + | + | NA | NA | + | 0,0533667676315145 | 9 | -526,347457902093 | 1072,11223863883 | 24,1506657235409 | 1.98210758364842e-07 |
| 1375 | 25,6972081136687 | NA | + | + | + | + | NA | + | NA | + | NA | + | NA | 0,0533223968984723 | 9 | -526,350667091559 | 1072,11865701776 | 24,157084102472 | 1.97575682072259e-07 |
| 3533 | 25,9271585446025 | NA | NA | + | + | NA | NA | + | + | + | NA | + | + | 0,0532843736579136 | 9 | -526,35341706811 | 1072,12415697087 | 24,1625840555746 | 1.97033099965425e-07 |
| 3096 | 25,6165428067663 | + | + | + | NA | + | NA | NA | NA | NA | NA | + | + | 0,037268235523002 | 8 | -527,502053661978 | 1072,12910732396 | 24,1675344086657 | 1.96546011321567e-07 |
| 1427 | 27,1723235889808 | NA | + | NA | NA | + | NA | NA | + | + | NA | + | NA | 0,0212550797627735 | 7 | -528,631538019607 | 1072,13129309348 | 24,1697201781876 | 1.9633132651521e-07 |
| 3562 | 25,9657519729726 | + | NA | NA | + | NA | + | + | + | + | NA | + | + | 0,0691307551476841 | 10 | -525,197677897373 | 1072,14138754078 | 24,1798146254866 | 1.95342894916865e-07 |
| 1240 | 25,7055901541424 | + | + | + | NA | + | NA | + | + | NA | NA | + | NA | 0,0530959692397838 | 9 | -526,367041519954 | 1072,15140587455 | 24,1898329592627 | 1.9436683641671e-07 |
| 2303 | 24,9160637670341 | NA | + | + | + | + | + | + | + | NA | NA | NA | + | 0,0690271960636601 | 10 | -525,20529452394 | 1072,15662079391 | 24,1950478786212 | 1.93860692888964e-07 |
| 2334 | 25,8990653470476 | + | NA | + | + | + | NA | NA | NA | + | NA | NA | + | 0,0370252385508599 | 8 | -527,519333306296 | 1072,16366661259 | 24,202093697302 | 1.93178940823703e-07 |
| 3471 | 25,8705135535017 | NA | + | + | + | NA | NA | NA | + | + | NA | + | + | 0,0529646548941667 | 9 | -526,376535905031 | 1072,17039464471 | 24,2088217294163 | 1.92530175604991e-07 |

| | | | | | | | | | | | | | | | | | | |
|---|---|---|---|---|---|---|---|---|---|---|---|---|---|---|---|---|---|---|
| **2296** | 25,0884466373781 | + | + | + | NA | + | + | + | + | NA | NA | NA | + | 0,0689205396543286 | 10 | -525,213138069297 | 1072,17230788462 | 24,2107349693342 | 1.92346085462495e-07 |
| **510** | 25,9034094817226 | + | NA | + | + | + | + | + | + | + | NA | NA | NA | 0,0688837837446912 | 10 | -525,215840902196 | 1072,17771355042 | 24,2161406351322 | 1.91826908075638e-07 |
| **1518** | 25,8958690831156 | + | NA | + | + | NA | + | + | + | + | NA | + | NA | 0,0688803286449515 | 10 | -525,216094966257 | 1072,17822167855 | 24,2166487632555 | 1.91778177942786e-07 |
| **1533** | 25,9115698347045 | NA | NA | + | + | + | + | + | + | + | NA | + | NA | 0,0688498251006908 | 10 | -525,218337944371 | 1072,18270763477 | 24,2211347194834 | 1.91348505739741e-07 |
| **2454** | 26,2262103439199 | + | NA | + | NA | + | NA | NA | + | + | NA | NA | + | 0,0368780666720659 | 8 | -527,529796658044 | 1072,18459331609 | 24,2230204007974 | 1.91168179611636e-07 |
| **3357** | 25,9650432333081 | NA | NA | + | + | + | NA | NA | NA | + | NA | + | + | 0,0368550349409524 | 8 | -527,531433980483 | 1072,18786796097 | 24,2262950456754 | 1.90855431765932e-07 |
| **224** | 25,5139386451566 | + | + | + | + | + | NA | + | + | NA | NA | NA | NA | 0,0528386243587166 | 9 | -526,385647018799 | 1072,18861687224 | 24,2270439569534 | 1.90783978252201e-07 |
| **352** | 25,6528588654671 | + | + | + | + | + | NA | + | NA | + | NA | NA | NA | 0,0527705368158971 | 9 | -526,390568761039 | 1072,19846035672 | 24,2368874414337 | 1.89847295633705e-07 |
| **1368** | 25,7519491557902 | + | + | + | NA | + | NA | + | NA | + | NA | + | NA | 0,0527615275317033 | 9 | -526,391219975183 | 1072,19976278501 | 24,2381898697206 | 1.89723704636167e-07 |
| **3342** | 25,8338239468433 | + | NA | + | + | NA | NA | NA | NA | + | NA | + | + | 0,0366710015644337 | 8 | -527,544515481117 | 1072,21403096223 | 24,2524580469435 | 1.88375015469758e-07 |
| **3462** | 26,275629143678 | + | NA | + | NA | NA | NA | NA | + | + | NA | + | + | 0,0366555025658196 | 8 | -527,545617070272 | 1072,21623414054 | 24,2546612252524 | 1.881676178502e-07 |
| **3162** | 25,5496648256716 | + | NA | NA | + | + | NA | + | NA | NA | NA | + | + | 0,0366438237014007 | 8 | -527,54644713222 | 1072,21789426444 | 24,2563213491489 | 1.88011491876809e-07 |
| **1372** | 25,7738236365356 | + | + | NA | + | + | NA | + | NA | + | NA | + | NA | 0,0526276910379713 | 9 | -526,400893290809 | 1072,21910941626 | 24,2575365009734 | 1.87897295318175e-07 |
| **1493** | 26,536094583956 | NA | NA | + | NA | + | NA | + | + | + | NA | + | NA | 0,0366323800541571 | 8 | -527,547260466619 | 1072,21952093324 | 24,2579480179484 | 1.87858637832078e-07 |
| **1175** | 26,5377314061836 | NA | + | + | NA | + | NA | NA | + | NA | NA | + | NA | 0,020575332531051 | 7 | -528,679074219171 | 1072,22636549261 | 24,2647925773149 | 1.8721683187864e-07 |
| **1421** | 26,8613275474245 | NA | NA | + | + | NA | NA | NA | + | + | NA | + | NA | 0,0204383594365945 | 7 | -528,688649052921 | 1072,24551516011 | 24,2839422448144 | 1.85432816294039e-07 |
| **3483** | 25,9459621382726 | NA | + | NA | + | + | NA | NA | + | + | NA | + | + | 0,0524060963241285 | 9 | -526,416906514716 | 1072,25113586408 | 24,2895629487875 | 1.84912416406334e-07 |
| **2080** | 25,2795733973612 | + | + | + | + | + | NA | NA | NA | NA | NA | NA | + | 0,0363814673377224 | 8 | -527,565091160657 | 1072,25518232131 | 24,2936094060226 | 1.84538674523851e-07 |
| **3564** | 25,4254958542254 | + | + | NA | + | NA | + | + | + | + | NA | + | + | 0,0842950088109582 | 11 | -524,073125646978 | 1072,25825129396 | 24,2966783786651 | 1.84255719602803e-07 |
| **3468** | 25,9885111728629 | + | + | NA | + | NA | NA | NA | + | + | NA | + | + | 0,0522878295545974 | 9 | -526,425451361212 | 1072,26822555707 | 24,3066526417786 | 1.83339099641039e-07 |
| **413** | 26,9358469416742 | NA | NA | + | + | + | NA | NA | + | + | NA | NA | NA | 0,0202641915168706 | 7 | -528,700821984974 | 1072,26986102421 | 24,3082881089213 | 1.83189238385962e-07 |
| **222** | 26,0221174662122 | + | NA | + | + | + | NA | + | + | NA | NA | NA | NA | 0,0362646100778224 | 8 | -527,573393843177 | 1072,27178768635 | 24,3102147710626 | 1.83012851473945e-07 |
| **1230** | 26,0173142159336 | + | NA | + | + | NA | NA | + | + | NA | NA | + | NA | 0,0362546684434006 | 8 | -527,574100147661 | 1072,27320029532 | 24,3116273800308 | 1.82883634315005e-07 |
| **3196** | 24,9746136219752 | + | + | NA | + | + | + | + | NA | NA | NA | + | + | 0,0681763397133204 | 10 | -525,267841781248 | 1072,28171530853 | 24,3201423932369 | 1.82106661192063e-07 |
| **1172** | 26,5936281934182 | + | + | NA | NA | + | NA | NA | + | NA | NA | + | NA | 0,0200942371297571 | 7 | -528,712698340117 | 1072,2936137345 | 24,3320408192071 | 1.81026486160183e-07 |
| **3530** | 26,0763146693965 | + | NA | NA | + | NA | NA | + | + | + | NA | + | + | 0,0520777354020577 | 9 | -526,440628164219 | 1072,29857916308 | 24,337006247792 | 1.80577606566141e-07 |
| **2520** | 25,4948503413606 | + | + | + | NA | + | NA | + | + | + | NA | NA | + | 0,0680437230635788 | 10 | -525,277585414494 | 1072,30120257502 | 24,3396296597296 | 1.80340897122319e-07 |
| **1425** | 27,7060084072789 | NA | NA | NA | NA | + | NA | NA | + | + | NA | + | NA | 0,0039563526391761 | 6 | -529,831130024753 | 1072,30841389566 | 24,3468409803691 | 1.79691819984586e-07 |
| **1530** | 26,0093993668882 | + | NA | NA | + | + | + | + | + | + | NA | + | NA | 0,0679668834585722 | 10 | -525,283230353037 | 1072,3124924521 | 24,3509195368142 | 1.79325751753664e-07 |
| **1414** | 26,7464682346541 | + | NA | + | NA | NA | NA | NA | + | + | NA | + | NA | 0,0198800694434088 | 7 | -528,727661373995 | 1072,32353980225 | 24,3619668869633 | 1.78337945241165e-07 |
| **3197** | 25,1989551104544 | NA | NA | + | + | + | + | + | NA | NA | NA | + | + | 0,0518866226820836 | 9 | -526,454430864568 | 1072,32618456378 | 24,3646116484915 | 1.78102270433124e-07 |
| **2550** | 25,8582146641801 | + | NA | + | NA | + | + | + | + | + | NA | NA | + | 0,0678615189764085 | 10 | -525,290970084655 | 1072,32797191534 | 24,3663990000514 | 1.77943175847764e-07 |
| **410** | 26,9775546589175 | + | NA | NA | + | + | NA | NA | + | + | NA | NA | NA | 0,0198152758059151 | 7 | -528,732187600622 | 1072,33259225551 | 24,3710193402173 | 1.7753257131222e-07 |
| **2518** | 25,9262610330943 | + | NA | + | NA | + | NA | + | + | + | NA | NA | + | 0,0518422312955311 | 9 | -526,457636537869 | 1072,33259591038 | 24,3710229950932 | 1.77532246882762e-07 |
| **1242** | 26,123299864109 | + | NA | NA | + | + | NA | + | + | NA | NA | + | NA | 0,0357336291013459 | 8 | -527,611107254434 | 1072,34721450887 | 24,3856415935782 | 1.76239341437711e-07 |
| **3558** | 25,8789759482825 | + | NA | + | NA | NA | + | + | + | + | NA | + | + | 0,0677246240610353 | 10 | -525,301024633156 | 1072,34808101234 | 24,3865080970529 | 1.76163001975136e-07 |
| **2174** | 25,1855757242513 | + | NA | + | + | + | + | + | NA | NA | NA | NA | + | 0,0517200837463349 | 9 | -526,466456509443 | 1072,35023585353 | 24,3886629382412 | 1.75973302540274e-07 |
| **406** | 26,8192373337959 | + | NA | + | NA | + | NA | NA | + | + | NA | NA | NA | 0,019688273355128 | 7 | -528,741058620761 | 1072,35033429578 | 24,3887613804941 | 1.7596464114926e-07 |
| **1294** | 26,3132753351877 | + | NA | + | + | NA | NA | NA | NA | + | NA | + | NA | 0,019683266796125 | 7 | -528,741408301388 | 1072,35103365704 | 24,3894607417487 | 1.75903120480076e-07 |
| **3517** | 25,8773245595388 | NA | NA | + | + | + | + | NA | + | + | NA | + | + | 0,0676931464170992 | 10 | -525,303336369352 | 1072,35270448474 | 24,3911315694456 | 1.75756229942979e-07 |
| **2494** | 25,8535465910733 | + | NA | + | + | + | + | NA | + | + | NA | NA | + | 0,0676582096287508 | 10 | -525,305902055751 | 1072,35783585753 | 24,3962629422429 | 1.75305872559299e-07 |
| **3573** | 25,9011280607642 | NA | NA | + | NA | + | + | + | + | + | NA | + | + | 0,0676412063416937 | 10 | -525,307150707668 | 1072,36033316137 | 24,398760246077 | 1.75087113151052e-07 |
| **3502** | 25,8882311425662 | + | NA | + | + | NA | + | NA | + | + | NA | + | + | 0,067591645389741 | 10 | -525,310790131592 | 1072,36761200922 | 24,4060390939253 | 1.74451055067216e-07 |
| **1418** | 27,0076344039953 | + | NA | NA | + | NA | NA | NA | + | + | NA | + | NA | 0,0195606625472717 | 7 | -528,749970977207 | 1072,36815900868 | 24,4065860933863 | 1.74403349274728e-07 |
| **3579** | 25,4095790310024 | NA | + | NA | + | + | + | + | + | + | NA | + | + | 0,0835544737895329 | 11 | -524,12847320694 | 1072,36894641388 | 24,4073734985905 | 1.74334699736944e-07 |
| **3567** | 25,405629179588 | NA | + | + | + | NA | + | + | + | + | NA | + | + | 0,0835409923228266 | 11 | -524,129480396897 | 1072,37096079379 | 24,4093878785031 | 1.74159199973963e-07 |
| **1245** | 26,1295719600052 | NA | NA | + | + | + | NA | + | + | NA | NA | + | NA | 0,035537177449641 | 8 | -527,625055152875 | 1072,37511030575 | 24,4135373904594 | 1.73798236717492e-07 |
| **1150** | 25,3713435300842 | + | NA | + | + | + | + | + | NA | NA | NA | + | NA | 0,0514761727277364 | 9 | -526,484065321847 | 1072,38545347834 | 24,4238805630491 | 1.72901744280492e-07 |
| **3190** | 25,3438274370922 | + | NA | + | NA | + | + | + | NA | NA | NA | + | + | 0,0514086735967095 | 9 | -526,488937526972 | 1072,39519788859 | 24,433624973298 | 1.72061380391011e-07 |
| **3526** | 25,9723568084029 | + | NA | + | NA | NA | NA | + | + | + | NA | + | + | 0,0512985716217939 | 9 | -526,496884136163 | 1072,41109110697 | 24,44951819168 | 1.70699494199606e-07 |
| **3541** | 26,0011399329328 | NA | NA | + | NA | + | NA | + | + | + | NA | + | + | 0,0512457861321487 | 9 | -526,500693601565 | 1072,41871003778 | 24,4571371224856 | 1.70050457408678e-07 |
| **3192** | 25,0357740346687 | + | + | + | NA | + | + | + | NA | NA | NA | + | + | 0,0671722520541709 | 10 | -525,341579826531 | 1072,42919139909 | 24,4676184838022 | 1.69161608386874e-07 |
| **3254** | 25,6915156733607 | + | NA | + | NA | + | + | NA | + | NA | NA | + | + | 0,0511401009261413 | 9 | -526,508320139325 | 1072,4339631133 | 24,4723901980053 | 1.68758494040028e-07 |
| **3182** | 25,1343960666751 | + | NA | + | + | NA | + | + | NA | NA | NA | + | + | 0,0510309323546686 | 9 | -526,516197154954 | 1072,44971714455 | 24,4881442292624 | 1.67434402536121e-07 |
| **3194** | 25,4140154547425 | + | NA | NA | + | + | + | + | NA | NA | NA | + | + | 0,0509495065150866 | 9 | -526,52207181489 | 1072,46146646443 | 24,4998935491353 | 1.66453665927155e-07 |
| **2463** | 25,8095380926492 | NA | + | + | + | + | NA | NA | + | + | NA | NA | + | 0,0509444853670019 | 9 | -526,522434061013 | 1072,46219095667 | 24,5006180413802 | 1.66393379651999e-07 |
| **2448** | 25,789421498613 | + | + | + | + | NA | NA | NA | + | + | NA | NA | + | 0,05087250799985 | 9 | -526,527626591597 | 1072,47257601784 | 24,511003102549 | 1.65531616241812e-07 |
| **2200** | 25,6090553632559 | + | + | + | NA | + | NA | NA | + | NA | NA | NA | + | 0,034811701783207 | 8 | -527,676538706083 | 1072,47807741217 | 24,5165044968751 | 1.65076914554645e-07 |
| **1302** | 26,6217018207653 | + | NA | + | NA | + | NA | NA | NA | + | NA | + | NA | 0,0187568193979325 | 7 | -528,806084851815 | 1072,48038675789 | 24,5188138426029 | 1.64886414724612e-07 |
| **3477** | 26,4089029351771 | NA | NA | + | NA | + | NA | NA | + | + | NA | + | + | 0,0347787027598044 | 8 | -527,678879569825 | 1072,48275913965 | 24,5211862243593 | 1.64690943919285e-07 |
| **3538** | 26,0561726530928 | + | NA | NA | NA | + | NA | + | + | + | NA | + | + | 0,0507835891630788 | 9 | -526,534040755976 | 1072,4854043466 | 24,5238314313071 | 1.64473267086535e-07 |
| **2556** | 25,4034938862881 | + | + | NA | + | + | + | + | + | + | NA | NA | + | 0,0827709354774511 | 11 | -524,186986180785 | 1072,48597236157 | 24,5243994462792 | 1.64426562080032e-07 |
| **1309** | 26,706814117181 | NA | NA | + | + | + | NA | NA | NA | + | NA | + | NA | 0,0186691071234902 | 7 | -528,812205001488 | 1072,49262705724 | 24,531054141949 | 1.63880366909277e-07 |
| **1238** | 26,0706427586454 | + | NA | + | NA | + | NA | + | + | NA | NA | + | NA | 0,0346610800808046 | 8 | -527,687222761965 | 1072,49944552393 | 24,5378726086401 | 1.63322611796617e-07 |
| **1152** | 25,0961256400542 | + | + | + | + | + | + | + | NA | NA | NA | + | NA | 0,0666353329151617 | 10 | -525,380977470172 | 1072,50798668638 | 24,5464137710853 | 1.62626616527402e-07 |
| **2544** | 25,3720926377512 | + | + | + | + | NA | + | + | + | + | NA | NA | + | 0,0825407689573093 | 11 | -524,204165033358 | 1072,52033006672 | 24,5587571514257 | 1.61626026279291e-07 |
| **3570** | 26,0040577773272 | + | NA | NA | NA | + | + | + | + | + | NA | + | + | 0,066489936582277 | 10 | -525,39164235381 | 1072,52931645365 | 24,5677435383607 | 1.60901438349755e-07 |
| **1526** | 25,9675217444188 | + | NA | + | NA | + | + | + | + | + | NA | + | NA | 0,0663671101226249 | 10 | -525,40065043389 | 1072,54733261381 | 24,5857596985215 | 1.59458533964257e-07 |
| **3514** | 26,0152560479817 | + | NA | NA | + | + | + | NA | + | + | NA | + | + | 0,066350851990057 | 10 | -525,401842714908 | 1072,54971717585 | 24,5881442605573 | 1.59268527873906e-07 |
| **1470** | 25,9756483823456 | + | NA | + | + | + | + | NA | + | + | NA | + | NA | 0,0662739052575307 | 10 | -525,407485278796 | 1072,56100230362 | 24,599429388333 | 1.58372375705669e-07 |
| **3350** | 26,075030256007 | + | NA | + | NA | + | NA | NA | NA | + | NA | + | + | 0,0342264683173297 | 8 | -527,718041759407 | 1072,56108351881 | 24,5995106035227 | 1.5836594471497e-07 |
| **1532** | 25,6080372597161 | + | + | NA | + | + | + | + | + | + | NA | + | NA | 0,0821250179407922 | 11 | -524,235184371586 | 1072,58236874317 | 24,6207958278806 | 1.56689454325511e-07 |
| **2559** | 25,3601302710906 | NA | + | + | + | + | + | + | + | + | NA | NA | + | 0,0821023928773622 | 11 | -524,236872033056 | 1072,58574406611 | 24,6241711508205 | 1.56425238586765e-07 |
| **2460** | 25,9083822956173 | + | + | NA | + | + | NA | NA | + | + | NA | NA | + | 0,0500196788560734 | 9 | -526,589120730758 | 1072,59556429616 | 24,6339913808702 | 1.55659055236051e-07 |
| **3134** | 25,3700377991732 | + | NA | + | + | + | + | NA | NA | NA | NA | + | + | 0,0499457259571247 | 9 | -526,594450581859 | 1072,60622399836 | 24,6446510830731 | 1.54831622654038e-07 |
| **3280** | 25,2401082657745 | + | + | + | + | NA | NA | + | + | NA | NA | + | + | 0,0659511456357069 | 10 | -525,431148431059 | 1072,60832860815 | 24,6467556928594 | 1.54668778275898e-07 |

| | | | | | | | | | | | | | | | | | | |
|---|---|---|---|---|---|---|---|---|---|---|---|---|---|---|---|---|---|---|
| 1520 | 25,5901126216628 | + | + | + | + | NA | + | + | + | + | NA | + | NA | 0,0818686987014768 | 11 | -524,254301448271 | 1072,62060289654 | 24,6590299812517 | 1.53722460495531e-07 |
| 1166 | 26,3503935548388 | + | NA | + | + | NA | NA | NA | + | NA | NA | + | NA | 0,0177502469959505 | 7 | -528,876285889541 | 1072,62078883335 | 24,6592159180559 | 1.53708169828312e-07 |
| 158 | 26,3256352848869 | + | NA | + | + | + | NA | NA | + | NA | NA | NA | NA | 0,0177366508023247 | 7 | -528,877233631982 | 1072,62268431823 | 24,661111402937 | 1.53562563082055e-07 |
| 350 | 26,1371353611247 | + | NA | + | + | + | NA | + | NA | + | NA | NA | NA | 0,0337733355200853 | 8 | -527,750159354263 | 1072,62531870853 | 24,6637457932347 | 1.533604243762e-07 |
| 3572 | 25,5552990361497 | + | + | NA | NA | + | + | + | + | + | NA | + | + | 0,0818359778580818 | 11 | -524,256741485376 | 1072,62548297075 | 24,6639100554623 | 1.53347829230967e-07 |
| 3560 | 25,5280229603448 | + | + | + | NA | NA | + | + | + | + | NA | + | + | 0,0818013970954228 | 11 | -524,259320124688 | 1072,63064024938 | 24,6690673340845 | 1.52952909886347e-07 |
| 1535 | 25,5579346919531 | NA | + | + | + | + | + | + | + | + | NA | + | NA | 0,0817436295603989 | 11 | -524,26362755281 | 1072,63925510562 | 24,6776821903304 | 1.5229549312501e-07 |
| 1366 | 26,1136570345691 | + | NA | + | NA | + | NA | + | NA | + | NA | + | NA | 0,0336733885592301 | 8 | -527,757241467241 | 1072,63948293448 | 24,6779100191914 | 1.5227814545874e-07 |
| 3575 | 25,50495172495 | NA | + | + | NA | + | + | + | + | + | NA | + | + | 0,0816191268736477 | 11 | -524,272910157274 | 1072,65782031455 | 24,6962473992569 | 1.50888335449027e-07 |
| 3229 | 25,8473023346194 | NA | NA | + | + | + | NA | NA | + | NA | NA | + | + | 0,0335429830557629 | 8 | -527,766480732557 | 1072,65796146511 | 24,6963885498237 | 1.50877686837757e-07 |
| 3359 | 25,6890160707249 | NA | + | + | + | + | NA | NA | NA | + | NA | + | + | 0,0495603098812858 | 9 | -526,622221150105 | 1072,66176513486 | 24,7001922195652 | 1.50591015079291e-07 |
| 3199 | 24,8390381609457 | NA | + | + | + | + | + | + | NA | NA | NA | + | + | 0,0654162213987296 | 10 | -525,47034846058 | 1072,68672866719 | 24,7251557519003 | 1.48723055212698e-07 |
| 400 | 26,3445726155203 | + | + | + | + | NA | NA | NA | + | + | NA | NA | NA | 0,0333215400455192 | 8 | -527,782167175673 | 1072,68933435135 | 24,7277614360557 | 1.4852941872471e-07 |
| 3260 | 25,3482020586976 | + | + | NA | + | + | + | NA | + | NA | NA | + | + | 0,0653473290441845 | 10 | -525,475395362766 | 1072,69682247156 | 24,7352495562727 | 1.47974355396763e-07 |
| 3248 | 25,3443010914389 | + | + | + | + | NA | + | NA | + | NA | NA | + | + | 0,0653256010678963 | 10 | -525,476987029333 | 1072,7000058047 | 24,7384328894072 | 1.47739016902357e-07 |
| 3292 | 25,1488244170722 | + | + | NA | + | + | NA | + | + | NA | NA | + | + | 0,0653025925094572 | 10 | -525,478672463666 | 1072,70337667336 | 24,7418037580735 | 1.47490222213378e-07 |
| 1373 | 26,1917918632051 | NA | NA | + | + | + | NA | + | NA | + | NA | + | NA | 0,0331171298336945 | 8 | -527,796643872276 | 1072,71828774455 | 24,756714829261 | 1.46394692566655e-07 |
| 3214 | 25,8666161440979 | + | NA | + | + | NA | NA | NA | + | NA | NA | + | + | 0,0330747049704308 | 8 | -527,799648093579 | 1072,72429618716 | 24,7627232718669 | 1.45955550482894e-07 |
| 512 | 25,5378983442576 | + | + | + | + | + | + | + | + | + | NA | NA | NA | 0,0811585949723636 | 11 | -524,307235319535 | 1072,72647063907 | 24,7648977237786 | 1.4579695005288e-07 |
| 415 | 26,5102524034249 | NA | + | + | + | + | NA | NA | + | + | NA | NA | NA | 0,0328874919538441 | 8 | -527,812903589085 | 1072,75080717817 | 24,7892342628795 | 1.44033603659814e-07 |
| 288 | 26,2406781865327 | + | + | + | + | + | NA | NA | NA | + | NA | NA | NA | 0,0326445663591653 | 8 | -527,830099954549 | 1072,7851999091 | 24,8236269938075 | 1.41577924068847e-07 |
| 3226 | 25,9662148375963 | + | NA | NA | + | + | NA | NA | + | NA | NA | + | + | 0,0326429373114736 | 8 | -527,830215258002 | 1072,785430516 | 24,8238576007134 | 1.41561600586434e-07 |
| 3184 | 24,7793605829394 | + | + | + | + | NA | + | + | NA | NA | NA | + | + | 0,0647397533192729 | 10 | -525,519888929464 | 1072,78580960496 | 24,8242366896684 | 1.41534770909605e-07 |
| 3474 | 26,5596866522025 | + | NA | NA | NA | + | NA | NA | + | + | NA | + | + | 0,032600374914432 | 8 | -527,833227741509 | 1072,79145548302 | 24,8298825677271 | 1.41135790294728e-07 |
| 1433 | 27,1719647293243 | NA | NA | NA | + | + | NA | NA | + | + | NA | + | NA | 0,0163885668647956 | 7 | -528,971138736779 | 1072,81049452782 | 24,848921612532 | 1.39798619687999e-07 |
| 3295 | 25,1499875688893 | NA | + | + | + | + | NA | + | + | NA | NA | + | + | 0,0645503201721666 | 10 | -525,533755459133 | 1072,8135426643 | 24,8519697490076 | 1.39585719330589e-07 |
| 1420 | 26,5885317647887 | + | + | NA | + | NA | NA | NA | + | + | NA | + | NA | 0,0323841601603398 | 8 | -527,848528954811 | 1072,82205790962 | 24,8604849943308 | 1.3899267937476e-07 |
| 2552 | 25,4818694862807 | + | + | + | NA | + | + | + | + | + | NA | NA | + | 0,0804940033137402 | 11 | -524,356739516376 | 1072,82547903275 | 24,8639061174617 | 1.38755127071385e-07 |
| 3354 | 26,1571483759188 | + | NA | NA | + | + | NA | NA | NA | + | NA | + | + | 0,0322886860573969 | 8 | -527,855284434772 | 1072,83556886954 | 24,8739959542531 | 1.38056881555082e-07 |
| 2206 | 25,7116446061231 | + | NA | + | + | + | NA | NA | + | NA | NA | NA | + | 0,0322760516219969 | 8 | -527,8561783621 | 1072,8373567242 | 24,8757838089086 | 1.37933523880489e-07 |
| 1120 | 25,443502596735 | + | + | + | + | + | NA | + | NA | NA | NA | + | NA | 0,0482551826637357 | 9 | -526,716176579434 | 1072,84967599351 | 24,8881030782227 | 1.37086515076855e-07 |
| 3344 | 25,5710819418976 | + | + | + | + | NA | NA | NA | NA | + | NA | + | + | 0,0482524042180519 | 9 | -526,71637646089 | 1072,85007575643 | 24,8885028411348 | 1.37059116762913e-07 |
| 1423 | 26,5445784764757 | NA | + | + | + | NA | NA | NA | + | + | NA | + | NA | 0,0321411799037768 | 8 | -527,865720247206 | 1072,85644049441 | 24,8948675791223 | 1.36623637374992e-07 |
| 3420 | 25,339686642718 | + | + | NA | + | + | NA | + | NA | + | NA | + | + | 0,0642163528177676 | 10 | -525,558195076026 | 1072,86242189808 | 24,9008489827934 | 1.36215647204493e-07 |
| 412 | 26,5748963546422 | + | + | NA | + | + | NA | NA | + | + | NA | NA | NA | 0,032008641178857 | 8 | -527,875095783341 | 1072,87519156668 | 24,9136186513913 | 1.35348703459405e-07 |
| 3263 | 25,2659477308217 | NA | + | + | + | + | + | NA | + | NA | NA | + | + | 0,0641235951873854 | 10 | -525,564981500772 | 1072,87599474758 | 24,9144218322845 | 1.3529435962585e-07 |
| 1311 | 26,4124153791527 | NA | + | + | + | + | NA | NA | NA | + | NA | + | NA | 0,0319586876917953 | 8 | -527,878629064624 | 1072,88225812925 | 24,9206852139569 | 1.3487132227646e-07 |
| 3356 | 25,8270873484683 | + | + | NA | + | + | NA | NA | NA | + | NA | + | + | 0,0479720233342153 | 9 | -526,736544070527 | 1072,8904109757 | 24,9288380604098 | 1.34322648758269e-07 |
| 2176 | 24,8182973950706 | + | + | + | + | + | + | + | NA | NA | NA | NA | + | 0,0639360142421596 | 10 | -525,578703427607 | 1072,90343860125 | 24,9418656859552 | 1.3345053963451e-07 |
| 3288 | 25,3092443193466 | + | + | + | NA | + | NA | + | + | NA | NA | + | + | 0,0639293261211611 | 10 | -525,579192626485 | 1072,904416999 | 24,9428440837105 | 1.33385271746093e-07 |
| 1181 | 26,5765631527402 | NA | NA | + | + | + | NA | NA | + | NA | NA | + | NA | 0,0155986545529697 | 7 | -529,026102853422 | 1072,92042276111 | 24,9588498458174 | 1.32322065313843e-07 |
| 1528 | 25,6763910346951 | + | + | + | NA | + | + | + | + | + | NA | + | NA | 0,0797746830495857 | 11 | -524,410280059336 | 1072,93256011867 | 24,9709872033809 | 1.31521476917686e-07 |
| 3293 | 25,5841841808866 | NA | NA | + | + | + | NA | + | + | NA | NA | + | + | 0,047669346703419 | 9 | -526,758308735694 | 1072,93394030603 | 24,9723673907438 | 1.31430746087479e-07 |
| 2336 | 25,6463305884714 | + | + | + | + | + | NA | NA | NA | + | NA | NA | + | 0,0476434498611574 | 9 | -526,760170587095 | 1072,93766400883 | 24,976091093544 | 1.3118626922927e-07 |
| 3416 | 25,3631584825593 | + | + | + | NA | + | NA | + | NA | + | NA | + | + | 0,0636845444803741 | 10 | -525,59709463956 | 1072,94022102515 | 24,9786481098604 | 1.31018653685571e-07 |
| 3290 | 25,6285531609045 | + | NA | NA | + | + | NA | + | + | NA | NA | + | + | 0,047624087126063 | 9 | -526,76156263632 | 1072,94044810729 | 24,9788751919953 | 1.31003778532264e-07 |
| 3510 | 25,9995719219752 | + | NA | + | NA | + | + | NA | + | + | NA | + | + | 0,0636616530081909 | 10 | -525,598768559554 | 1072,94356886514 | 24,9819959498498 | 1.3079952239678e-07 |
| 1216 | 25,4732514748728 | + | + | + | + | + | + | NA | + | NA | NA | + | NA | 0,0635768113156392 | 10 | -525,604972181519 | 1072,95597610907 | 24,9944031937796 | 1.29990603314995e-07 |
| 3423 | 25,2486444843143 | NA | + | + | + | + | NA | + | NA | + | NA | + | + | 0,063575716101231 | 10 | -525,60505225989 | 1072,95613626581 | 24,9945633505217 | 1.29980194296001e-07 |
| 2430 | 25,6006012050089 | + | NA | + | + | + | + | + | NA | + | NA | NA | + | 0,0635614032875775 | 10 | -525,606098755781 | 1072,95822925759 | 24,9966563423036 | 1.29844241706089e-07 |
| 3453 | 25,5870405294722 | NA | NA | + | + | + | + | + | NA | + | NA | + | + | 0,0635214428877045 | 10 | -525,609020416332 | 1072,9640725787 | 25,0024996634058 | 1.29465434549729e-07 |
| 3464 | 26,0305242061044 | + | + | + | NA | NA | NA | NA | + | + | NA | + | + | 0,047454870687727 | 9 | -526,773726945901 | 1072,96477672645 | 25,003203811156 | 1.29419861175565e-07 |
| 1358 | 25,9071259365237 | + | NA | + | + | NA | NA | + | NA | + | NA | + | NA | 0,0313565417010182 | 8 | -527,921205375271 | 1072,96741075054 | 25,0058378352512 | 1.29249525850657e-07 |
| 1435 | 26,7246787967231 | NA | + | NA | + | + | NA | NA | + | + | NA | + | NA | 0,0313521349075497 | 8 | -527,92151687163 | 1072,96803374326 | 25,0064608279699 | 1.29209271363795e-07 |
| 1360 | 25,5133517083883 | + | + | + | + | NA | NA | + | NA | + | NA | + | NA | 0,0472826940779231 | 9 | -526,786101833892 | 1072,98952650243 | 25,0279535871389 | 1.27828173679143e-07 |
| 2270 | 25,5173478121847 | + | NA | + | + | + | NA | + | + | NA | NA | NA | + | 0,04710592170101 | 9 | -526,798804709014 | 1073,01493225267 | 25,0533593373827 | 1.26214658191461e-07 |
| 1178 | 26,6471509684955 | + | NA | NA | + | + | NA | NA | + | NA | NA | + | NA | 0,0149169676291802 | 7 | -529,073500943773 | 1073,01521894181 | 25,0536460265182 | 1.26196567302487e-07 |
| 1054 | 26,1820097008404 | + | NA | + | + | + | NA | NA | NA | NA | NA | + | NA | 0,0148294292641914 | 7 | -529,079585161956 | 1073,02738737818 | 25,0658144628851 | 1.25431090877157e-07 |
| 3479 | 26,1007430428037 | NA | + | + | NA | + | NA | NA | + | + | NA | + | + | 0,0469415913993051 | 9 | -526,810611383336 | 1073,03854560132 | 25,0769726860276 | 1.24733245315895e-07 |
| 3278 | 25,6491553549143 | + | NA | + | + | NA | NA | + | + | NA | NA | + | + | 0,0469370459347445 | 9 | -526,810937933379 | 1073,0391987014 | 25,0776257861132 | 1.24692520319033e-07 |
| 1306 | 26,8075973865252 | + | NA | NA | + | + | NA | NA | NA | + | NA | + | NA | 0,0143389959513705 | 7 | -529,11366197683 | 1073,09554100792 | 25,1339680926324 | 1.21228805704401e-07 |
| 1406 | 25,7017739851552 | + | NA | + | + | + | + | + | NA | + | NA | + | NA | 0,0625863461052176 | 10 | -525,677353433392 | 1073,10073861282 | 25,1391656975245 | 1.20914165009191e-07 |
| 2400 | 25,2409769482187 | + | + | + | + | + | NA | + | NA | + | NA | NA | + | 0,0624935785788465 | 10 | -525,684128786464 | 1073,11428931896 | 25,1527164036695 | 1.20097697899609e-07 |
| 3390 | 25,6841357539193 | + | NA | + | + | + | + | NA | NA | + | NA | + | + | 0,06241808334737 | 10 | -525,68964214862 | 1073,12531604327 | 25,1637431279814 | 1.19437377761674e-07 |
| 3446 | 25,6776881833774 | + | NA | + | NA | + | + | + | NA | + | NA | + | + | 0,0623229973314881 | 10 | -525,696585580228 | 1073,13920290649 | 25,1776299911976 | 1.18610944957841e-07 |
| 3438 | 25,4610396304162 | + | NA | + | + | NA | + | + | NA | + | NA | + | + | 0,0620692282219119 | 10 | -525,715113024794 | 1073,17625779562 | 25,2146848803284 | 1.16433619670228e-07 |
| 3352 | 25,8950561839501 | + | + | + | NA | + | NA | NA | NA | + | NA | + | + | 0,0458925057722283 | 9 | -526,885937336099 | 1073,18919750684 | 25,2276245915534 | 1.15682742614831e-07 |
| 2456 | 25,9695556322379 | + | + | + | NA | + | NA | NA | + | + | NA | NA | + | 0,0458379860713016 | 9 | -526,889849670742 | 1073,19702217613 | 25,2354492608383 | 1.15231037200898e-07 |
| 1370 | 26,2982373005432 | + | NA | NA | + | + | NA | + | NA | + | NA | + | NA | 0,0297130032261715 | 8 | -528,037281559623 | 1073,19956311925 | 25,237990203955 | 1.15084732403221e-07 |
| 408 | 26,5334713954269 | + | + | + | NA | + | NA | NA | + | + | NA | NA | NA | 0,0295670105595766 | 8 | -528,04758289959 | 1073,22016579918 | 25,2585928838901 | 1.13905290794095e-07 |
| 1308 | 26,490978420432 | + | + | NA | + | + | NA | NA | NA | + | NA | + | NA | 0,0295512477700436 | 8 | -528,048695039882 | 1073,22239007976 | 25,2608171644736 | 1.13778682546876e-07 |
| 2272 | 25,0871355654359 | + | + | + | + | + | NA | + | + | NA | NA | NA | + | 0,0616684774166814 | 10 | -525,744361263458 | 1073,23475427295 | 25,273181357657 | 1.13077461482758e-07 |
| 3452 | 25,3370842120313 | + | + | NA | + | + | + | + | NA | + | NA | + | + | 0,077736734475419 | 11 | -524,561742082803 | 1073,23548416561 | 25,2739112503145 | 1.13036201807564e-07 |
| 3286 | 25,6729443673915 | + | NA | + | NA | + | NA | + | + | NA | NA | + | + | 0,0455424353012194 | 9 | -526,911054510832 | 1073,23943185631 | 25,2778589410193 | 1.12813305879389e-07 |
| 3476 | 26,2225368949937 | + | + | NA | NA | + | NA | NA | + | + | NA | + | + | 0,0454894221238405 | 9 | -526,914857345704 | 1073,24703752605 | 25,2854646107626 | 1.12385110200567e-07 |
| 1174 | 26,4884270178224 | + | NA | + | NA | + | NA | NA | + | NA | NA | + | NA | 0,0132205960009577 | 7 | -529,191308495132 | 1073,25083404453 | 25,2892611292361 | 1.12171976482577e-07 |

| | | | | | | | | | | | | | | | | | | |
|---|---|---|---|---|---|---|---|---|---|---|---|---|---|---|---|---|---|---|
| **3504** | 25,6492754477818 | + | + | + | + | NA | + | NA | + | + | NA | + | + | 0,0775970718344577 | 11 | -524,572109674059 | 1073,25621934812 | 25,2946464328265 | 1.11870342688365e-07 |
| **3408** | 25,1289441385319 | + | + | + | + | NA | NA | + | NA | + | NA | + | + | 0,0614684668590798 | 10 | -525,758954084606 | 1073,26393991524 | 25,3023669999529 | 1.11439323905882e-07 |
| **2398** | 25,6630745608532 | + | NA | + | + | + | NA | + | NA | + | NA | NA | + | 0,0453613851979606 | 9 | -526,92404104602 | 1073,26540492668 | 25,303832011394 | 1.11357723853532e-07 |
| **3421** | 25,6771217305717 | NA | NA | + | + | + | NA | + | NA | + | NA | + | + | 0,0453472309397154 | 9 | -526,925056212454 | 1073,26743525955 | 25,3058623442621 | 1.11244734591259e-07 |
| **1296** | 26,1020926652092 | + | + | + | + | NA | NA | NA | NA | + | NA | + | NA | 0,0292221099779675 | 8 | -528,071913163543 | 1073,26882632709 | 25,3072534117946 | 1.11167387024074e-07 |
| **1304** | 26,4722640513643 | + | + | + | NA | + | NA | NA | NA | + | NA | + | NA | 0,0291973413840602 | 8 | -528,073660077776 | 1073,27232015555 | 25,3107472402608 | 1.10973356659973e-07 |
| **3136** | 25,1755129416334 | + | + | + | + | + | + | NA | NA | NA | NA | + | + | 0,0613814277894214 | 10 | -525,765303506607 | 1073,27663875925 | 25,3150658439545 | 1.10733990211198e-07 |
| **3516** | 25,7161609920766 | + | + | NA | + | + | + | NA | + | + | NA | + | + | 0,0773835792753693 | 11 | -524,58795485722 | 1073,28790971444 | 25,3263367991501 | 1.10111706370273e-07 |
| **3519** | 25,6230921610955 | NA | + | + | + | + | + | NA | + | + | NA | + | + | 0,0773137399534049 | 11 | -524,593137459446 | 1073,29827491889 | 25,3367020036007 | 1.09542517409796e-07 |
| **1118** | 25,7992530159596 | + | NA | + | + | + | NA | + | NA | NA | NA | + | NA | 0,0287502187237921 | 8 | -528,105187715089 | 1073,33537543018 | 25,3738025148866 | 1.07529207151415e-07 |
| **1416** | 26,5500759168827 | + | + | + | NA | NA | NA | NA | + | + | NA | + | NA | 0,0287445751840997 | 8 | -528,10558556121 | 1073,33617112242 | 25,3745982071289 | 1.07486435582257e-07 |
| **3448** | 25,3633374013534 | + | + | + | NA | + | + | + | NA | + | NA | + | + | 0,07698861938119 | 11 | -524,617258687637 | 1073,34651737527 | 25,3849444599844 | 1.06931830419861e-07 |
| **3455** | 25,226796649552 | NA | + | + | + | + | + | + | NA | + | NA | + | + | 0,0769544721210501 | 11 | -524,619791636006 | 1073,35158327201 | 25,3900103567219 | 1.06661320353076e-07 |
| **3102** | 25,6396879874136 | + | NA | + | + | + | NA | NA | NA | NA | NA | + | + | 0,0286264015103615 | 8 | -528,113915783923 | 1073,35283156785 | 25,3912586525562 | 1.06594768683339e-07 |
| **2240** | 25,2010003719747 | + | + | + | + | + | + | NA | + | NA | NA | NA | + | 0,06084455678928 | 10 | -525,804454758484 | 1073,354941263 | 25,3933683477098 | 1.06482386733226e-07 |
| **1429** | 27,1813626132528 | NA | NA | + | NA | + | NA | NA | + | + | NA | + | NA | 0,012410226205633 | 7 | -529,247514638457 | 1073,36324633118 | 25,4016734158879 | 1.06041131789599e-07 |
| **1426** | 27,235566779996 | + | NA | NA | NA | + | NA | NA | + | + | NA | + | NA | 0,0123921500938484 | 7 | -529,248767847223 | 1073,36575274871 | 25,4041798334181 | 1.05908323349512e-07 |
| **3222** | 25,9727185508163 | + | NA | + | NA | + | NA | NA | + | NA | NA | + | + | 0,0285035499716529 | 8 | -528,122574681849 | 1073,3701493637 | 25,4085764484075 | 1.05675760005067e-07 |
| **3414** | 25,7253442728994 | + | NA | + | NA | + | NA | + | NA | + | NA | + | + | 0,0445788846869593 | 9 | -526,98014069355 | 1073,37760422175 | 25,4160313064547 | 1.05282594313952e-07 |
| **3216** | 25,6025885128841 | + | + | + | + | NA | NA | NA | + | NA | NA | + | + | 0,0445633167467617 | 9 | -526,981256336679 | 1073,379835508 | 25,4182625927135 | 1.05165202007094e-07 |
| **3228** | 25,6335531051615 | + | + | NA | + | + | NA | NA | + | NA | NA | + | + | 0,0444973538693615 | 9 | -526,985983223583 | 1073,38928928181 | 25,4277163665217 | 1.04669271019206e-07 |
| **3256** | 25,4950469894862 | + | + | + | NA | + | + | NA | + | NA | NA | + | + | 0,0605698226593194 | 10 | -525,824481059149 | 1073,39499386433 | 25,4334209490396 | 1.0437114913973e-07 |
| **1408** | 25,4200791306487 | + | + | + | + | + | + | + | NA | + | NA | + | NA | 0,0766594612099891 | 11 | -524,641670822184 | 1073,39534164437 | 25,4337687290783 | 1.04353001616467e-07 |
| **2432** | 25,2354773343868 | + | + | + | + | + | + | + | NA | + | NA | NA | + | 0,0766479673908518 | 11 | -524,642523108247 | 1073,39704621649 | 25,435473301203 | 1.04264100897428e-07 |
| **3406** | 25,5520917457552 | + | NA | + | + | NA | NA | + | NA | + | NA | + | + | 0,0444106783418113 | 9 | -526,992193879095 | 1073,40171059283 | 25,440137677544 | 1.04021220725926e-07 |
| **3231** | 25,5532025738305 | NA | + | + | + | + | NA | NA | + | NA | NA | + | + | 0,0443015689607455 | 9 | -527,000011211334 | 1073,41734525731 | 25,4557723420221 | 1.0321122242149e-07 |
| **3450** | 25,7753933734807 | + | NA | NA | + | + | + | + | NA | + | NA | + | + | 0,0603398007749479 | 10 | -525,841243631017 | 1073,42851900807 | 25,4669460927753 | 1.02636201963857e-07 |
| **3322** | 25,4946869330081 | + | NA | NA | + | + | + | + | + | NA | NA | + | + | 0,0602527953410369 | 10 | -525,847582979705 | 1073,44119770544 | 25,4796247901518 | 1.01987613278573e-07 |
| **2496** | 25,6023090383649 | + | + | + | + | + | + | NA | + | + | NA | NA | + | 0,0761805796384923 | 11 | -524,677171717838 | 1073,46634343568 | 25,5047705203849 | 1.00713364040896e-07 |
| **3325** | 25,4164713588157 | NA | NA | + | + | + | + | + | + | NA | NA | + | + | 0,0600267992512459 | 10 | -525,864046659119 | 1073,47412506427 | 25,5125521489797 | 1.00322268377827e-07 |
| **3310** | 25,4972024784631 | + | NA | + | + | NA | + | + | + | NA | NA | + | + | 0,0599893979824258 | 10 | -525,866770937513 | 1073,47957362106 | 25,5180007057672 | 1.00049334532363e-07 |
| **1472** | 25,7823377837131 | + | + | + | + | + | + | NA | + | + | NA | + | NA | 0,0758660649746331 | 11 | -524,700477606499 | 1073,512955213 | 25,5513822977077 | 9.83932902962131e-08 |
| **1183** | 26,2459200663285 | NA | + | + | + | + | NA | NA | + | NA | NA | + | NA | 0,0274242825679377 | 8 | -528,198597263762 | 1073,52219452752 | 25,5606216122326 | 9.79397953203134e-08 |
| **1168** | 26,1386674075151 | + | + | + | + | NA | NA | NA | + | NA | NA | + | NA | 0,0272981253565771 | 8 | -528,207478154161 | 1073,53995630832 | 25,5783833930316 | 9.70738535905319e-08 |
| **2302** | 25,3728948821761 | + | NA | + | + | + | + | + | + | NA | NA | NA | + | 0,0594967437974385 | 10 | -525,902645363649 | 1073,55132247333 | 25,58974955804 | 9.65237395239927e-08 |
| **1500** | 25,8991120268392 | + | + | NA | + | + | NA | + | + | + | NA | + | NA | 0,0592076322292732 | 10 | -525,923689340998 | 1073,59341042803 | 25,6318375127378 | 9.45137197188338e-08 |
| **1180** | 26,3100226587471 | + | + | NA | + | + | NA | NA | + | NA | NA | + | NA | 0,0269056654474766 | 8 | -528,235098175961 | 1073,59519635192 | 25,6336234366318 | 9.44293602341669e-08 |
| **1503** | 25,9061946278099 | NA | + | + | + | + | NA | + | + | + | NA | + | NA | 0,0591863864561754 | 10 | -525,925235532512 | 1073,59650281106 | 25,6349298957657 | 9.43676963266076e-08 |
| **3312** | 25,1519620296655 | + | + | + | + | NA | + | + | + | NA | NA | + | + | 0,0751183974746014 | 11 | -524,755848795964 | 1073,62369759193 | 25,6621246766374 | 9.30932262847356e-08 |
| **480** | 25,8424387114409 | + | + | + | + | + | NA | + | + | + | NA | NA | NA | 0,0586874732610275 | 10 | -525,961534624448 | 1073,66910099493 | 25,7075280796371 | 9.10036597482592e-08 |
| **1488** | 25,876244443153 | + | + | + | + | NA | NA | + | + | + | NA | + | NA | 0,0586854628832585 | 10 | -525,961680853248 | 1073,66939345253 | 25,707820537237 | 9.09903533652139e-08 |
| **160** | 26,0194658498996 | + | + | + | + | + | NA | NA | + | NA | NA | NA | NA | 0,0263250166966283 | 8 | -528,275941885442 | 1073,67688377088 | 25,7153108555931 | 9.06502173375466e-08 |
| **1278** | 25,5833518498136 | + | NA | + | + | + | + | + | + | NA | NA | + | NA | 0,0586144959261753 | 10 | -525,966842575125 | 1073,67971689628 | 25,7181439809917 | 9.05218965296739e-08 |
| **3440** | 25,1039179920776 | + | + | + | + | NA | + | + | NA | + | NA | + | + | 0,0747283213278342 | 11 | -524,784719521825 | 1073,68143904365 | 25,7198661283601 | 9.04439840557349e-08 |
| **3324** | 25,0772634209376 | + | + | NA | + | + | + | + | + | NA | NA | + | + | 0,0745934593564161 | 11 | -524,794698237808 | 1073,70139647562 | 25,7398235603252 | 8.95459572554241e-08 |
| **1056** | 26,0052439598632 | + | + | + | + | + | NA | NA | NA | NA | NA | + | NA | 0,0260932127378548 | 8 | -528,292240526416 | 1073,70948105283 | 25,7479081375423 | 8.91847172625184e-08 |
| **3418** | 25,837704683058 | + | NA | NA | + | + | NA | + | NA | + | NA | + | + | 0,0421824288062109 | 9 | -527,15166405019 | 1073,72065093503 | 25,7590780197345 | 8.86880141897136e-08 |
| **1431** | 26,8384688017277 | NA | + | + | NA | + | NA | NA | + | + | NA | + | NA | 0,025727074402646 | 8 | -528,317976612121 | 1073,76095322424 | 25,7993803089516 | 8.69187355558385e-08 |
| **3168** | 25,0366627228307 | + | + | + | + | + | NA | + | NA | NA | NA | + | + | 0,0580036021322511 | 10 | -526,011259343405 | 1073,76855043284 | 25,8069775175511 | 8.65891919723249e-08 |
| **2462** | 26,0695400330689 | + | NA | + | + | + | NA | NA | + | + | NA | NA | + | 0,0417545975742115 | 9 | -527,182240408671 | 1073,78180365199 | 25,8202307366969 | 8.60172961617724e-08 |
| **3318** | 25,549877963648 | + | NA | + | NA | + | + | + | + | NA | NA | + | + | 0,0577881109530859 | 10 | -526,026920372596 | 1073,79987249122 | 25,8382995759337 | 8.52436796637631e-08 |
| **3392** | 25,489523247161 | + | + | + | + | + | + | NA | NA | + | NA | + | + | 0,0738323329389269 | 11 | -524,850988318035 | 1073,81397663607 | 25,8524037207799 | 8.46446497449117e-08 |
| **1496** | 25,9475566361425 | + | + | + | NA | + | NA | + | + | + | NA | + | NA | 0,0576740556978357 | 10 | -526,035207999372 | 1073,81644774478 | 25,8548748294861 | 8.45401312618812e-08 |
| **1428** | 26,8974850771782 | + | + | NA | NA | + | NA | NA | + | + | NA | + | NA | 0,0253124649381455 | 8 | -528,347108093565 | 1073,81921618713 | 25,8576432718387 | 8.44231899769155e-08 |
| **3470** | 26,1375584341063 | + | NA | + | + | NA | NA | NA | + | + | NA | + | + | 0,0414391538654021 | 9 | -527,204775887065 | 1073,82687460878 | 25,8653016934854 | 8.41005339373668e-08 |
| **3485** | 26,1566946360261 | NA | NA | + | + | + | NA | NA | + | + | NA | + | + | 0,0413980560944616 | 9 | -527,207711389446 | 1073,83274561354 | 25,8711726982476 | 8.38540186199153e-08 |
| **1486** | 26,2618626535347 | + | NA | + | + | NA | NA | + | + | + | NA | + | NA | 0,0412038234051757 | 9 | -527,22158320203 | 1073,8604892387 | 25,8989163234144 | 8.26988421091735e-08 |
| **3104** | 25,4301464995209 | + | + | + | + | + | NA | NA | NA | NA | NA | + | + | 0,04107975850815 | 9 | -527,230442264242 | 1073,87820736313 | 25,9166344478392 | 8.1969443585922e-08 |
| **3512** | 25,8031379637025 | + | + | + | NA | + | + | NA | + | + | NA | + | + | 0,0731097845903034 | 11 | -524,904382542507 | 1073,92076508501 | 25,9591921697229 | 8.02436537333842e-08 |
| **3327** | 25,0564556222035 | NA | + | + | + | + | + | + | + | NA | NA | + | + | 0,0730987776448099 | 11 | -524,905195602062 | 1073,92239120412 | 25,9608182888335 | 8.01784373799466e-08 |
| **478** | 26,3284960056922 | + | NA | + | + | + | NA | + | + | + | NA | NA | NA | 0,0407670622911397 | 9 | -527,252765780753 | 1073,92285439615 | 25,9612814808613 | 8.01598705235351e-08 |
| **1422** | 26,616018384606 | + | NA | + | + | NA | NA | NA | + | + | NA | + | NA | 0,0243960630587624 | 8 | -528,41145282533 | 1073,94790565066 | 25,9863327353701 | 7.91620798867536e-08 |
| **1280** | 25,3020235472882 | + | + | + | + | + | + | + | + | NA | NA | + | NA | 0,0727966578603639 | 11 | -524,927508778297 | 1073,96701755659 | 26,0054446413028 | 7.84092136807305e-08 |
| **414** | 26,6735095358688 | + | NA | + | + | + | NA | NA | + | + | NA | NA | NA | 0,0242360699513152 | 8 | -528,422680472222 | 1073,97036094444 | 26,0087880291542 | 7.82782469731268e-08 |
| **3320** | 25,2364437988169 | + | + | + | NA | + | + | + | + | NA | NA | + | + | 0,0727713031092928 | 11 | -524,929381032929 | 1073,97076206586 | 26,0091891505676 | 7.82625490068425e-08 |
| **1498** | 26,4052404461072 | + | NA | NA | + | + | NA | + | + | + | NA | + | NA | 0,0398922263124561 | 9 | -527,315182074264 | 1074,04768698317 | 26,0861140678821 | 7.53095331160645e-08 |
| **3482** | 26,2805559156767 | + | NA | NA | + | + | NA | NA | + | + | NA | + | + | 0,0398417775308955 | 9 | -527,318779673397 | 1074,05488218144 | 26,0933092661498 | 7.50390863761268e-08 |
| **2208** | 25,4452865846729 | + | + | + | + | + | NA | NA | + | NA | NA | NA | + | 0,0398408575307495 | 9 | -527,318845278614 | 1074,05501339187 | 26,0934404765826 | 7.50341635821105e-08 |
| **3536** | 25,4895249242763 | + | + | + | + | NA | NA | + | + | + | NA | + | + | 0,0719428895755743 | 11 | -524,99052490958 | 1074,09304981916 | 26,1314769038688 | 7.36206318577922e-08 |
| **1494** | 26,3178500096737 | + | NA | + | NA | + | NA | + | + | + | NA | + | NA | 0,0395725161333757 | 9 | -527,337978024004 | 1074,09327888265 | 26,1317059673634 | 7.36122004410319e-08 |
| **1501** | 26,4033274425461 | NA | NA | + | + | + | NA | + | + | + | NA | + | NA | 0,0395218561133083 | 9 | -527,341589484545 | 1074,10050180374 | 26,1389288884457 | 7.3346832355402e-08 |
| **3548** | 25,4551484503632 | + | + | NA | + | + | NA | + | + | + | NA | + | + | 0,0718164205674172 | 11 | -524,99985458443 | 1074,11170916886 | 26,1501362535687 | 7.29369694333612e-08 |
| **3262** | 25,5602197165494 | + | NA | + | + | + | + | NA | + | NA | NA | + | + | 0,0554080503316146 | 10 | -526,199655825584 | 1074,1453433972 | 26,1837704819097 | 7.17206363635355e-08 |
| **1176** | 26,2961733927223 | + | + | + | NA | + | NA | NA | + | NA | NA | + | NA | 0,0228761201746565 | 8 | -528,518041828016 | 1074,16108365603 | 26,1995107407417 | 7.11584010080798e-08 |
| **3551** | 25,4474697184272 | NA | + | + | + | + | NA | + | + | + | NA | + | + | 0,0714011075598156 | 11 | -525,030483474275 | 1074,17296694855 | 26,2113940332597 | 7.07368565351593e-08 |
| **3224** | 25,7599542420357 | + | + | + | NA | + | NA | NA | + | NA | NA | + | + | 0,0389222645962465 | 9 | -527,384318814999 | 1074,18596046464 | 26,2243875493521 | 7.02787858915535e-08 |

| | | | | | | | | | | | | | | | | | | |
|---|---|---|---|---|---|---|---|---|---|---|---|---|---|---|---|---|---|---|
| **2304** | 25,0078710405996 | + | + | + | + | + | + | + | + | NA | NA | NA | + | 0,0708774985323337 | 11 | -525,069079572277 | 1074,25015914455 | 26,2885862292635 | 6.80587054318929e-08 |
| **3166** | 25,3944703224239 | + | NA | + | + | + | NA | + | NA | NA | NA | + | + | 0,0384301054233383 | 9 | -527,419372172358 | 1074,25606717936 | 26,2944942640718 | 6.78579554869421e-08 |
| **1248** | 25,6507482606934 | + | + | + | + | + | NA | + | + | NA | NA | + | NA | 0,0546093586760371 | 10 | -526,257524254497 | 1074,26108025503 | 26,2995073397356 | 6.76880799429983e-08 |
| **3581** | 25,7709474669791 | NA | NA | + | + | + | + | + | + | + | NA | + | + | 0,0705412937767034 | 11 | -525,093850327944 | 1074,29970065589 | 26,3381277405972 | 6.63935486078622e-08 |
| **2558** | 25,7641022286482 | + | NA | + | + | + | + | + | + | + | NA | NA | + | 0,0705310720494267 | 11 | -525,09460329987 | 1074,30120659974 | 26,3396336844489 | 6.63435749464452e-08 |
| **1310** | 26,4612365933692 | + | NA | + | + | + | NA | NA | NA | + | NA | + | NA | 0,0217211743061807 | 8 | -528,598923941094 | 1074,32284788219 | 26,3612749668964 | 6.56295649143605e-08 |
| **3566** | 25,7995790034113 | + | NA | + | + | NA | + | + | + | + | NA | + | + | 0,0703356671435411 | 11 | -525,108995988983 | 1074,32999197797 | 26,3684190626757 | 6.53955511684894e-08 |
| **2526** | 25,8418678501577 | + | NA | + | + | + | NA | + | + | + | NA | NA | + | 0,0540403317179907 | 10 | -526,298722749131 | 1074,34347724429 | 26,381904329003 | 6.49560961632866e-08 |
| **3549** | 25,8760290752014 | NA | NA | + | + | + | NA | + | + | + | NA | + | + | 0,0539500605965825 | 10 | -526,305256251269 | 1074,35654424857 | 26,3949713332797 | 6.45330887348019e-08 |
| **2528** | 25,4217568317484 | + | + | + | + | + | NA | + | + | + | NA | NA | + | 0,0698965873404892 | 11 | -525,141325700365 | 1074,39465140073 | 26,4330784854385 | 6.33151425803947e-08 |
| **3578** | 25,8619225266492 | + | NA | NA | + | + | + | + | + | + | NA | + | + | 0,0698915210281315 | 11 | -525,141698646975 | 1074,39539729395 | 26,4338243786597 | 6.32915338152489e-08 |
| **3544** | 25,5600121129212 | + | + | + | NA | + | NA | + | + | + | NA | + | + | 0,0698474603579234 | 11 | -525,14494200086 | 1074,40188400172 | 26,4403110864289 | 6.3086589506303e-08 |
| **3546** | 25,9285927616601 | + | NA | NA | + | + | NA | + | + | + | NA | + | + | 0,0535708283247195 | 10 | -526,332696925992 | 1074,41142559802 | 26,4498526827249 | 6.27863329231972e-08 |
| **3534** | 25,9028184427562 | + | NA | + | + | NA | NA | + | + | + | NA | + | + | 0,0533861645985673 | 10 | -526,34605493591 | 1074,43814161785 | 26,4765687025604 | 6.19532092904489e-08 |
| **3358** | 25,9183144749106 | + | NA | + | + | + | NA | NA | NA | + | NA | + | + | 0,037107110615578 | 9 | -527,513511826992 | 1074,44434648863 | 26,482773573339 | 6.17613013064515e-08 |
| **1376** | 25,7056439148067 | + | + | + | + | + | NA | + | NA | + | NA | + | NA | 0,0533307365780407 | 10 | -526,350063921386 | 1074,4461595888 | 26,4845866735122 | 6.17053369644761e-08 |
| **3478** | 26,24541314909 | + | NA | + | NA | + | NA | NA | + | + | NA | + | + | 0,0369442702332135 | 9 | -527,525090038502 | 1074,46750291165 | 26,5059299963598 | 6.10503396711757e-08 |
| **1437** | 26,8711596702943 | NA | NA | + | + | + | NA | NA | + | + | NA | + | NA | 0,0206321774011888 | 8 | -528,675100197749 | 1074,4752003955 | 26,5136274802064 | 6.0815824253596e-08 |
| **3487** | 25,8673216143503 | NA | + | + | + | + | NA | NA | + | + | NA | + | + | 0,0529699510382174 | 10 | -526,376153004848 | 1074,49833775573 | 26,5367648404365 | 6.0116319412213e-08 |
| **3472** | 25,8737152915233 | + | + | + | + | NA | NA | NA | + | + | NA | + | + | 0,0529670484811365 | 10 | -526,376362853944 | 1074,49875745392 | 26,5371845386285 | 6.01037053804976e-08 |
| **1534** | 25,8894300765008 | + | NA | + | + | + | + | + | + | + | NA | + | NA | 0,0689052917699954 | 11 | -525,214259330057 | 1074,54051866011 | 26,5789457448234 | 5.88617156364659e-08 |
| **1430** | 26,7581131345105 | + | NA | + | NA | + | NA | NA | + | + | NA | + | NA | 0,0200449806658337 | 8 | -528,716139980344 | 1074,55727996069 | 26,5957070453974 | 5.83704775051022e-08 |
| **1246** | 26,0131697072945 | + | NA | + | + | + | NA | + | + | NA | NA | + | NA | 0,0362688261545602 | 9 | -527,573094309353 | 1074,56351145335 | 26,6019385380616 | 5.81888929365378e-08 |
| **1434** | 26,9452651673975 | + | NA | NA | + | + | NA | NA | + | + | NA | + | NA | 0,0199442043828578 | 8 | -528,723180866852 | 1074,5713617337 | 26,6097888184124 | 5.79609410393515e-08 |
| **3198** | 25,2725115487376 | + | NA | + | + | + | + | + | NA | NA | NA | + | + | 0,0524598574547177 | 10 | -526,413021887698 | 1074,57207552143 | 26,610502606138 | 5.79402588261111e-08 |
| **3484** | 25,9533342597689 | + | + | NA | + | + | NA | NA | + | + | NA | + | + | 0,0524242304670329 | 10 | -526,415596217537 | 1074,5772241811 | 26,6156512658142 | 5.77912933144355e-08 |
| **3200** | 24,9661318724729 | + | + | + | + | + | + | + | NA | NA | NA | + | + | 0,0681853110488447 | 11 | -525,267182587692 | 1074,64636517538 | 26,6847922600925 | 5.58275587845671e-08 |
| **3580** | 25,4450244923954 | + | + | NA | + | + | + | + | + | + | NA | + | + | 0,0843499297331896 | 12 | -524,069019077433 | 1074,65416718712 | 26,6925942718335 | 5.56101993858038e-08 |
| **3568** | 25,4504798891768 | + | + | + | + | NA | + | + | + | + | NA | + | + | 0,0843454752894185 | 12 | -524,069352156089 | 1074,65483334444 | 26,6932604291462 | 5.55916798997068e-08 |
| **3542** | 25,927779659178 | + | NA | + | NA | + | NA | + | + | + | NA | + | + | 0,0518425956544136 | 10 | -526,457610226719 | 1074,66125219947 | 26,6996792841792 | 5.54135484354632e-08 |
| **3574** | 25,8572900310162 | + | NA | + | NA | + | + | + | + | + | NA | + | + | 0,0678616535993014 | 11 | -525,290960196254 | 1074,69392039251 | 26,7323474772172 | 5.45157703445653e-08 |
| **3518** | 25,8690267240196 | + | NA | + | + | + | + | NA | + | + | NA | + | + | 0,0677041757754067 | 11 | -525,302526375841 | 1074,71705275168 | 26,7554798363915 | 5.38888636011857e-08 |
| **3583** | 25,4107504450312 | NA | + | + | + | + | + | + | + | + | NA | + | + | 0,0835546875711523 | 12 | -524,128457235362 | 1074,77304350298 | 26,8114705876908 | 5.24011463866313e-08 |
| **2464** | 25,8057978075073 | + | + | + | + | + | NA | NA | + | + | NA | NA | + | 0,0509478359447575 | 10 | -526,522192336867 | 1074,79041641977 | 26,8288435044747 | 5.19479372497305e-08 |
| **416** | 26,3802538119077 | + | + | + | + | + | NA | NA | + | + | NA | NA | NA | 0,0343605788861272 | 9 | -527,708533249078 | 1074,8343893328 | 26,8728164175106 | 5.08182505970037e-08 |
| **1374** | 26,0678443379368 | + | NA | + | + | + | NA | + | NA | + | NA | + | NA | 0,0342328033555711 | 9 | -527,717592631541 | 1074,85250809773 | 26,8909351824379 | 5.0359947733671e-08 |
| **1182** | 26,3561506396595 | + | NA | + | + | + | NA | NA | + | NA | NA | + | NA | 0,017789570997942 | 8 | -528,87354467901 | 1074,87208935802 | 26,9105164427285 | 4.98692979160446e-08 |
| **2560** | 25,4005164881315 | + | + | + | + | + | + | + | + | + | NA | NA | + | 0,0827722782227038 | 12 | -524,186885950168 | 1074,88990093259 | 26,9283280173036 | 4.94271443418416e-08 |
| **1424** | 26,4037551356636 | + | + | + | + | NA | NA | NA | + | + | NA | + | NA | 0,0338837169248544 | 9 | -527,742337019064 | 1074,90199687277 | 26,9404239574835 | 4.91291126031981e-08 |
| **3230** | 25,8240492656735 | + | NA | + | + | + | NA | NA | + | NA | NA | + | + | 0,0335947215200864 | 9 | -527,762815197447 | 1074,94295322954 | 26,9813803142481 | 4.81332692017685e-08 |
| **3296** | 25,2355136589335 | + | + | + | + | + | NA | + | + | NA | NA | + | + | 0,0659613170434471 | 11 | -525,430402838067 | 1074,97280567613 | 27,011232760844 | 4.74201565565013e-08 |
| **3360** | 25,709306778591 | + | + | + | + | + | NA | NA | NA | + | NA | + | + | 0,0496272037720835 | 10 | -526,617402020081 | 1074,98083578619 | 27,0192628709024 | 4.72301437296734e-08 |
| **1536** | 25,6031805975327 | + | + | + | + | + | + | + | + | + | NA | + | NA | 0,0821273403375623 | 12 | -524,235011135705 | 1074,98615130367 | 27,0245783883774 | 4.71047840641654e-08 |
| **1439** | 26,5514961830573 | NA | + | + | + | + | NA | NA | + | + | NA | + | NA | 0,0332183654429962 | 9 | -527,789474567859 | 1074,99627197036 | 27,034699055072 | 4.68670202440107e-08 |
| **3576** | 25,5389260644263 | + | + | + | NA | + | + | + | + | + | NA | + | + | 0,0818632781394493 | 12 | -524,254705672846 | 1075,02554037795 | 27,0639674626602 | 4.61861528458334e-08 |
| **1312** | 26,2844286664836 | + | + | + | + | + | NA | NA | NA | + | NA | + | NA | 0,0329815591912626 | 9 | -527,806243540334 | 1075,02980991531 | 27,0682370000238 | 4.6087661258987e-08 |
| **3264** | 25,3506332205244 | + | + | + | + | + | + | NA | + | NA | NA | + | + | 0,0653477078118288 | 11 | -525,475367616099 | 1075,0627352322 | 27,101162316907 | 4.53351470211462e-08 |
| **1436** | 26,6173262925959 | + | + | NA | + | + | NA | NA | + | + | NA | + | NA | 0,032550798572036 | 9 | -527,836736491376 | 1075,0907958174 | 27,1292229021069 | 4.47035229398999e-08 |
| **3424** | 25,3039617706887 | + | + | + | + | + | NA | + | NA | + | NA | + | + | 0,0644812345256384 | 11 | -525,538811838512 | 1075,18962367702 | 27,2280507617331 | 4.25482353457466e-08 |
| **3294** | 25,5961329384926 | + | NA | + | + | + | NA | + | + | NA | NA | + | + | 0,0476848040146775 | 10 | -526,757197409422 | 1075,26042656488 | 27,2988536495848 | 4.10683165661545e-08 |
| **3480** | 26,0332164551218 | + | + | + | NA | + | NA | NA | + | + | NA | + | + | 0,0474587236404753 | 10 | -526,77344999619 | 1075,29293173841 | 27,3313588231222 | 4.04062449495153e-08 |
| **3454** | 25,6032045245322 | + | NA | + | + | + | + | + | NA | + | NA | + | + | 0,0635626310944918 | 11 | -525,60600898406 | 1075,32401796812 | 27,3624450528293 | 3.97830616940246e-08 |
| **1432** | 26,5663125336284 | + | + | + | NA | + | NA | NA | + | + | NA | + | NA | 0,0297467852077986 | 9 | -528,034897659734 | 1075,48711815411 | 27,525545238823 | 3.66675124299272e-08 |
| **3456** | 25,2908901599299 | + | + | + | + | + | + | + | NA | + | NA | + | + | 0,0781785132220404 | 12 | -524,528937139616 | 1075,57400331149 | 27,6124303962001 | 3.51086860068624e-08 |
| **3422** | 25,6666993522653 | + | NA | + | + | + | NA | + | NA | + | NA | + | + | 0,0453637686086654 | 10 | -526,923870102445 | 1075,59377195092 | 27,6321990356319 | 3.47633699481675e-08 |
| **3520** | 25,6594510020169 | + | + | + | + | + | + | NA | + | + | NA | + | + | 0,0776491808616164 | 12 | -524,56824164277 | 1075,6526123178 | 27,6910394025078 | 3.37555234358492e-08 |
| **3232** | 25,5973490599775 | + | + | + | + | + | NA | NA | + | NA | NA | + | + | 0,0445764110900111 | 10 | -526,980317959798 | 1075,70666766563 | 27,7450947503362 | 3.28554089494733e-08 |
| **1184** | 26,1517153592046 | + | + | + | + | + | NA | NA | + | NA | NA | + | NA | 0,027944940490872 | 9 | -528,161933142076 | 1075,7411891188 | 27,7796162035079 | 3.22931670298726e-08 |
| **3326** | 25,4646256364309 | + | NA | + | + | + | + | + | + | NA | NA | + | + | 0,0603051451944434 | 11 | -525,8437687598 | 1075,7995375196 | 27,8379646043099 | 3.1364649932072e-08 |
| **1504** | 25,8929165415243 | + | + | + | + | + | NA | + | + | + | NA | + | NA | 0,0592113990300195 | 11 | -525,923415203046 | 1075,95883040609 | 27,9972574908011 | 2.89634594676794e-08 |
| **3328** | 25,1556662441331 | + | + | + | + | + | + | + | + | NA | NA | + | + | 0,0751255304964394 | 12 | -524,755320745947 | 1076,02677052415 | 28,0651976088614 | 2.79960928148049e-08 |
| **3486** | 26,0965450741952 | + | NA | + | + | + | NA | NA | + | + | NA | + | + | 0,0419002120377985 | 10 | -527,171835126758 | 1076,08970199955 | 28,1281290842567 | 2.71288902359018e-08 |
| **1502** | 26,2605595975692 | + | NA | + | + | + | NA | + | + | + | NA | + | NA | 0,0412049343567234 | 10 | -527,221503867494 | 1076,18903948102 | 28,2274665657287 | 2.58143484723321e-08 |
| **1438** | 26,6251154507784 | + | NA | + | + | + | NA | NA | + | + | NA | + | NA | 0,0244751180664015 | 9 | -528,405904396264 | 1076,22913162717 | 28,2675587118824 | 2.53020243558904e-08 |
| **3552** | 25,4865012911661 | + | + | + | + | + | NA | + | + | + | NA | + | + | 0,0719470127871254 | 12 | -524,990220717044 | 1076,49657046635 | 28,5349975510546 | 2.21351089228208e-08 |
| **3582** | 25,7718259028655 | + | NA | + | + | + | + | + | + | + | NA | + | + | 0,0705414315801698 | 12 | -525,093840176752 | 1076,70380938576 | 28,7422364704705 | 1.99563128587809e-08 |
| **3550** | 25,8511441032758 | + | NA | + | + | + | NA | + | + | + | NA | + | + | 0,0540553134584687 | 11 | -526,297638363838 | 1076,70727672768 | 28,7457038123853 | 1.99217451519341e-08 |
| **3488** | 25,8703913743319 | + | + | + | + | + | NA | NA | + | + | NA | + | + | 0,0529720049238499 | 11 | -526,376004512619 | 1076,86400902524 | 28,9024361099468 | 1.84201598262435e-08 |
| **3584** | 25,4581290811035 | + | + | + | + | + | + | + | + | + | NA | + | + | 0,0843725739090978 | 13 | -524,067325846217 | 1077,09400128593 | 29,1324283706404 | 1.64191703708392e-08 |
| **1440** | 26,4210761527161 | + | + | + | + | + | NA | NA | + | + | NA | + | NA | 0,0346699481918997 | 10 | -527,686593766015 | 1077,11921927806 | 29,1576463627703 | 1.62134408632017e-08 |
