## Appendix S3 for "Richness and composition of phyllosphere *Methylobacterium* communities cause variation in *Arabidopsis thaliana* growth"

**Table S1** – Parameters and statistical results for all models explored ($n$ = 167) in the context of our third hypothesis, based on the saturated model: dry leaf biomass ~ strain richness + J-067 + E-045 + J-092 + strain richness:J-067 + strain richness:E-045 + strain richness:J-092 + J-067:E-045 + J-067:J-092 + E-045:J-092 + strain richness:J-067:E-045 + strain richness:J-067:J-092 + strain richness:E-045:J-092 + J-067:E-045:J-092 + strain richness:J-067:E-045:J-092. Intercept indicates the mean biomass of one-strain samples that do not contain the strains included in the corresponding model. ID: model number; D2: polynomial strain richness; +: parameters considered in the corresponding model; adj.R2: adjusted $R^2$; df: degrees of freedom; LL: log-likelihood.

(beginning on the next page)

models.H3

| ID | Intercept | J-067 | E-045 | J-092 | D2 | J-067:E-045 | J-067:J-092 | J-067:D2 | E-045:J-092 | E-045:D2 | J-092:D2 | J-067:E-045:J-092 | J-067:E-045:D2 | J-067:J-092:D2 | E-045:J-092:D2 | J-067:E-045:J-092:D2 | adj.R2 | df | LL | AICc | ΔAICc | weight |
|---|---|---|---|---|---|---|---|---|---|---|---|---|---|---|---|---|---|---|---|---|---|---|
| 928 | 32,6262197584058 | + | + | + | + | + | NA | NA | + | + | + | NA | NA | NA | NA | NA | 0,339347093181402 | 13 | -501,722807549167 | 1032,40496469183 | 0 | 0.109861474710245 |
| 784 | 32,262668956707 | + | + | + | + | NA | NA | NA | NA | + | + | NA | NA | NA | NA | NA | 0,313464270276543 | 11 | -504,353537524073 | 1032,81907504815 | 0,414110356314723 | 0.0893146093645038 |
| 800 | 32,2164036494231 | + | + | + | + | + | NA | NA | NA | + | + | NA | NA | NA | NA | NA | 0,324257747581785 | 12 | -503,268742959462 | 1033,05361495118 | 0,648650259352507 | 0.0794315084729773 |
| 9120 | 35,4845151437256 | + | + | + | + | + | NA | NA | + | + | + | NA | NA | NA | + | NA | 0,358194239159225 | 15 | -499,741545394188 | 1033,45003293714 | 1,04506824530654 | 0.0651495977092129 |
| 912 | 32,5304921482878 | + | + | + | + | NA | NA | NA | + | + | + | NA | NA | NA | NA | NA | 0,320073325822968 | 12 | -503,691338210708 | 1033,89880545367 | 1,49384076184401 | 0.052054948908924 |
| 960 | 32,8995248326605 | + | + | + | + | + | + | NA | + | + | + | NA | NA | NA | NA | NA | 0,342807409444323 | 14 | -501,363318463396 | 1034,16925987761 | 1,76429518578107 | 0.045470904436314 |
| 9104 | 35,4187598477672 | + | + | + | + | NA | NA | NA | + | + | + | NA | NA | NA | + | NA | 0,340834547911976 | 14 | -501,568508368209 | 1034,57963968724 | 2,17467499540749 | 0.0370357163451811 |
| 783 | 32,0648499717577 | NA | + | + | + | NA | NA | NA | NA | + | + | NA | NA | NA | NA | NA | 0,291928936447551 | 10 | -506,467919890029 | 1034,68187152609 | 2,27690683425999 | 0.0351901718297468 |
| 528 | 31,5776912993557 | + | + | + | + | NA | NA | NA | NA | NA | + | NA | NA | NA | NA | NA | 0,277902131384645 | 9 | -507,810812526475 | 1035,03894788759 | 2,63398319576436 | 0.0294363013796482 |
| 832 | 32,39420466518 | + | + | + | + | + | + | NA | NA | + | + | NA | NA | NA | NA | NA | 0,326068121640128 | 13 | -503,085097650315 | 1035,12954489413 | 2,72458020229578 | 0.0281326310576875 |
| 816 | 32,2146559202282 | + | + | + | + | NA | + | NA | NA | + | + | NA | NA | NA | NA | NA | 0,31361722596745 | 12 | -504,338284153736 | 1035,19269733973 | 2,78773264789947 | 0.0272581873015536 |
| 911 | 32,3670638741174 | NA | + | + | + | NA | NA | NA | + | + | + | NA | NA | NA | NA | NA | 0,299982369687415 | 11 | -505,684837252699 | 1035,4816745054 | 3,07670981356682 | 0.0235910014441833 |
| 1984 | 33,9156087992359 | + | + | + | + | + | + | NA | + | + | + | + | NA | NA | NA | NA | 0,348186129380012 | 15 | -500,800755599395 | 1035,56845334755 | 3,16348865571877 | 0.0225892905276067 |
| 9152 | 35,480397555742 | + | + | + | + | + | + | NA | + | + | + | NA | NA | NA | + | NA | 0,359905453433302 | 16 | -499,558787659171 | 1035,65090865168 | 3,24594395984514 | 0.0216769237150927 |
| 9103 | 35,4822572693599 | NA | + | + | + | NA | NA | NA | + | + | + | NA | NA | NA | + | NA | 0,322307710194349 | 13 | -503,466006555638 | 1035,89136270477 | 3,48639801294053 | 0.0192213423976863 |
| 320 | 32,5207476231935 | + | + | + | + | + | + | NA | NA | + | NA | NA | NA | NA | NA | NA | 0,297173307121717 | 11 | -505,958996767067 | 1036,02999353413 | 3,62502884230366 | 0.0179341341052857 |
| 944 | 32,4685528896821 | + | + | + | + | NA | + | NA | + | + | + | NA | NA | NA | NA | NA | 0,320351519764753 | 13 | -503,663323465206 | 1036,28599652391 | 3,88103183207704 | 0.0157793849061587 |
| 368 | 31,7816987840682 | + | + | + | + | NA | + | + | NA | + | NA | NA | NA | NA | NA | NA | 0,306687894165439 | 12 | -505,025919569407 | 1036,56796817107 | 4,16300347924061 | 0.0137044214143946 |
| 526 | 30,995203924435 | + | NA | + | + | NA | NA | NA | NA | NA | + | NA | NA | NA | NA | NA | 0,256373349666363 | 8 | -509,822037614038 | 1036,76907522808 | 4,36411053624579 | 0.0123934112829056 |
| 1472 | 34,2259645553768 | + | + | + | + | + | + | NA | + | + | NA | + | NA | NA | NA | NA | 0,317913384768524 | 13 | -503,908460116641 | 1036,77626982678 | 4,37130513494731 | 0.0123489085653333 |
| 992 | 32,5498106441971 | + | + | + | + | + | NA | + | + | + | + | NA | NA | NA | NA | NA | 0,342251486428645 | 15 | -501,421200147814 | 1036,80934244439 | 4,40437775255759 | 0.0121463823330864 |
| 110 | 30,4429903231817 | + | NA | + | + | NA | + | + | NA | NA | NA | NA | NA | NA | NA | NA | 0,268350165664038 | 9 | -508,7104555329 | 1036,83823390045 | 4,43326920861455 | 0.0119721802645744 |
| 384 | 32,2134066722498 | + | + | + | + | + | + | + | NA | + | NA | NA | NA | NA | NA | NA | 0,317197980703383 | 13 | -503,980222612684 | 1036,91979481886 | 4,51483012703352 | 0.0114937704414043 |
| 9136 | 35,4284718069735 | + | + | + | + | NA | + | NA | + | + | + | NA | NA | NA | + | NA | 0,341684285975778 | 15 | -501,480205653773 | 1036,92735345631 | 4,52238876447609 | 0.0114504138004854 |
| 656 | 31,4265182403441 | + | + | + | + | NA | NA | NA | + | NA | + | NA | NA | NA | NA | NA | 0,28003405636756 | 10 | -507,608396031761 | 1036,96282380955 | 4,55785911772409 | 0.0112491288770769 |
| 112 | 30,5320363386445 | + | + | + | + | NA | + | + | NA | NA | NA | NA | NA | NA | NA | NA | 0,279149029567912 | 10 | -507,692497890599 | 1037,13102752723 | 4,72606283539835 | 0.0103417472414281 |
| 848 | 31,975738380491 | + | + | + | + | NA | NA | + | NA | + | + | NA | NA | NA | NA | NA | 0,315507670004406 | 13 | -504,149480353598 | 1037,25831030069 | 4,85334560886213 | 0.00970409000785917 |
| 560 | 31,539988448569 | + | + | + | + | NA | + | NA | NA | NA | + | NA | NA | NA | NA | NA | 0,27826699018339 | 10 | -507,776213253579 | 1037,29845825319 | 4,89349356136017 | 0.0095112325200218 |
| 544 | 31,5802276169344 | + | + | + | + | + | NA | NA | NA | NA | + | NA | NA | NA | NA | NA | 0,277902872244974 | 10 | -507,810742288976 | 1037,36751632398 | 4,96255163215392 | 0.00918842404022818 |
| 10176 | 35,4726931389677 | + | + | + | + | + | + | NA | + | + | + | + | NA | NA | + | NA | 0,361602728285704 | 17 | -499,377035717825 | 1037,89692857851 | 5,49196388667656 | 0.00705148552947106 |
| 864 | 32,3124981098845 | + | + | + | + | + | NA | + | NA | + | + | NA | NA | NA | NA | NA | 0,324412932343054 | 14 | -503,253020213285 | 1037,94866337739 | 5,54369868555864 | 0.00687142087907428 |
| 9184 | 35,4972937076271 | + | + | + | + | + | NA | + | + | + | + | NA | NA | NA | + | NA | 0,361332579059507 | 17 | -499,405996820002 | 1037,95485078286 | 5,54988609102952 | 0.00685019559479327 |
| 448 | 32,7563914780649 | + | + | + | + | + | + | NA | + | + | NA | NA | NA | NA | NA | NA | 0,298463418292182 | 12 | -505,833220443912 | 1038,18256992008 | 5,77760522825065 | 0.00611299991560235 |
| 976 | 31,9831092085733 | + | + | + | + | NA | NA | + | + | + | + | NA | NA | NA | NA | NA | 0,322032445045536 | 14 | -503,493806371924 | 1038,43023569467 | 6,02527100283714 | 0.00540100339364558 |
| 240 | 30,4434211670764 | + | + | + | + | NA | + | + | + | NA | NA | NA | NA | NA | NA | NA | 0,283403300878898 | 11 | -507,287277111237 | 1038,68655422247 | 6,28158953064417 | 0.00475133431035962 |
| 512 | 32,4150036037376 | + | + | + | + | + | + | + | + | + | NA | NA | NA | NA | NA | NA | 0,320455049843283 | 14 | -503,652894831199 | 1038,74841261322 | 6,34344792138609 | 0.00460662871250187 |
| 558 | 30,9430202665662 | + | NA | + | + | NA | + | NA | NA | NA | + | NA | NA | NA | NA | NA | 0,257445159721225 | 9 | -509,72329332543 | 1038,86390948551 | 6,45894479367462 | 0.00434813863936154 |
| 592 | 31,1750486941863 | + | + | + | + | NA | NA | + | NA | NA | + | NA | NA | NA | NA | NA | 0,28245709529156 | 11 | -507,377611026963 | 1038,86722205393 | 6,46225736209567 | 0.00434094284677182 |
| 1536 | 33,5466128746353 | + | + | + | + | + | + | + | + | + | NA | + | NA | NA | NA | NA | 0,332220762638628 | 15 | -502,457264678734 | 1038,88147150623 | 6,47650681439814 | 0.0043101247334244 |
| 496 | 31,8073654807658 | + | + | + | + | NA | + | + | + | + | NA | NA | NA | NA | NA | NA | 0,306911286863213 | 13 | -505,003858438021 | 1038,96706646954 | 6,56210177770731 | 0.00412955380048144 |
| 1024 | 32,5793731252786 | + | + | + | + | + | + | + | + | + | + | NA | NA | NA | NA | NA | 0,343217436305459 | 16 | -501,320595857017 | 1039,17452504737 | 6,76956035553644 | 0.00372266598261008 |
| 688 | 31,3954406310124 | + | + | + | + | NA | + | NA | + | NA | + | NA | NA | NA | NA | NA | 0,280323554337074 | 11 | -507,580863344327 | 1039,27372668865 | 6,86876199682274 | 0.00354252324019258 |
| 672 | 31,4224702438704 | + | + | + | + | + | NA | NA | + | NA | + | NA | NA | NA | NA | NA | 0,28003575762204 | 11 | -507,608234266375 | 1039,32846853275 | 6,92350384091856 | 0.00344687605973033 |
| 9168 | 35,079288876567 | + | + | + | + | NA | NA | + | + | + | + | NA | NA | NA | + | NA | 0,342142115924708 | 16 | -501,432581848617 | 1039,39849703057 | 6,9935323387374 | 0.00332827476782058 |
| 880 | 32,0102991994539 | + | + | + | + | NA | + | + | NA | + | + | NA | NA | NA | NA | NA | 0,317165333743096 | 14 | -503,983495650612 | 1039,40961425204 | 7,00464956021369 | 0.00330982550758866 |
| 144 | 30,2752526782313 | + | + | + | + | NA | NA | NA | + | NA | NA | NA | NA | NA | NA | NA | 0,241738183798851 | 8 | -511,156303219831 | 1039,43760643966 | 7,03264174783089 | 0.00326382355406635 |
| 128 | 30,5197975925903 | + | + | + | + | + | + | + | NA | NA | NA | NA | NA | NA | NA | NA | 0,279337539563349 | 11 | -507,674592914913 | 1039,46118582983 | 7,0562211379945 | 0.00322557001145735 |
| 590 | 30,6075514354938 | + | NA | + | + | NA | NA | + | NA | NA | + | NA | NA | NA | NA | NA | 0,266587178322165 | 10 | -508,875216930375 | 1039,49646560678 | 7,09150091495098 | 0.00316917022164333 |
| 576 | 31,5614666499993 | + | + | + | + | + | + | NA | NA | NA | + | NA | NA | NA | NA | NA | 0,278349742526219 | 11 | -507,768363482069 | 1039,64872696414 | 7,24376227230823 | 0.00293685451342 |
| 527 | 31,1269548228082 | NA | + | + | + | NA | NA | NA | NA | NA | + | NA | NA | NA | NA | NA | 0,239437709682972 | 8 | -511,363690255369 | 1039,85238051074 | 7,44741581890639 | 0.00265252586307922 |
| 896 | 32,3723973606295 | + | + | + | + | + | + | + | NA | + | + | NA | NA | NA | NA | NA | 0,32691487381606 | 15 | -502,999033371695 | 1039,96500889215 | 7,56004420032059 | 0.0025072791249347 |
| 304 | 32,1895119861138 | + | + | + | + | NA | + | NA | NA | + | NA | NA | NA | NA | NA | NA | 0,263501323780431 | 10 | -509,162657466848 | 1040,07134667973 | 7,66638198789633 | 0.00237745183704549 |
| 176 | 30,6852469175899 | + | + | + | + | NA | + | NA | + | NA | NA | NA | NA | NA | NA | NA | 0,250594497228643 | 9 | -510,351995928353 | 1040,12131469135 | 7,71634999952084 | 0.00231878942889731 |
| 3040 | 35,3827907269311 | + | + | + | + | + | NA | + | + | + | + | NA | + | NA | NA | NA | 0,350125332803591 | 17 | -500,596794914127 | 1040,33644697111 | 7,93148227928032 | 0.00208231261813902 |
| 9216 | 35,4975754515051 | + | + | + | + | + | + | + | + | + | + | NA | NA | NA | + | NA | 0,361332584588586 | 18 | -499,405996227387 | 1040,60860262427 | 8,20363793243473 | 0.00181738978055078 |
| 2048 | 33,5648174452381 | + | + | + | + | + | + | + | + | + | + | + | NA | NA | NA | NA | 0,348607052121639 | 17 | -500,756535546927 | 1040,65592823671 | 8,25096354487982 | 0.00177489005280242 |
| 624 | 31,0548206409431 | + | + | + | + | NA | + | + | NA | NA | + | NA | NA | NA | NA | NA | 0,285617367466946 | 12 | -507,075434278758 | 1040,66699758977 | 8,26203289794421 | 0.00176509374518411 |
| 622 | 30,5034831013052 | + | NA | + | + | NA | + | + | NA | NA | + | NA | NA | NA | NA | NA | 0,272404259413078 | 11 | -508,330066707003 | 1040,77213341401 | 8,36716872217607 | 0.00167470309069406 |
| 48 | 31,4525818884348 | + | + | + | + | NA | + | NA | NA | NA | NA | NA | NA | NA | NA | NA | 0,234220044149725 | 8 | -511,831747132264 | 1040,78849426453 | 8,38352957269808 | 0.00166105918976883 |
| 1008 | 32,0062115789868 | + | + | + | + | NA | + | + | + | + | + | NA | NA | NA | NA | NA | 0,322778684199408 | 15 | -503,418415359489 | 1040,80377286774 | 8,39880817590756 | 0.00164841820326525 |
| 720 | 31,1964034613792 | + | + | + | + | NA | NA | + | + | NA | + | NA | NA | NA | NA | NA | 0,284364983926254 | 12 | -507,195343338764 | 1040,90681570979 | 8,50185101795637 | 0.00156564008815172 |
| 256 | 30,4343688278778 | + | + | + | + | + | + | + | + | NA | NA | NA | NA | NA | NA | NA | 0,283521541043987 | 12 | -507,275980386185 | 1041,06808980463 | 8,66312511279671 | 0.00144434754691087 |
| 608 | 31,1634825796366 | + | + | + | + | + | NA | + | NA | NA | + | NA | NA | NA | NA | NA | 0,283421746108625 | 12 | -507,285514967319 | 1041,0871589669 | 8,68219427506529 | 0.00143064174142667 |
| 288 | 31,3651820181602 | + | + | + | + | + | NA | NA | NA | + | NA | NA | NA | NA | NA | NA | 0,256989733488528 | 10 | -509,765268469145 | 1041,27656868432 | 8,87160399249183 | 0.00130137091018427 |
| 525 | 30,5571694051641 | NA | NA | + | + | NA | NA | NA | NA | NA | + | NA | NA | NA | NA | NA | 0,218563212024955 | 7 | -513,217365143422 | 1041,30294734111 | 8,89798264927663 | 0.00128431939807302 |
| 2432 | 33,4602691922087 | + | + | + | + | + | + | + | NA | + | NA | NA | + | NA | NA | NA | 0,320212887045812 | 15 | -503,677285518907 | 1041,32151318657 | 8,91654849474412 | 0.00127245232610928 |
| 655 | 30,9317374542623 | NA | + | + | + | NA | NA | NA | + | NA | + | NA | NA | NA | NA | NA | 0,243538386163588 | 9 | -510,993576482489 | 1041,40447579962 | 8,9995111077933 | 0.00122074911313689 |
| 16 | 31,0667692034895 | + | + | + | + | NA | NA | NA | NA | NA | NA | NA | NA | NA | NA | NA | 0,217562371315243 | 7 | -513,304992906306 | 1041,47820286688 | 9,07323817504562 | 0.00117656733850166 |
| 160 | 30,2039511132295 | + | + | + | + | + | NA | NA | + | NA | NA | NA | NA | NA | NA | NA | 0,242393123499465 | 9 | -511,097145602035 | 1041,61161403872 | 9,20664934688511 | 0.00110064412644746 |
| 272 | 31,3926160436827 | + | + | + | + | NA | NA | NA | NA | + | NA | NA | NA | NA | NA | NA | 0,242349993017976 | 9 | -511,10104294708 | 1041,6194087288 | 9,2144440369741 | 0.0010963628846728 |
| 704 | 31,4107720090008 | + | + | + | + | + | + | NA | + | NA | + | NA | NA | NA | NA | NA | 0,280358533903952 | 12 | -507,577535865193 | 1041,67120076264 | 9,26623607081342 | 0.00106833591256492 |
| 432 | 31,9270595343823 | + | + | + | + | NA | + | NA | + | + | NA | NA | NA | NA | NA | NA | 0,266263223153888 | 11 | -508,905449366924 | 1041,92289873385 | 9,51793404201749 | 0.000942003047151379 |
| 192 | 30,5964417418858 | + | + | + | + | + | + | NA | + | NA | NA | NA | NA | NA | NA | NA | 0,253257245969347 | 10 | -510,108313820751 | 1041,96265938753 | 9,5576946957035 | 0.000923460643906833 |
| 9200 | 35,0944758093087 | + | + | + | + | NA | + | + | + | + | + | NA | NA | NA | + | NA | 0,342157504728029 | 17 | -501,430980518775 | 1042,00481818041 | 9,5998534885764 | 0.000904198382567662 |
| 208 | 30,1854485239026 | + | + | + | + | NA | NA | + | + | NA | NA | NA | NA | NA | NA | NA | 0,252061699649725 | 10 | -510,217831781513 | 1042,18169530906 | 9,77673061722749 | 0.000827666438970626 |
| 11232 | 36,062193683134 | + | + | + | + | + | NA | + | + | + | + | NA | + | NA | + | NA | 0,366389754541736 | 19 | -498,861805589171 | 1042,21933767407 | 9,81437298223727 | 0.000812234457484992 |
| 400 | 31,103111085616 | + | + | + | + | NA | NA | NA | + | + | NA | NA | NA | NA | NA | NA | 0,251412955533236 | 10 | -510,27718674934 | 1042,30040524471 | 9,8954405528807 | 0.000779969841277627 |
| 2912 | 34,5289137623521 | + | + | + | + | + | NA | + | NA | + | + | NA | + | NA | NA | NA | 0,327952608740794 | 16 | -502,893409839501 | 1042,32015301233 | 9,91518832050406 | 0.000772306405884784 |
| 64 | 31,3165773098426 | + | + | + | + | + | + | NA | NA | NA | NA | NA | NA | NA | NA | NA | 0,238038622265143 | 9 | -511,489511196155 | 1042,39634522695 | 9,99138053512434 | 0.000743437918026361 |
| 5120 | 37,4011311431876 | + | + | + | + | + | + | + | + | + | + | NA | NA | + | NA | NA | 0,352789843793774 | 18 | -500,31555395549 | 1042,42771808047 | 10,0227533886405 | 0.000731867023832277 |
| 752 | 31,0739385898564 | + | + | + | + | NA | + | + | + | NA | + | NA | NA | NA | NA | NA | 0,287689465673585 | 13 | -506,876579800533 | 1042,71250919456 | 10,3075445027309 | 0.00063473226431562 |
| 3072 | 35,1334198399039 | + | + | + | + | + | + | + | + | + | + | NA | + | NA | NA | NA | 0,350897381632078 | 18 | -500,515423379079 | 1042,82745692765 | 10,4224922358192 | 0.000599280286335204 |
| 416 | 31,1729681668034 | + | + | + | + | + | NA | NA | + | + | NA | NA | NA | NA | NA | NA | 0,260792042615427 | 11 | -509,414030485267 | 1042,94006097053 | 10,5350962787033 | 0.000566471851613489 |
| 10240 | 35,7075153260996 | + | + | + | + | + | + | + | + | + | + | + | NA | NA | + | NA | 0,363031155615699 | 19 | -499,22369859403 | 1042,94312368379 | 10,5381589919568 | 0.000565605045054035 |
| 2560 | 33,6239109406041 | + | + | + | + | + | + | + | + | + | NA | NA | + | NA | NA | NA | 0,324683737521162 | 16 | -503,225574587443 | 1042,98448250822 | 10,5795178163878 | 0.000554028772962421 |
| 640 | 31,052648545826 | + | + | + | + | + | + | + | NA | NA | + | NA | NA | NA | NA | NA | 0,286186261583201 | 13 | -507,020896287734 | 1043,00114216896 | 10,5961774771336 | 0.000549432974944579 |
| 736 | 31,184984083391 | + | + | + | + | + | NA | + | + | NA | + | NA | NA | NA | NA | NA | 0,285132586325403 | 13 | -507,121874396295 | 1043,20309838609 | 10,7981336942557 | 0.000496661487924768 |
| 80 | 30,3093271125432 | + | + | + | + | NA | NA | + | NA | NA | NA | NA | NA | NA | NA | NA | 0,232569665705856 | 9 | -511,979132959861 | 1043,37558875437 | 10,9706240625376 | 0.000455621991206448 |
| 143 | 30,2601756642449 | NA | + | + | + | NA | NA | NA | + | NA | NA | NA | NA | NA | NA | NA | 0,206611519837353 | 7 | -514,256532783558 | 1043,38128262138 | 10,9763179295496 | 0.000454326710357684 |
| 46 | 31,443248752285 | + | NA | + | + | NA | + | NA | NA | NA | NA | NA | NA | NA | NA | NA | 0,206387901484669 | 7 | -514,275826347782 | 1043,41986974983 | 11,0149050579969 | 0.000445645147242241 |
| 1280 | 30,4381242449544 | + | + | + | + | + | + | + | + | NA | NA | + | NA | NA | NA | NA | 0,283531269103751 | 13 | -507,275050879496 | 1043,50945135249 | 11,1044866606569 | 0.000426124773966858 |
| 32 | 30,9860687645861 | + | + | + | + | + | NA | NA | NA | NA | NA | NA | NA | NA | NA | NA | 0,218353621061948 | 8 | -513,23572498482 | 1043,59644996964 | 11,1914852778086 | 0.000407986012791805 |
| 14 | 31,0840648943539 | + | NA | + | + | NA | NA | NA | NA | NA | NA | NA | NA | NA | NA | NA | 0,191964114460232 | 6 | -515,508954581656 | 1043,66406300947 | 11,2590983176358 | 0.000394423960055044 |
| 78 | 30,193171000276 | + | NA | + | + | NA | NA | + | NA | NA | NA | NA | NA | NA | NA | NA | 0,217540267027239 | 8 | -513,306926963926 | 1043,73885392785 | 11,3338892360207 | 0.00037994667439642 |
| 3584 | 33,4569545616102 | + | + | + | + | + | + | + | + | + | NA | + | + | NA | NA | NA | 0,33305996419007 | 17 | -502,371182460718 | 1043,88522206429 | 11,4802573724633 | 0.000353133737202922 |
| 1728 | 31,3820983363244 | + | + | + | + | + | + | NA | + | NA | + | + | NA | NA | NA | NA | 0,280584502394744 | 13 | -507,556036407834 | 1044,07142240916 | 11,6664577173335 | 0.000321740934887349 |
| 336 | 30,8626532717054 | + | + | + | + | NA | NA | + | NA | + | NA | NA | NA | NA | NA | NA | 0,254144416543433 | 11 | -510,02693127859 | 1044,16586255718 | 11,7608978653498 | 0.00030690142280619 |
| 4976 | 33,2098680099546 | + | + | + | + | NA | + | + | NA | + | + | NA | NA | + | NA | NA | 0,318709662271606 | 16 | -503,828496661793 | 1044,19032665692 | 11,785361965088 | 0.000303170255730658 |
| 1216 | 30,5202378636059 | + | + | + | + | + | + | NA | + | NA | NA | + | NA | NA | NA | NA | 0,253992822863524 | 11 | -510,040844225388 | 1044,19368845077 | 11,7887237589443 | 0.000302661085830031 |
| 271 | 31,5120622543132 | NA | + | + | + | NA | NA | NA | NA | + | NA | NA | NA | NA | NA | NA | 0,214921130905754 | 8 | -513,535707657819 | 1044,19641531564 | 11,791450623808 | 0.000302248709077554 |
| 224 | 30,172037828769 | + | + | + | + | + | NA | + | + | NA | NA | NA | NA | NA | NA | NA | 0,252357412758479 | 11 | -510,190759291184 | 1044,49351858237 | 12,0885538905384 | 0.000260524937265617 |
| 2944 | 34,1511363758275 | + | + | + | + | + | + | + | NA | + | + | NA | + | NA | NA | NA | 0,330080174470636 | 17 | -502,676349729389 | 1044,49555660163 | 12,0905919098043 | 0.00026025959506101 |
| 6144 | 36,9050828747169 | + | + | + | + | + | + | + | + | + | + | + | NA | + | NA | NA | 0,355317256482314 | 19 | -500,047712465128 | 1044,59115142598 | 12,1861867341511 | 0.000248112472939254 |
| 4992 | 34,4737508547825 | + | + | + | + | + | + | + | NA | + | + | NA | NA | + | NA | NA | 0,329112576071725 | 17 | -502,775152062225 | 1044,69316126731 | 12,2881965754757 | 0.000235774830756049 |
| 399 | 31,2164888208172 | NA | + | + | + | NA | NA | NA | + | + | NA | NA | NA | NA | NA | NA | 0,22432159709534 | 9 | -512,711005054731 | 1044,83933294411 | 12,4343682522776 | 0.000219157665419029 |
| 11264 | 35,959308162504 | + | + | + | + | + | + | + | + | + | + | NA | + | NA | + | NA | 0,366504830104603 | 20 | -498,849372121104 | 1044,94012355255 | 12,5351588607225 | 0.000208386827474689 |
| 4720 | 33,6898868358137 | + | + | + | + | NA | + | + | NA | NA | + | NA | NA | + | NA | NA | 0,288958966562543 | 14 | -506,754462938797 | 1044,95154882841 | 12,5465841365829 | 0.000207199782786353 |

| | | | | | | | | | | | | | | | | | | | | | | |
|---|---|---|---|---|---|---|---|---|---|---|---|---|---|---|---|---|---|---|---|---|---|---|
| **768** | 31,0713908632359 | + | + | + | + | + | + | + | + | NA | + | NA | NA | NA | NA | NA | 0,288093822955644 | 14 | -506,837707158499 | 1045,11803726782 | 12,7130725759866 | 0.000190649991621104 |
| **464** | 30,8756276443305 | + | + | + | + | NA | NA | + | + | + | NA | NA | NA | NA | NA | NA | 0,261220775162414 | 12 | -509,374313130286 | 1045,26475529283 | 12,859790601 | 0.000177164774666793 |
| **4718** | 32,2893723478372 | + | NA | + | + | NA | + | + | NA | NA | + | NA | NA | + | NA | NA | 0,273845569002392 | 13 | -508,194320441165 | 1045,34799047583 | 12,9430257839963 | 0.000169942923804488 |
| **4096** | 34,9891201215451 | + | + | + | + | + | + | + | + | + | + | + | + | NA | NA | NA | 0,351693818896769 | 19 | -500,431379965876 | 1045,35848642748 | 12,9535217356479 | 0.000169053403580454 |
| **352** | 31,1393310865095 | + | + | + | + | + | NA | + | NA | + | NA | NA | NA | NA | NA | NA | 0,259716580436692 | 12 | -509,513559008532 | 1045,54324704932 | 13,1382823574922 | 0.000154135849258643 |
| **96** | 30,2879249357144 | + | + | + | + | + | NA | + | NA | NA | NA | NA | NA | NA | NA | NA | 0,233230317799553 | 10 | -511,920171971052 | 1045,58637568814 | 13,1814109963052 | 0.000150847596464268 |
| **5104** | 33,9412768621828 | + | + | + | + | NA | + | + | + | + | + | NA | NA | + | NA | NA | 0,324553815535171 | 17 | -503,238743323258 | 1045,62034378937 | 13,2153790975431 | 0.000148307227195819 |
| **13312** | 36,8192691332095 | + | + | + | + | + | + | + | + | + | + | NA | NA | + | + | NA | 0,363077216422669 | 20 | -499,218748396026 | 1045,6788761024 | 13,2739114105652 | 0.000144029742741653 |
| **7168** | 38,571353492113 | + | + | + | + | + | + | + | + | + | + | NA | + | + | NA | NA | 0,362383503963946 | 20 | -499,293264471731 | 1045,82790825381 | 13,4229435619754 | 0.000133687334564153 |
| **136** | 23,5581781507762 | + | + | + | NA | NA | NA | NA | + | NA | NA | NA | NA | NA | NA | NA | 0,176035108803875 | 6 | -516,84546010944 | 1046,33707406503 | 13,9321093732021 | 0.000103639739662121 |
| **15** | 31,177573415374 | NA | + | + | + | NA | NA | NA | NA | NA | NA | NA | NA | NA | NA | NA | 0,174479806022408 | 6 | -516,974569496292 | 1046,59529283874 | 14,1903281469069 | 9.10866687299697e-05 |
| **152** | 24,0413659481715 | + | + | + | NA | + | NA | NA | + | NA | NA | NA | NA | NA | NA | NA | 0,184854686809788 | 7 | -516,108687416684 | 1047,08559188763 | 14,6806271958001 | 7.1283289596955e-05 |
| **4848** | 33,3191307176173 | + | + | + | + | NA | + | + | + | NA | + | NA | NA | + | NA | NA | 0,290776101870955 | 15 | -506,579287886025 | 1047,12551792081 | 14,7205532289788 | 6.98743700366104e-05 |
| **13296** | 33,7824167151982 | + | + | + | + | NA | + | + | + | + | + | NA | NA | + | + | NA | 0,343244301322474 | 19 | -501,317795735578 | 1047,13131796688 | 14,7263532750515 | 6.96720262964214e-05 |
| **480** | 31,078069872184 | + | + | + | + | + | NA | + | + | + | NA | NA | NA | NA | NA | NA | 0,264030873317796 | 13 | -509,113416750207 | 1047,18618309391 | 14,7812184020795 | 6.77867216101553e-05 |
| **4736** | 33,5189967467393 | + | + | + | + | + | + | + | NA | NA | + | NA | NA | + | NA | NA | 0,289171454984431 | 15 | -506,734001793963 | 1047,43494573669 | 15,0299810448562 | 5.98585936354605e-05 |
| **1792** | 31,0266047598804 | + | + | + | + | + | + | + | + | NA | + | + | NA | NA | NA | NA | 0,288264299711313 | 15 | -506,821311866038 | 1047,60956588084 | 15,2046011890052 | 5.48539904345977e-05 |
| **14336** | 36,230927089994 | + | + | + | + | + | + | + | + | + | + | + | NA | + | + | NA | 0,366572440313025 | 21 | -498,842066047796 | 1047,71891470429 | 15,3139500124566 | 5.19353940551755e-05 |
| **12288** | 36,0480446253318 | + | + | + | + | + | + | + | + | + | + | + | + | NA | + | NA | 0,366533592599677 | 21 | -498,846264092548 | 1047,72731079379 | 15,3223461019616 | 5.17178239507433e-05 |
| **8** | 24,2851289472049 | + | + | + | NA | NA | NA | NA | NA | NA | NA | NA | NA | NA | NA | NA | 0,152770250944867 | 5 | -518,751778824058 | 1047,96157291529 | 15,55660822346 | 4.60013810386511e-05 |
| **168** | 23,5950991697417 | + | + | + | NA | NA | + | NA | + | NA | NA | NA | NA | NA | NA | NA | 0,176093482648054 | 7 | -516,840609616398 | 1048,54943628706 | 16,1444715952298 | 3.42860900059291e-05 |
| **24** | 24,7763078276055 | + | + | + | NA | + | NA | NA | NA | NA | NA | NA | NA | NA | NA | NA | 0,162349077122578 | 6 | -517,973305435372 | 1048,5927647169 | 16,1878000250681 | 3.35512968800524e-05 |
| **8192** | 38,6162759773794 | + | + | + | + | + | + | + | + | + | + | + | + | + | NA | NA | 0,362387537257581 | 21 | -499,292831464152 | 1048,620445537 | 16,2154808451689 | 3.3090131891371e-05 |
| **13** | 31,1968771090695 | NA | NA | + | + | NA | NA | NA | NA | NA | NA | NA | NA | NA | NA | NA | 0,147648317318856 | 5 | -519,164437687107 | 1048,78689064139 | 16,3819259495588 | 3.04477638771206e-05 |
| **184** | 23,9639570008149 | + | + | + | NA | + | + | NA | + | NA | NA | NA | NA | NA | NA | NA | 0,185373891821718 | 8 | -516,065065920986 | 1049,25513184197 | 16,8501671501406 | 2.40922919991231e-05 |
| **15360** | 37,8412328990334 | + | + | + | + | + | + | + | + | + | + | NA | + | + | + | NA | 0,371413192076156 | 22 | -498,316930573558 | 1049,51105412957 | 17,106089437741 | 2.11985082042348e-05 |
| **2400** | 34,2396430915765 | + | + | + | + | + | NA | + | NA | + | NA | NA | + | NA | NA | NA | 0,264543740018286 | 14 | -509,065693539599 | 1049,57401003002 | 17,169045338188 | 2.05416156732557e-05 |
| **7040** | 34,9870338310248 | + | + | + | + | + | + | + | NA | + | + | NA | + | + | NA | NA | 0,331077979282979 | 19 | -502,57431352049 | 1049,64435353671 | 17,2393888448753 | 1.98316889013714e-05 |
| **4864** | 33,1512090850878 | + | + | + | + | + | + | + | + | NA | + | NA | NA | + | NA | NA | 0,290982723670793 | 16 | -506,559340784848 | 1049,65201490303 | 17,2470502111983 | 1.97558653054397e-05 |
| **40** | 24,1070282143905 | + | + | + | NA | NA | + | NA | NA | NA | NA | NA | NA | NA | NA | NA | 0,153693208590343 | 6 | -518,677153824121 | 1050,00046149439 | 17,5954968025642 | 1.65970858552934e-05 |
| **56** | 24,4917316692668 | + | + | + | NA | + | + | NA | NA | NA | NA | NA | NA | NA | NA | NA | 0,166203363295307 | 7 | -517,657553429084 | 1050,18332391243 | 17,7783592206015 | 1.51469004467024e-05 |
| **2528** | 34,0078470227678 | + | + | + | + | + | NA | + | + | + | NA | NA | + | NA | NA | NA | 0,268304630951042 | 15 | -508,714716005476 | 1051,39637415971 | 18,9914094678813 | 8.25872978297775e-06 |
| **1208** | 23,9615384615385 | + | + | + | NA | + | + | NA | + | NA | NA | + | NA | NA | NA | NA | 0,18537445871751 | 9 | -516,065018277519 | 1051,54735938968 | 19,1423946978539 | 7.65820928348456e-06 |
| **135** | 24,9304347826087 | NA | + | + | NA | NA | NA | NA | + | NA | NA | NA | NA | NA | NA | NA | 0,127593519035655 | 5 | -520,756685963656 | 1051,97138719449 | 19,5664225026574 | 6.19513482906315e-06 |
| **6** | 23,469998397228 | + | NA | + | NA | NA | NA | NA | NA | NA | NA | NA | NA | NA | NA | NA | 0,11299343679854 | 4 | -521,893011899873 | 1052,08905410278 | 19,6840894109455 | 5.84116828056443e-06 |
| **5888** | 33,4098727336551 | + | + | + | + | + | + | + | + | NA | + | + | NA | + | NA | NA | 0,291824339730989 | 17 | -506,478031783728 | 1052,09892071031 | 19,6939560184817 | 5.81242298600979e-06 |
| **16384** | 37,7789308145617 | + | + | + | + | + | + | + | + | + | + | + | + | + | + | NA | 0,371416431675097 | 23 | -498,316577783229 | 1052,40306707088 | 19,9981023790513 | 4.99243786629943e-06 |
| **267** | 32,5068323935604 | NA | + | NA | + | NA | NA | NA | NA | + | NA | NA | NA | NA | NA | NA | 0,151294510915347 | 7 | -518,870929941549 | 1052,61007693736 | 20,2051122455312 | 4.50153931184237e-06 |
| **284** | 32,4715060580938 | + | + | NA | + | + | NA | NA | NA | + | NA | NA | NA | NA | NA | NA | 0,17830714304436 | 9 | -516,656414659158 | 1052,73015215296 | 20,3251874611312 | 4.23923063929228e-06 |
| **268** | 32,548977515279 | + | + | NA | + | NA | NA | NA | NA | + | NA | NA | NA | NA | NA | NA | 0,157693825761524 | 8 | -518,352743896458 | 1053,83048779292 | 21,4255231010848 | 2.44541288992679e-06 |
| **7** | 25,7709296077378 | NA | + | + | NA | NA | NA | NA | NA | NA | NA | NA | NA | NA | NA | NA | 0,100513293995182 | 4 | -522,84961977992 | 1054,00226986287 | 21,5973051710387 | 2.24414128033227e-06 |
| **38** | 23,3180327868853 | + | NA | + | NA | NA | + | NA | NA | NA | NA | NA | NA | NA | NA | NA | 0,113643441483382 | 5 | -521,842820883433 | 1054,14365703404 | 21,7386923422118 | 2.09097268919949e-06 |
| **11** | 31,1482986487476 | NA | + | NA | + | NA | NA | NA | NA | NA | NA | NA | NA | NA | NA | NA | 0,112770477986076 | 5 | -521,910219507689 | 1054,27845428255 | 21,8734895907221 | 1.95468828358267e-06 |
| **12** | 31,0840392254571 | + | + | NA | + | NA | NA | NA | NA | NA | NA | NA | NA | NA | NA | NA | 0,126740467645958 | 6 | -520,823600095075 | 1054,2933540363 | 21,8883893444736 | 1.94018020533845e-06 |
| **5** | 24,9033333333334 | NA | NA | + | NA | NA | NA | NA | NA | NA | NA | NA | NA | NA | NA | NA | 0,0796930196936916 | 3 | -524,41635579295 | 1055,01316271372 | 22,6081980218894 | 1.35374729259751e-06 |
| **10** | 31,0989380126039 | + | NA | NA | + | NA | NA | NA | NA | NA | NA | NA | NA | NA | NA | NA | 0,106545923220789 | 5 | -522,388884135645 | 1055,23578353847 | 22,8308188466358 | 1.21114489568544e-06 |
| **32768** | 38,2245714285711 | + | + | + | + | + | + | + | + | + | + | + | + | + | + | + | 0,371492771520516 | 24 | -498,308263895169 | 1055,33081350462 | 22,9258488127934 | 1.15494315227283e-06 |
| **9** | 31,1668567639434 | NA | NA | NA | + | NA | NA | NA | NA | NA | NA | NA | NA | NA | NA | NA | 0,0911977810082743 | 4 | -523,555050014779 | 1055,41313033259 | 23,0081656407572 | 1.10837248537837e-06 |
| **28** | 30,9696914662184 | + | + | NA | + | + | NA | NA | NA | NA | NA | NA | NA | NA | NA | NA | 0,128329018187633 | 7 | -520,698940216255 | 1056,26609748677 | 23,8611327949432 | 7.23546208372309e-07 |
| **74** | 29,7547731437867 | + | NA | NA | + | NA | NA | + | NA | NA | NA | NA | NA | NA | NA | NA | 0,125458184909957 | 7 | -520,92406071187 | 1056,716338478 | 24,3113737861727 | 5.77693768647352e-07 |
| **332** | 31,4153029000218 | + | + | NA | + | NA | NA | + | NA | + | NA | NA | NA | NA | NA | NA | 0,167479953814838 | 10 | -517,552650501471 | 1056,85133274897 | 24,4463680571434 | 5.39987926382915e-07 |
| **76** | 29,8536010967209 | + | + | NA | + | NA | NA | + | NA | NA | NA | NA | NA | NA | NA | NA | 0,137750172663482 | 8 | -519,954925647269 | 1057,03485129454 | 24,6298866027078 | 4.92644346975942e-07 |
| **348** | 31,7933983426173 | + | + | NA | + | + | NA | + | NA | + | NA | NA | NA | NA | NA | NA | 0,179461165884903 | 11 | -516,560193592164 | 1057,23238718433 | 24,8274224924965 | 4.46312582019915e-07 |
| **92** | 29,8497735497526 | + | + | NA | + | + | NA | + | NA | NA | NA | NA | NA | NA | NA | NA | 0,137766960373611 | 9 | -519,953592636409 | 1059,32450810746 | 26,9195434156331 | 1.56798117407319e-07 |
| **2** | 25,523595505618 | + | NA | NA | NA | NA | NA | NA | NA | NA | NA | NA | NA | NA | NA | NA | 0,0419917217098653 | 3 | -527,165295242648 | 1060,51104161312 | 28,1060769212847 | 8.66340000407547e-08 |
| **2396** | 35,5176009128511 | + | + | NA | + | + | NA | + | NA | + | NA | NA | + | NA | NA | NA | 0,188119302769368 | 13 | -515,833944721722 | 1060,62723903694 | 28,2222743451086 | 8.17441000700169e-08 |
| **4** | 26,1945672410541 | + | + | NA | NA | NA | NA | NA | NA | NA | NA | NA | NA | NA | NA | NA | 0,0534170806611641 | 4 | -526,343818743911 | 1060,99066779085 | 28,5857030990219 | 6.8161457092201e-08 |
| **20** | 26,6253731343284 | + | + | NA | NA | + | NA | NA | NA | NA | NA | NA | NA | NA | NA | NA | 0,0599289479658493 | 5 | -525,871173839129 | 1062,20036294543 | 29,7953982536039 | 3.72269023934419e-08 |
| **1** | 27,2678832116789 | NA | NA | NA | NA | NA | NA | NA | NA | NA | NA | NA | NA | NA | NA | NA | 0 | 2 | -530,102559176374 | 1064,29467059155 | 31,8897058997225 | 1.30642286716515e-08 |
| **3** | 27,6566666666667 | NA | + | NA | NA | NA | NA | NA | NA | NA | NA | NA | NA | NA | NA | NA | 0,0021544621755788 | 3 | -529,954883739105 | 1066,09021860603 | 33,6852539141985 | 5.3233555778431e-09 |
